# Population matched (PM) germline allelic variants of immunoglobulin (*IG*) loci: New pmIG database to better understand *IG* repertoire and selection processes in disease and vaccination

**DOI:** 10.1101/2020.04.09.033530

**Authors:** Indu Khatri, Magdalena A. Berkowska, Erik B. van den Akker, Cristina Teodosio, Marcel J.T. Reinders, Jacques J.M. van Dongen

## Abstract

At the population level, immunoglobulin (*IG*) loci harbor inter-individual allelic variants in the many different germline *IG* variable (*V*), Diversity (*D*) and Joining (*J*) genes of the *IG* heavy (*IGH*), *IG* kappa (*IGK*) and *IG* lambda (*IGL*) loci, which together form the genetic basis of the highly diverse antigen-specific B-cell receptors. These inter-individual allelic variants can be shared between or be specific to human populations. The current *IG* databases IMGT, VBASE2 and IgPdb hold information about germline alleles, most of which are partial sequences, obtained from a mixture of human (B-cell) samples, many with sequence errors and/or acquired (non-germline) *IG* variations, induced by somatic hypermutation (SHM) during antigen-specific B-cell responses. We systematically identified true germline alleles (without SHM) from 26 different human populations around the world, profiled by the “1000 Genomes data”. Our resource is uniquely enriched with complete *IG* allele sequences and their frequencies across human populations. We identified 409 *IGHV*, 179 *IGKV*, and 199 *IGLV* germline alleles *supported by at least seven haplotypes* (= minimum of four individuals), after removal of potential false-positives, based on using other genomic databases, i.e. ENSEMBL, TopMed, ExAC, ProjectMine. Remarkably, the positions of the identified variant nucleotides of the different alleles are not at random (as observed in case of SHM), but show striking patterns, restricted to limited nucleotide positions, the same as found in other *IG* data bases, suggesting over-time evolutionary selection processes. The identification of these specific patterns provides extra evidence that the identified variant nucleotides are not sequencing errors, but genuine allelic variants. The diversity of germline allelic variants in *IGH* and *IGL* loci is the highest in Africans, while the *IGK* locus is most diverse in Europeans. We also report on the presence of recombination signal sequences (RSS) in *V* pseudogenes, explaining their usage in *V(D)J* rearrangements. We propose that this new set of genuine germline *IG* sequences can serve as a new population-matched *IG* (pmIG) database for better understanding B-cell repertoire and B-cell receptor selection processes in disease and vaccination within and between different human populations. The database in format of fasta is available via GitHub (https://github.com/InduKhatri/pmIG).

**Contribution to the Field Statement:** We present a catalogue of immunoglobulin (*IG*) germline-alleles of unprecedented completeness and accuracy from 26 different human populations belonging to five different large ethnicities (Source: 1000 Genomes). We identified the population distribution of several known germline alleles and identified multiple new alleles, especially in African populations, indicative of high allelic diversity of *IG* genes in Africa. Strikingly, the identified variant nucleotides of the different alleles are not at random, but show striking patterns, restricted to limited nucleotide positions, the same as found in other *IG* databases, suggesting over-time evolutionary selection processes. Furthermore, we identified recombination signal sequences in pseudogenes (previously not known). We provide an overview of *IG* germline alleles shared with and between known databases and also point to potential sources of non-germline variation and incompleteness of the existing *IG* databases. More importantly, we believe that this information can serve as a novel population-matched *IG* (pmIG) database, highly valuable for the research community in supporting the dissection and understanding of differences in effectiveness of antibody-based immune responses in infectious diseases, other (immune) diseases and vaccination within and between human populations. Such knowledge might be used in developing population-specific vaccination strategies e.g. for currently ongoing SARS-CoV2 pandemic.

## Introduction

The complex mechanism of antibody production from immunoglobulin (*IG*) genes is key to the development of the broad repertoire of the antigen-specific B-cell receptors of the adaptive immune system (1–5). These Ig proteins (antibodies) are assembled in B cells from two pairs of polypeptide chains, the so-called Ig heavy (IgH) and Ig light (Igκ or Igλ) chains that are assembled from different genes present in the *IG* loci, termed variable (*V*), diversity (*D*), joining (*J*) and constant (*C*) (**Figure 1A**). The *IG* heavy chain locus (*IGH*) on chromosome 14q32.3 consists of multiple different functional genes: ∼44 *V*, ∼27 *D*, ∼6 *J* and ∼9 *C* genes (**Figure 1B**). During recombination, one of each *V*, *D* and *J* genes recombine to a *V*-*D*-*J* exon to code for the antigen-binding domain of the IgH chain (**Figure 1A**). The rearrangements in both *IG* light chain loci (kappa: *IGK*, on chromosome 2p11.2; lambda: *IGL,* on chromosome 22q11.2) take place in an analogical way with direct rearrangement between *V* and *J* genes, as *D* genes are absent. This process itself can produce up to three million different antibodies (6). Additionally, most of the *IG* genes harbor inter-individual germline allelic variants, which can be shared between or be specific to human populations (7–9). Consequently, different individuals can produce different antibodies (derived from different allelic variants), implying that at the population level the diversity of antibodies is even more extensive than at the individual level.

**Figure 1:**
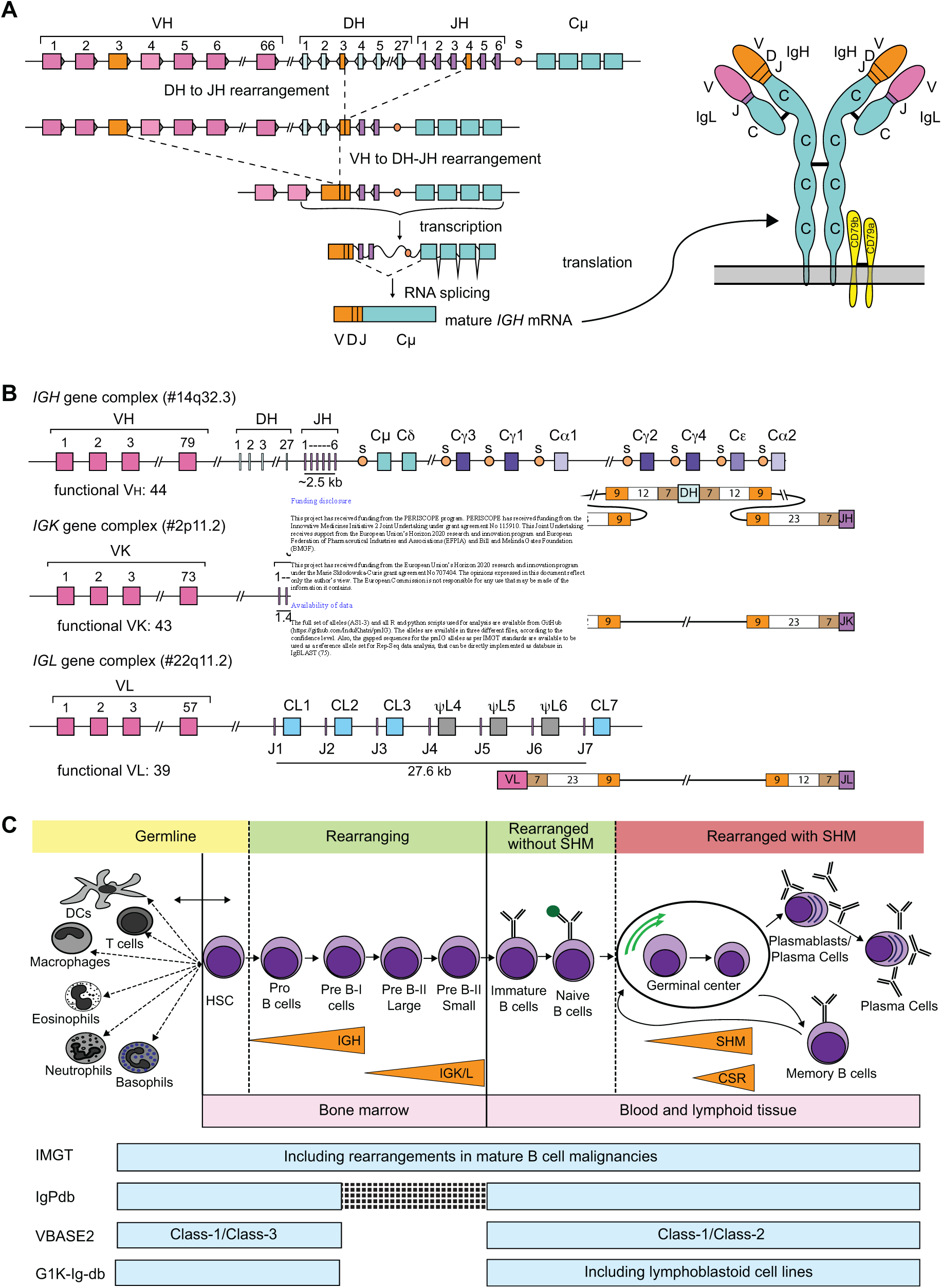
Generation and assessment of diversity in *IG* loci. **(A)** In the first step of *V(D)J* recombination in the *IGH* locus a *D* gene is coupled to a *J* gene. Subsequently, a *V* gene is coupled to the *DJ* joint. The VDJ exon is transcribed and spliced to the *IGHM* exons. An analogical process takes place in the *IG* light chain genes. When a functional IgM protein is transported to the plasma membrane with anchoring molecules CD79a and CD79b, and a functional Ig light chain it forms a complete antibody molecule. **(B)** Schematic overview of the three *IG* loci: *IGH*, *IGK*, *IGL* and the structure of their corresponding Recombination Signal Sequences (RSS). Genomic position of the loci is indicated in brackets. In *IG* loci each rectangle depicts one of the variable (*V*), diversity (*D*), joining (*J*) and constant (*C*) genes, and circles depict switch regions. The number of known functional genes, as listed in the IMGT, is indicated underneath each scheme. RSS structure schemes depict the position of heptamers (7), nonamers (9) and spacers (12/23) relative to *V*, *D* and *J* genes. **(C)** Hematopoietic stem cells in bone marrow, give rise to cells of both myeloid and lymphoid origin. While most of the cell types retain *IG* genes in their germline configuration, precursor B cells rearrange first Ig heavy chain and then Ig light chain genes to form a functional antibody. B cells with the functional B-cell receptor migrate to the periphery where they can recognize antigen. Upon antigen recognition and receiving help from T cells, B cells enter germinal center reaction during which they undergo intensive proliferation, improve affinity for antigen by the introduction of somatic hypermutations (SHM) in rearranged *IG* genes, and may change their effector functions in the process of class-switch recombination (CSR). This results in the formation of memory B cells and antibody-secreting plasma cells. *IG* genes can be sequenced from any B-cell type. However, in (virtually) all cells other than B cells, they will be in their germline configuration. Precursor B cells and naive mature B cells carry rearranged *IG* genes, which can be further modified by the presence of SHM in post-germinal center cells. Light blue block underneath B-cell maturation scheme depicts the sources of *IG* sequencing in the three existing *IG* databases: IMGT (ImMunoGeneTics, http://www.imgt.org/), IgPdb (http://cgi.cse.unsw.edu.au/~ihmmune/IgPdb), VBASE2 (http://www.vbase2.org), and the *IG* gene sequence data from the 1000 genome project (G1K-Ig-db, http://www.internationalgenome.org). VBASE2 has classified the alleles in different classes based on their genomic and rearrangement evidence. Class-1 alleles in VBASE2 have evidence from both; Class-2, and Class-3 alleles of VBASE2 are supported either by germline occurrence or rearranged repertoires.

The *V(D)J* recombination process is guided via short highly conserved DNA stretches, called “recombination signal sequences” (RSS), present at each recombination site of the *IG* gene segments, i.e. downstream to *V*, upstream to *J,* and at both sites of *D* (RSS panel in **Figure 1B**) (6,10–12). Each RSS comprises a palindromic heptamer (7bp: CACAGTG), a spacer (12 or 23 bp) and a nonamer (9bp: ACAAAAACC), which are signals for recognition and binding by recombination proteins (5). The heptamer juxtaposes the site of DNA cleavage (13), the spacer length restricts recombination according to the 12-23 recombination rule (5), and the nonamer provides the binding site for the RAG1 recombination protein (14, 15).

The B-cell receptor (BCR) remains the same during the developmental stages of B cells from bone marrow (BM) to naive mature B cells (**Figure 1C**). However, upon antigen recognition, generally taking place in germinal centers (GC) during interaction with T cells, B cells proliferate and modify the antigen-binding domain of their BCR via somatic hypermutation (SHM), randomly occurring in the *V(D)J* exon region. B-cells with SHM that induce better antigen-binding of their antibodies will be positively selected and contribute to improved B-cell responses such as in vaccination. Thus, *IG* genes sequenced from antigen-experienced B cells, most likely carry SHMs; if used for databases, such sequences might unwantedly result in false identification of allelic variants. The identification of new germline alleles from B-cell repertoire transcriptomics data have changed the perspective of the field (16, 17), but they may at any stage also suffer from sequencing errors.

Variations in individual germline genes can result in inter-individual differences in immune response (18–21). Consequently, reliable information on true germline allelic variants is important to understand such inter-individual differences. Several *IG* databases, such as IMGT (22–27), IgPdb (http://cgi.cse.unsw.edu.au/~ihmmune/IgPdb/), and VBASE2 (28), report germline variations, present in different individuals. The sources of these three databases differ and each of them comprises specific errors depending on the origin of the included *IG* sequences. In **Figure 1C**, the sources of the three different databases are aligned with the B-cell differentiation & maturation pathway, including the GC and post-GC stages, where SHMs take place or have taken place. IMGT is the most widely used database, because of its early availability, longstanding experience and the most complete structure, but is at least in part derived from mature post-GC B-cells (25,25,26). IgPdb does not comprise complete *IG* gene sequences, owing to short read sequencing technologies. VBASE2 has better strategies for identification of true germline alleles, but all the alleles are partial in sequence; this database is drawn from genome databases, namely the EMBL nucleotide sequence database, Ensembl, and supported by evidence from the rearranged repertoires.

Completeness and accuracy of germline sequences will greatly influence downstream analyses in repertoire sequencing (Rep-Seq), as unreported *IG* allelic variants can appear as recurrent SHMs and skew estimated segment distributions and/or estimated mutation frequencies in clinically relevant decision processes e.g. infection response, vaccination studies (17,29–31) and prognosis in chronic lymphocytic leukemia (32). Accordingly, the open germline receptor database (OGRDB) aims to stringently assess and classify the germline alleles (33, 34). OGRDB’s endeavor does not include *IG* alleles called from high-throughput whole genome mapping studies, as OGRDB believes that the inferred sequences may contain false positives and should be supported by rearrangements (34). Nevertheless, whole genome sequencing provides the opportunity to search for allelic variants, which might be shared or be unique to human populations. With the application of stringent parameters on whole genome studies, a new transitional resource can be created which will have accurate germline alleles with population information for each allele on top of complete *IG* genes similar to IMGT, as well as a proper classification of alleles like VBASE2.

The “1,000 Genomes (G1K)” dataset, derived from cell samples of 2,504 individuals, has been used to call alleles (9). Yu *et al* developed a method named alleleminer to determine alleles for the *IG* and *TR* loci from the G1K data creating the Lym1k database. The Lym1k database has provided a base to use G1K data, but the information provided may not be relevant for the research community because the reliability of the new alleles identified is not provided. Moreover, they did not retain population information, which is critical to develop a rich resource (9). Furthermore, Yu *et al* also did not include all relevant components of each *IG* locus namely *D*, *J*, *C* genes and RSS, which is a strong feature of the IMGT database. To obtain an accurate set of alleles with enriched information, we have profiled all alleles for the *V*, *D*, *J*, *C* regions as well as the RSS regions for all three *IG* loci from the G1K whole-genome sequencing (WGS) data, and the reliability and population information of each allele is mentioned meticulously in a database called pmIG

Using the G1K resource, or using short read data, to profile alleles for the *IG* loci raises potential pitfalls (35–37). The repetitive and complex nature of *IG* loci makes it difficult regions to identify germline alleles from short read sequencing data in these regions. The level of haplotype diversity of *IG* loci is another limitation (37). The main complexity with analyzing the G1K data lies in the sample origin from the Epstein-Barr virus (EBV) transfected B-cell lines (**Figure 1C**). As ∼75% of the genomes (1,941 genomes) in G1K are derived from EBV-transformed B-cell cultures, we framed a set of rules to obtain high-fidelity alleles (detailed in Methods) that prevent them from being the result of sequencing errors or SHM. We have manually interfered with the resource to obtain highly accurate alleles, submitted them to GenBank and made them also available via GitHub with population information and confidence levels to provide access for the community. The compiled resource comprises of *IG* germline allelic variants that cover a wide range of ethnicities as investigated for five superpopulations, namely Africans (AFR), Americans (AMR), East Asians (EAS), Europeans (EUR) and South Asians (SAS).

## Materials and methods

### Data source

The 1,000 Genomes data (G1K) (May, 2013 release; http://www.1000genomes.org; GRCh37 assembly) in the form of phased variant cell format (VCF) was used for this study. Phased variants for GRCh38 (a recent release for the 1,000 genomes) are available, however, we used the GRCh37 version as the SNP IDs are not yet available for the GRCh38 mapping. These SNP IDs are relevant to perform mapping to other databases and assess false negatives. Both genome assemblies do not comprise of all the *V* genes mentioned in the IMGT database as multiple duplicated genes are not present in all individuals. The full release of the data set was collected from 2,504 cell samples from diverse ethnic groups that have a uniform distribution of individuals across populations. The samples are classified in five superpopulations i.e. Africa, America, East Asia, Europe and South Asia, that are further subdivided into 26 populations (7 African, 4 American, 5 East Asian, 5 European and 5 South Asian populations) with a minimum of 61 and a maximum of 113 samples per population (**Table S1**). The VCF format of the data comprises information of both parental and maternal chromosomes for each sample.

### Identification of alleles from G1K data

The genes (*V*, *D*, *J* and *C*) from *IG* loci were retrieved from the VCF files for Chromosome 14 (*IGH*), Chromosome 2 (*IGK*), and Chromosome 22 (*IGL*). Only SNPs, deletions or insertions were processed from the VCF files. Copy number variations were not considered. Software, such as Plink (http://zzz.bwh.harvard.edu/plink/), to retrieve haplotypes from VCF files cannot process multi-allelic SNPs. Therefore, special python scripts were written, while R scripting was used to obtain the 5,008 independent haplotypes from the 2,504 cell samples (all available via GitHub). Identical haplotypes are merged, and the number of times a particular haplotype appears is counted and marked as an allele. The IMGT nomenclature is used to name genes, and alleles extend this name with a numbering, for example the 01 and 02 alleles of the *IGHV1-8* gene are referred to as *IGHV1-8_01*, *IGHV1-8_02*. IMGT alleles are denoted with an asterix, such as *IGHV1-8*01*, *IGHV1-8*02*. The alleles are sorted in descending order such that it the first allele is supported by the maximum number of haplotypes.

### Terminology

With the term *“haplotype*” we refer to an operationally distinguishable gene (segment) present on one strand (inherited from a single parent) in one individual. There are two haplotypes, one on each positive and negative strand, with exactly the same or different polymorphisms. With *“allele”* we refer to the profile of variants across one haplotype.

*“Mutations”* are genetic mutations that occurred to form different alleles of the gene and “*Somatic Hypermutation Mutation (SHM)*” are the mutation that arises in *IGH/IGK/IGL* sequences in post-GC cells.

### Mapping the alleles to existing databases

The alleles obtained from the G1K samples are mapped to three different databases namely IMGT, IgPdb and VBASE2 (28) using Muscle (38) and manually checked to ensure accurate mapping. IMGT (www.imgt.org) is the global reference in immunogenetics and immunoinformatics studies and is maintained since 1989. IgPdb (http://cgi.cse.unsw.edu.au/~ihmmune/IgPdb) is a repository of suspected allelic variants of human *IG* germline genes. VBASE2 (http://www.vbase2.org) presents *V* gene sequences extracted from the EMBL nucleotide sequence database and Ensembl together with links to the respective source sequences. VBASE2 classifies the *V* genes into three different classes: Class-1, genomic and rearranged evidence; Class-2, genomic evidence only; and Class-3, rearranged evidence only (28). This evidence classification of alleles was only performed by VBASE2 database; none of the other databases had such feature, implying that the capability for rearrangement is formally not included in the IMGT and IgPdb databases.

### Classifying alleles into confidence levels

The alleles in our study were classified into three major confidence levels (allele set (AS) 1-3) as described below:

*AS1 (known)*: G1K alleles with a minimum support of 4 haplotypes and identified in either the IMGT, IgPdb and/or VBASE2 (Class-1) databases. This AS1 allele set obviously has the highest level of confidence as the alleles are observed in the G1K resource as well as in at least one of the three existing databases. This set of alleles also validates G1K as a solid resource since there is substantial overlap with the existing databases.

The alleles that did not classify as AS1 were divided into two categories:

*AS2 (frequent new alleles):* G1K alleles with a minimum support of 19 haplotype (minimum of ten individuals). These alleles would represent a set of newly identified alleles that are frequent.

*AS3 (rare new alleles):* G1K alleles that have a haplotype support between 7 and 18 (minimum four individuals). This group of new alleles with less confidence in terms of haplotype support is called rare alleles. Despite the rarity of these alleles, we believe that they are genuine allelic variants, because the chance that 7 identical haplotypes within 5,008 independent haplotypes are caused by sequencing errors is highly unlikely.

As alleles can be duplicated or diverged from each other, we further subdivided all alleles into three other categories (**Figure S1**):

*Alleles for Group genes:* Alleles for which the genes are marked as duplicated in the *IG* loci. These are *IGHV1-69*, *IGHV1-69D*; *IGHV3-43*, *IGHV 3-43D; IGHV3-23*, *IGHV3-23D*; *IGHV3-64*, *IGHV3-64D* and *IGHV2-70*, *IGHV2-70D* pairs.

*Alleles for operationally indistinguishable (OI) genes:* As multiple *V* genes are paralogous (39, 40), the mapping of short reads to such genes can be erroneous, influencing the subsequently derived alleles. Mutations on the alleles of such genes can thus easily be false positives, even after using stringent parameters. We denote these genes as operationally indistinguishable (OI) genes. As these genes can be recognized based on their similarity (41), we generated a neighbor-joining (NJ) tree for all *V* genes on the *IGH*, *IGK* and *IGL* loci, separately. The genes sharing a clade with a short branch length i.e. 0.02, are called OI genes (**Figure S1**); and the corresponding alleles as OI alleles.

*Alleles for self-evident (SE) genes:* Alleles that are *not* annotated as group or OI alleles.

The alleles that fall into AS1 category i.e. Known alleles are also termed as group or OI alleles as these resources also contain false positives.

### Filtering out false positive alleles

The G1K alleles were scrutinized manually.

1. Alleles with stop codons were removed from the final set.
2. Alleles with mutations or frameshift mutations absent in any of the following resources were removed: ESP (https://evs.gs.washington.edu/EVS/), TOPMed (https://www.nhlbi.nih.gov/science/trans-omics-precision-medicine-topmed-program), gnomAD (https://gnomad.broadinstitute.org/), and ProjectMine (https://www.projectmine.com/).
3. All the alleles of group genes or OI genes (e.g. *IGHV1-69D* gene from IMGT and alleles *for IGHV1-69* gene (group genes); alleles for all *IGHA1* and *IGHA2* genes (OI genes)) were aligned. We removed alleles within group genes and OI genes when a mutation of an allele is shared between alleles belonging to different genes within the group (pointing towards a mis-alignment of a read) except when this mutation is present in one of the databases across multiple alleles. For example, an allele of *IGHA1* has a position mutated i.e. A->C exclusively and two alleles from *IGHA2* also has similar mutation at the same aligned position, then all the three alleles were considered as false positives and filtered out.

### Identifying mutation patterns in the filtered alleles

To identify the mutation patterns, we performed alignments of all alleles per gene. The alleles were compiled from our database pmIG and the other existing databases i.e. IMGT, IgPdb and VBASE2. In the alignments per gene, the mutating positions are identified for all alleles. In the complete set of alignments per gene, the mutated positions for all alleles of that gene are compared and characterized as *new* (when the mutation is only seen in our resource) or *known* (when the mutation is seen in one of the other resources). We have done this for mutations in the *V* region as well as the leader sequences. The positions added by our resource are mentioned and the pattern of the mutations i.e. conserved (= at fixed positions) or random (= scattered, as caused by SHM) is identified.

### Mapping population information to the identified alleles

G1K alleles are annotated with superpopulation information (**Tables S2-S4**) into four categories: 1) ALL, present in all superpopulations; 2) AFR, only present in Africans; 3) AFR SHARED, present in African and at least one of the other superpopulations, but not all; and 4) NON-AFR, present in at least one of the superpopulations but not in Africans.

### Variants in RSS haplotypes

We retrieved the RSS variants from the 40 bases adjacent to 3’ *IGHV* genes and the 5’ *IGHJ* genes (having 23-bp spacers), and from the 30 bases adjacent to 5’ and 3’ *IGHD* genes (having 12-bp spacers). Similarly, variants were retrieved from the 30 bases adjacent to 3’ *IGKV* and 5’ *IGLJ* genes (having 12-bp spacers) and from the 40 bases adjacent to 5’ *IGKJ* and 3’ *IGLV* genes (having 23-bp spacers). The perfect RSS sequence has a conserved heptamer “CACAGTG”, a conserved nonamer “ACAAAAACC”, and a specific length of the spacer sequence (23 bp or 12 bp). Mutations in heptamer and nonamer sequences as well as a deviating length of the spacer (less than 23 bp or 12 bp) directly affect the recombination frequency of the linked genes (6,29,42,43).

### Phylogenetic trees for alleles

Maximum Likelihood trees were built for the alleles using RAxML (44). The PROTGAMMAJTT model was used to build the trees with 100 bootstraps. The trees were visualized using the iTOL server (45). The trees taxa were colored as per AS classification; the population level annotation is displayed in binary format and the frequency of alleles as text.

### Independent validation of the mutation patterns in the pmIG alleles

To validate that the mutations that we detect are caused by somatic hypermutations (SHM), we aligned the alleles to the rearranged *IGH* sequences derived from the transcriptomics data of antigen-experienced B cells i.e. sorted HBsAg^+^ B cells, sampled after primary Hepatitis B vaccination (46). The raw fastq files were obtained from SRA (SRP068400). Paired-end reads were joined using fastq-join (ea-utils) with default settings and filtered for minimum Phred quality of 30 over at least 75 % of bases. IMGT/HighV-Quest (17) was used for sequence annotation and functional *IGH* sequences were retained. We selected 20 B-cell receptor sequences from affinity-maturated B-cells after Hepatitis B vaccination for each gene randomly and aligned those sequences to the all alleles (detected by us (pmIG), IMGT, IgPdb and VBASE2) for the respective genes. The mutating positions are marked and the mutation patterns were compared between germline alleles and antigen-experienced *IGH* sequences from HepB study.

### Genetic diversity and migration events of population based on *IG* loci

The VCF file of the complete individual locus, i.e. *IGH* (Chr14 [106032614, 107288051, complement]), *IGK* (Chr2 [89890568, 90274235]), [89156874, 89630436, complement]) and *IGL* (Chr22 [22380474, 23265085]), was subjected to a principal component analysis (PCA) using the R Bioconductor package ‘SNPRelate’ (47). We then calculated the pairwise population differentiation, which is based on levels of differentiation in polymorphism frequencies across populations, as quantified by the fixation index (F_ST_). F_ST_ is proportional to the evolutionary branch length between each pair of populations. F_ST_ distances between populations were visualized with a Neighbor joining tree. We used TreeMix (48) that uses the composite likelihood to build the population trees. Six migration edges are tested for significance using 500 SNPs per block (-k 500). As “Out of Africa” is the most accepted theory (49) we used Yoruba population (YRI) of the African superpopulation an outgroup for building the migration trees.

## Results

The 1,000 Genomes (G1K) database from 2,504 individuals is a resource covering 26 populations representing five continents (**Table S1**). We identified population-specific alleles in all three *IG* loci (*IGH*, *IGK* and *IGL*), where each allele is supported by at least seven haplotypes (four haplotypes for known alelles). We have collected the alleles identified from G1K to create a population matched *IG* (pmIG) database. These alleles were divided into three allele sets (AS1, AS2, or AS3) based on different confidence levels (see Methods). AS1 are known alleles; AS2 are novel alleles that are frequent in the populations (supported by at least 19 haplotypes); and AS3 are novel rare alleles (supported by at least 7 haplotypes and at max by 18 haplotypes) (**Figure S1**). The population matched *IG* (pmIG) database further contains meta information about the alleles such as the support of haplotypes for each (sub)population (**Tables S2, S3 and S4**).

### The alleles from G1K samples are not affected by SHM from EBV transfected cell-lines

The G1K samples originate from different sources, including EBV-transfected B-cell cultures and blood as well as an unreported source in a few cases. SHMs might be present in mature B-cells immortalized by EBV transfection. It should be noted that such EBV-transfected B-cells are mainly polyclonal, unless cultured for (very) long time (= many months or more) or single cell subcloned; this has not been the case for the G1K samples. Polyclonal B-cell cultures will not likely have dominant SHM-based nucleotide variants detectable (50). Nevertheless, we tested whether more allelic variants are found in samples stemming from an EBV origin. The metadata of the G1K samples reports EBV coverage which was obtained by mapping the sequencing reads to the EBV genome. Using these annotations, we divided the samples into two groups: 1) **Set-1**, the non-EBV samples (563 samples), comprising of samples derived from blood (i.e. samples with EBV coverage <20X) and samples where the source was not reported; and 2) **Set-2**, the EBV samples, containing the remaining samples (1,941 samples). We used filtered *IGHV* alleles for this analysis. Of 410 different *IGHV* alleles, 186 (45%) alleles are supported by samples in Set-1 (**Figure 2A**). From the 410 *IGHV* alleles, 145 (35%) are known to existing databases (AS1 category, Methods), and 103 (55%) of those are supported by Set-1 samples. There are 196 (47%) frequent novel alleles (AS2 category, Methods) from which 77 (39%) are also supported by Set-1 and 119 (61%) by Set-2. These relative frequencies comply with the distribution of the number of samples between the two sets, as Set-1 has about 3 times fewer samples than Set-2. Furthermore, rare alleles are not supported by Set-1 samples (**Figure 2B**).

**Figure 2:**
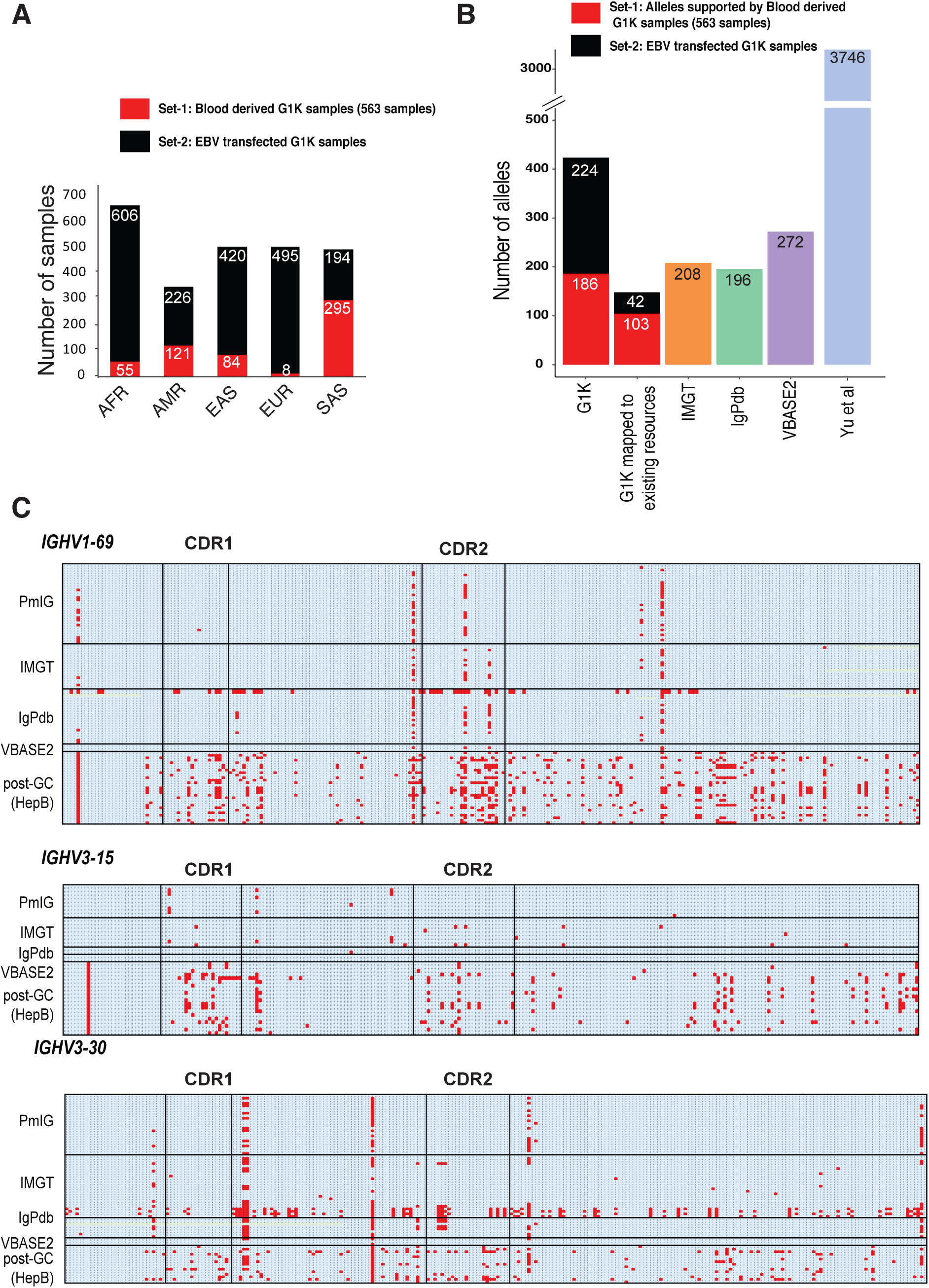
pmIG alleles are free of somatic-hypermutations (SHM). **A)** The distribution of samples in Set-1 and Set-2 from each super-population in G1K resource. **B)** The count of *IGHV* alleles in different resources. **C) Mutations mapped to the alleles from pmIG, IMGT, IgPdb and VBASE2 databases with the corresponding *IGHV* sequences from the HepB vaccination study for *IGHV1-69* and *IGHV3-30* genes.** The blue shade represents the conserved nucleotides and red shades indicate the mutated positions. Strikingly, the positions of the identified variant nucleotides of the *IGHV1-69* and *IGHV3-30* alleles in the pmIG database are not at random, but show restricted patterns, restricted to limited nucleotide positions, the same as found in other *IG* databases. However, several sequences in the IMGT and IgPdb databases show additional variant nucleotides at positions, which are mutated in the post germinal center (GC) B-cells after Hepatitis B vaccination. This comparison indicates that most likely the IMGT and IgPdb databases are contaminated with SHM-mutations, which is supported by the fact that several IMGT and IgPdb sequences seem to have comparable SHM patterns and are therefore most likely derived from the same postGC B-cell populations.

These observations on the novel alleles convince us that the *alleles that we detected in the G1K samples are not influenced by SHMs related to EBV-transfection* of mature (post-GC) B-cells, likely because we used the strict requirement of at least seven haplotypes for calling an allele. The Lym1k database by *Yu et al.* seems to have suffered from the contamination by EBV transfection as Yu et al. reported ∼3-4,000 alleles for *IGH* and *IGL* loci (9).

### Conserved mutation pattern of pmIG alleles differs from SHM-affected *IGH* sequences

Germinal center (GC) reactions result in affinity maturation of the Ig molecules, based on SHM and subsequent selection processes (51). To further substantiate that the alleles identified from the G1K resource with its stringent filtering criteria are correct, we aligned the alleles from all the resources with the *IG* sequences obtained after Hepatitis B (HepB) vaccination in naive individuals (***Methods***). **Figure 2C** shows the mutating positions for selected genes i.e. *IGHV1-69, IGHV3-15* and *IGHV3-30* (mutated positions in red).

Interestingly, the mutations in the pmIG alleles for these genes are not random and thus follow the above described strictly conserved pattern. In contrast, IMGT alleles for *IGHV1-69 and IGHV3-23* are heavily mutated, as well as the IgPdb alleles for *IGHV1-69*. From this, we conclude that the novel alleles that we identify in our database, pmIG, are generally free from SHMs and mostly represents combinations of already known mutating positions with only a few new mutations, if at all **(see also Figure S2).** Moreover, it is highly likely that the existing databases thus suffer from the false-positive alleles as a result of SHMs.

### Most known alleles are frequent and present in all ethnicities

The alleles that map to the IMGT database as well as to the other databases were instrumental in identifying two groups of alleles, i.e. known and novel alleles (see Methods). We found that 35% of the *IGHV* alleles mapped to the known alleles (**Table 1**) with 60% of them present in all the superpopulations with support of at least 100 haplotypes (**Figure 3A**). Most of these alleles are shared with the IMGT database, indicating that the IMGT database contains universal alleles. This was similar for the *IGKV* (34%) and *IGLV* (31%) alleles.

**Figure 3:**
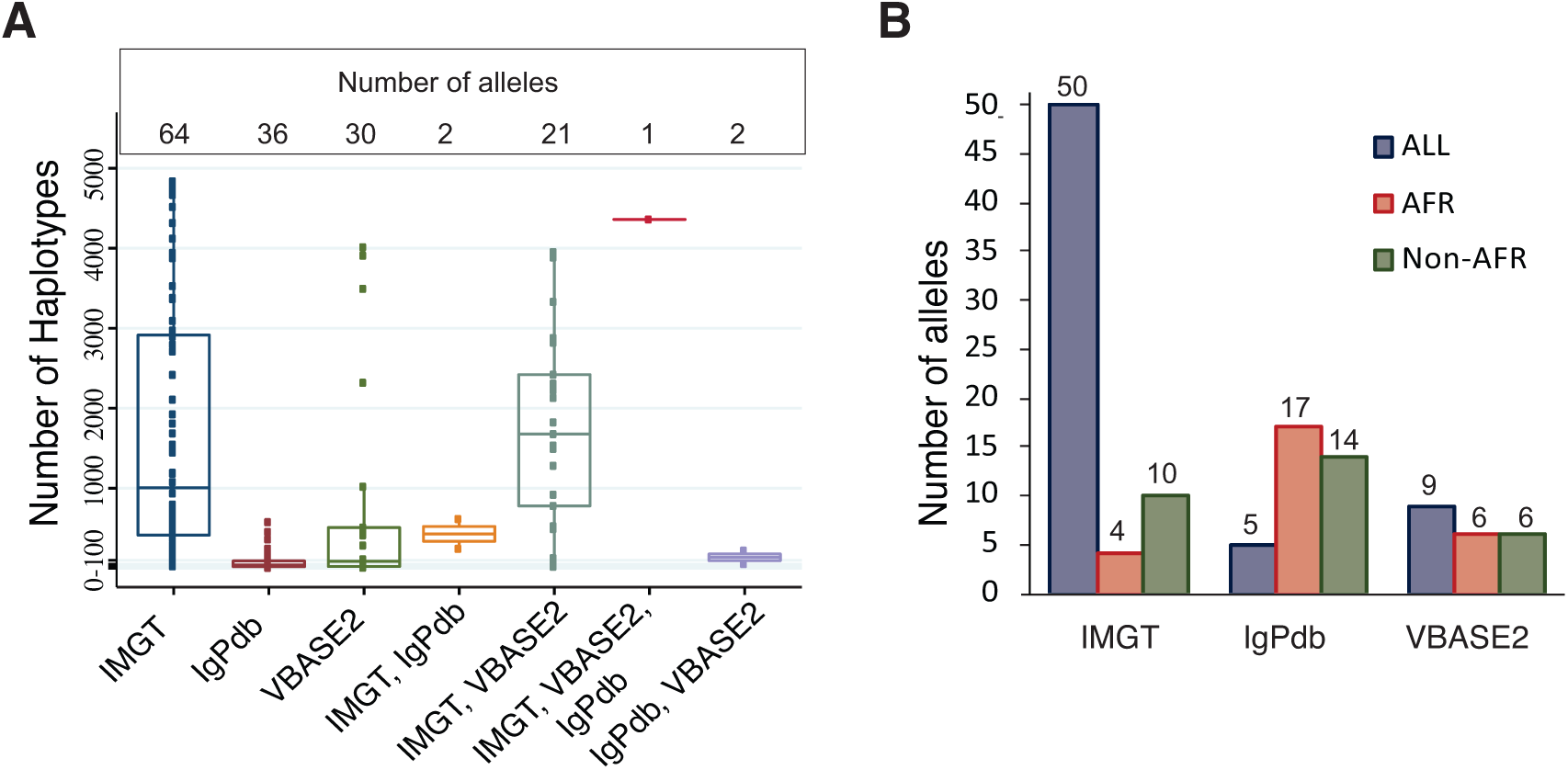
The known alleles are frequent and are present in all the ethnicities. **A)** Haplotypes support of the alleles mapped to the existing databases. Each dot is an allele. The alleles mapped to the alleles in multiple resources are also indicated. **B)** The population distribution of the AS1 (known) alleles. The alleles shared between IMGT and other resources are considered to be an IMGT alleles. Similarly, alleles are considered to be belonging to IgPdb if mapped to IgPdb and VBASE2 databases. IgPdb and VBASE2 harbor alleles unique to specific populations.

**Table 1:** Number of alleles in different functional gene segments in IG loci. AS1 (Known), AS2 (Frequent) and AS3 (Rare) are major confidence levels. AS2 and AS3 alleles are further subdivided into SE (self-evident alleles), GA (group alleles) and OI (operationally indistinguishable) alleles. * 5 AS1 alleles were identified as false positives, based on mapping to the paralogs in *IGHG* alleles.

|  | AS1 |  |  |  | AS2 |  |  |  | AS3 |  |  |  |  |  |  |  |  |
| --- | --- | --- | --- | --- | --- | --- | --- | --- | --- | --- | --- | --- | --- | --- | --- | --- | --- |
|  | Sum | SE | GA | OI | Sum | SE | GA | OI | Sum | SE | GA | OI | TOTAL |  | IMGT | IgPdb | VBASE2 |
| ***IGHV*** | 145 | 95 | 31 | 19 | 196 | 91 | 47 | 58 | 68 | 25 | 18 | 25 | 409 |  | 236 | 196 | 272 |
| ***IGHD*** | 29 | 29 | 0 | 0 | 8 | 8 | 0 | 0 | 5 | 5 | 0 | 0 | 42 |  | 33 | 2 | 0 |
| ***IGHJ*** | 5 | 5 | 0 | 0 | 4 | 4 | 0 | 0 | 0 | 0 | 0 | 0 | 9 |  | 12 | 1 | 0 |
| ***IGHC*** | 16* | 5 | 11 | 0 | 71 | 24 | 47 | 0 | 44 | 16 | 28 | 0 | 125 |  | 49 | 0 | 0 |
| ***IGKV*** | 83 | 10 | 73 | 0 | 70 | 5 | 65 | 0 | 26 | 3 | 23 | 0 | 179 |  | 80 | 23 | 168 |
| ***IGKJ*** | 7 | 7 | 0 | 0 | 2 | 2 | 0 | 0 | 0 | 0 | 0 | 0 | 9 |  | 9 | 0 | 0 |
| ***IGKC*** | 2 | 2 | 0 | 0 | 0 | 0 | 0 | 0 | 1 | 1 | 0 | 0 | 3 |  | 5 | 0 | 0 |
| ***IGLV*** | 84 | 56 | 0 | 28 | 81 | 59 | 0 | 23 | 34 | 26 | 0 | 8 | 200 |  | 74 | 9 | 136 |
| ***IGLJ*** | 10 | 10 | 0 | 0 | 11 | 11 | 0 | 0 | 6 | 6 | 0 | 0 | 27 |  | 10 | 0 | 0 |
| ***IGLC*** | 5 | 5 | 0 | 0 | 16 | 16 | 0 | 0 | 8 | 8 | 0 | 0 | 29 |  | 14 | 0 | 0 |

Out of 29 African *IGHV* alleles that mapped to known alleles (**Figure 3B, 4**), 17% (5) map to IMGT and the majority of them (60%;16) map to IgPdb with a minimum haplotype support of 5 (**Figure 3B**). The lower haplotype support of the alleles mapped to IgPdb databases suggests that the IgPdb includes rare alleles, whereas IMGT is comprised of frequent alleles. Similarly, known African alleles in *IGK* and *IGL* loci had a larger overlap with IgPdb and VBASE2 than with IMGT, suggesting that alleles private to specific populations are not present in IMGT. IgPdb and VBASE2, however, do not capture the complete diversity of African ethnicity as they contain ∼20% of the total African frequent *IGHV* alleles found from ∼600 African individuals in G1K. *IGHV* alleles private to Asian populations are mostly absent in all three existing databases. *IGKV* and *IGLV* alleles private to any population including Africans are not profiled in any of the current resources (**Figures 5, 6**). These findings indicate a biased sampling by the current databases.

**Figure 4:**
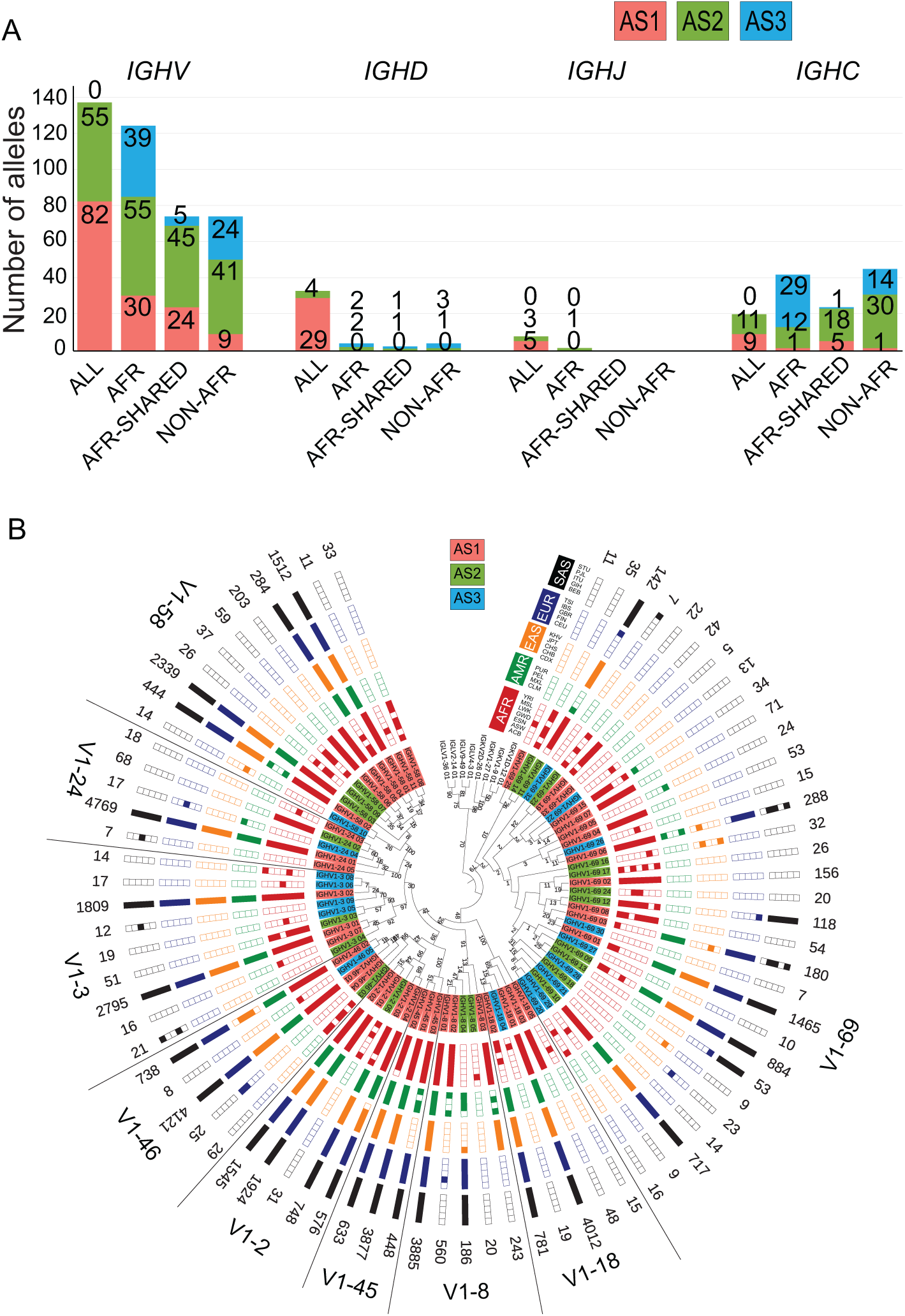
Population distribution of alleles in *IGH* heavy chain locus. **A)** The relative superpopulation distribution of *VDJ* genes and *C* genes for *IGH* locus. The population plots are represented for all superpopulations (ALL), Africans only (AFR), Africans shared with one of the other superpopulations (AFR Shared) and ‘Non-AFR where alleles are present in at least one of the populations other than Africans. **B)** Maximum Likelihood tree of the population distribution of *IGHV1* family alleles. The *IGHV1* family genes are indicated in the legends. Red label background indicates AS1 alleles, green AS2 and blue AS3 alleles. The population distribution is plotted in a binary format where each block is a population. Filled block represents the presence of that allele in at least four haplotypes in that population, otherwise the block is unfilled. For the population distribution of other *IGHV* families refer to Figures S3 and S4.

**Figure 5:**
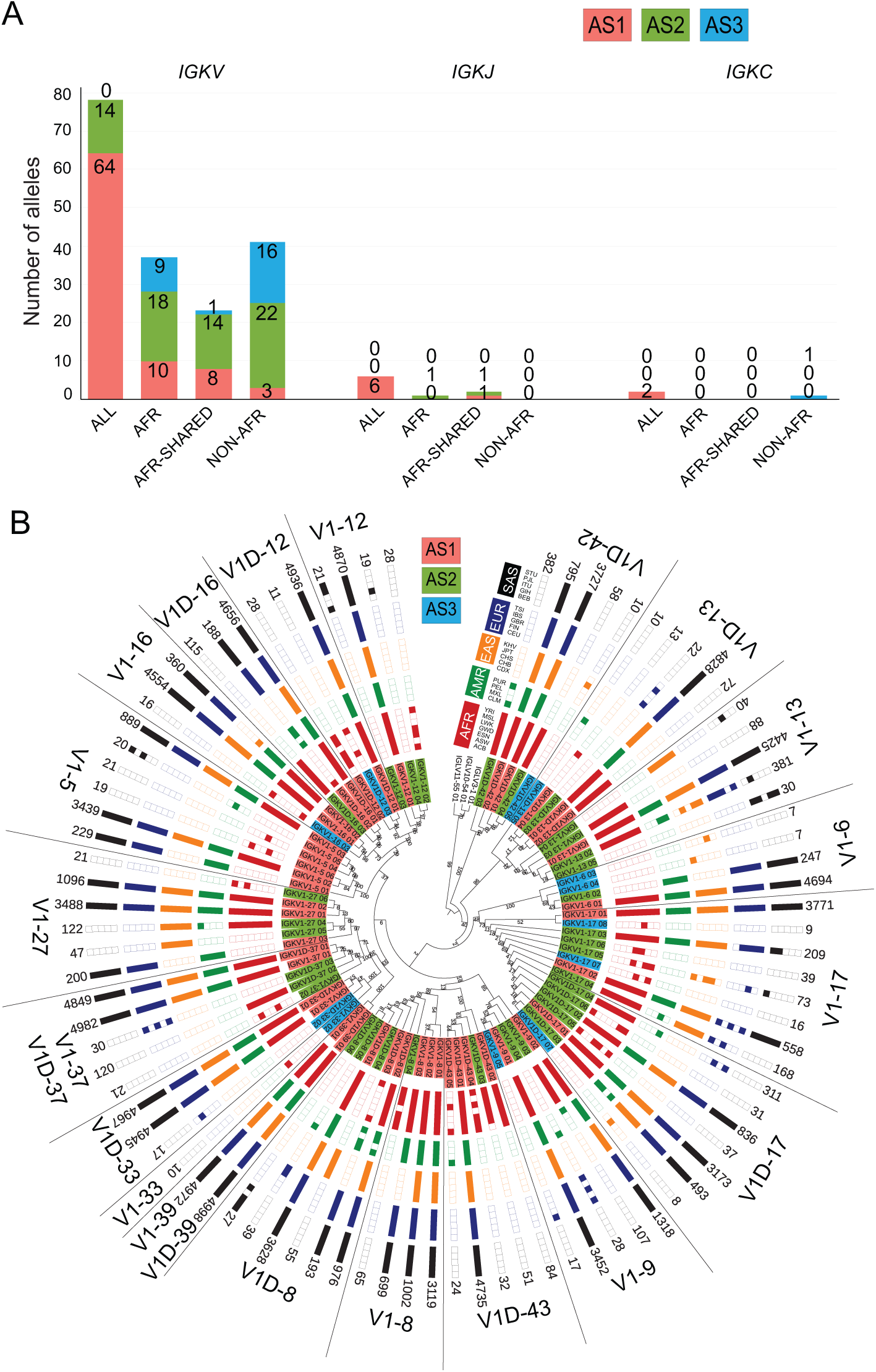
Population distribution of alleles in *IGK* light chain locus. **A)** The relative superpopulation distribution of *VJ* genes and *C* genes for *IGK* locus. The population plots are represented for all superpopulations (ALL), Africans only (AFR), Africans shared with one of the other superpopulations (AFR Shared) and ‘Non-AFR where alleles are present in one of the populations other than Africans. **B)** Maximum Likelihood tree of the population distribution of *IGKV1* family alleles. For the population distribution of other *IGKV* families refer to Figure S7.

**Figure 6:**
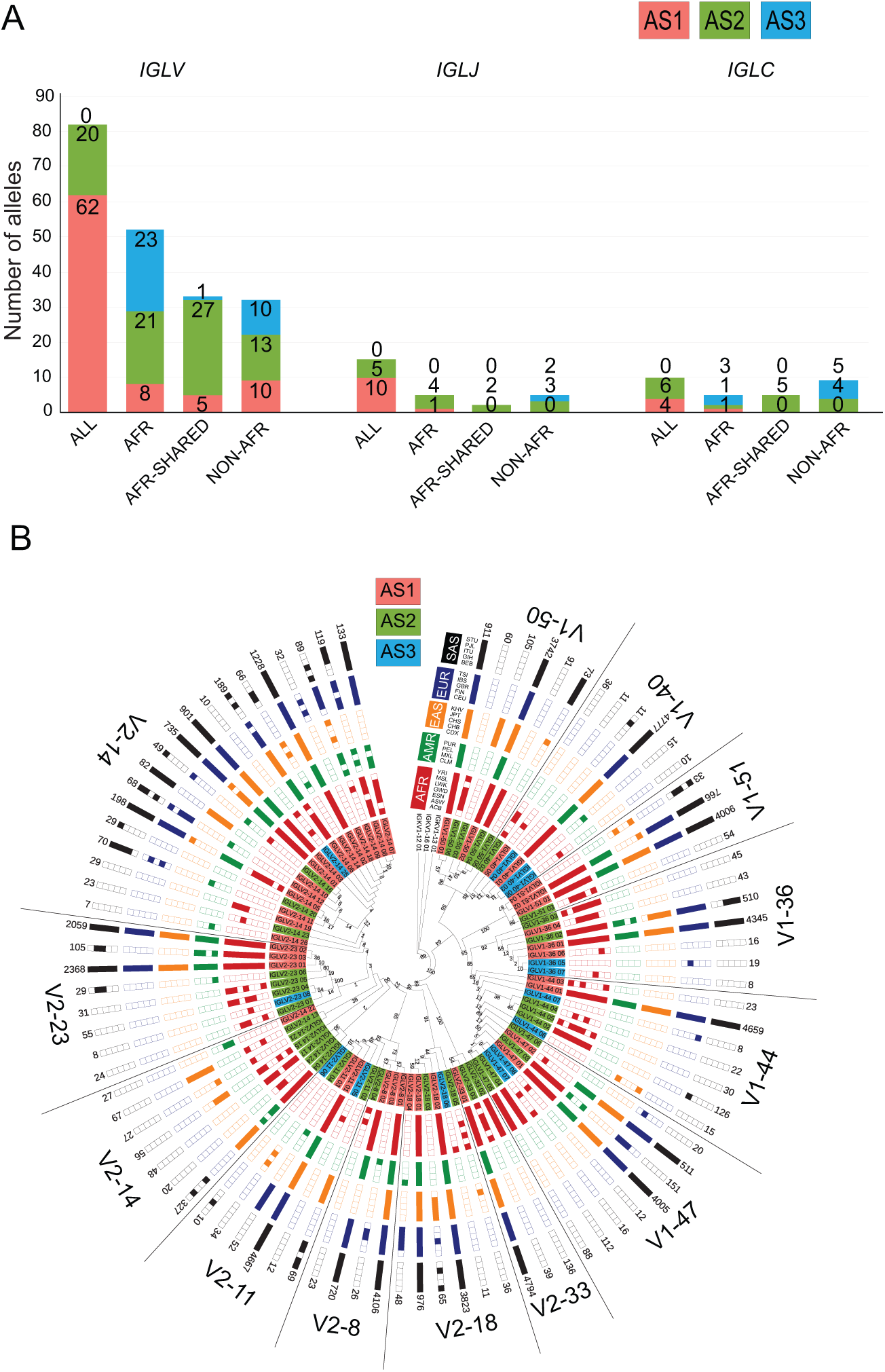
Population distribution of alleles in *IGL* light chain locus. **A)** The relative superpopulation distribution of *VJ* genes and *C* genes for *IGL* locus. The population plots are represented for all superpopulations (All), Africans only (AFR), Africans shared with one of the other superpopulations (AFR Shared) and ‘Non-AFR where alleles are present in one of the populations other than Africans. **B)** Maximum Likelihood tree of the population distribution of *IGLV1* and *V2* family alleles. For the population distribution of other *IGLV* families refer to Figure S8.

### Conserved mutation patterns in the filtered alleles as compared to the existing databases

Even after stringent filtering, “novel alleles” might suffer from SHM or sequencing errors (37). As SHMs are introduced randomly and do not have a fixed pattern, we have already eliminated possible false positives by putting a threshold of seven haplotypes (= at least four persons). Furthermore, 118 potential false-positive alleles were eliminated that were arising due to the lack of support in other databases or the same mutating patterns between the members of group and OI genes (**Table S5**). Moreover, when we performed the alignments of the novel alleles with the known alleles present in the pmIG, IMGT, IgPdb and VBASE2 databases, it appeared that the novel alleles revealed highly conserved mutation patterns at fixed nucleotide positions, the same positions as found in the other databases (**Figure S2**). The most spectacular examples from *IGH* locus are that the novel alleles of the *IGHV2-5*, *3-15*, *3-20*, *3-49*, *3-64*, *3-72*, *3-74*, *4-39*, *6-1* genes that did not gain any new polymorphisms, suggesting that evolutionary pressure and selection play an important role for the remaining locations in these genes (**Figure S2**). A comparable observation was made for the *IGHV1-69* gene, where only one new mutation was found in the CDR1 region, which was specific to Asian populations (1 allele supported by 10 haplotypes) (**Figure S2, Table S2**). The identification of these specific mutation patterns provides extra evidence that the identified variant nucleotides are not sequencing errors, but genuine allelic variants.

On the contrary, we found that several alleles for the *IGKV* genes (*V1-5, V1-33, V1-39, V3-11, V3-15, V3D-20*) in the IgPdb database suffer from possible SHMs as several new mutations across the reported alleles are concentrated towards the end of the *V* gene (near the CDR3 region). Also, the IMGT alleles mentioned to be false positives by Wang et al., 2008 (31), are too heavily mutated (**Figure S2**, alleles with yellow background) to be considered as genuine germline alleles. Accordingly, we did not find these alleles back in our pmIG database.

Based on the comparisons of different resources we realized that each *IG* germline database has certain unique features as well as disadvantages (**Table 2**). Our database does not contain alleles from the genes that are duplicated, as they are not present in human chromosome GRCh37 assembly. Elegantly, IMGT database has profiled all such genes to its completeness. In contrast to any of the existing databases, we have devised an elegant strategy to exclude SHMs possibly arising in group and OI genes along with the enriched information regarding confidence levels, haplotype and population information.

**Table 2:** Comparison of key features in different databases that profile IG germline alleles.

|  | IMGT | VBASE2 | IgPdb | pmIG |
| --- | --- | --- | --- | --- |
| Completeness of *IGHV* alleles | 138 Complete and 98 Partial | V-region only | V region only | All complete (including leader) |
| Completeness of *IGKV* alleles | 71 Complete and 9 Partial | V-region only | V region only | All complete (including leader) |
| Completeness of *IGLV* alleles | 39 Complete and 35 Partial | V-region only | V region only | All complete (including leader) |
| Presence of *IGHD* and *IGHJ* genes | Present | None | A few | Present |
| Presence of *IGKJ* genes | Present | None | A few | Present |
| Presence of *IGLJ* genes | Present | None | A few | Present |
| Presence of constant genes | Present | None | None | Present |
| RSS | Present | None | None | Present |
| Completeness in terms of alternate duplicated genes | Complete | No (Not all genes present) | No (Not all genes present) | Complete as per availability from GRCh37 (does not include alternate duplicated genes) |
| Strategy for exclusion of SHM | Not reported | Not reported | Not reported | Explicitly reported |
| Confidence Levels | NONE | Class-1, 2, 3 | None | Double layer of confidence levels (Confidence levels with information on group genes and OI genes) |
| Haplotype/Population frequency | None | None | None | Present |

### Novel alleles are ‘maximally two mutations away’ from the known alleles

Frequent new alleles were either a combination of known polymorphisms or gained polymorphisms. Many of the newly detected *IGHV* alleles (AS2/3 category) have no new mutations or only one new previously unreported mutation (**Table 3**), or in other words, the 134 novel *IGHV* alleles each contain a new unique combination of already observed (individual) mutations. 7 *IGHV* alleles gained more than three new mutations which belonged to the lower confidence category AS3. These observations were corroborated in the IgPdb and VBASE2 database, where novel alleles indeed also had one to ten new mutations as compared to the IMGT alleles (note that frequency of alleles is not reported in both databases) (**Figure S2**).

**Table 3:** Number of alleles with count of new mutating positions as compared to the existing databases. Complete sequence includes leader sequence and the *V* region for all the *V* genes/ This is important to realize that 98 *IGHV*, 9 *IGKV* and 35 *IGLV* alleles are partial and do not contain the leader sequence.

|  | *IGHV* |  | *IGKV* |  | *IGLV* |  |
| --- | --- | --- | --- | --- | --- | --- |
| **Mutation Counts** | *Complete Sequence* | *V region only* | *Complete Sequence* | *V region only* | *Complete Sequence* | *V region only* |
| *Zero* | 279 | 312 | 85 | 95 | 68 | 91 |
| *One* | 102 | 72 | 85 | 78 | 102 | 93 |
| *Two* | 21 | 19 | 7 | 4 | 26 | 14 |
| *Three* | 5 | 5 | 1 | 1 | 3 | 2 |
| *Four* | 1 | 1 | 1 | 1 | 1 | 0 |
| *Five* | 1 | 1 | 0 | 0 | 0 | 0 |

The *IGKV* and *IGLV* alleles did show a slightly different pattern as in *IGHV* alleles, i.e. the 85 novel *IGKV* and 102 novel *IGLV* alleles all have at least one new mutation as compared to the ones in known alleles (**Table 3**). However, the position of these new mutations is important, i.e. they can occur in the *V* region or in the leader region (16% *IGHV*, 18% *IGKV* and 25% *IGLV* of total mutating positions). Most of the light chain alleles (9 *IGKV* and 35 *IGLV* alleles) in the current databases are only partial i.e. they do not comprise of leader region. We believe this to be the reason why we observe for the *IGKV* and *IGLV* novel alleles have a new mutation instead of a novel combination of existing mutations.

### Population Distribution of the *IGH* alleles

The first allele of each *IGHV* gene (_01), sorted such that it is supported by the maximum number of haplotypes, is known and present across all superpopulations (denoted by the annotation “ALL”). Approximately 100 new *IGHV* alleles (frequent or rare) are unique to the African populations (denoted as “AFR”) while only 29 African *IGHV* alleles are known (**Figure 4A**). Alleles not observed in Africa (“Non-AFR”) or observed in Africa and other superpopulations but not all (“AFR Shared”) occur less (∼40% less). Together, this suggests that Africans have a huge diversity that has not been captured so far.

*IGHV1-69* (**Figures 4B**), *IGHV2-70*, *IGHV3-53* and *IGHV3-23* (**Figures S3**) genes have the highest allelic diversity in the African superpopulation. *IGHV3-48* alleles are highly diverse in South Asians (**Figure S3**), whereas new alleles in the *IGHV4* family are highly diverse in Africans and East Asian populations (**Figure S4**). Similarly, *IGHV7-81* alleles mostly belong to the African and American superpopulations (**Figure S4**). These differential diversities suggest an environmental adaptability or population drift of these *IGHV* genes.

Most of the *IGHD* alleles are present in all populations and only a few are private to (super-)populations. Even rare variants are shared among different ethnicities, which hints that the *IGHD* genes are evolutionary conserved across populations. Also, all *IGHJ* alleles are shared between all the ethnicities (**Figure 4A**).

The constant genes of the *IG* loci are responsible for the effector functions of the antibodies and have been considered to be more conserved as compared to the *V*, *D*, *J* genes. In contrast, we found 196 alleles for the nine *IGHC* genes. As *IGHA1,2* and *IGHG1,2,3,4* genes are highly conserved, also visible from the alignment of the alleles within these groups based on their CH1-3 domains (**Figure S6**), we grouped the alleles within these two groups (group alleles, Methods). This resulted in that only 125 alleles were retained, where the majority of the alleles that were filtered out are from the *IGHA1*, *IGHG1*, *IGHG3* and *IGHG4* genes (**Table S2, S5**).

To further understand the contrast between the supposedly conserved constant genes and the many alleles found, we converted the nucleotide sequences of the *IGHC* alleles into protein sequences and mapped them to the known allotype sequences. This shows that multiple mutations in the allelic sequences for *IGHC* genes are synonymous and, therefore, the diversity at the protein level is quite low as compared to the nucleotide level (**Table S2, Figure S6**). This suggests a high evolutionary pressure on the *IGH* constant genes to conserve the structural and functional properties of the Ig proteins.

Three new allotypes were identified for the IgA1, IgE and IgD proteins, respectively, all specific to African populations (**Figure S6**). The amino acid change in the IgG1 allotypes did not result in a change of either structural properties (aliphatic <-> aromatic) or the charge (neutral <-> negative <-> positive).

### Population Distribution of the *IGK* alleles

76 *IGKV* alleles are present in all superpopulations, of which 62 alleles are already known (**Figure 5A**). Only 10 of the 41 African alleles map to known alleles. A large number of alleles is observed outside Africa (“Non-AFR”), suggesting that the diversity in the *IGK* locus does not only prevail in the African superpopulation (**Figure 5B, S7**). All *IGKJ* alleles are shared between ethnicities except one allele that was unique to Africans. Of the two major *IGKC* alleles, one is present in all superpopulations and one is unique to Africans. A third rare allele is present in Europeans and South Asians.

### Population Distribution of the *IGL* alleles

The population distribution of the *IGL* alleles is similar to that of the *IGH* locus. Most of the known *IGLV* alleles are present in all superpopulations. The majority of the new alleles in both the frequent and rare alleles is unique to African populations (**Figure 6A**). Similarly, known *IGLJ* alleles are present in all the superpopulations, whereas rare *IGLJ* alleles are either unique to African populations or are not observed in the African population (“Non-AFR”). The first *IGLC* alleles (_01) are present in all superpopulations, except *IGLC2_01* that is unique to Africans. New rare *IGLC* alleles belong to either the African populations or populations outside Africa (“Non-AFR”). Of the new frequent *IGLC* alleles, only a few are unique to European and South Asian populations, and the majority exists in African populations (**Figure 6B, S8**).

### Recombinant Signal Sequence (RSS) variants in *IG* (pseudo)genes influence their recombination frequencies

RSS regulates the recombination process in which the conservation of RSS heptamers and nonamers play a significant role (15,52,53). We did not find variations in conserved heptamers and nonamers of allelic RSSs that may explain population-specific recombination frequency of the respective genes. Also the conservation of RSSs in *IGHV* genes was reported to be related to differences in recombination frequencies (29, 54). We found that the heptamers and nonamers in all *IGHV* RSSs are conserved, except the ones related to *IGHV3-16* and *IGHV7-81*. Interestingly, a relatively lower recombination frequency of *IGHV7-81* has been reported before (29).

Several *IGHD* genes have mutated heptamer sequences at 3’D-RS and 5’D-RS (**Figure 7, Table S6**), which might explain their reported reduced recombination frequencies in healthy individuals (29). All *V* genes in *IGK* and *IGL* locus have conserved heptamers, except the *IGKV1D-13* and *IGKV2D-30* genes in the *IGK* locus (**Table S7**), and the *IGLV5-48*, *IGLV2-33*, *IGLV3-22*, *IGLV3-19* and *IGLV2-14* genes in the *IGL* locus (**Table S8**). These genes have mutations in the first three bases of their heptamers, which consequently should result in less efficient (=reduced) recombination frequencies (15,52,53).

**Figure 7:**
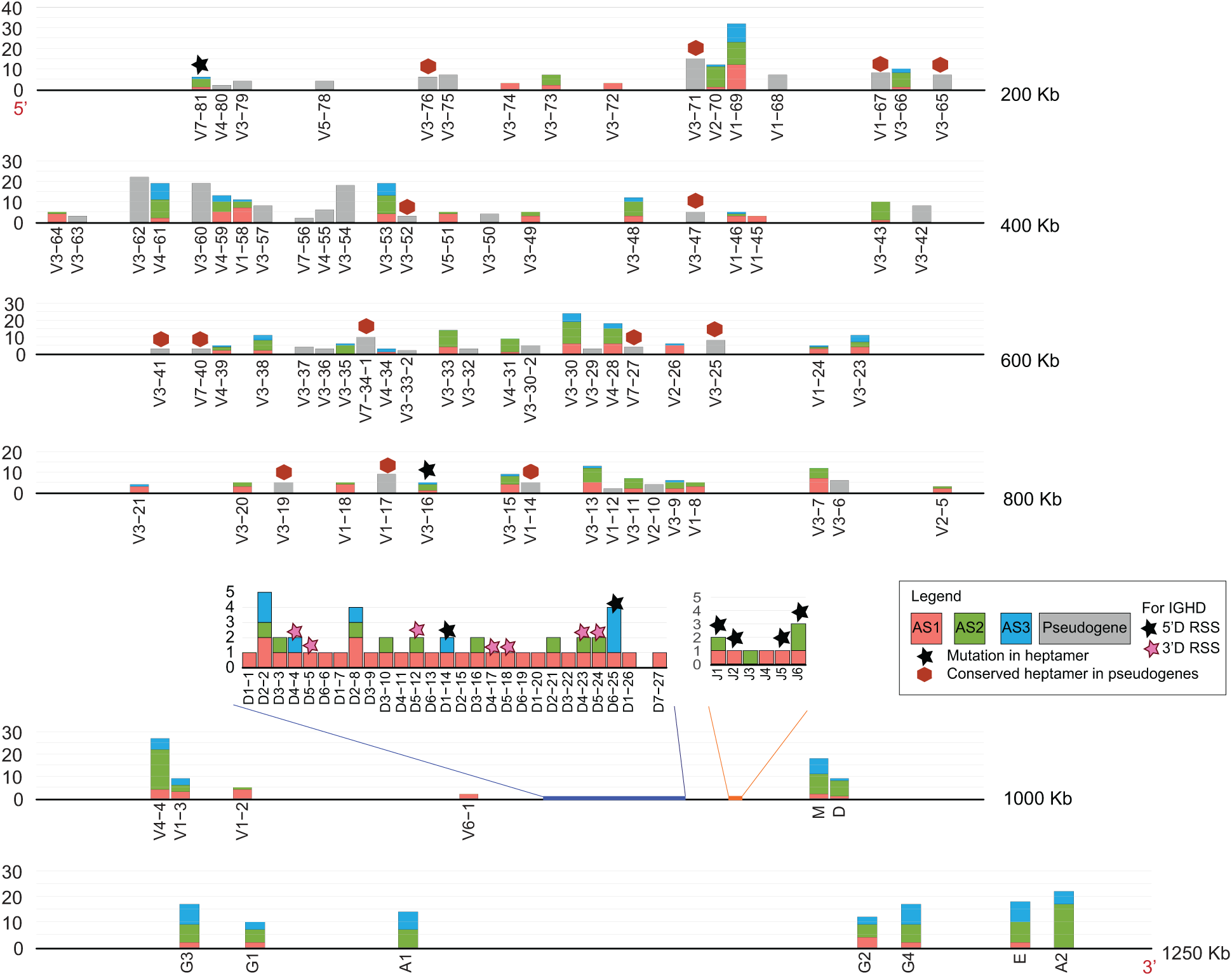
*IGH* alleles per gene with confidence levels in the locus representation. The alleles are marked on the locus with a confidence level AS1 (red), AS2 (green) and AS3 (blue), where the bar height represents the number of alleles. Pseudogenes are represented by grey colored bars. The Y-axis on the locus shows the absolute number of alleles. *IGHD* and *IGHJ* are projected outwards. A star indicates that the RSS sequence is mutated and it could have effect on the recombination frequency of these genes. *D* have stars colored with black and blue which indicates the mutation in 5’ RSS and 3’ RSS, respectively. A red hexagon on the pseudogenes indicates the presence of conserved heptamers.

The RSS spacer length also plays a role in the recombination frequency (11,43,55). We found that the spacer length in RSSs of most *IGHJ* genes is 22bp, except for the *IGHJ3* and *IGHJ4* genes that have a spacer of 23bp. In addition, *IGHJ4* has the most conserved heptamers followed by *IGHJ6* (**Table S6**). These observations could explain why the *IGHJ4* and *IGHJ6* genes have the maximum recombination frequency among *IGHJ* genes (56).

We also found conserved RSS heptamers adjacent to fifteen and eight pseudogenes in the heavy and light *IG* loci, respectively (**Table S9**). The location of the RSS in few of these pseudogenes is 10-30 bases more distant from the *V* pseudogene boundaries than for regular *IGHV* genes in which the RSS is generally 0-3 bases adjacent to the *V* gene boundary. Also, five *IGLV* functional genes had RSS sequence 10-25 bases downstream to the gene (**Table S8**). We found that only *IGLV3-7* (pseudogene) has a stop codon in between RSS heptamer and *V* gene boundary which could impact the recombination frequency of this gene. The impact of distance between RSS and gene boundaries is not yet known.

### Relation of *IG* alleles to variable immune responses in populations

Although we acknowledge that the immune repertoire is individual-specific, the efficiency of the response in different populations can be driven by the germline allelic variants. Therefore, we set out to understanding the diversity in immune responses to infections or diseases by investigating the allele distribution in the *IG* loci. To do so, we annotated alleles with their impact on human health based on their polymorphisms and whether they contain at least one known disease-associated variant (based on a literature search with keyword “*IG* gene name + disease/vaccine”). We then checked the frequencies of the disease alleles across the different ethnicities. Here, we report four examples that show different frequencies across the different (super-)populations.

**EXAMPLE-1:** The *IGKC* gene mutation rs232230 (C->G) results in a nonsynonymous variant (V->L) that is a risk factor in both gastric cancer and breast cancer (odds ratio 1.64 and 1.94, respectively) as well as *Helicobacter pylori* infection in gastric cancer and age in breast cancer (57). The IMGT allele *IGKC*04* is equal to the same variant, and we found this allele to be present in 1,066 Haplotypes of the G1K samples. The distribution of the alleles in different populations was not known before, but we found the allele to be evenly distributed across all populations with a median of 38 (Min: 7 – Max: 90) haplotypes (**Figure 8A**).

**Figure 8:**
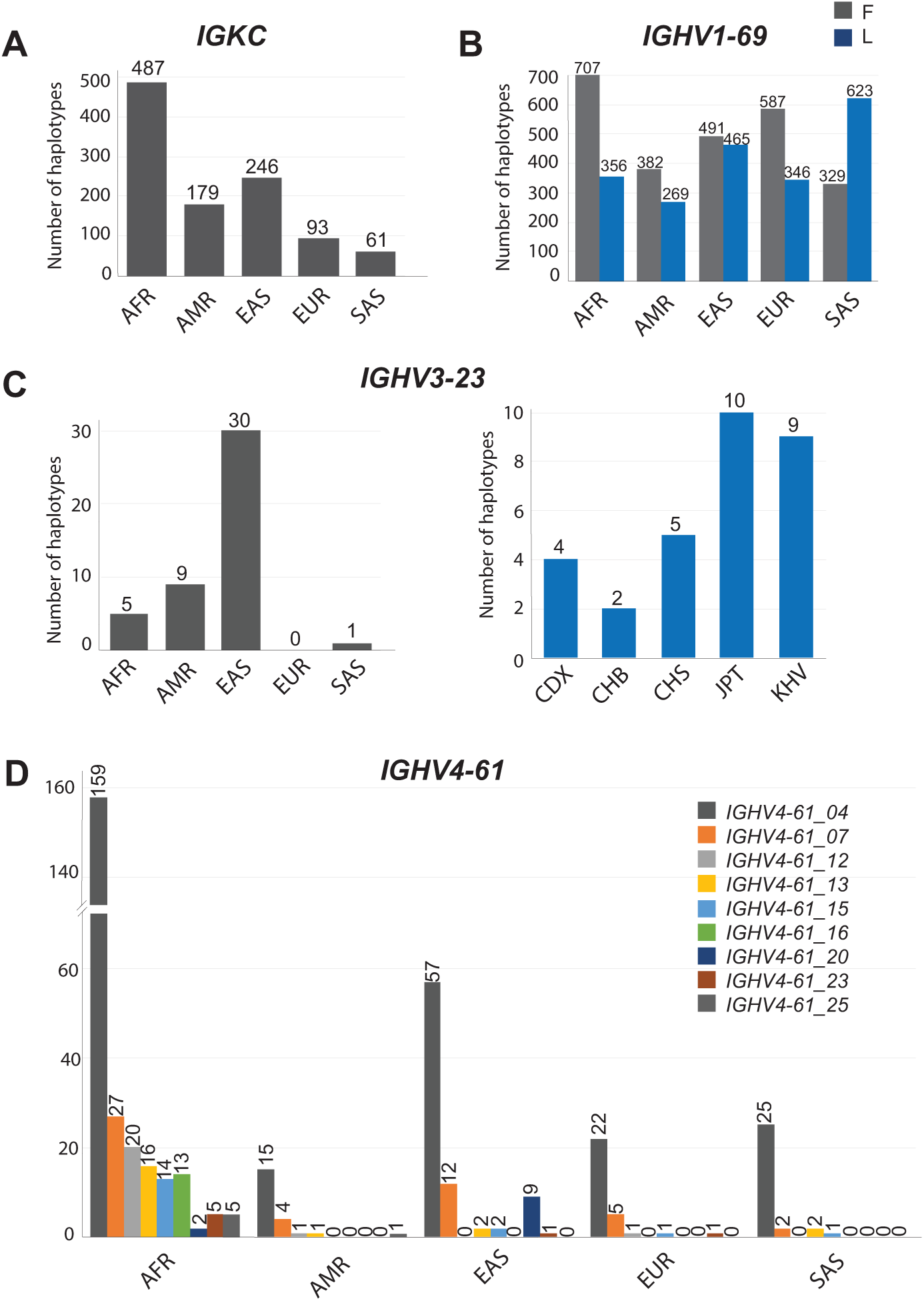
Frequency of *IG* alleles associated with immune responses in populations. **A) *IGKC***: One allele supported by 1066 haplotypes is distributed in all the superpopulations. **B) *IGHV1-69*:** All the alleles were divided into two groups i.e. one with F “Phe” in CDR2 region and other with “L”. The combined distribution of the two groups is represented in the figure. **C) *IGHV3-23*:** One allele is supported by 45 haplotypes. The plot on the left represents the superpopulation distribution of the allele. The right plot represents the population distribution of the allele in the East Asian population. CDX: Chinese Dai in Xishuangbanna, China; CHB: Han Chinese in Beijing, China; CHS: Han Chinese South; JPT: Japanese in Tokyo, Japan; KHV: Kinh in Ho Chi Minh City, Vietnam **D) *IGHV4-61*:** Nine alleles i.e. *IGHV4-61_04, IGHV4-61_07, IGHV4-61_12, IGHV4-61_13, IGHV4-61_15, IGHV4-61_16, IGHV4-61_20, IGHV4-61_23* and *IGHV4-61_25* were found to have the mutations in combination with other mutations. Separate bar plots are drawn for each allele.

**EXAMPLE-2:** The F->L polymorphism in in the CDR2 region of *IGHV1-69* gene is known to have a potential role in modulating the anti-influenza antibody repertoire (18). We not only detected a high diversity of the *IGHV1-69* gene in the African (super-)populations that bear the amino-acid “Phe: F” polymorphism (10 AFR alleles with F variant whereas only 5 with L variant) but also that the F variant is overrepresented in African (super-)populations (**Figure 8B** grey bars). For Non-African populations, we found that F and L amino-acid mutations occur in equal ratios, except for South Asians where most alleles have the “Leu: L” polymorphism (**Figure 8B** blue bars). These findings are concordant with those of Avnir et al. (18). The understanding of the population distribution of this polymorphism in *IGHV1-69* gene and its role in flu, can have an immediate implication in the implementation of “influenza” vaccines in different superpopulations.

**EXAMPLE-3:** The *IGHV3-23*03* IMGT allele is known to be four-fold more effective than *IGHV3-23*01* against *Haemophilus influenza* type b (Hib) (20). Recent studies suggest that meningitis caused by Hib is a common and serious disease in children in China (21, 58). We observed that the *IGHV3-23*03* allele is very rare and is present frequently only in the East Asian superpopulations. Only 30 haplotypes support this allele in the East Asian (super-)populations (CDX:4, CHB:2, CHS:5, JPT:10, KHV:9) (30 haplotypes) (**Figure 8C**).

**EXAMPLE-4:** The *IGHV4-61*02* IMGT allele is related to higher risk of rheumatic heart disease (RHD) in Oceanic populations where four polymorphisms (rs201076896, rs201691548, rs200931578 and rs202166511) increase the susceptibility (59). In this study, the relationship was drawn only within the Oceanic populations. Therefore, we profiled the alleles carrying these four mutating positions in our population matched database which comprises five different ethnicities. We found nine alleles comprising of these four mutating positions and their frequency was highest in African populations followed by Asian populations (**Figure 8D)**. This might suggest that RHD is more frequent in African populations as compared to Asian populations.

### Evolutionary dynamics of variation patterns in different populations

The genetic diversity of individual genes does not reflect on the diversity of the complete *IG* loci including coding and non-coding regions. Therefore, we used the SNPs in the complete locus to identify the existing variations between (super-)populations. In **Figure 9A** we found African populations to be unique and highly diverse for *IGH* and *IGL* loci, whereas the *IGK* loci are much more condensed due to some large outlier samples in African and American populations.

**Figure 9:**
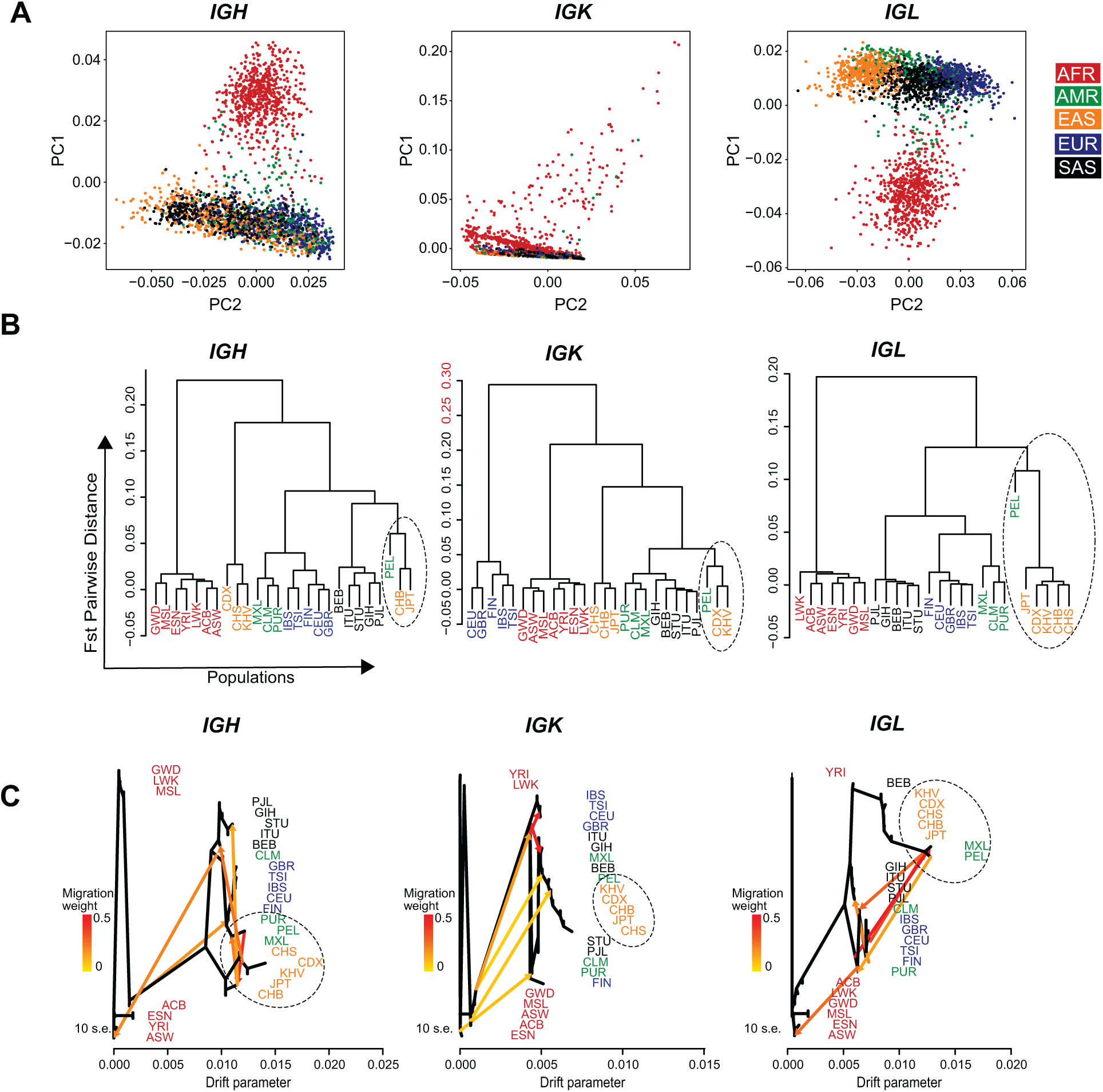
Genetic diversity, population structure and migration events in five superpopulations for *IG* loci. A) Separate PCA plot of the heavy and light chains of *IG* loci based on the single-nucleotide polymorphisms in the complete locus. Each dot represents a sample and each sample is colored based on the superpopulation they belong. In *IGH* and *IGL* locus Africans have higher diversity as compared to Non-African superpopulations, while *IGK* has more diversity in part of the European population. **B)** Pairwise population distribution calculated by Fst Matrix is represented as a cladogram for each locus namely *IGH*, *IGK* and *IGL*. 26 populations are colored as per the superpopulations *i.e.* Africans in red; Americans in green; East Asians in orange; Europeans in blue and South Asians in black. **C)** Migration events in *IGH*, *IGK*, and *IGL* locus. Six migration events are marked in the ML tree where edge color represents the migration weight; red suggest higher migration weight and yellow the lowest.

To completely capture the variability of the locus, we looked at the pairwise population differentiation (F_ST_) in the *IG* loci. The *IGH* and *IGL* loci again point towards higher diversity in African populations (F_ST_ ∼0.20; **Figure 9B**). Interestingly, we found a different pattern in the *IGK* locus that has highest variability within Europeans (Fst ∼0.30) rather than Africans (**Figure 9B).** The lower diversity of the *IGK* locus in Africans is consistently visible in the population distribution of the alleles (**Figures 4, S6**). This might suggest that the variability in this locus is more recent in evolution, as compared to the *IGH* and *IGL*.

Furthermore, the cladogram (**Figure 9B**, marked by dotted circles) reveals a closer relationship between the Peruvian (PEL) population and East Asian populations, especially the Han Chinese in Beijing in China (CHB), and Japanese in Tokyo (JPT) in each *IG* locus, suggestive of a mixture between the Peruvian and East Asian populations. To obtain insight into the migration patterns between populations, we plotted the tree for 26 populations from the G1K resource where horizontal branch lengths are proportional to the amount of genetic drift that has occurred on the branch (**Figure 9C**). Six migration links are plotted with a directed connection between populations wherein orange colored connections depict higher migration rate. We indeed observed the highest migration weights between the Peruvian and East Asian populations in all loci, supporting the mixture hypothesis (**Figure 9C**, marked by dotted circle).

## Discussion

We performed an extensive analysis of *IG* germline alleles from 2,504 individuals, representing 26 populations, and created a population matched *IG* germline database (pmIG), that comprises a comprehensive overview of haplotypes across five main different ethnicities. We enriched pmIG by including information on rare and frequent germline alleles per population, facilitating identification of genuine germline alleles and excluding SHM. This will be important when studying differences between populations in immune responses as a consequence of germline differences (7,42,60–64).

Similar to the Lym1K resource by *Yu et al*., we have used the 1,000 Genomes dataset to derive all *IG* alleles. But there are important differences. We report information on haplotype support, including a minimal support of seven haplotypes (four haplotypes for known alleles), categorized them into confidence levels, scrutinized each allele manually for their use in repertoire studies, and also profiled population information for each allele. As a result, we report ten times less alleles than the ones reported by Lym1K. It is fair to say that the “alleleminer tool” (used to create Lym1k) has a cutoff option for the minimal haplotype support. However, to identify potential sources of non-germline errors, the processes of B-cell development and SHM-based affinity maturation of antibodies should be understood carefully. For example, the existence of duplicated genes reduces the confidence in associated alleles. Therefore, we adopted a manual curation of alleles and assigned confidence levels to each allele. Currently, alleles are categorized into three different confidence levels based on the observed haplotype support and further grouping based on the duplications of *IG* genes. Additional support from sanger sequencing of the alleles from independent sources, can further increase the confidence level of the pmIG database.

One confidence level, the AS1 level, indicates whether the alleles that we detected are previously reported in other databases. Aligning alleles with tools like Muscle (38) is, however, not trivial, but a manual check was also performed to ensure accuracy. We found that the known alleles that we detect (and thus present in existing databases) are mainly present in the European-Caucasian populations, suggesting a population bias in the current IMGT, VBASE2, and IgPdb databases.

The largest numbers of novel frequent and rare alleles are identified in African populations of which 70-90% of diversity is not captured by the existing databases. Several studies analyzing genome wide patterns, genetic variations, demographic history and immune responses have also reported higher immune diversity in Africans (39,61,65–70). The sampling of more individuals from multiple African populations can further reveal the genetic diversity in Africa and can thereby substantiate our understanding of allelic diversity in *IG* loci. The recent completion of the African reference genome (69) can put this divergence in a different perspective, particularly when compared with the more narrow diversity of the European reference genomes (71, 72).

The accuracy of our novel alleles is supported by the highly conserved mutation patterns, restricted to a limited number of nucleotide positions, remarkably the same as found in other *IG* databases. This also allowed us to identify possible false-positive alleles in the IMGT and IgPdb databases. Our findings were consistent with a previous study that indicated that 104 IMGT alleles are false-positives (31). We believe the reliability of our pmIG database is mostly due to the stringent threshold for novel alleles to have been seen in at least 4 individuals (7 haplotypes).

Different germline alleles have shown to result in different responses and effectiveness against infections in individuals (18,20,32). The population information of these alleles can provide better understanding of these response differences at the population level. For example, the *IGHV3-23*03* IMGT allele is known to be four-fold more effective than *IGHV3-23*01* against *Haemophilus influenza* type b (Hib) (20). The *IGHV3-23*03* IMGT allele is supported by only 30 haplotypes and frequent in East Asian populations. This allele might a potential therapeutic antibody for meningitis caused by Hib in China (21, 58). Similarly, the under-representation of *IGHV1-69* alleles in European populations and their higher copy number in Africans might explain why European populations are more prone to pandemic flu outbreaks as compared to African populations. Together, these examples underpin the significant role of germline alleles in different populations and their protective nature against infection with the consequent potential impact of population-specific therapeutic antibodies. Detailed studies on differences in clinical disease course and final outcome in different regions of the world during pandemic outbreaks, such as the currently ongoing SARS-CoV2 pandemic, might at least in part show a role of the here presented diversity of *IG* gene alleles within and between human populations.

VBASE2 and IgPdb do not report pseudogenes or rearrangeable pseudogenes. With the extensive genomic information from the G1K resource, we could identify conserved heptamer sequences adjacent to these pseudogenes. Interestingly, these conserved heptamers were 10-30 bases juxtaposed to the pseudogene boundaries. Also, a similar pattern was observed in a few *IGLV* genes. The pseudogenes are reported to be rearranged in several repertoire analysis studies (29, 73), which also relates to our unique above-mentioned finding. This suggests a possible role for the position of RSSs on the recombination frequency of these genes.

The high variability in the *IGH* and *IGL* loci in African populations, as compared to Non-African superpopulations, could indicate the migration of human populations out of Africa based on *IG* loci. On the other hand, the *IGK* locus represents higher variability in European populations, as compared to Non-African populations. We did not find many new alleles in the *IGK* locus from G1K, which hints to current databases being biased towards a sampling from mostly European populations. Pairwise differences between alleles in the *IGK* locus show that Non-European populations are closer to each other than to the European population. Although we cannot follow the trend of mutations over time, the migration analysis and the alleles statistics support the variability and environmental adaptation of this locus over time.

The IMGT database is the most used database in research because of its completeness. Also, the database hosts several tools to support researchers in analyzing and understanding next generation sequencing data of *IG* loci. With our pmIG database of *IG* alleles, we report identified and curated alleles across 5 superpopulations, containing 26 populations, resulting in 170% more allelic variation. Having a richer source of *IG* alleles improves the interpretation of repertoire sequences. For example, determining whether an observed sequence belongs to a germline allele or is the result of SHM in response to an antigen. Or, whether measured sequences of naive B cells are considered to be the result of a sequencing error, which now is determined based on the presence of the observed allele in databases (74). Alternatively, pmIG can be exploited for applications in immune response dynamic analysis and clonal lineage analysis. Perhaps the most clinically relevant application of pmIG is understanding differences between populations and to help implementation of population-specific vaccination studies.

## Acknowledgements

We are grateful to Ms. W.M. Bitter for her high-quality figures.

We acknowledge 1,000 Genomes project for making the data publicly available. As we have used a specific locus of the chromosomes 2, 14 and 22, we have complied with the 1,000 Genomes policies for the publication of data and the submission of alleles to GenBank.

The authors would like to thank the Project MinE GWAS Consortium to provide us access to the variant data for *IG* loci for 1,007 healthy individuals from the Netherlands.

## Authors contributions statement

Concept and design of the study: IK, MAB, EBvdA, CIT, MJTR, and JJMvD

Data acquisition, data analysis and organization of the database: IK

Wrote the manuscript and designed the figures: IK, MAB, MJTR, and JJMvD

Manuscript revisions and approval of the submitted version: All authors

## Availability of data

The full set of alleles (AS1-3) and all R and python scripts used for analysis are available from GitHub (https://github.com/InduKhatri/pmIG). The alleles are available in three different files, according to the confidence level. Also, the gapped sequences for the pmIG alleles as per IMGT standards are available to be used as a reference allele set for Rep-Seq data analysis, that can be directly implemented as database in IgBLAST (75).

---

**Figure S1:**
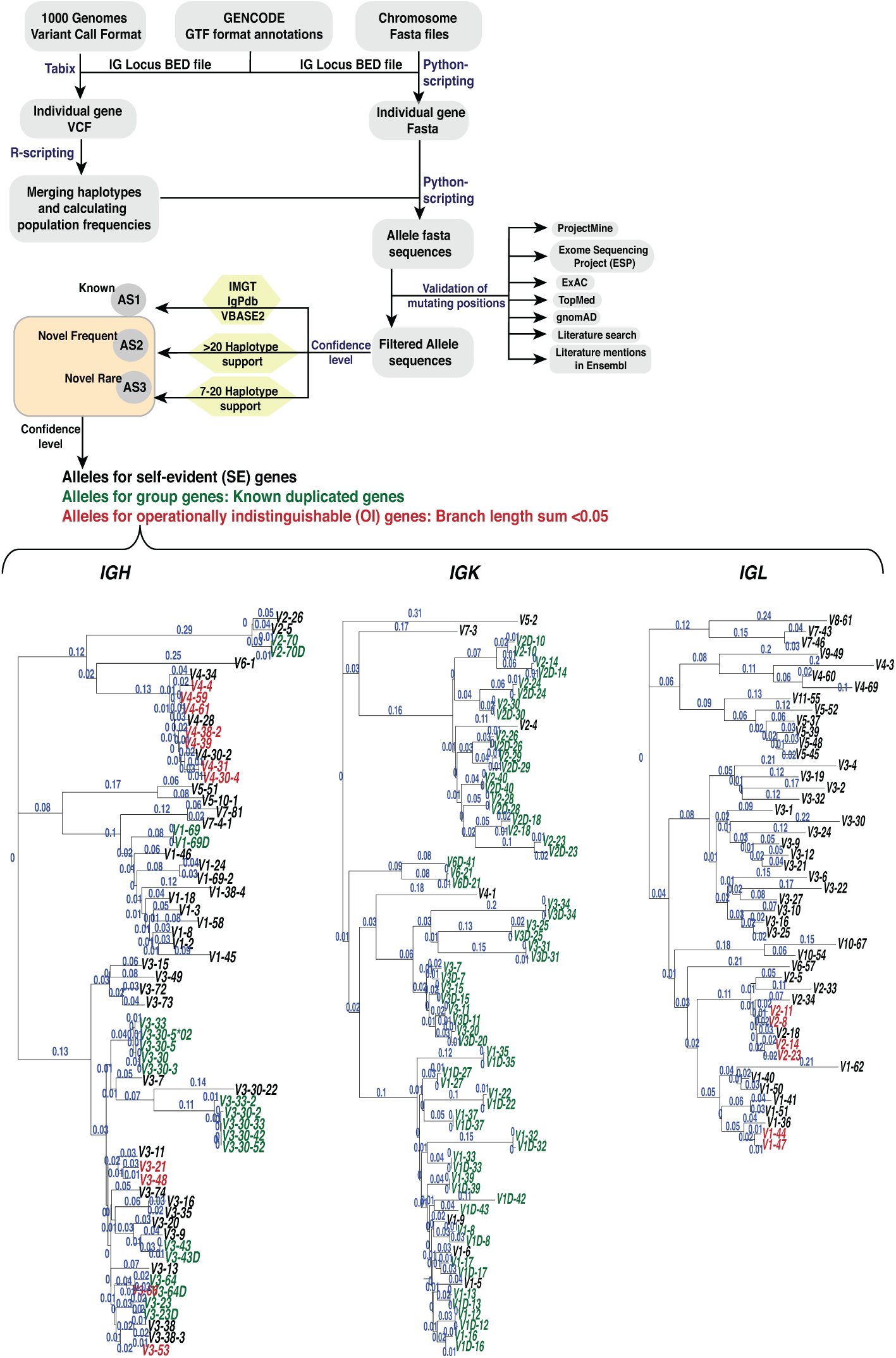
**The pipeline developed to call alleles and assign confidence levels.** The assigned confidence levels AS1-3 were identified based on known (AS1) and novel (AS2 and AS3) alleles. The novel alleles were further divided into self-evident, group and OI alleles highlighted in black, green and red colors, respectively on the NJ trees for *IGHV*, *IGKV* and *IGLV*. All the genes were obtained from GRCh37 version of human genomes. The genes absent in the human genome assembly were retrieved from the IMGT database.

**Figure S2:**
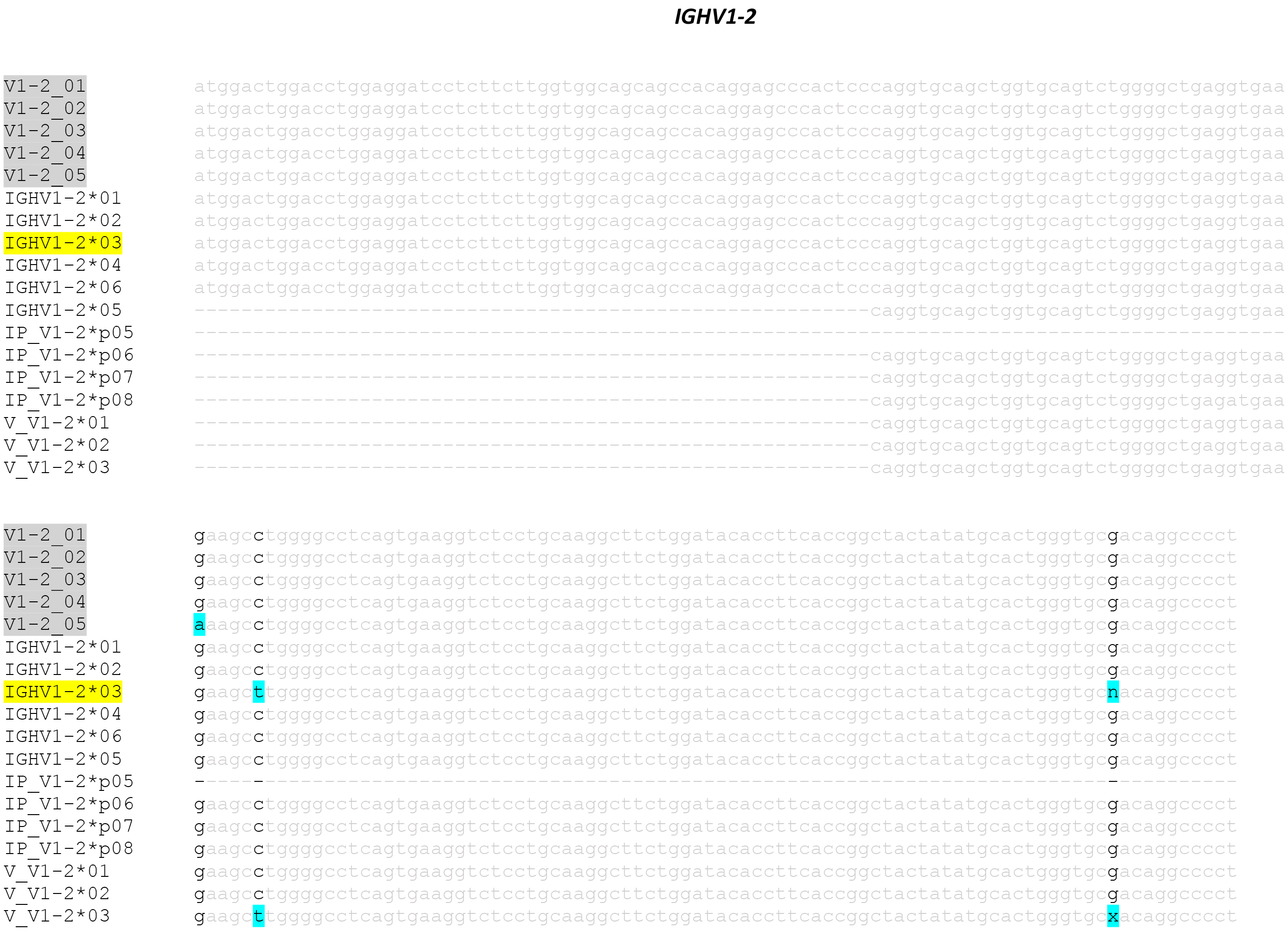
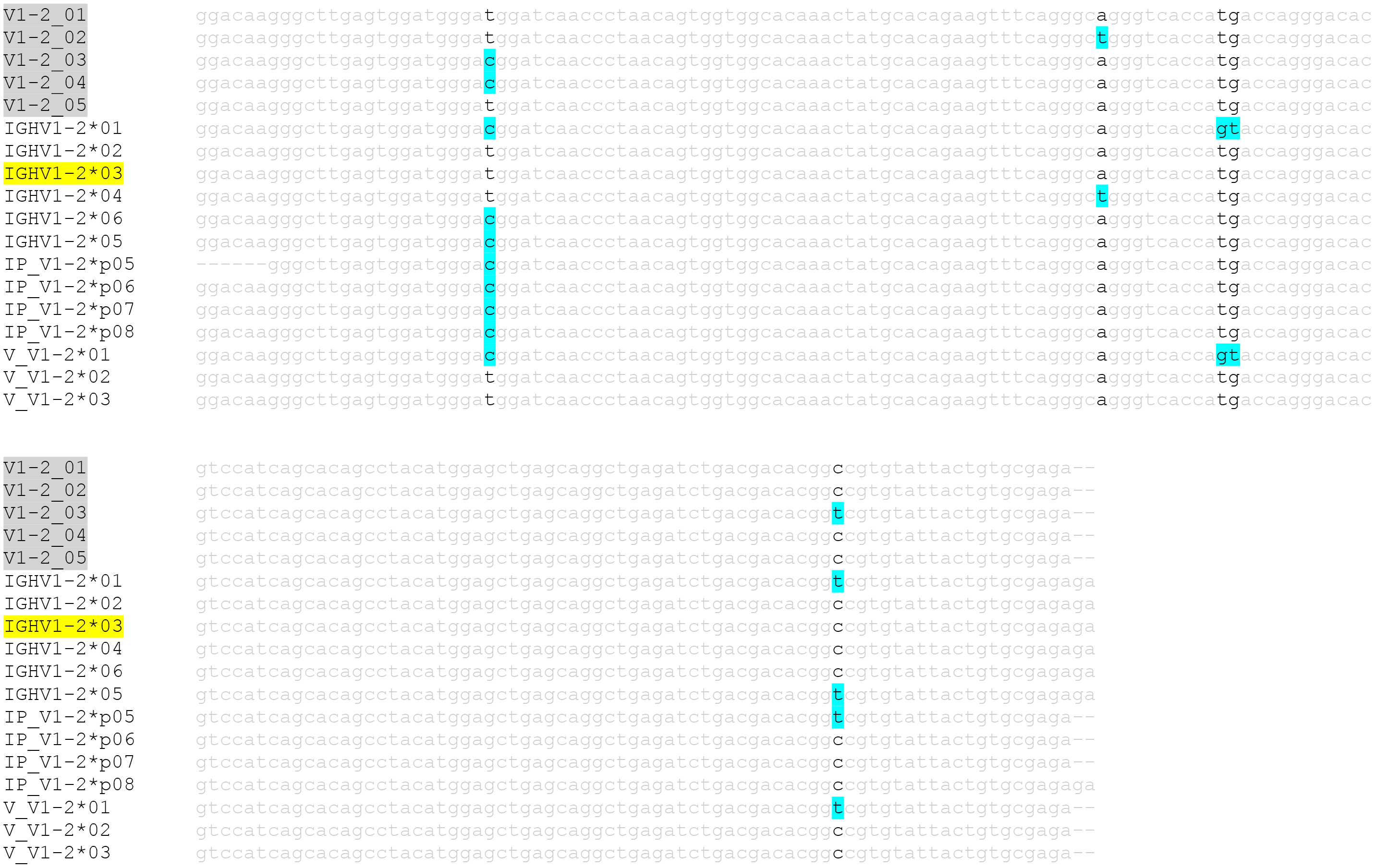
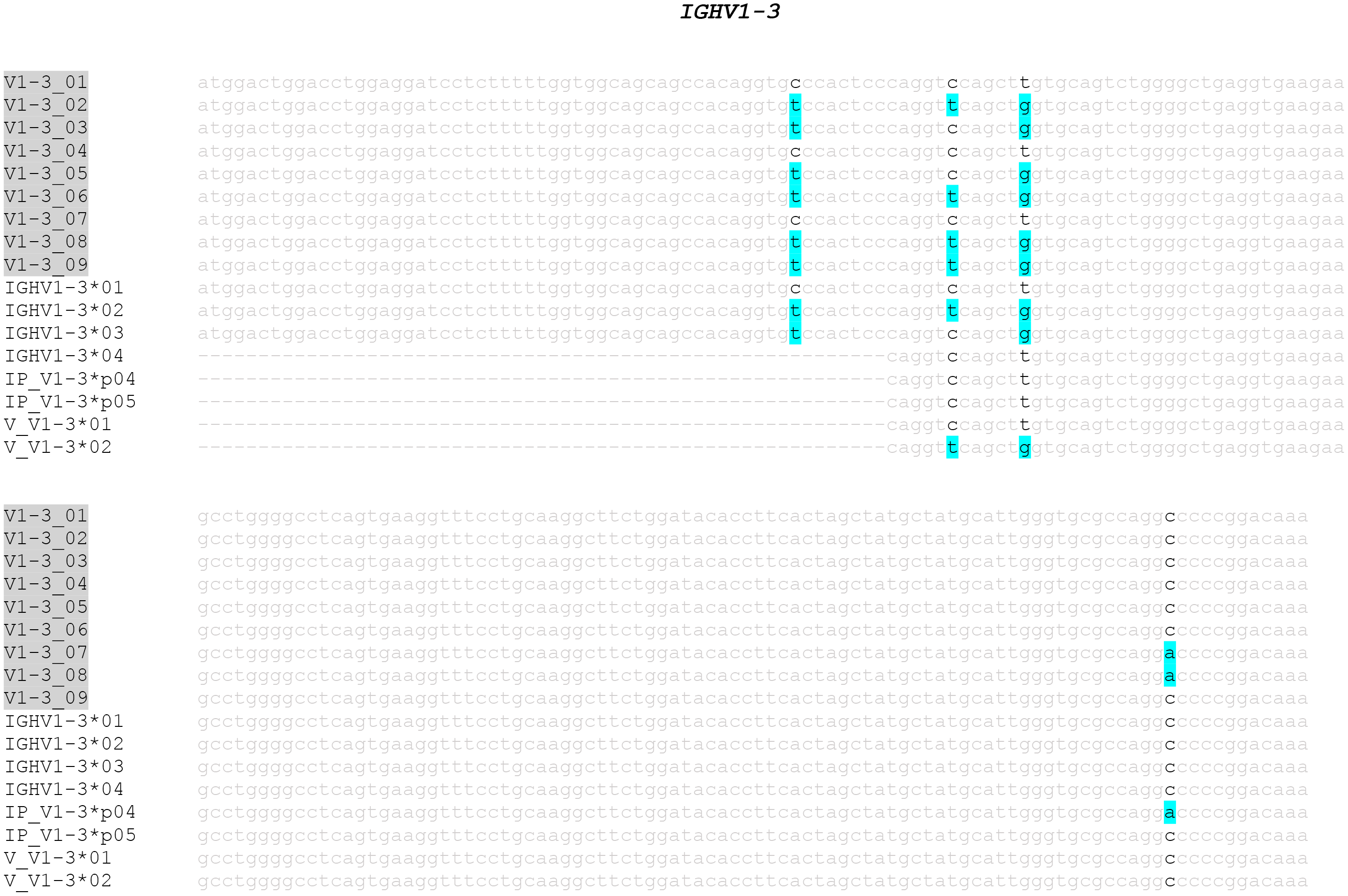
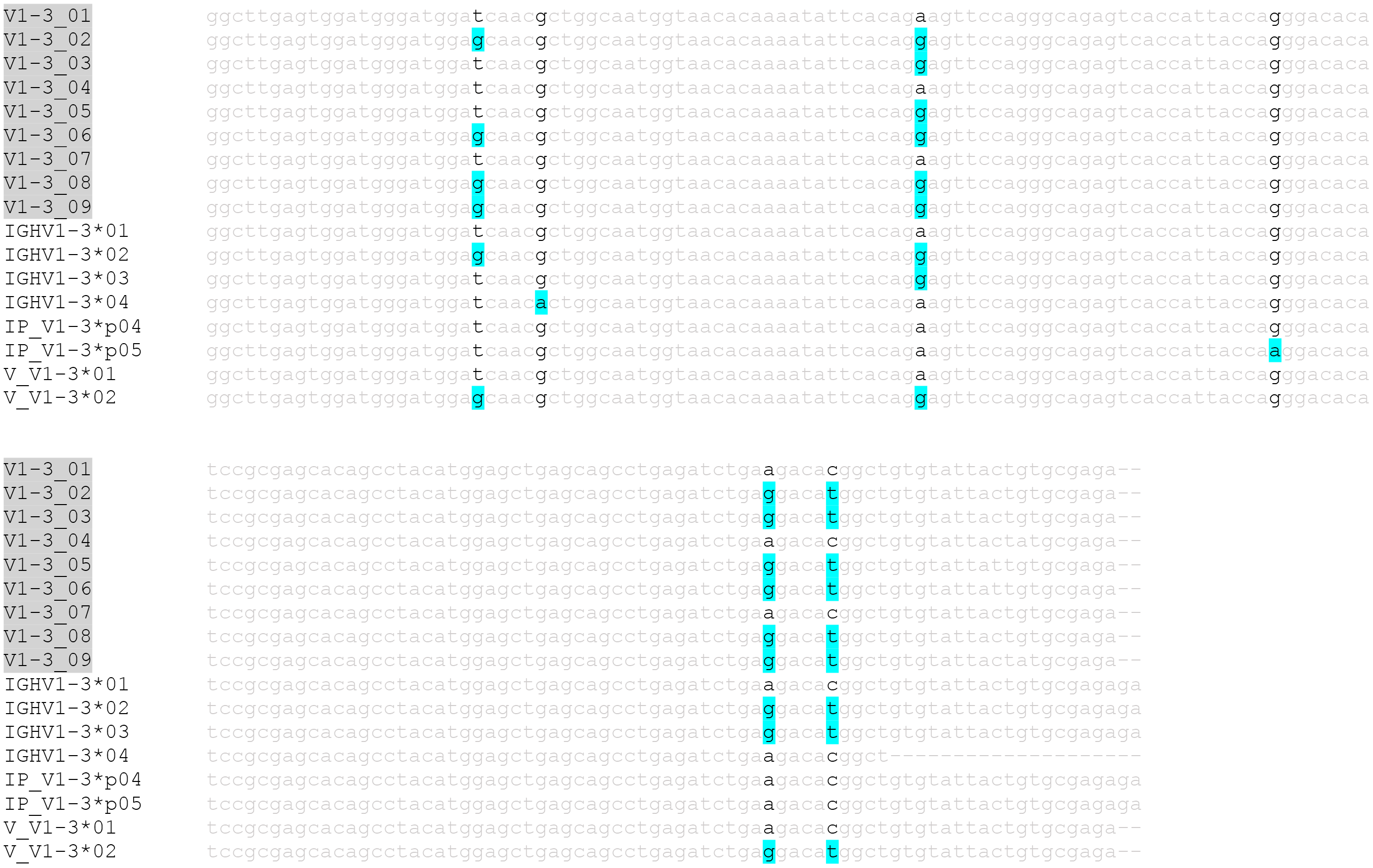
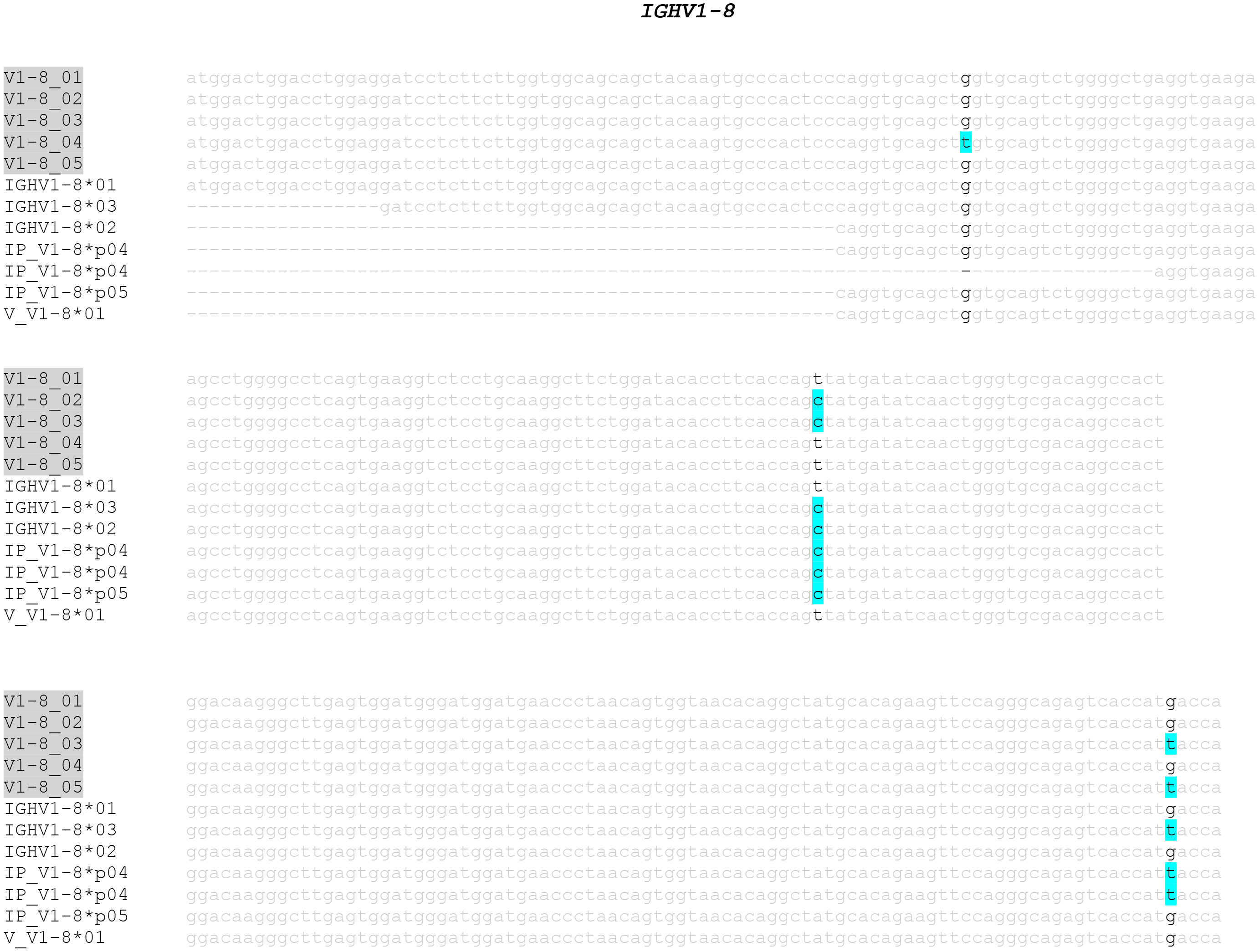
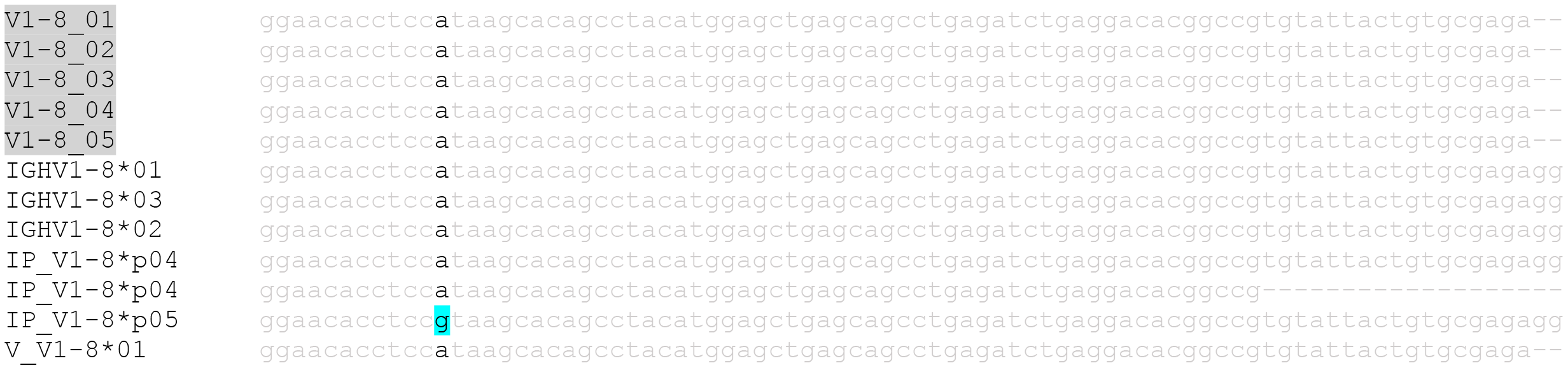
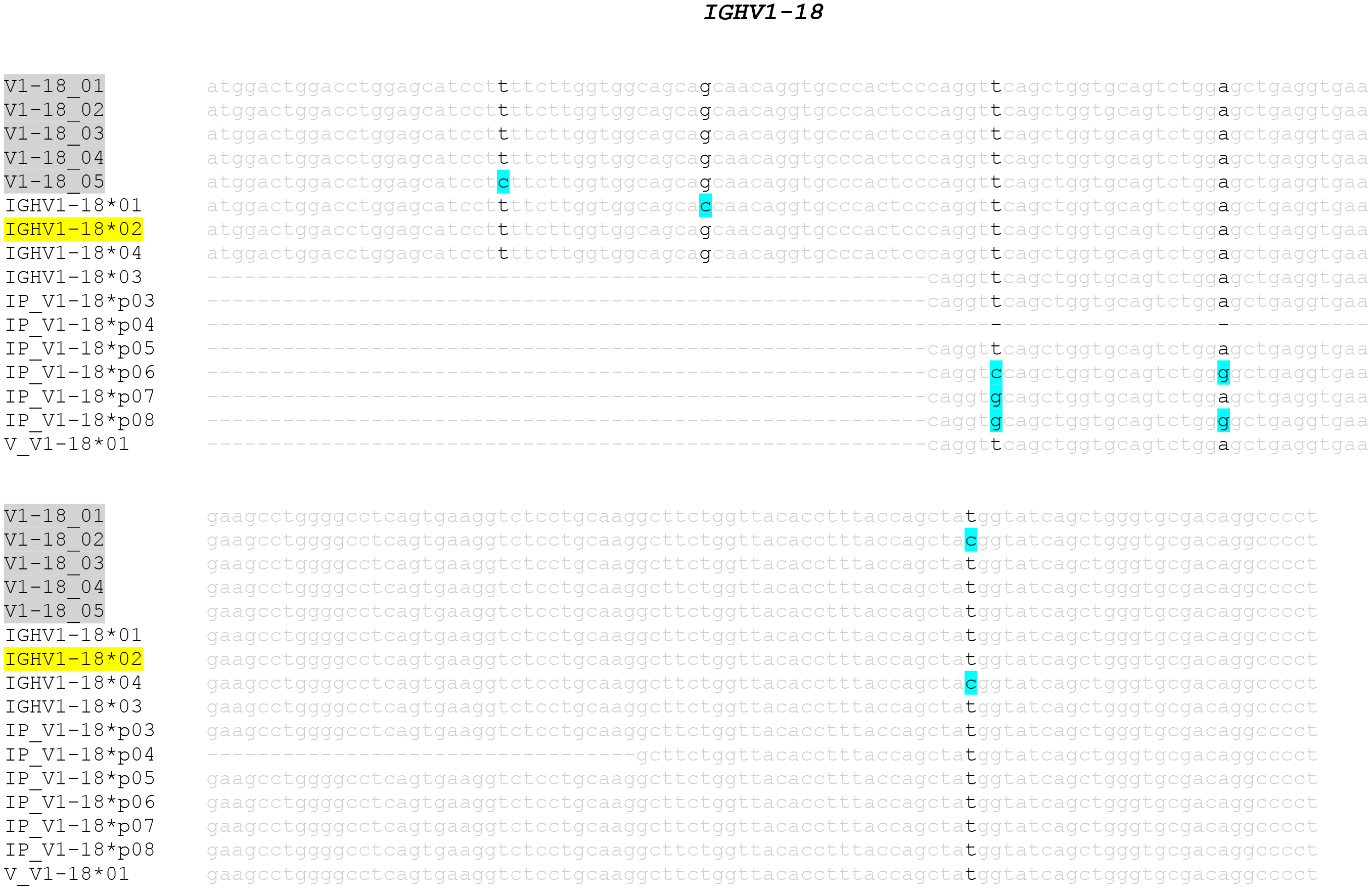
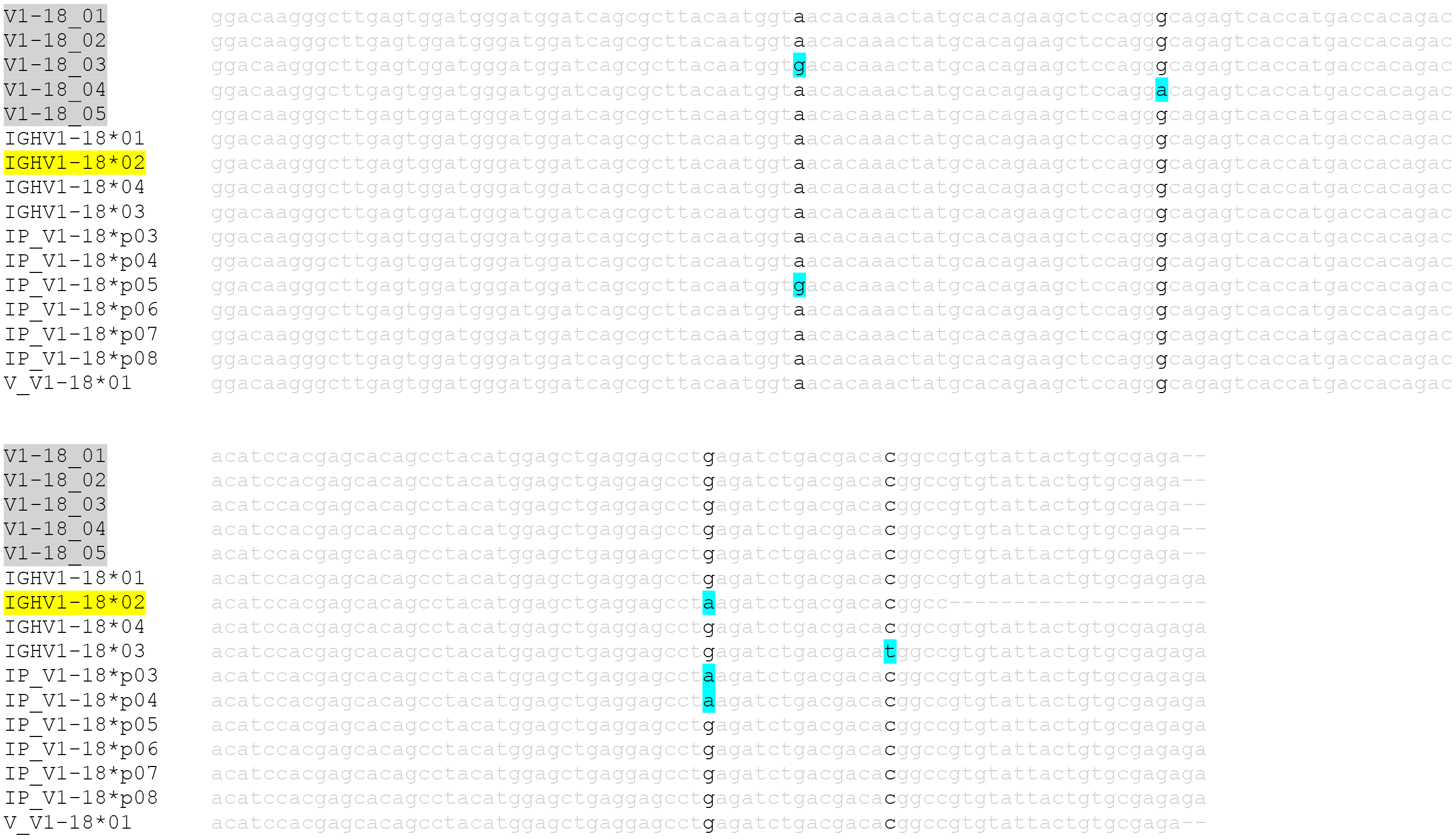
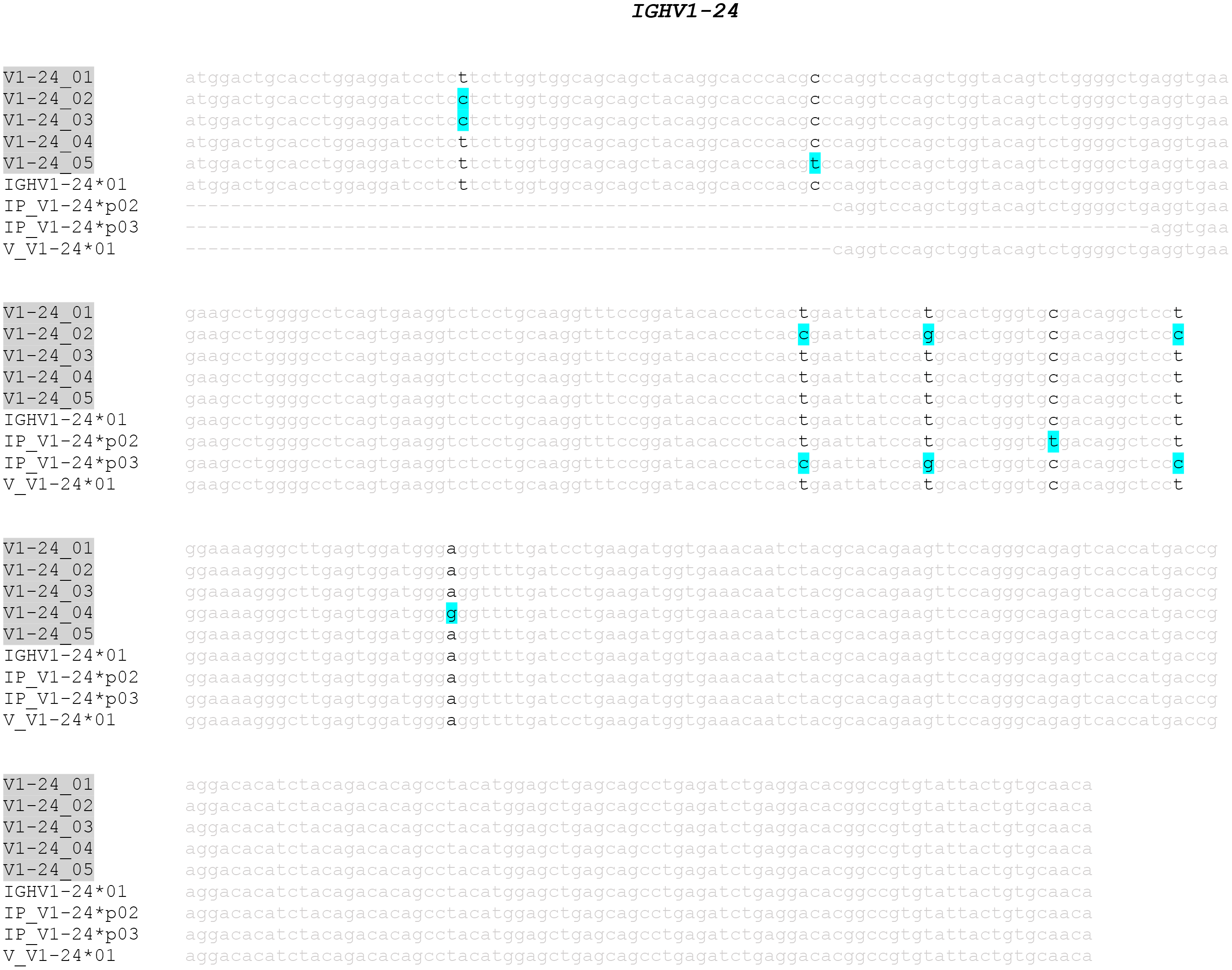
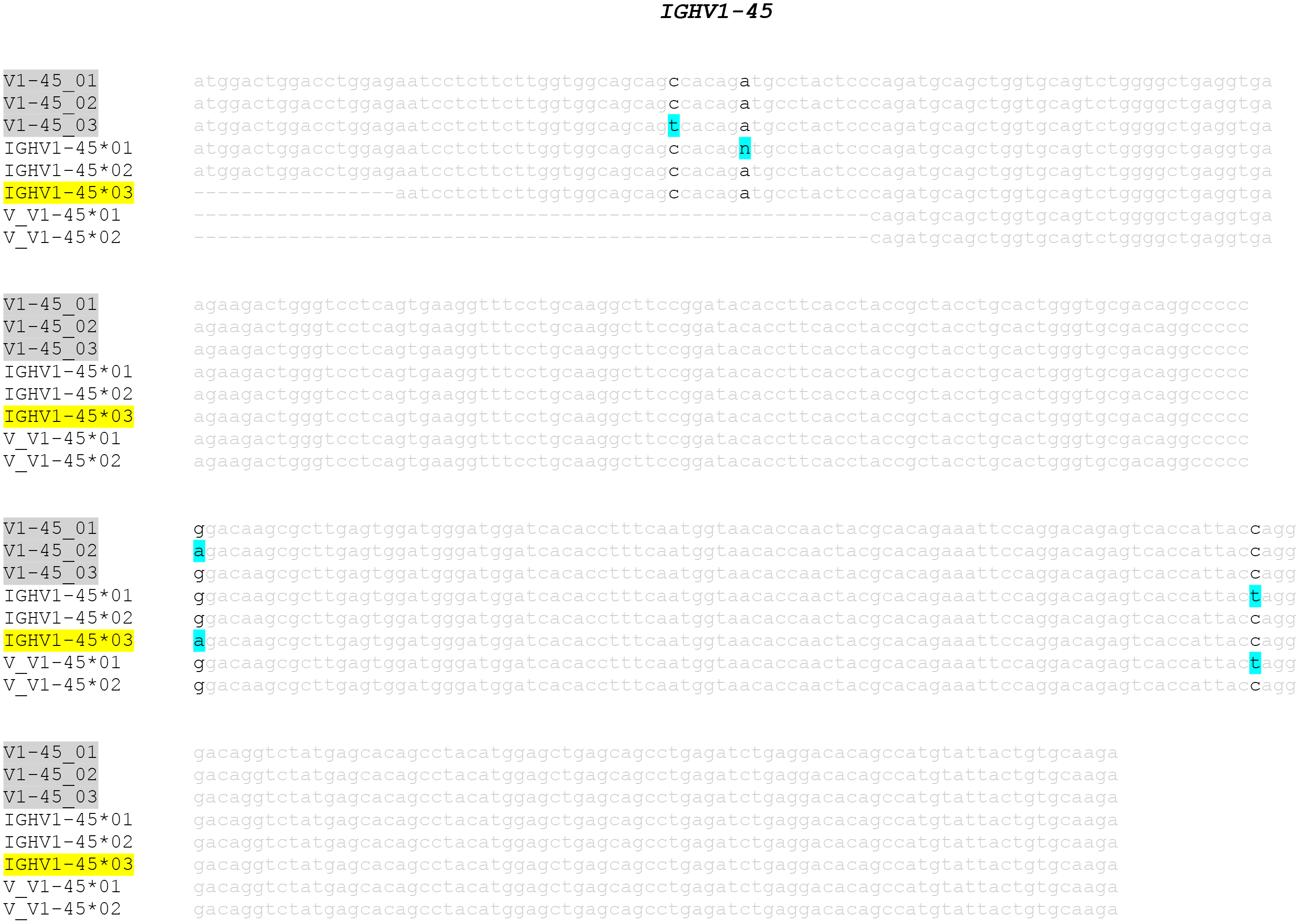
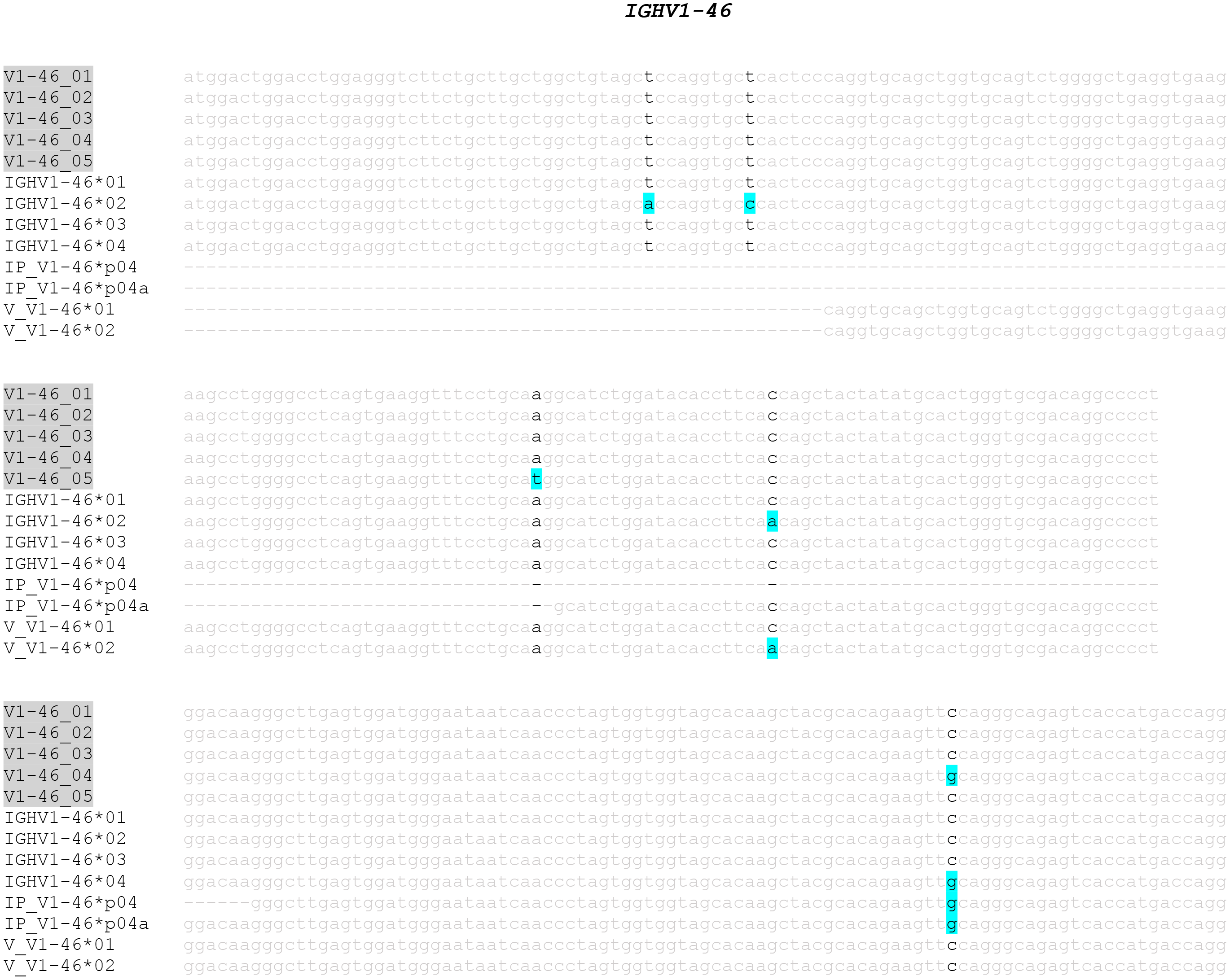
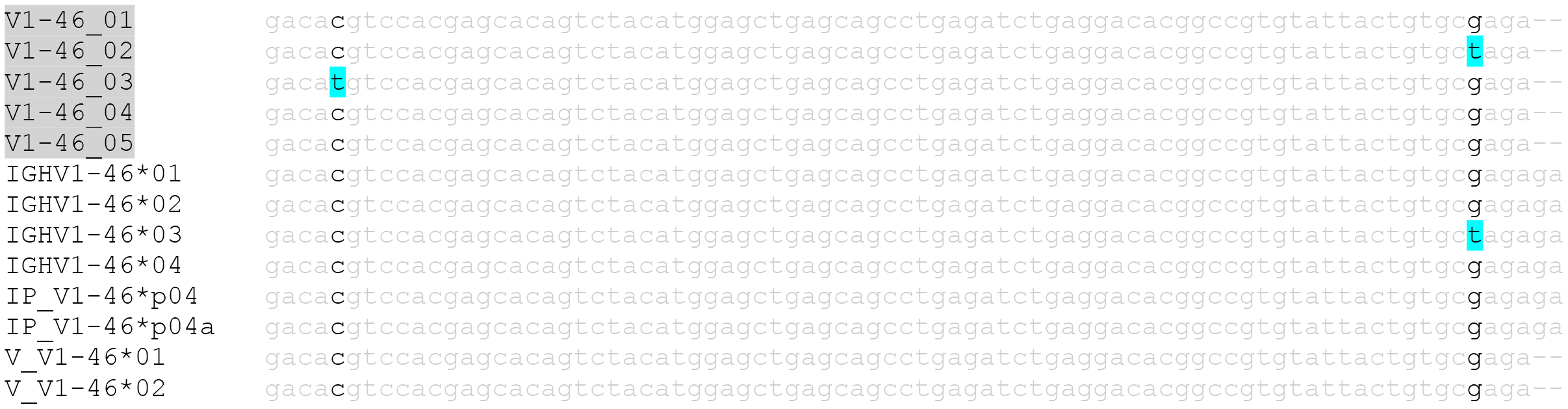
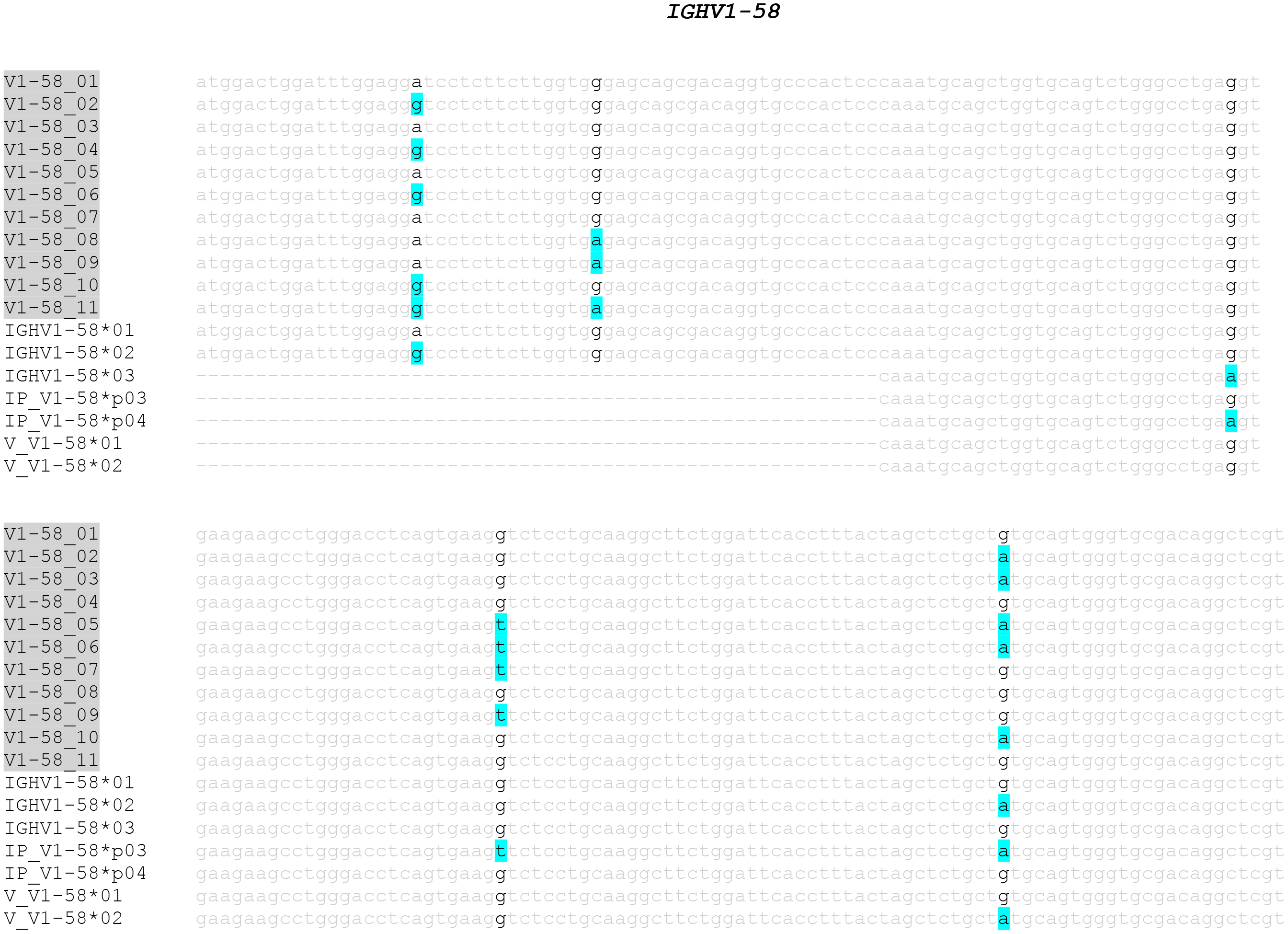
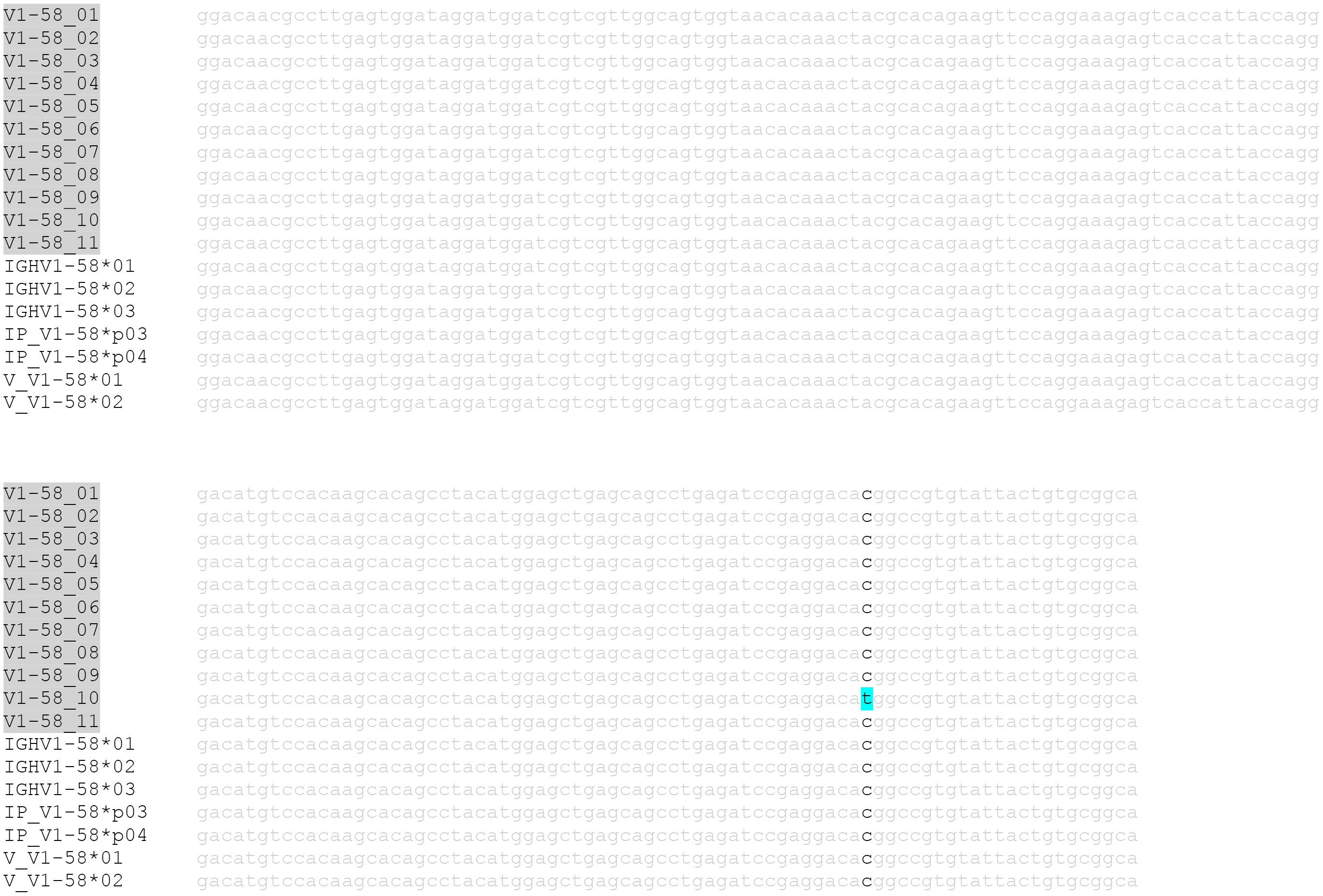
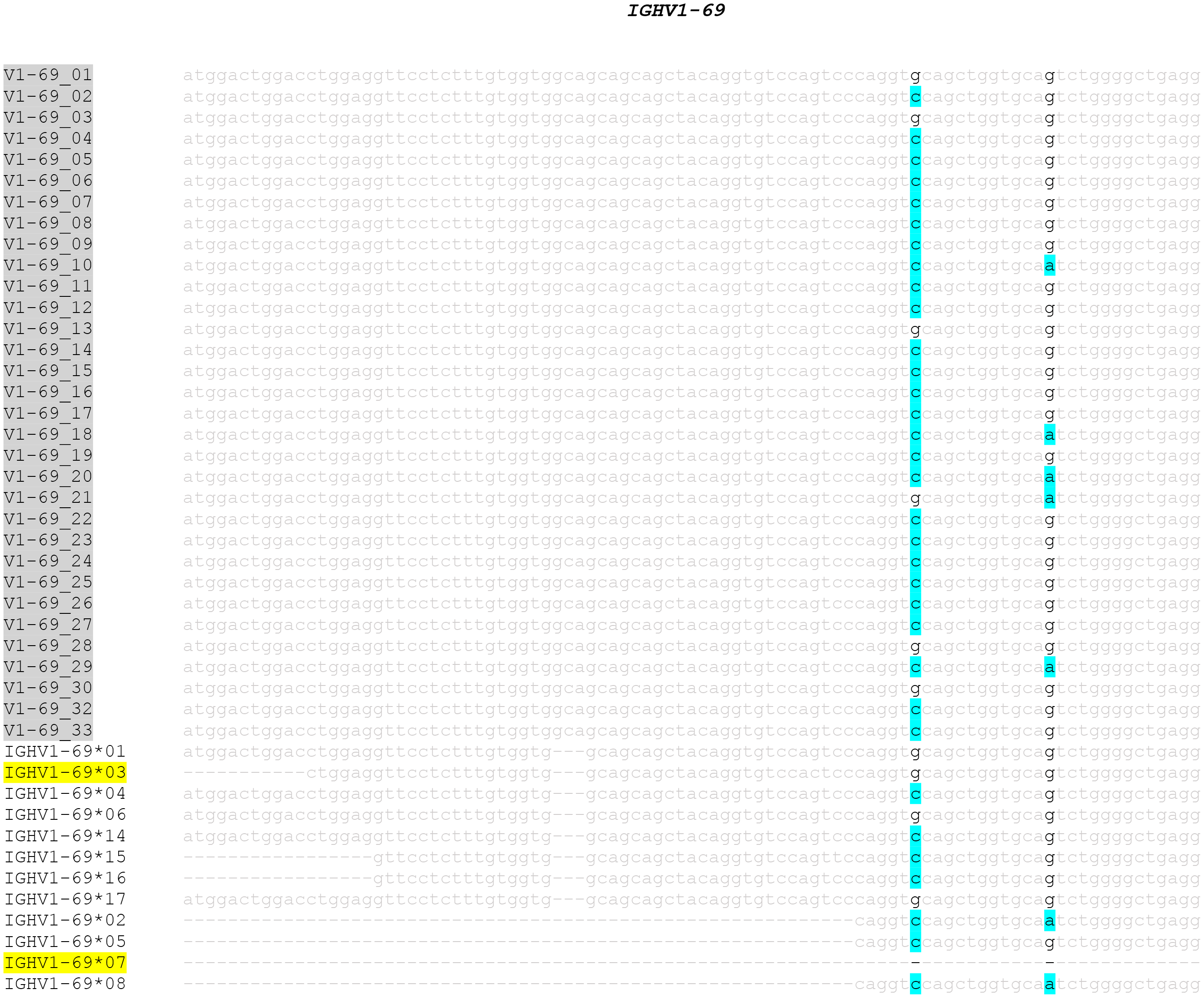
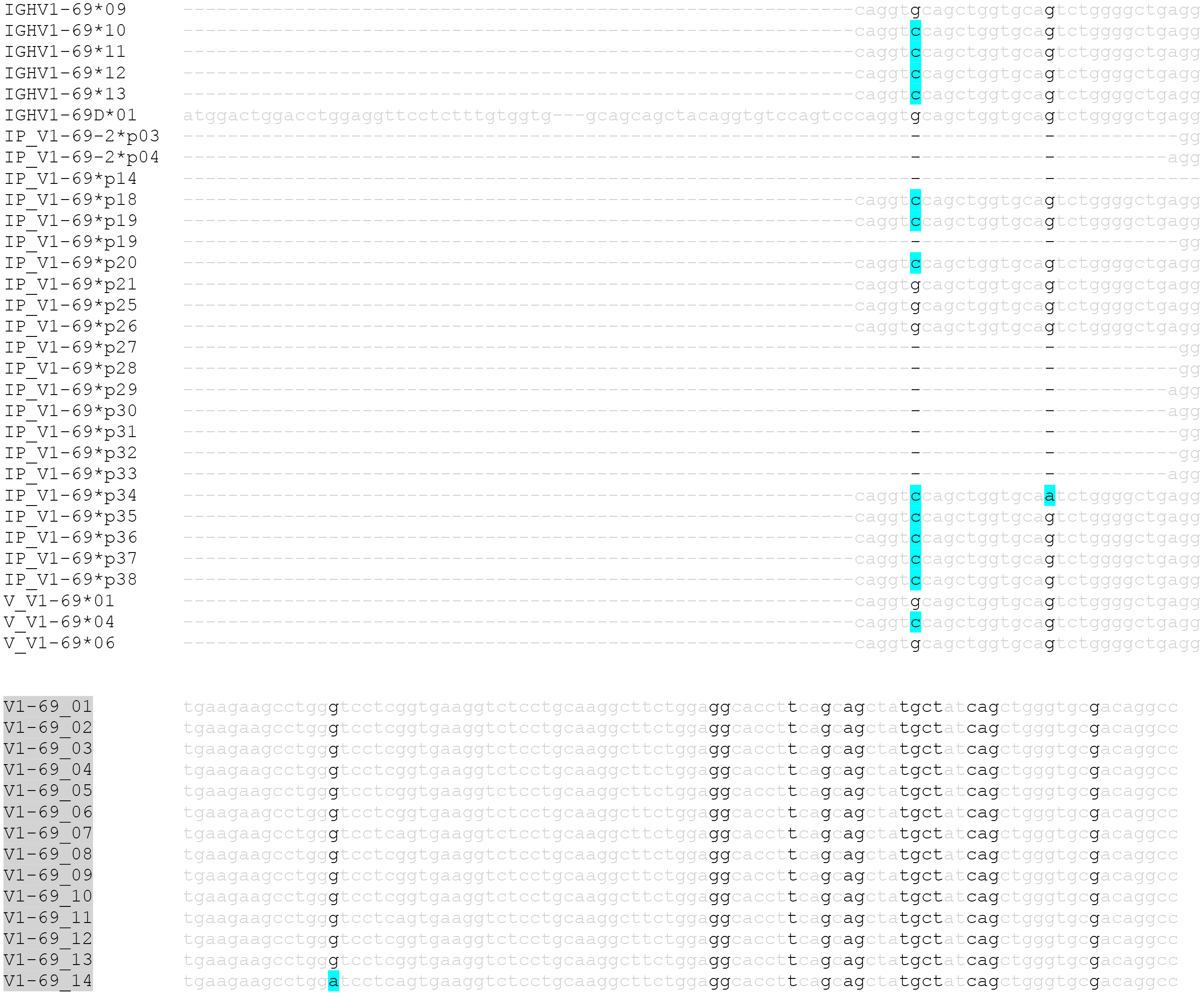
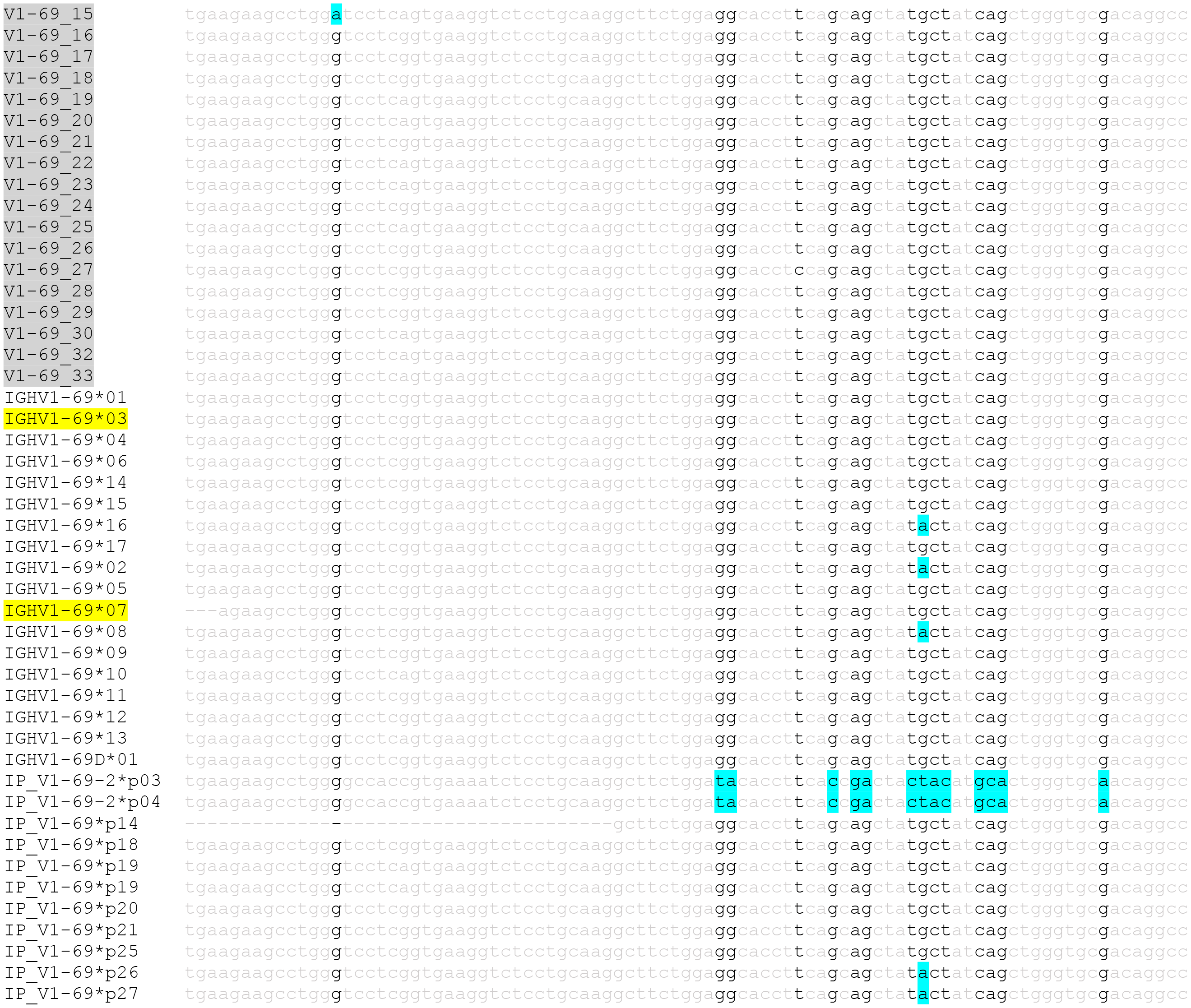
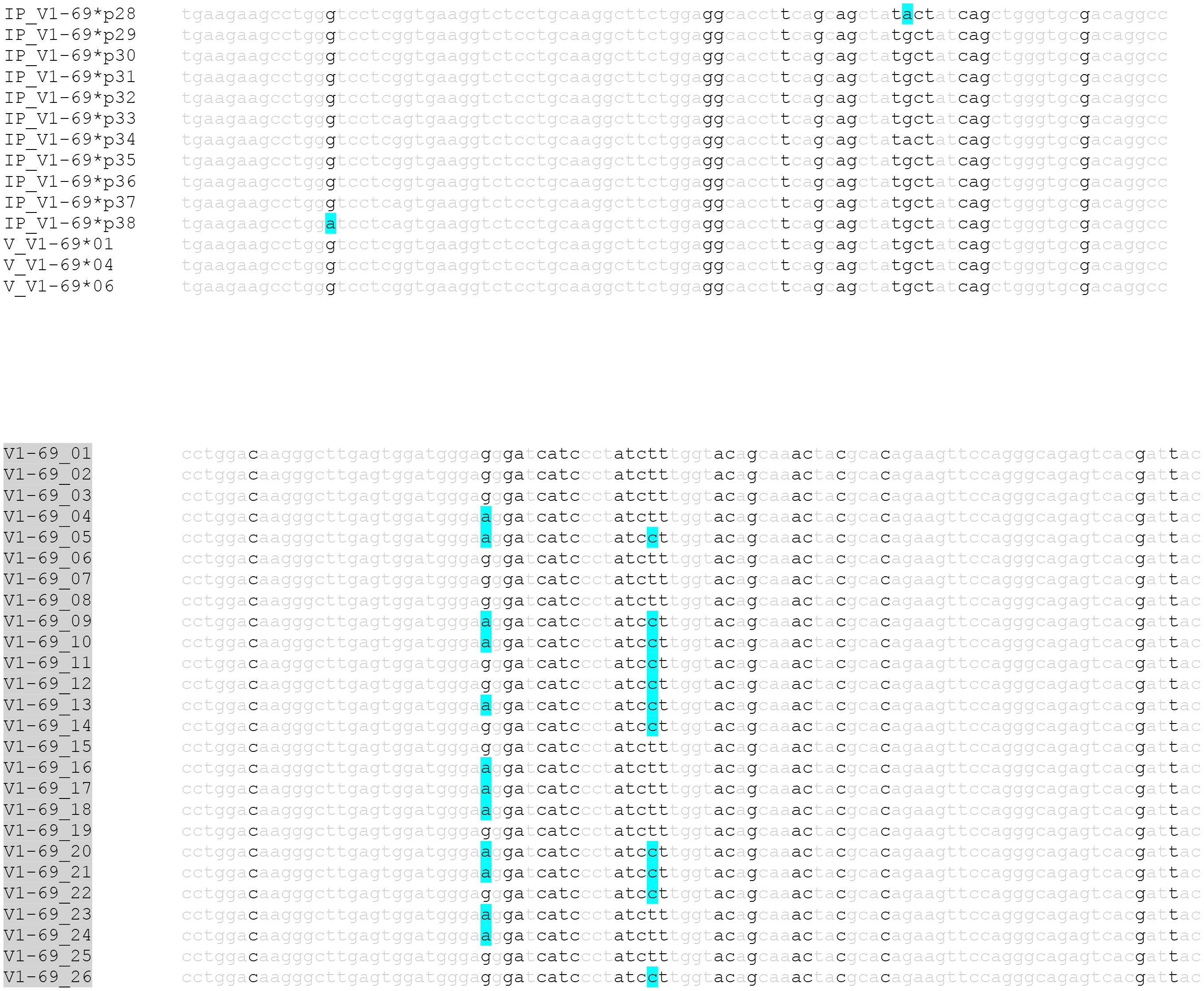
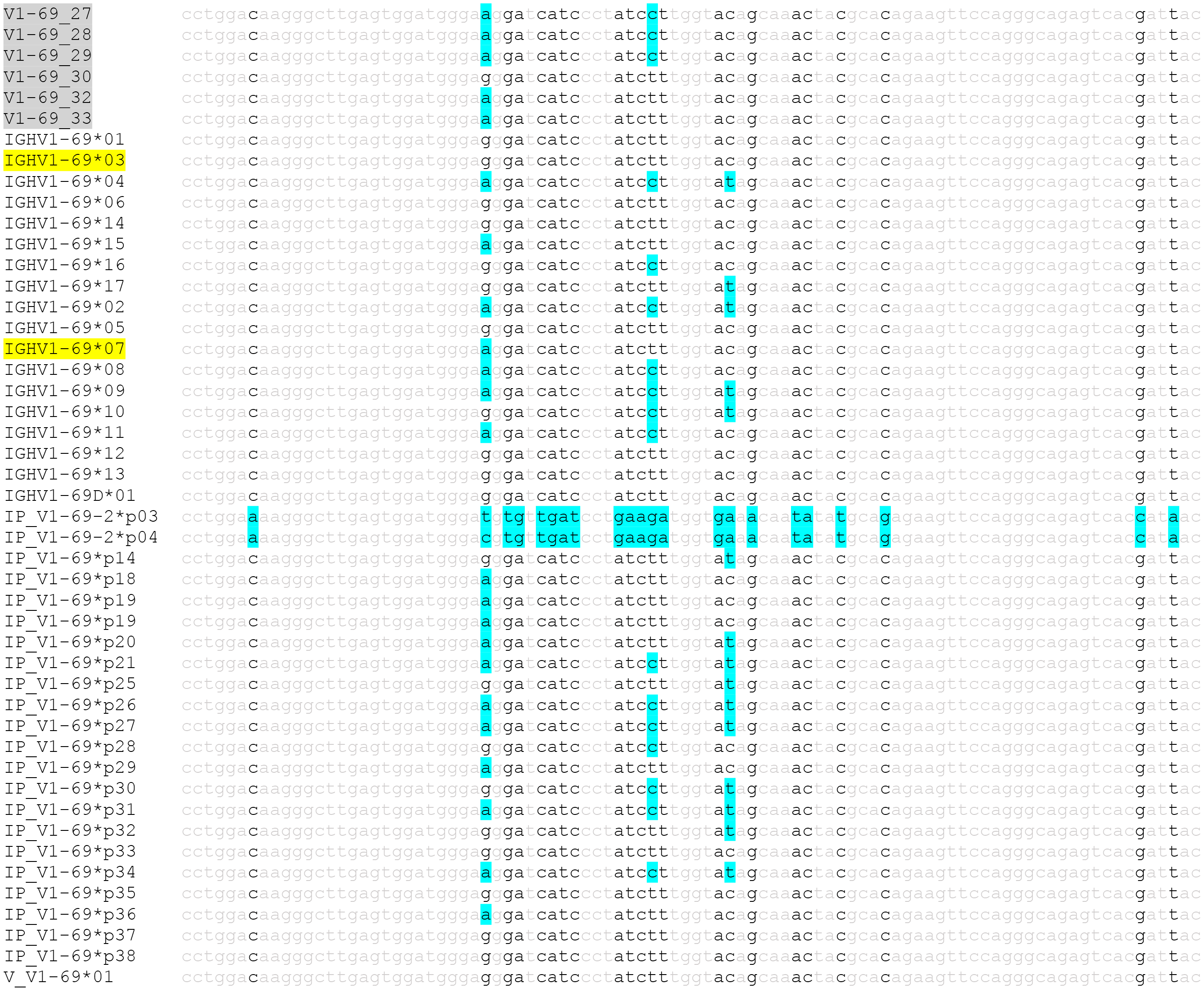
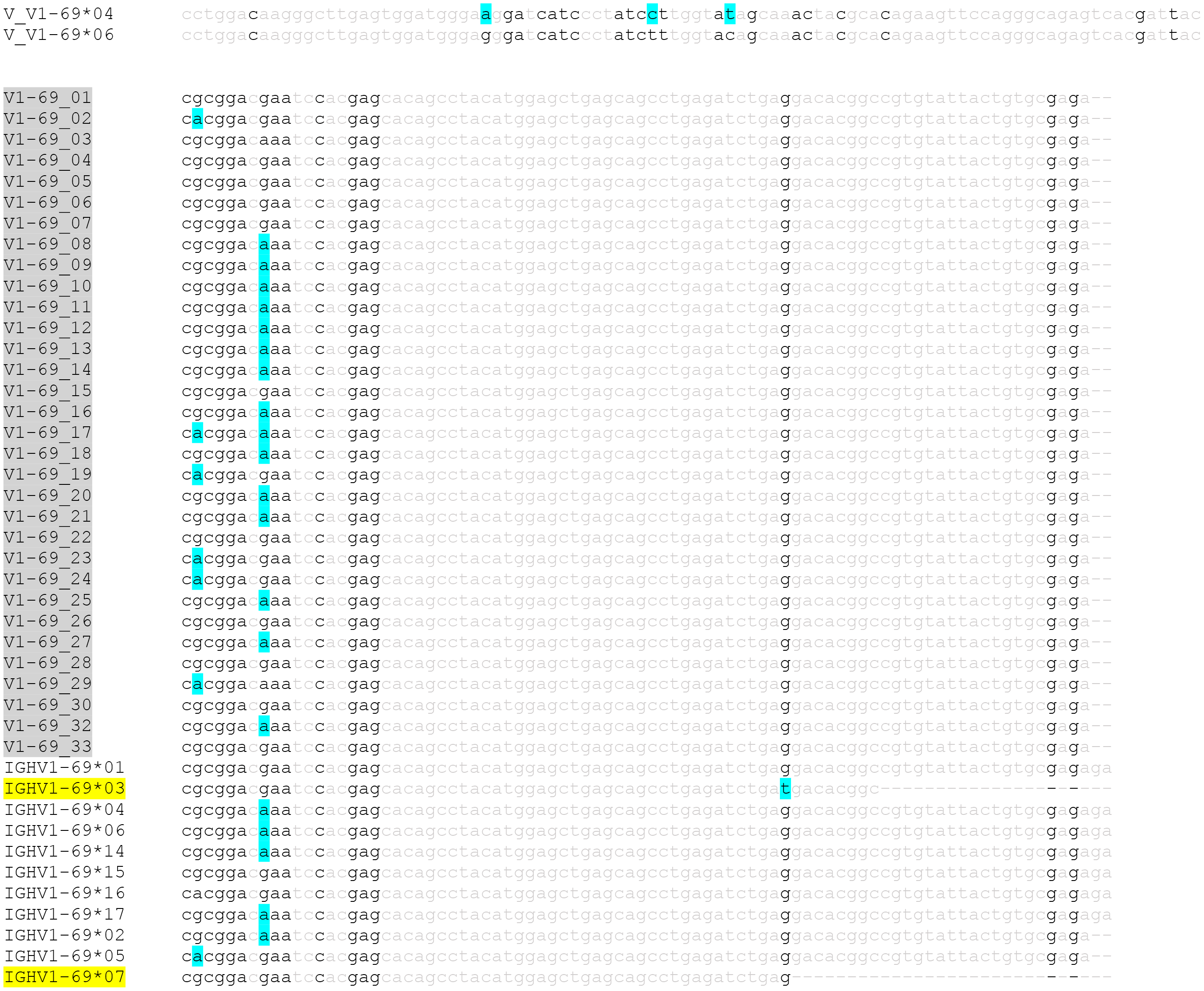
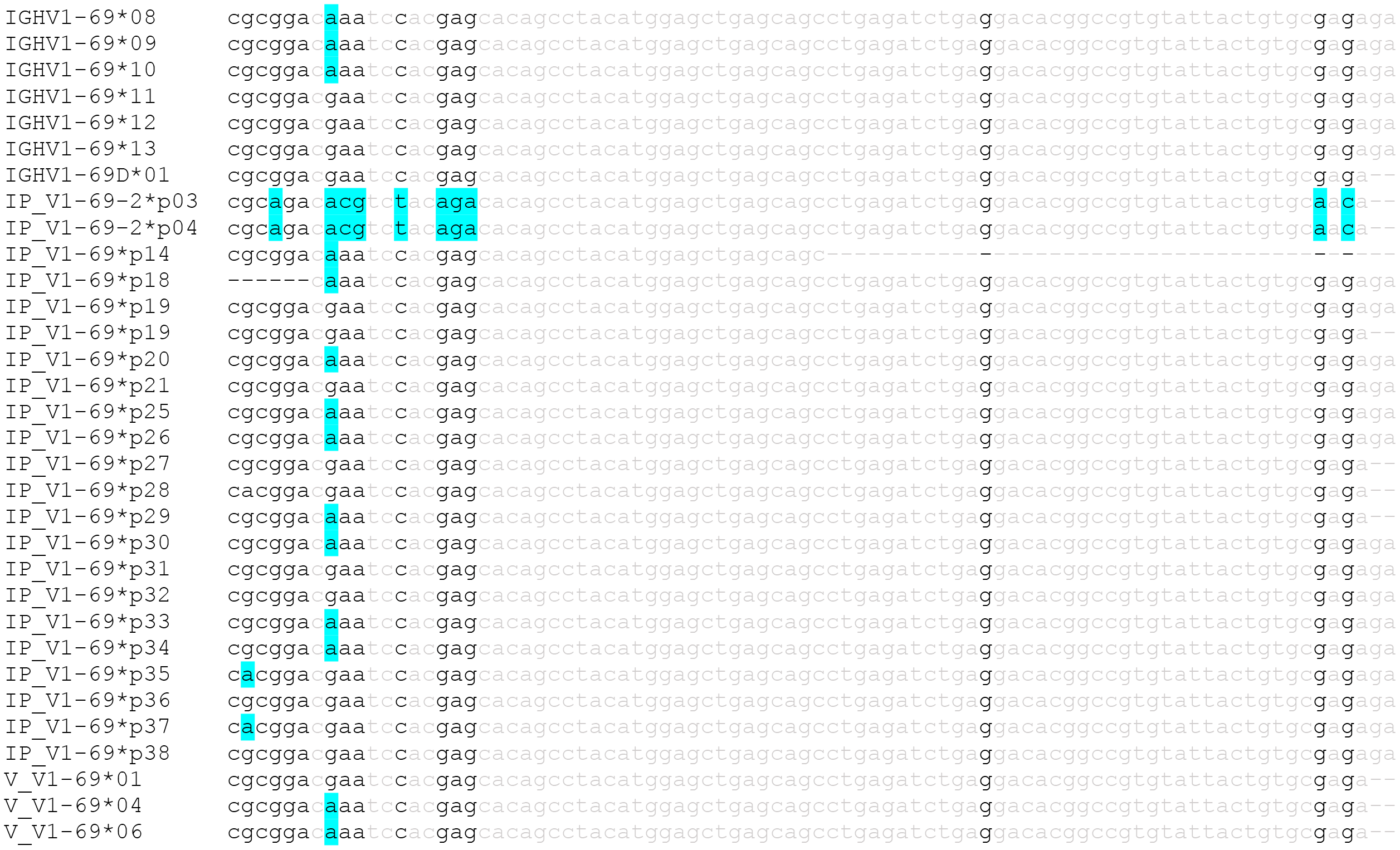
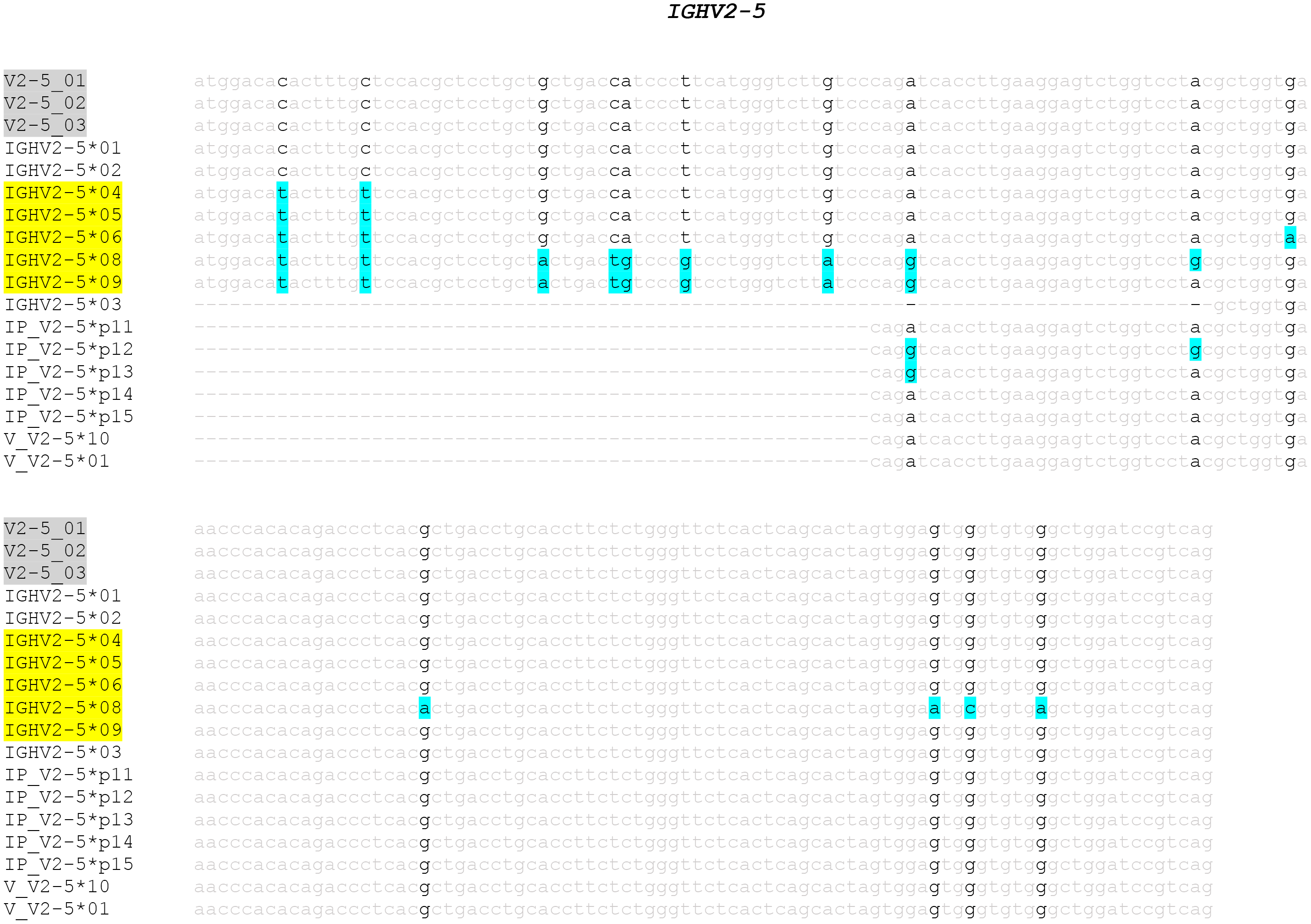
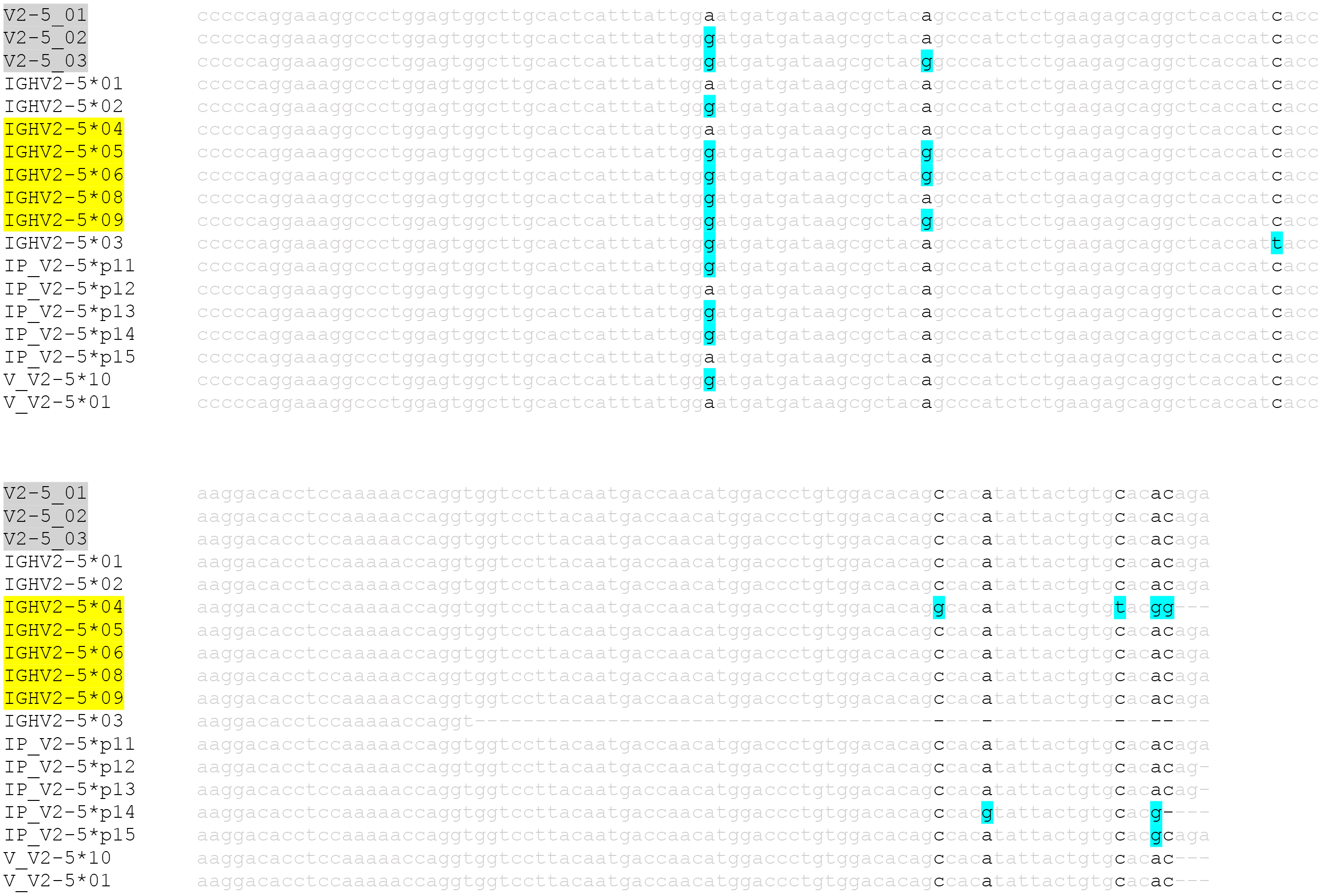
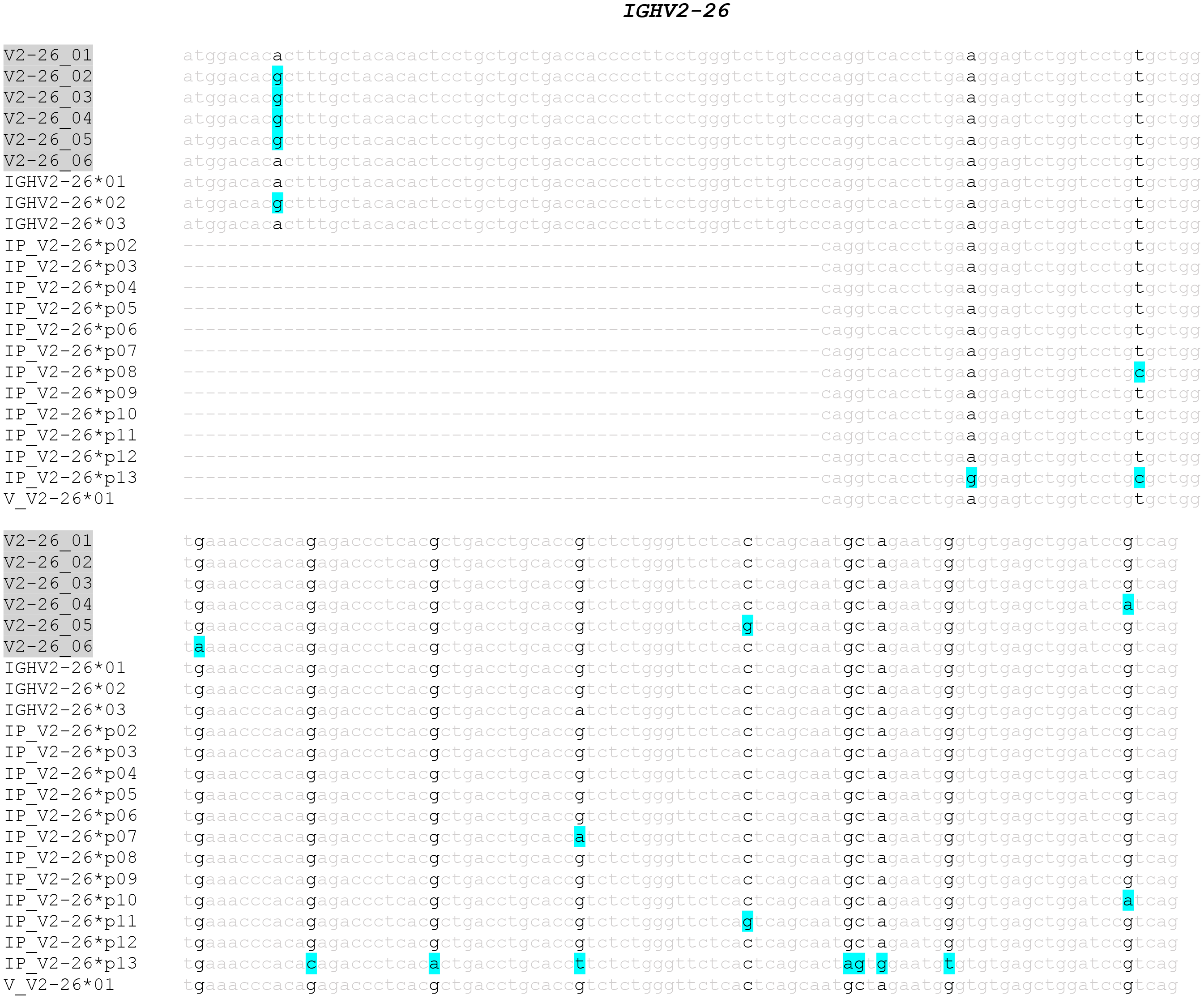
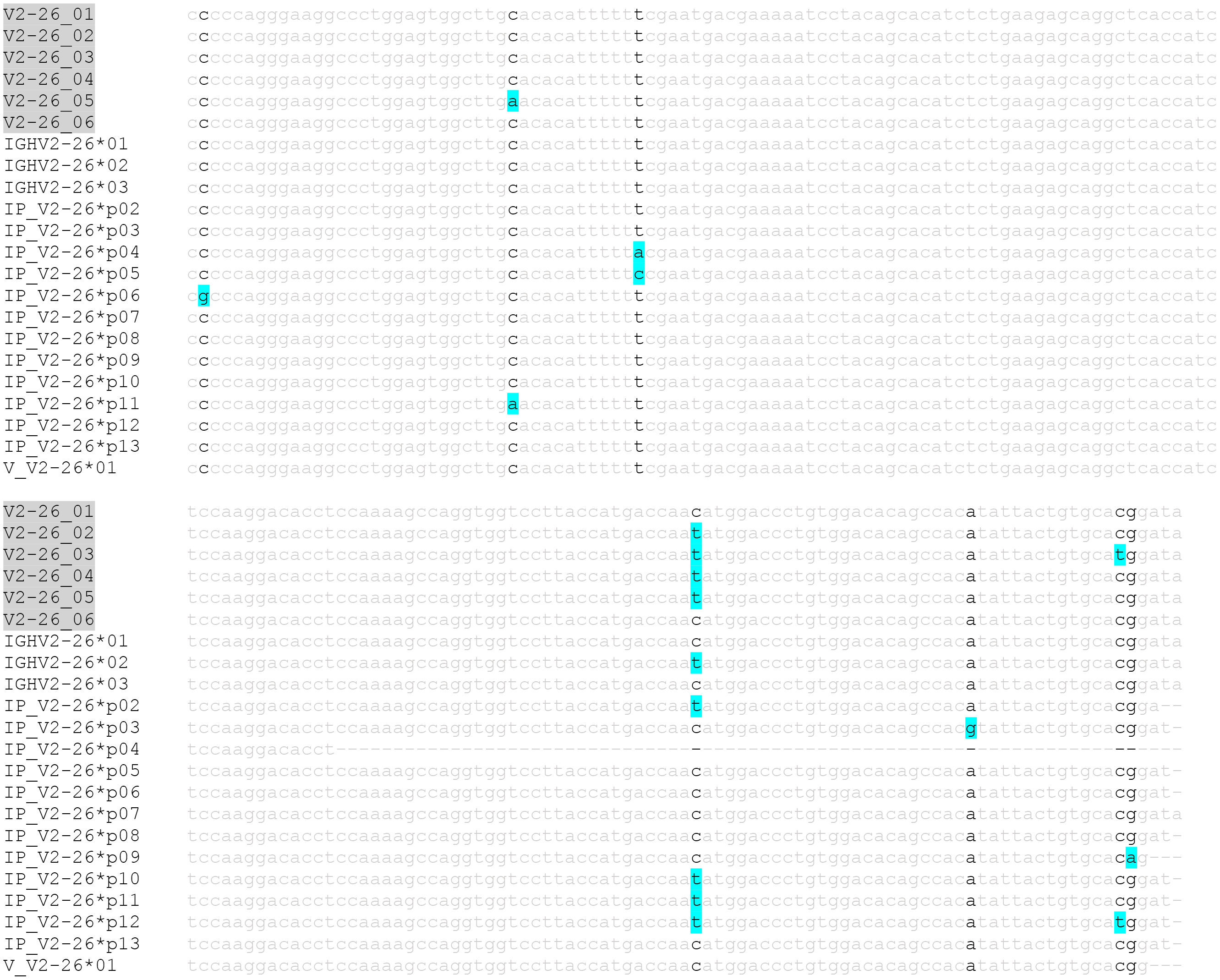
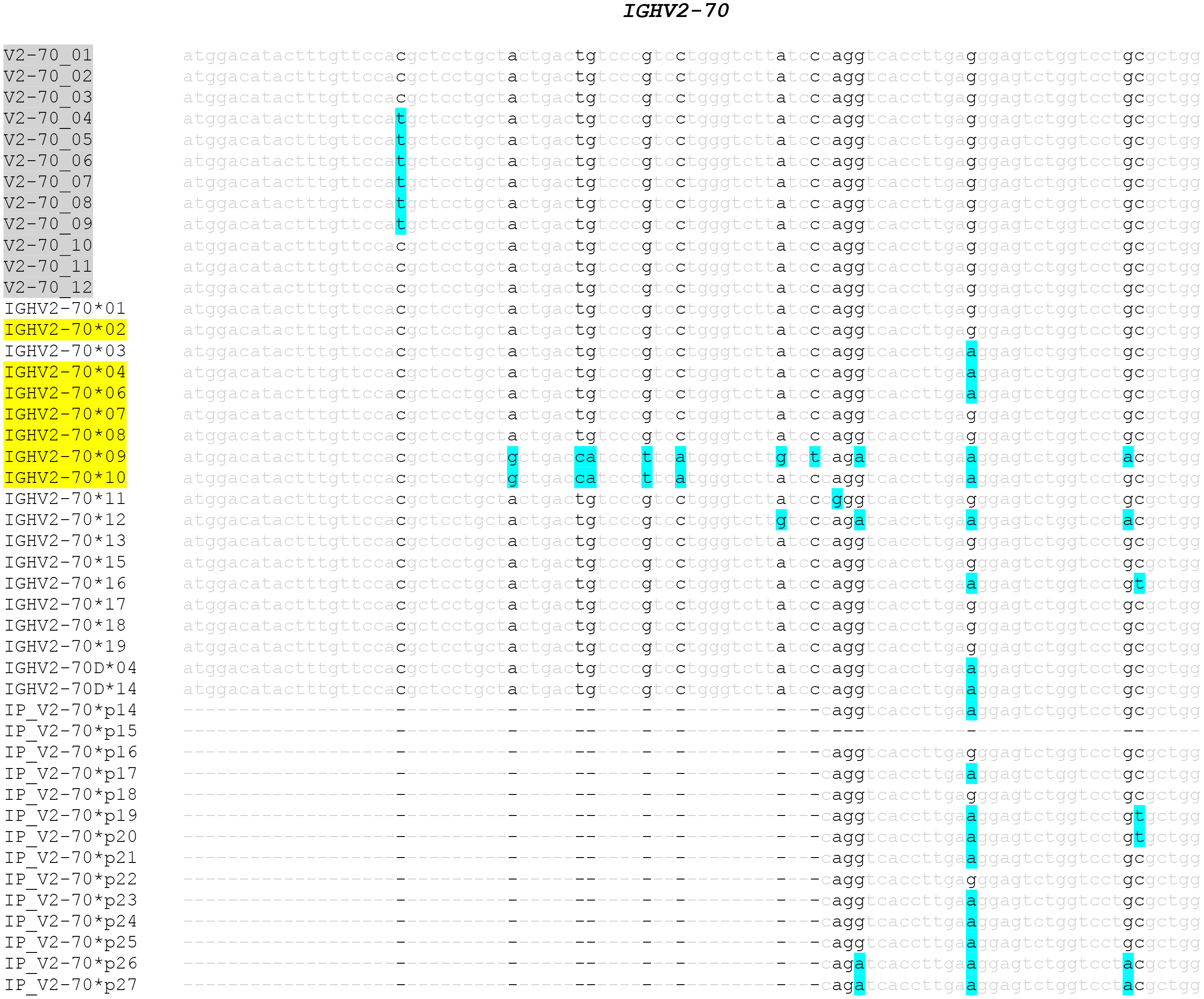
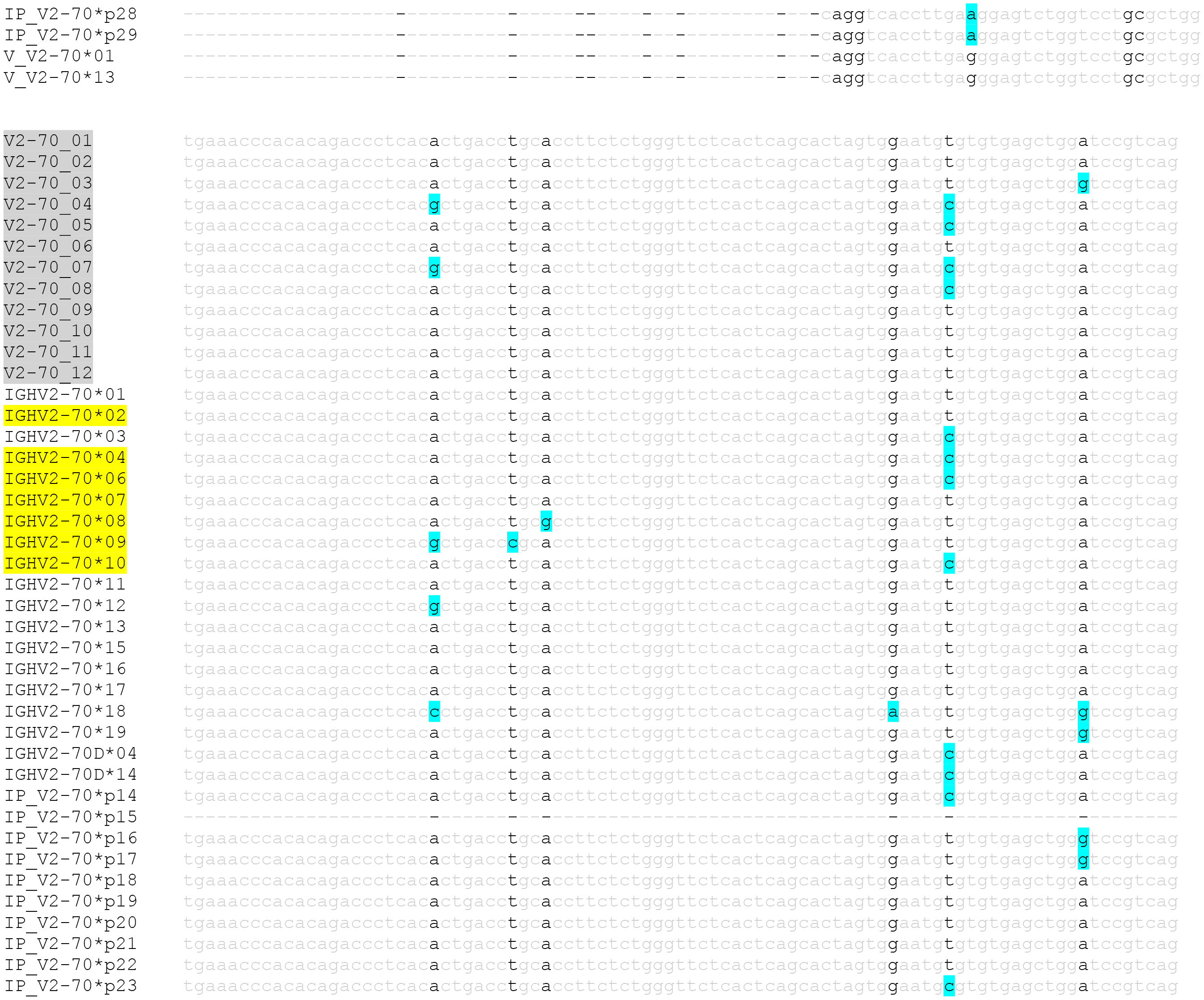
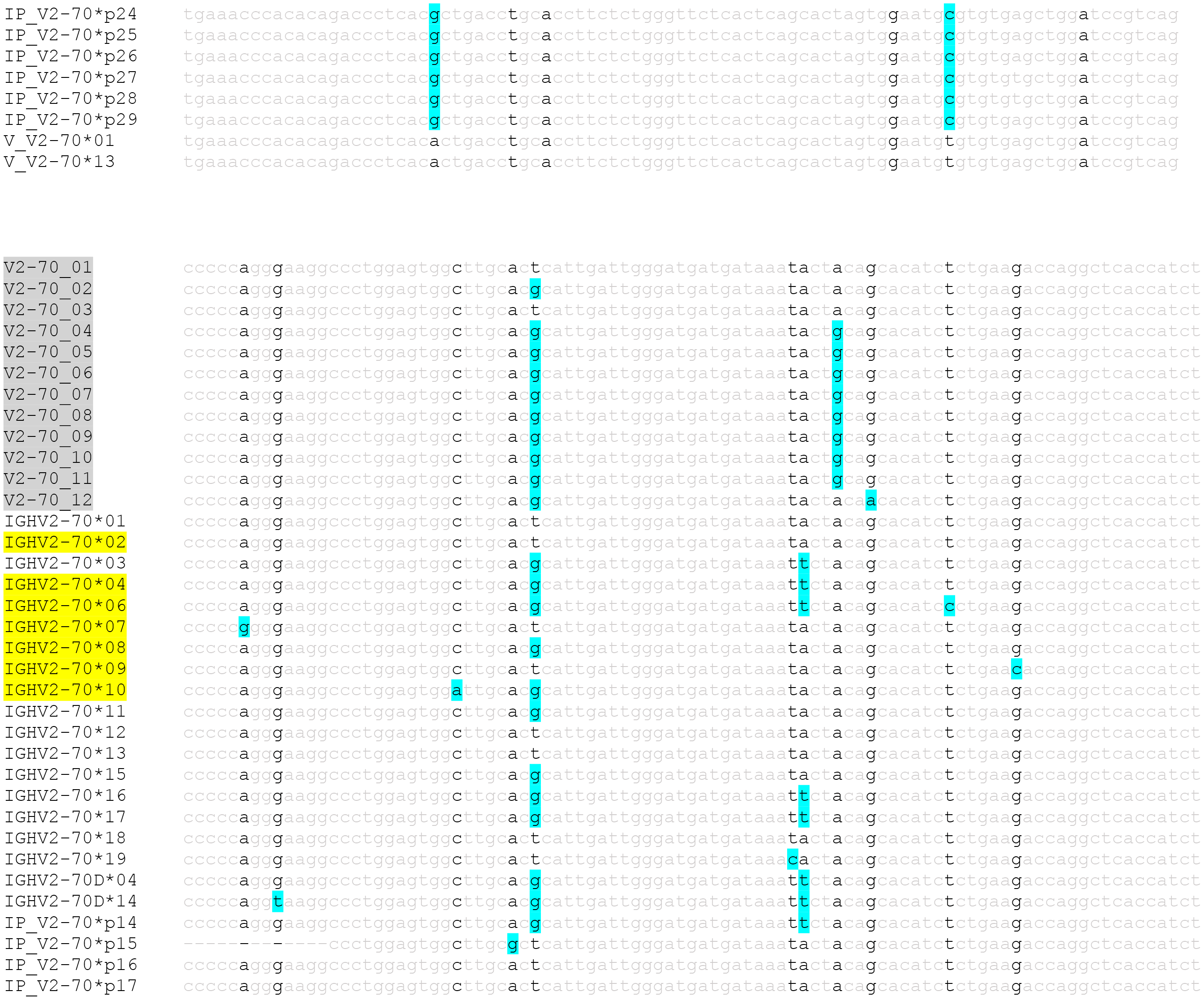
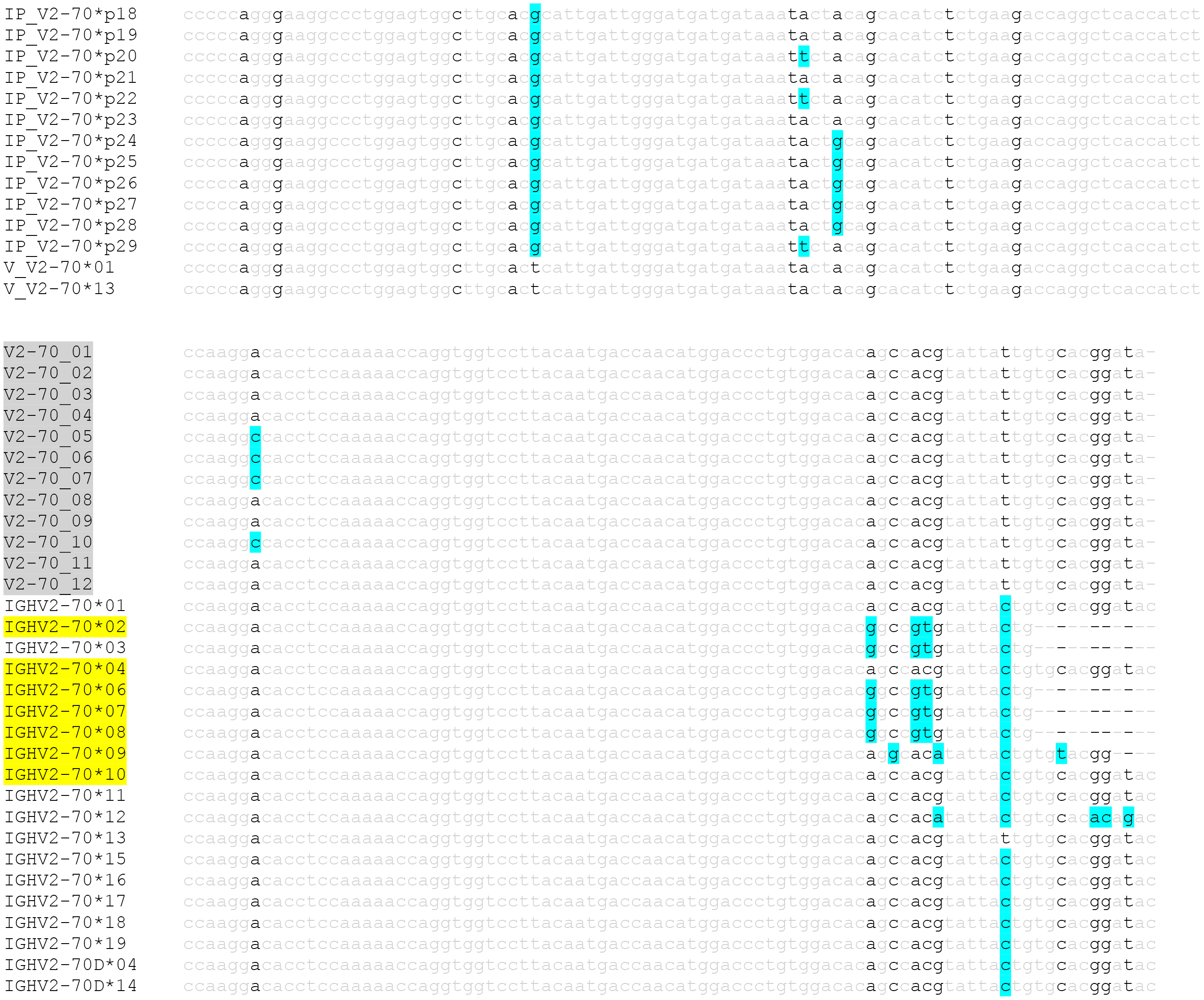
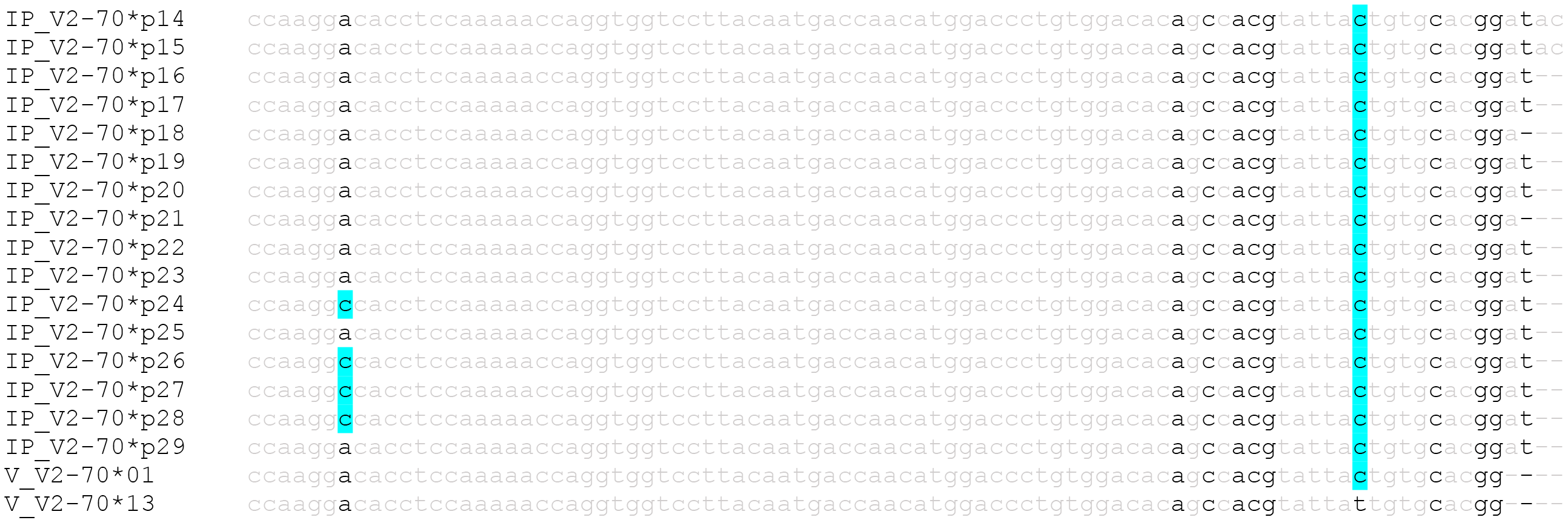
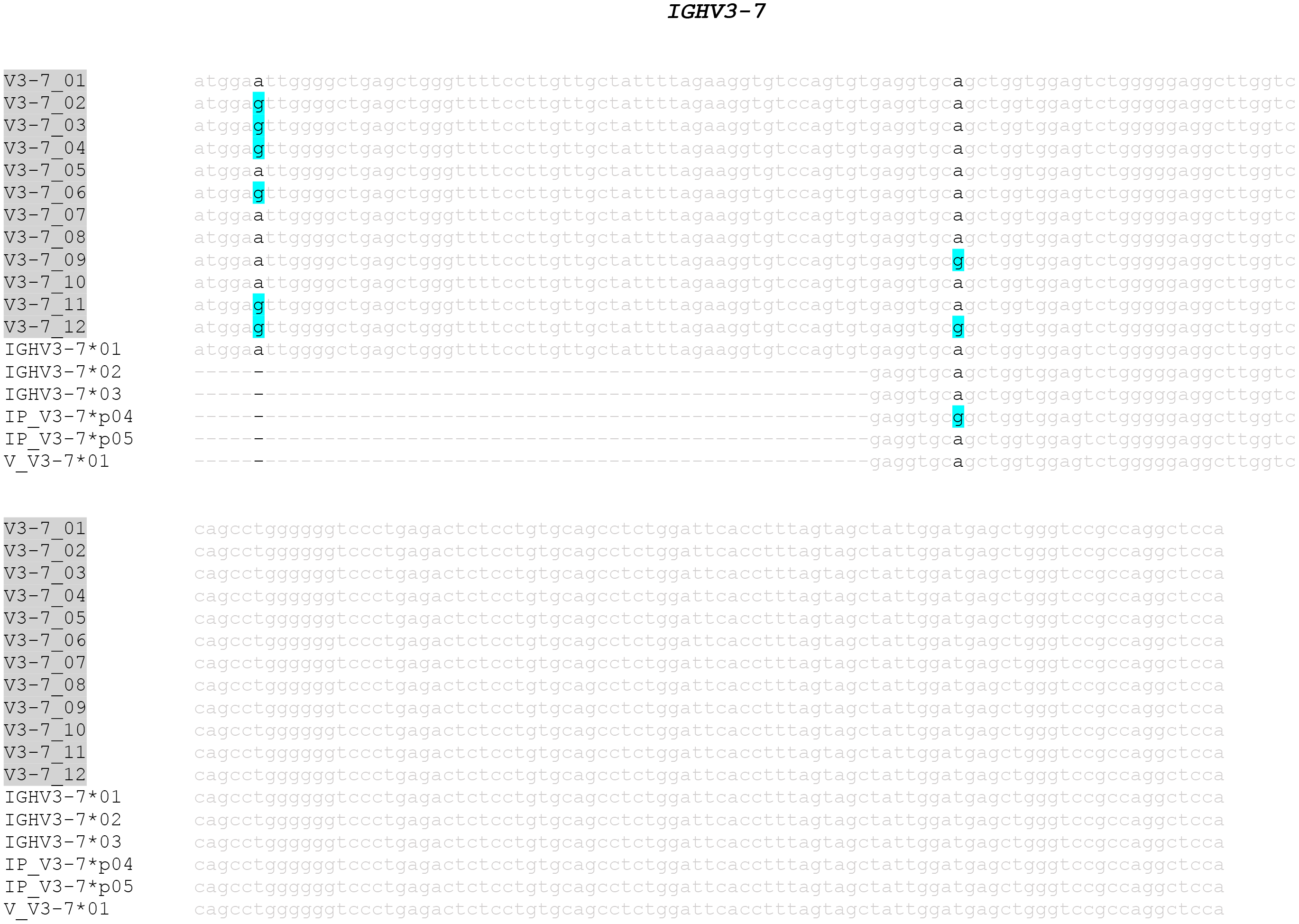
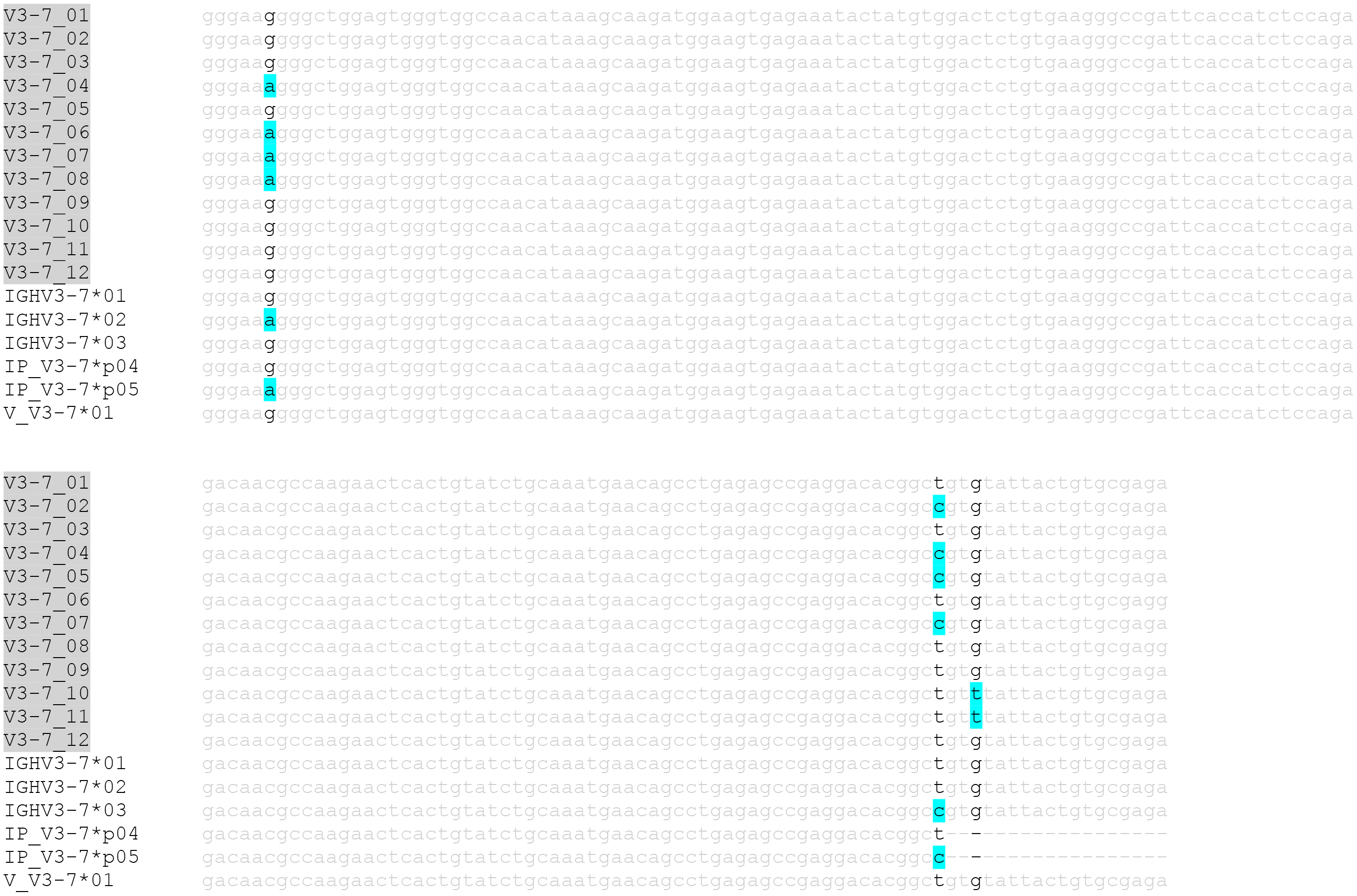
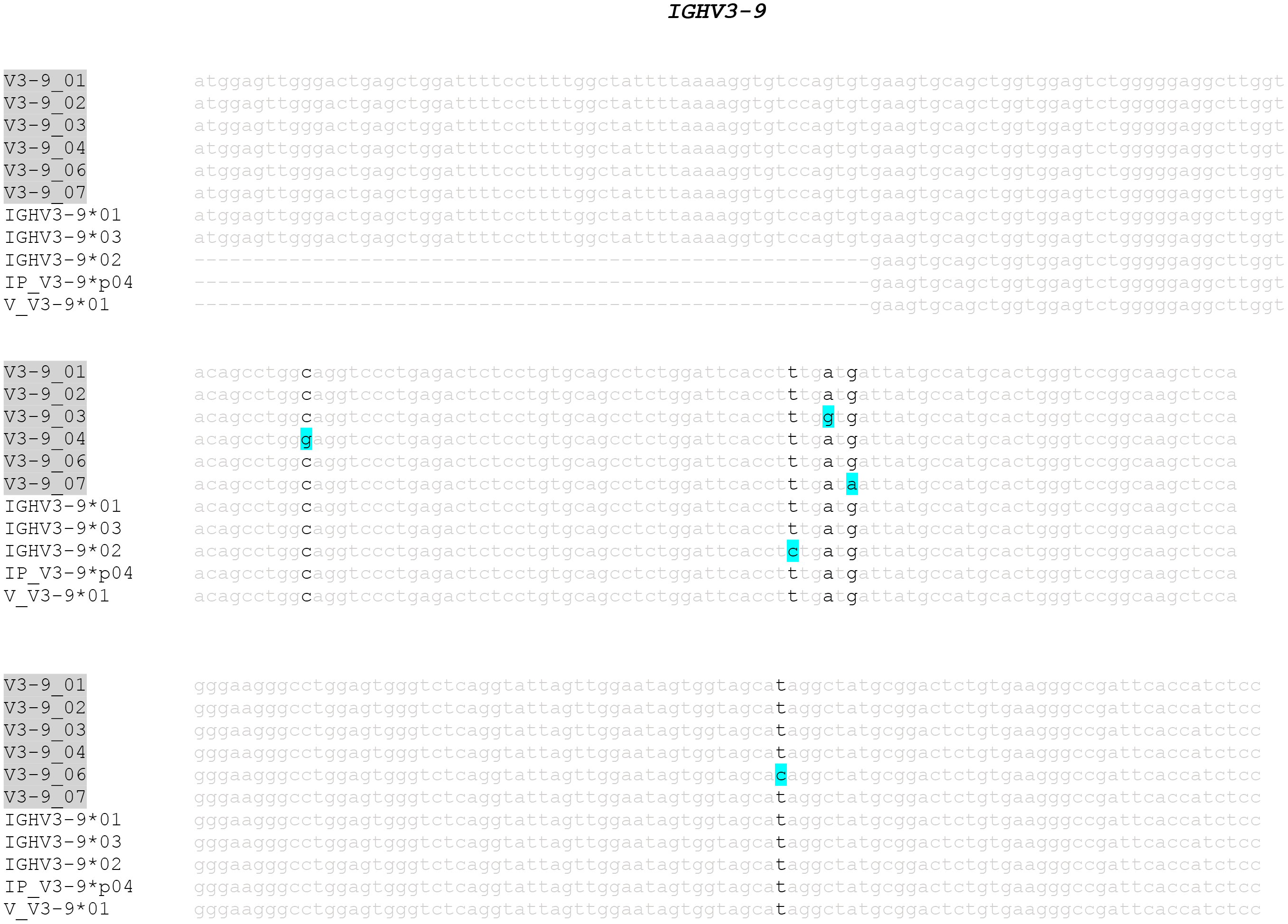
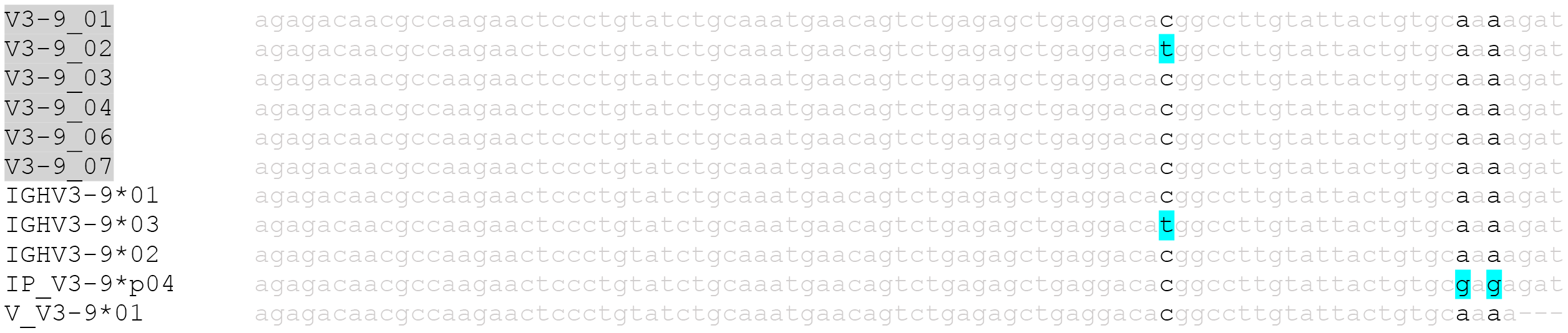
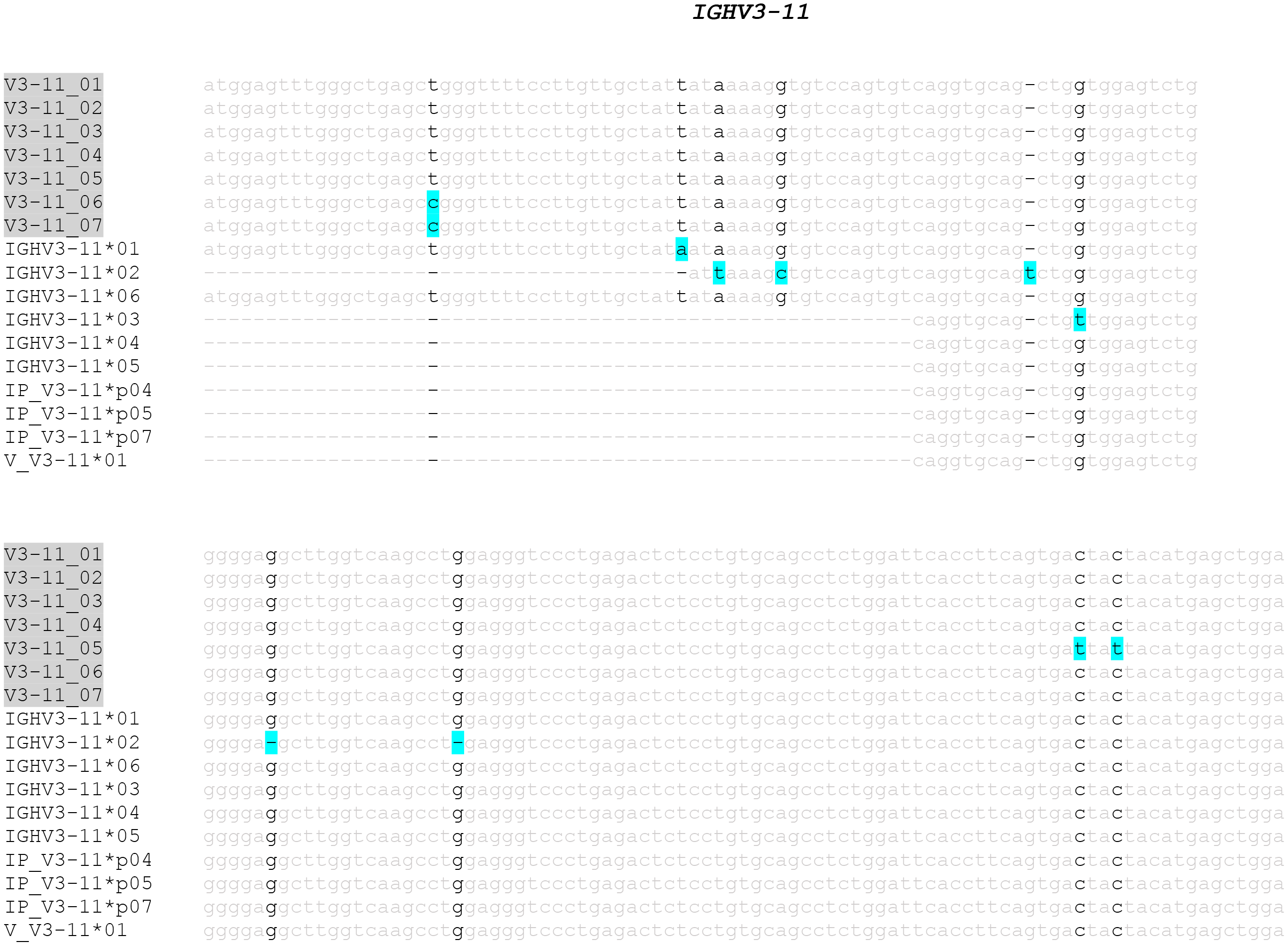
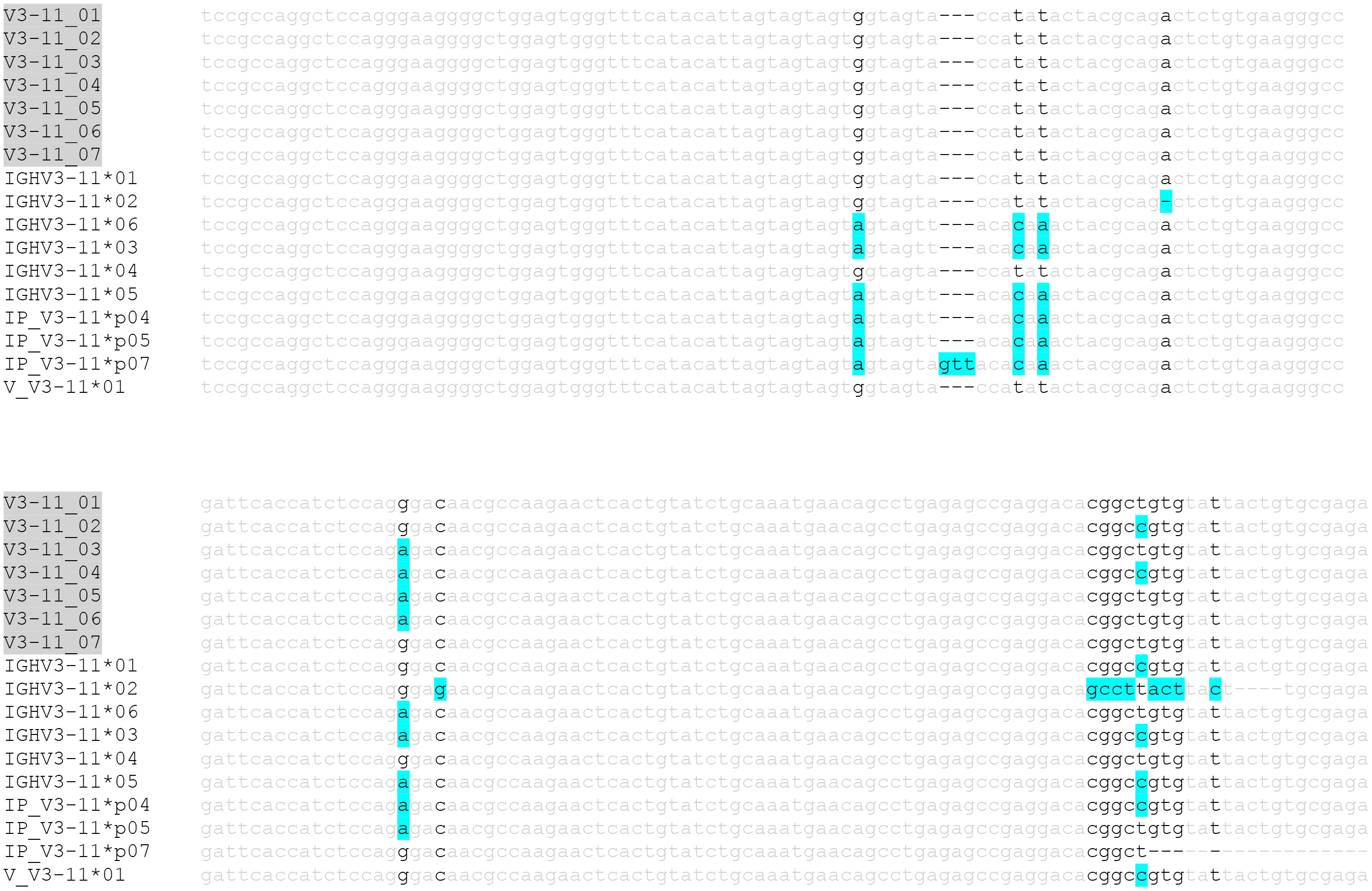
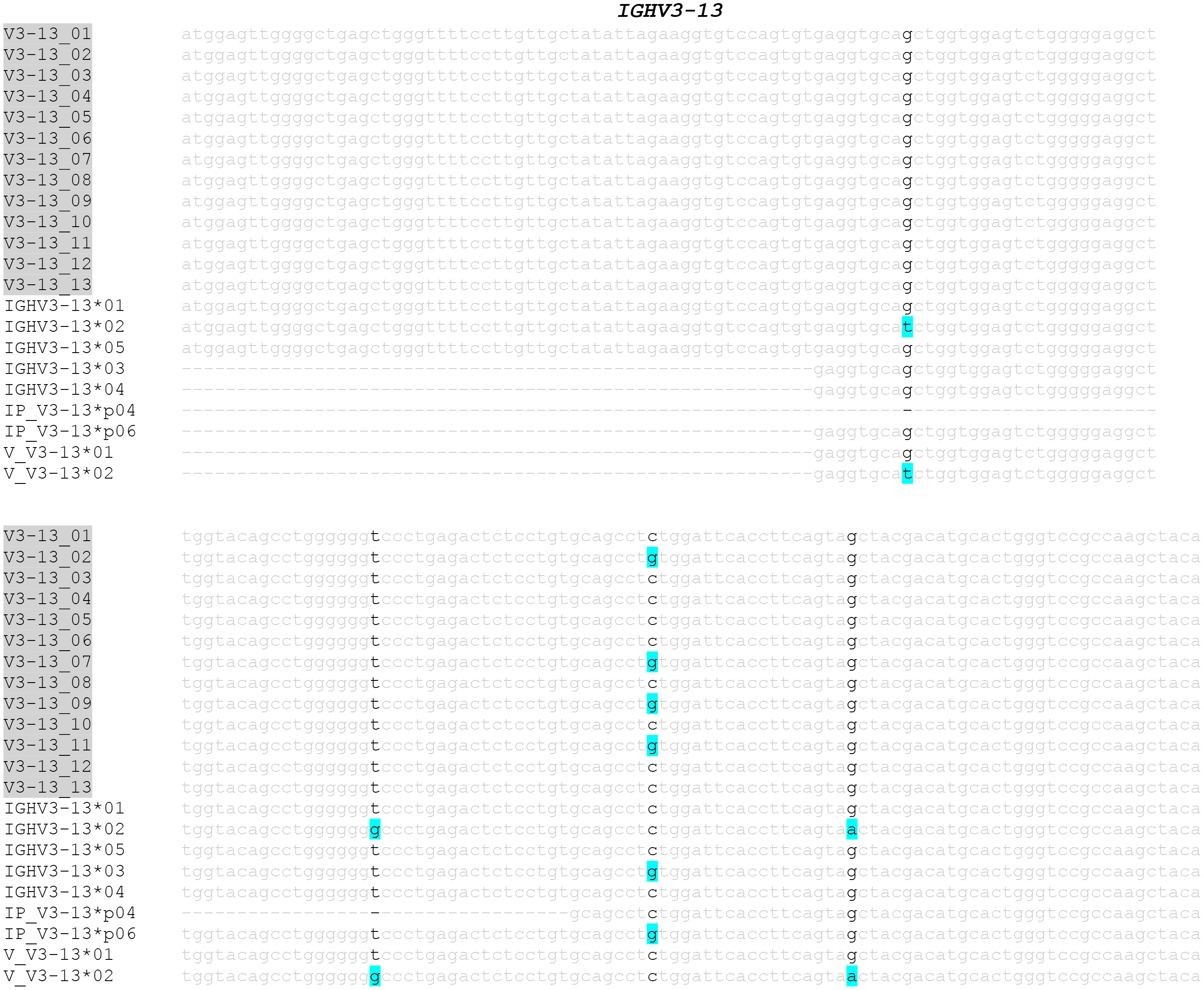
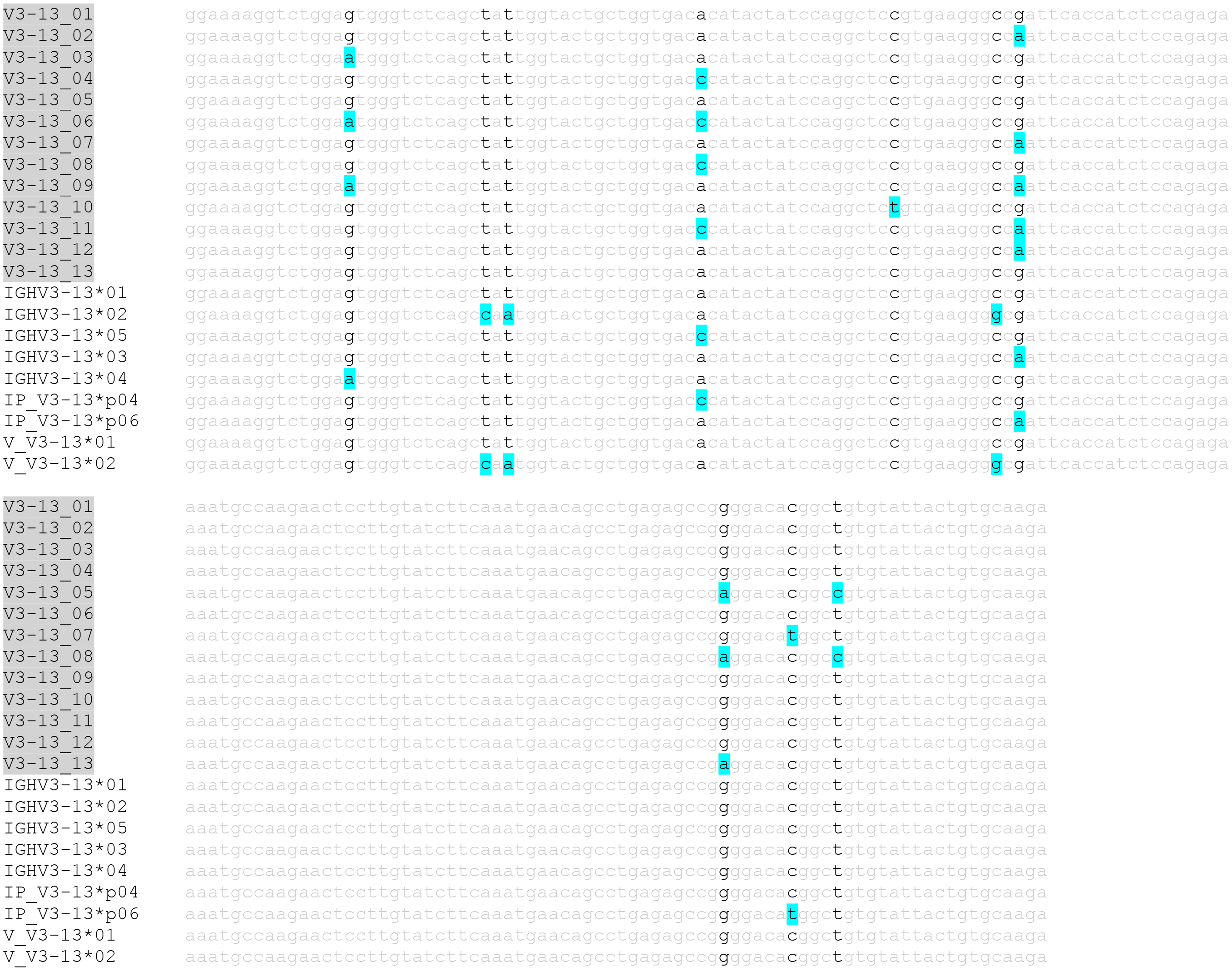
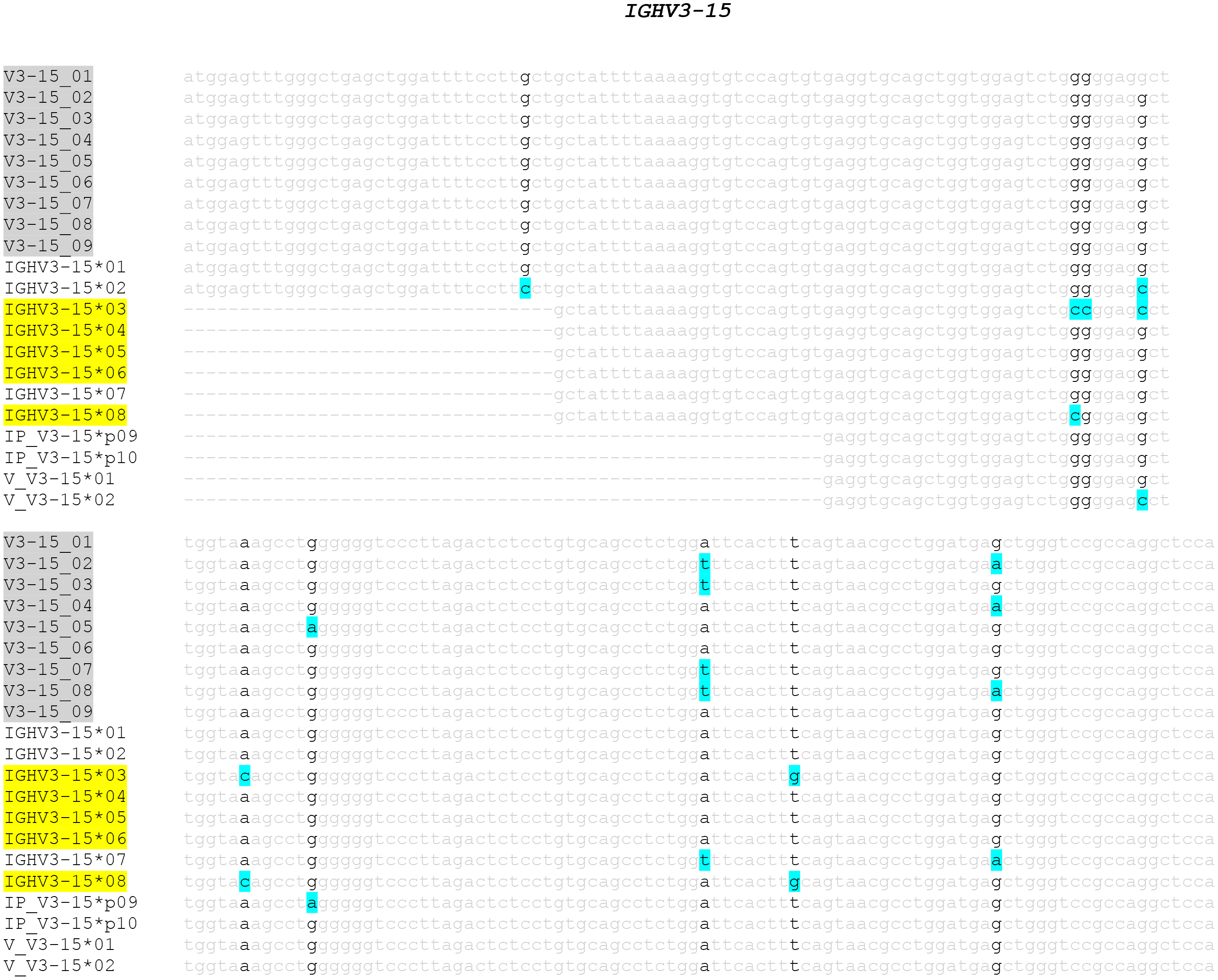
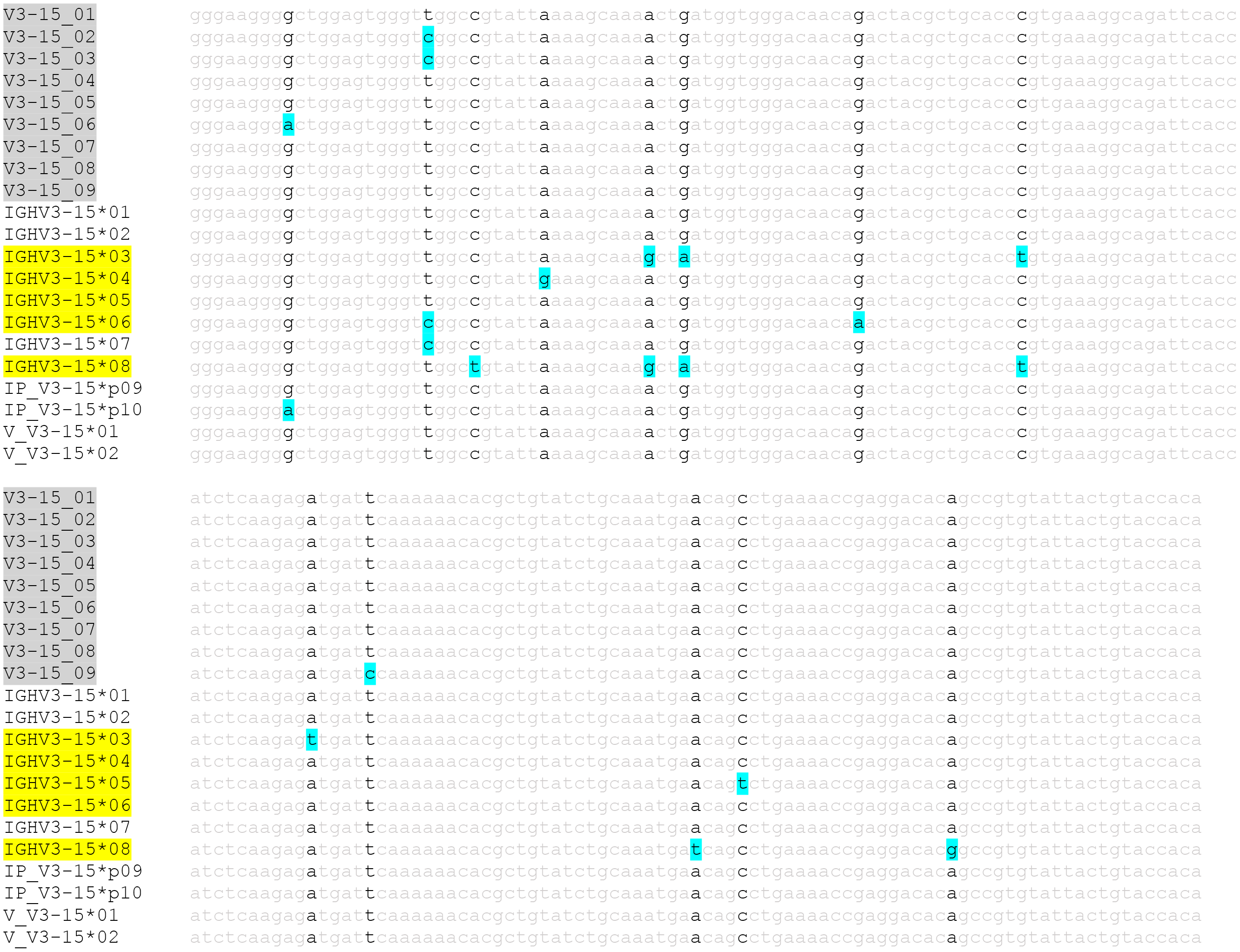
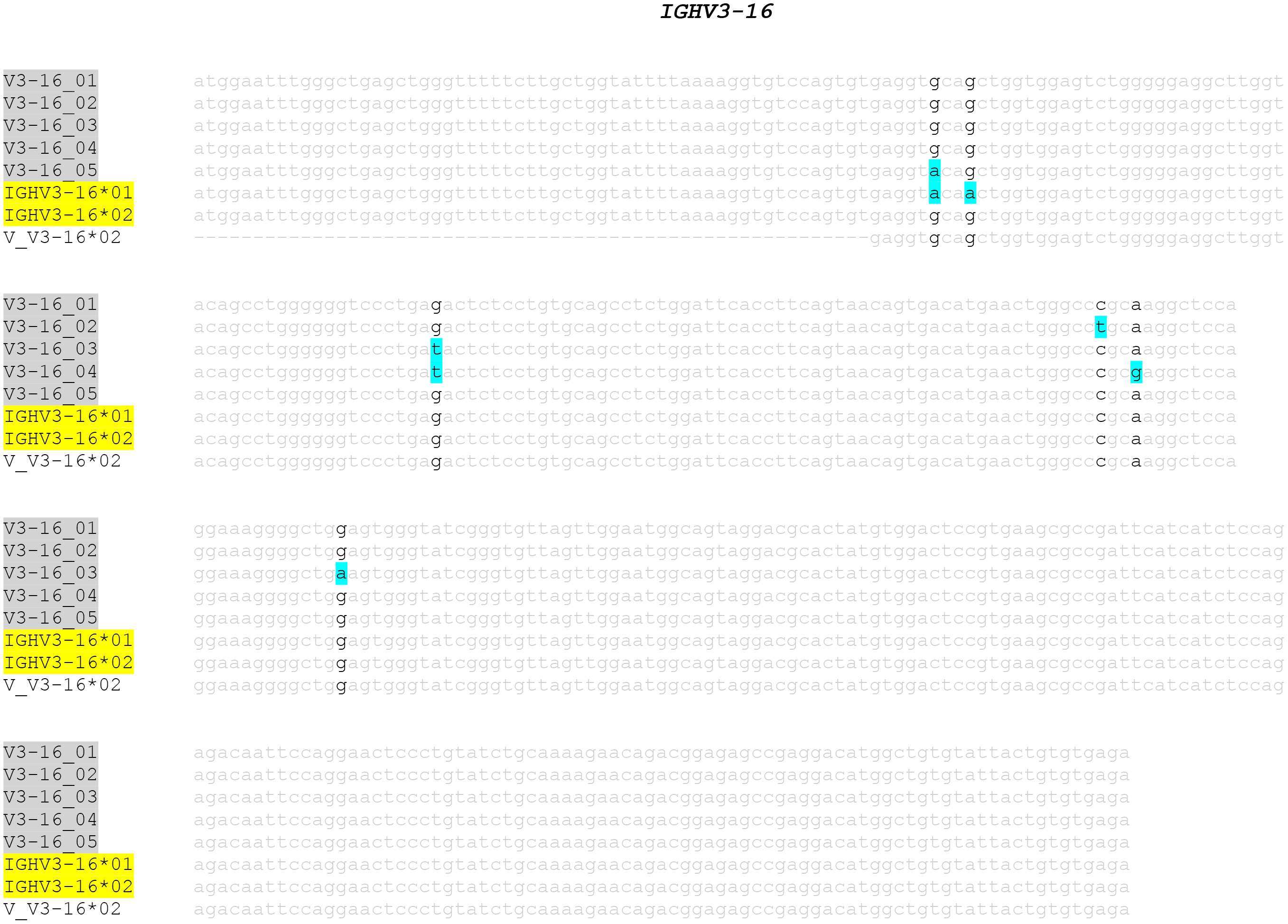
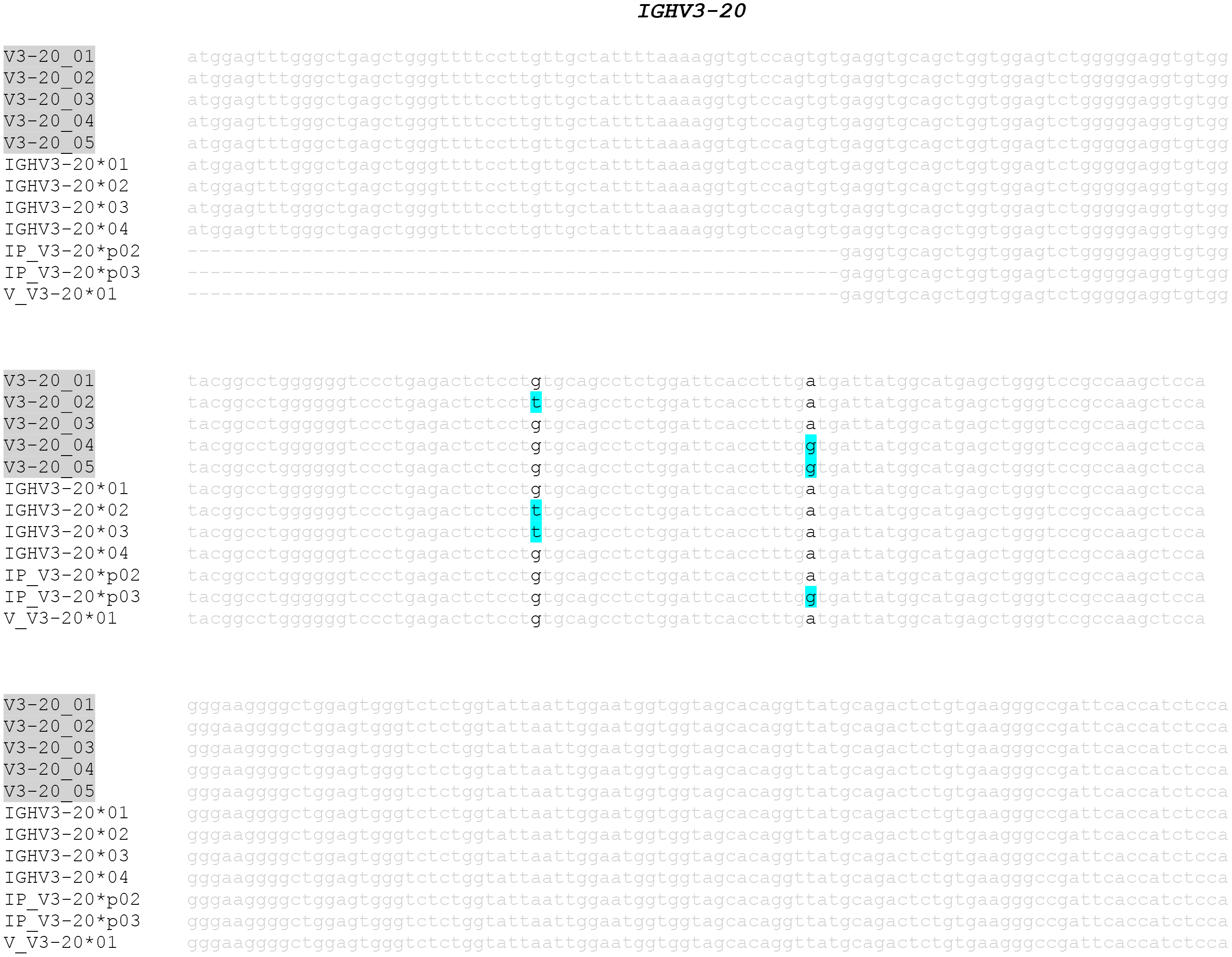
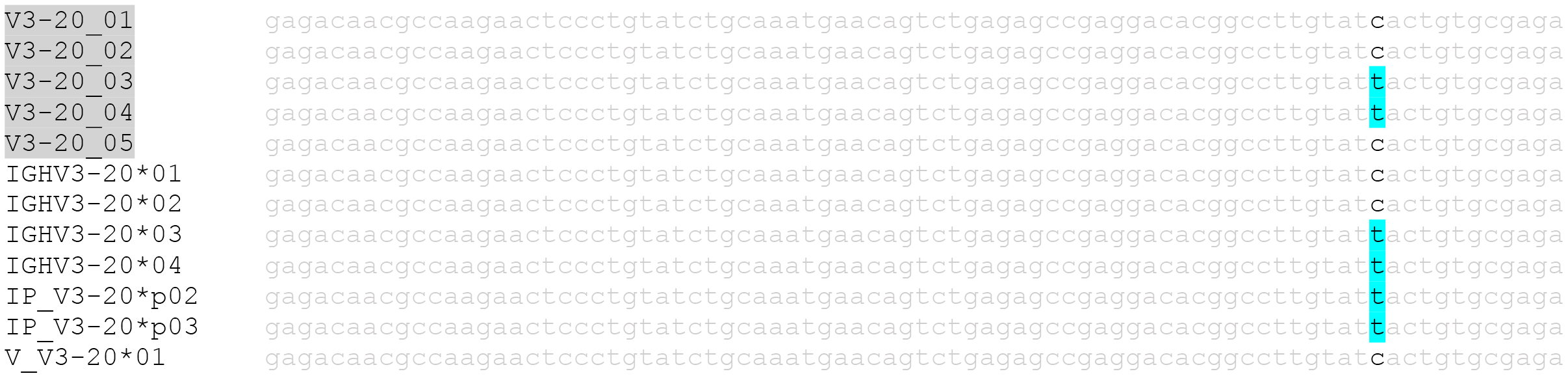
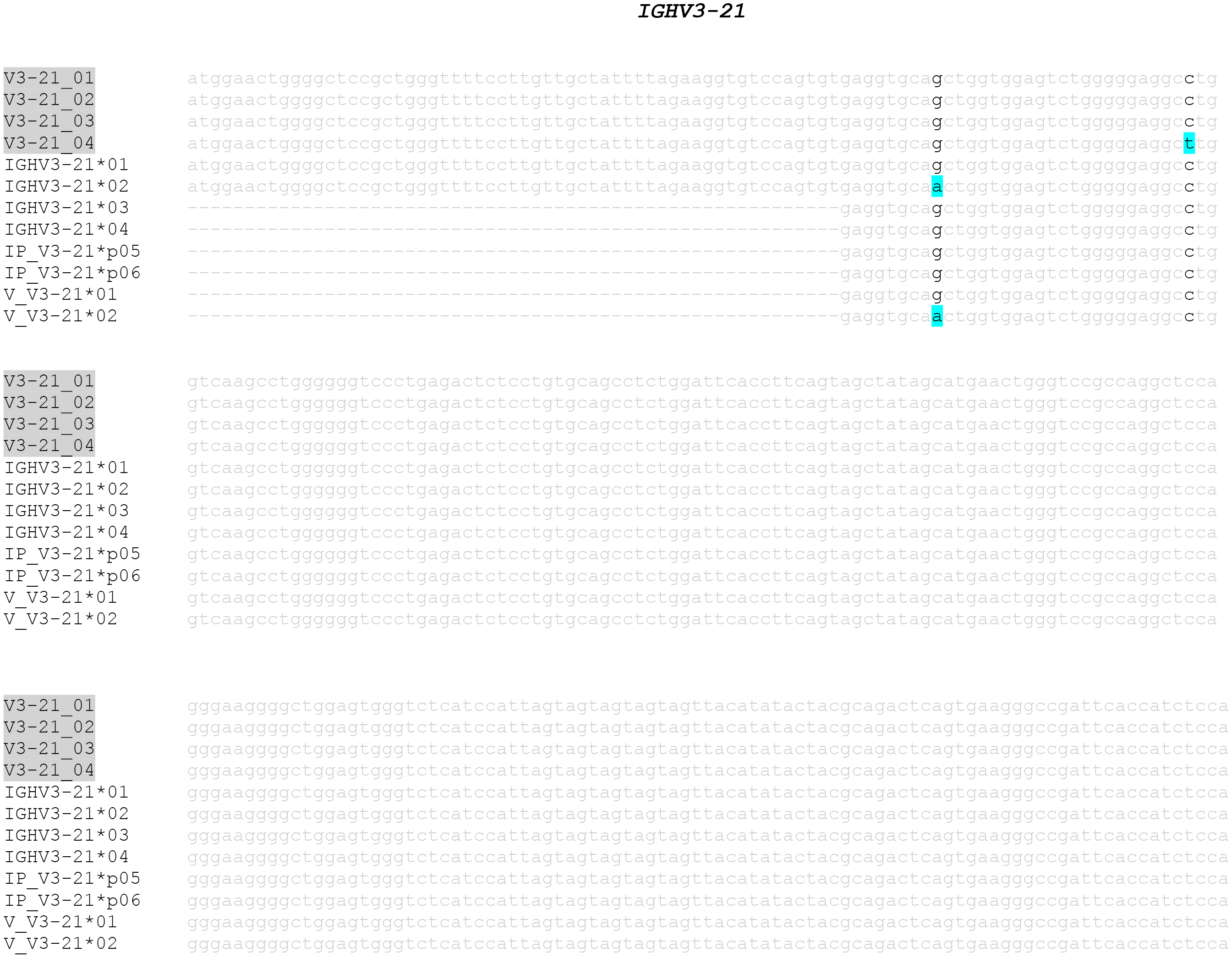
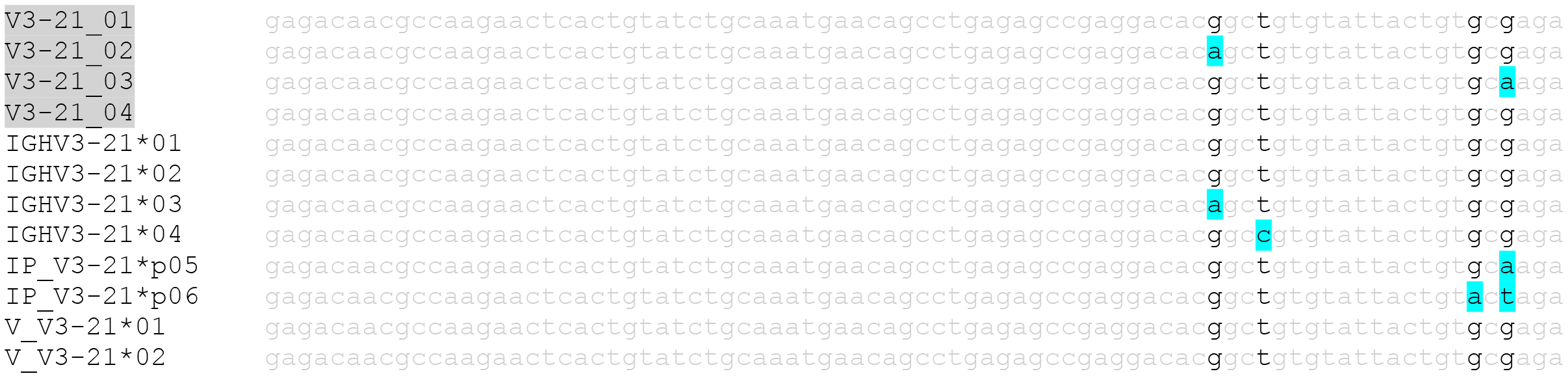
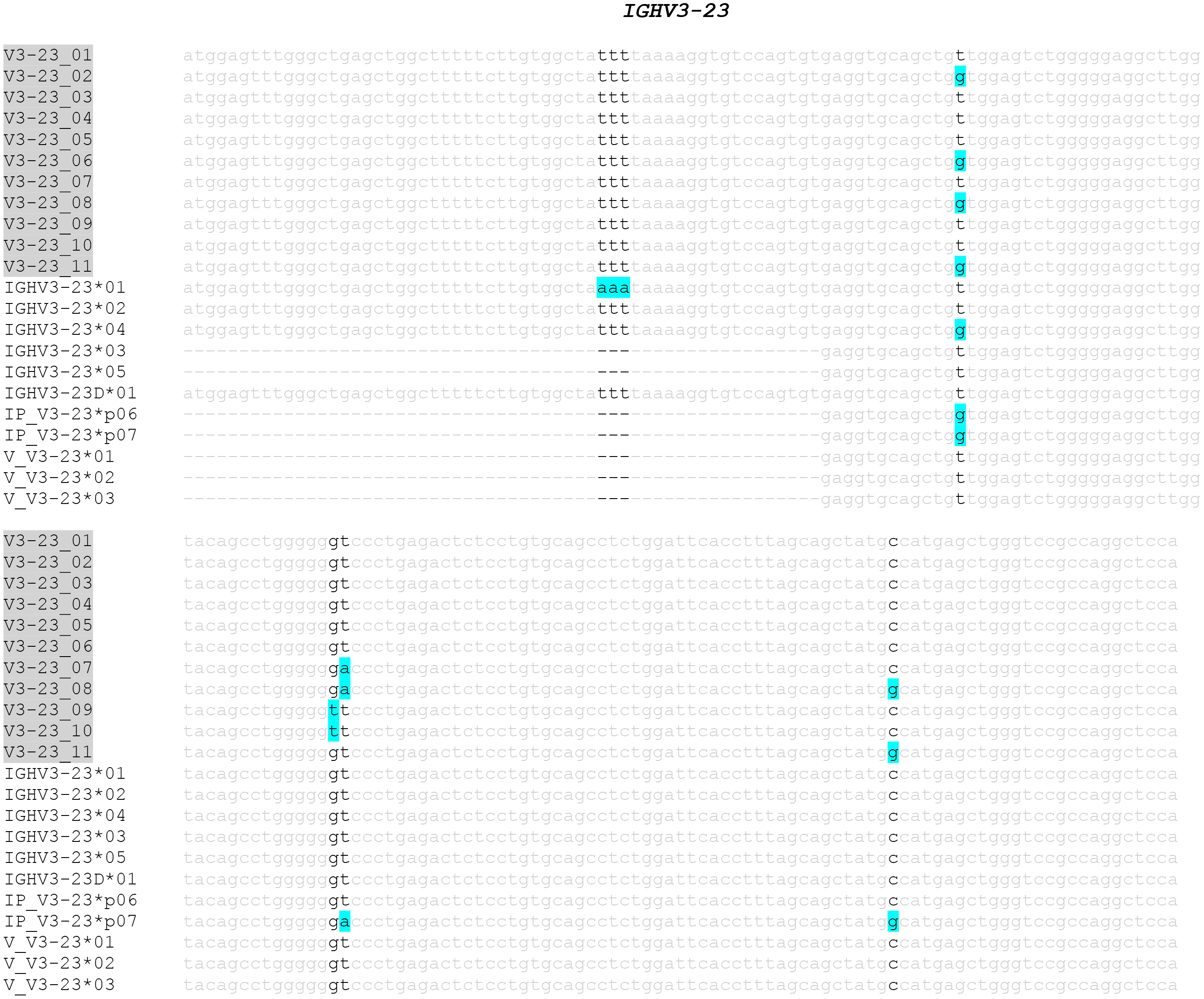
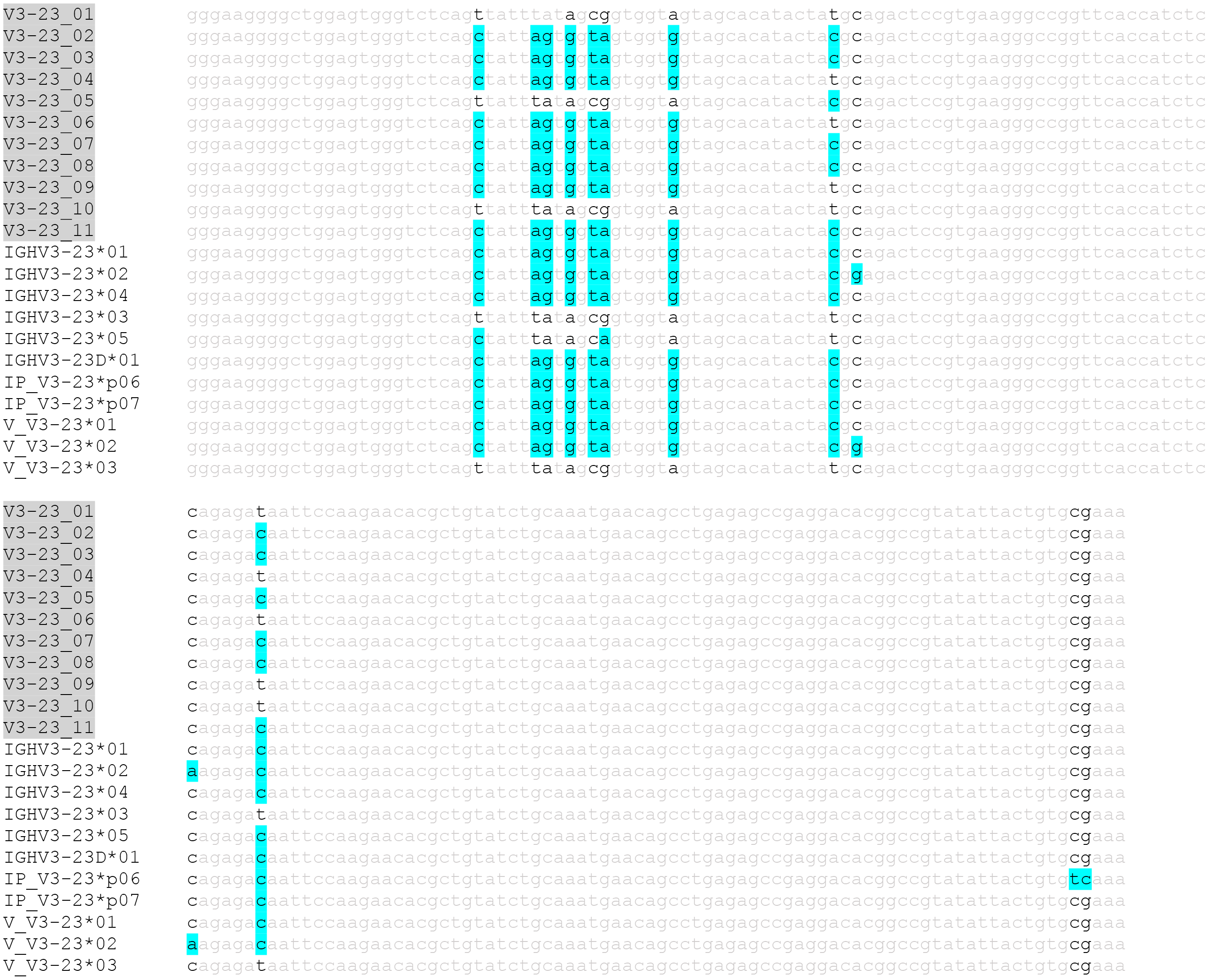
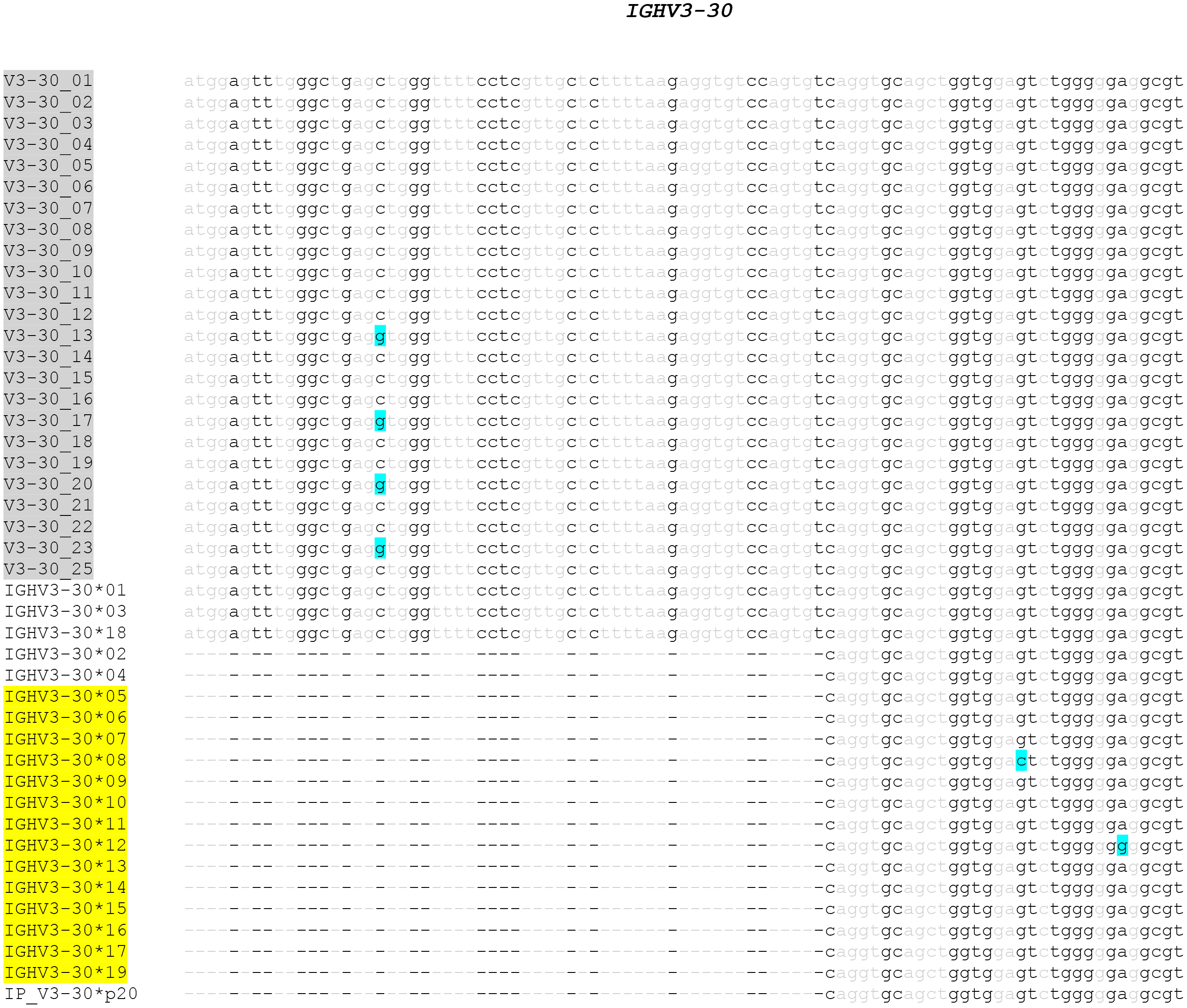
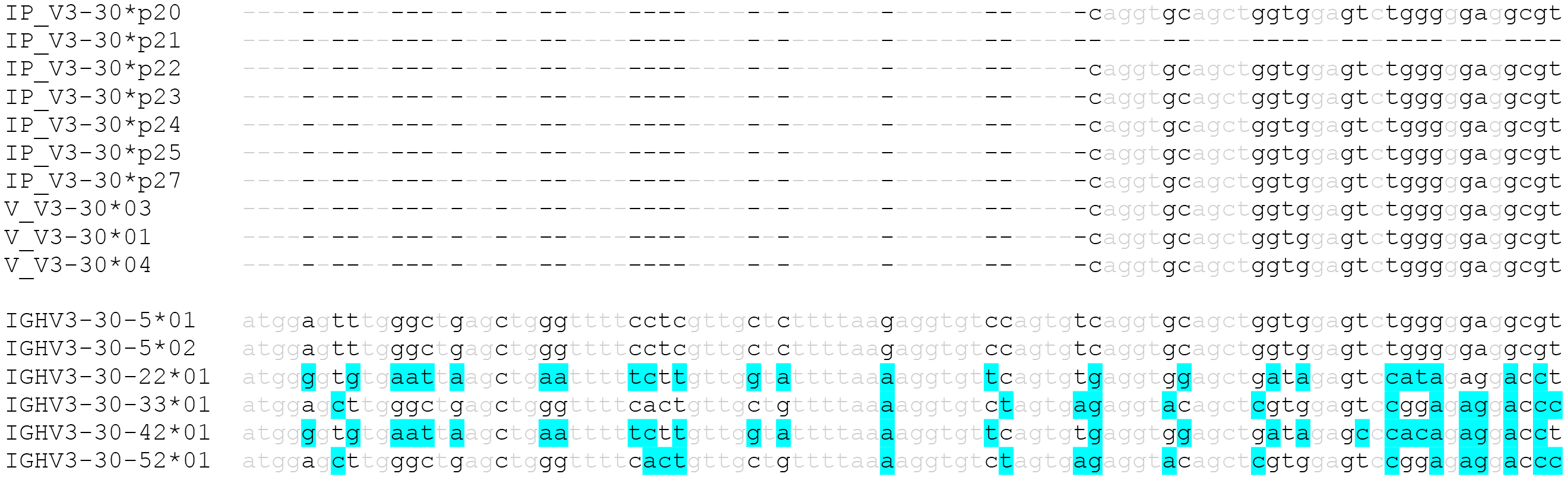
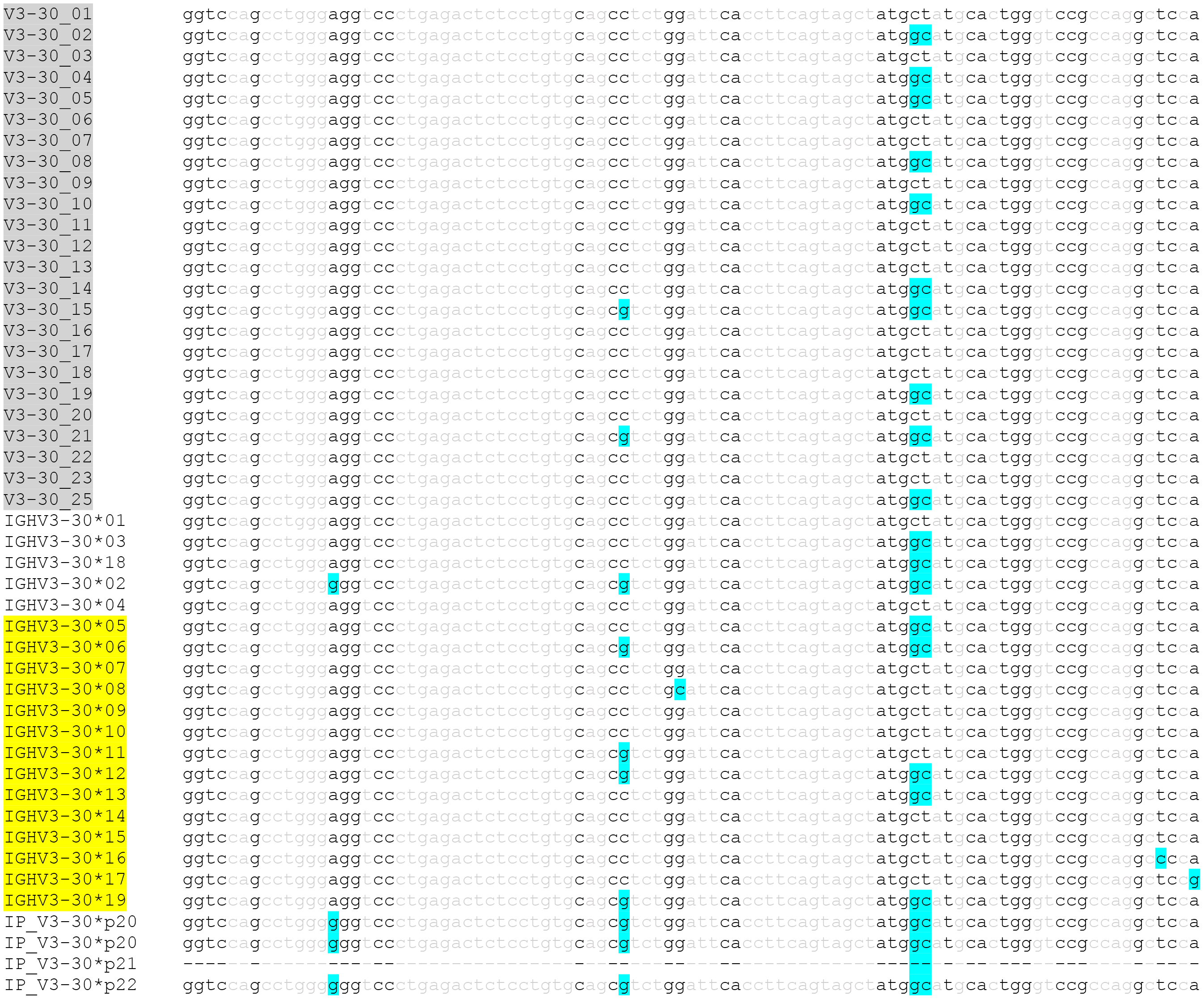
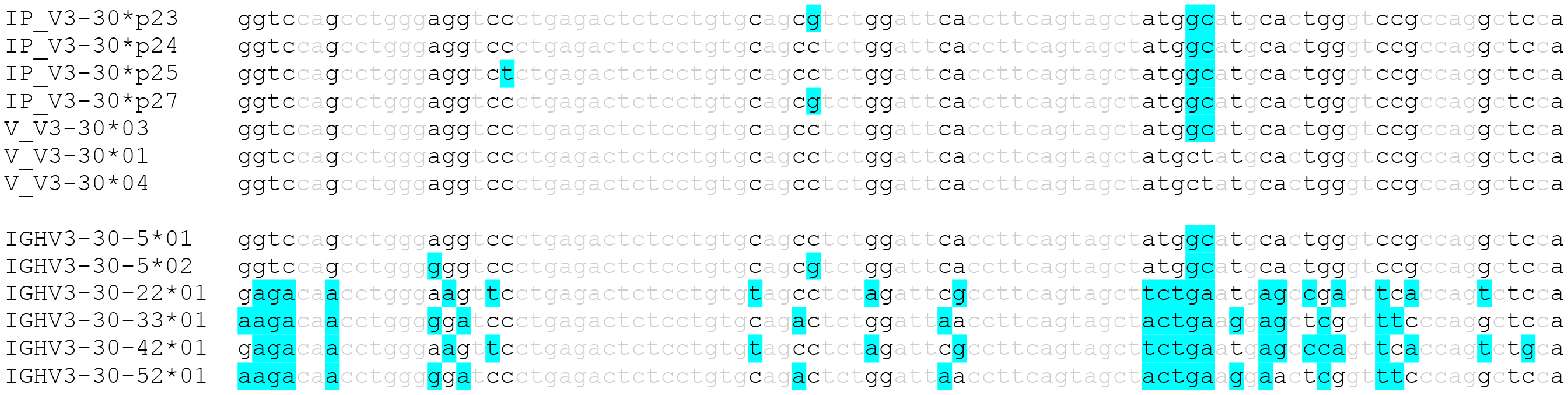
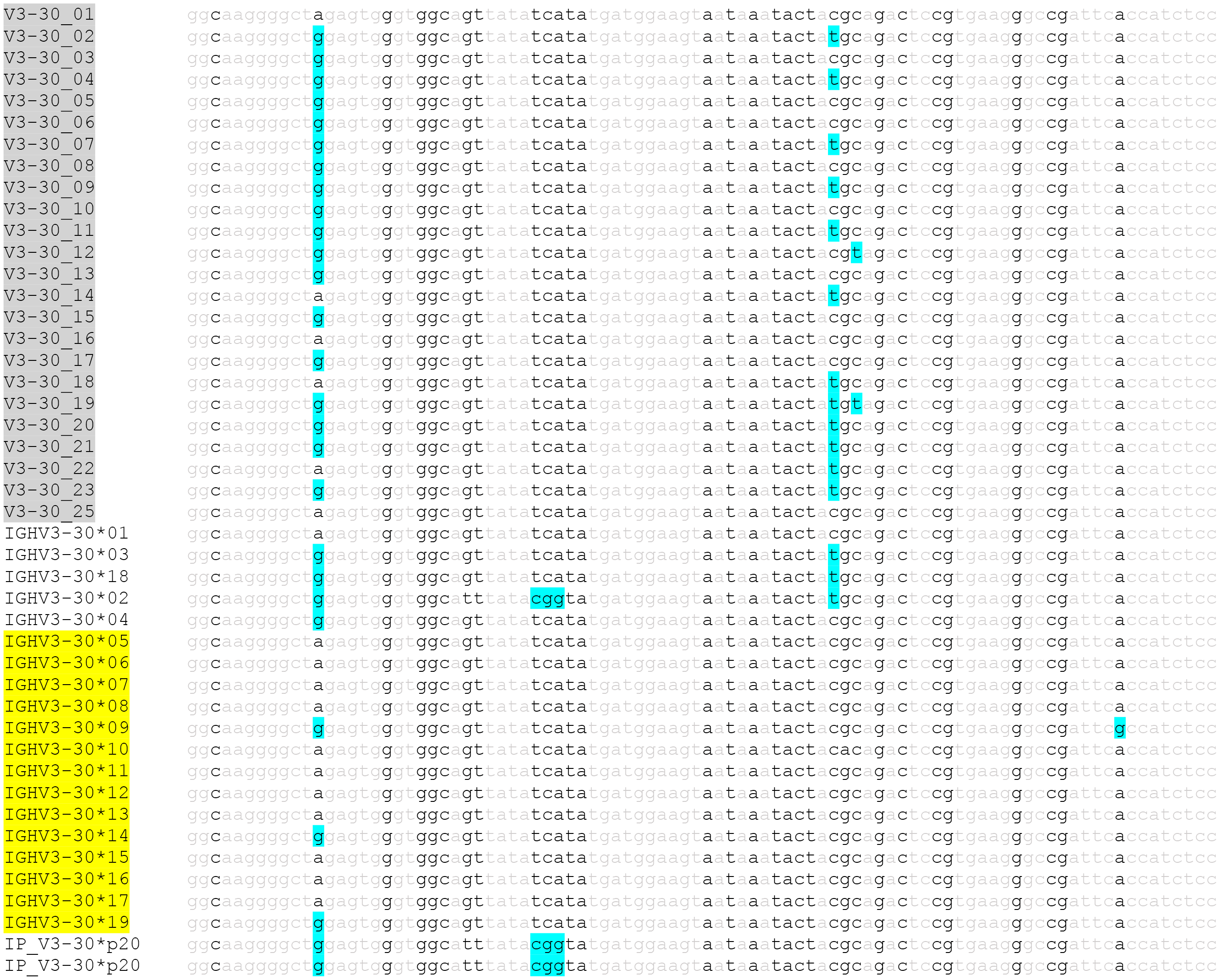
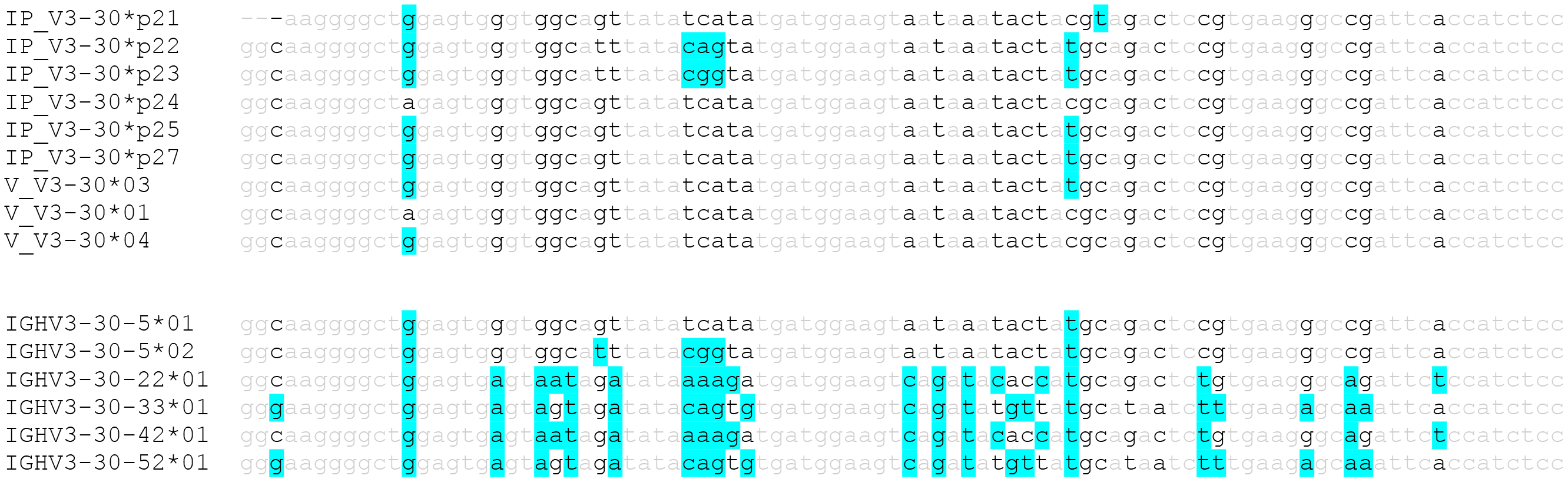
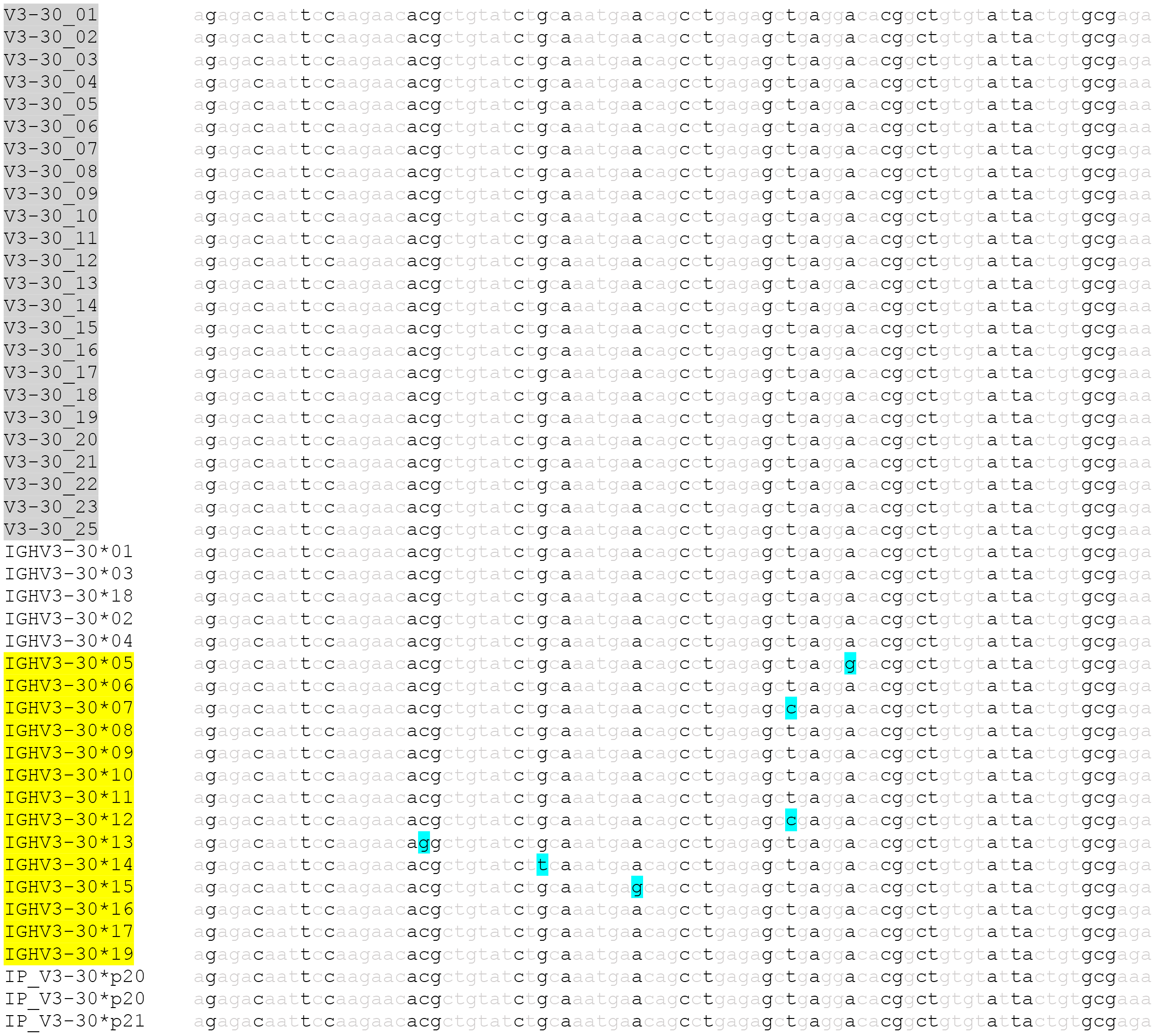
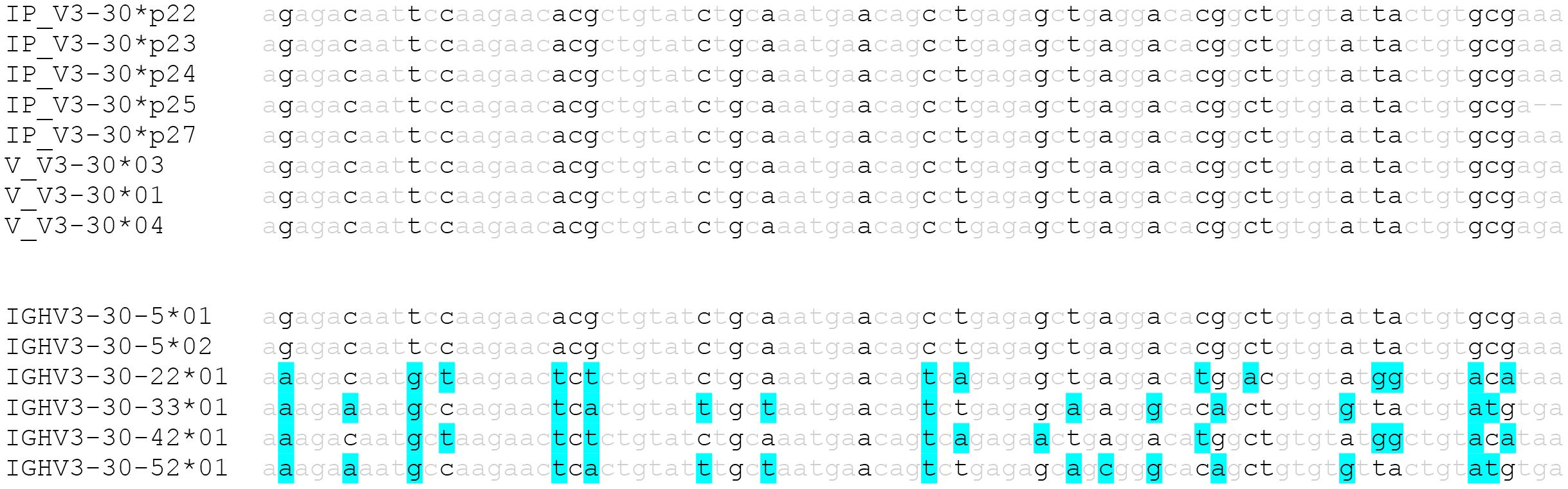
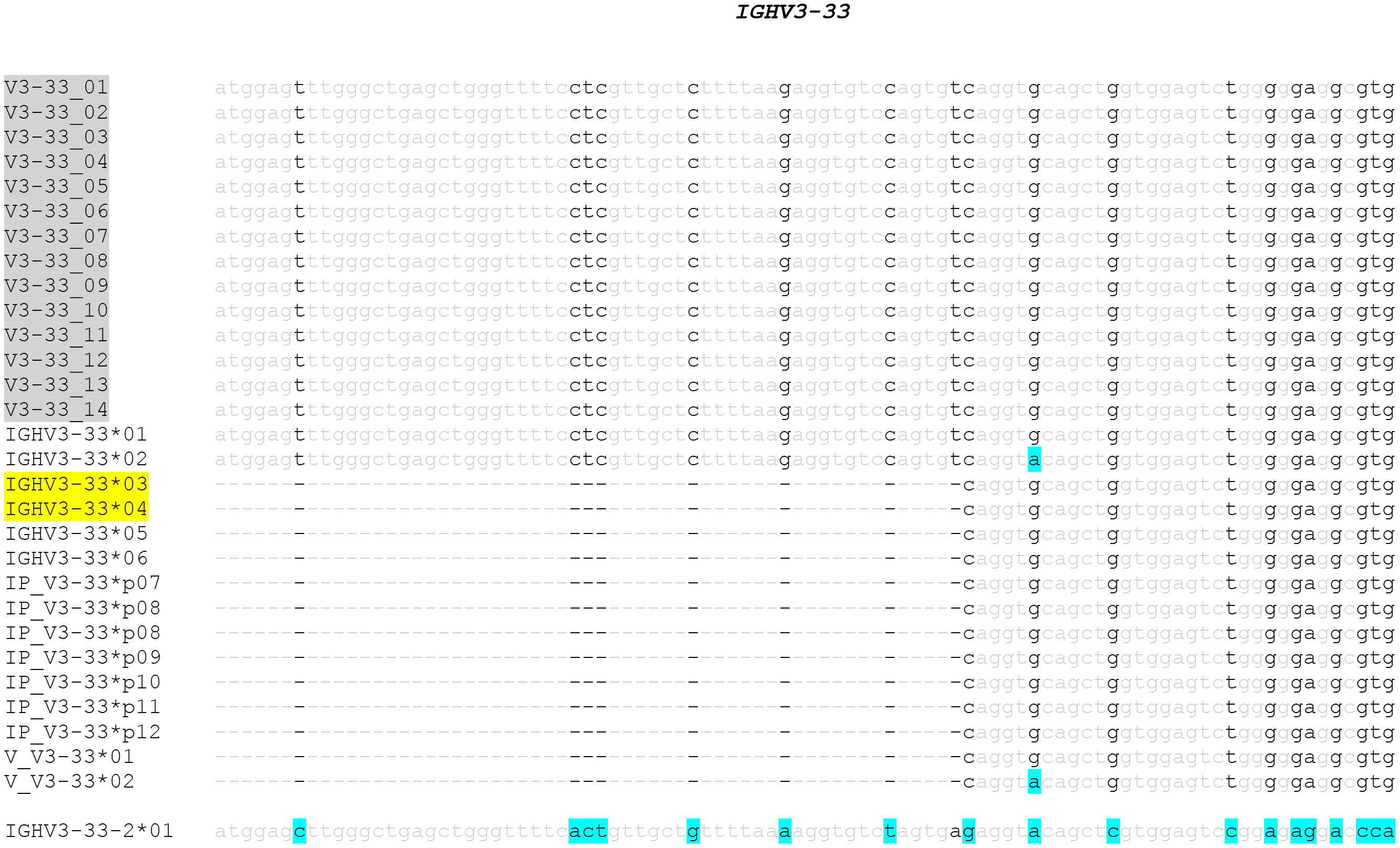
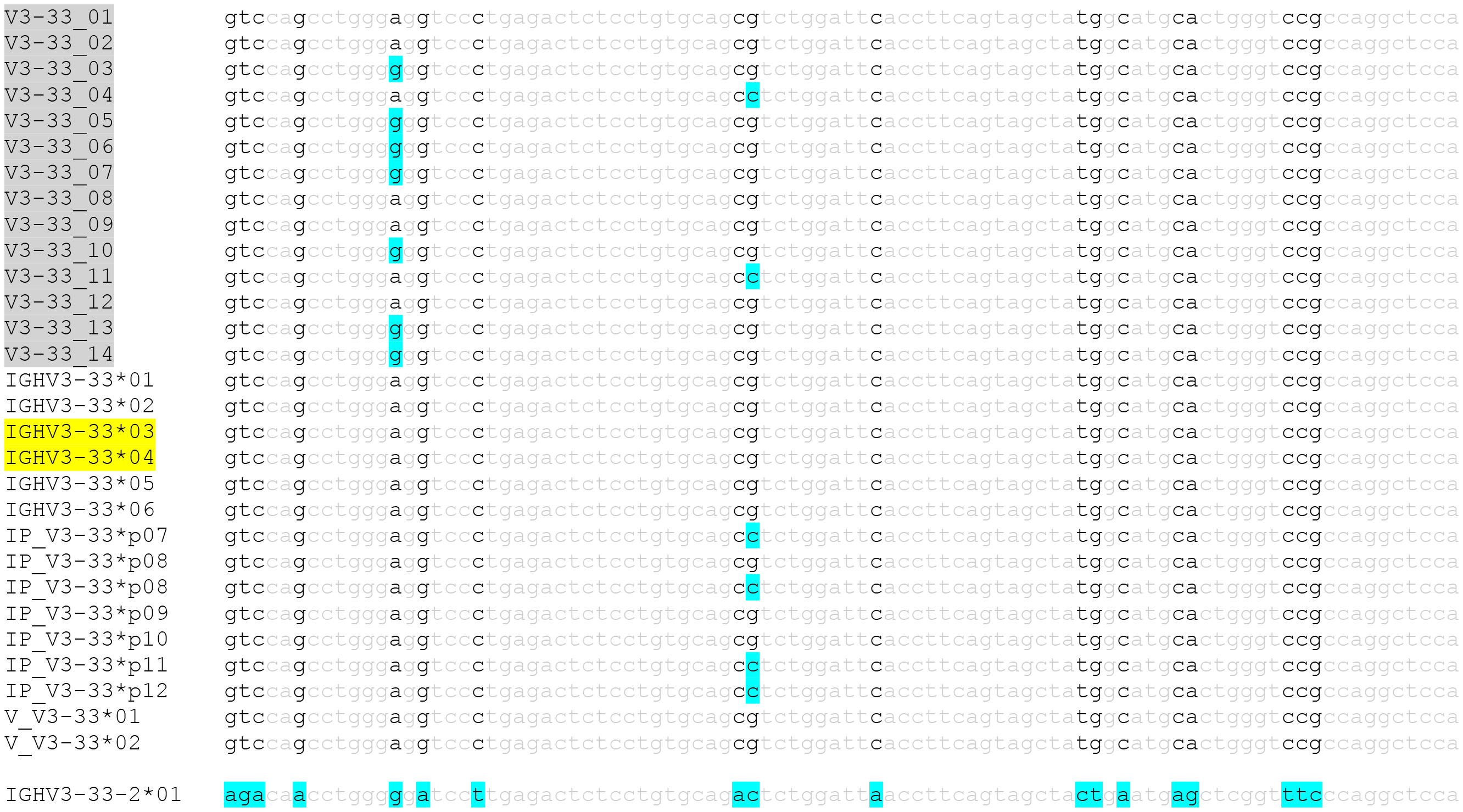
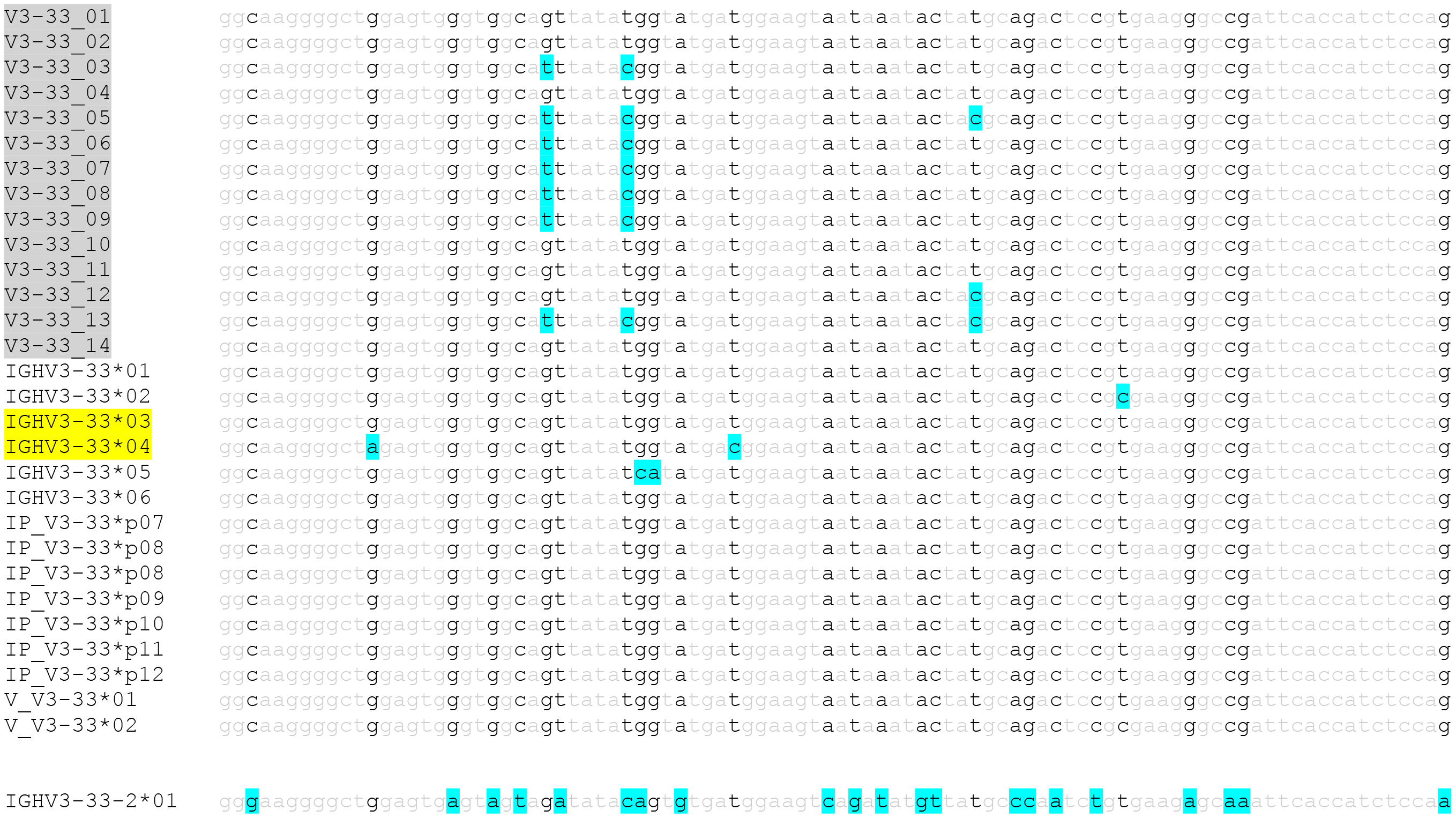
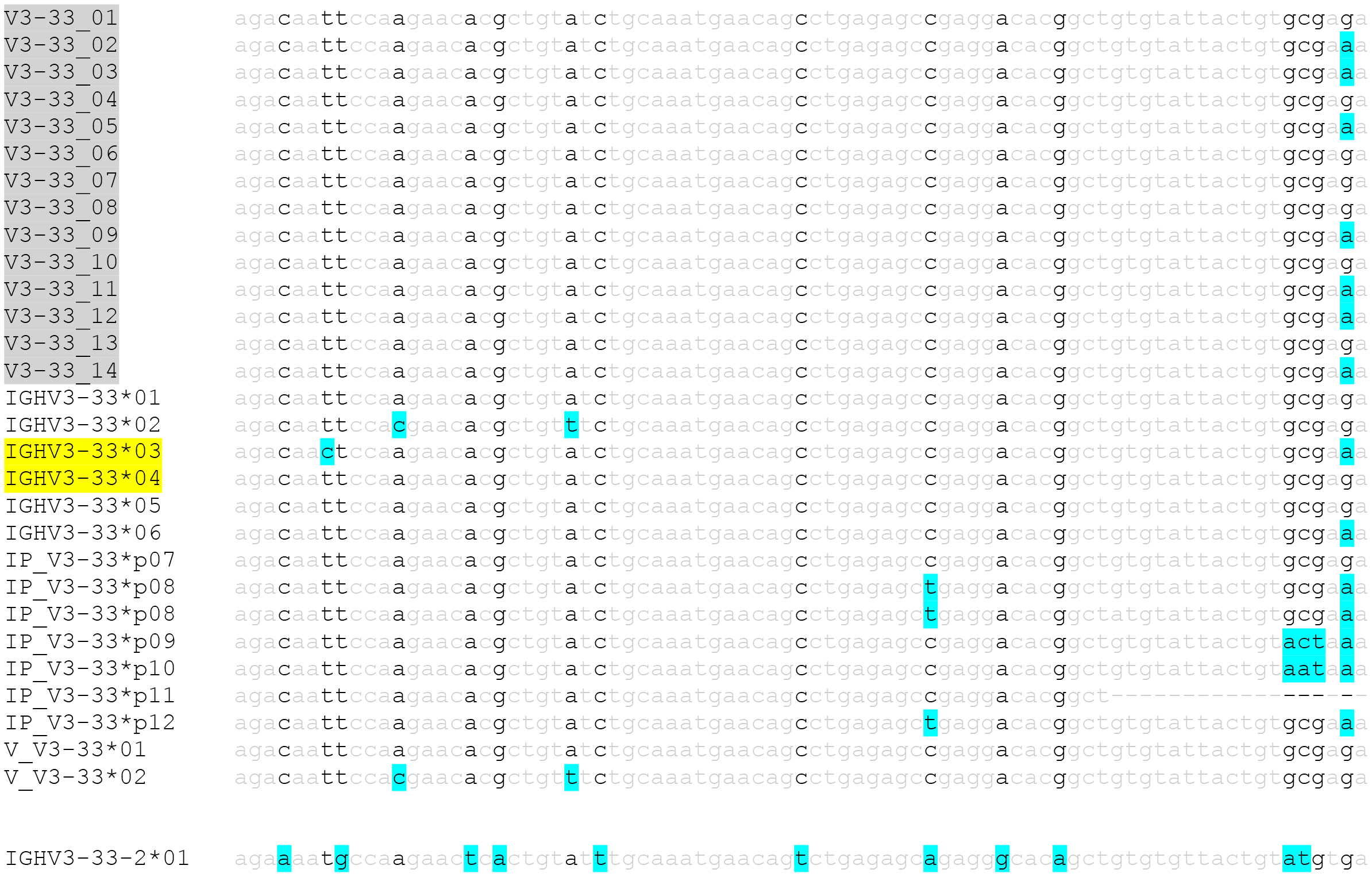
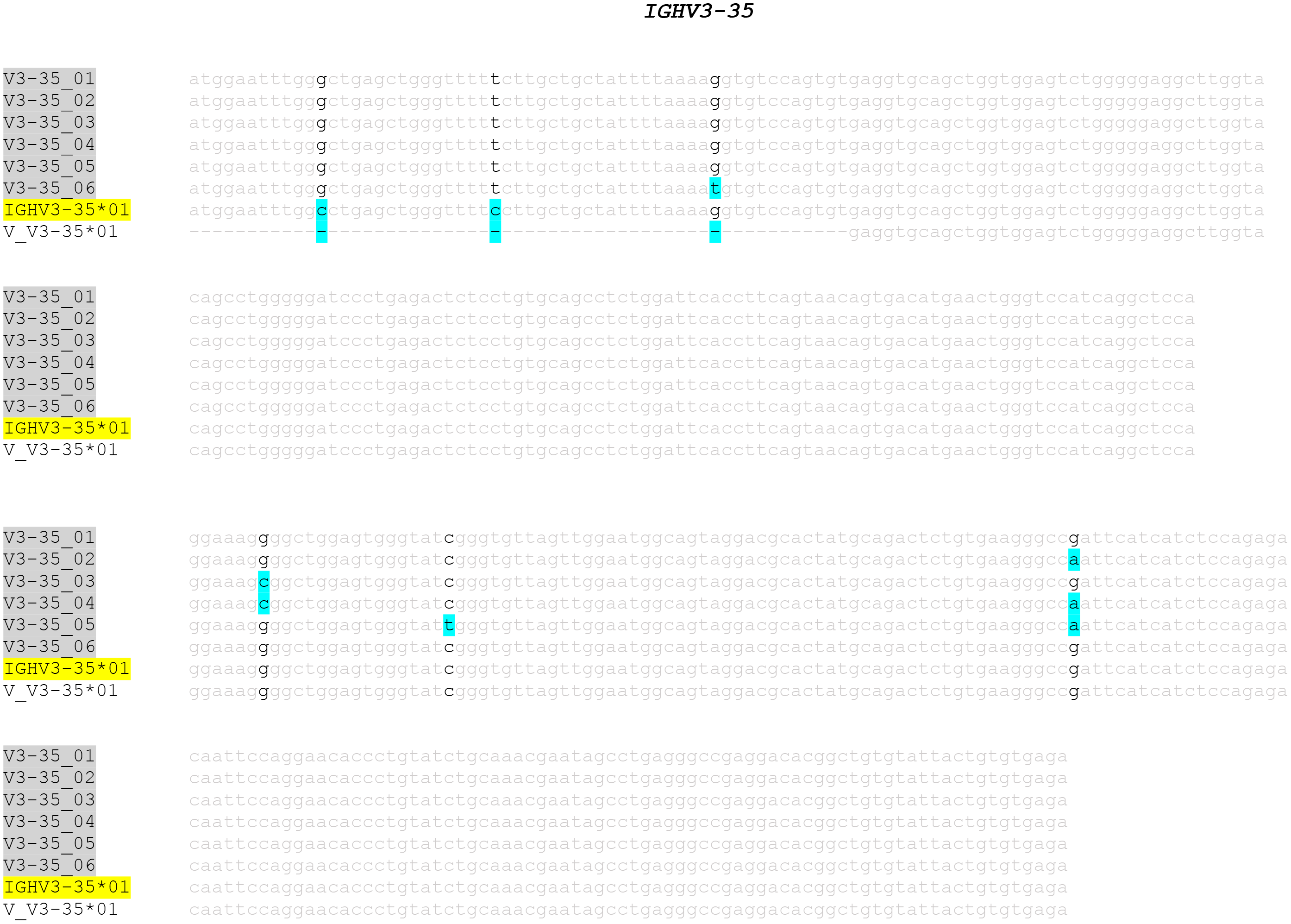
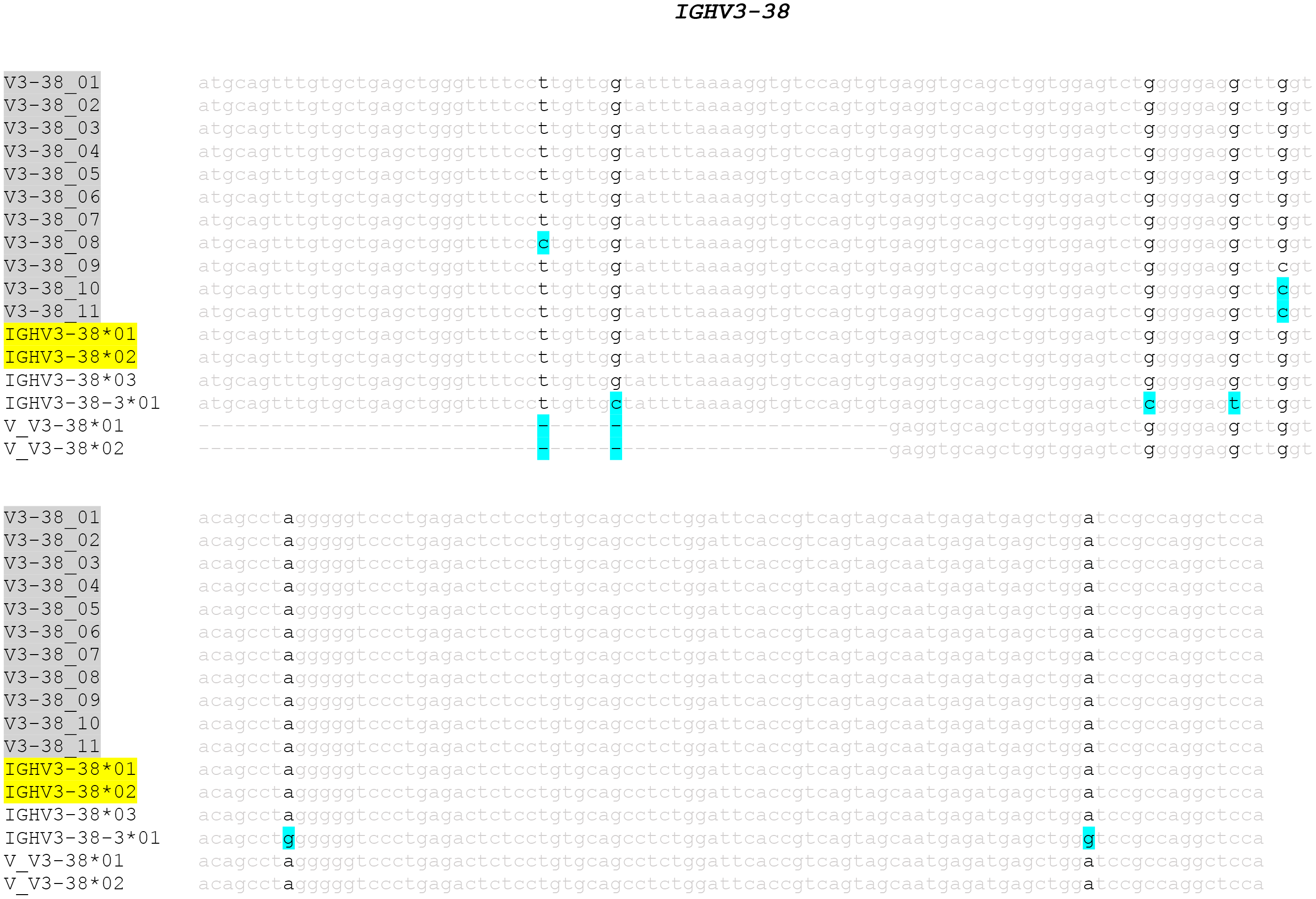
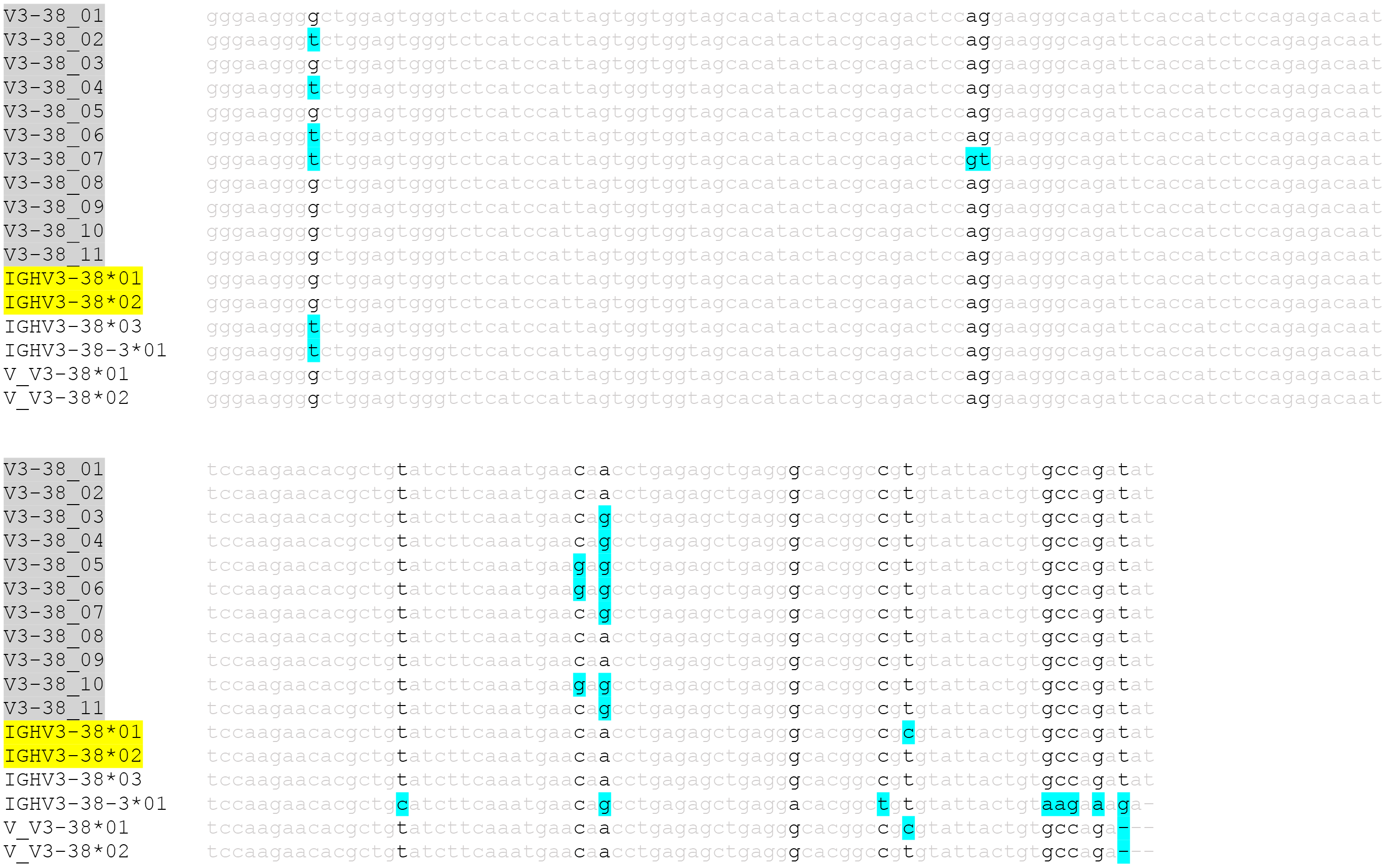
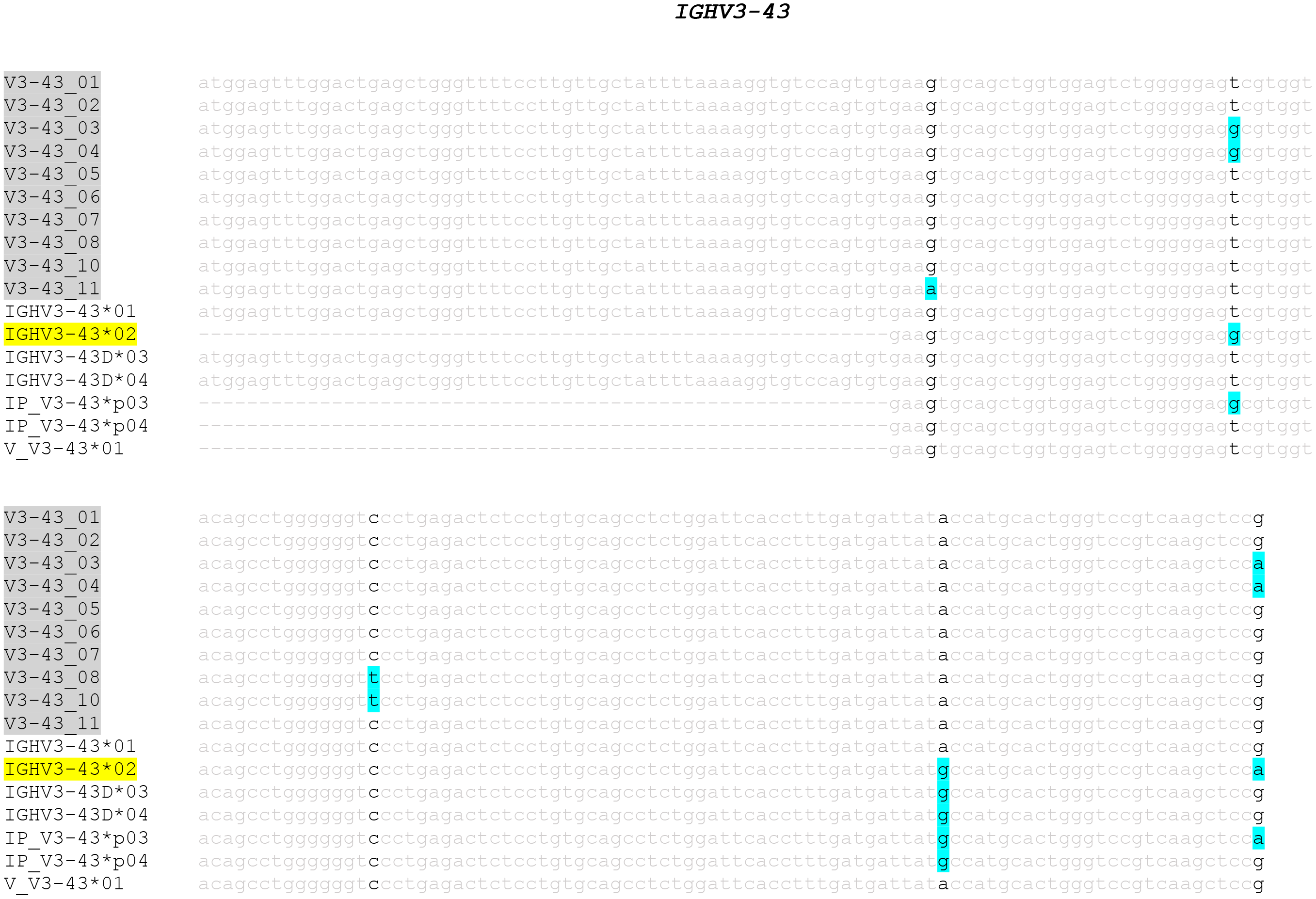
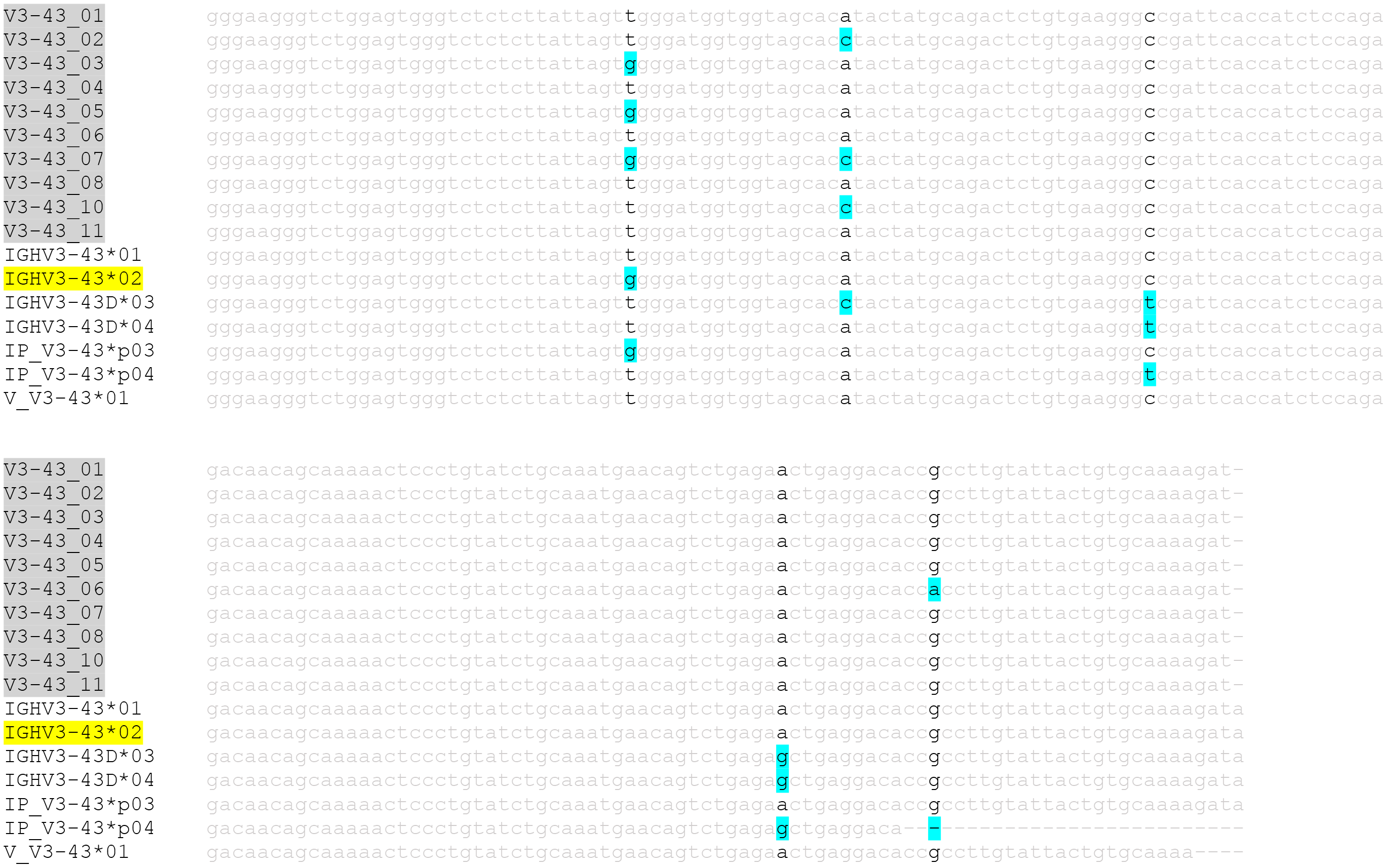
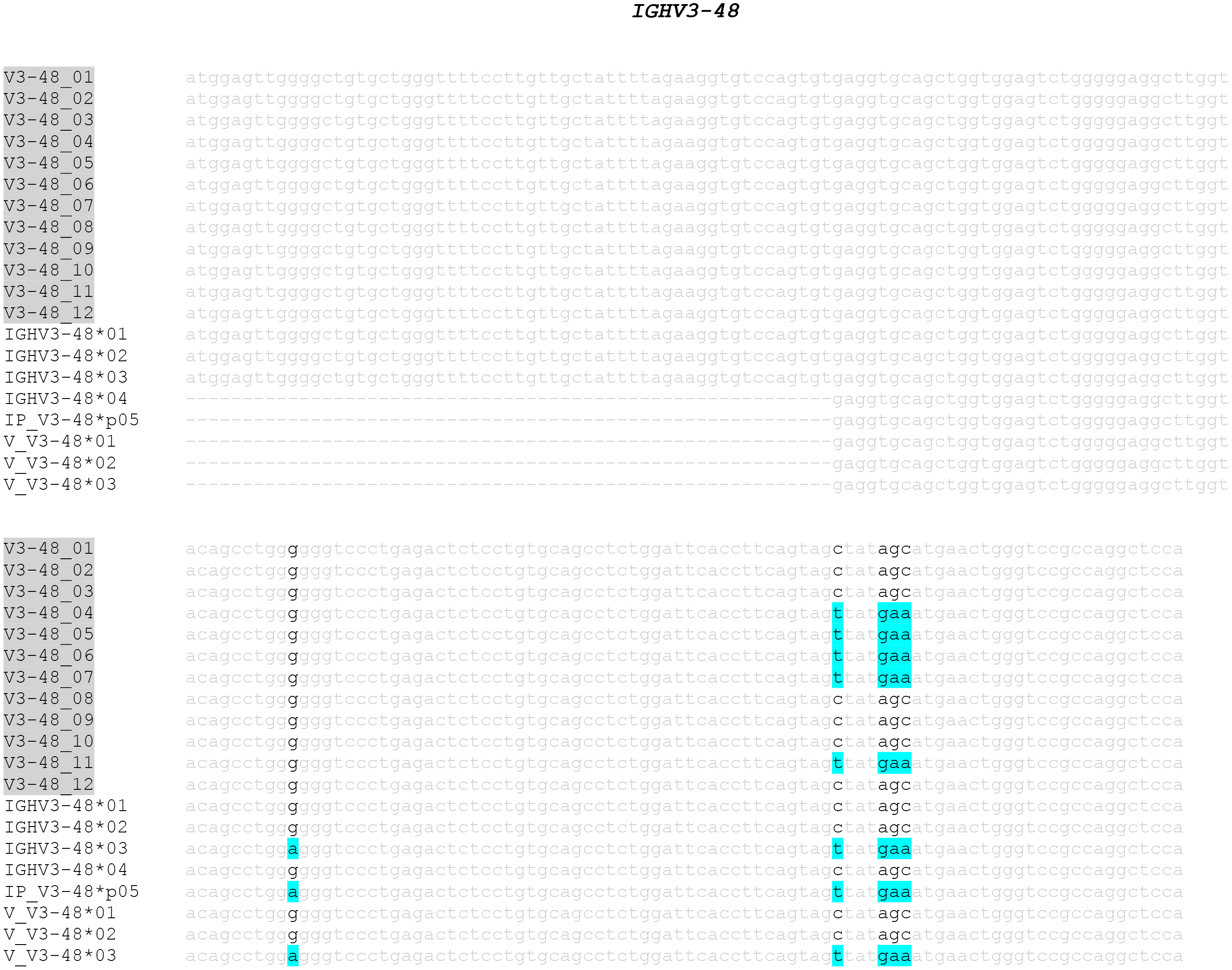
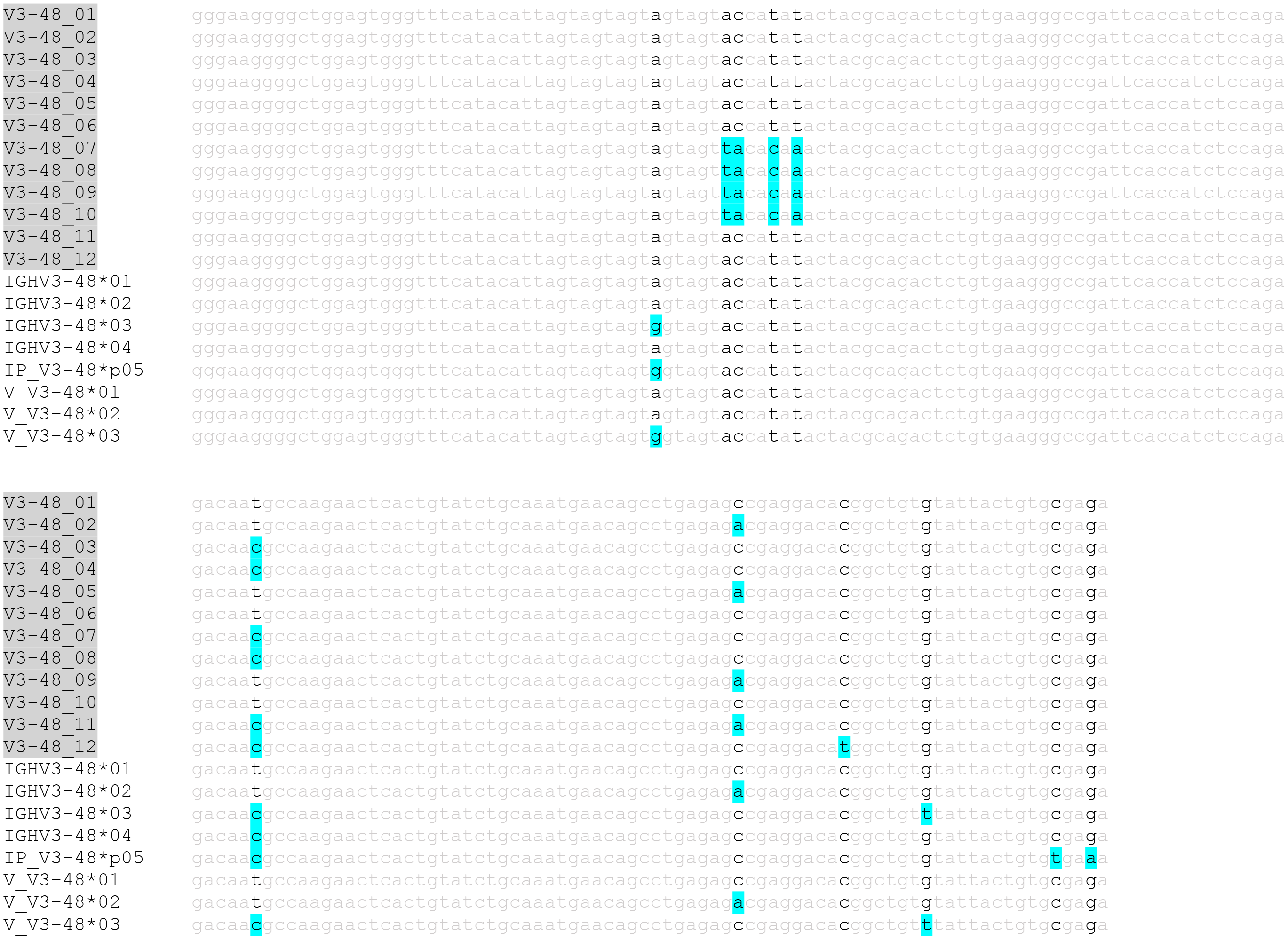
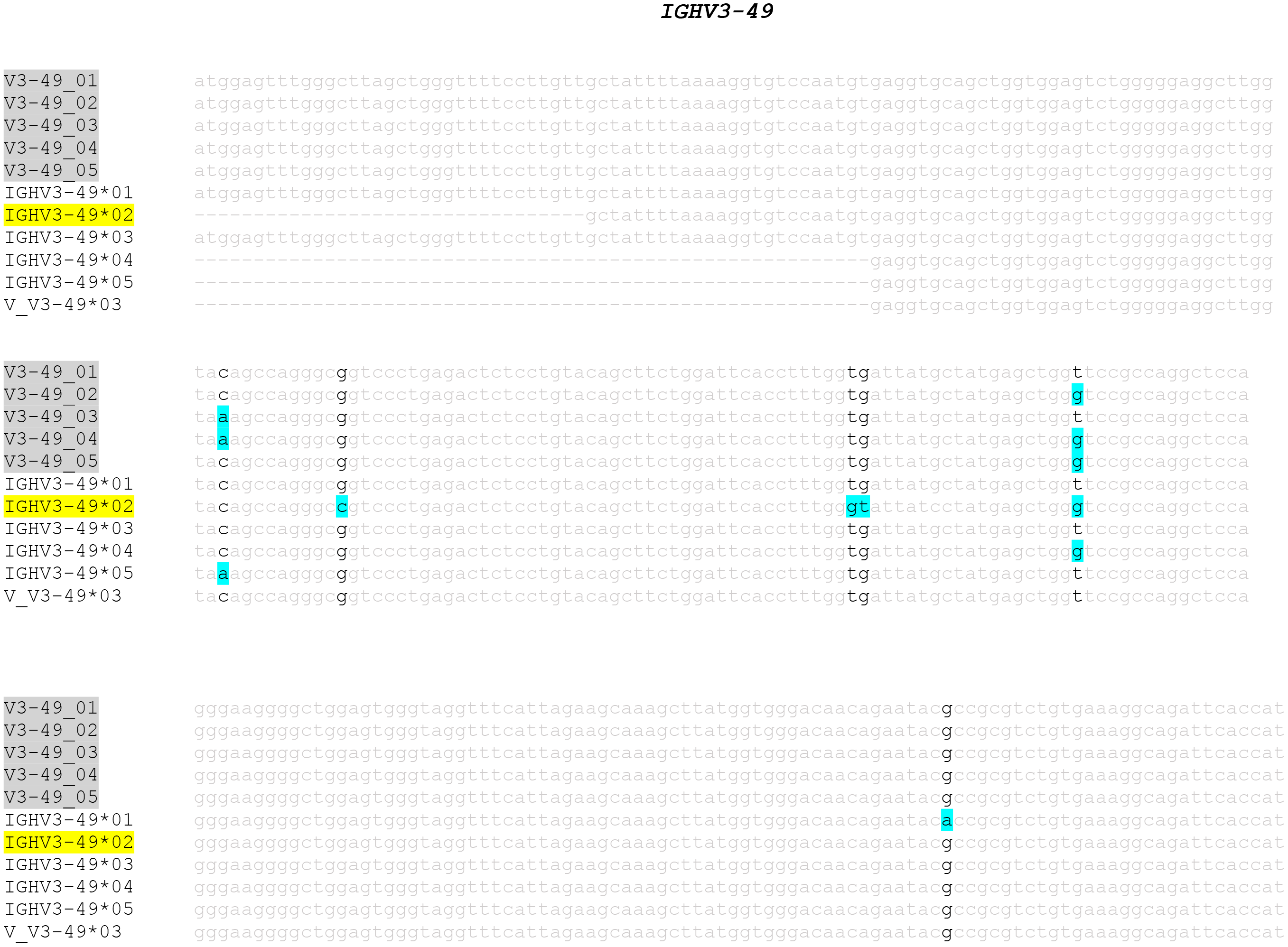
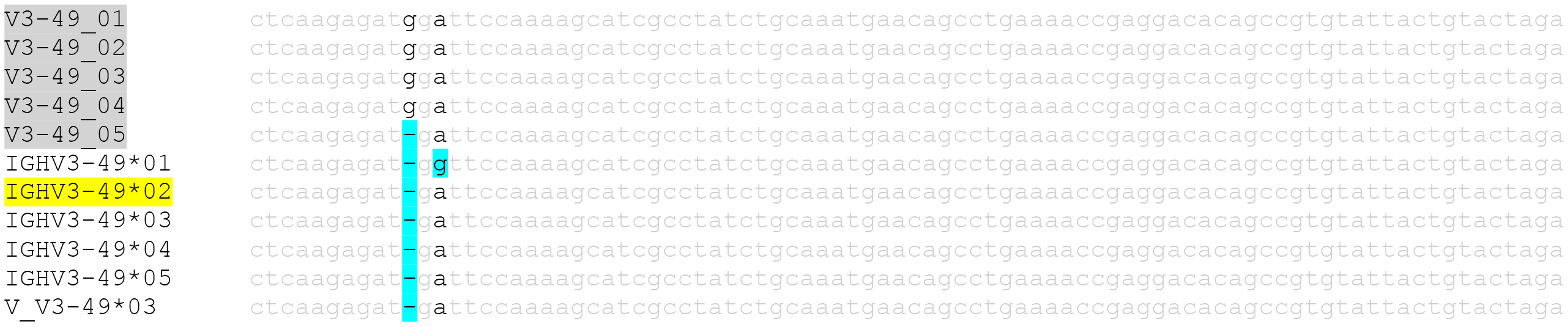
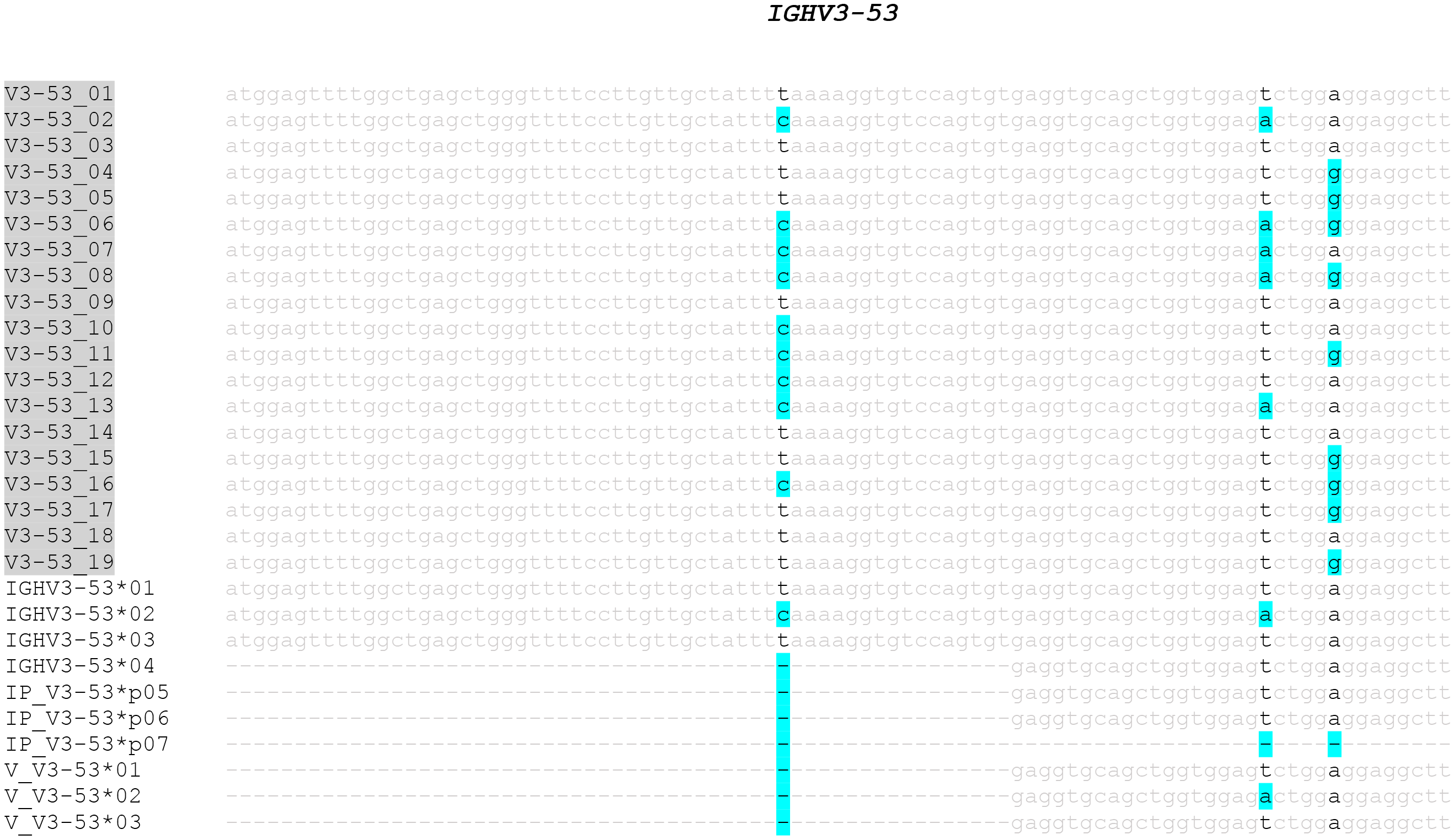
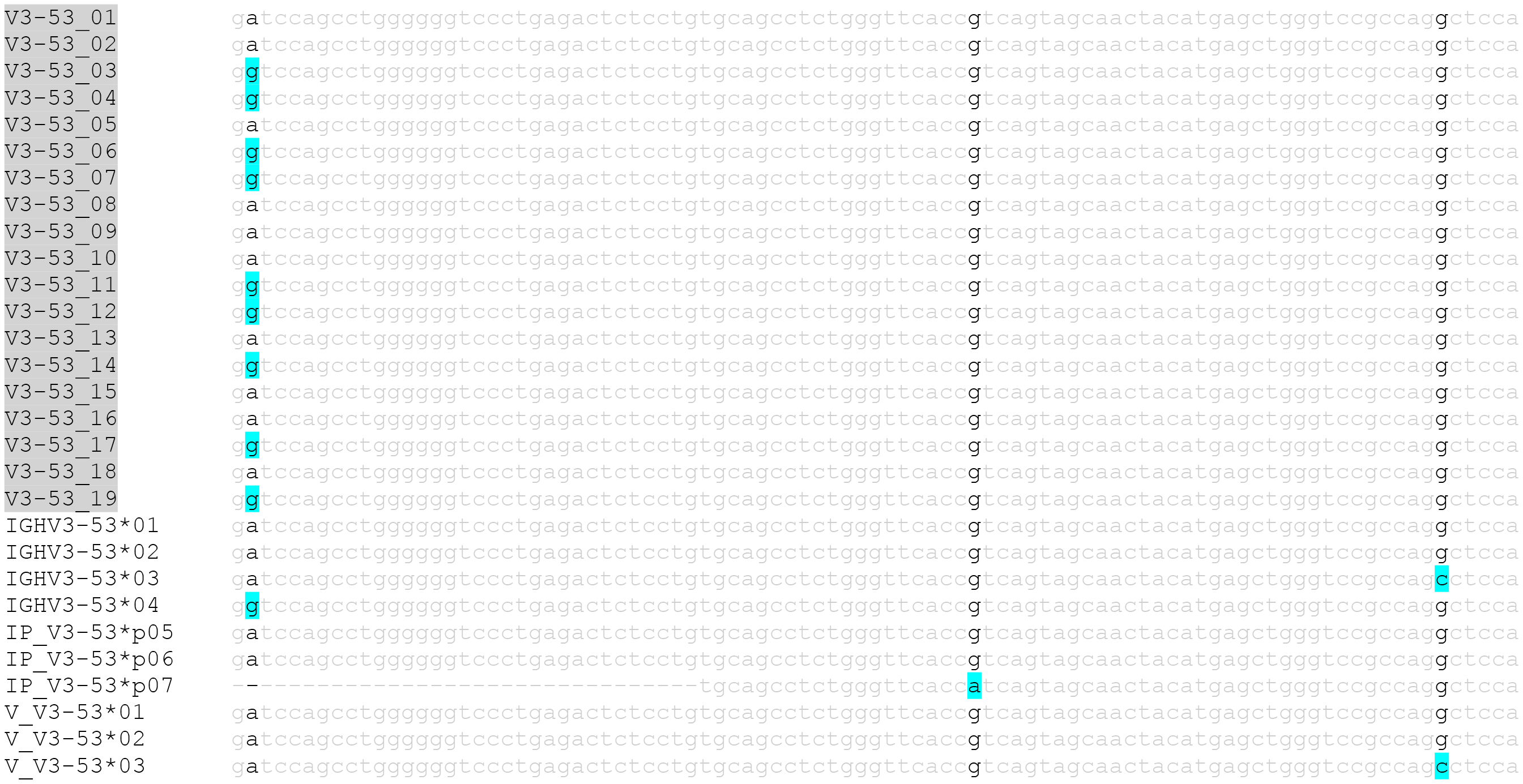
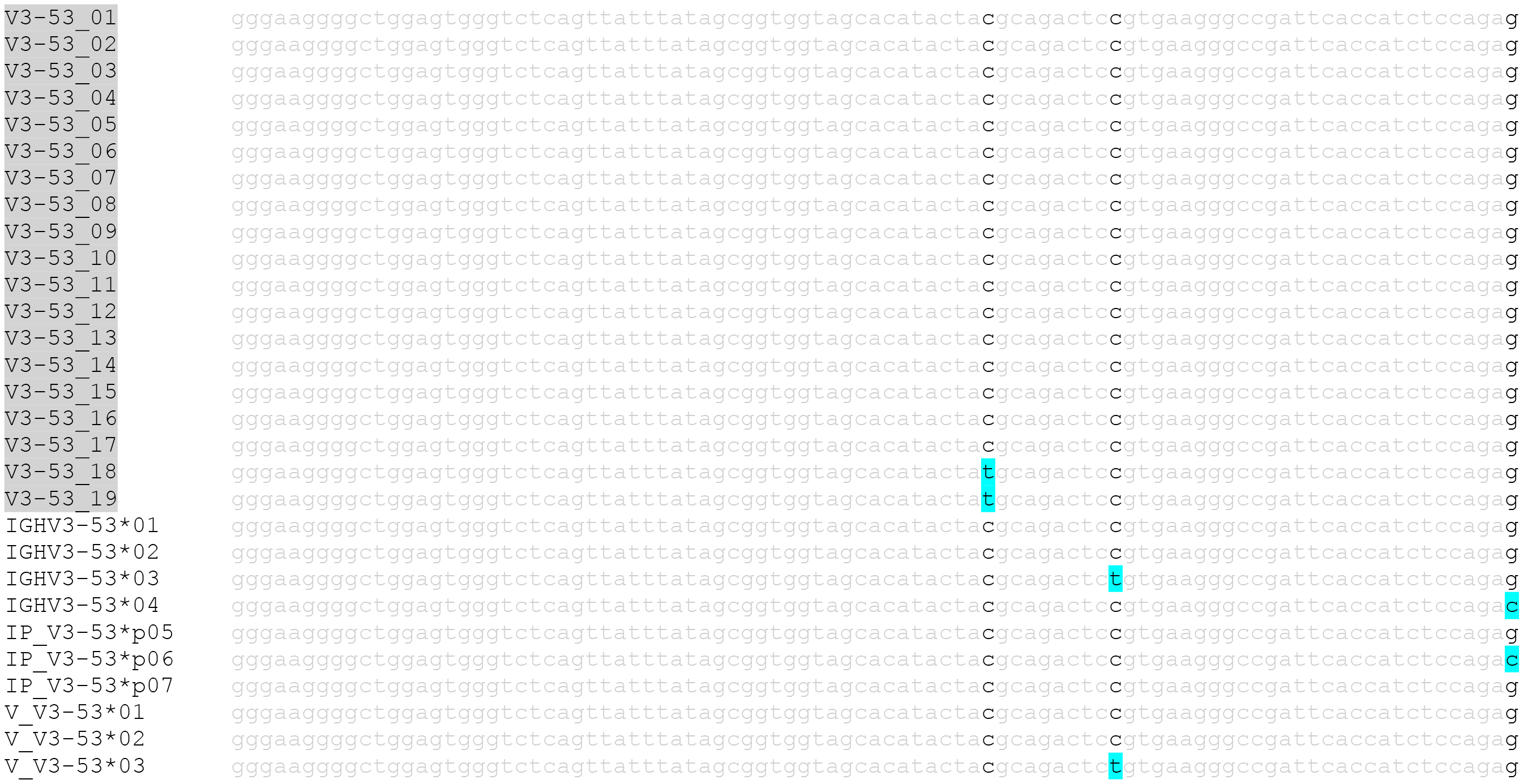
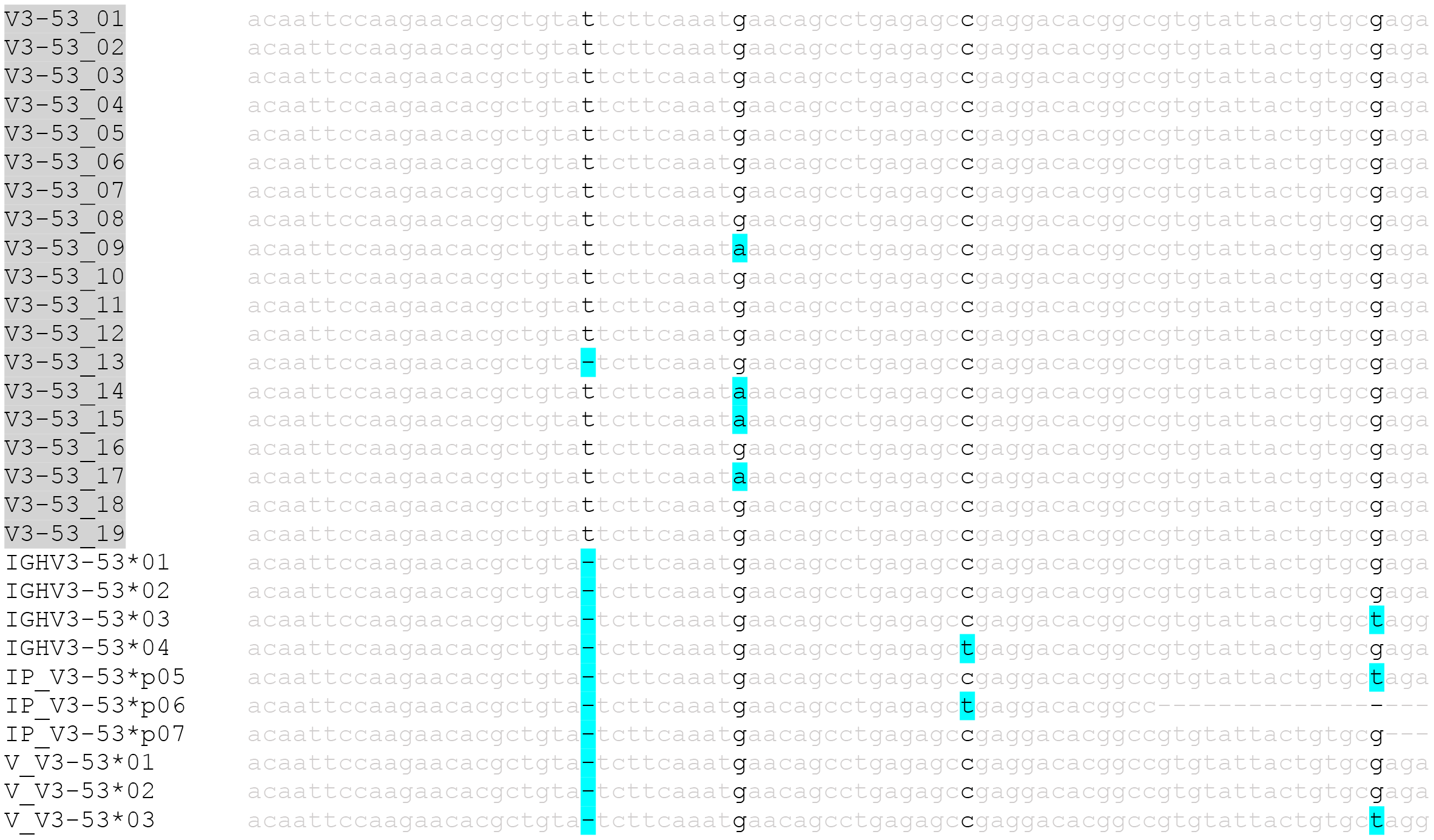
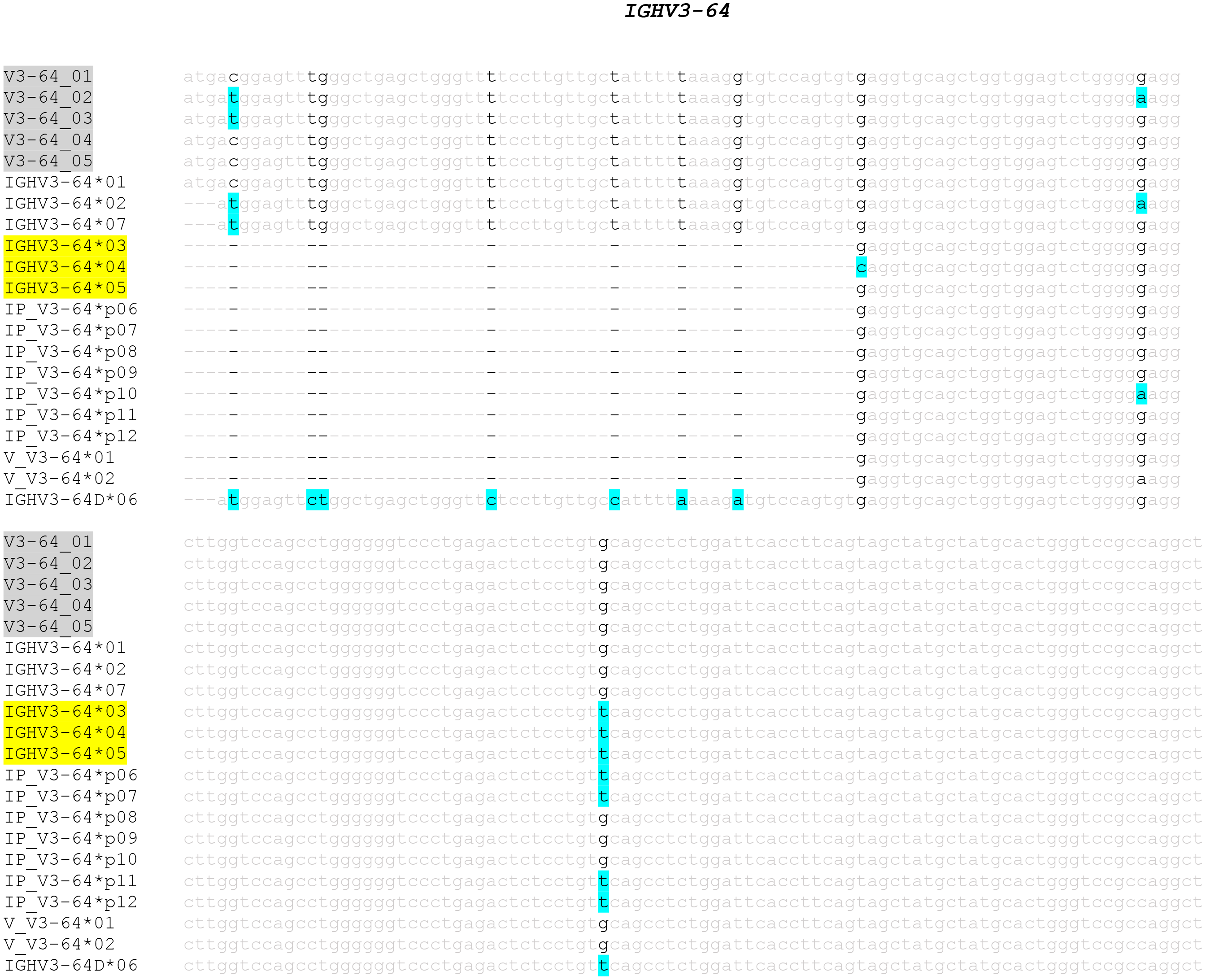
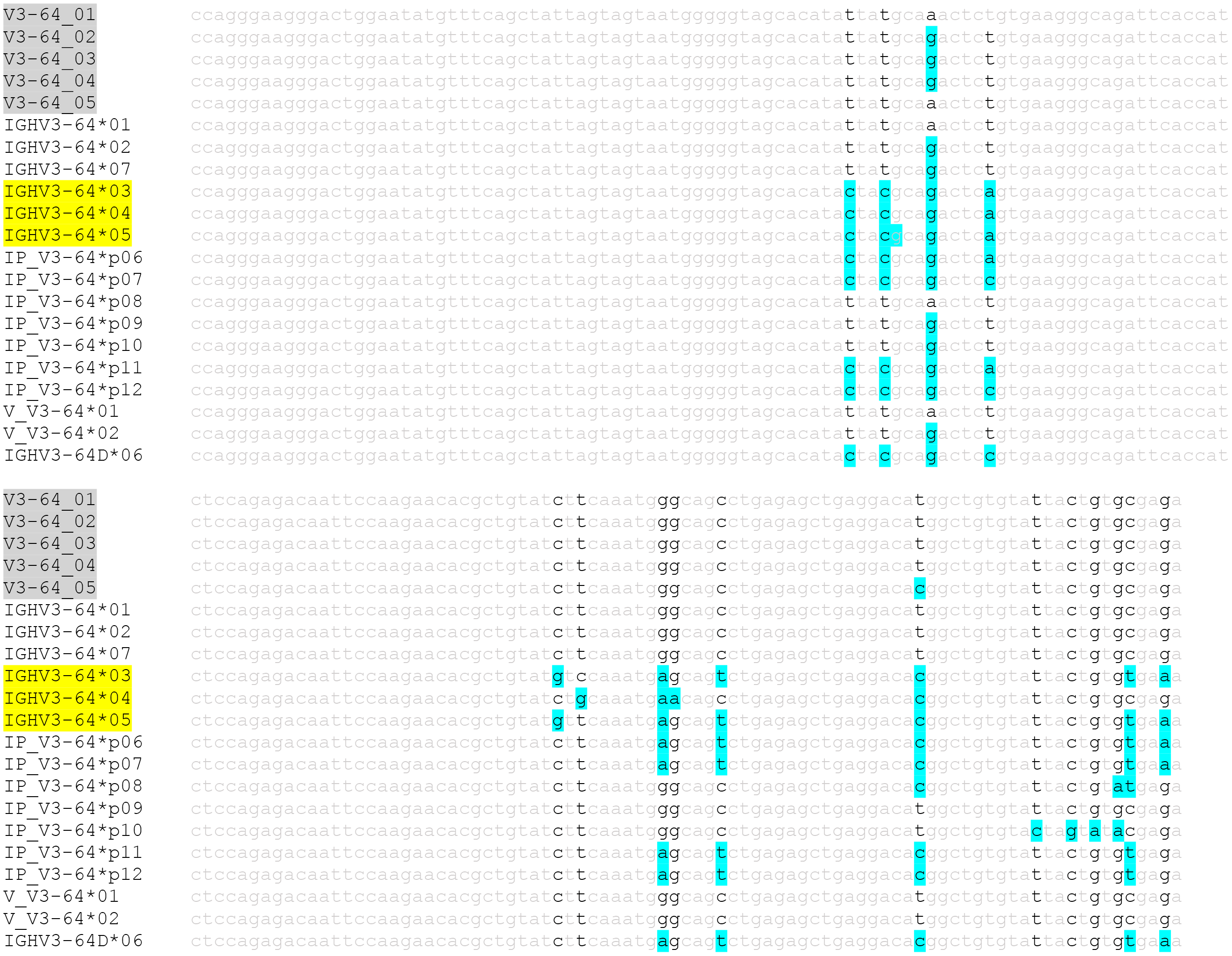
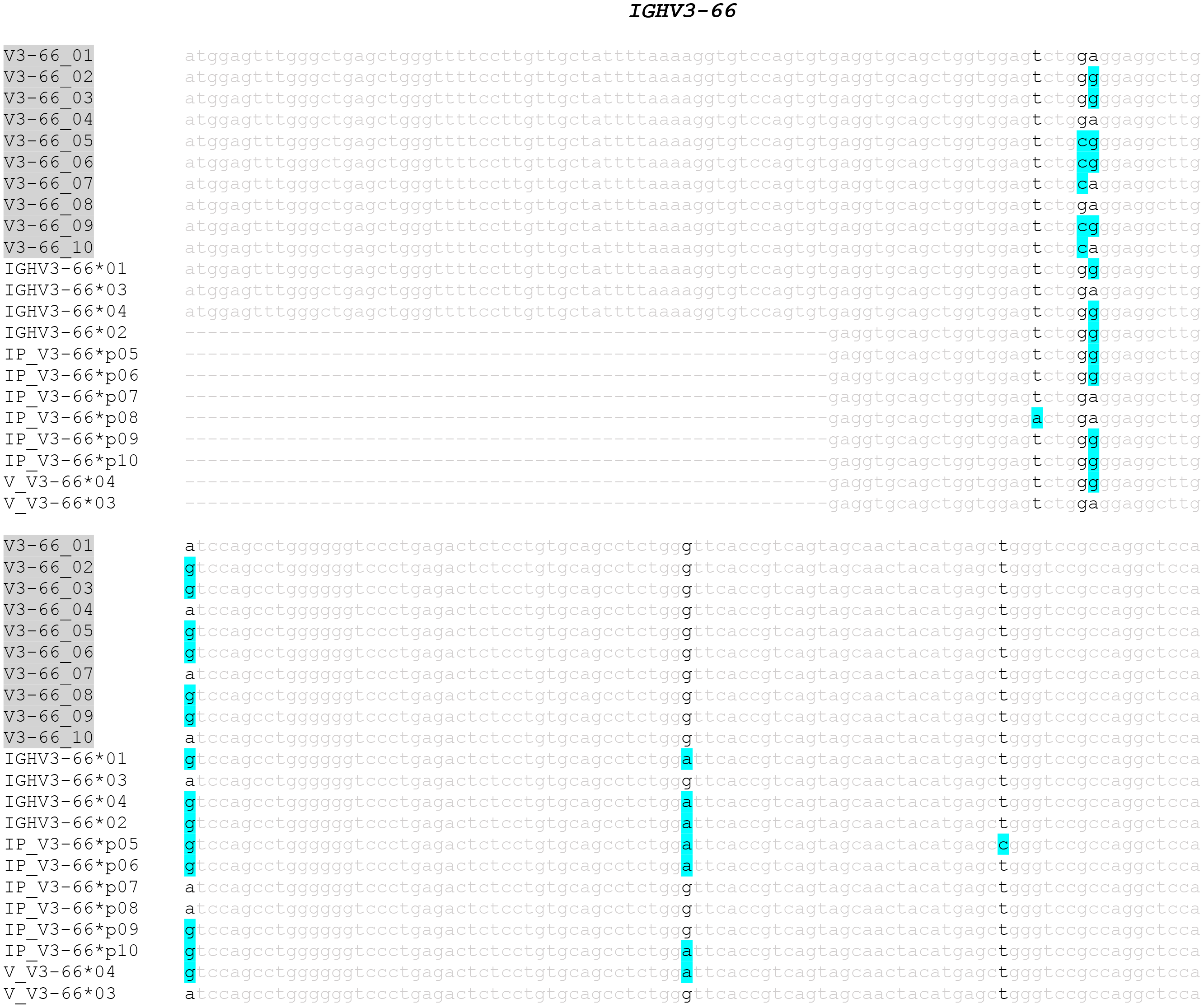
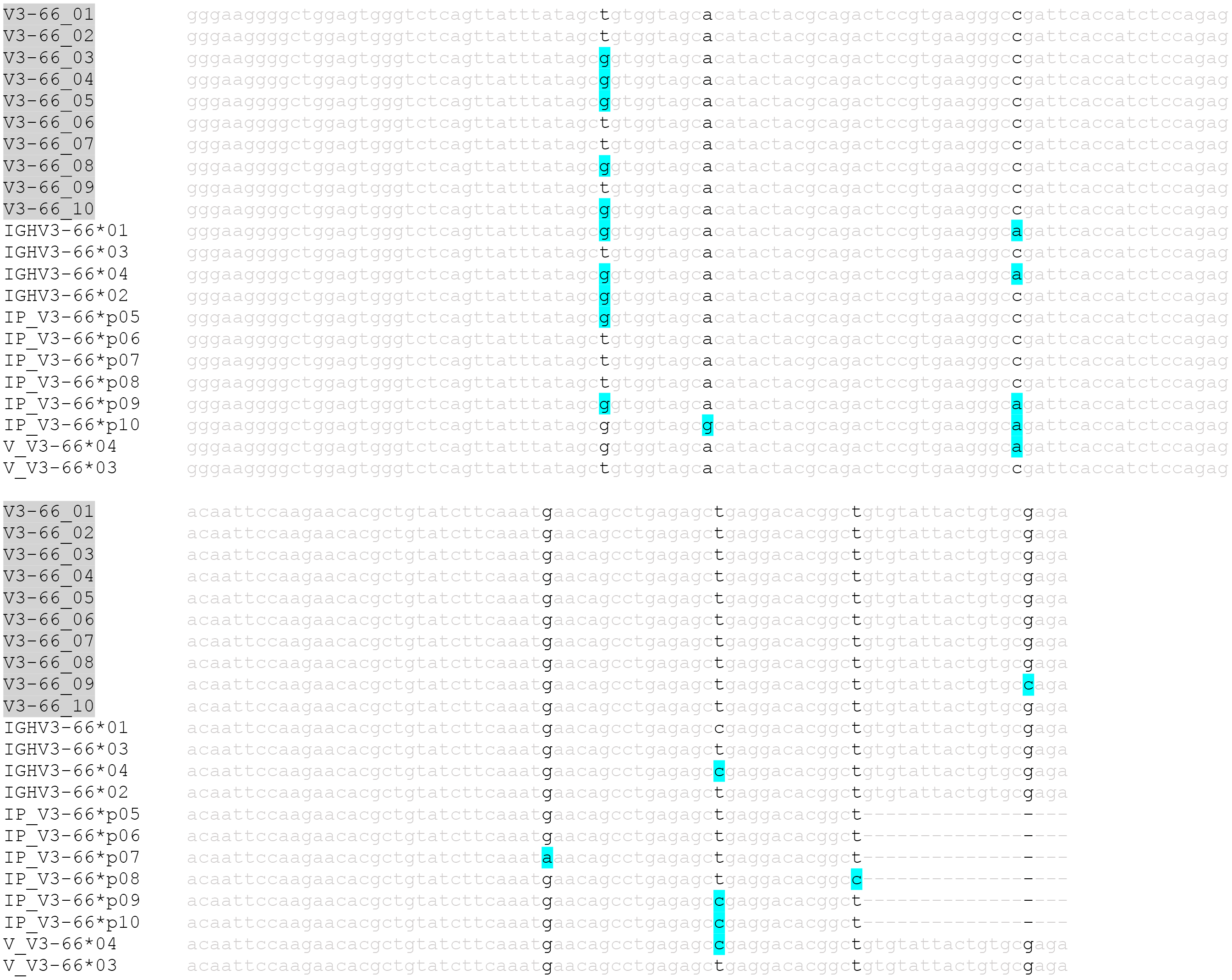
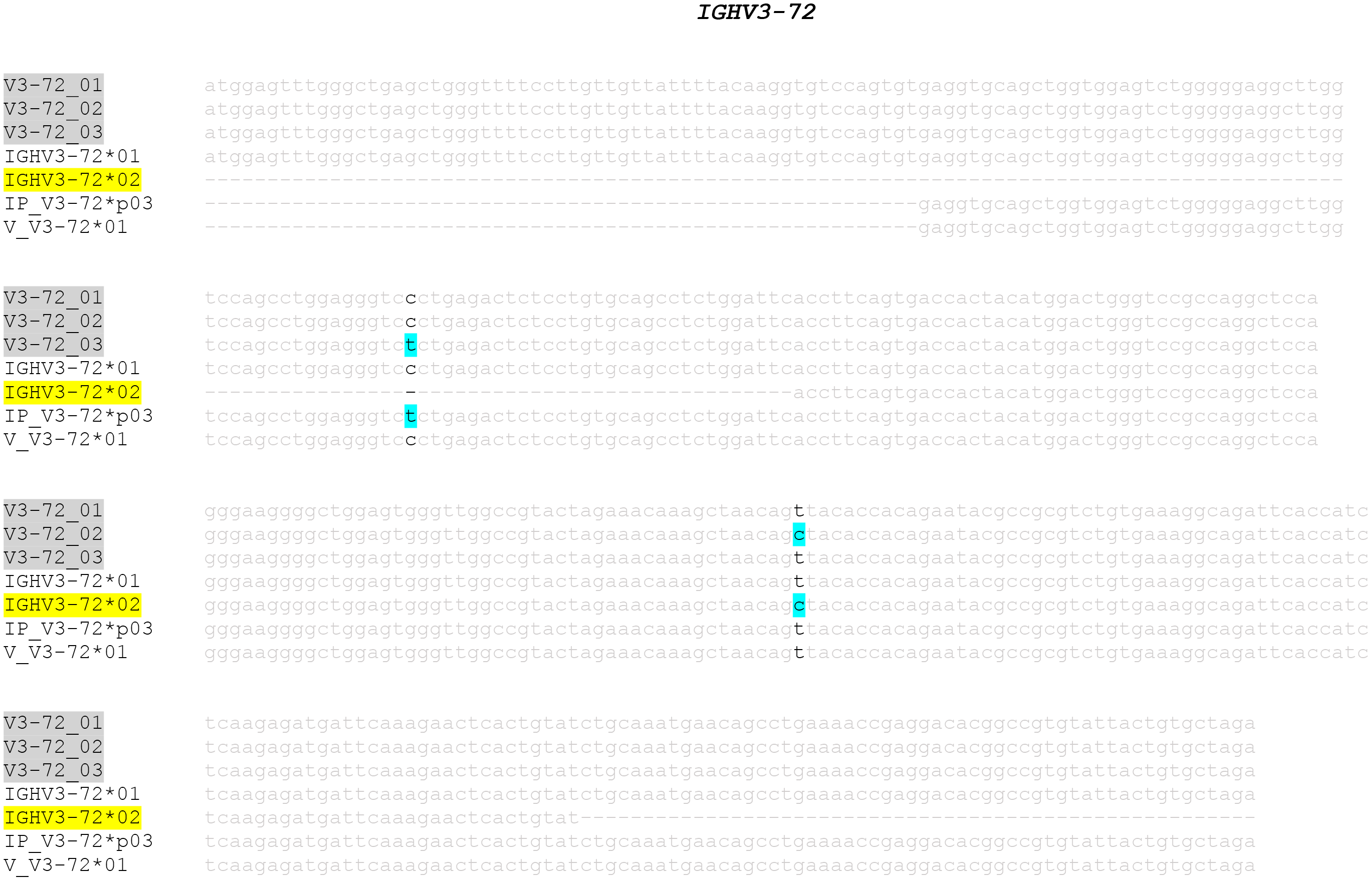
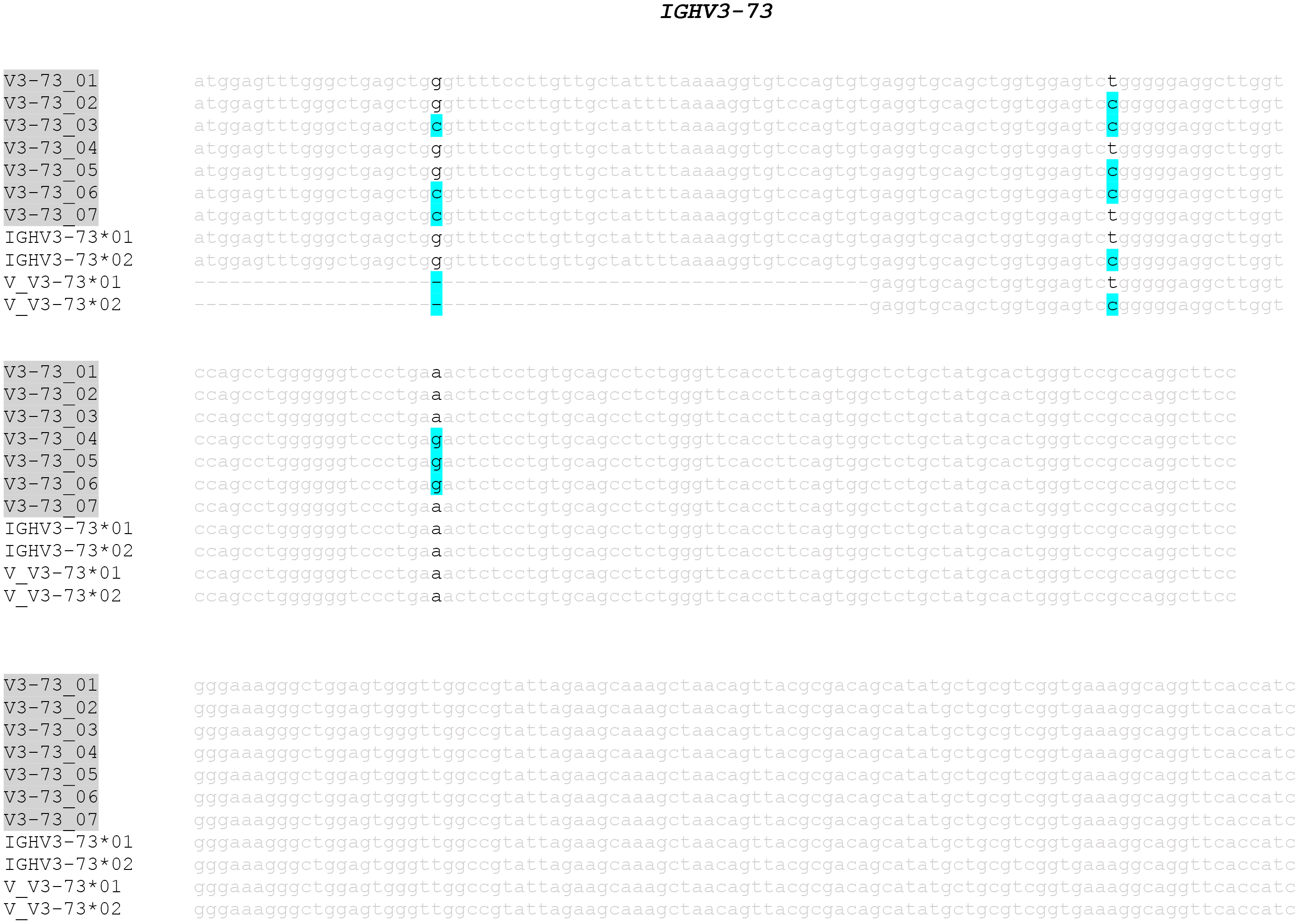
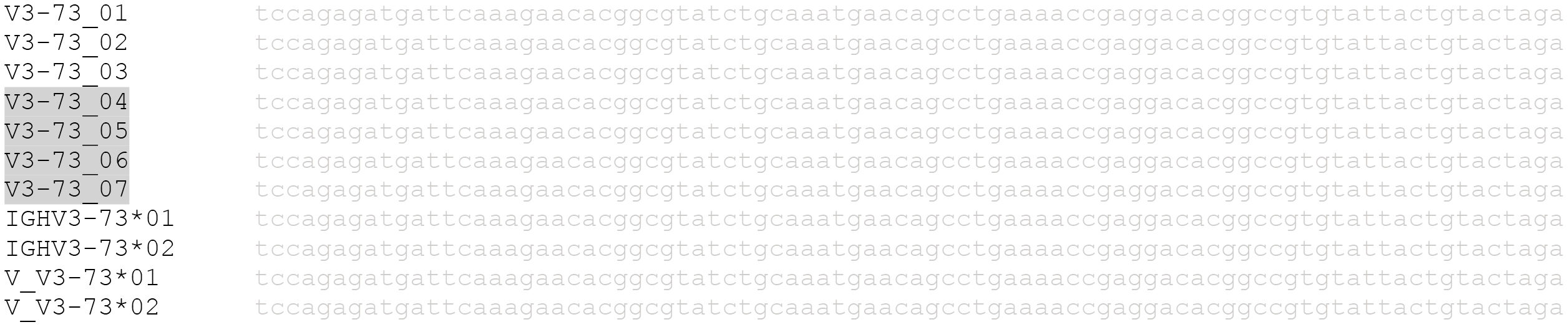
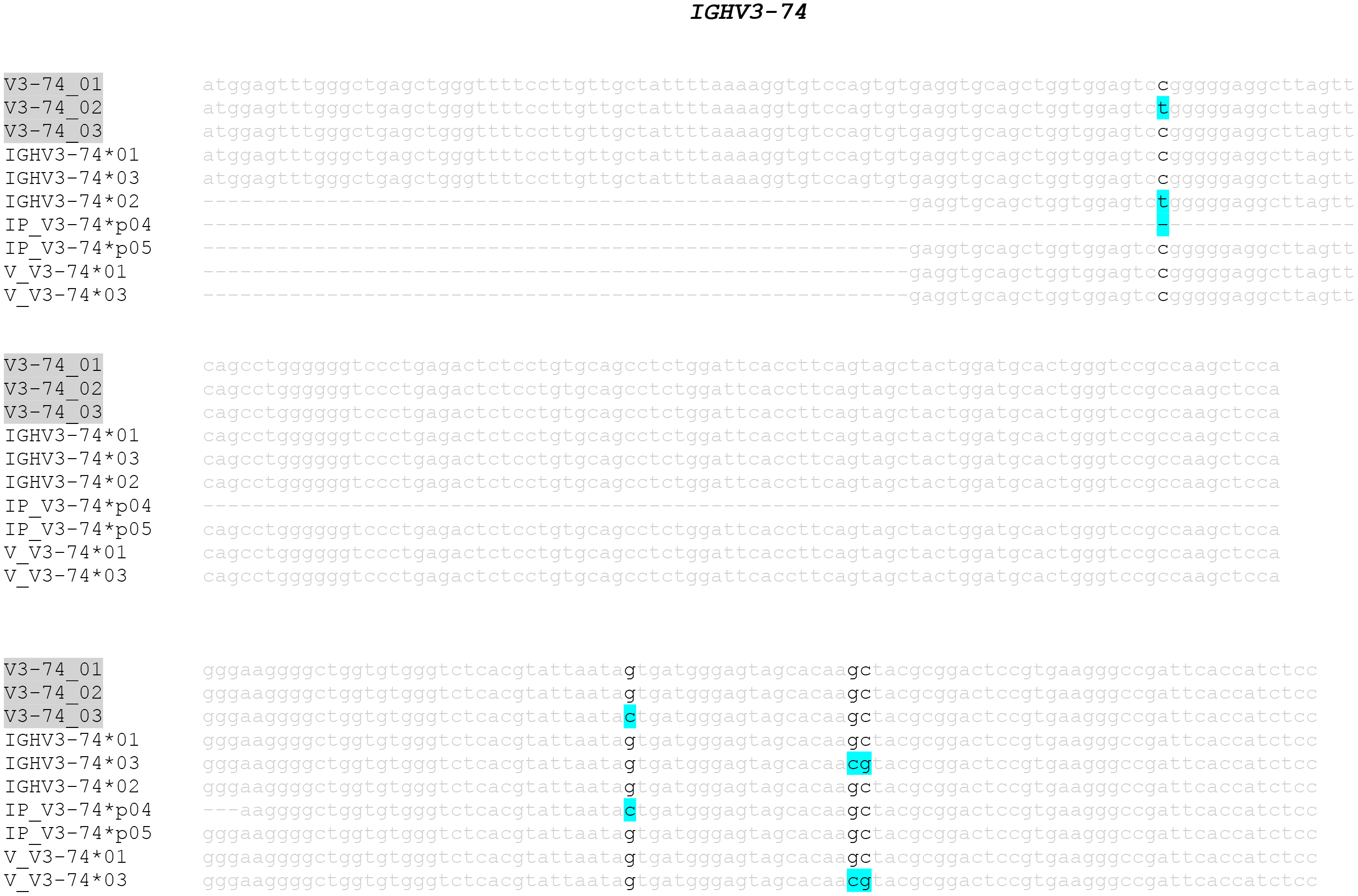
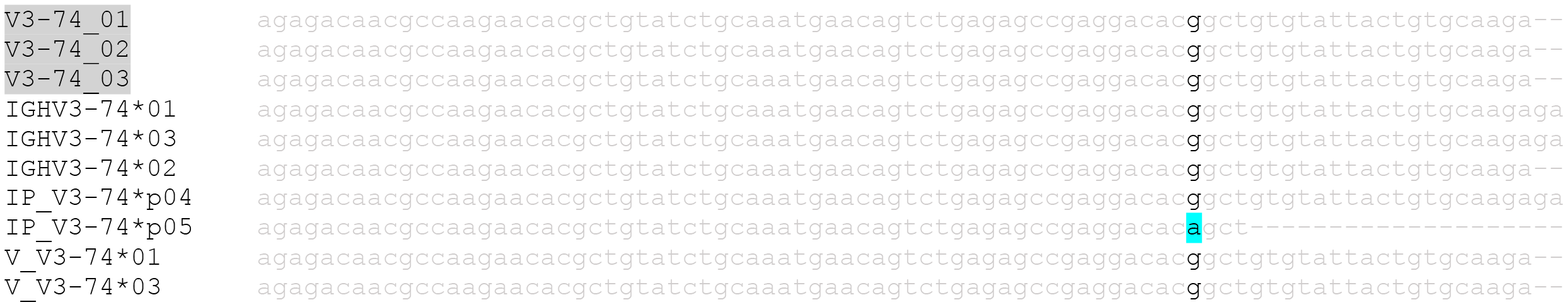
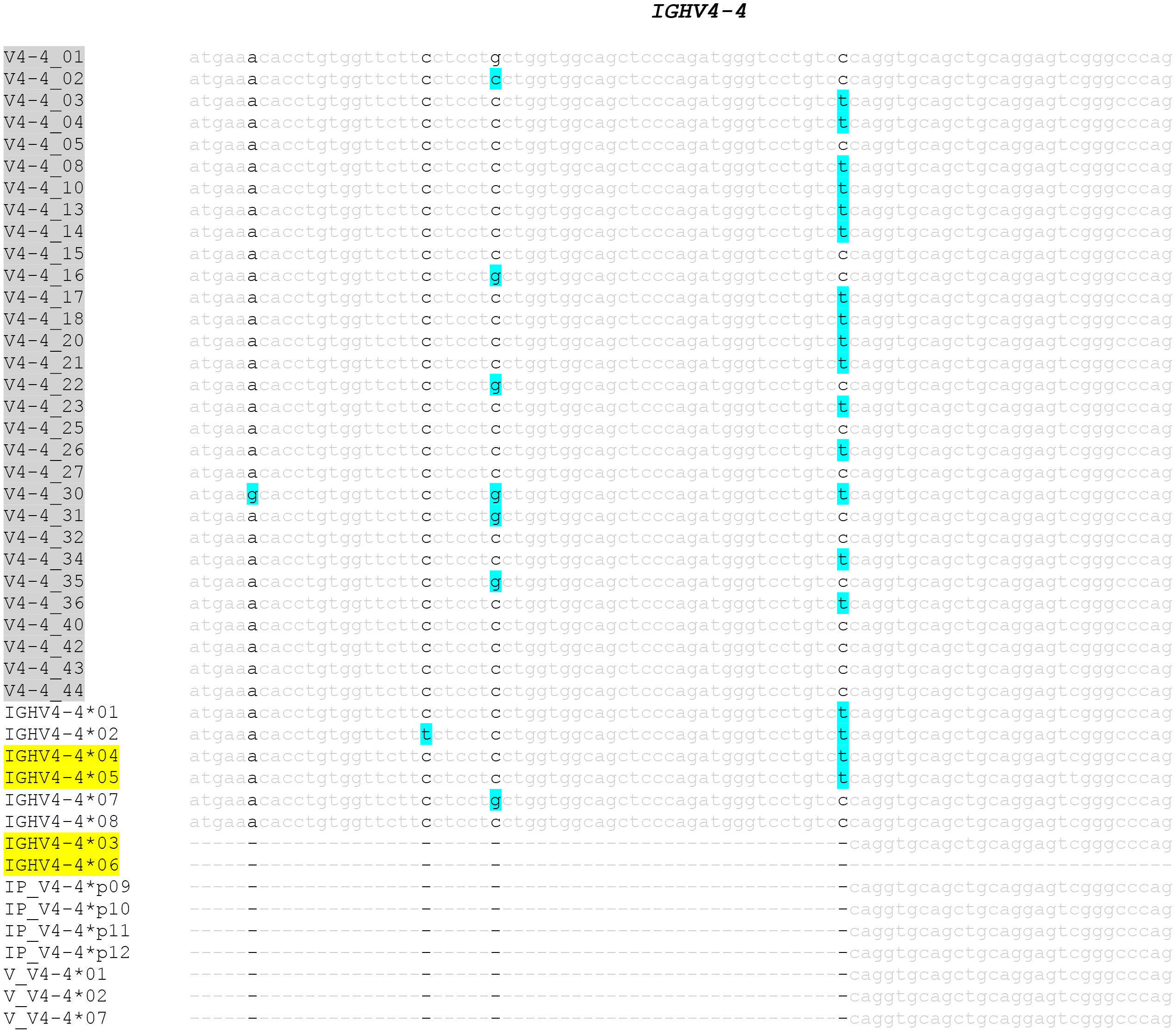
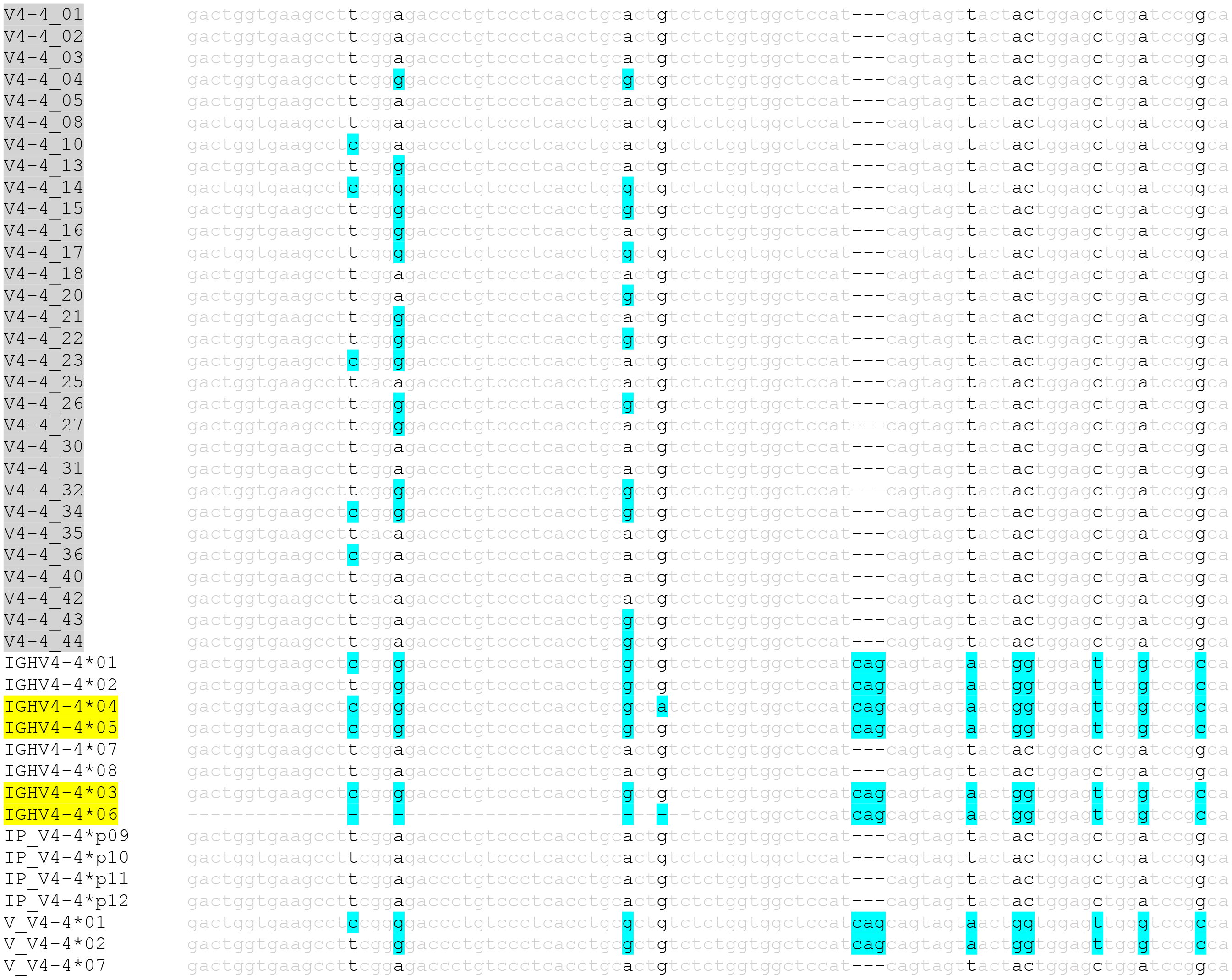
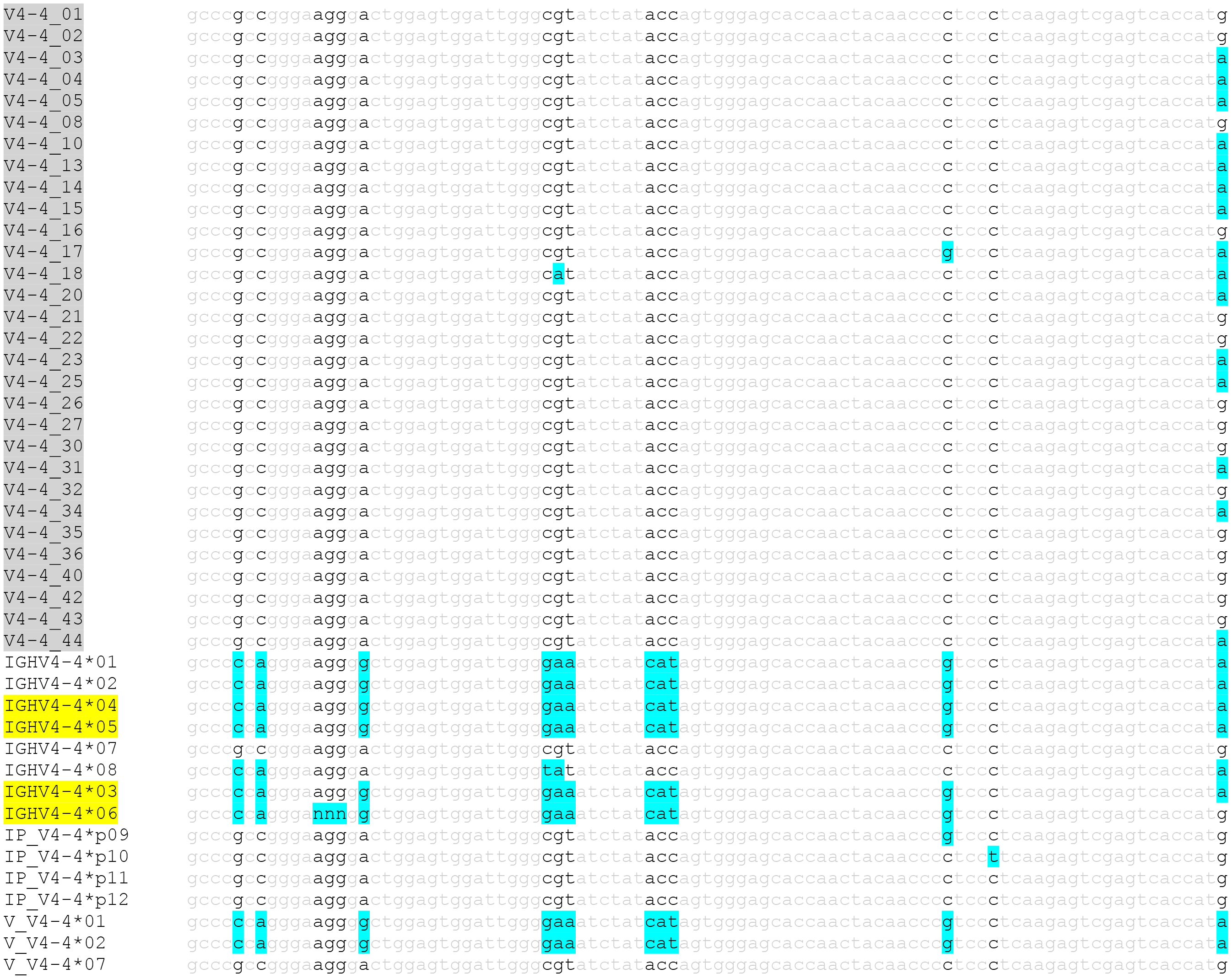
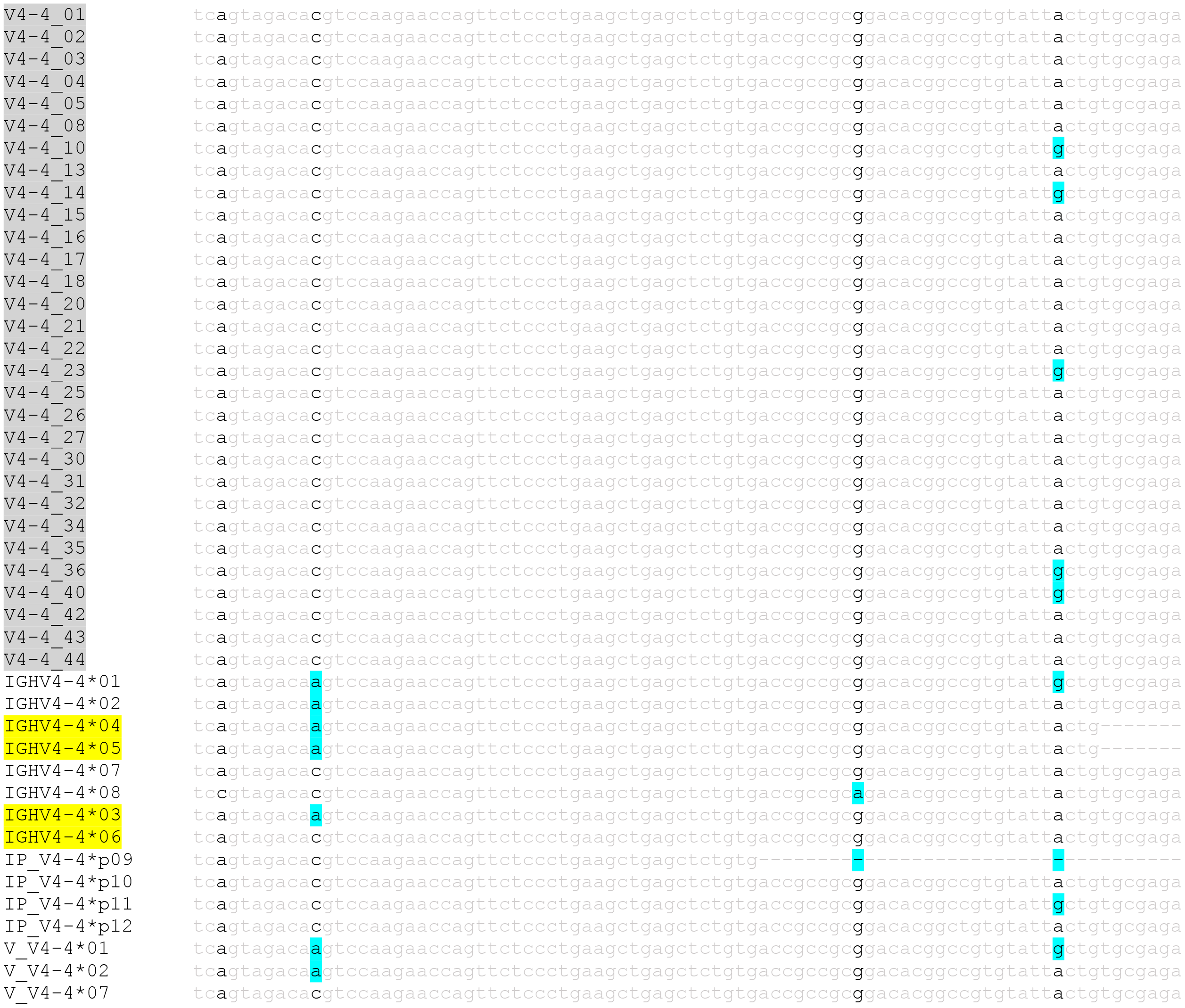
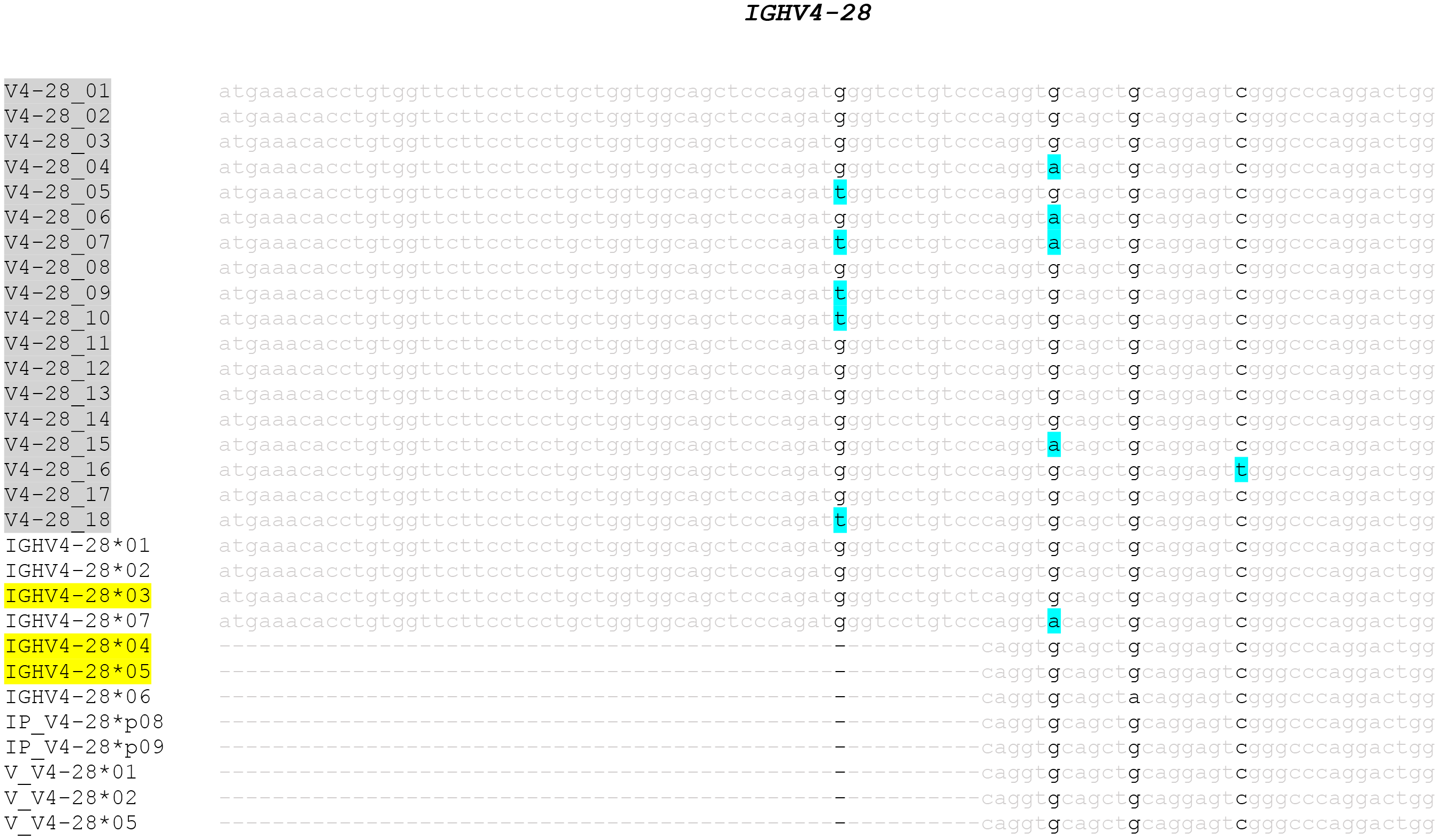
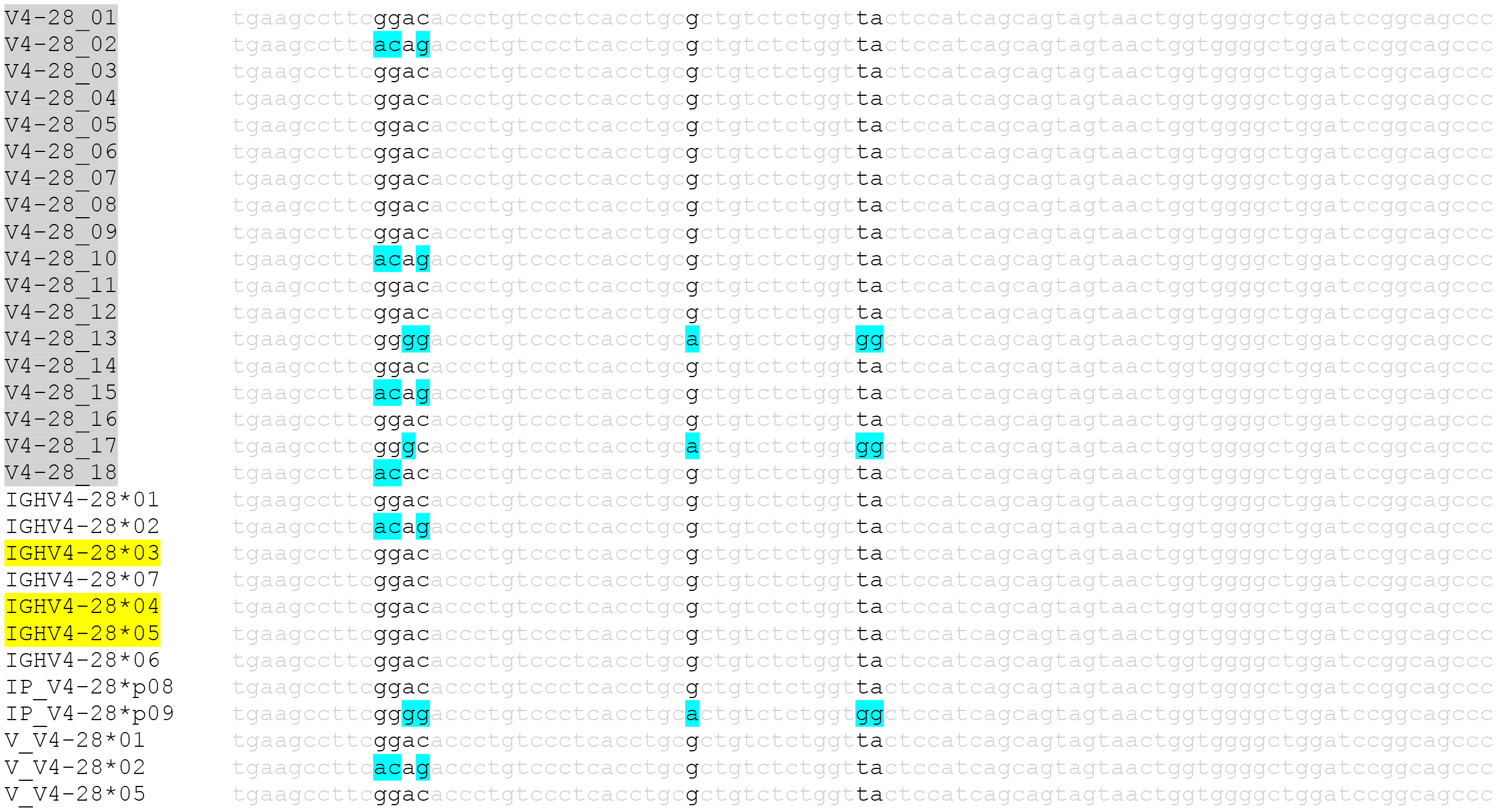
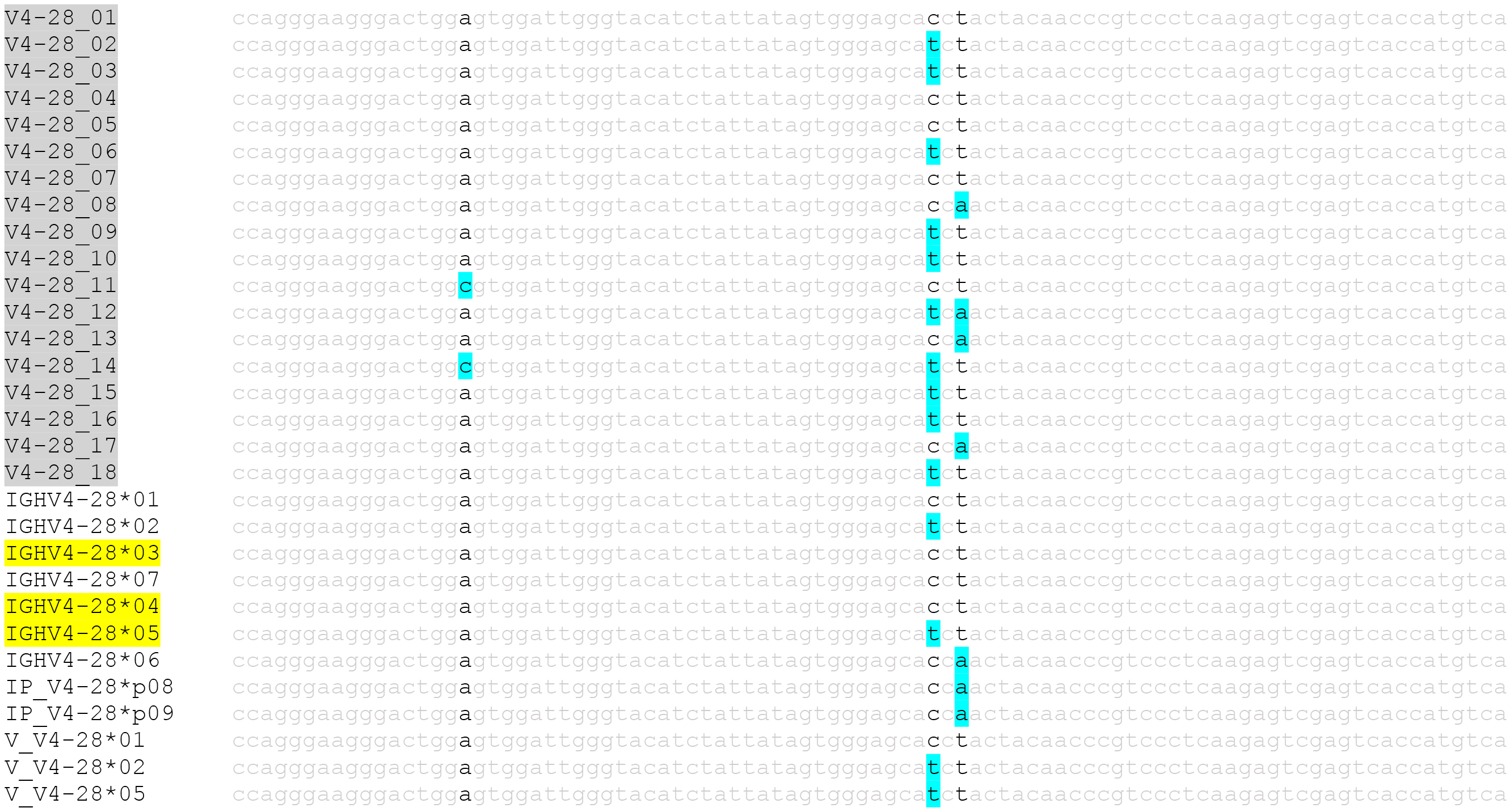
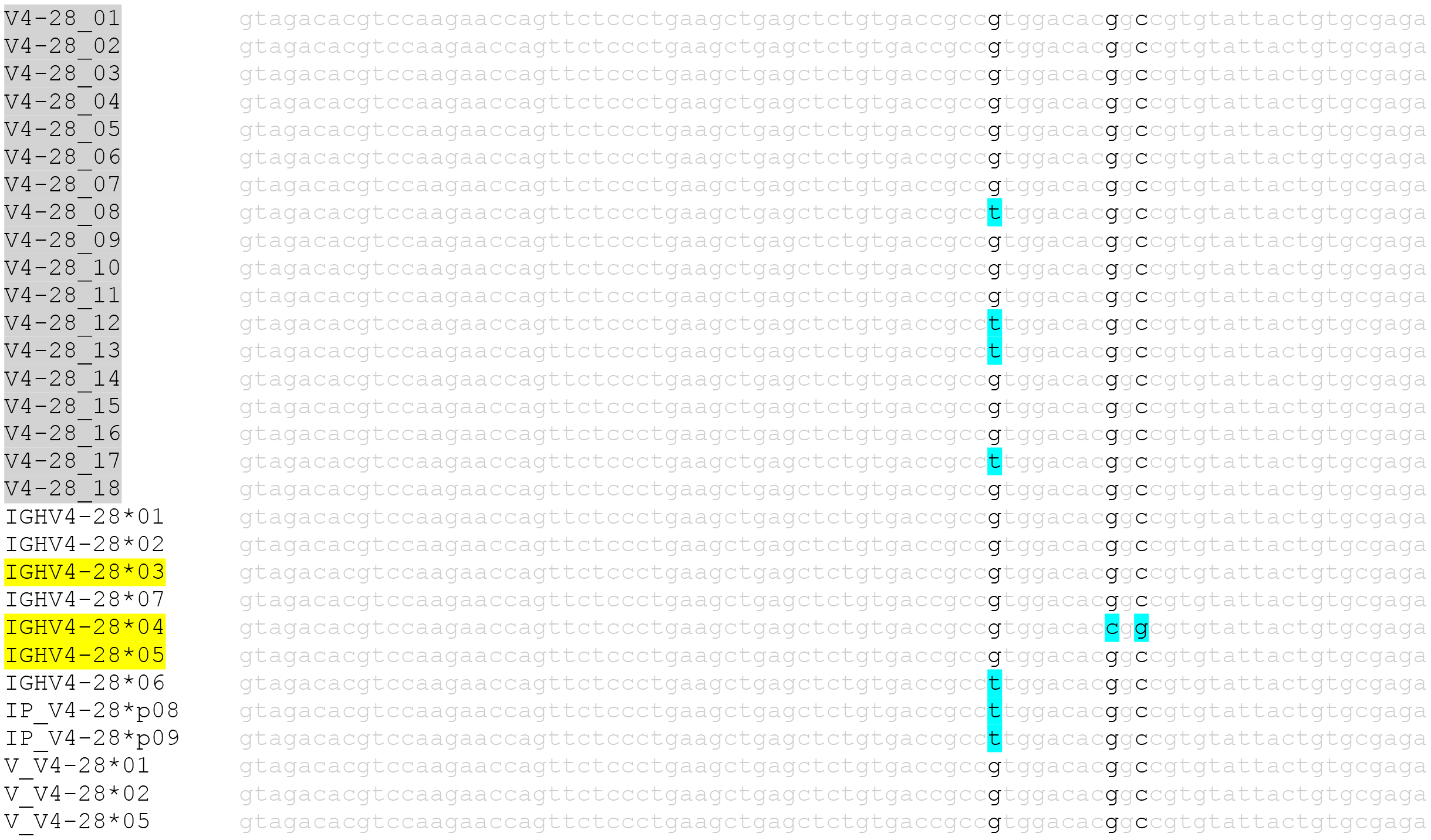
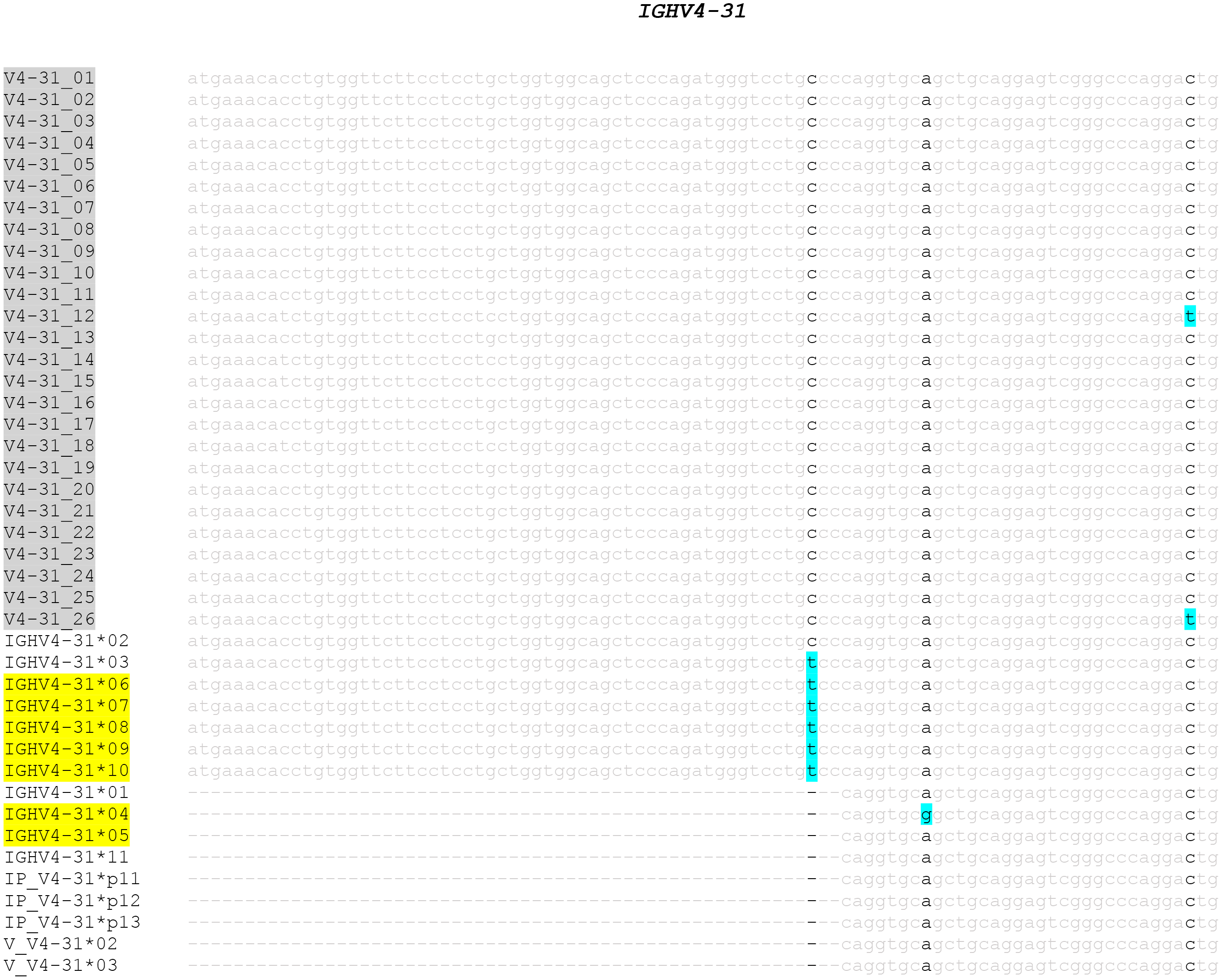
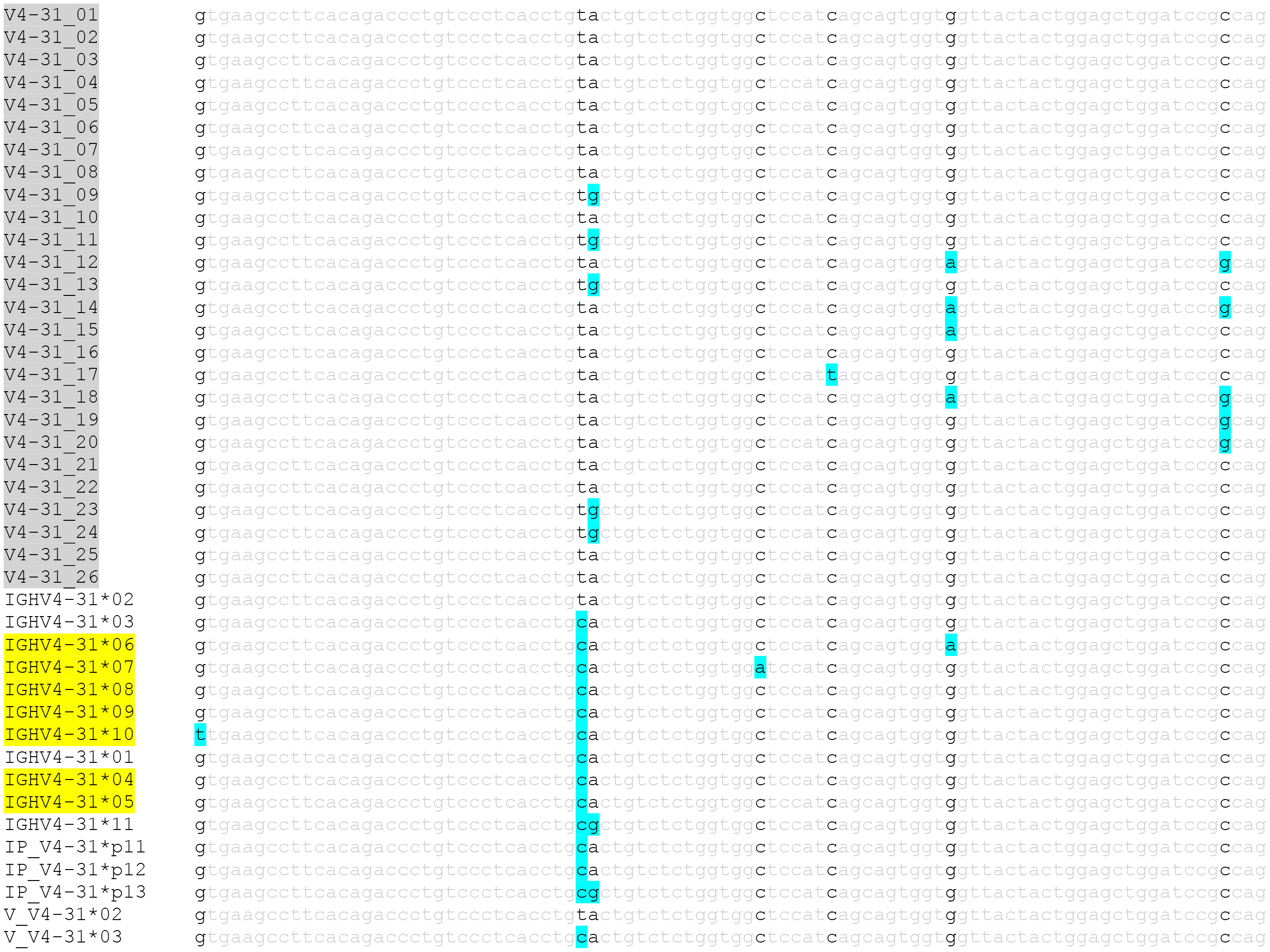
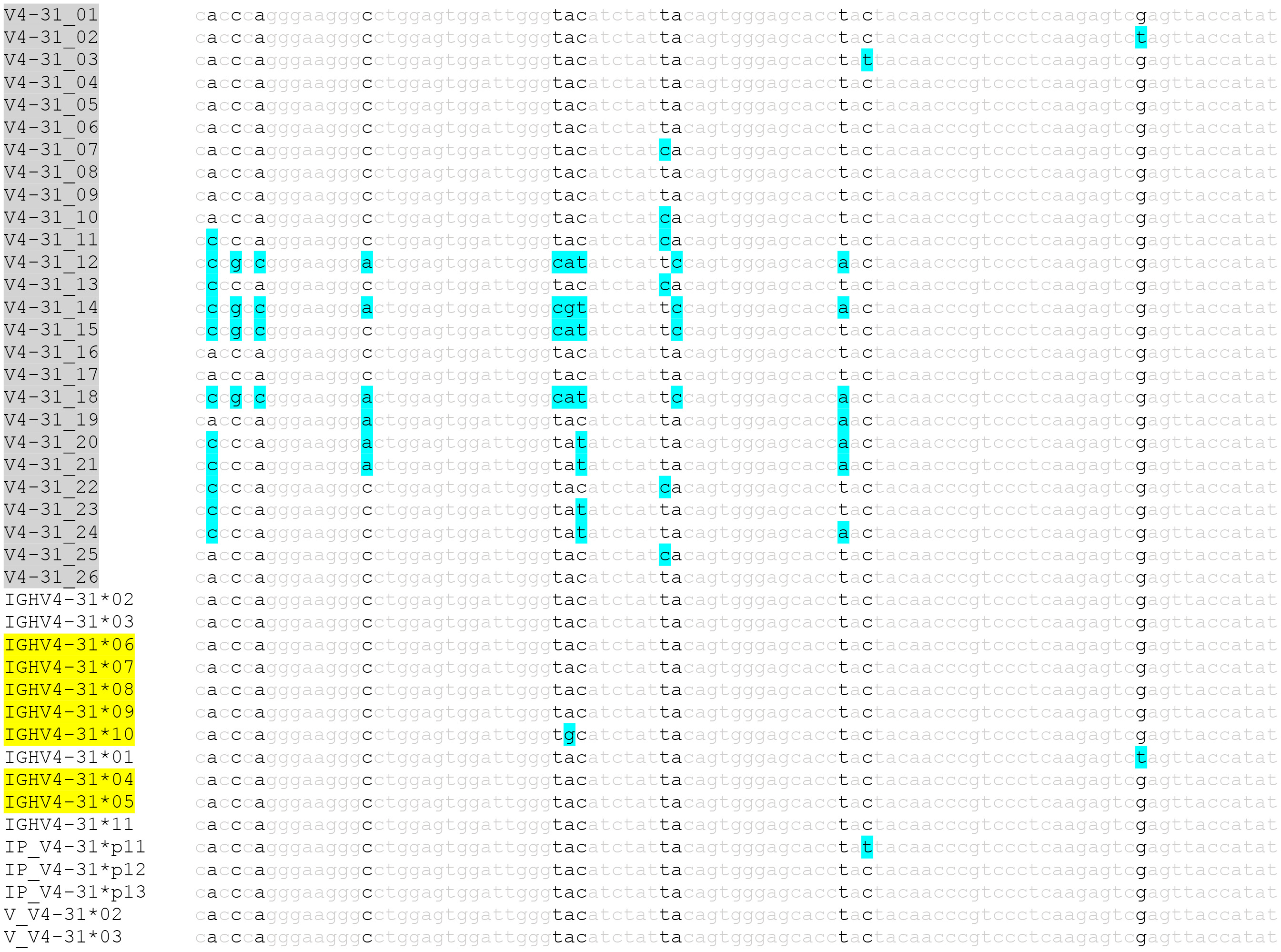
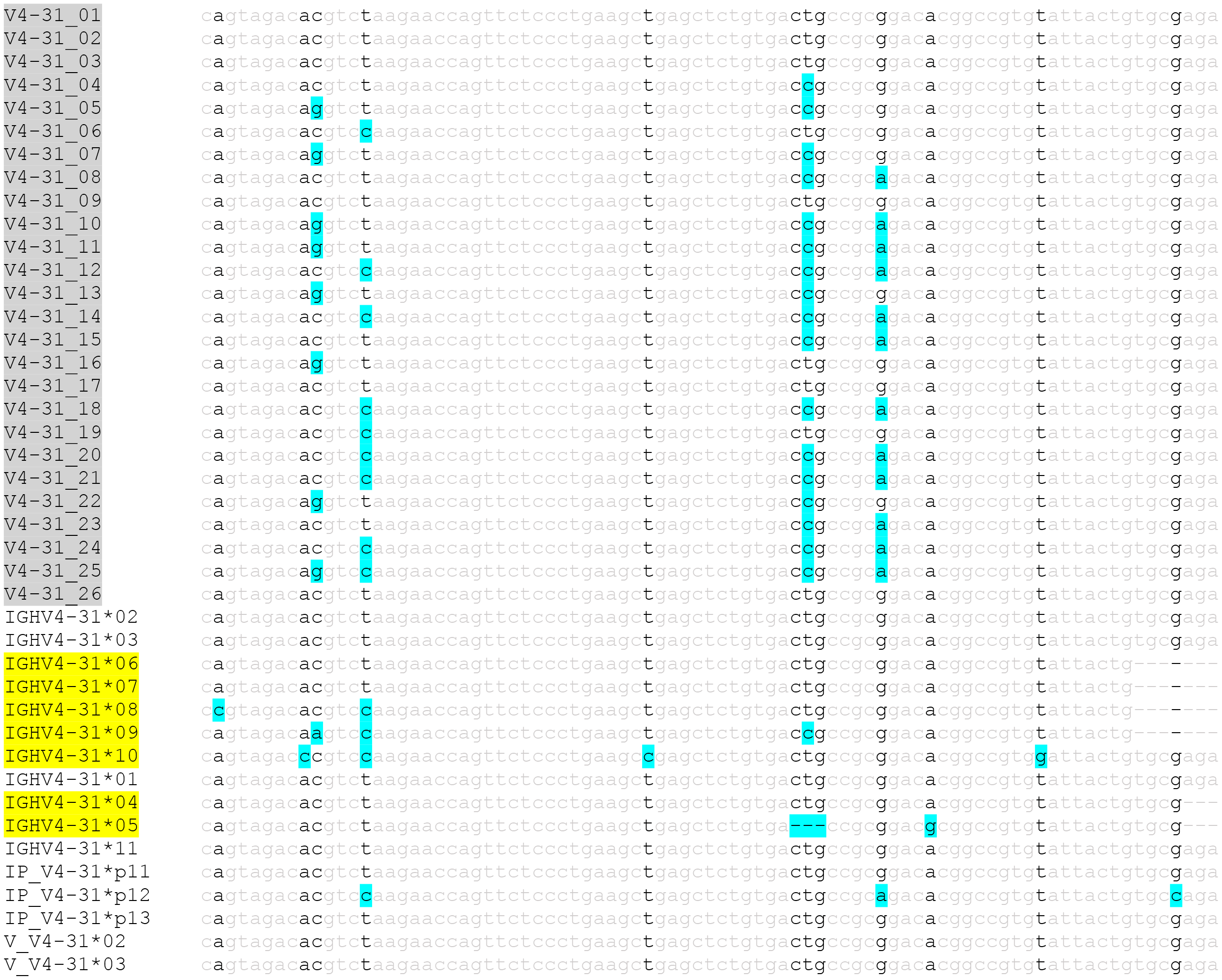
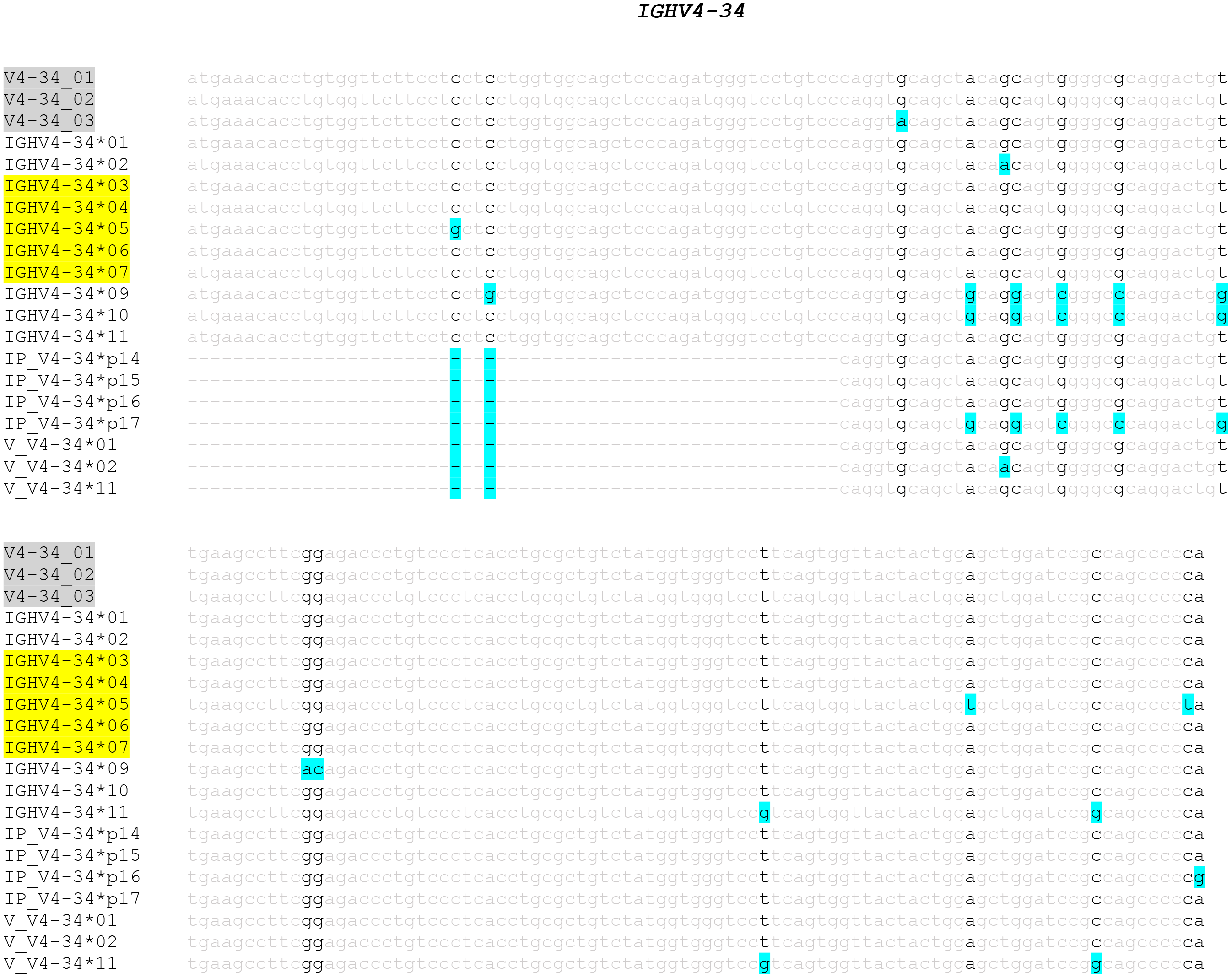
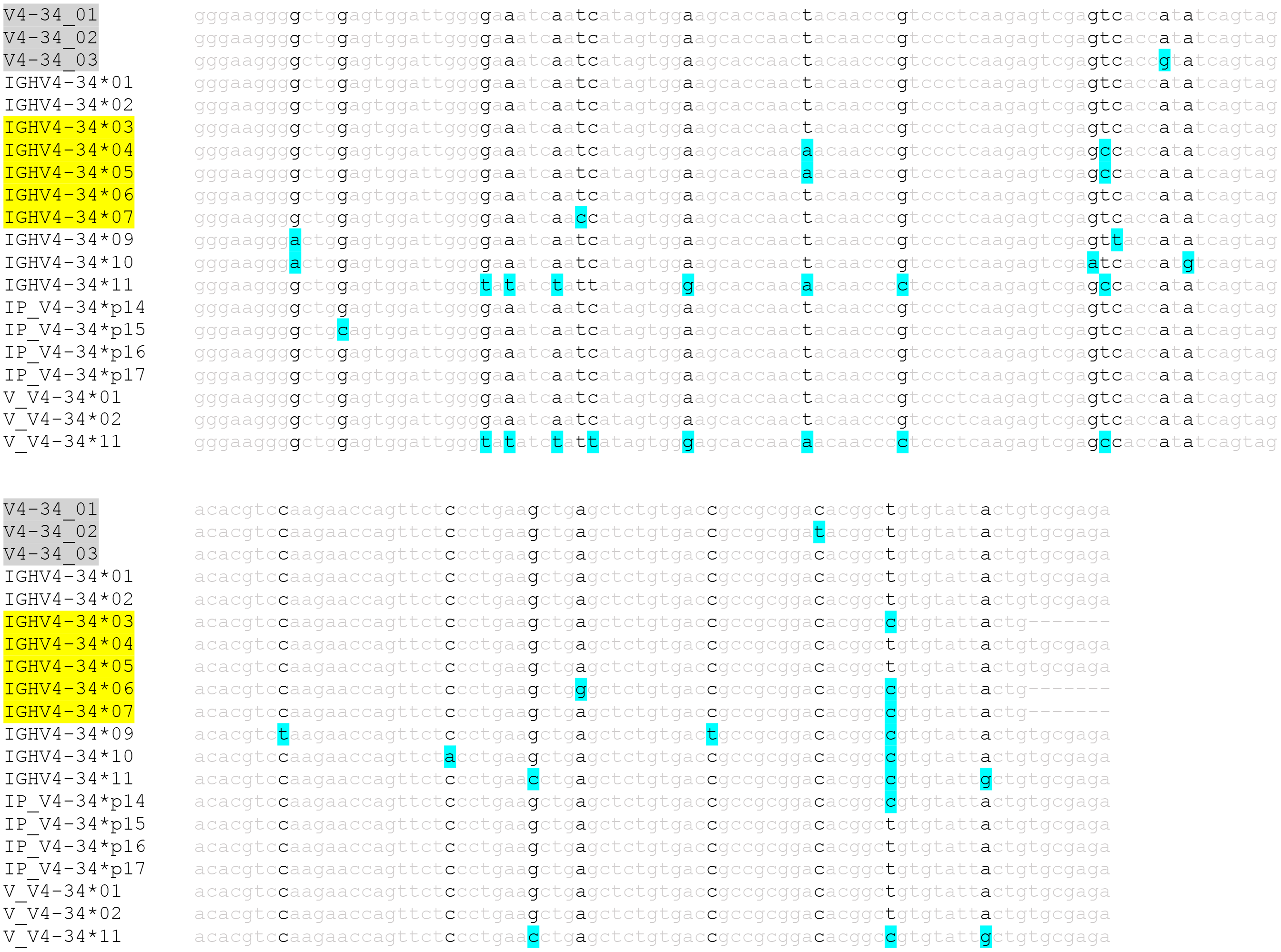
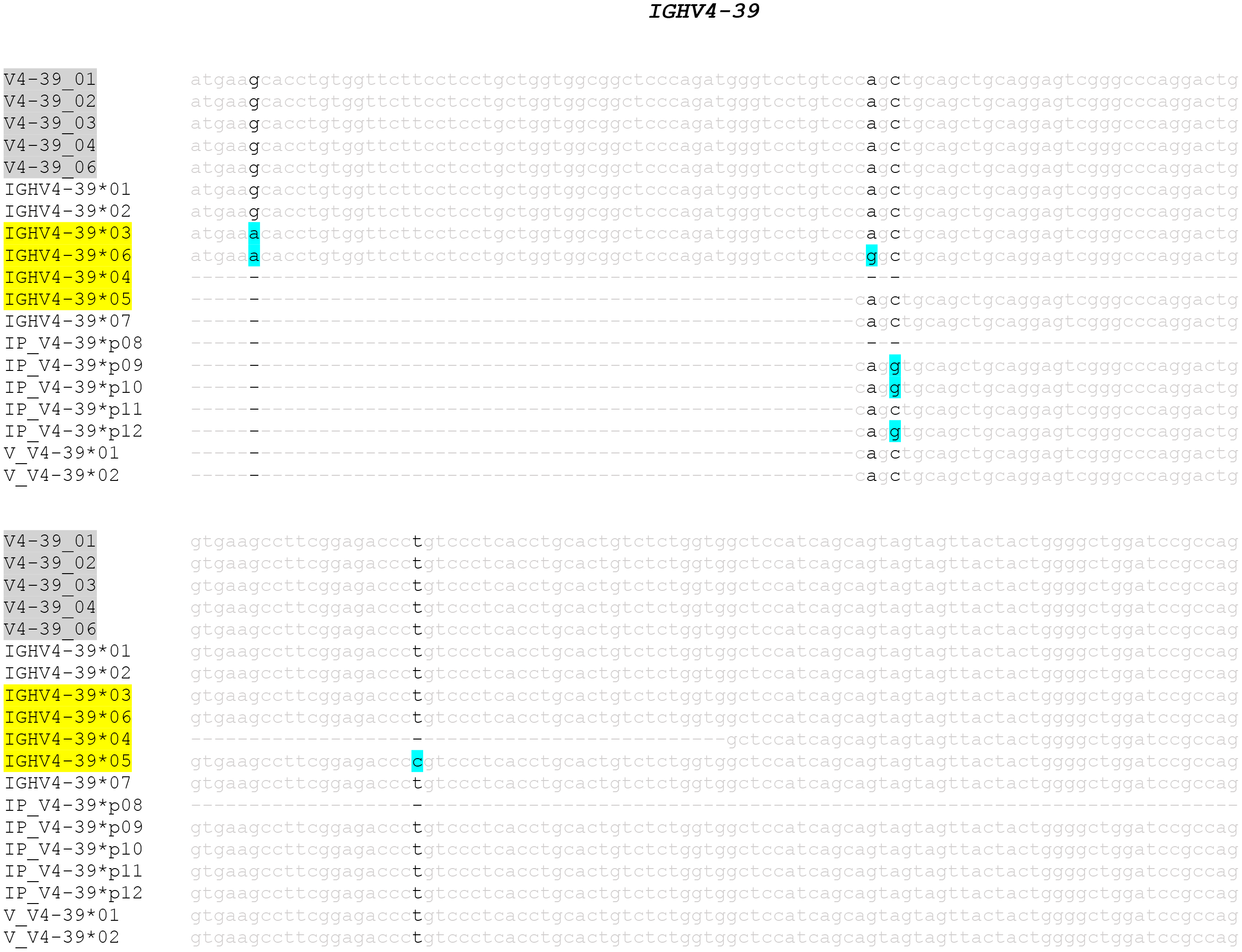
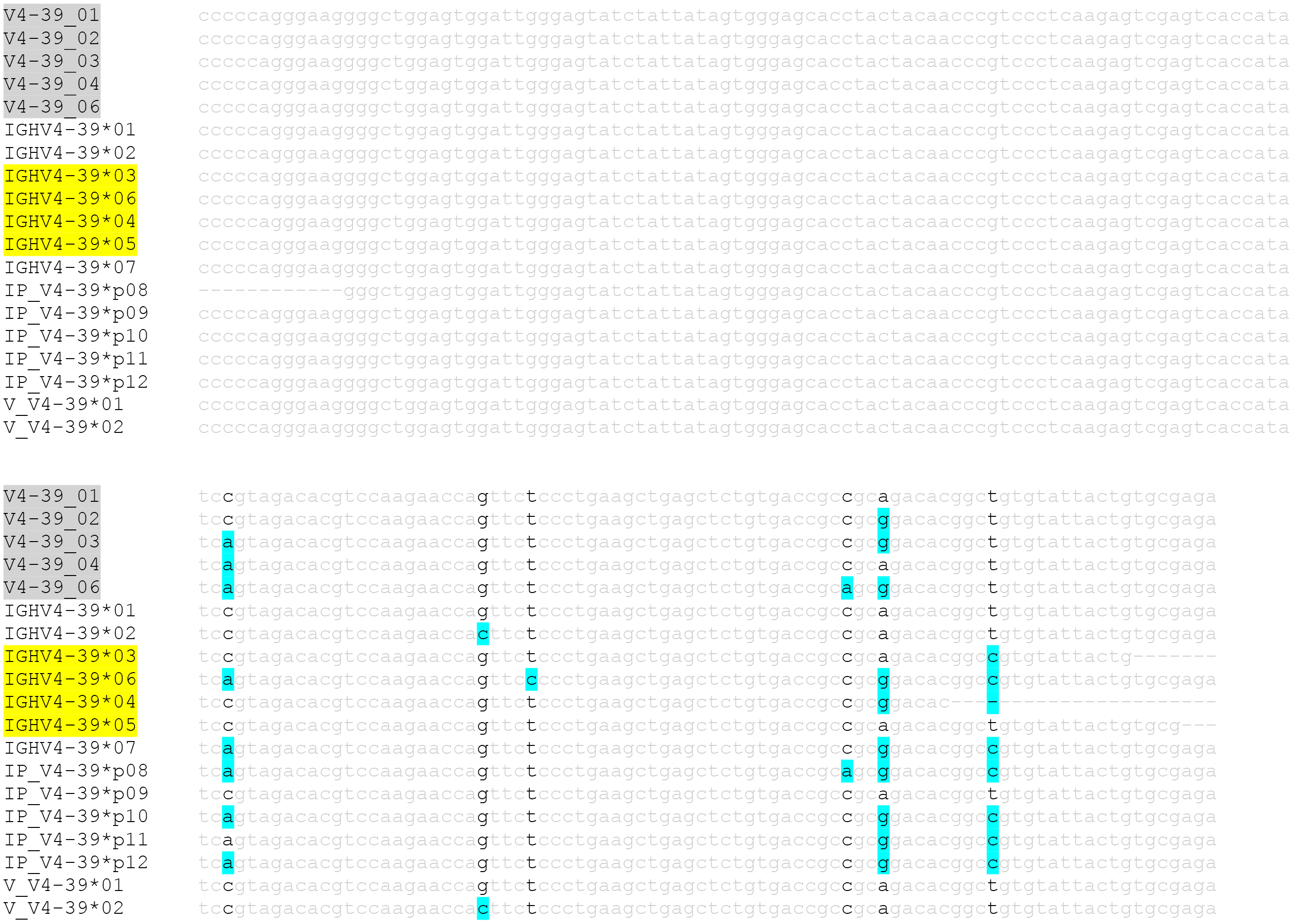
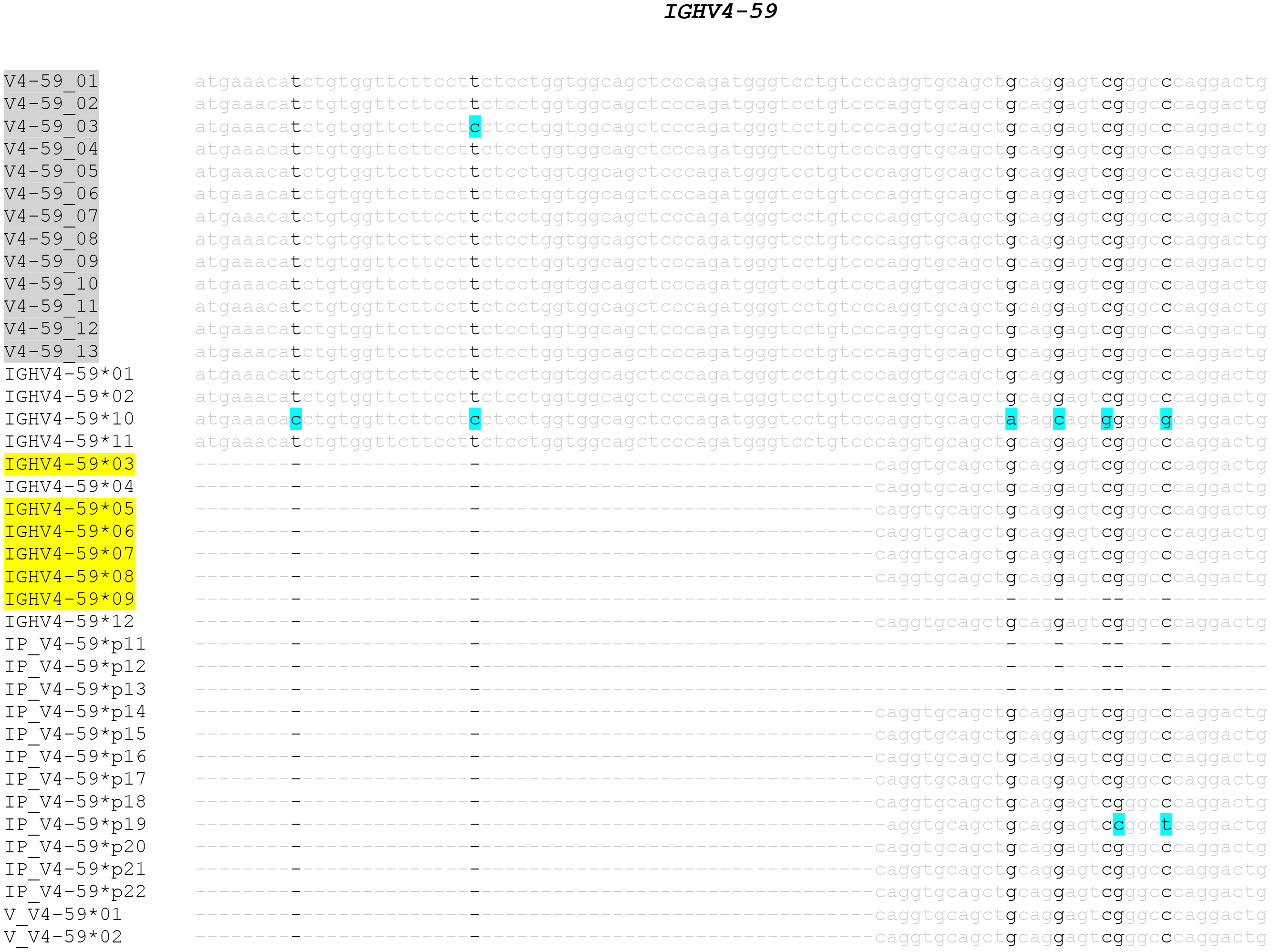
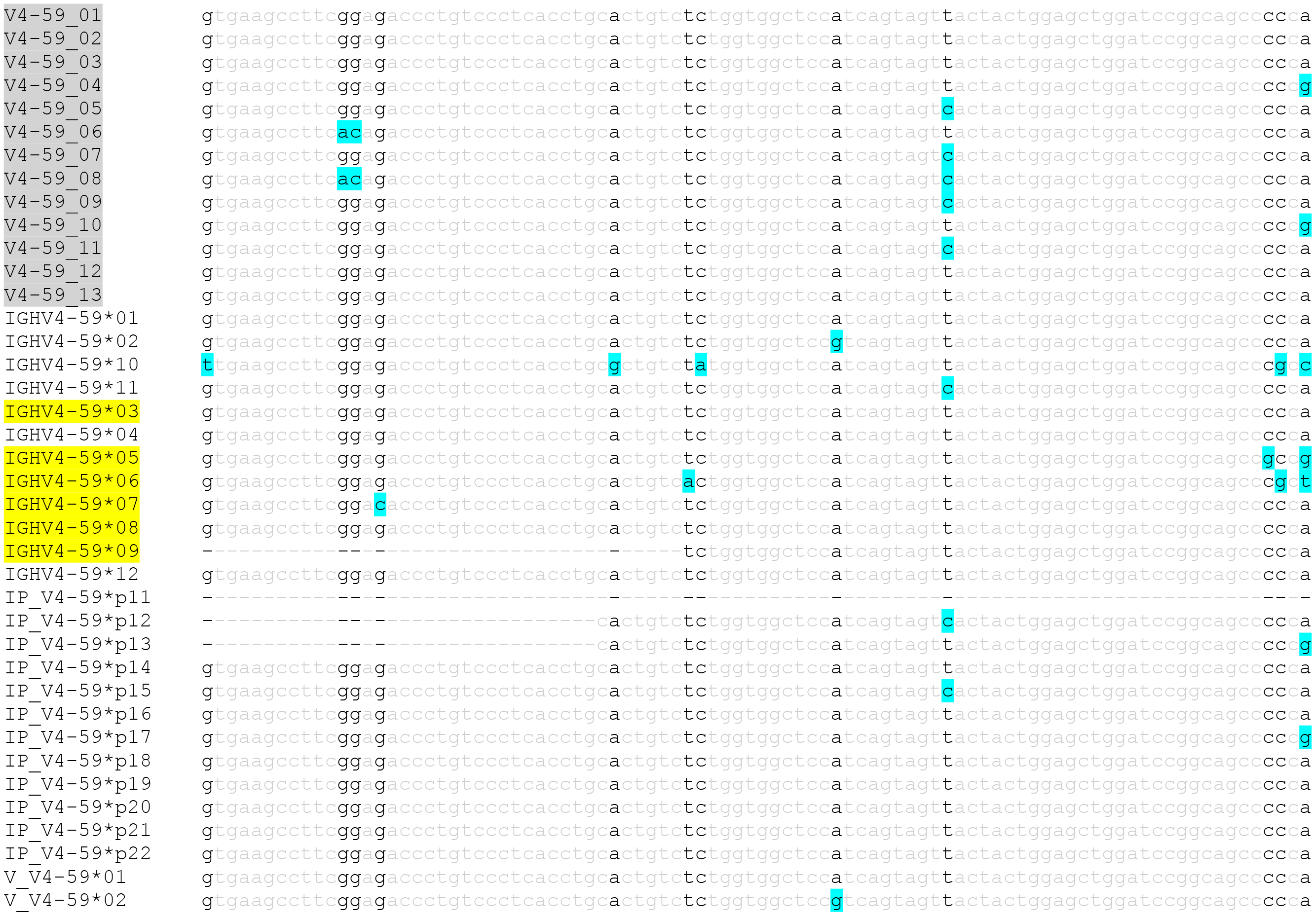
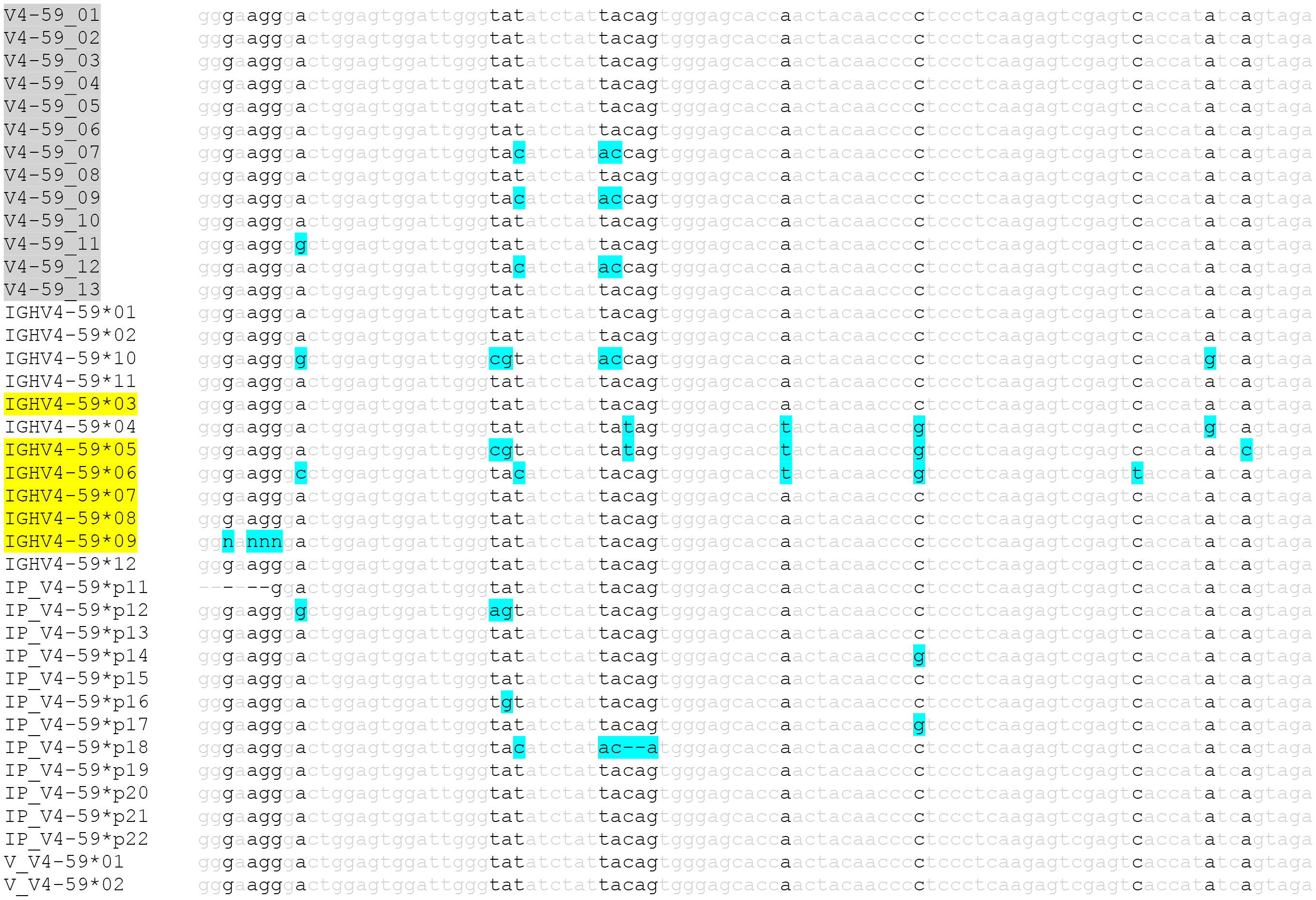
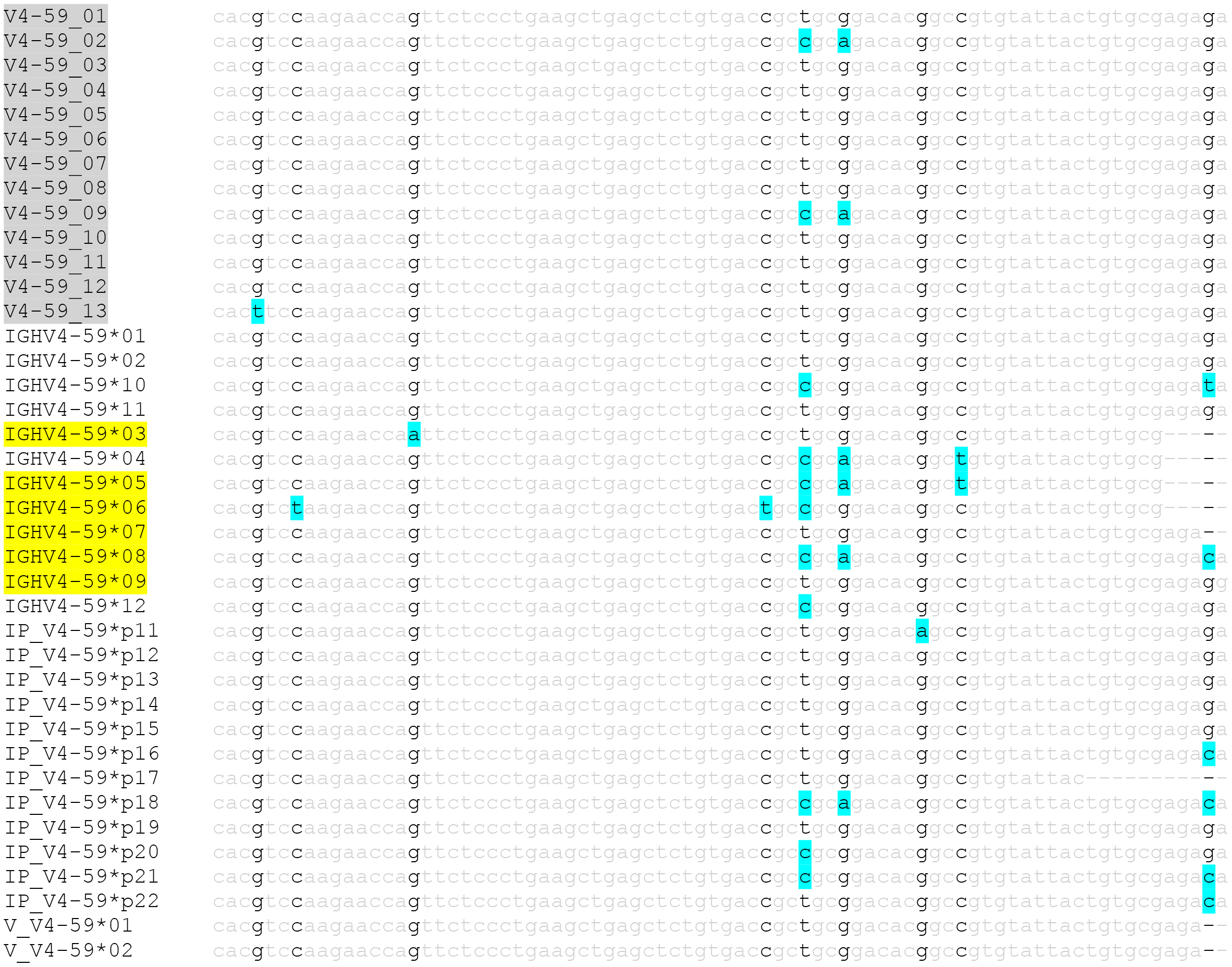
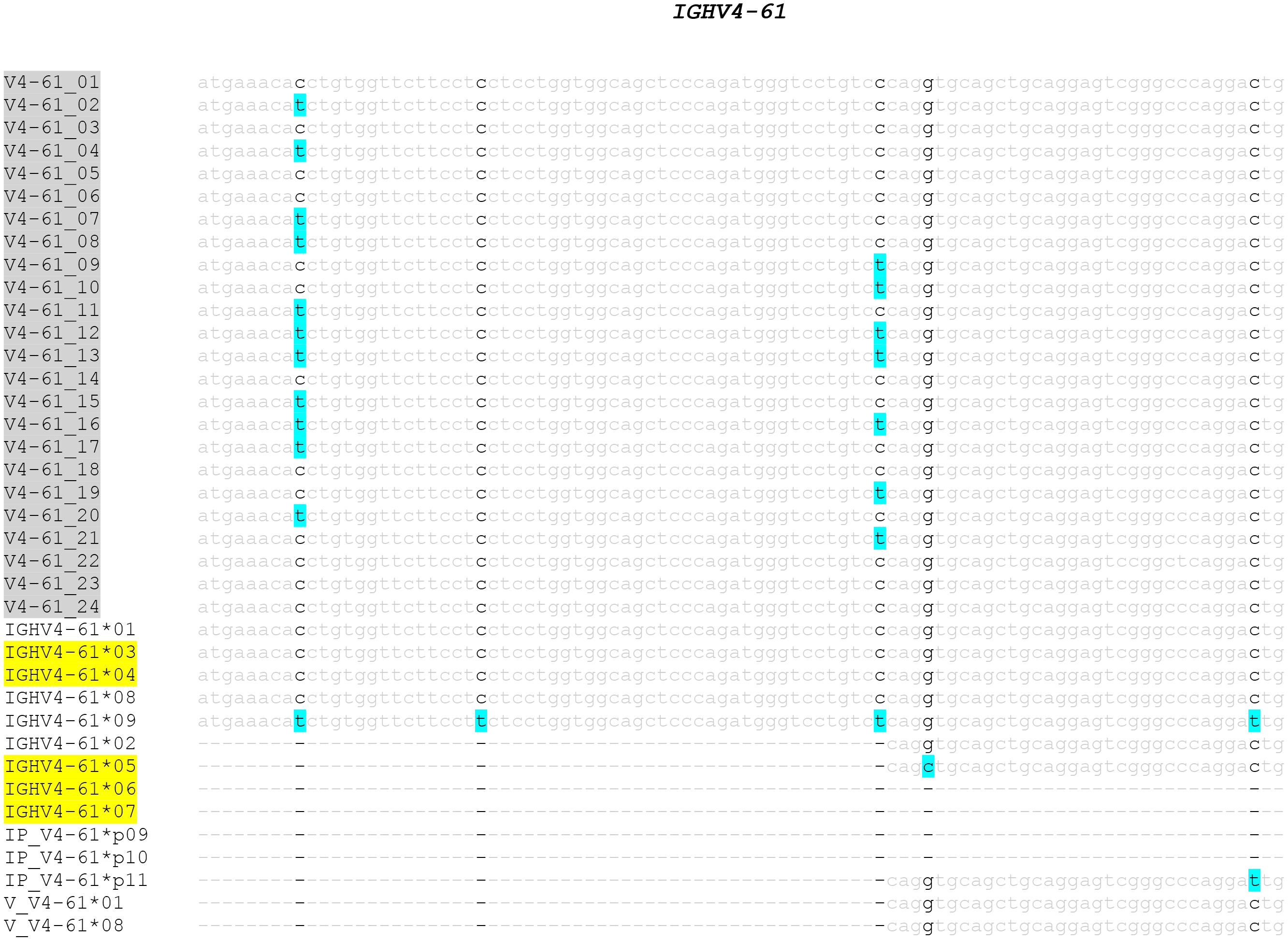
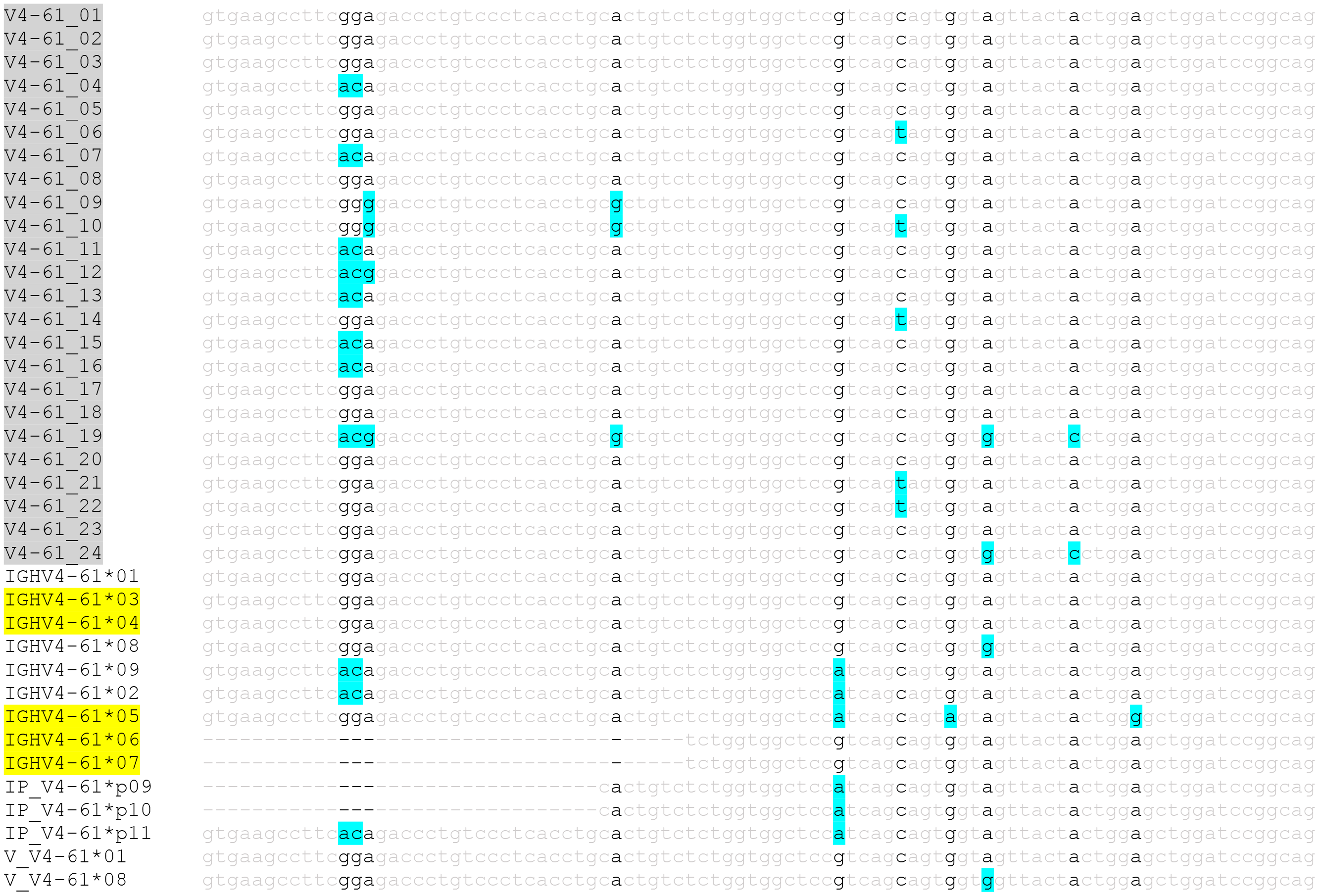
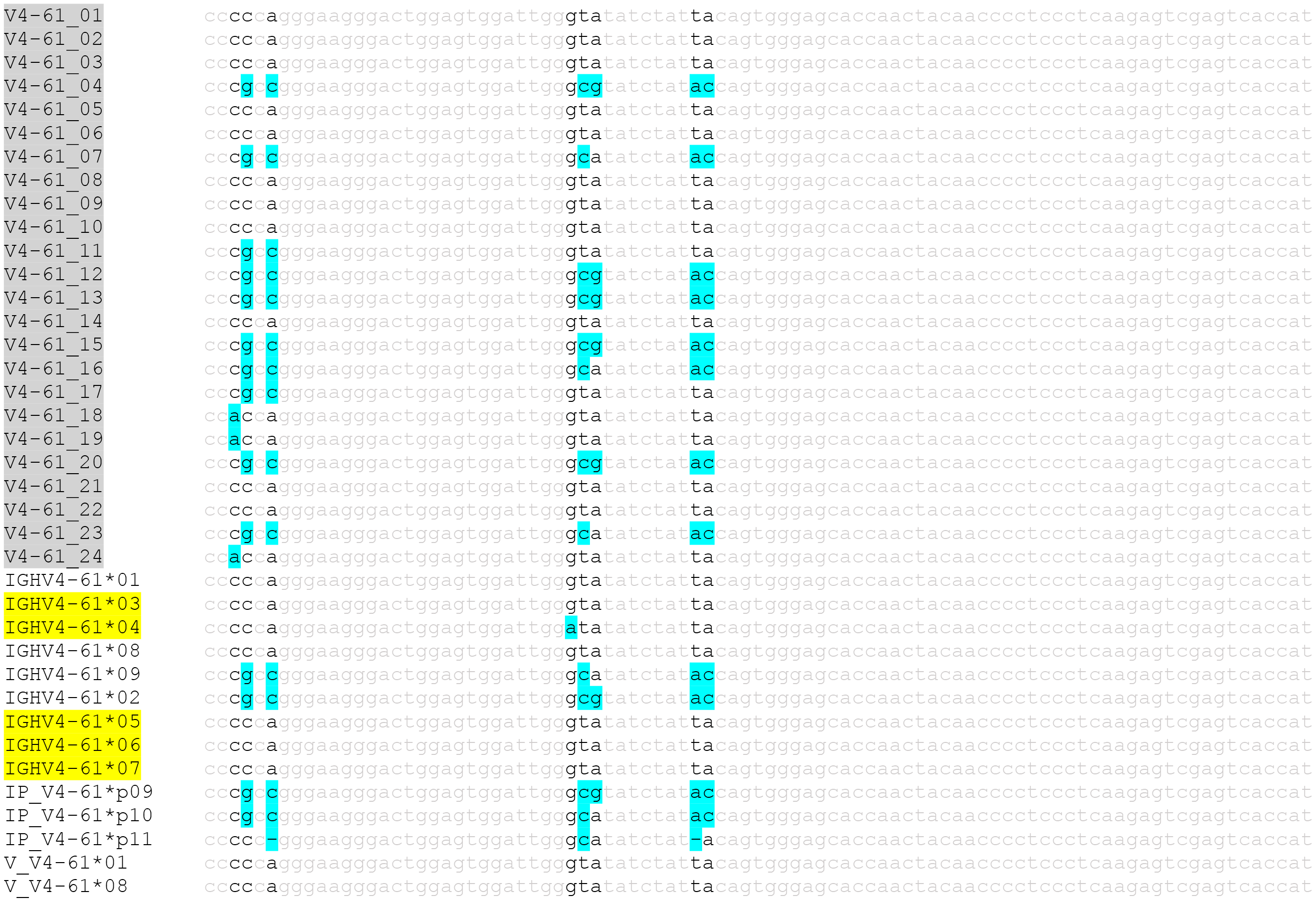
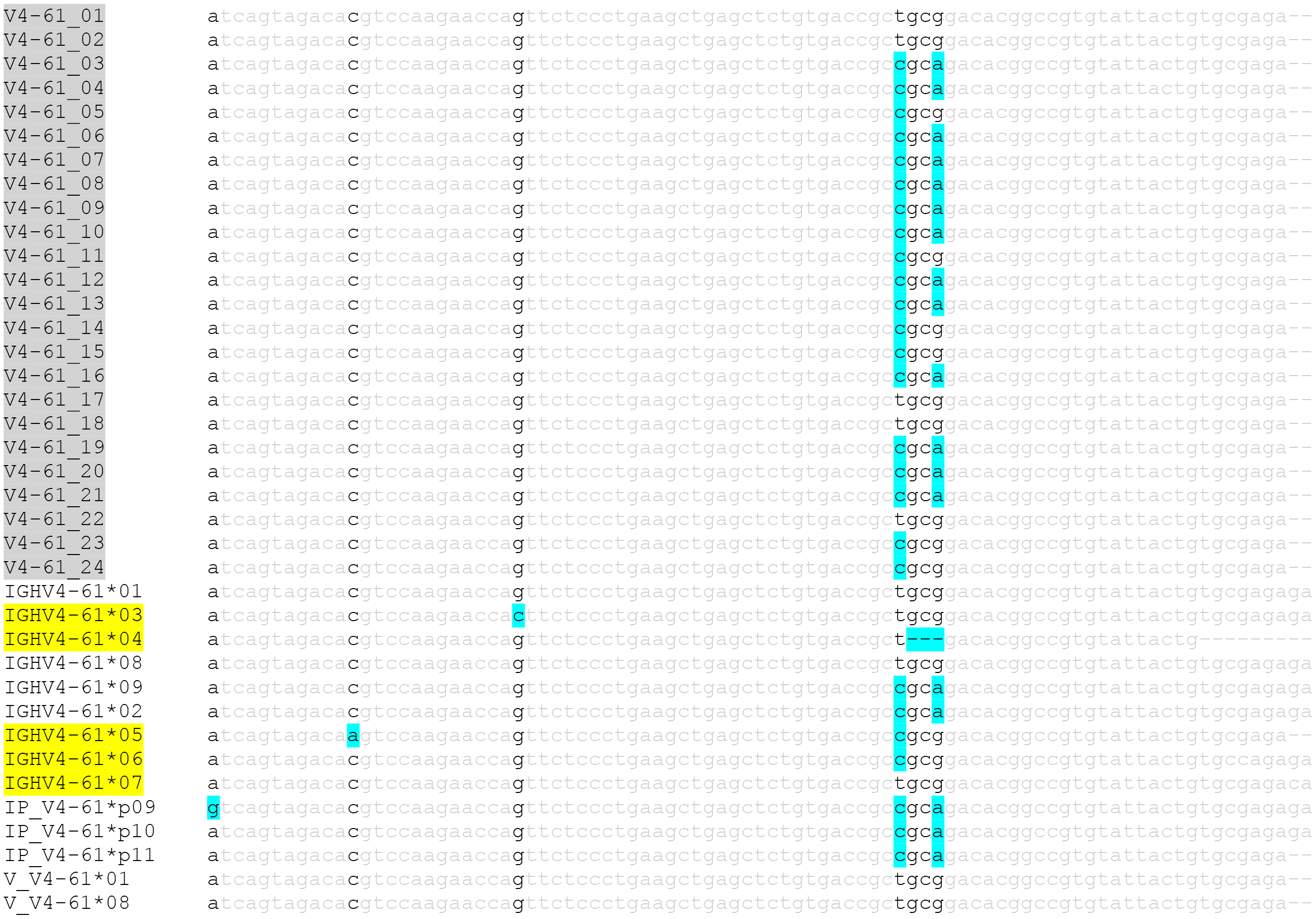
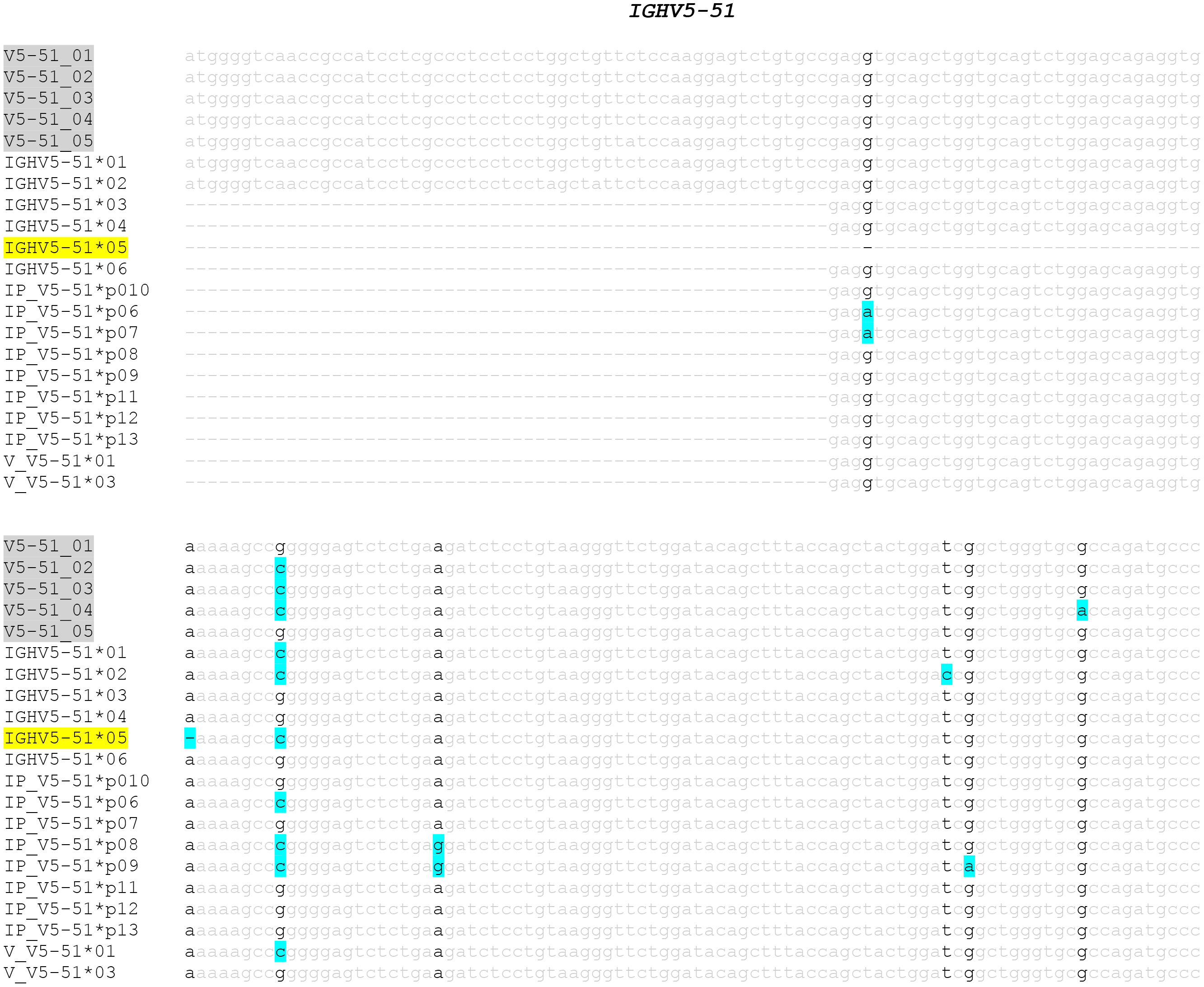
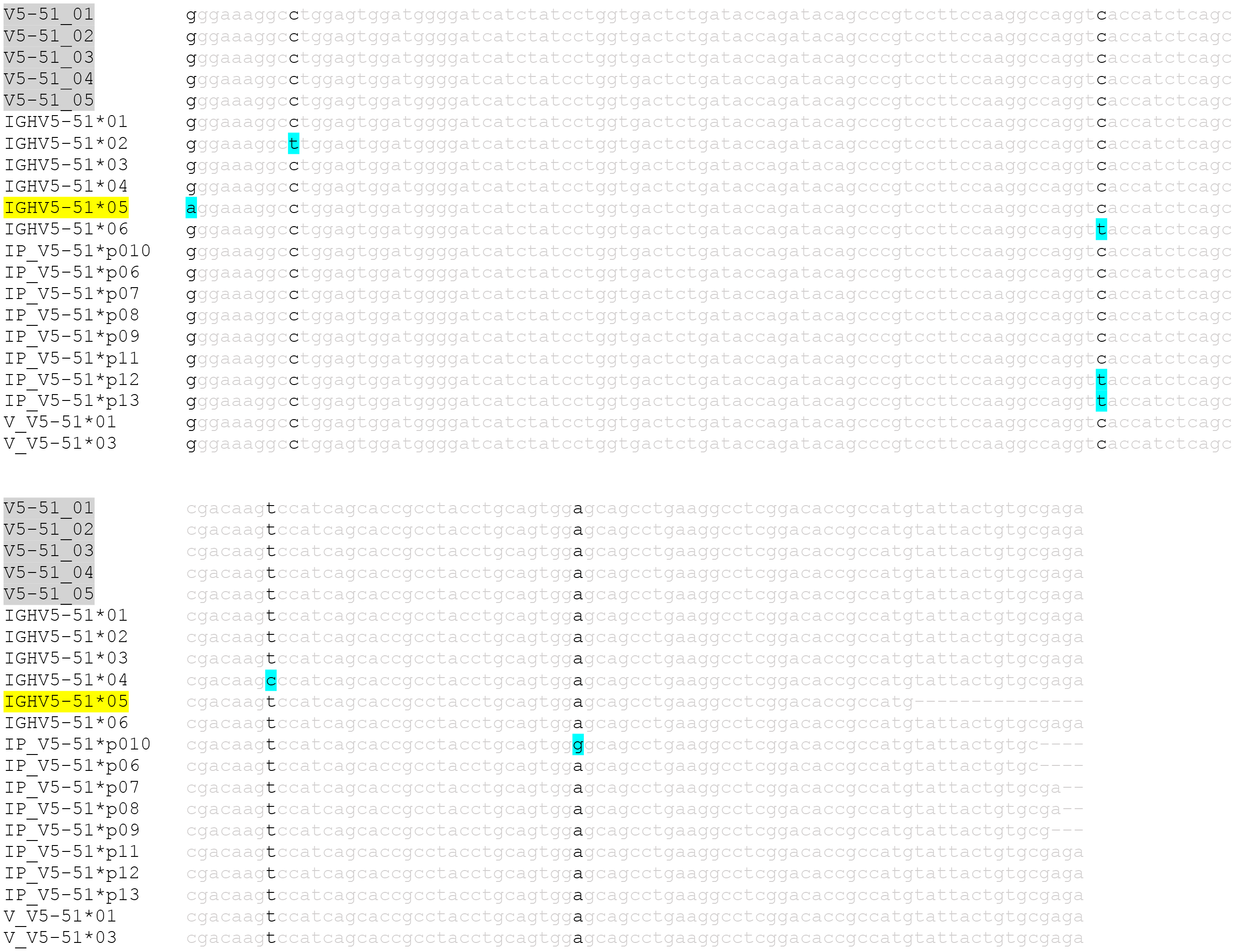
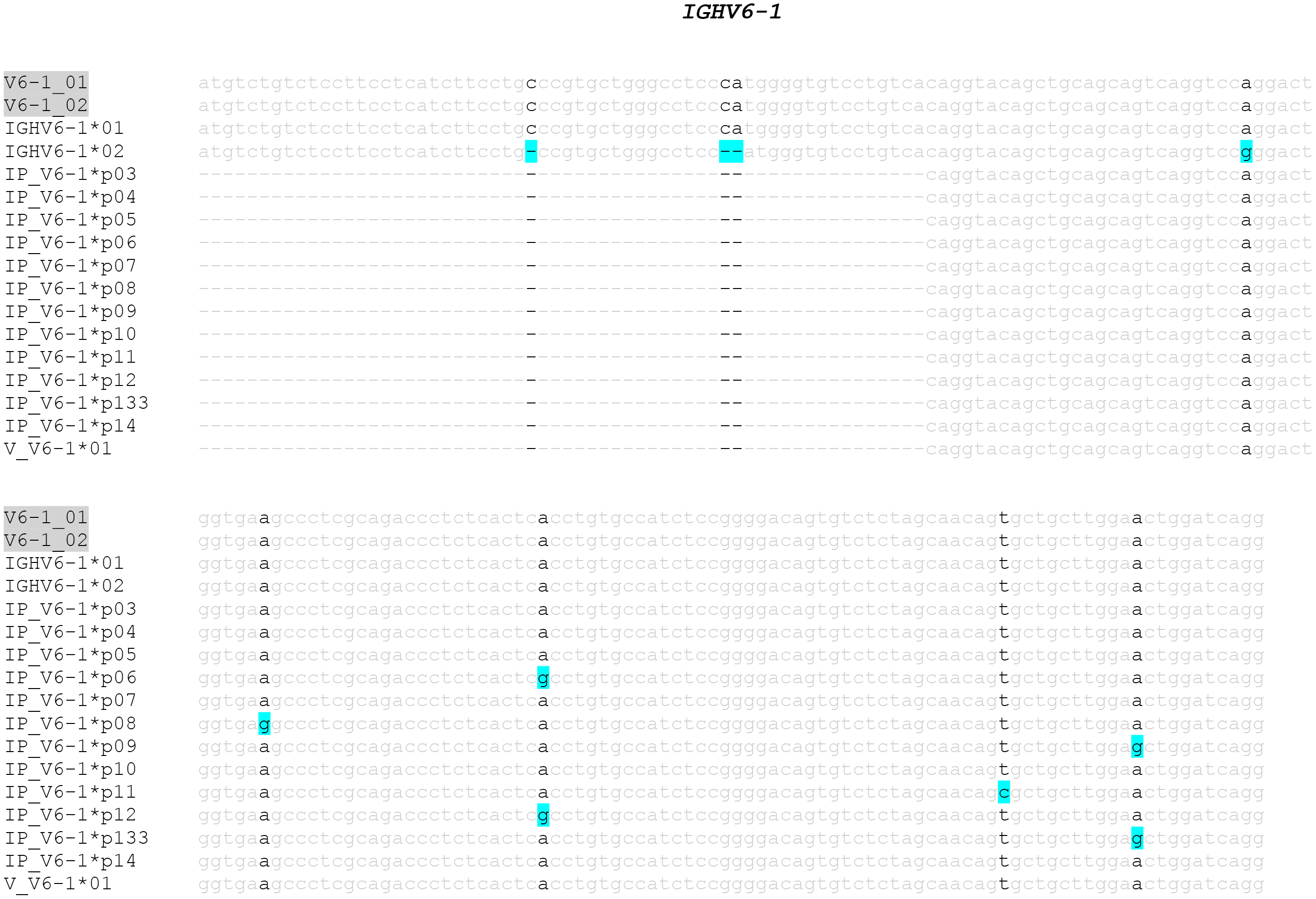
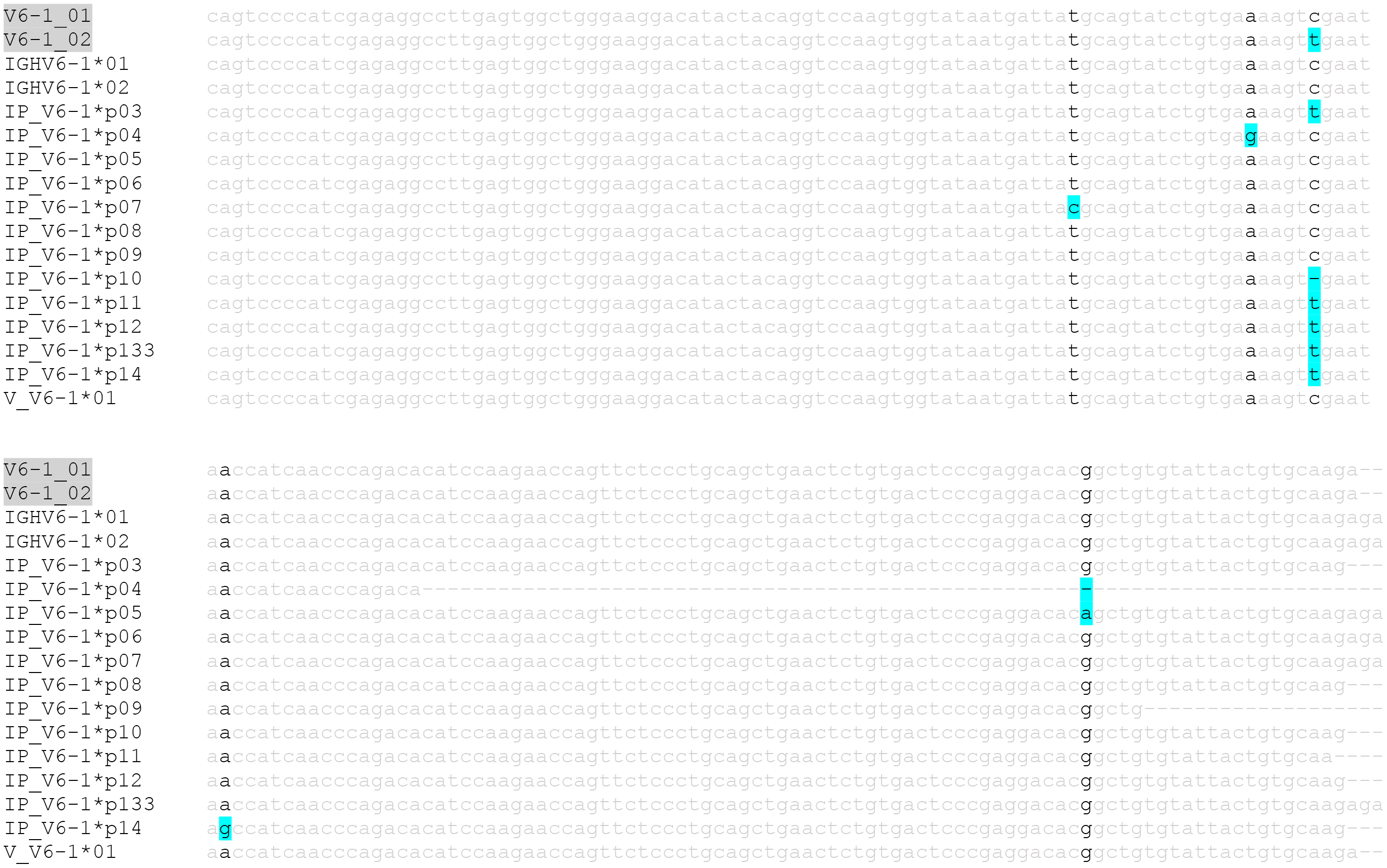
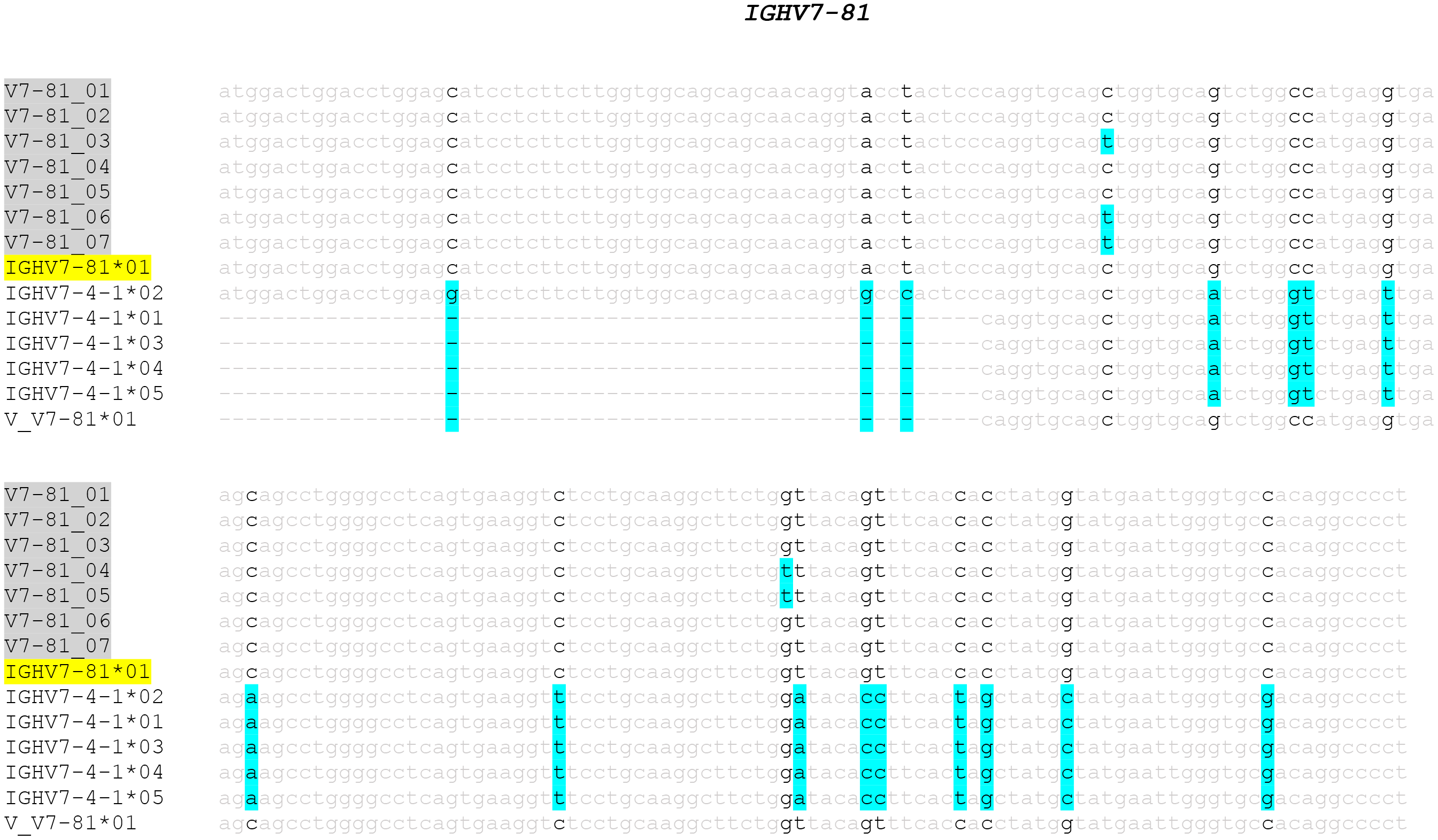
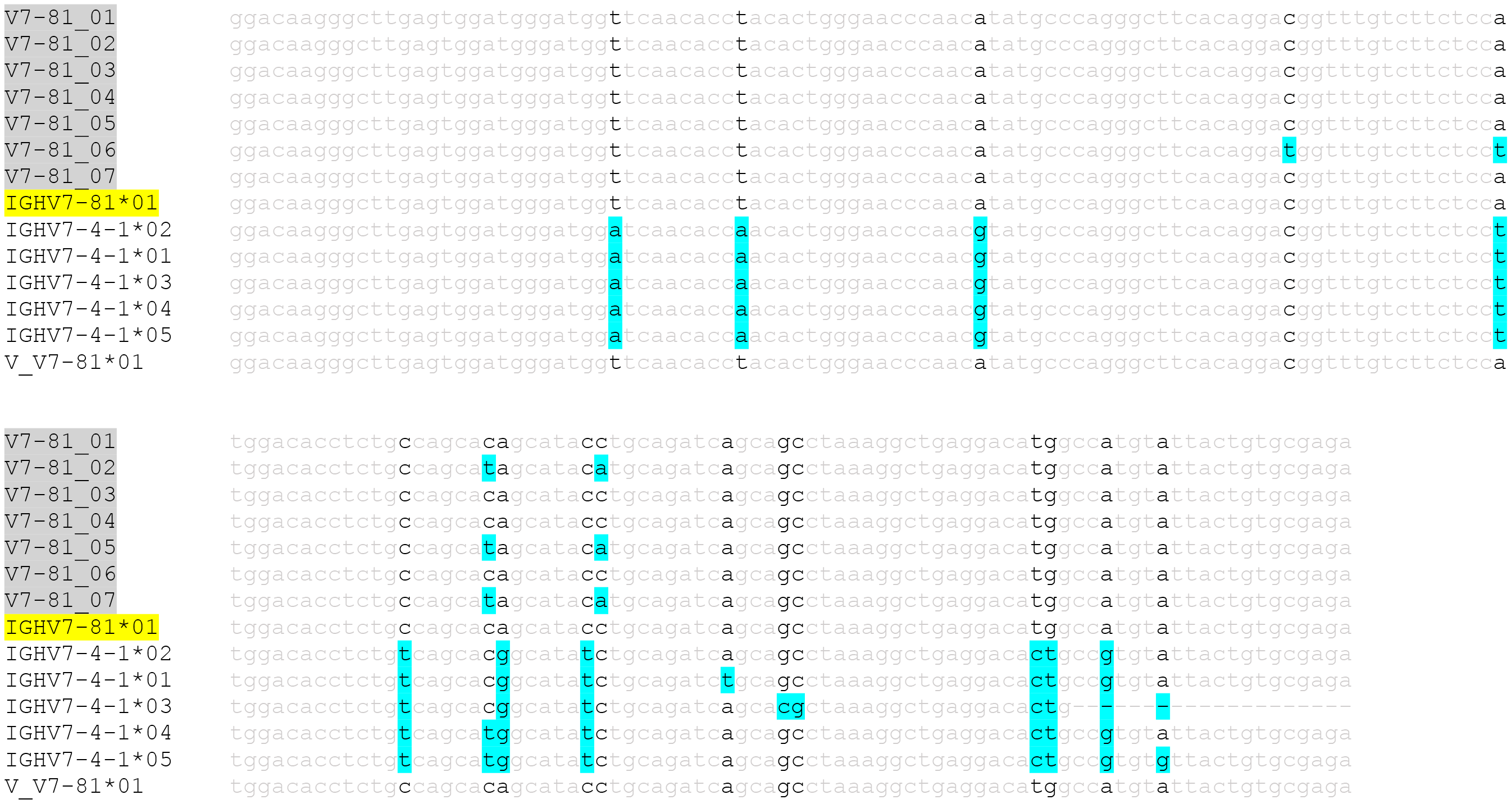
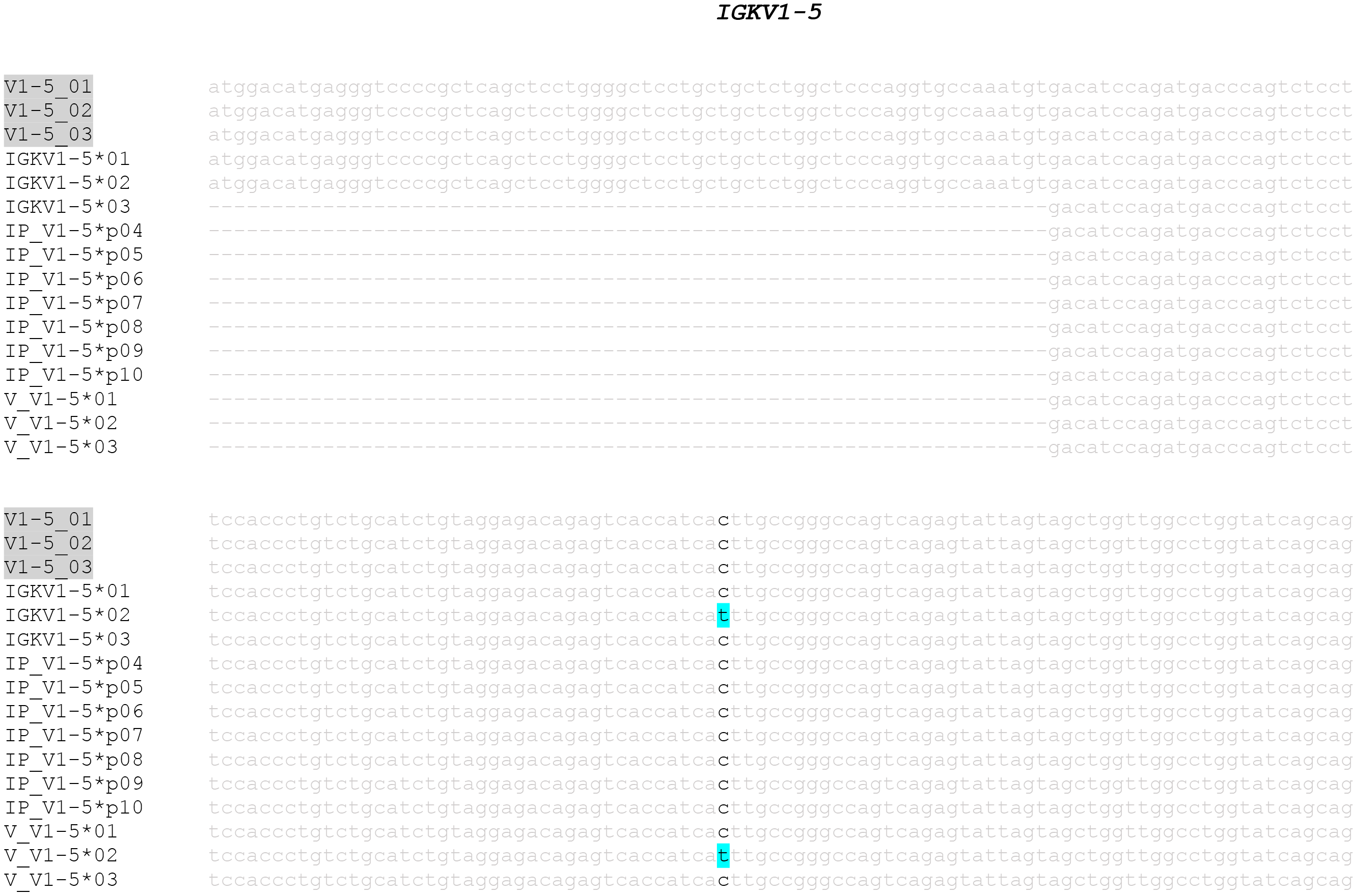
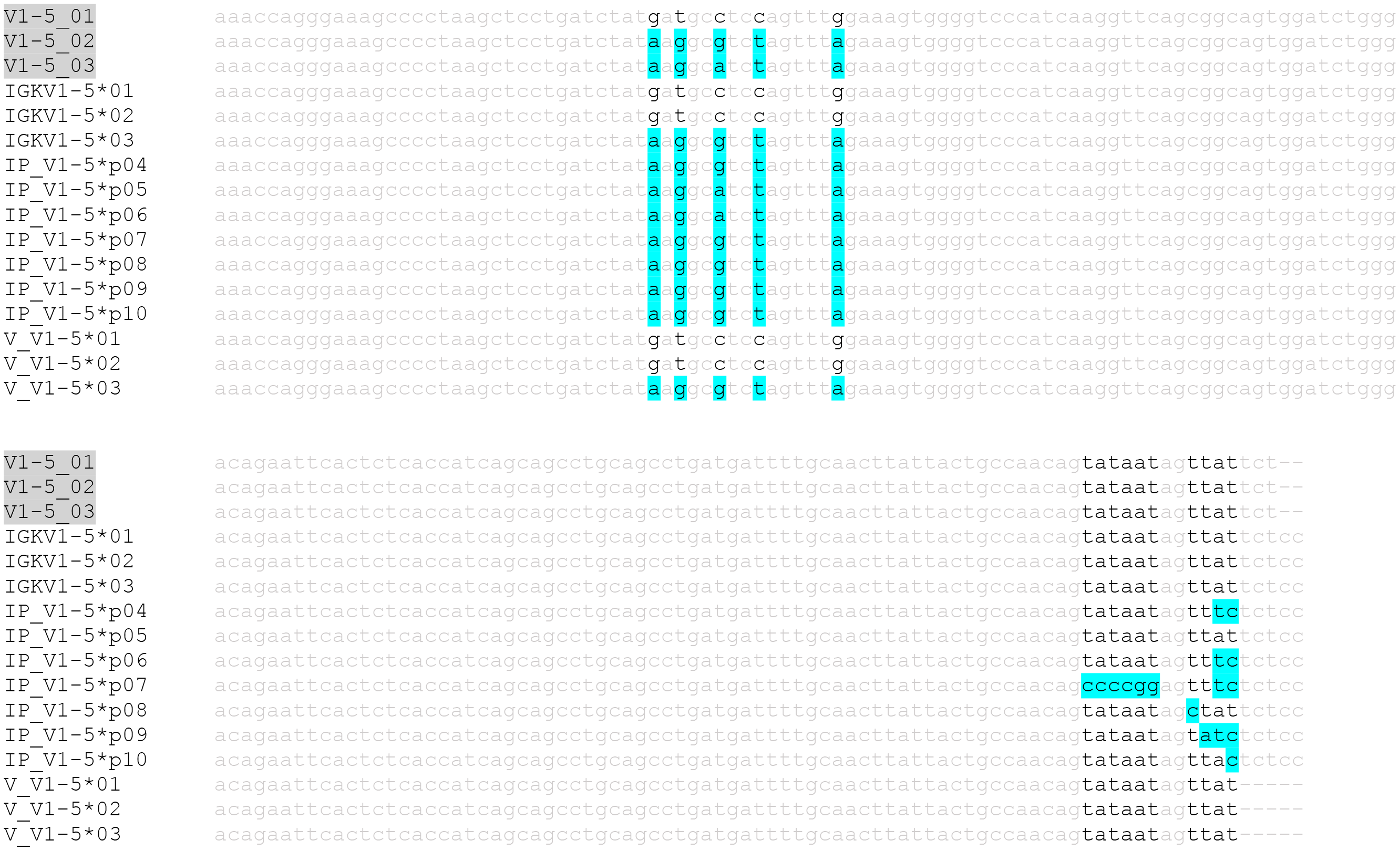
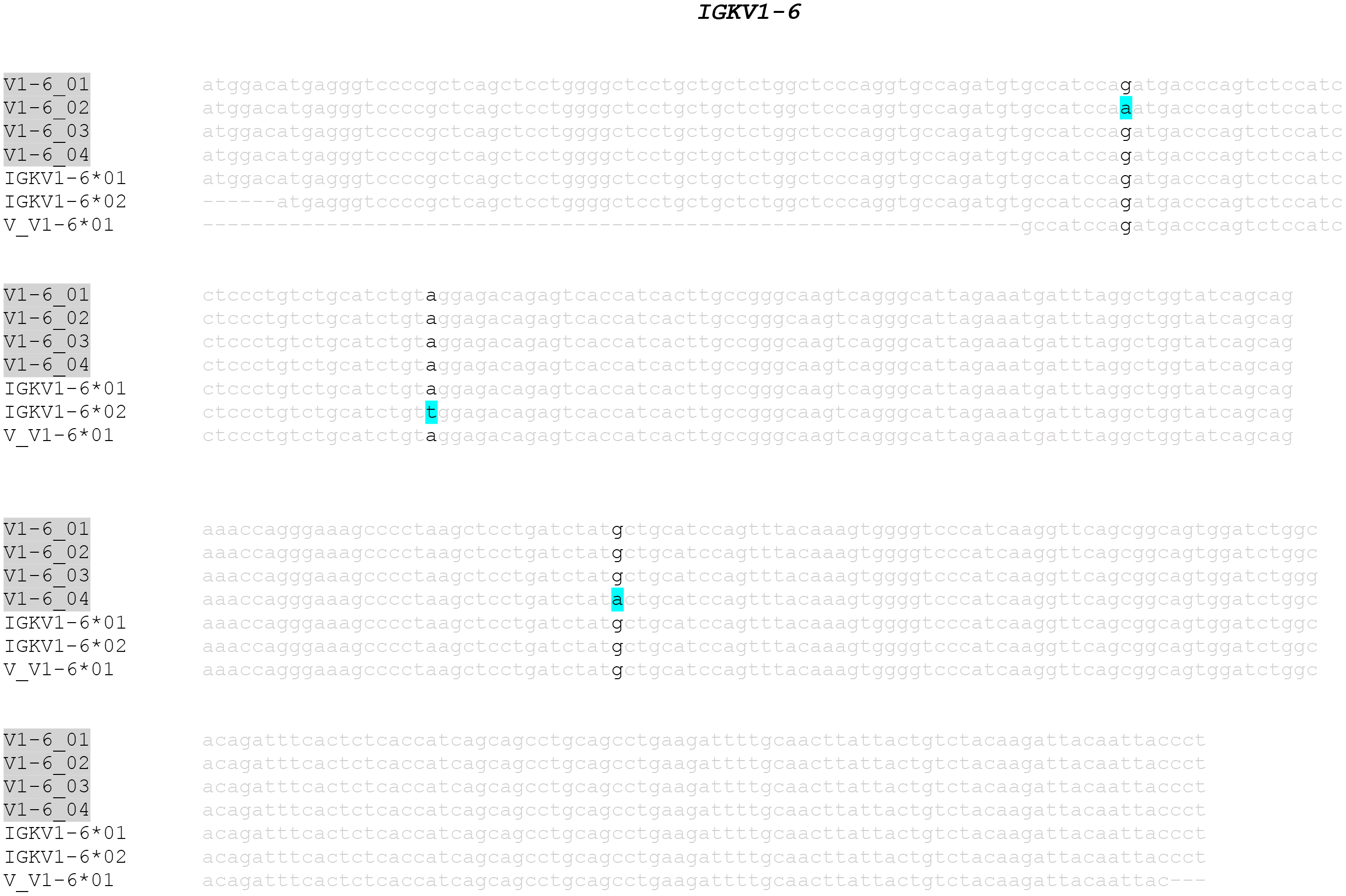
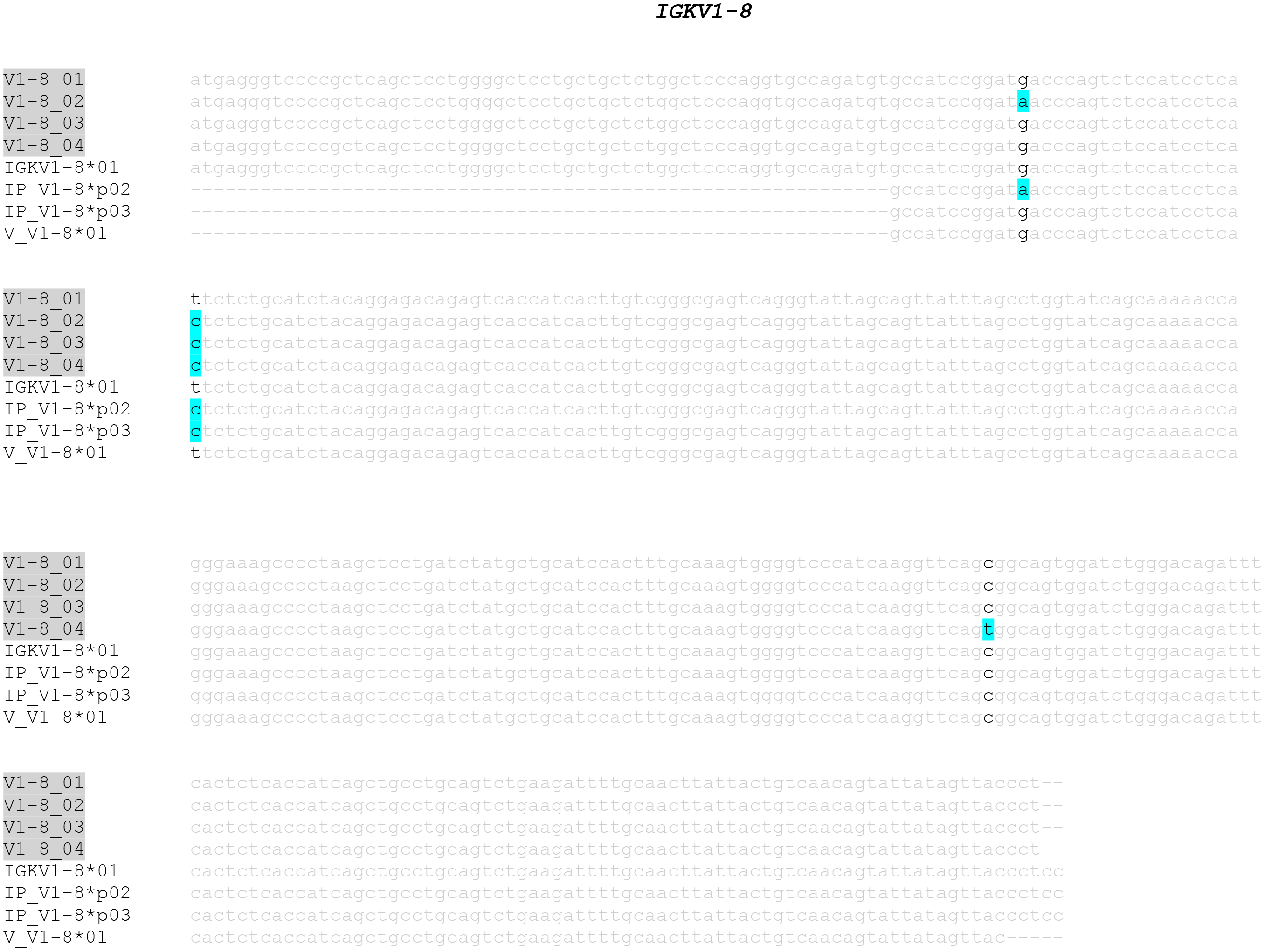
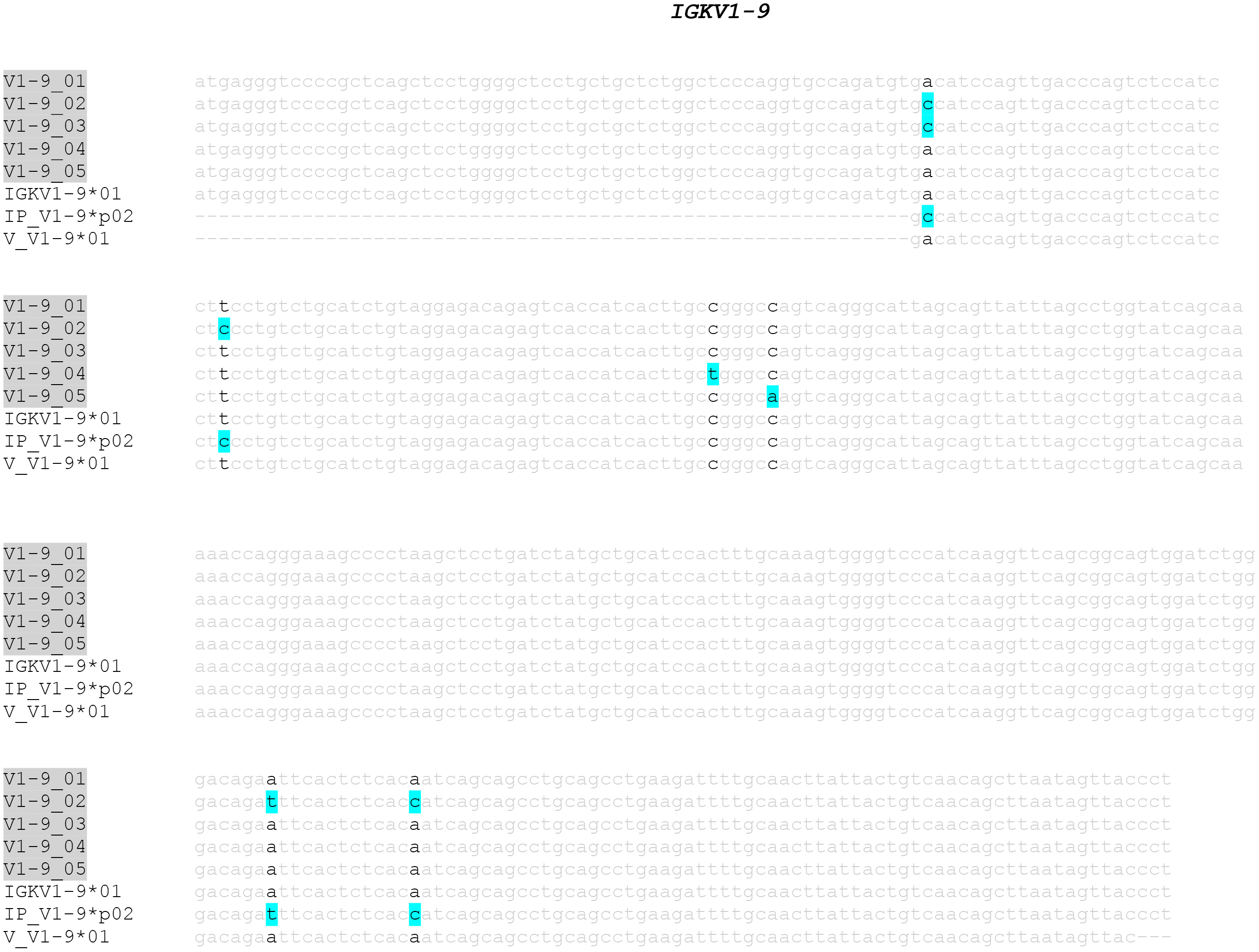
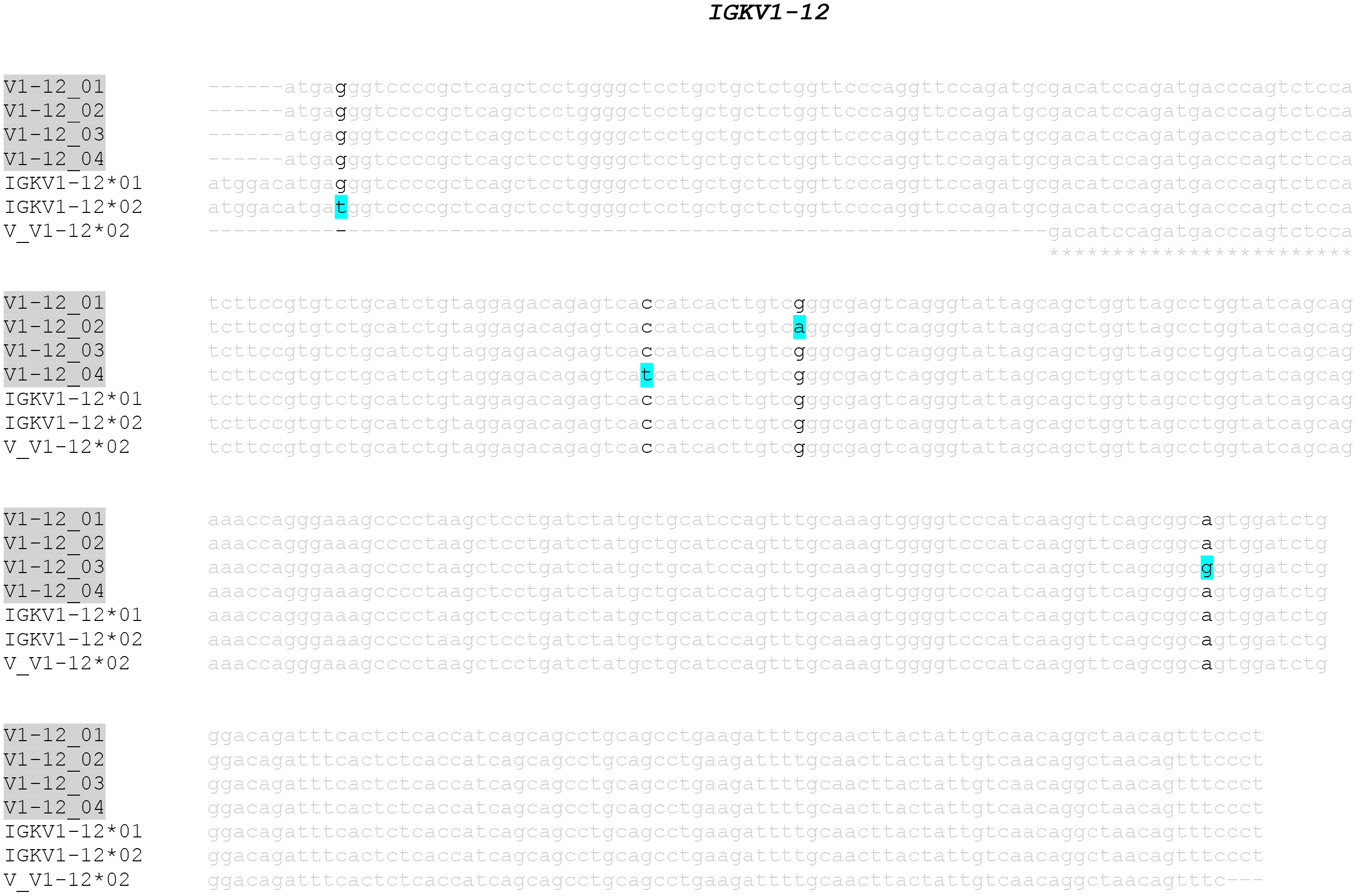
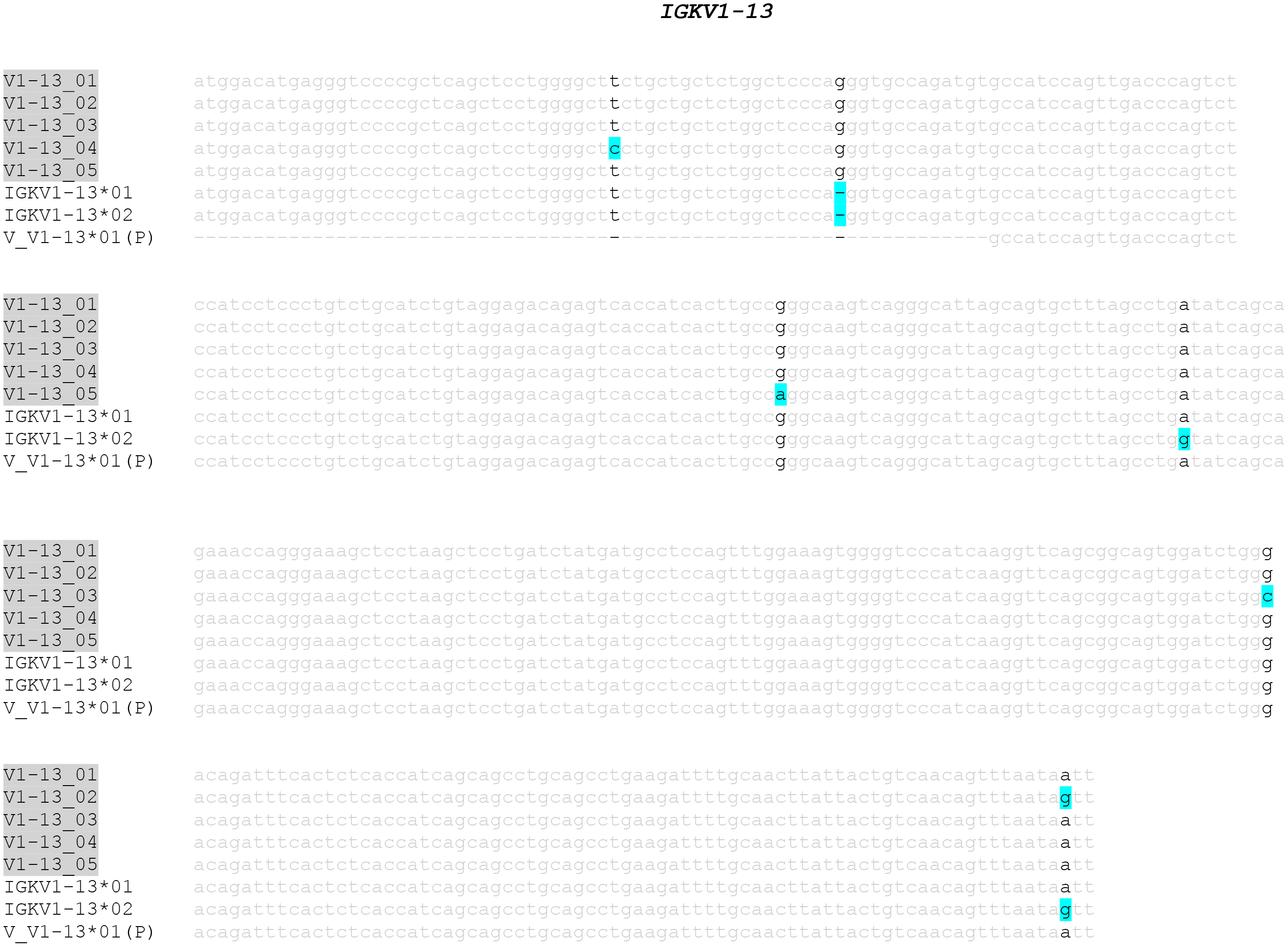
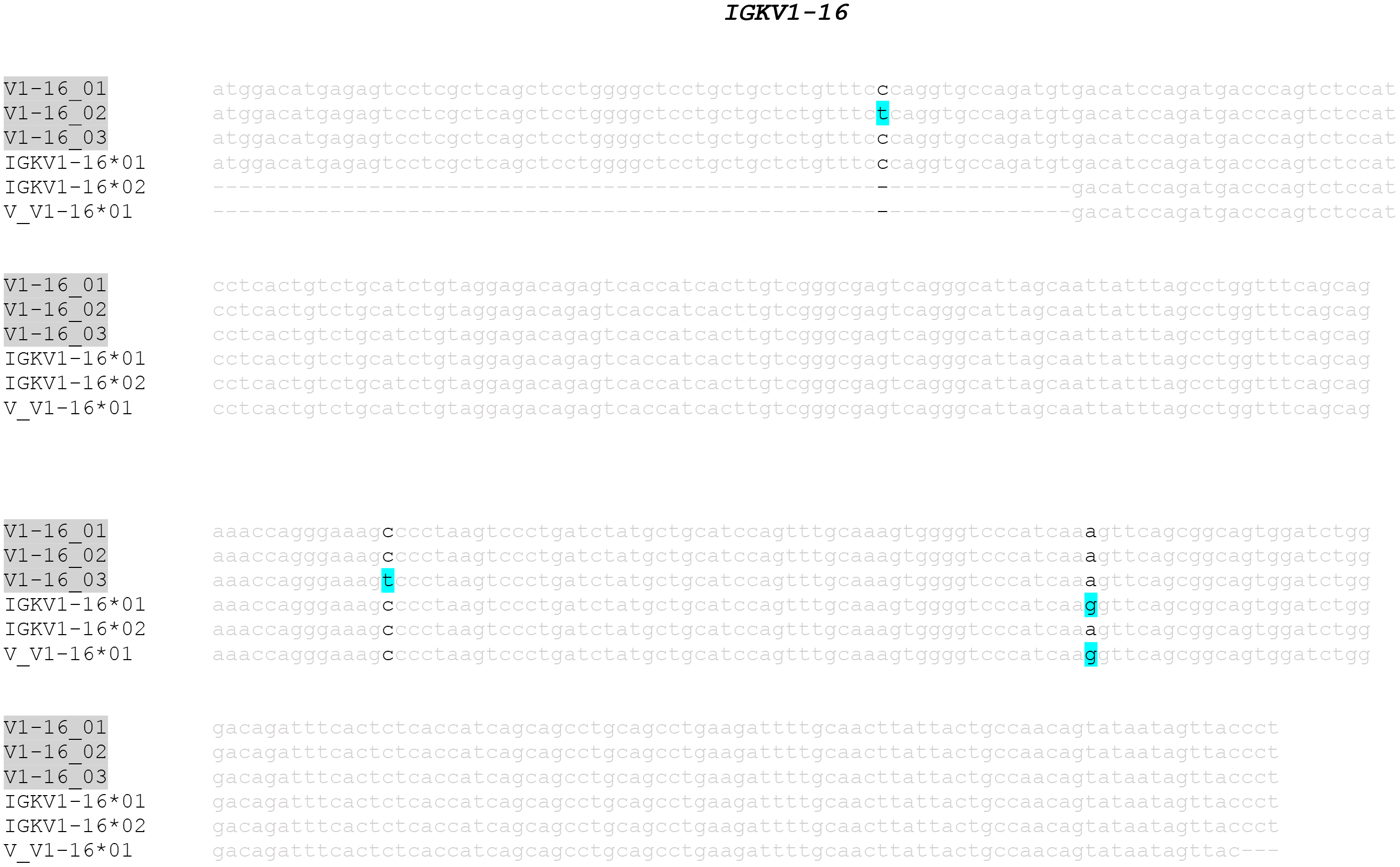
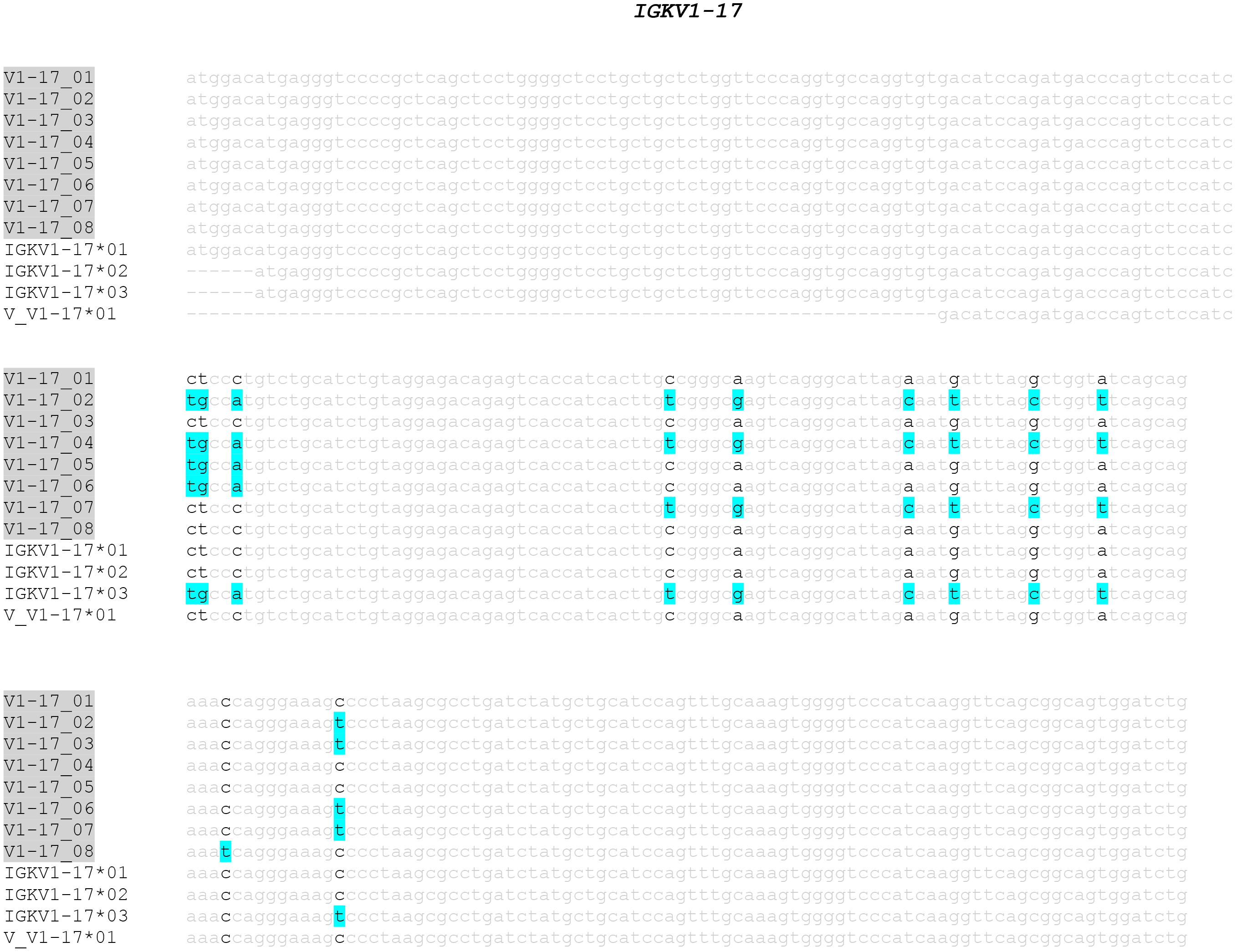
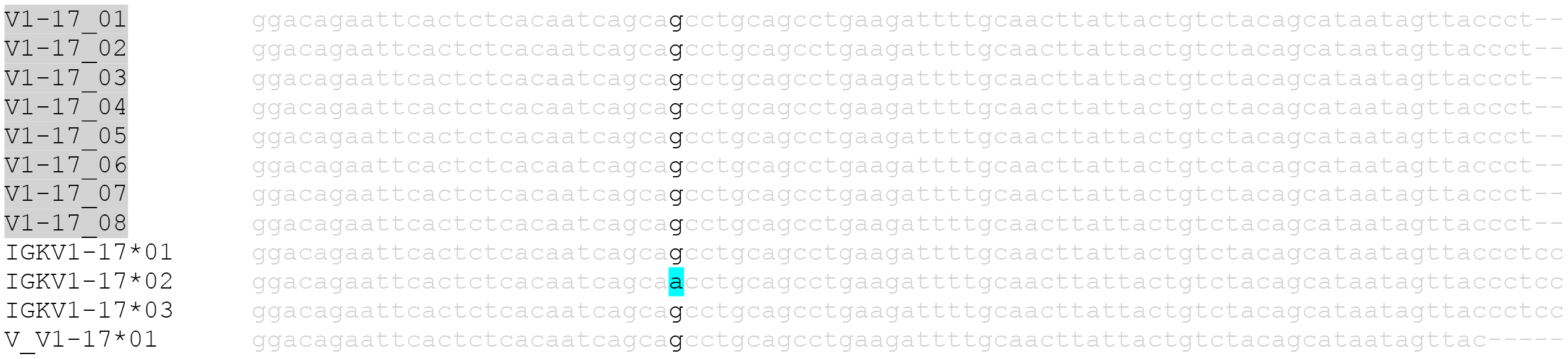
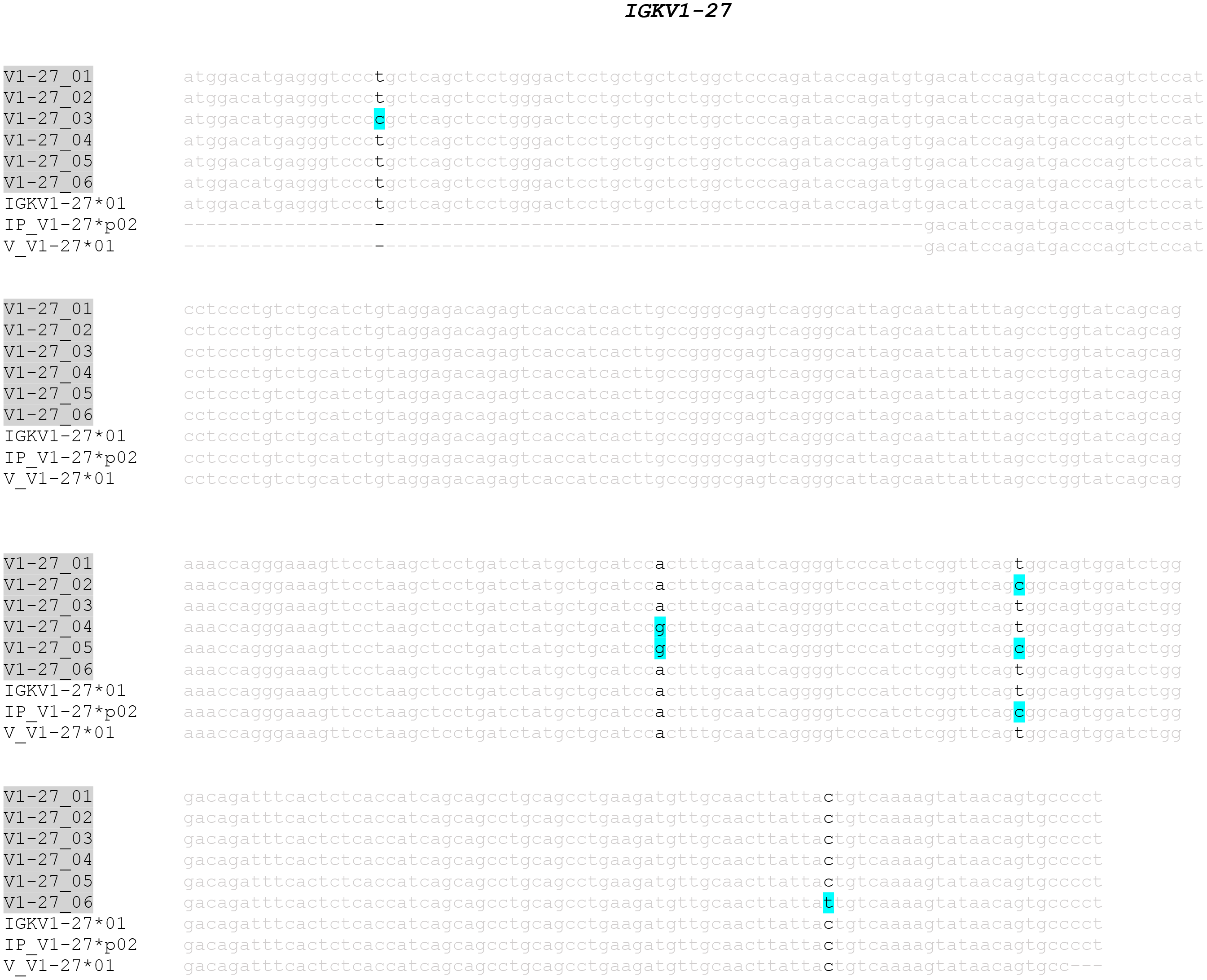
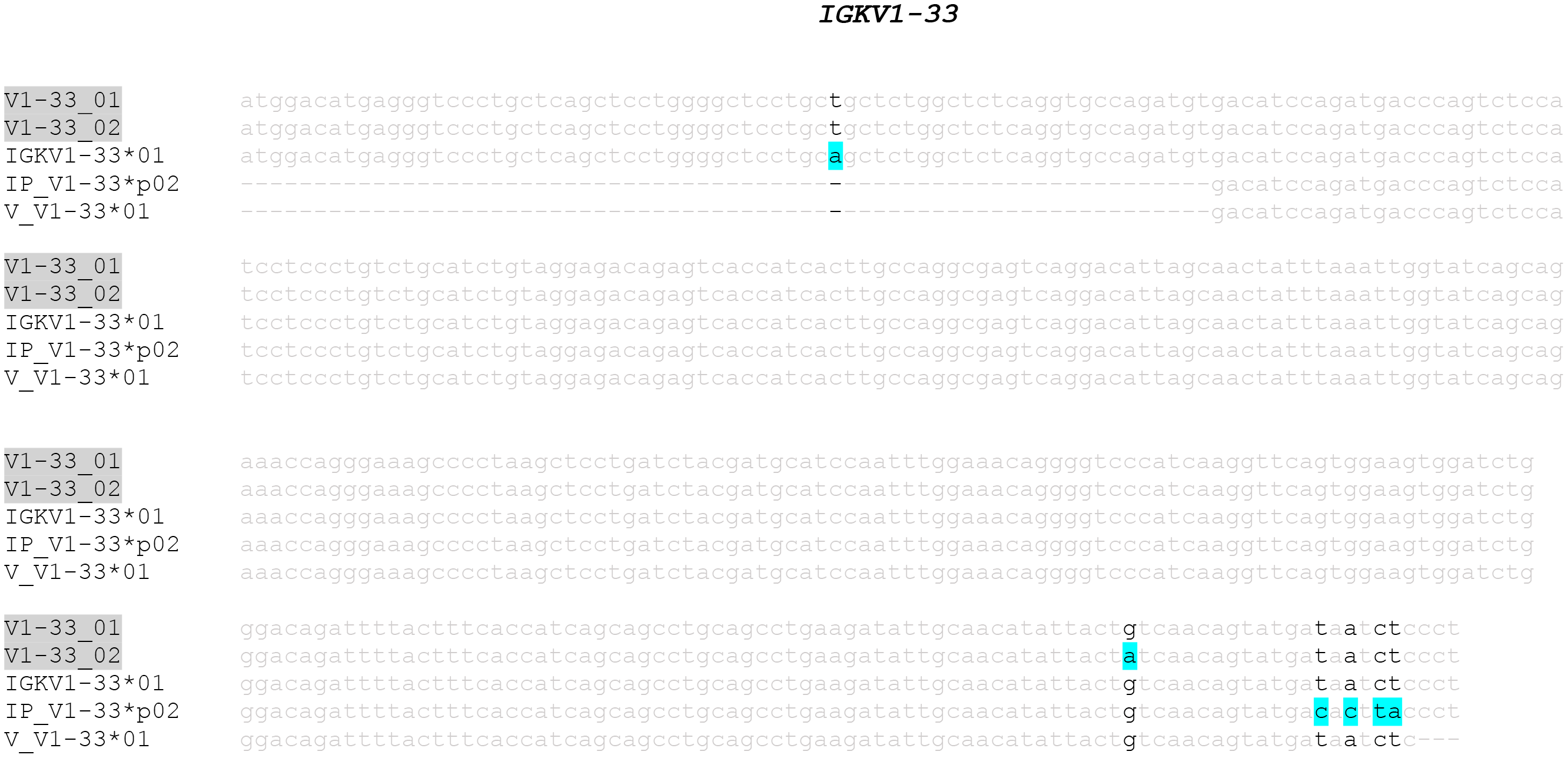
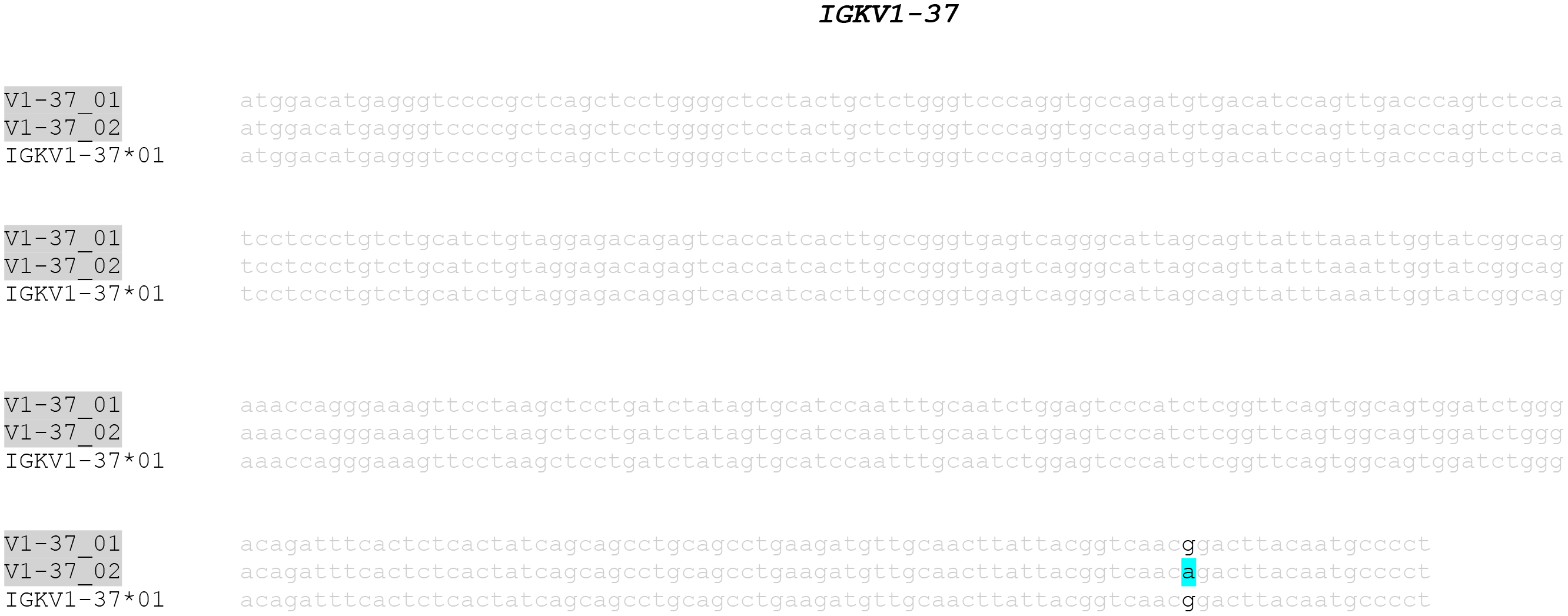
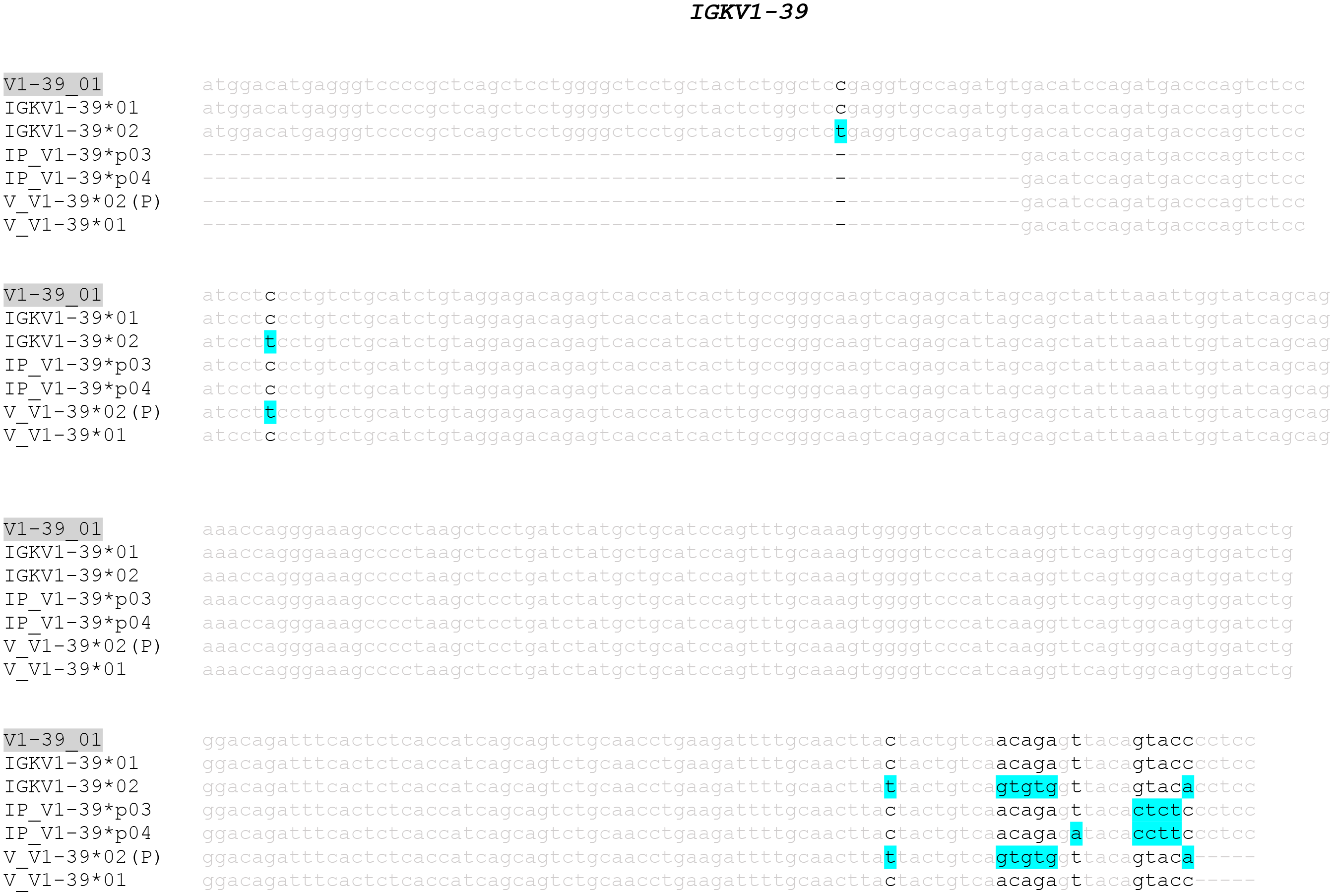
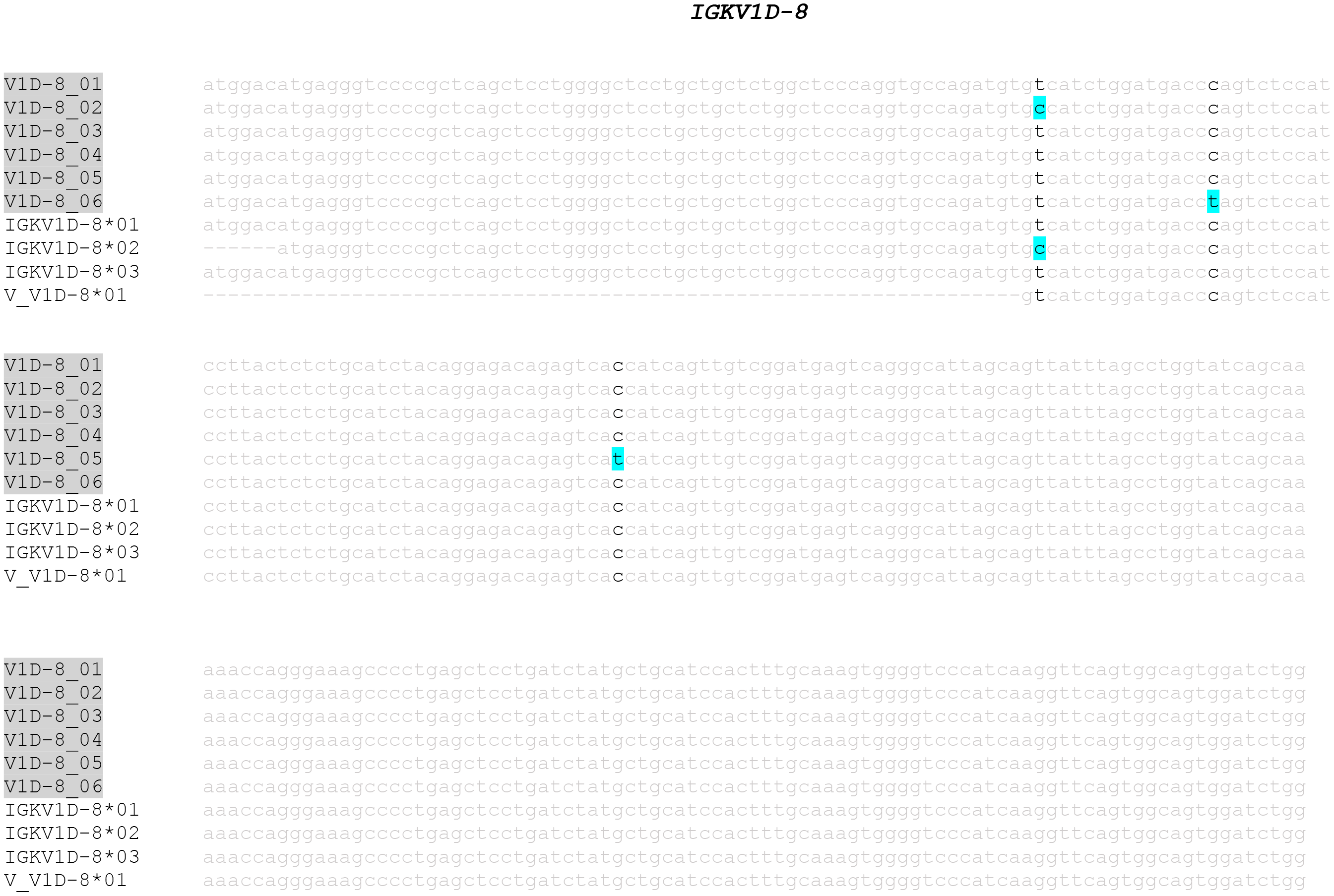
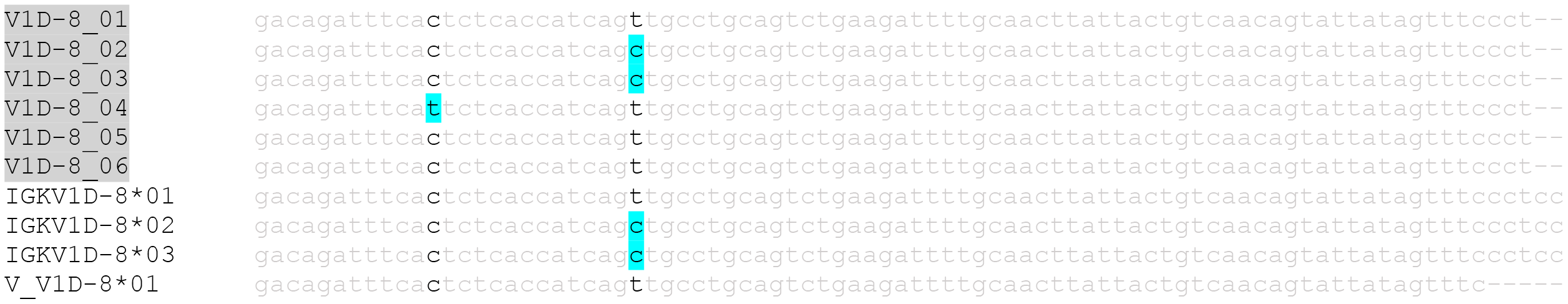
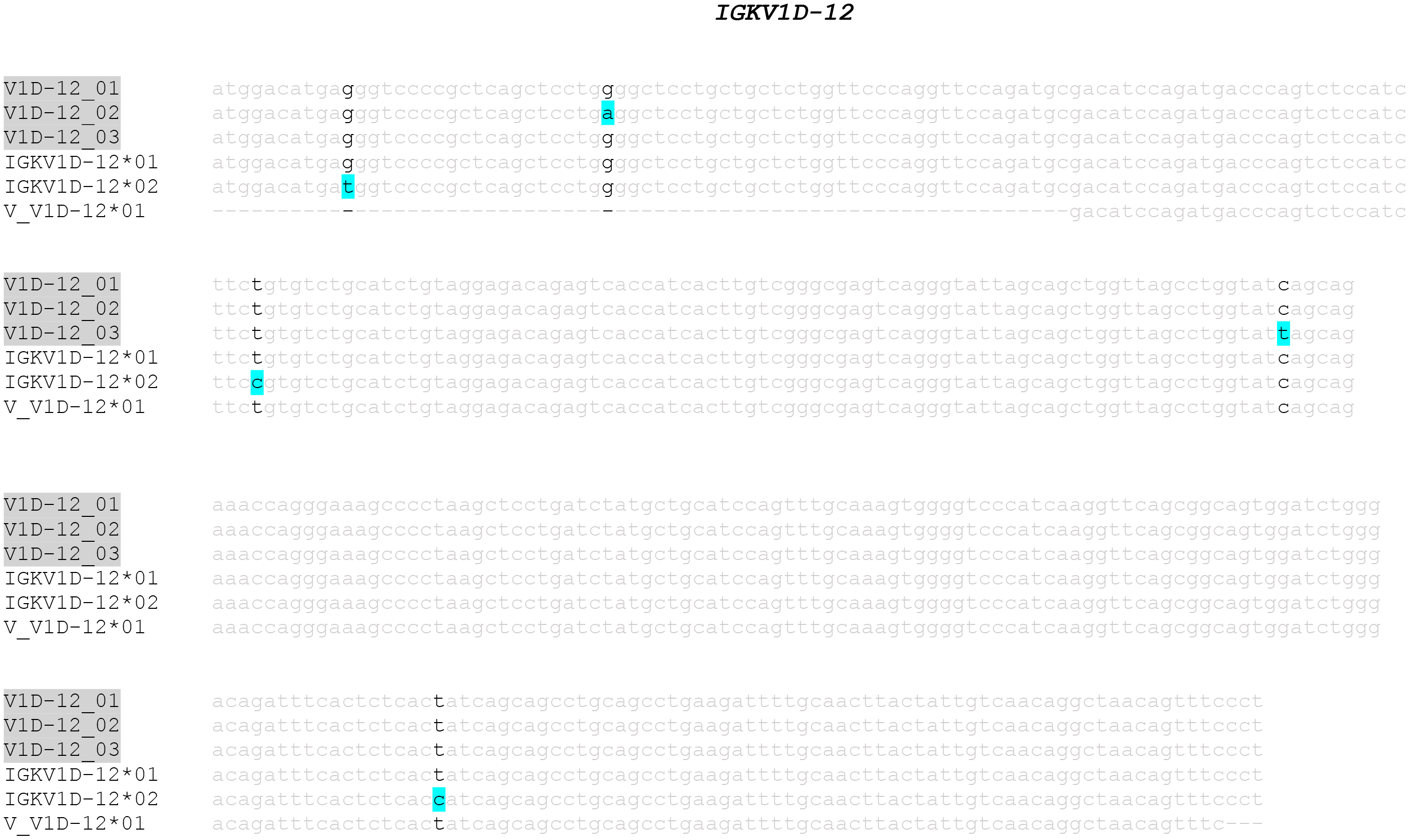
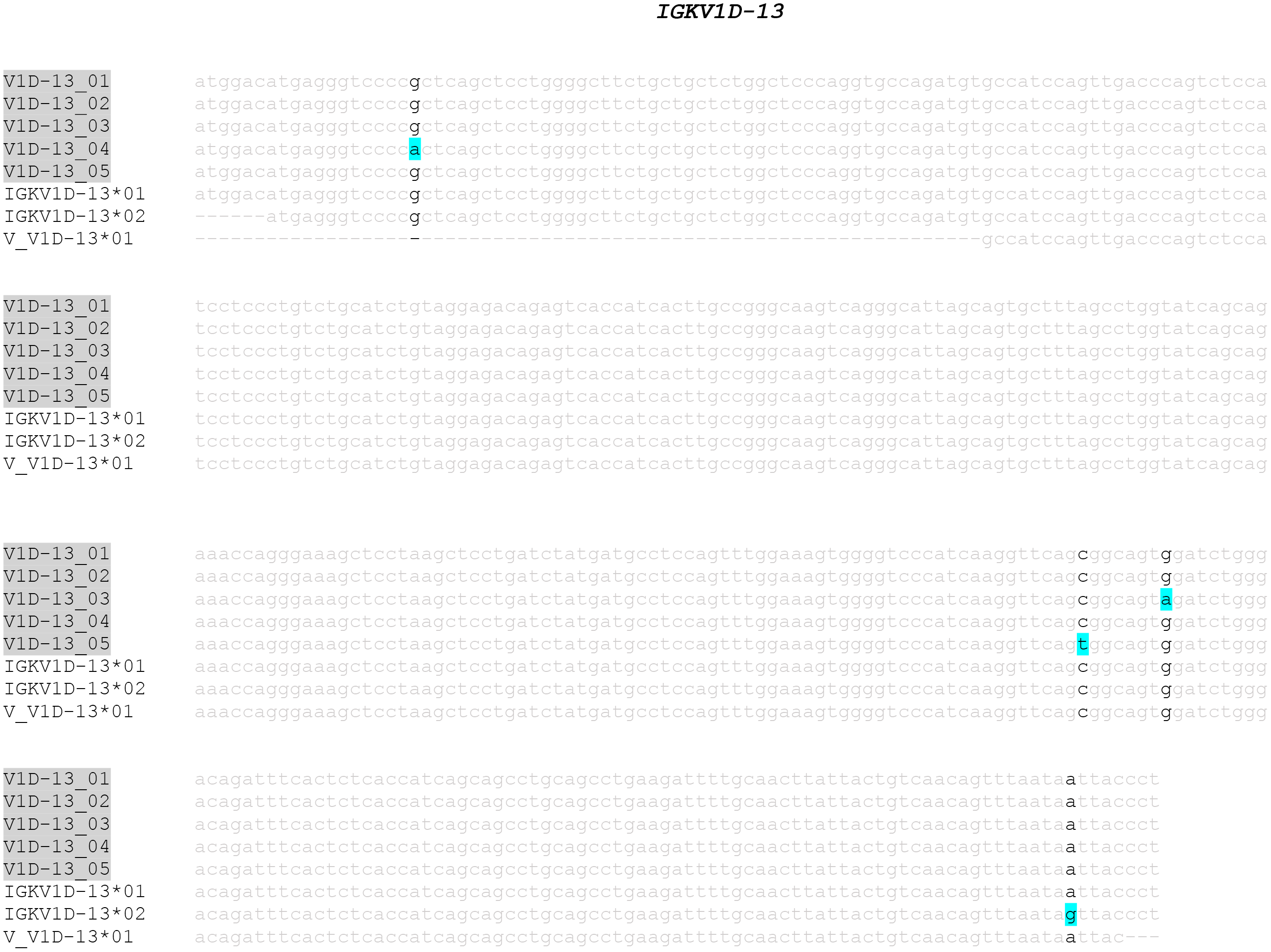
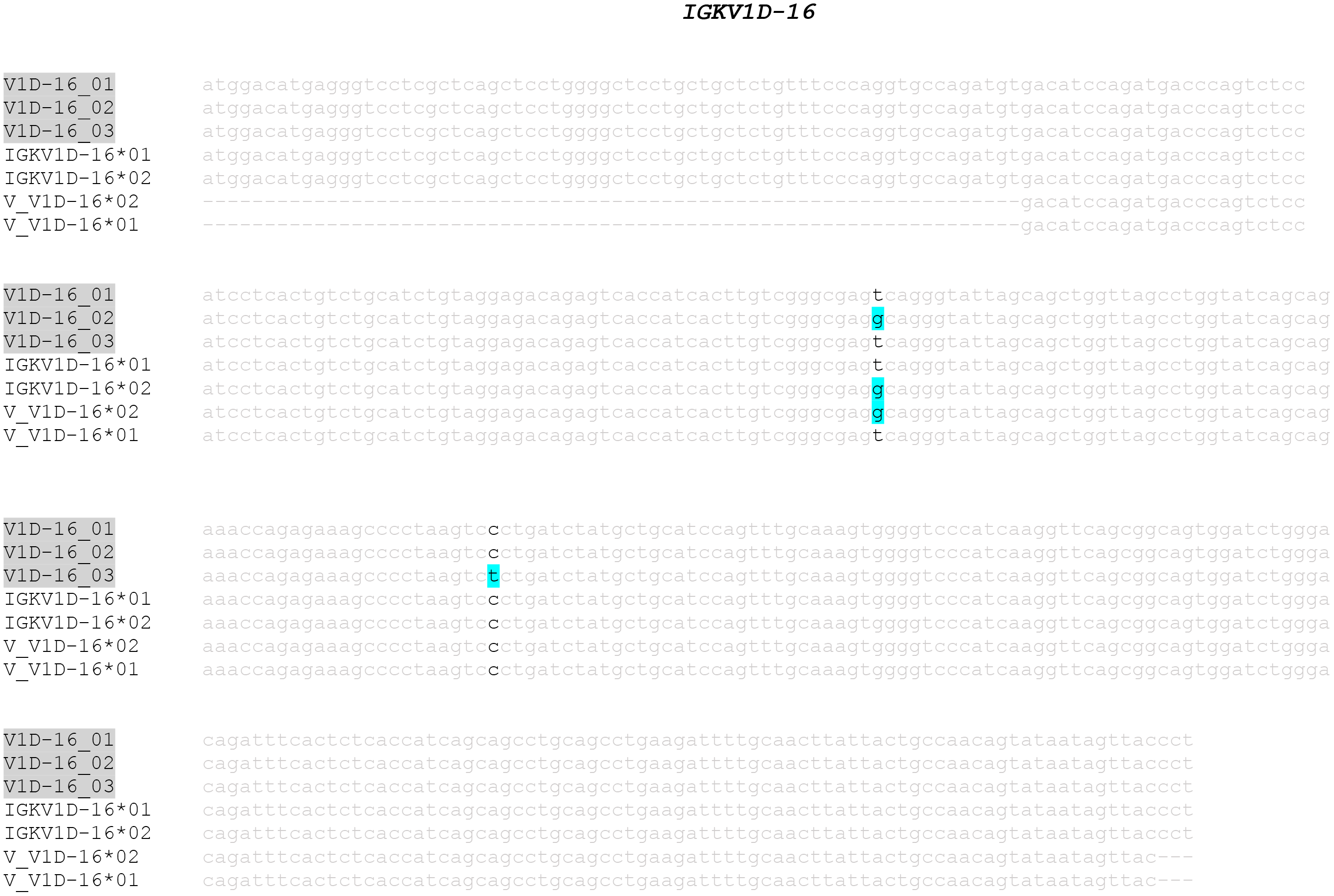
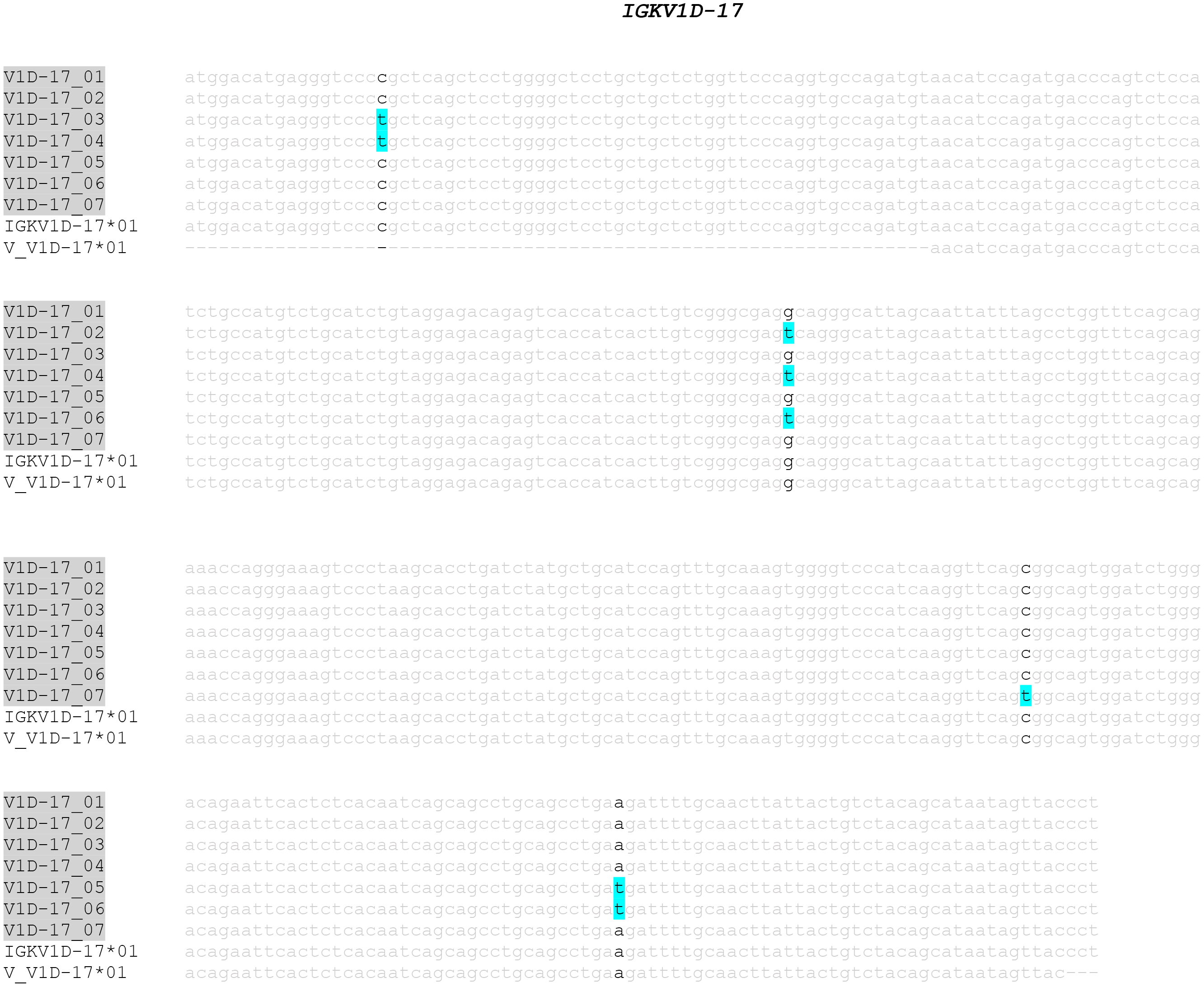
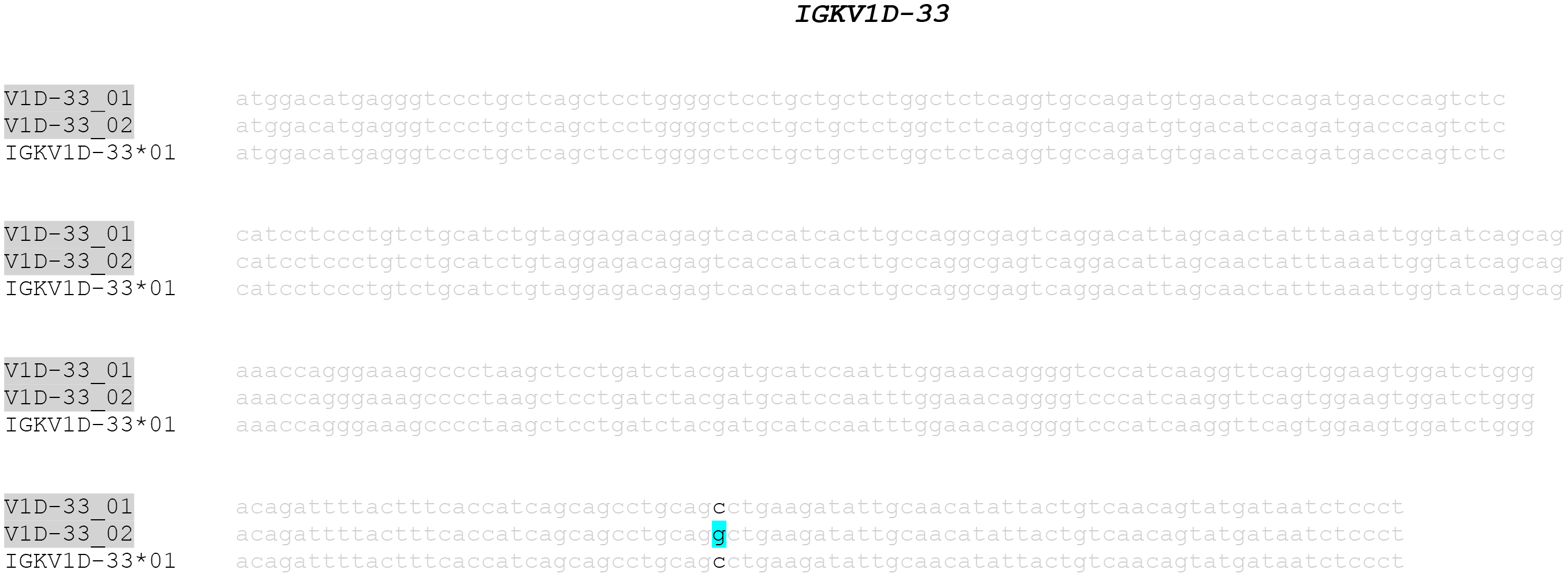
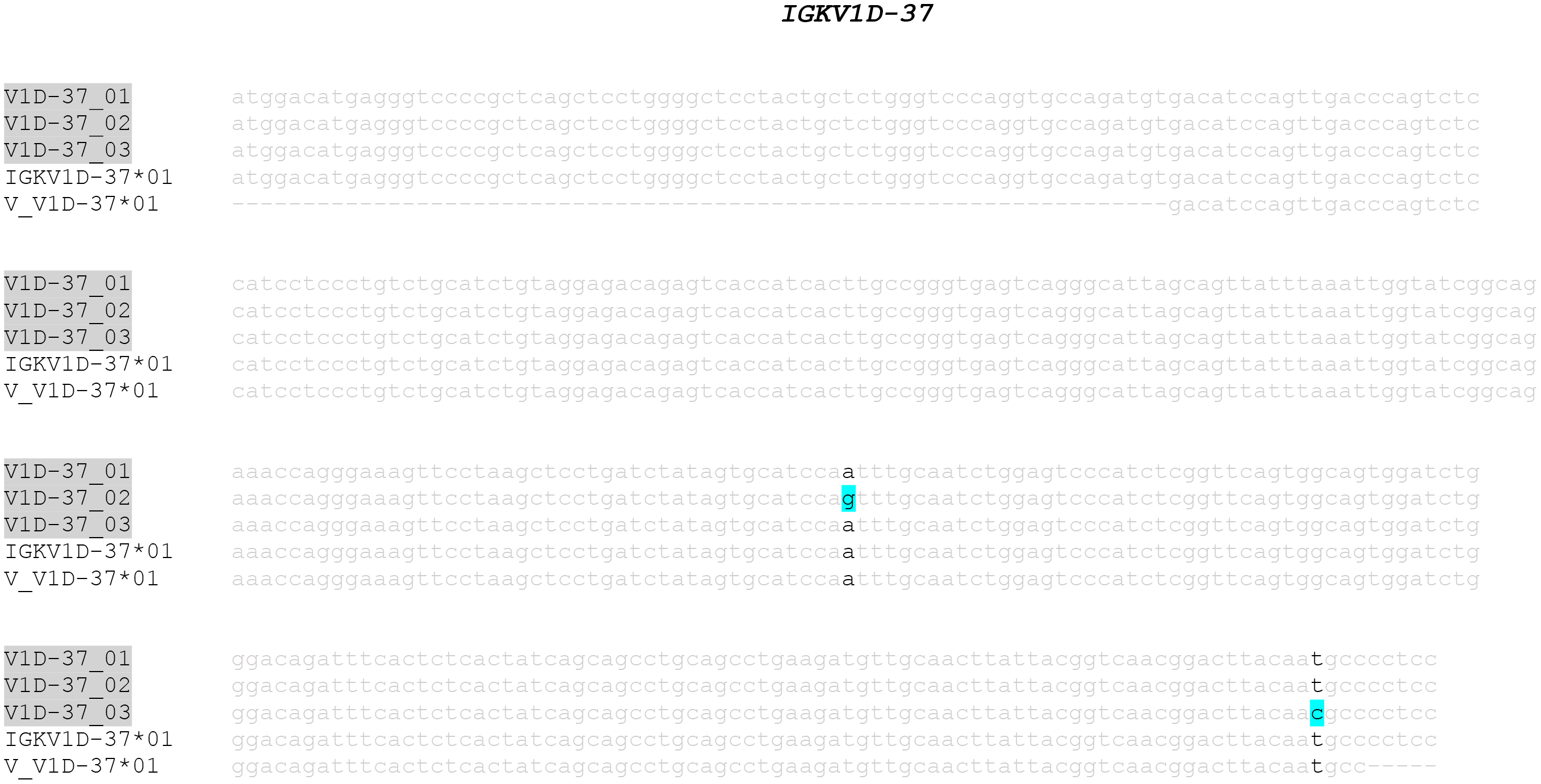
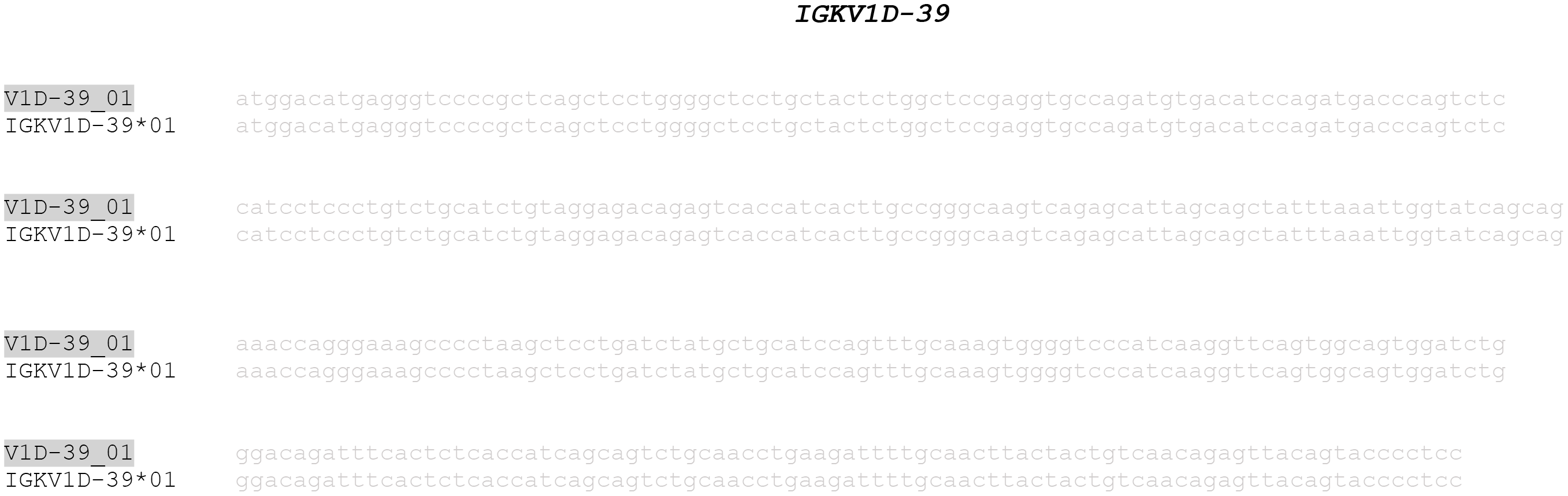
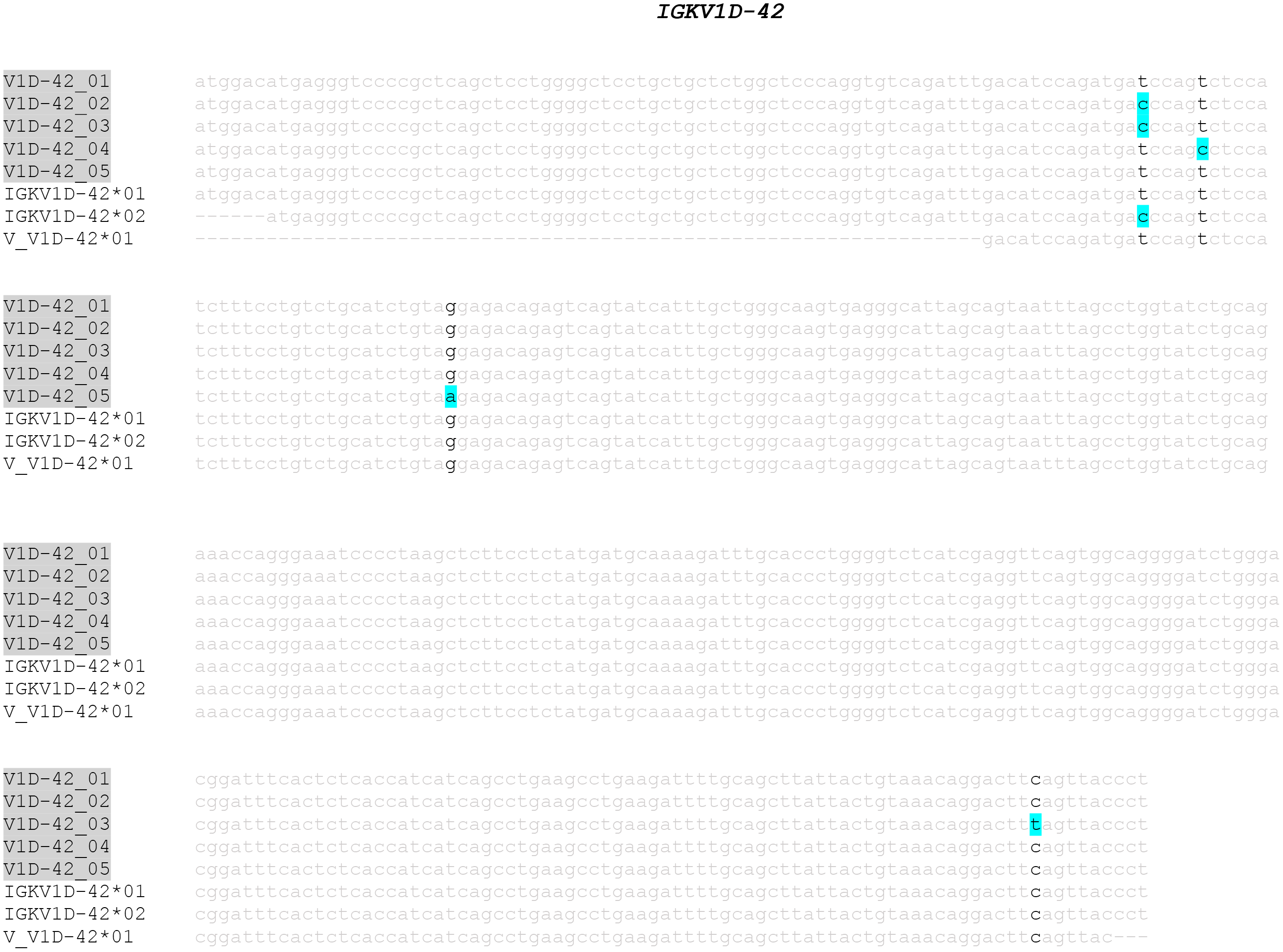
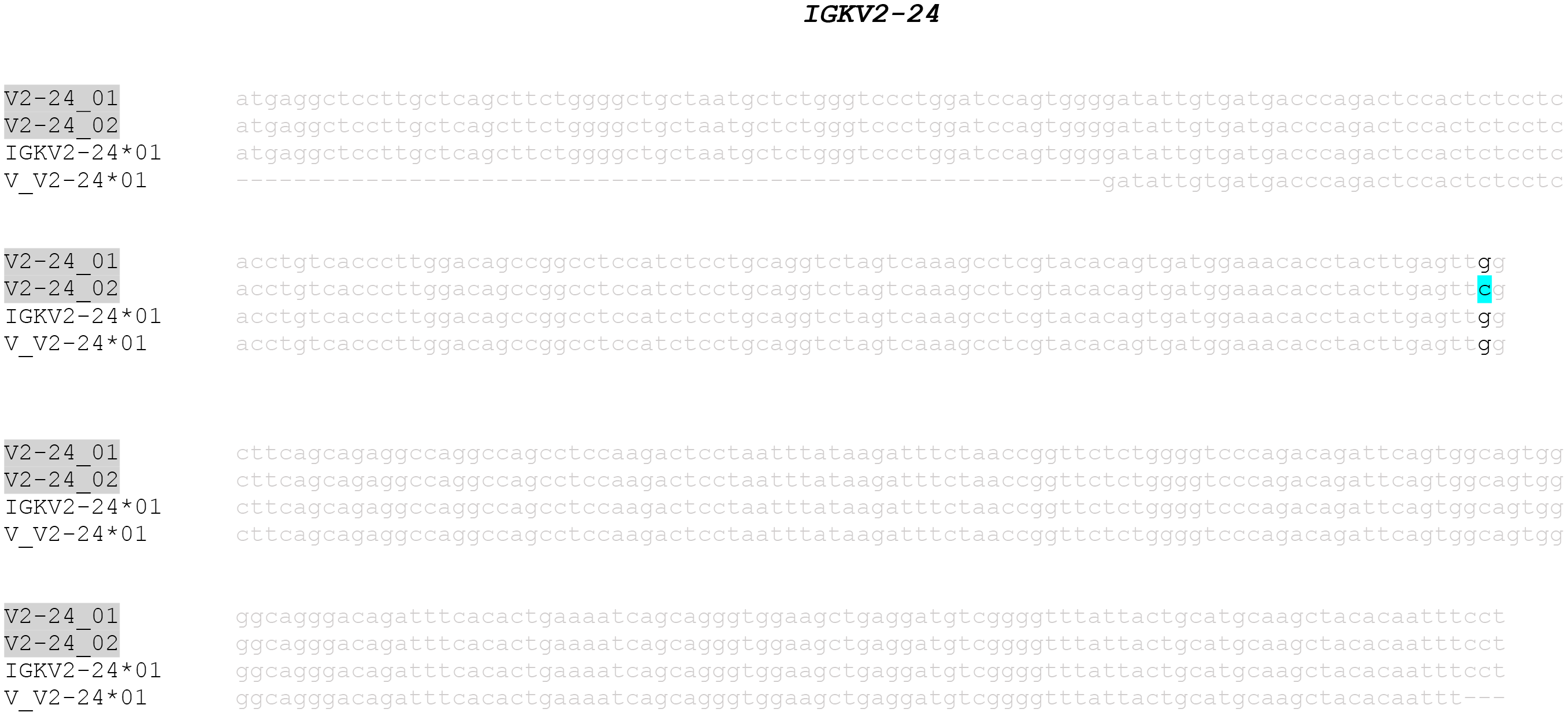
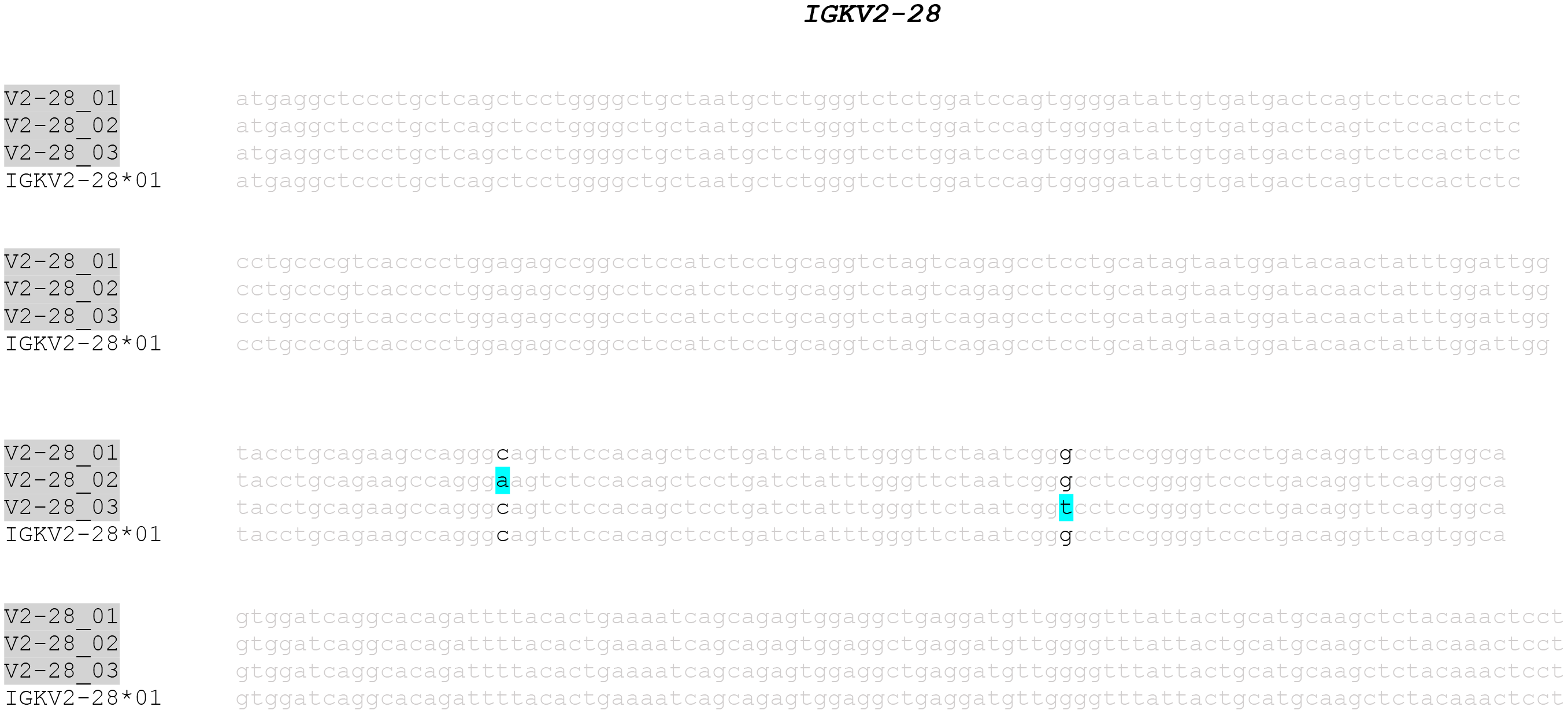
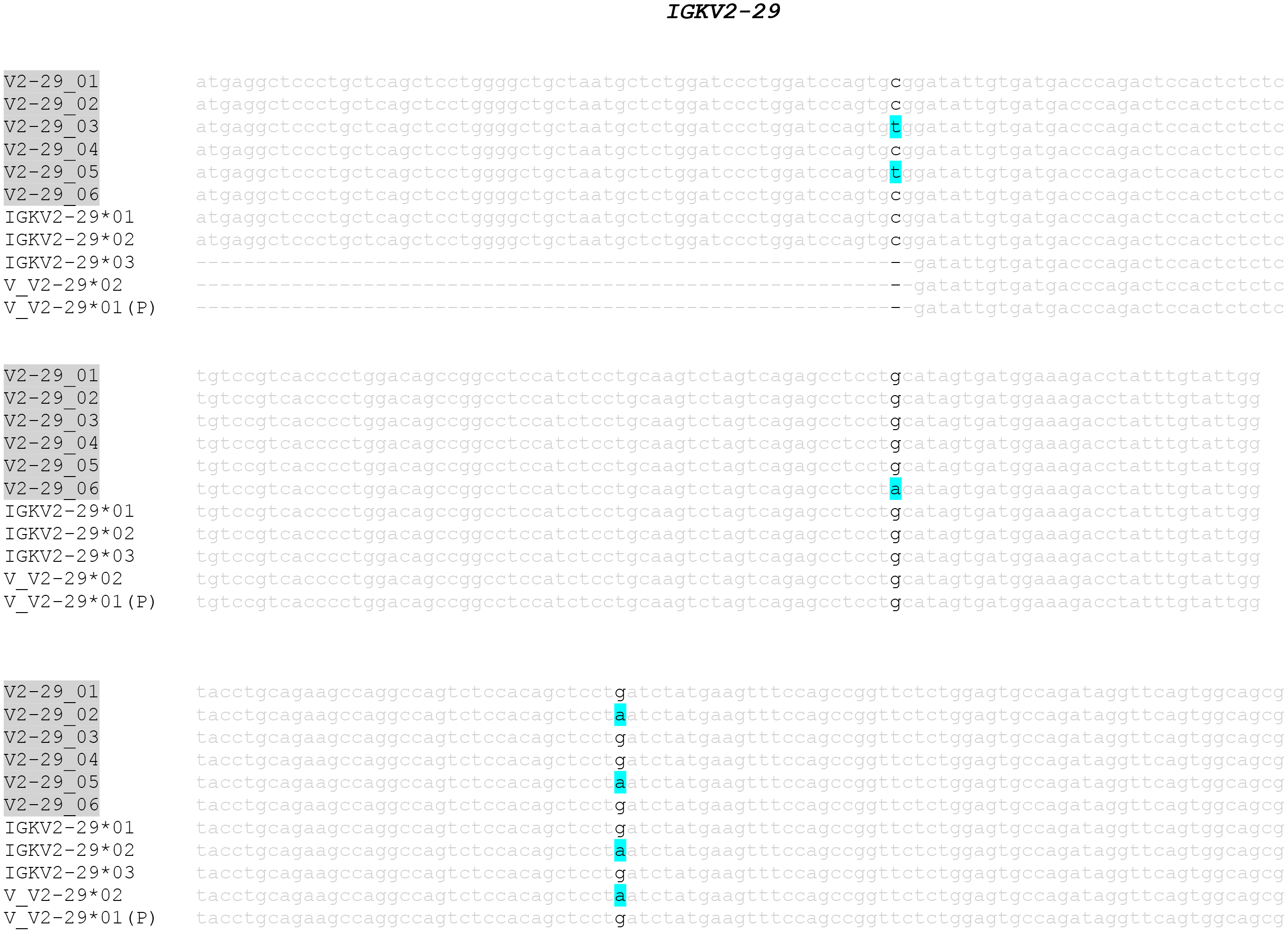
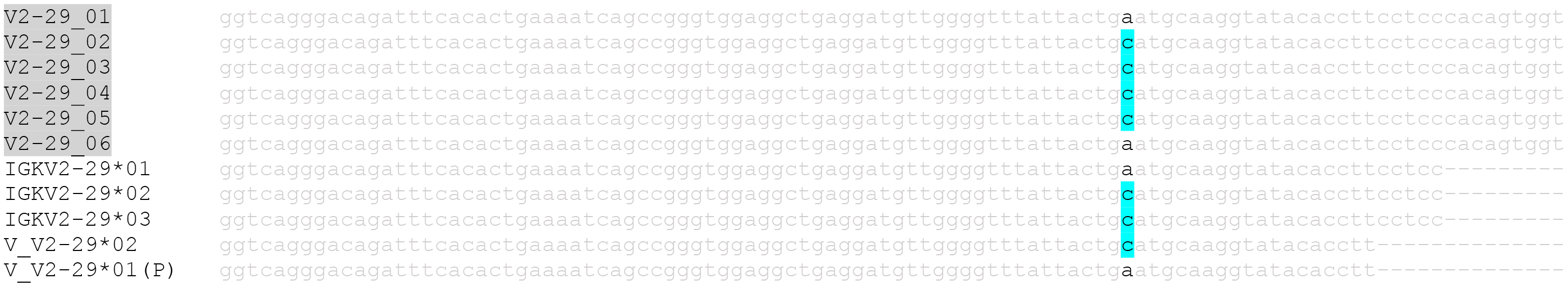
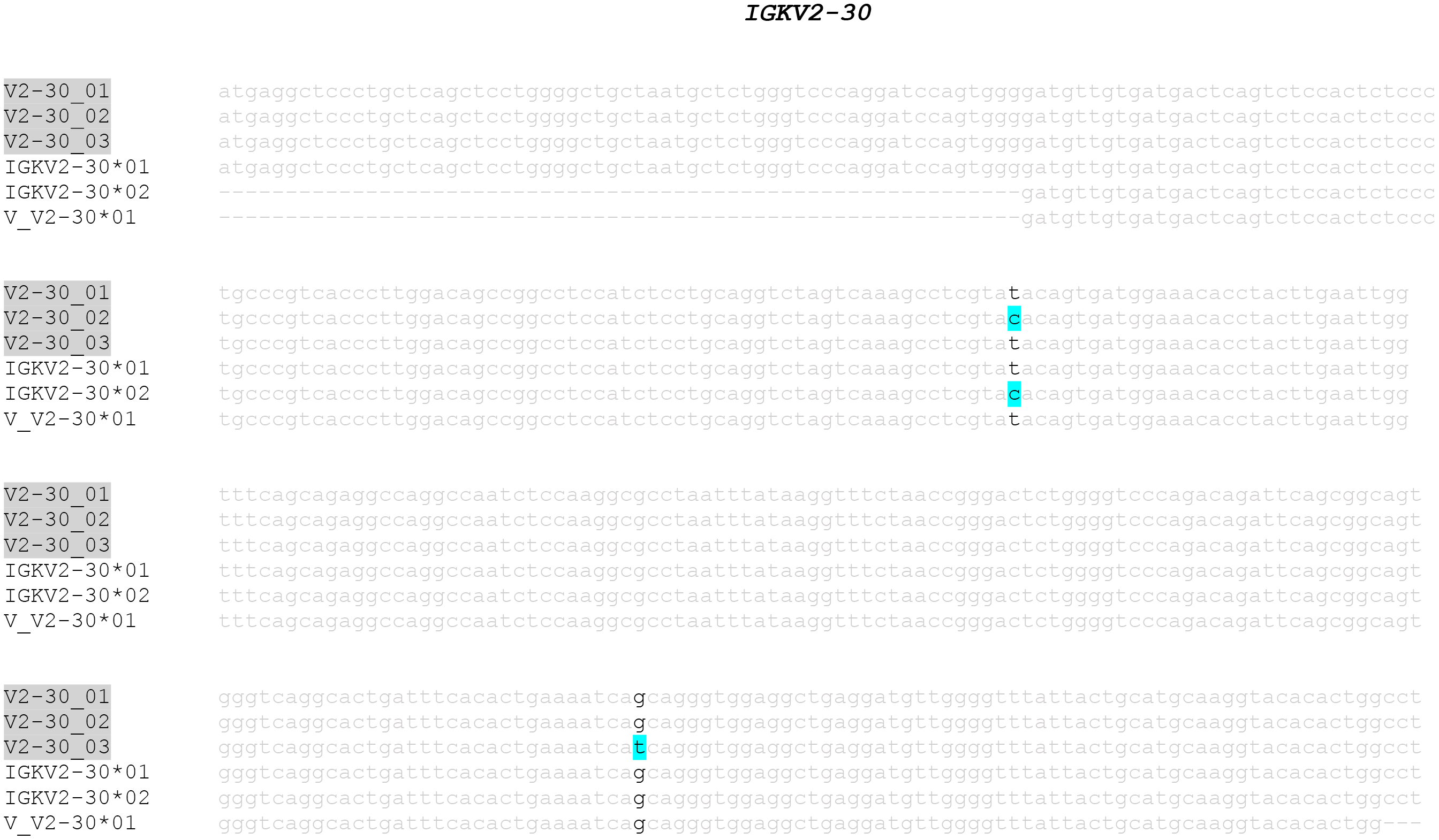
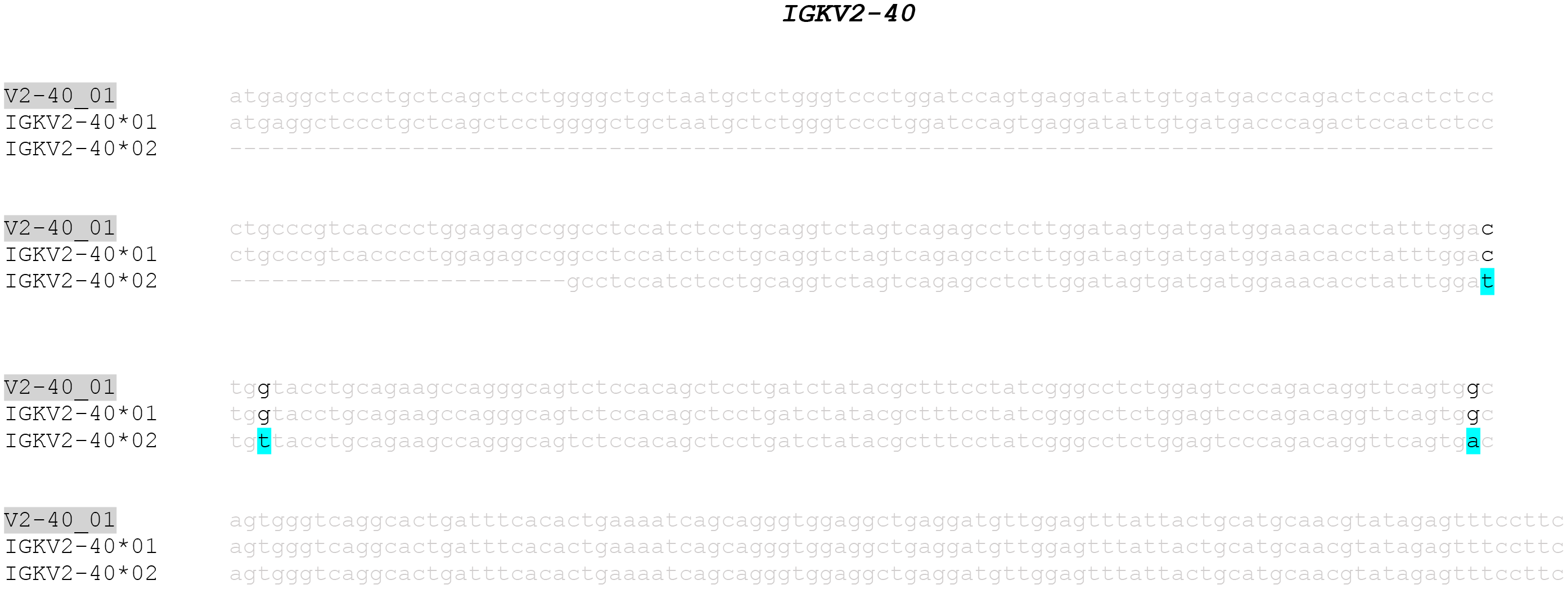
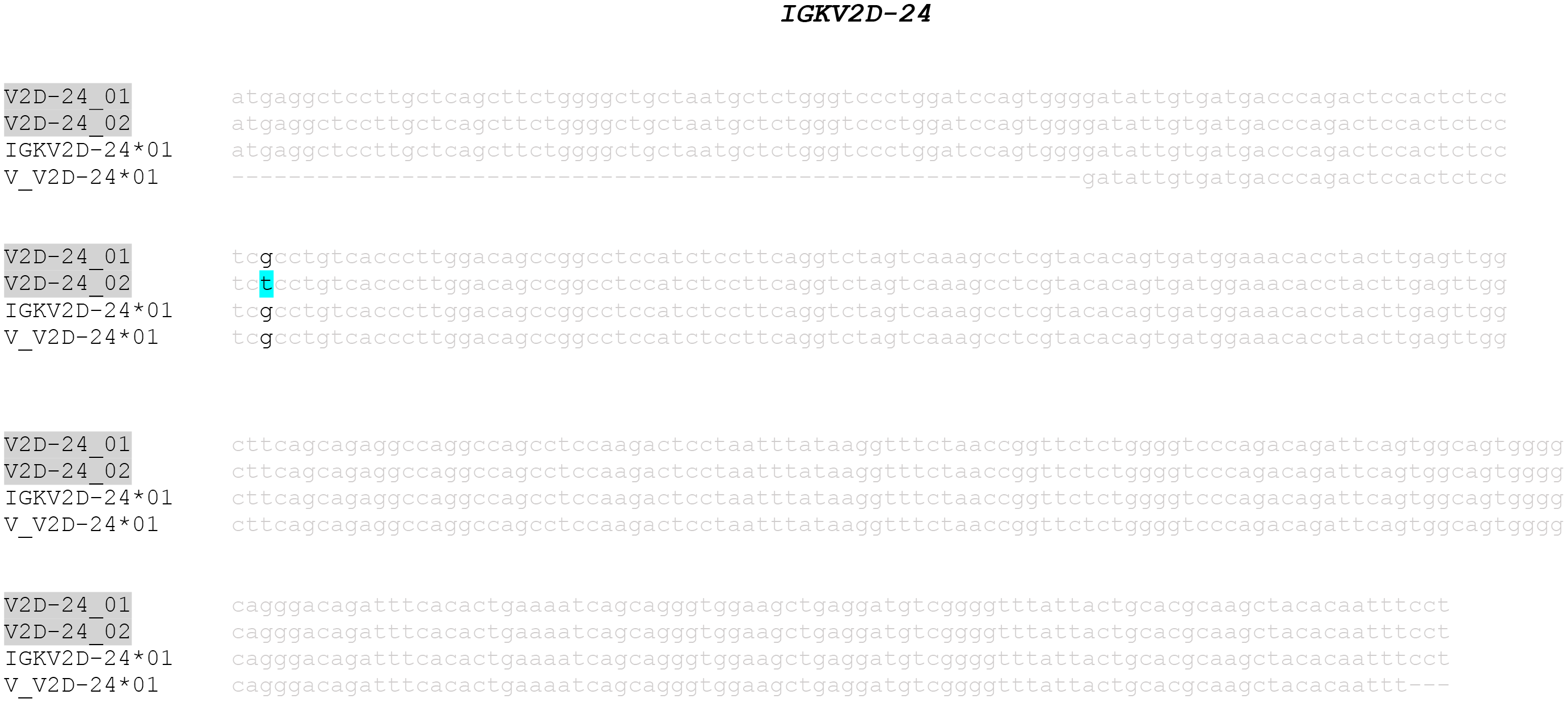
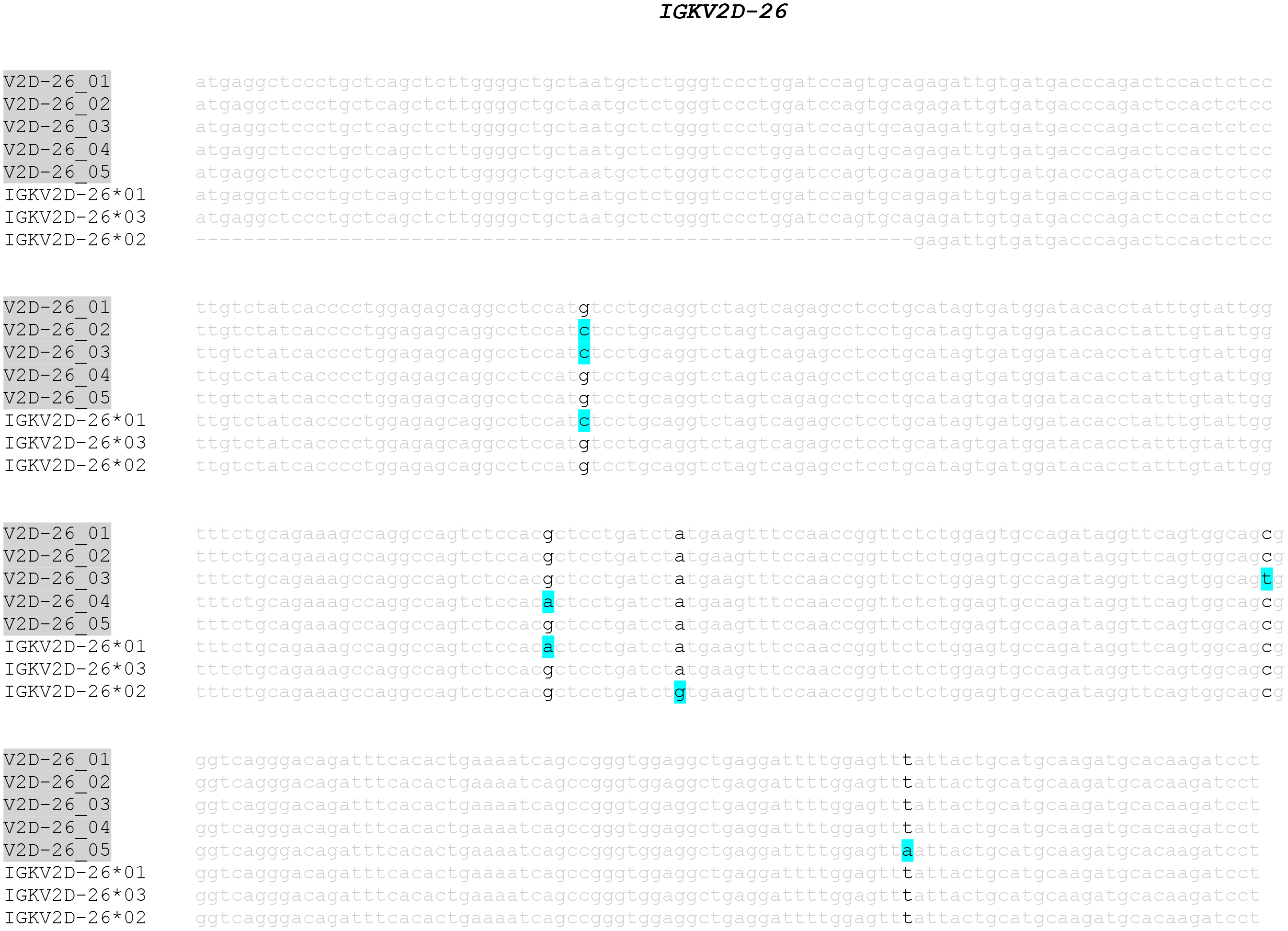
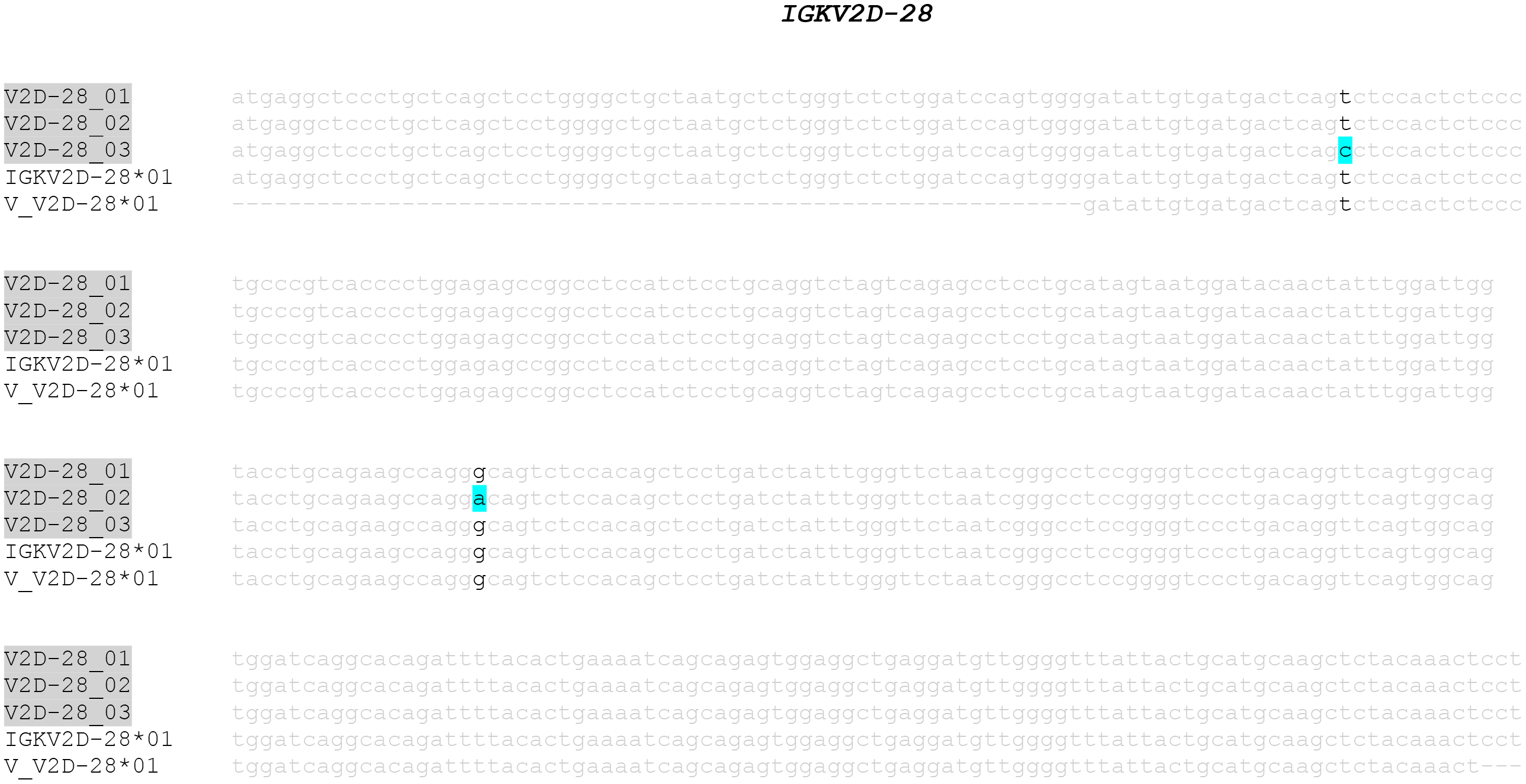
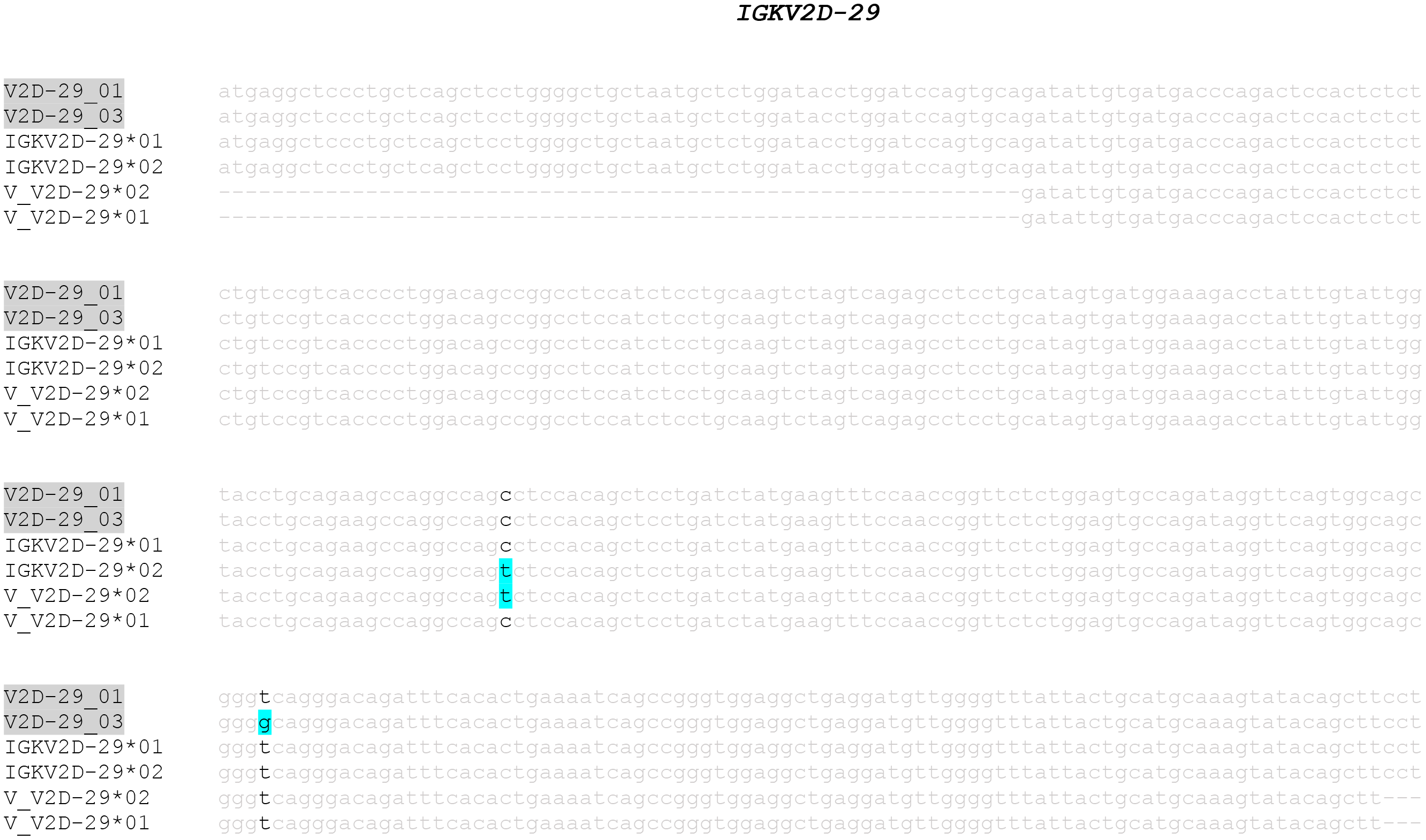
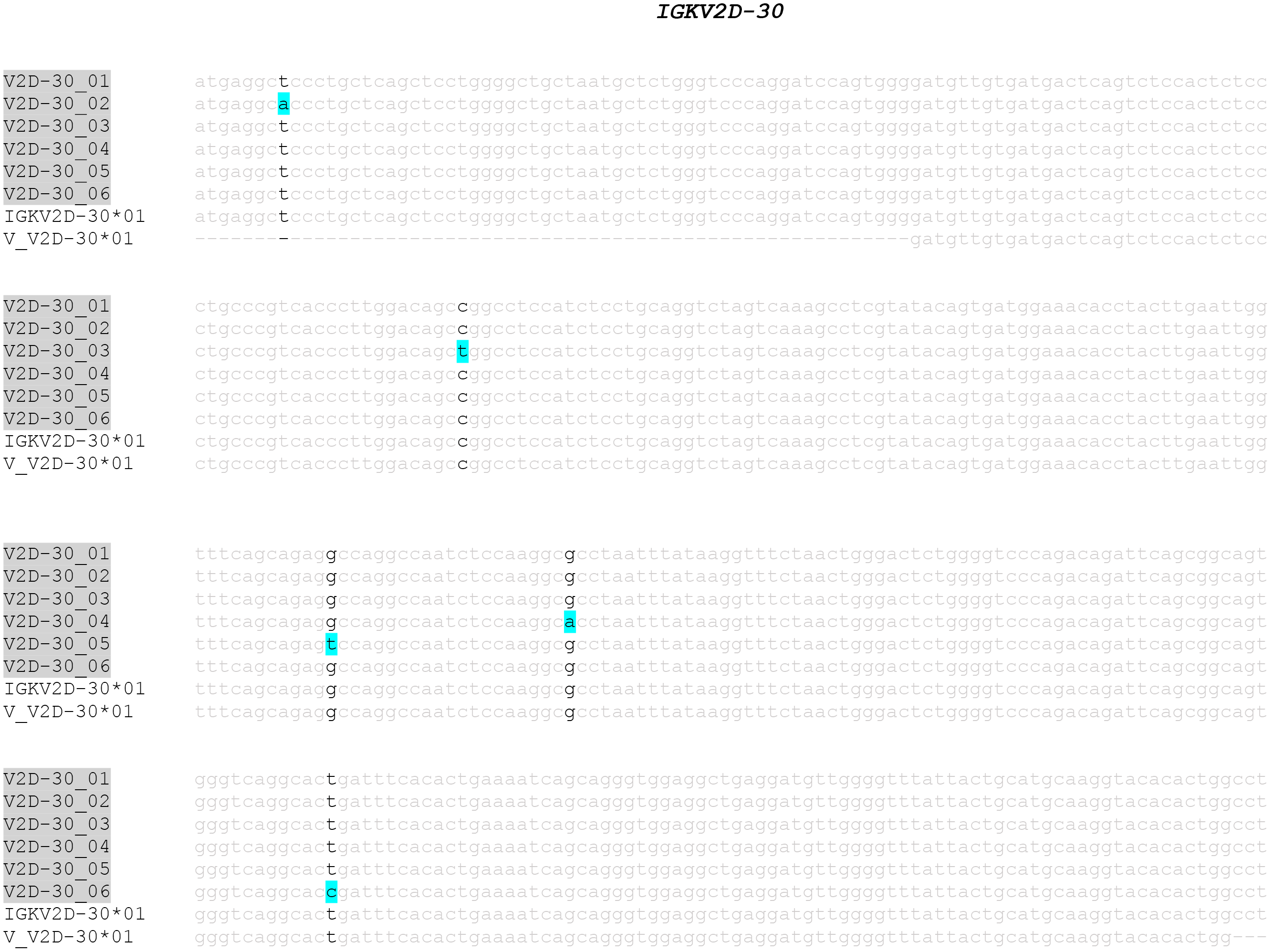
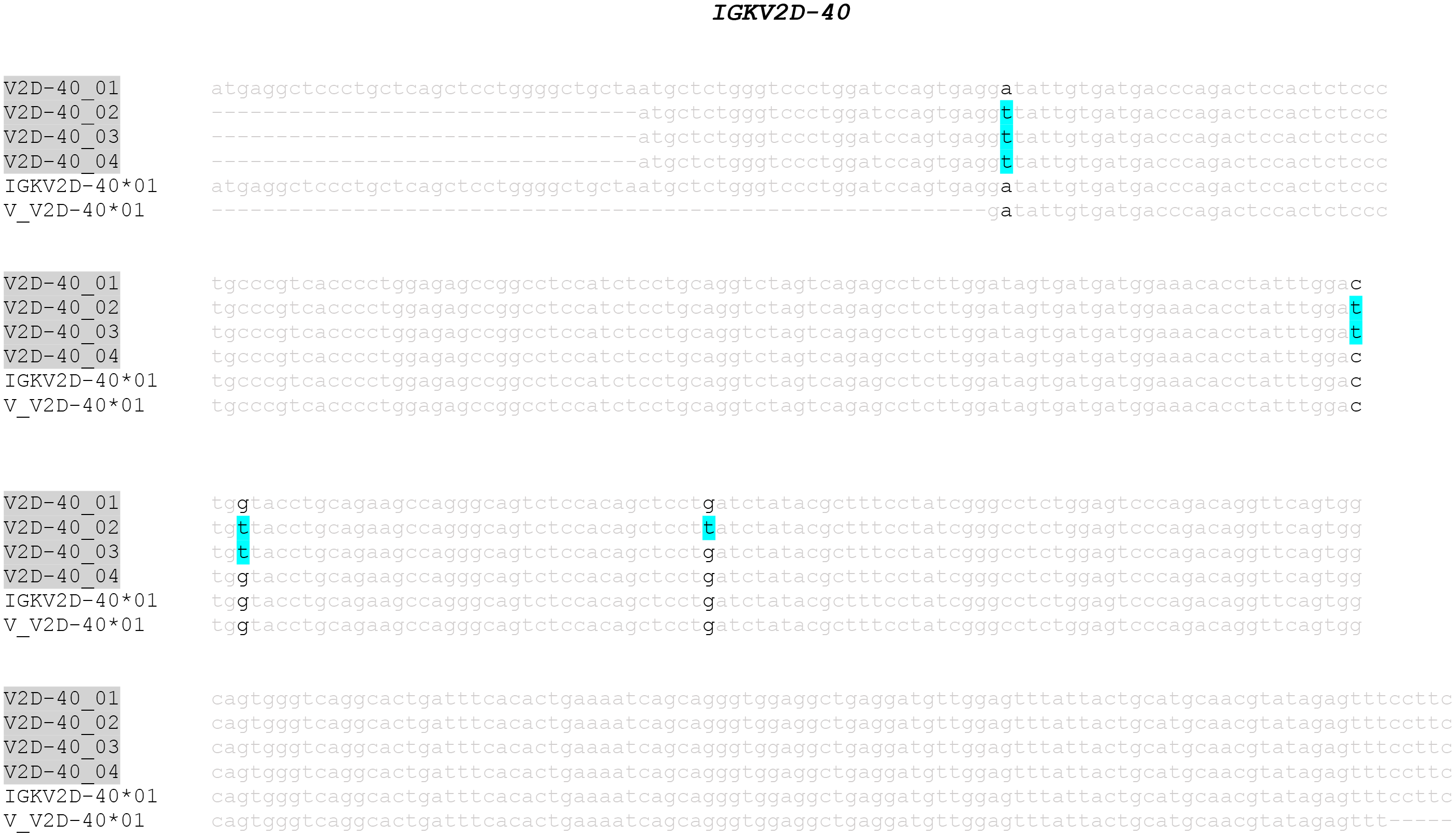
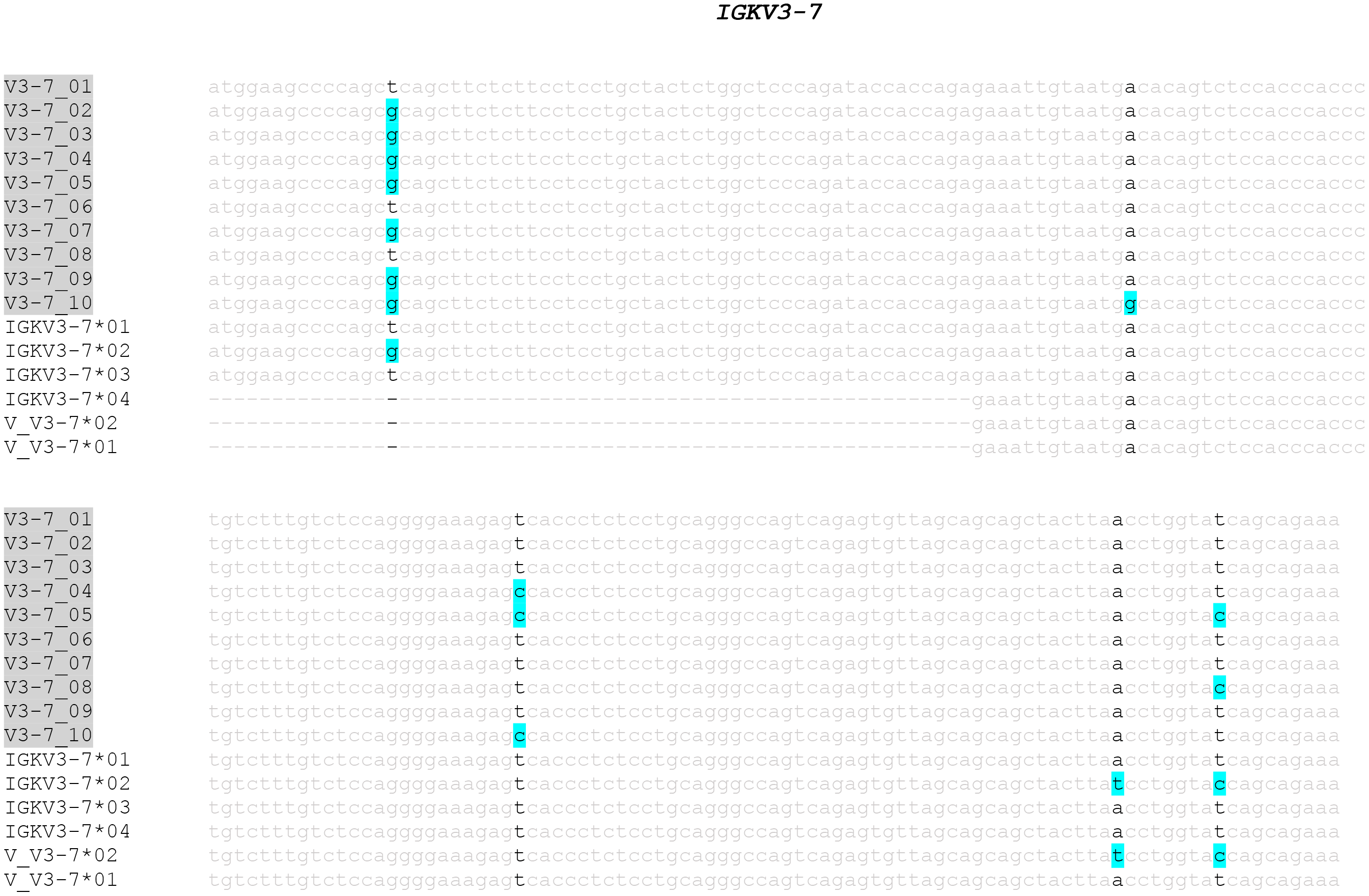
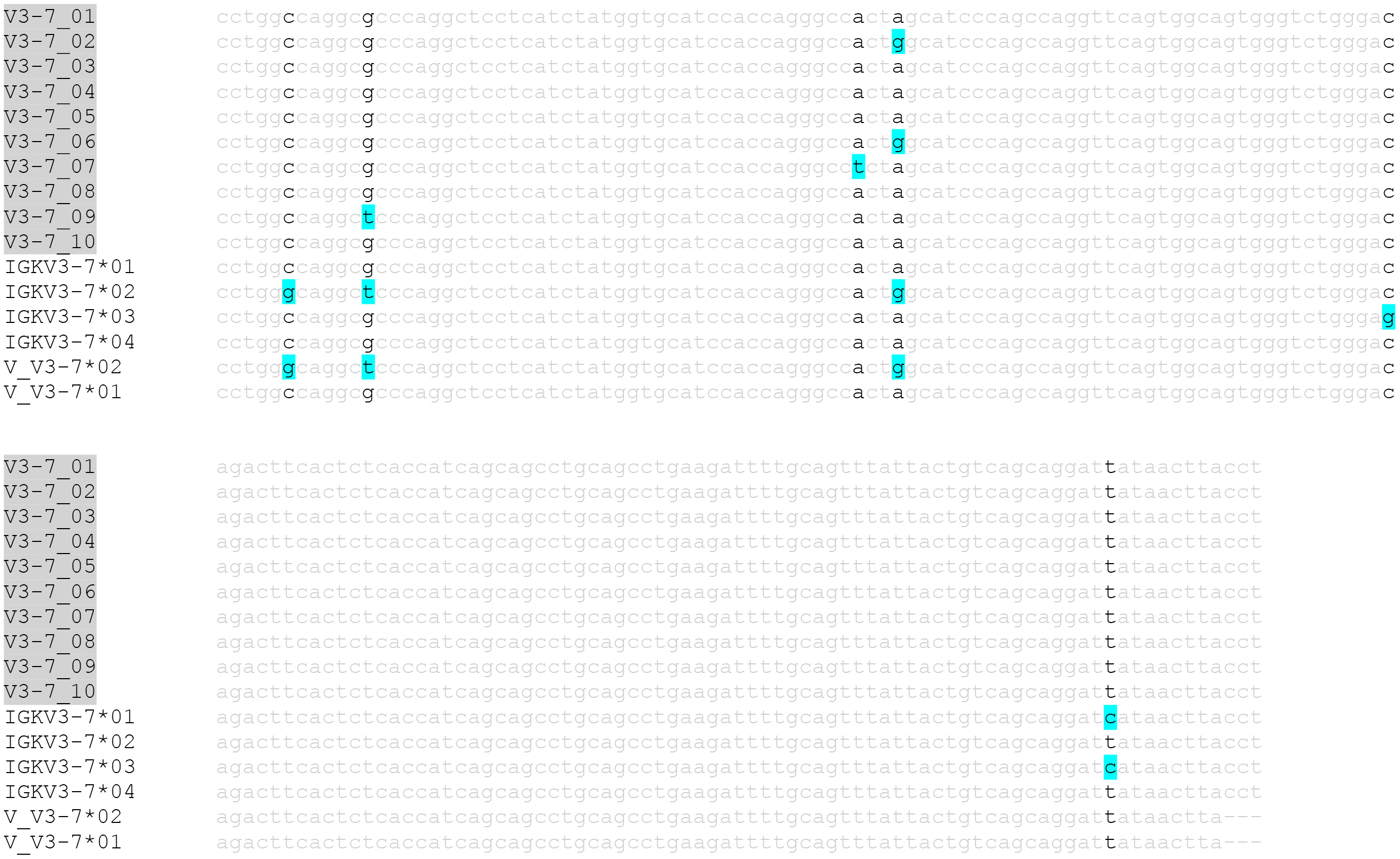
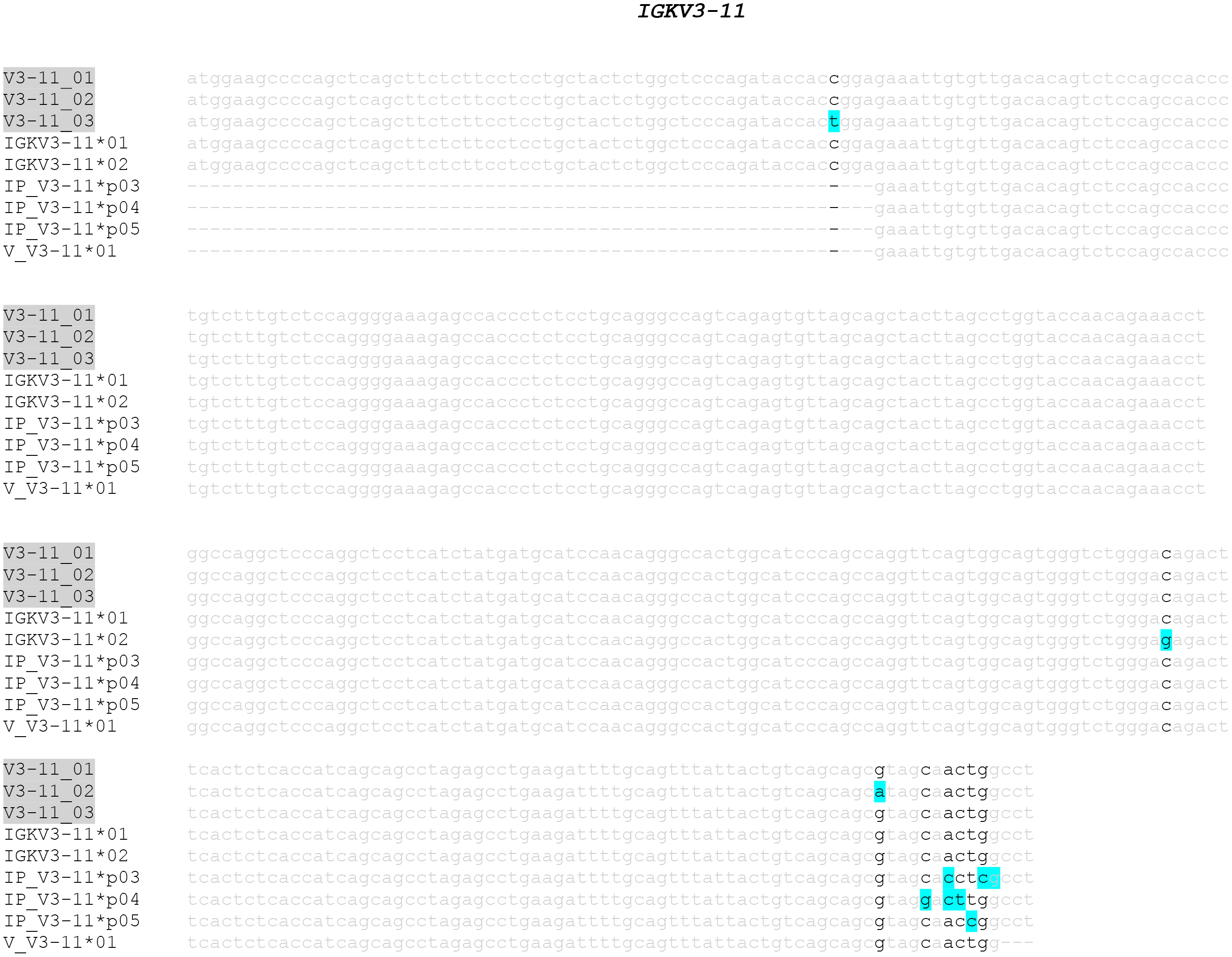
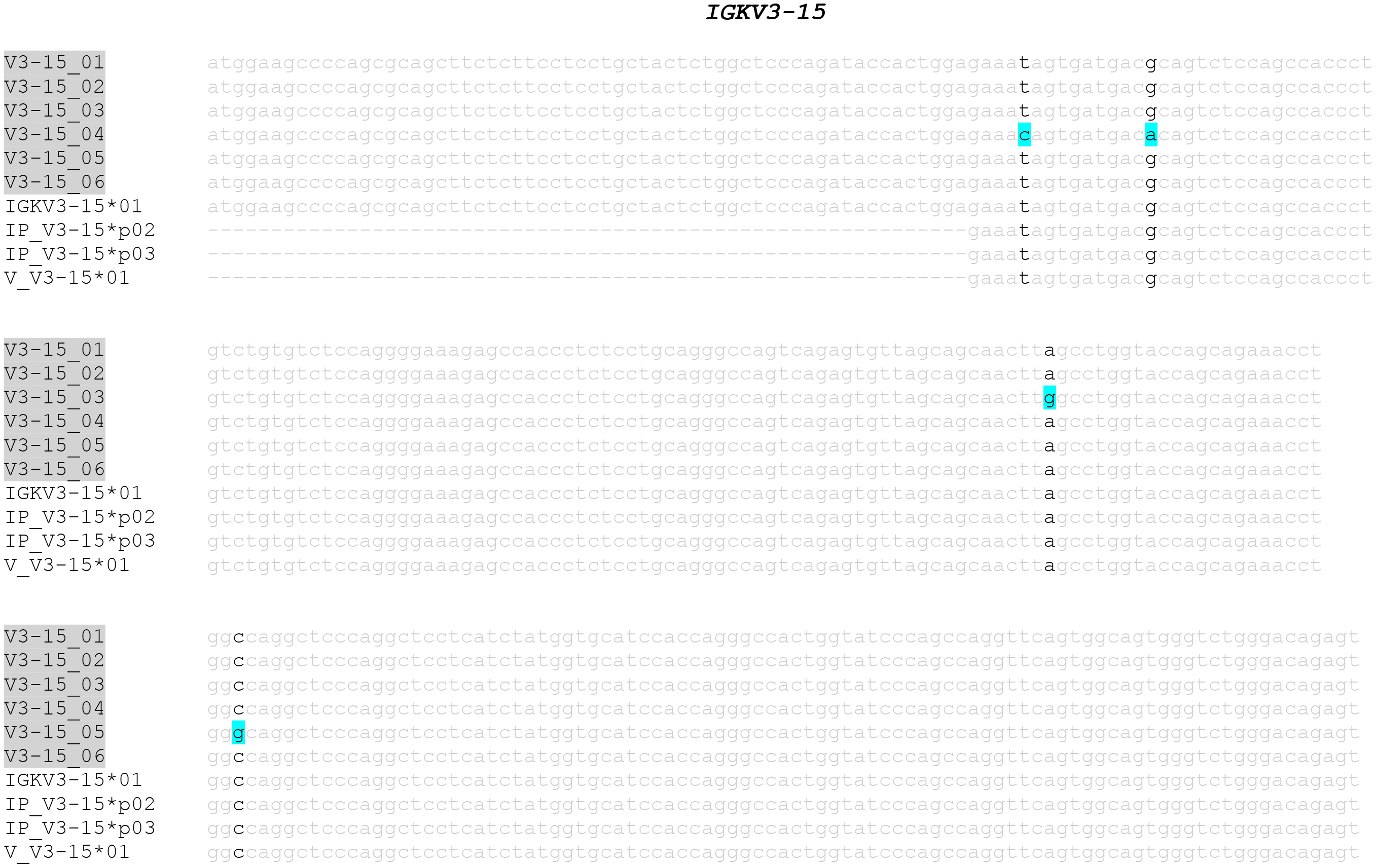
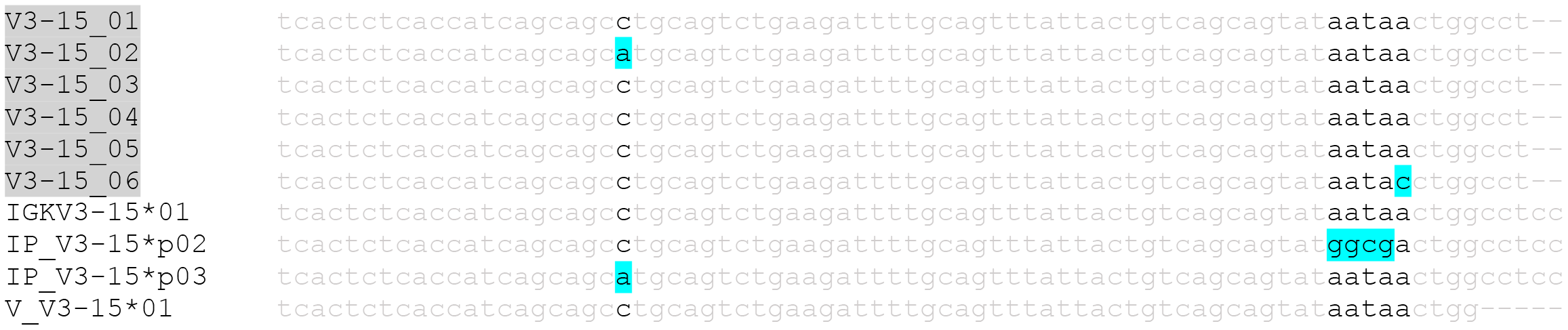
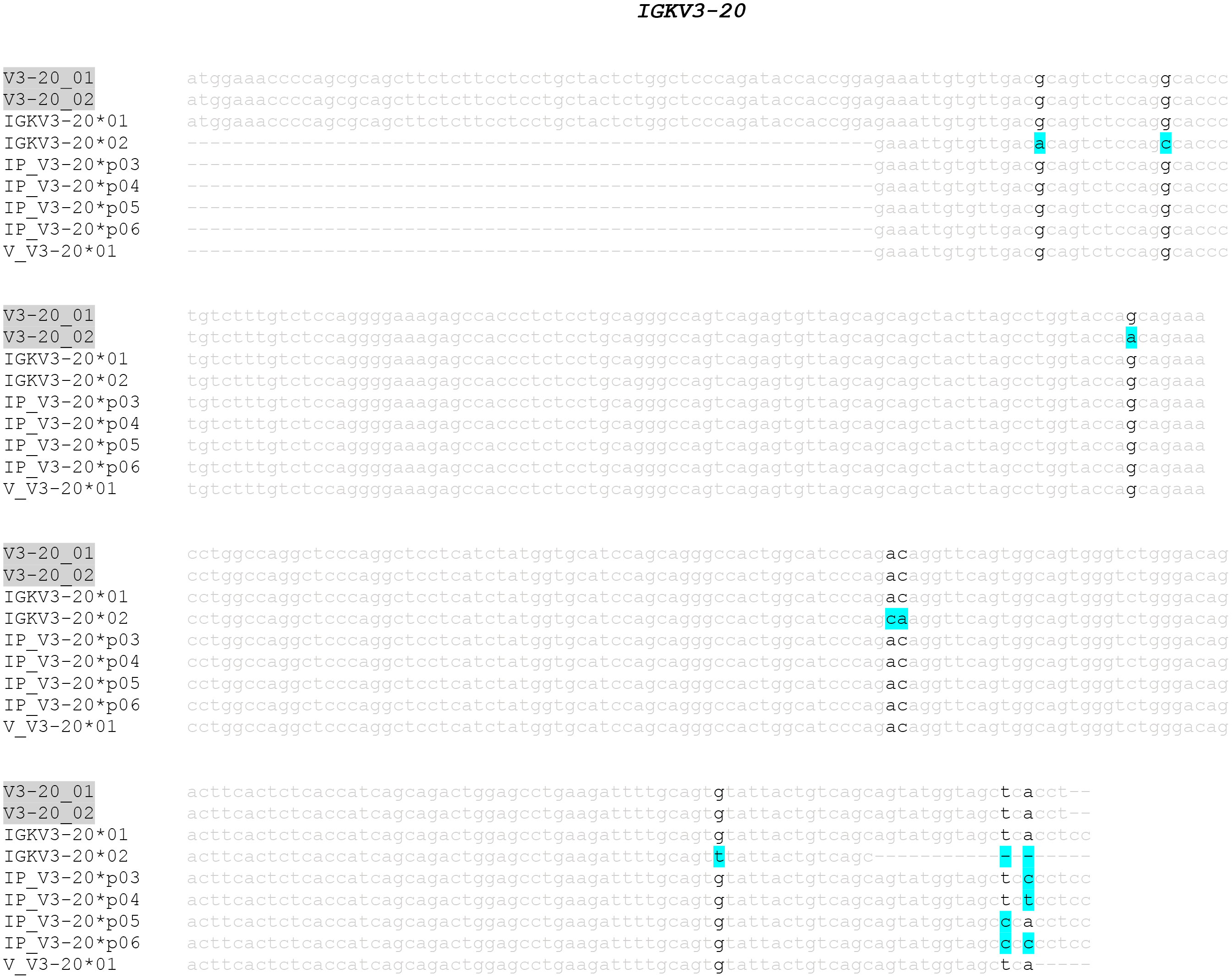
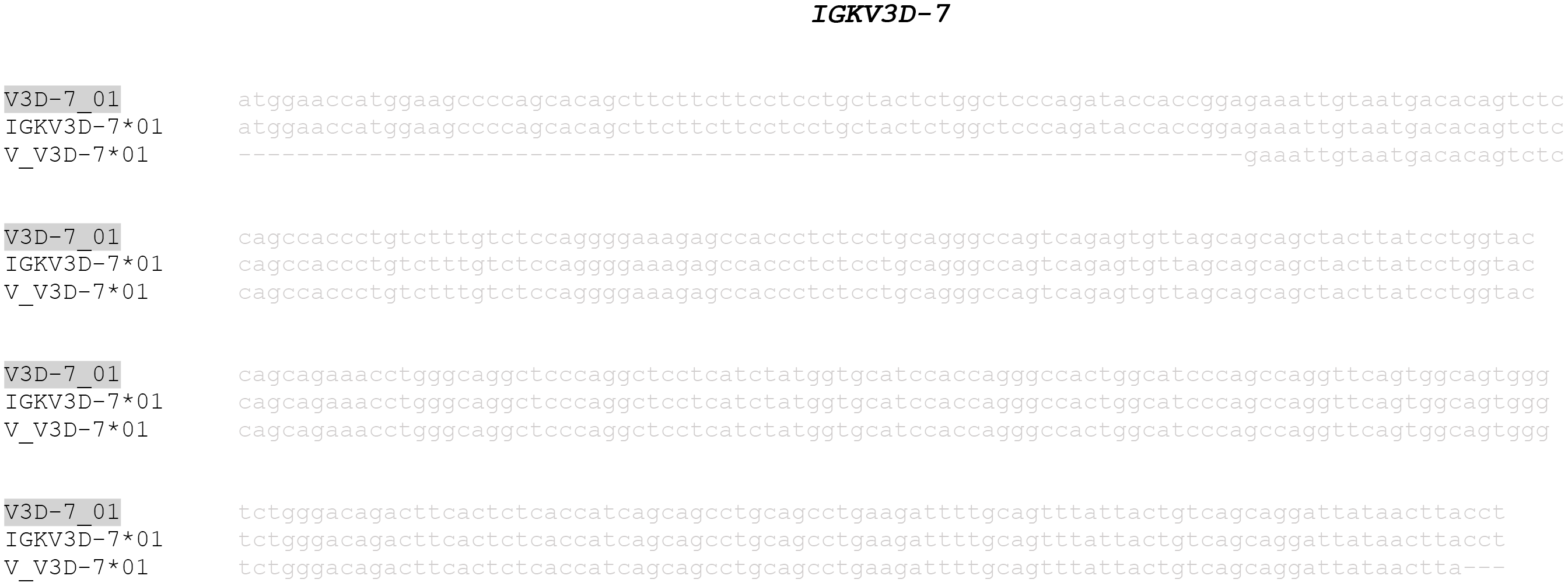
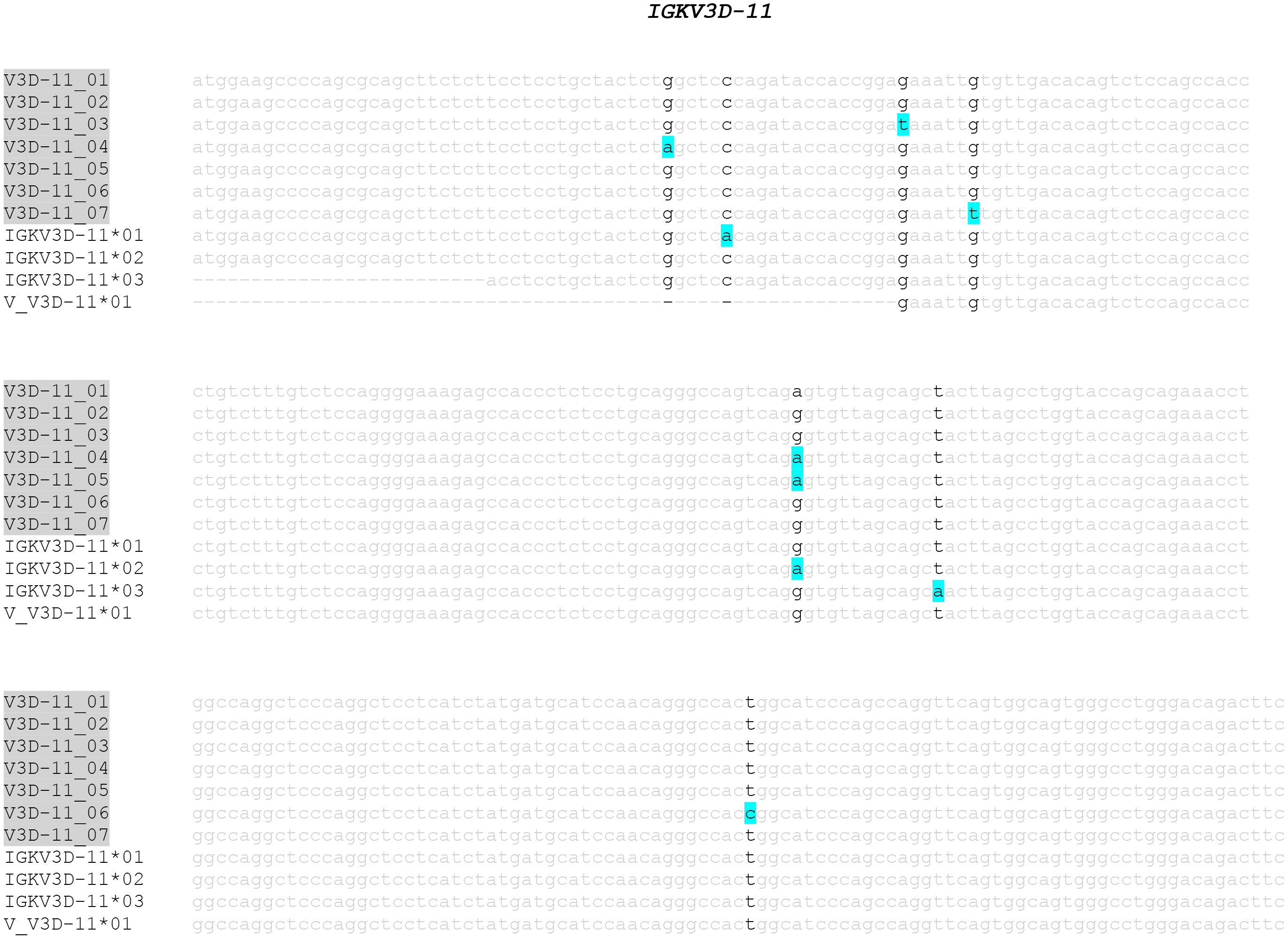
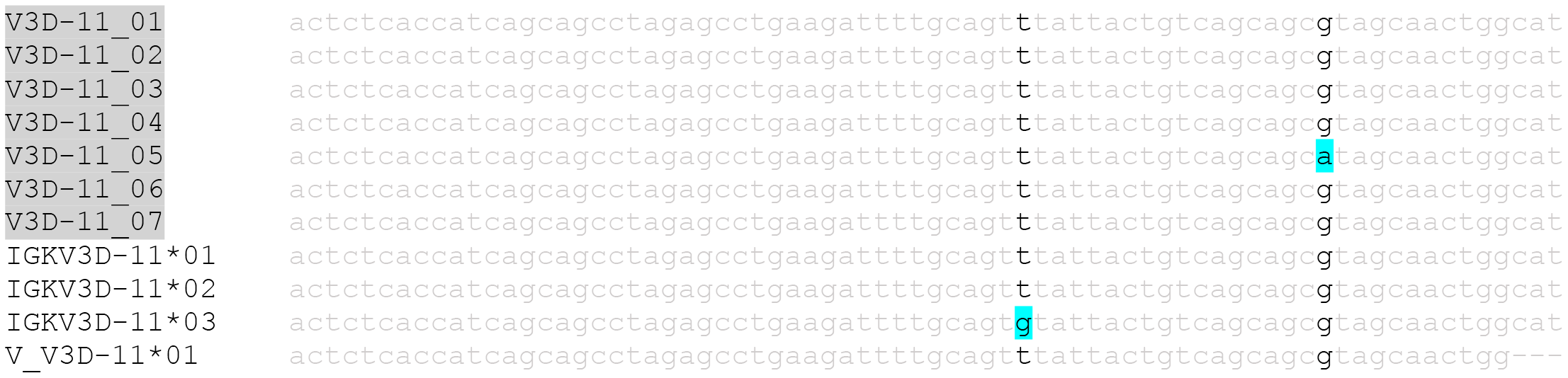
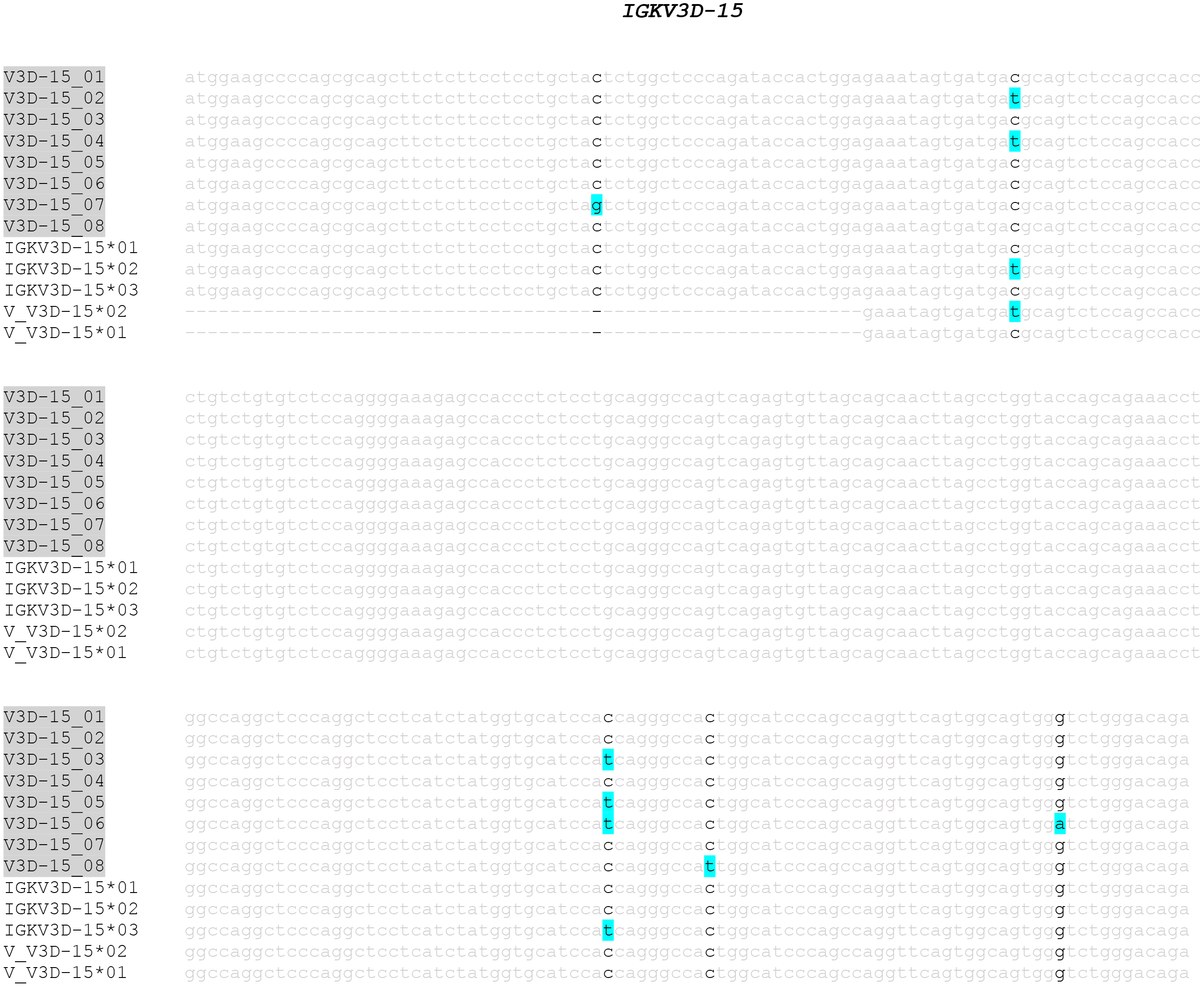
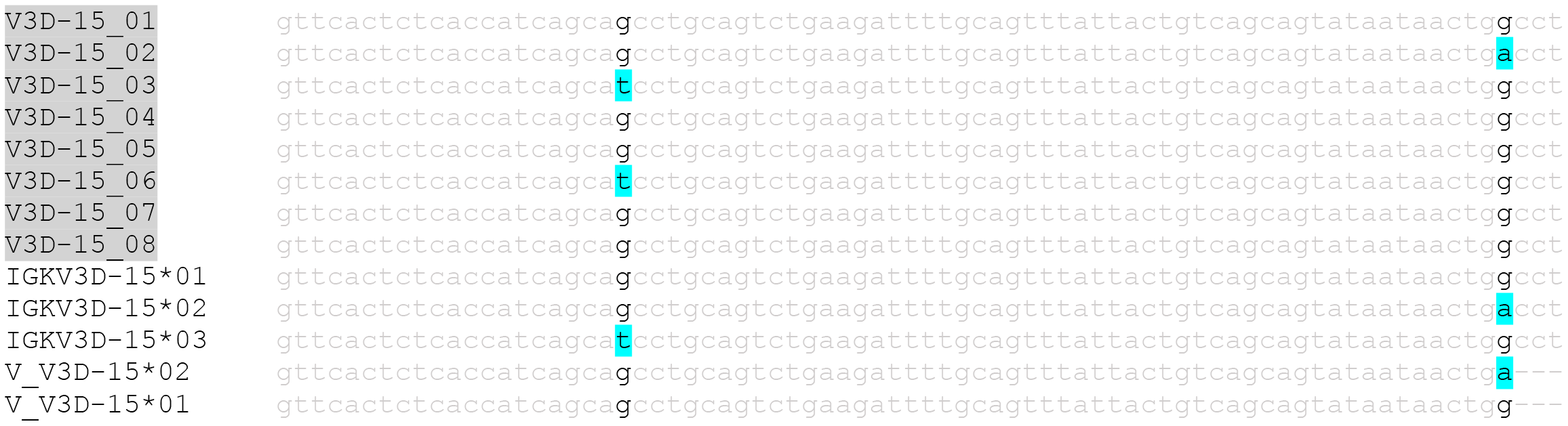
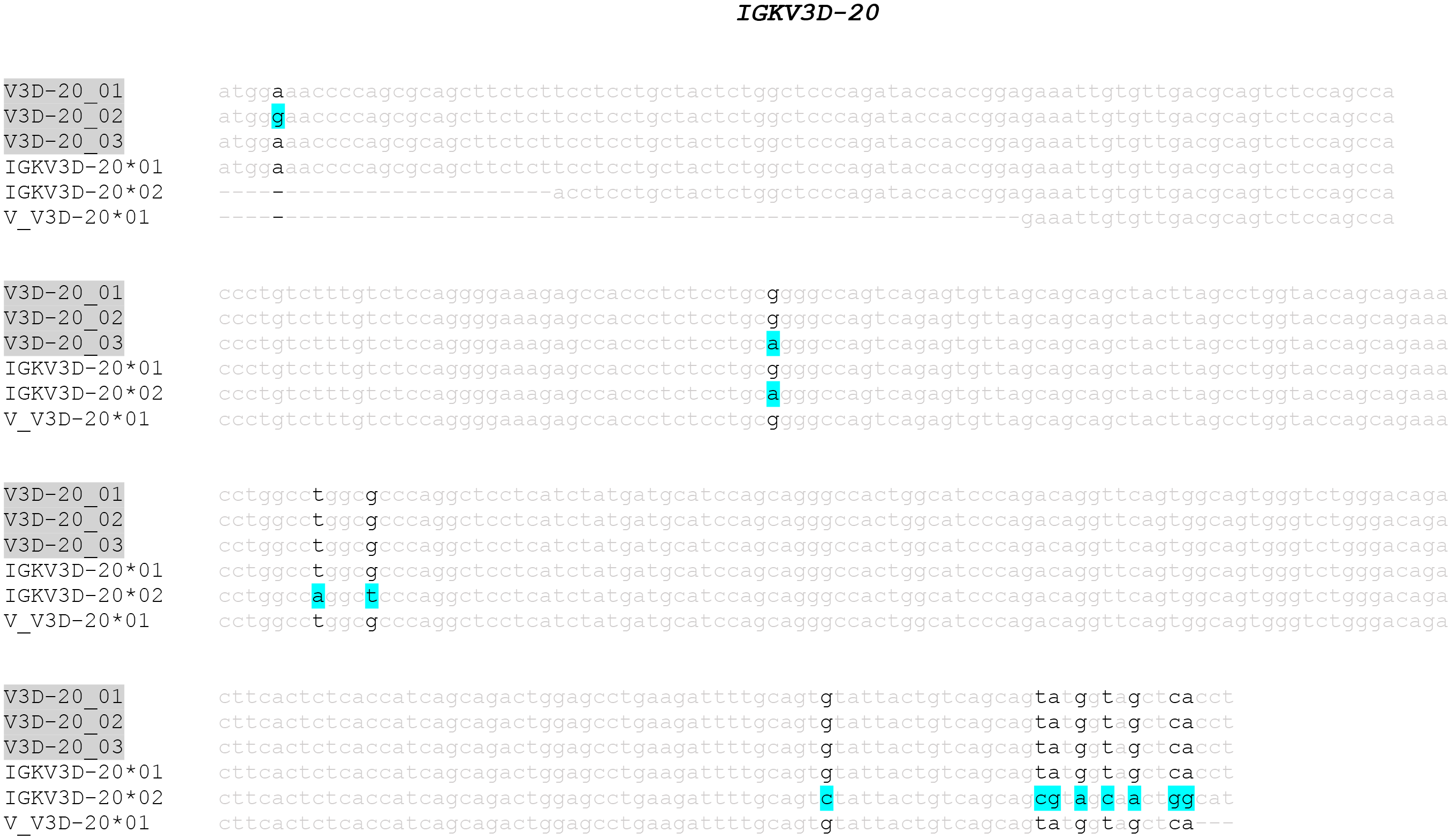
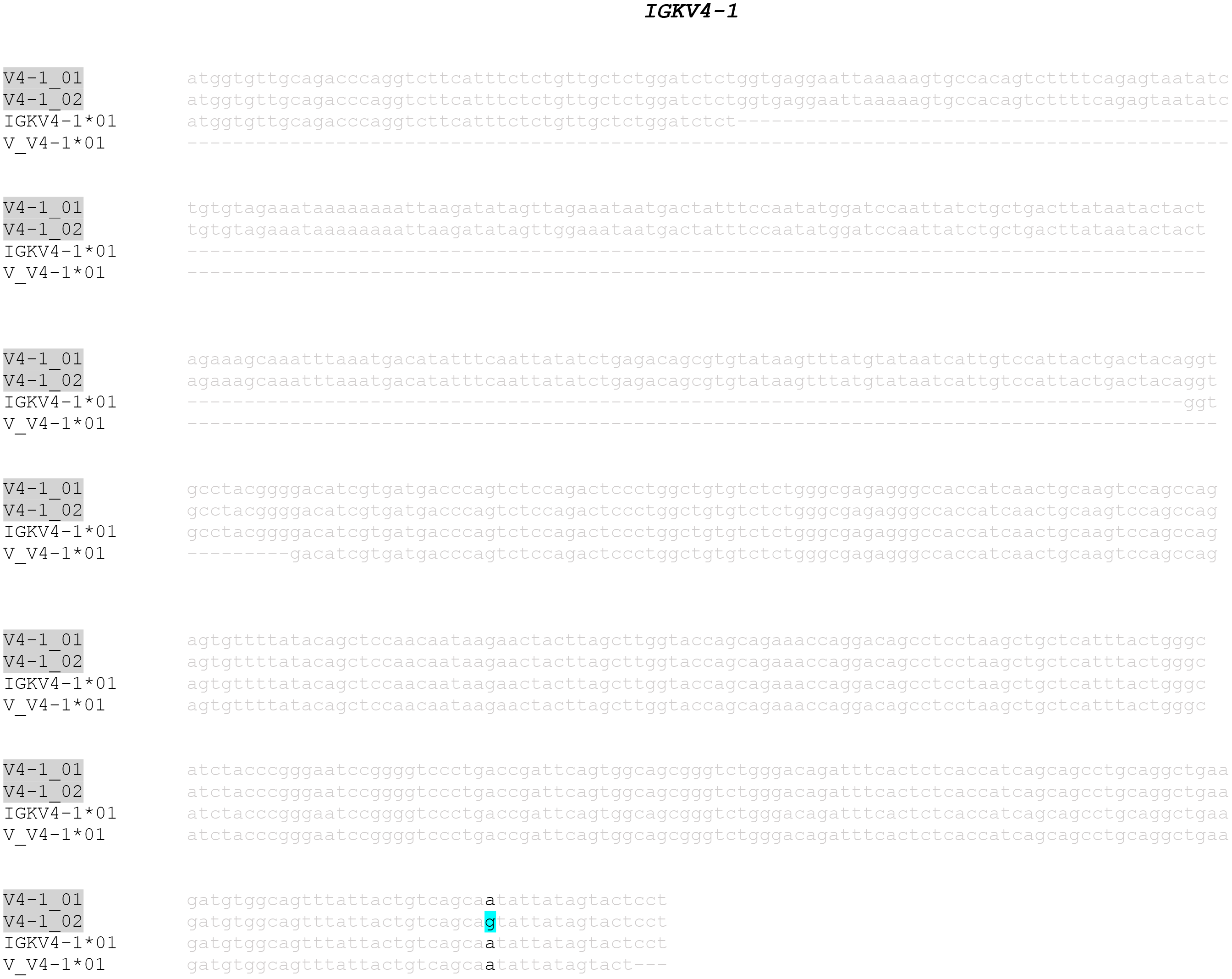
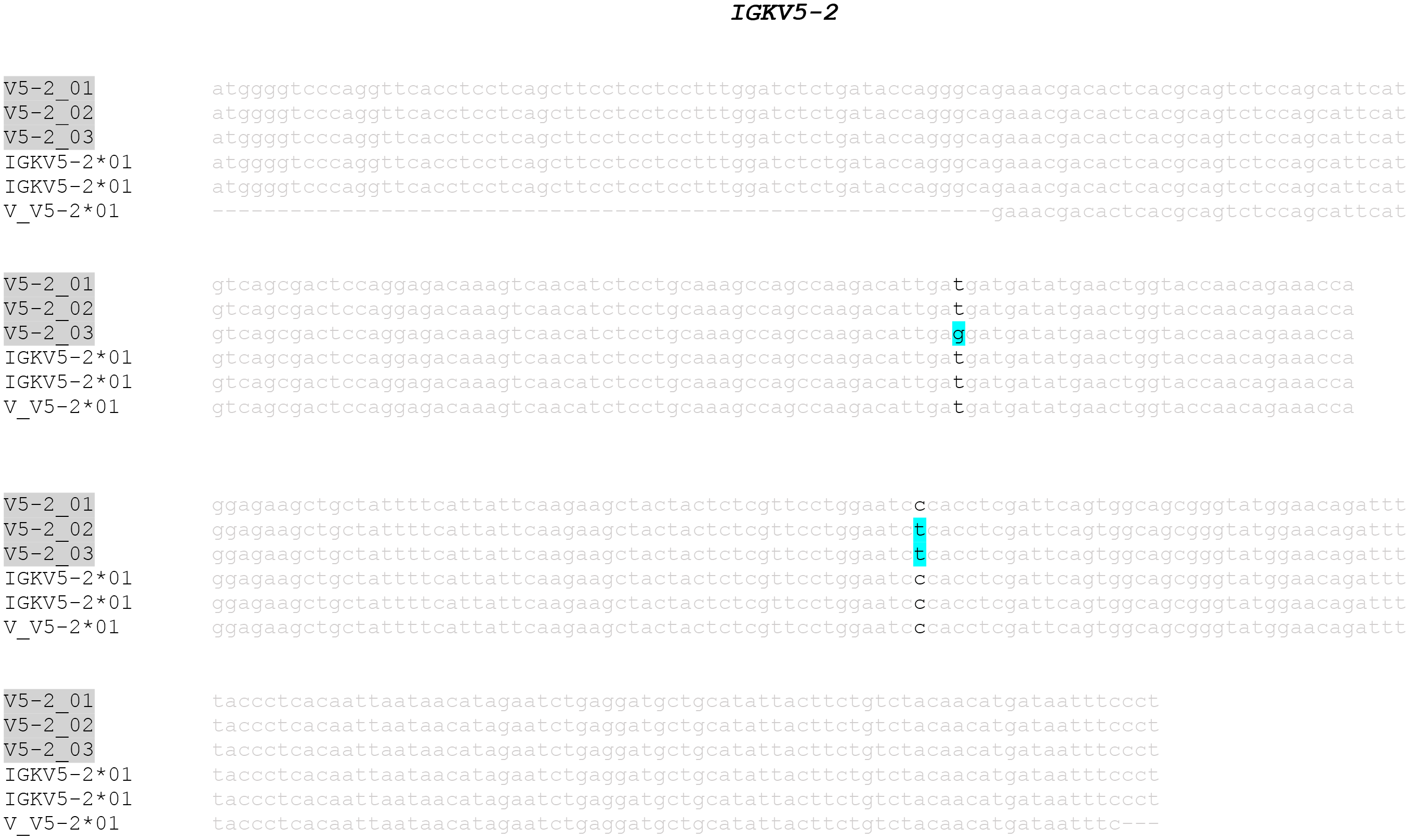
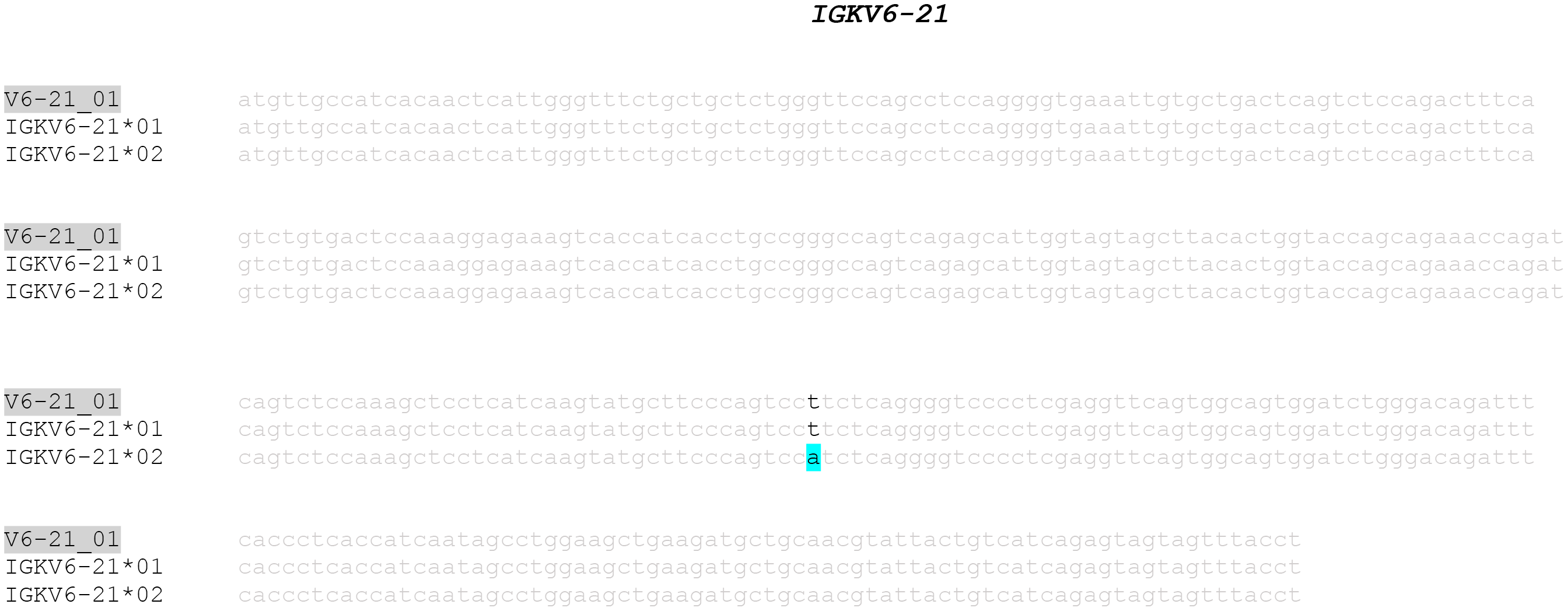
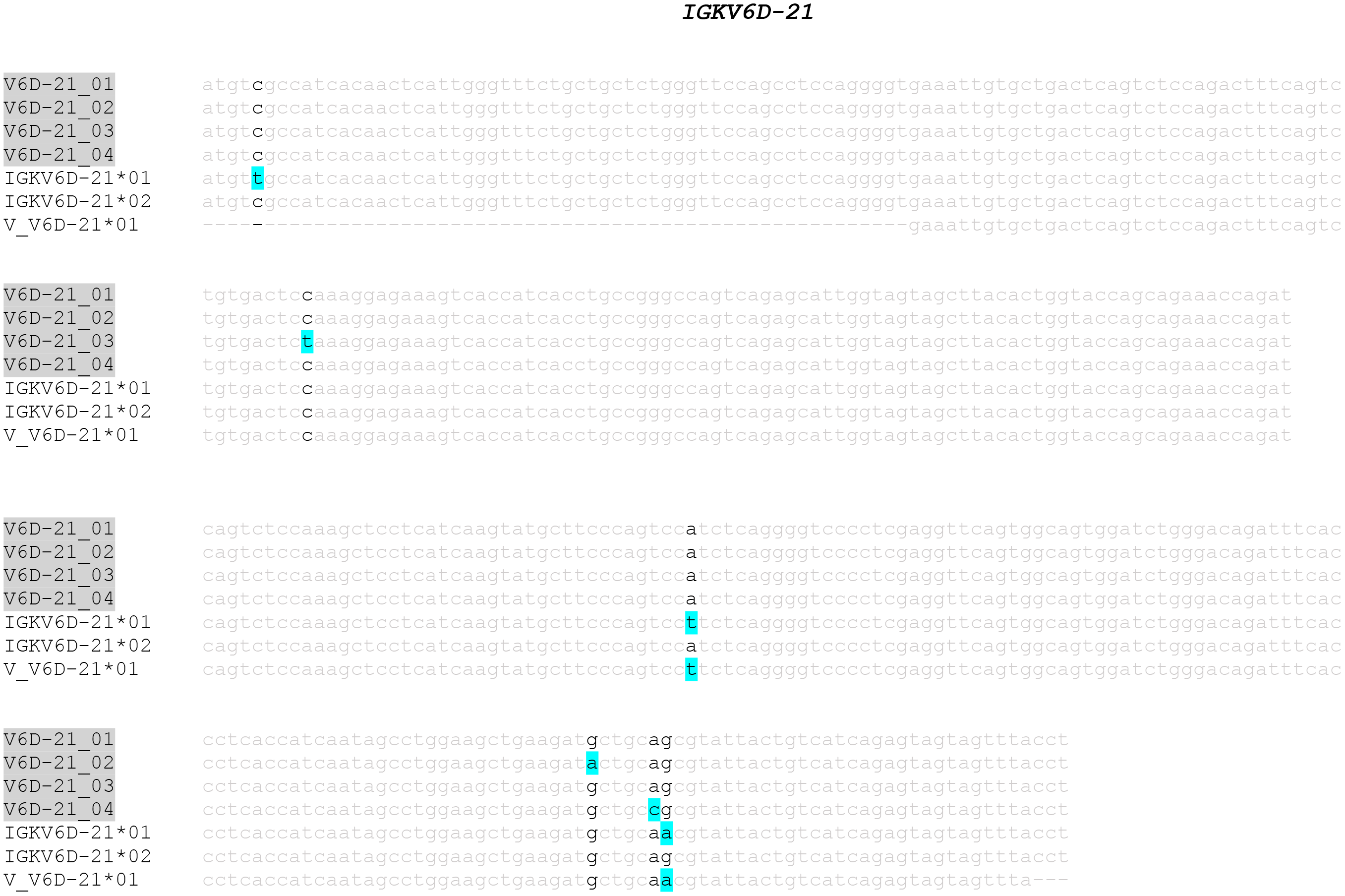
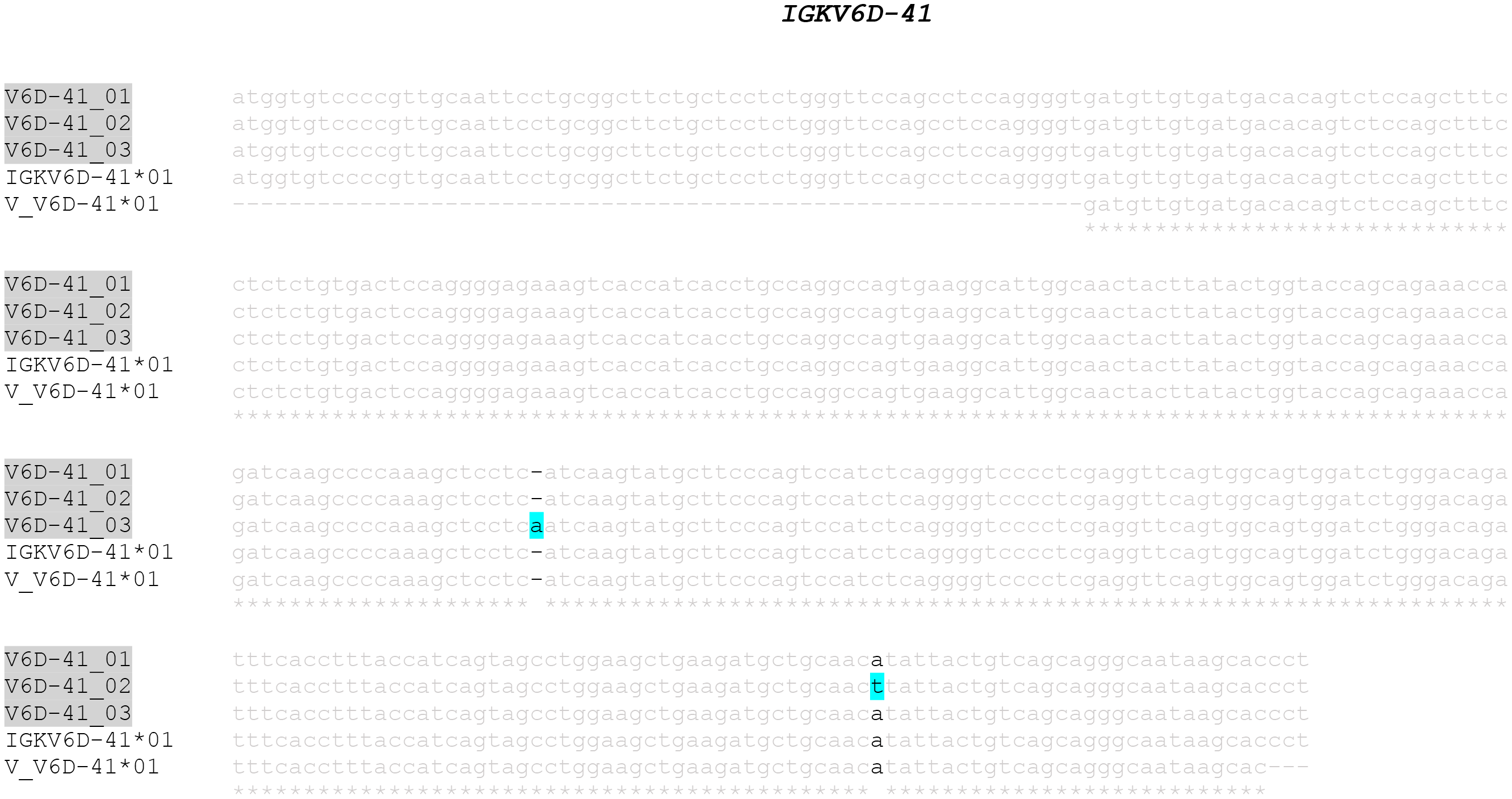
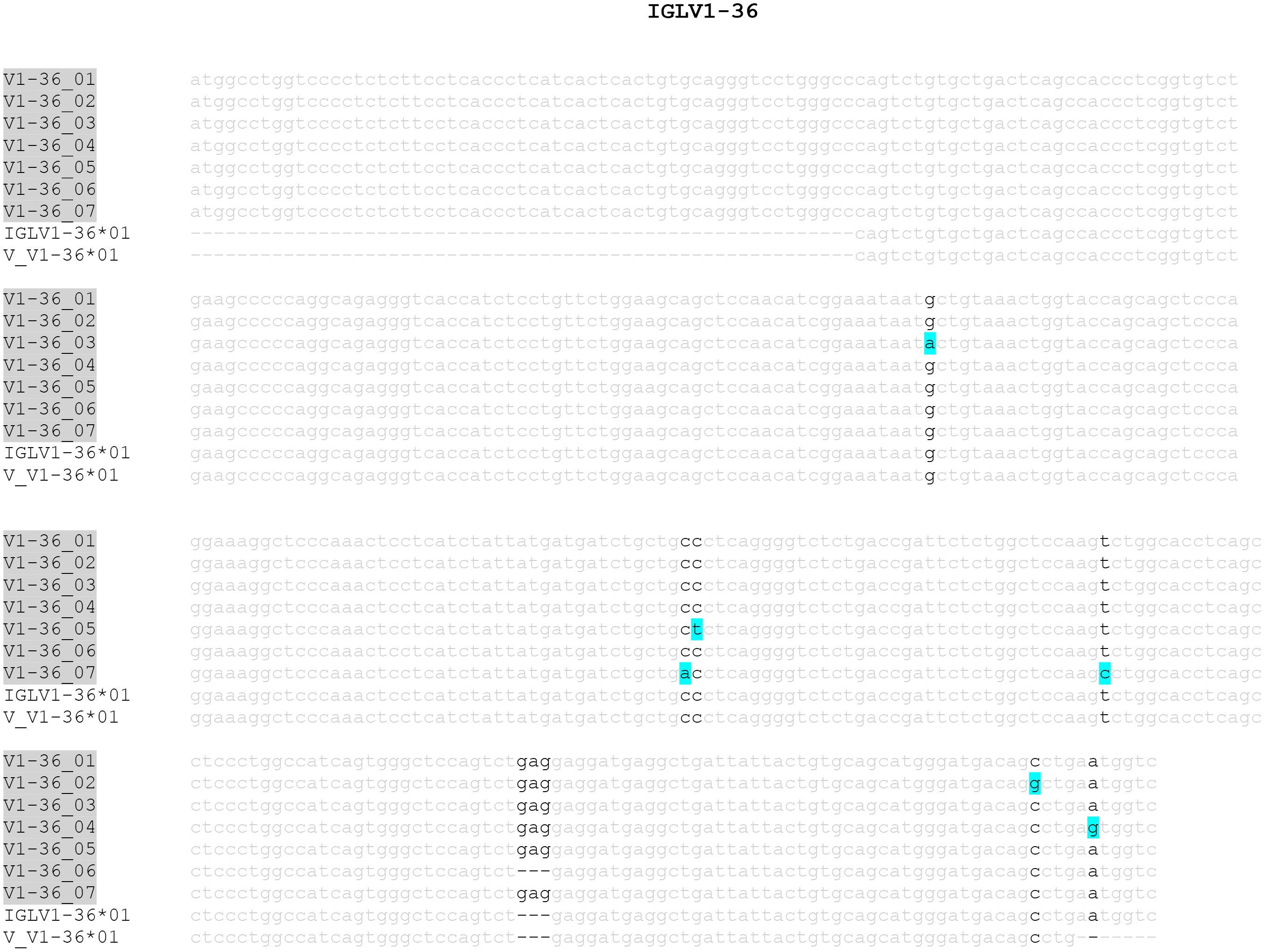
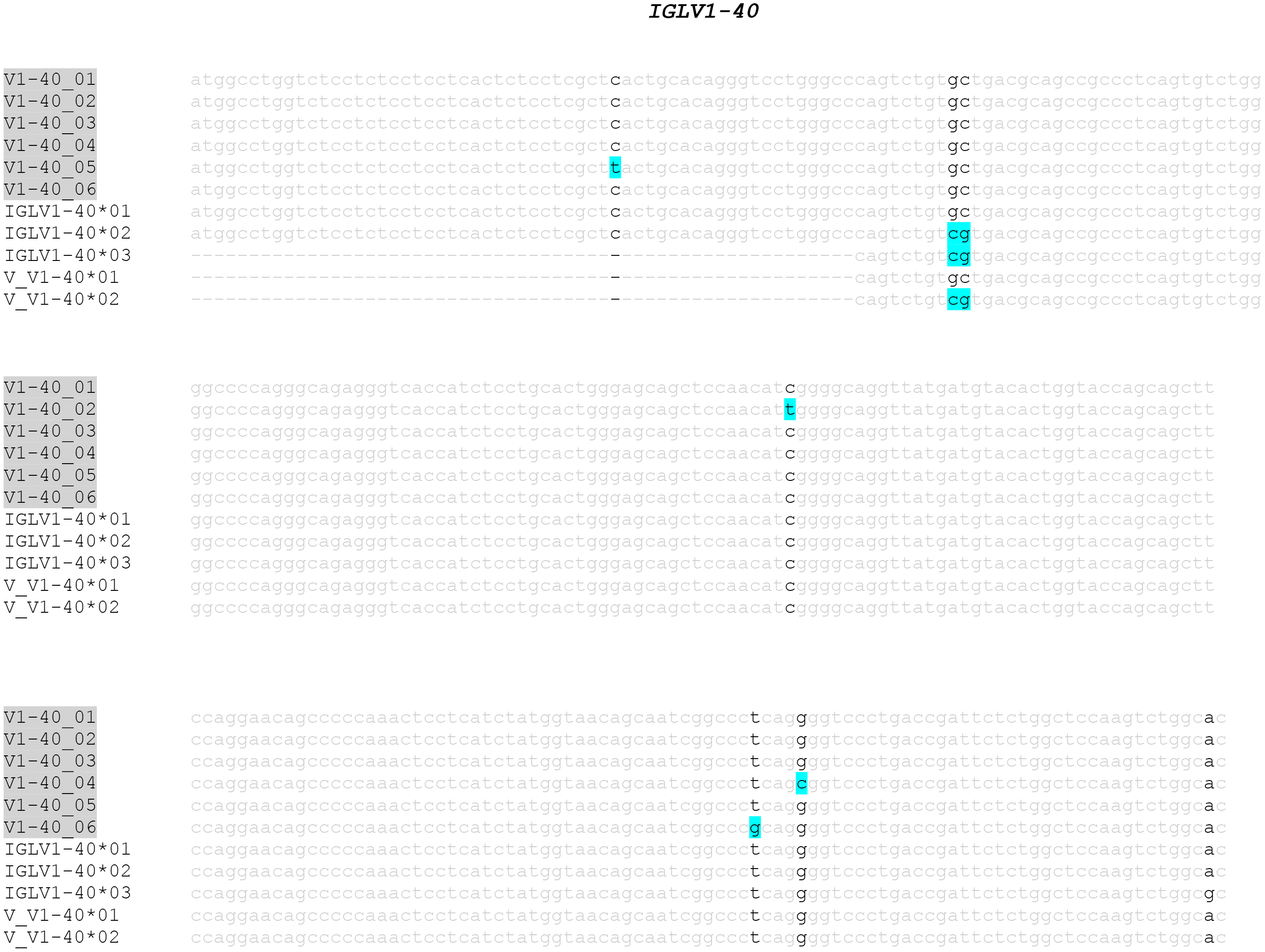
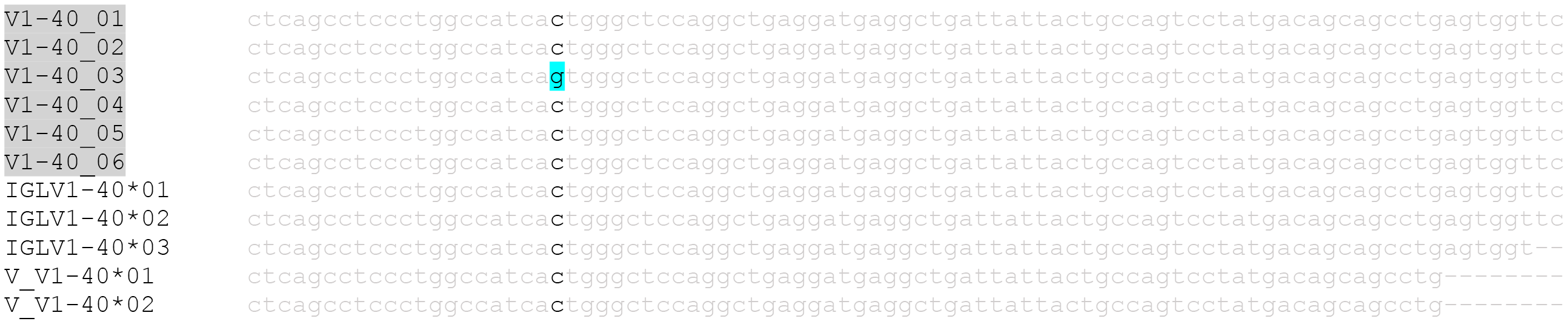
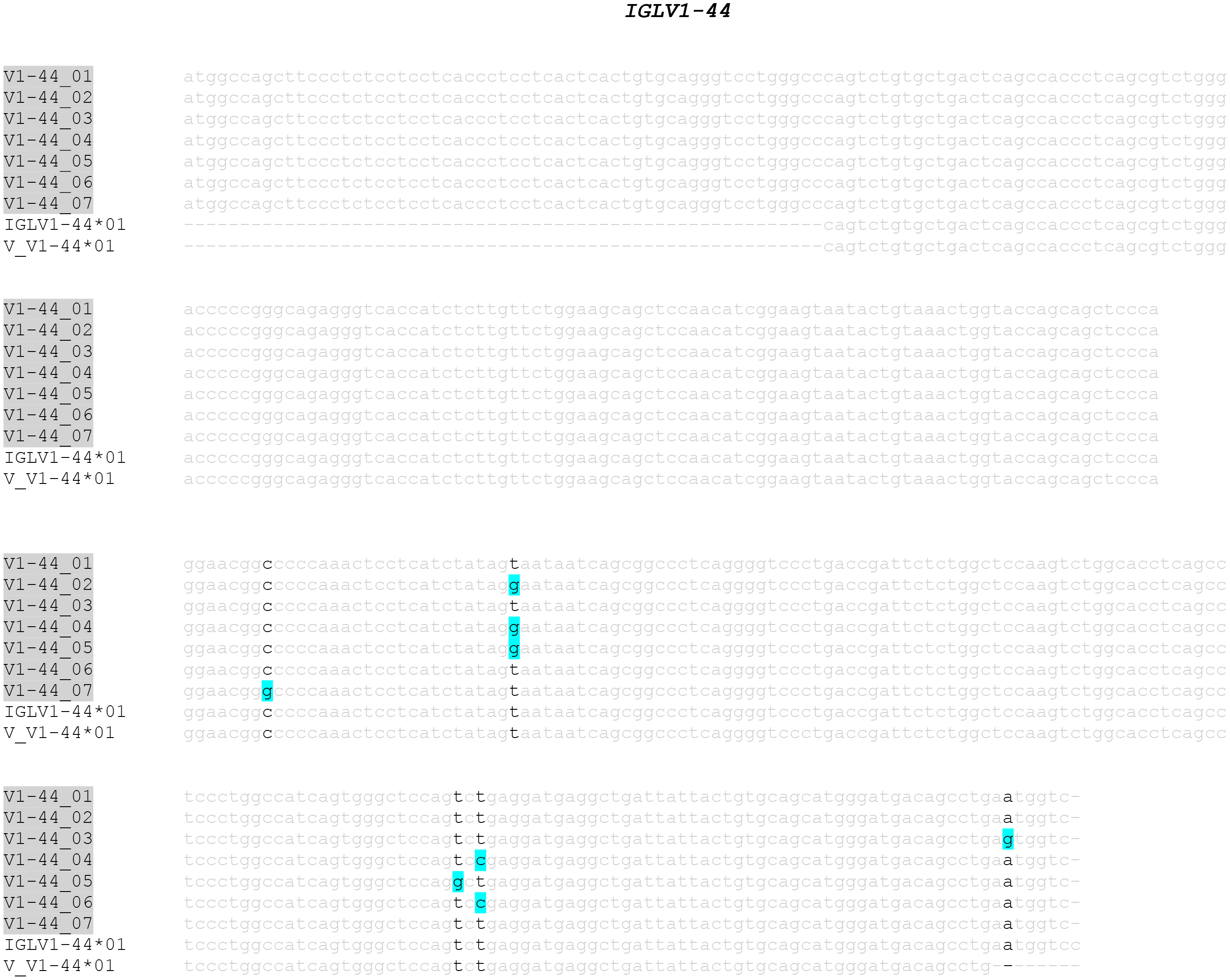
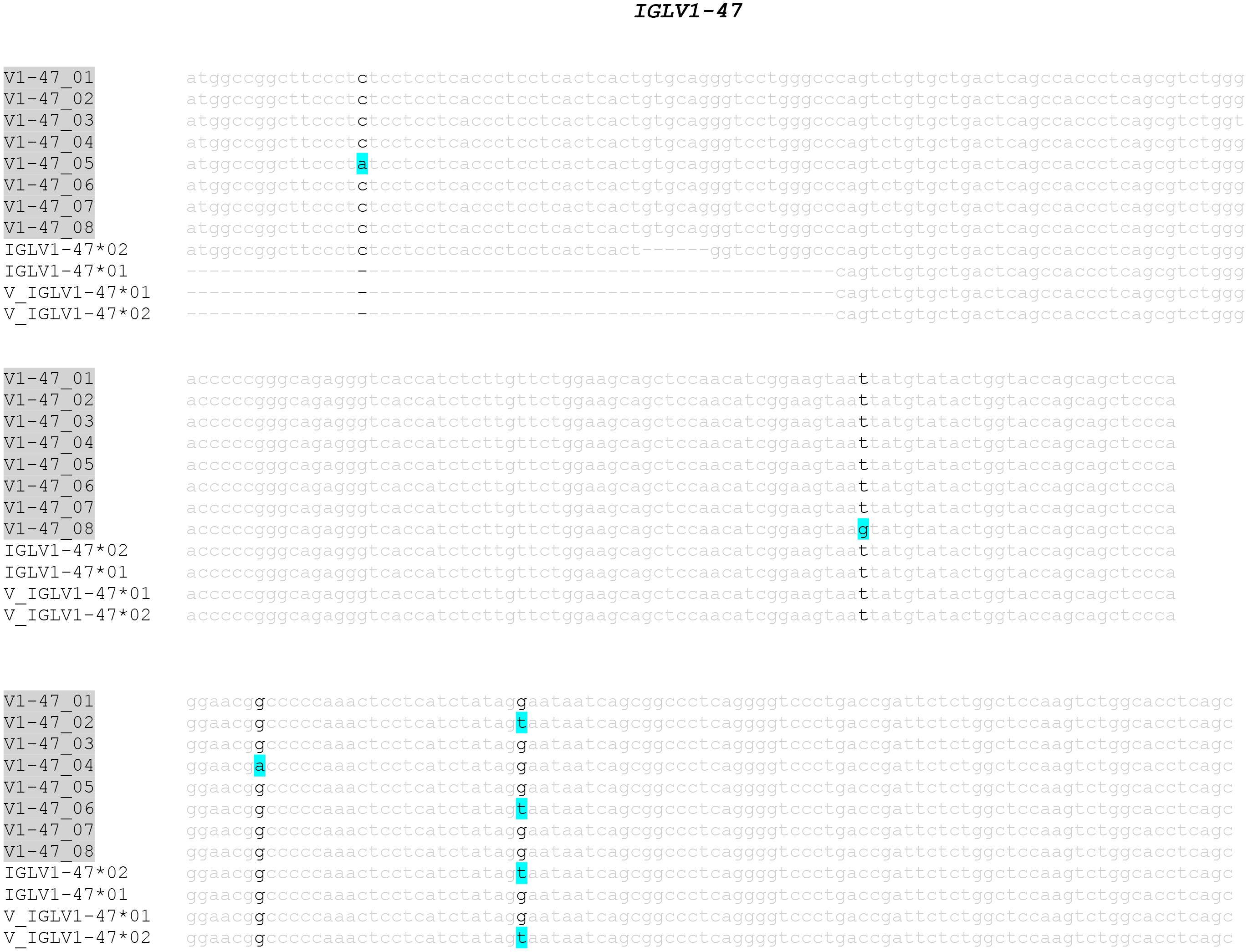
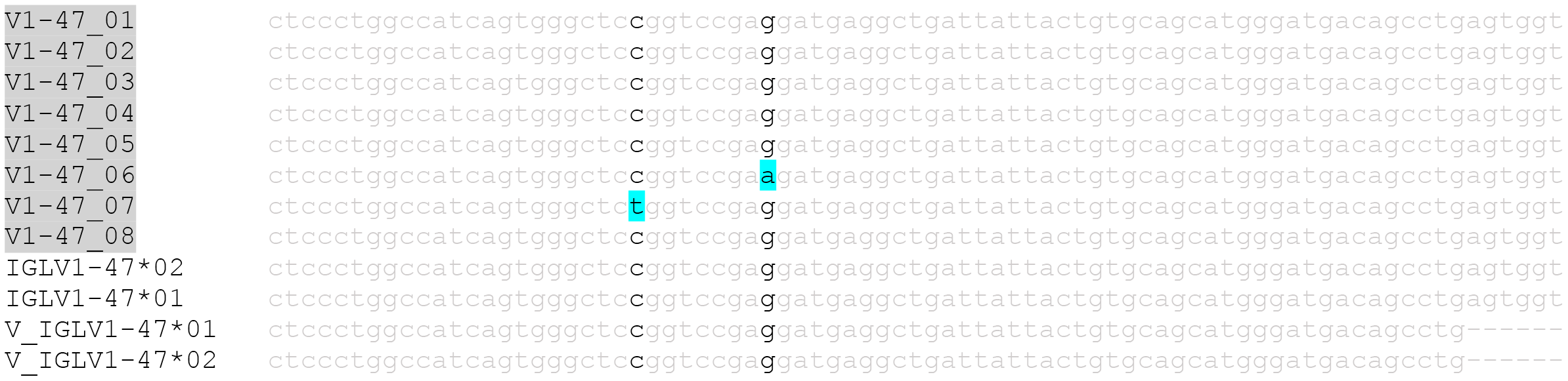
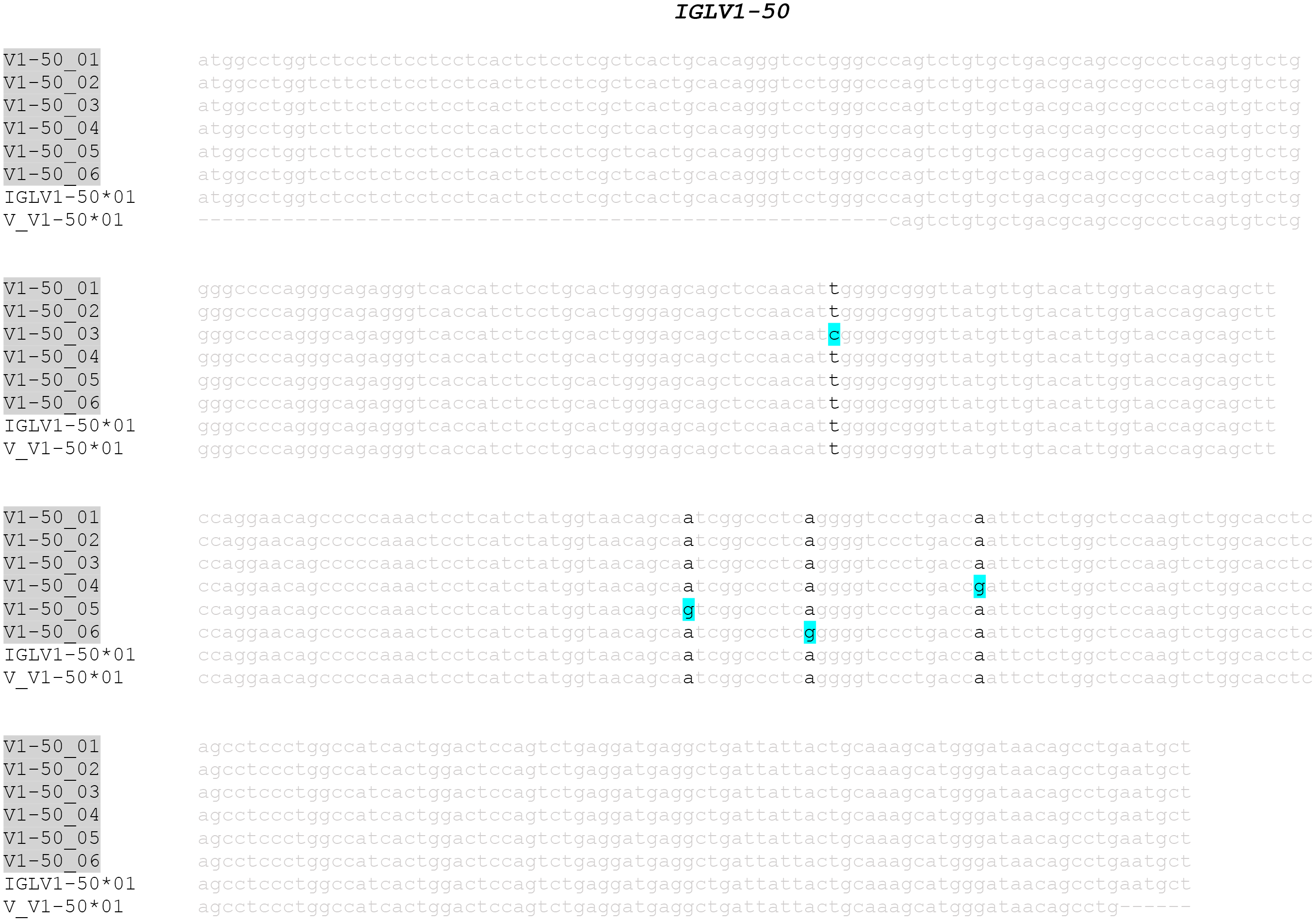
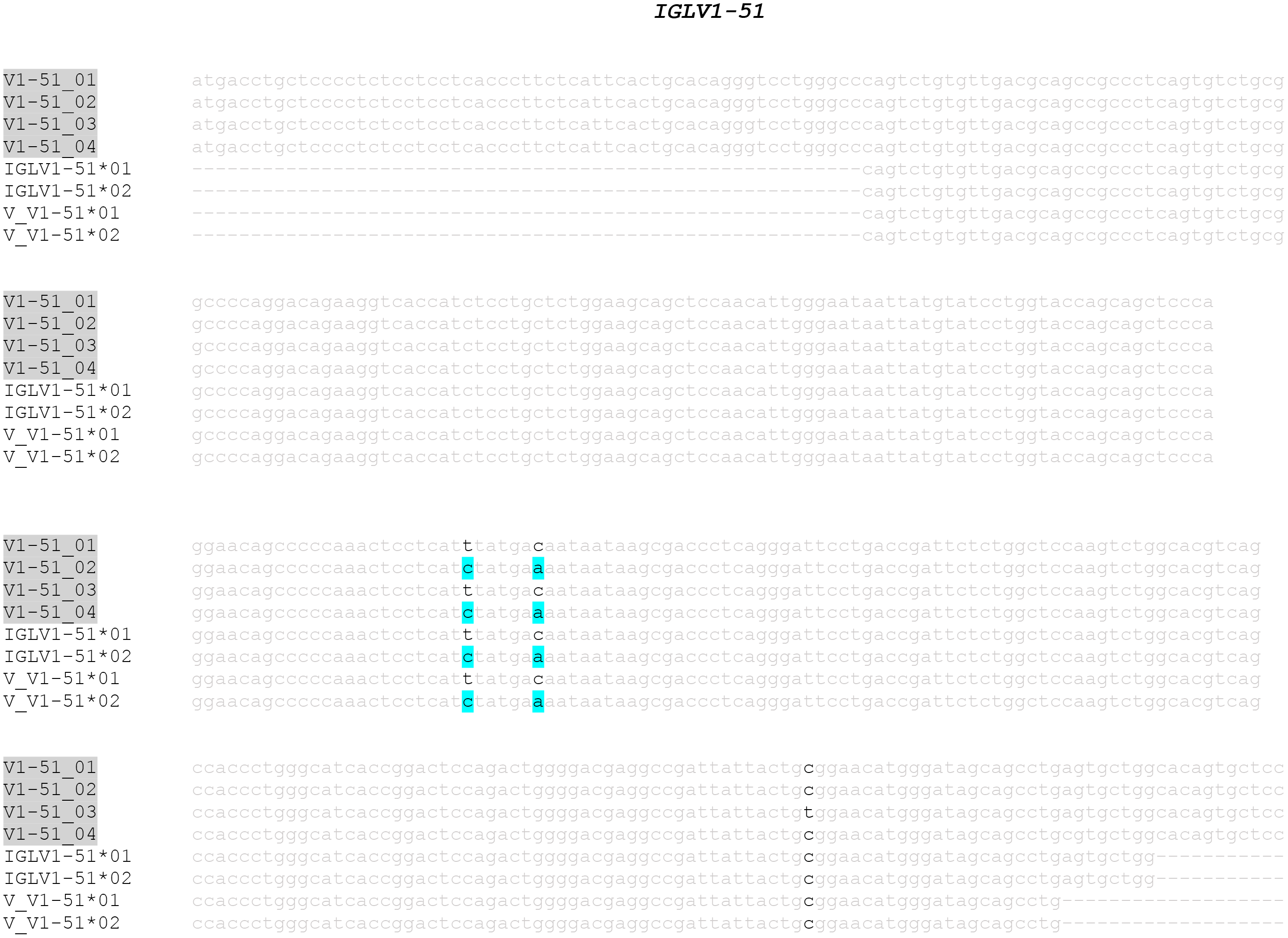
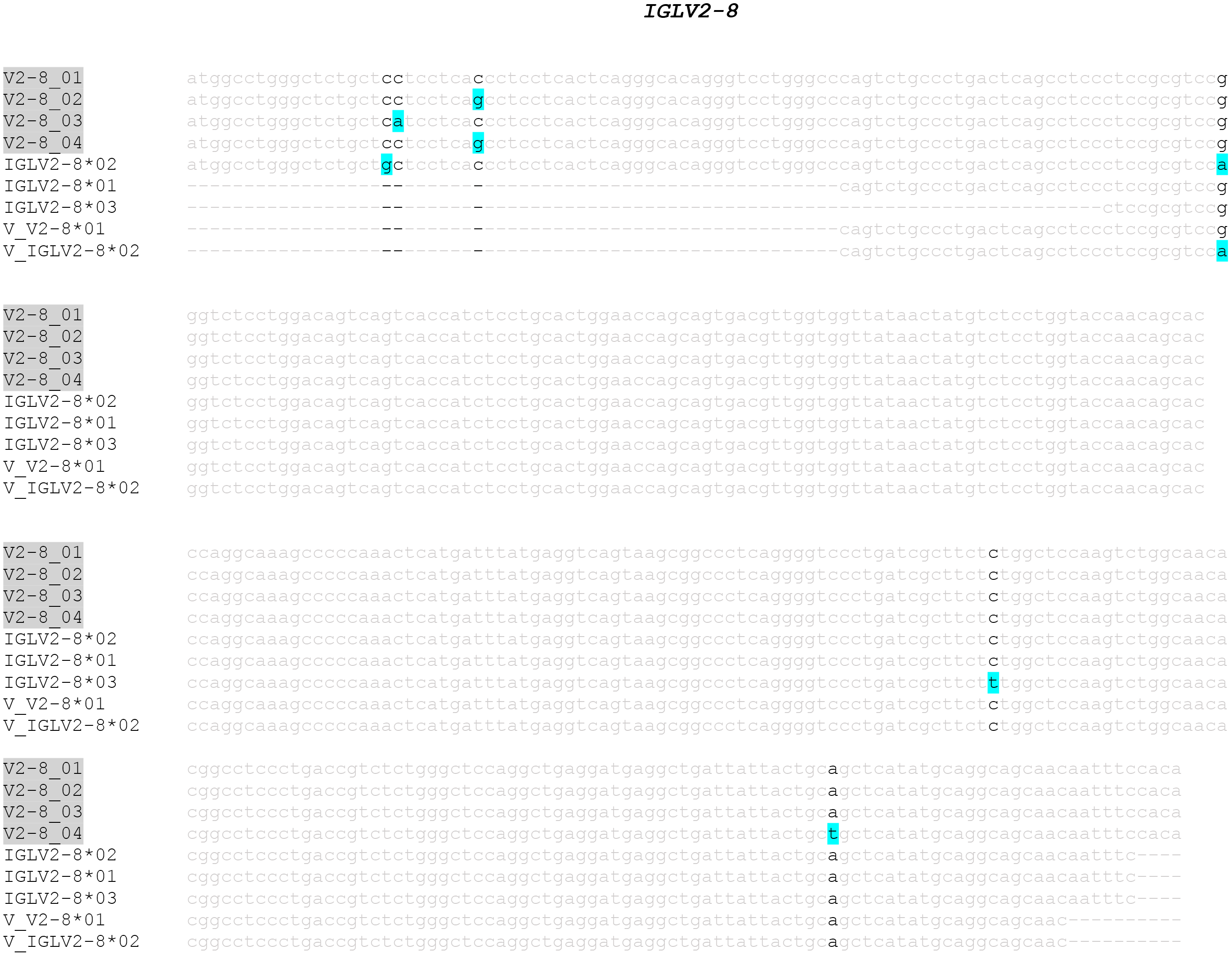
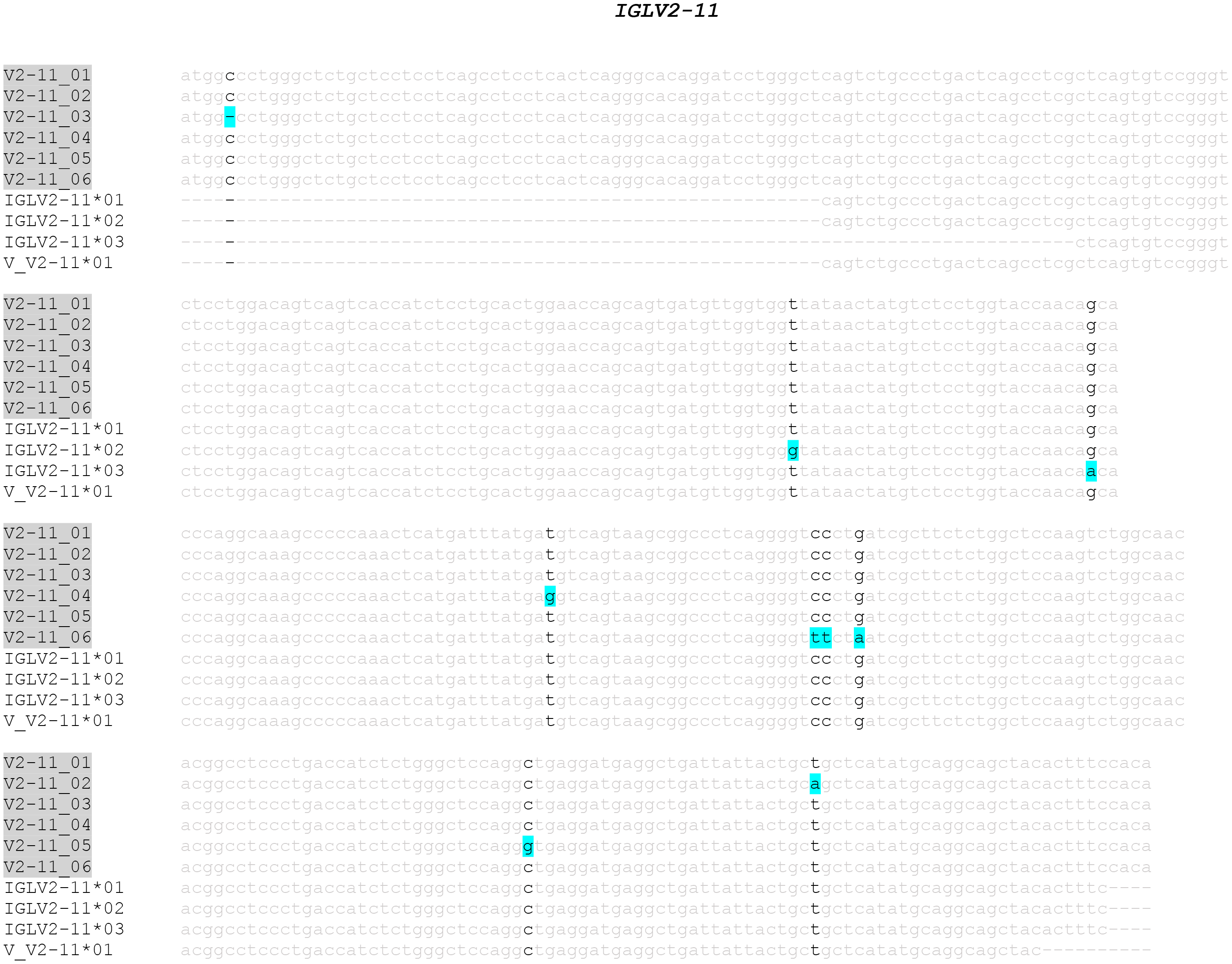
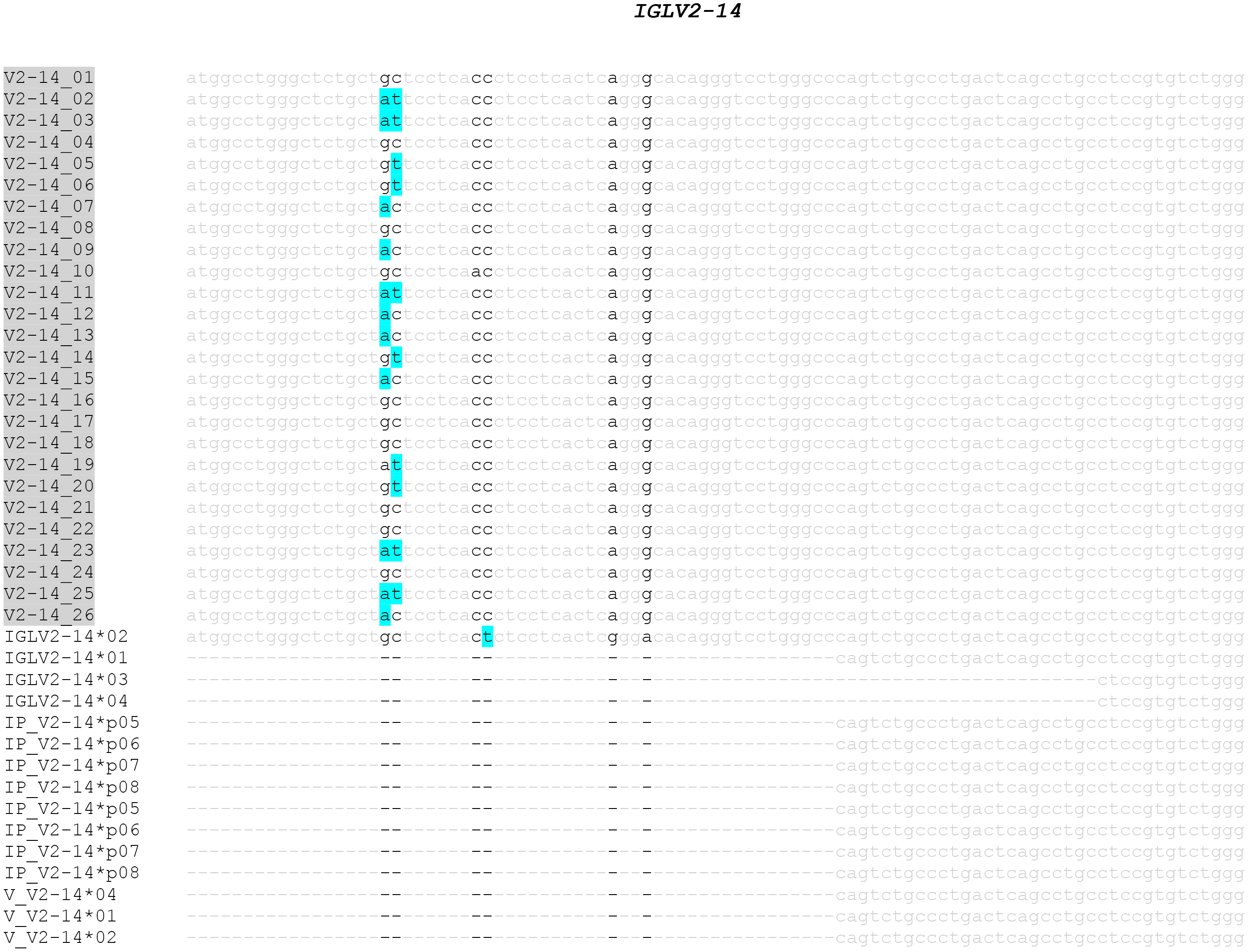
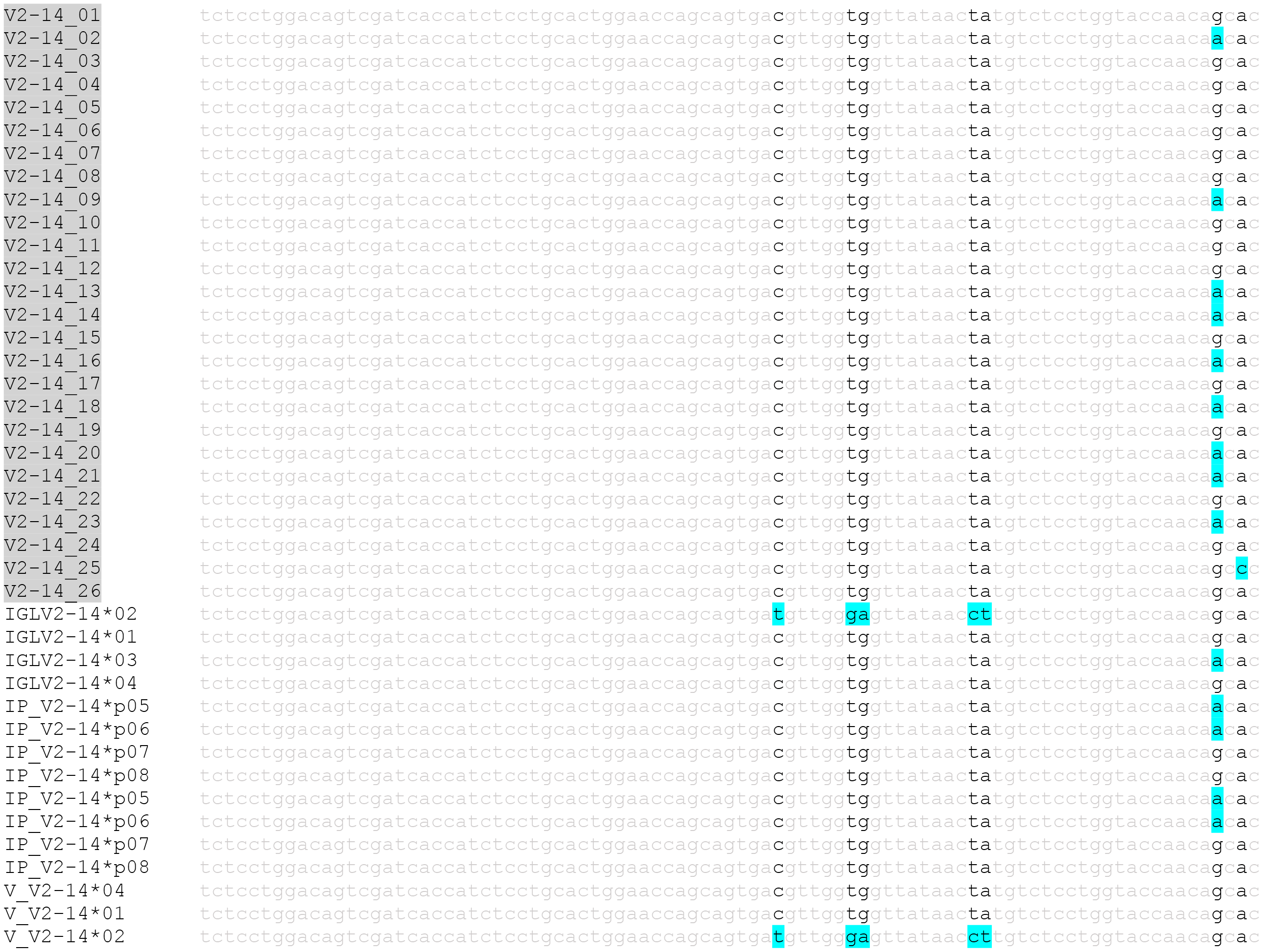
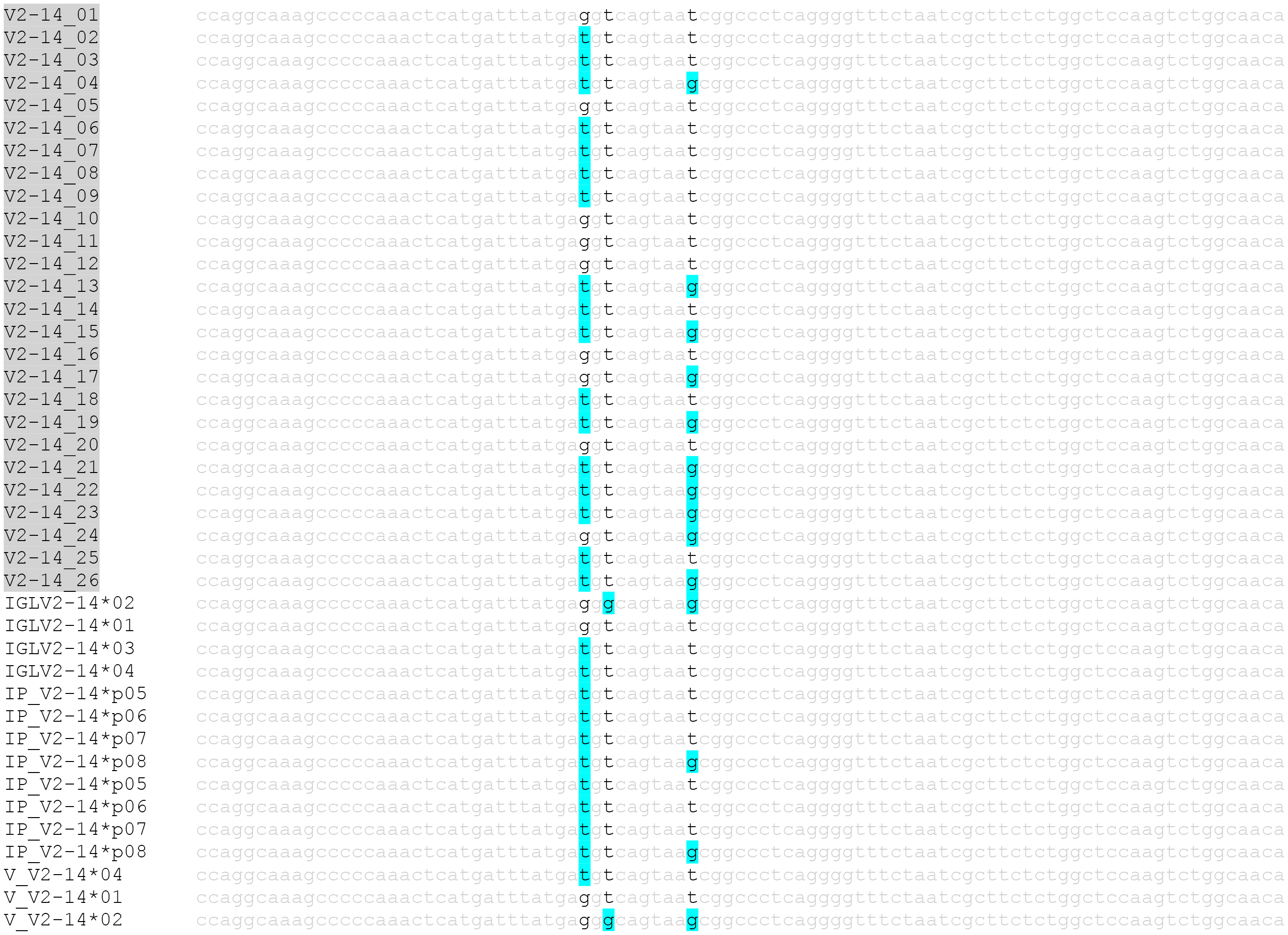
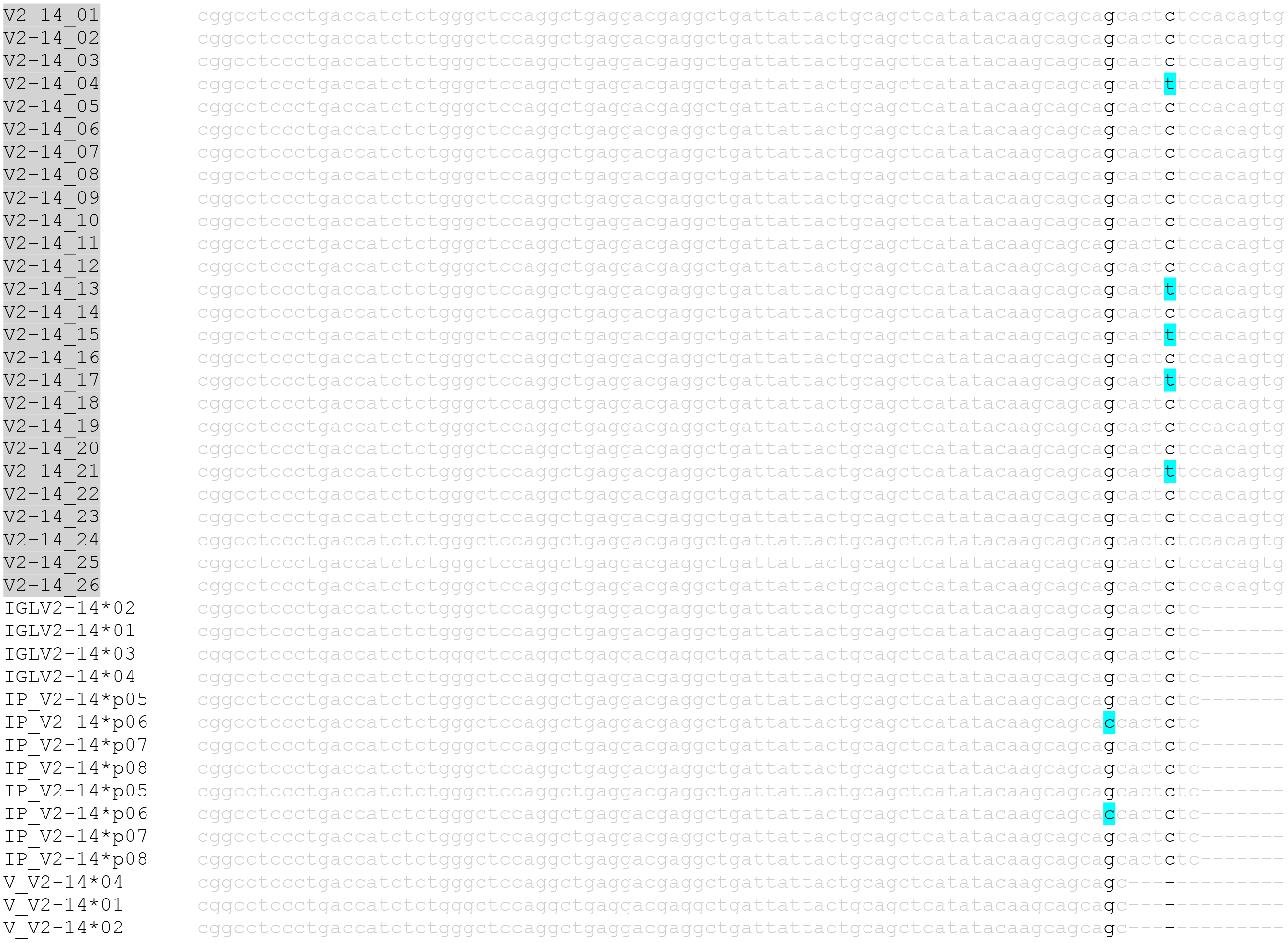
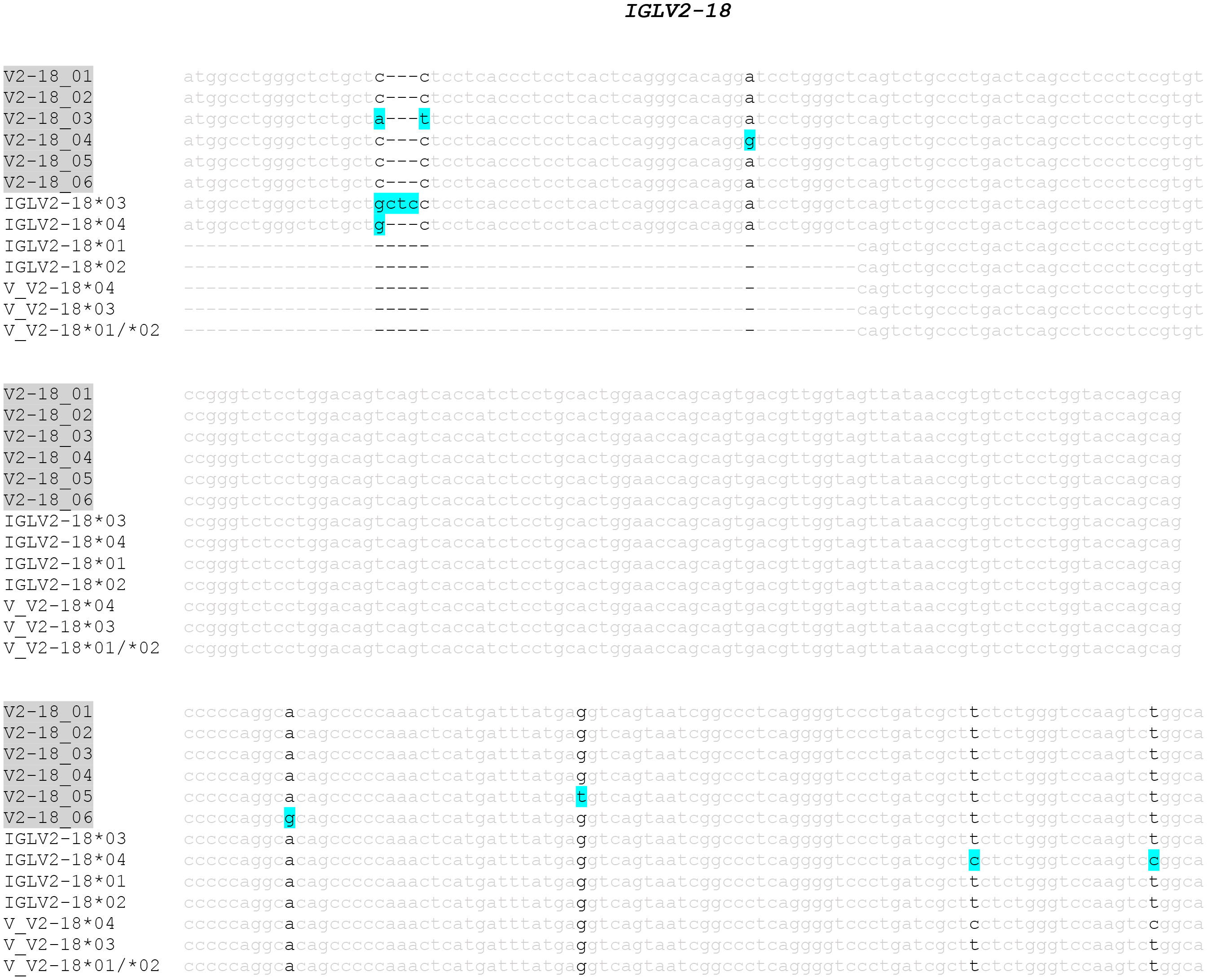
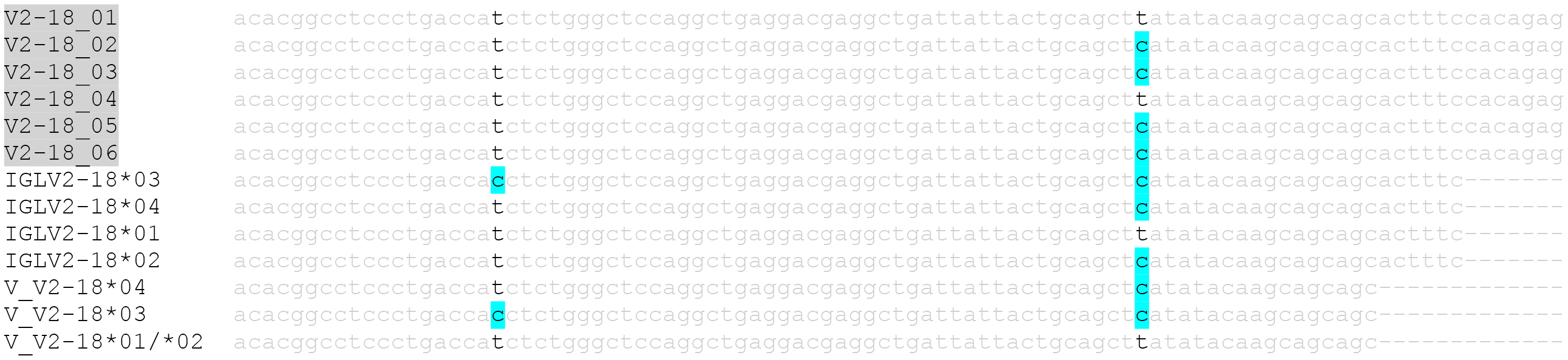
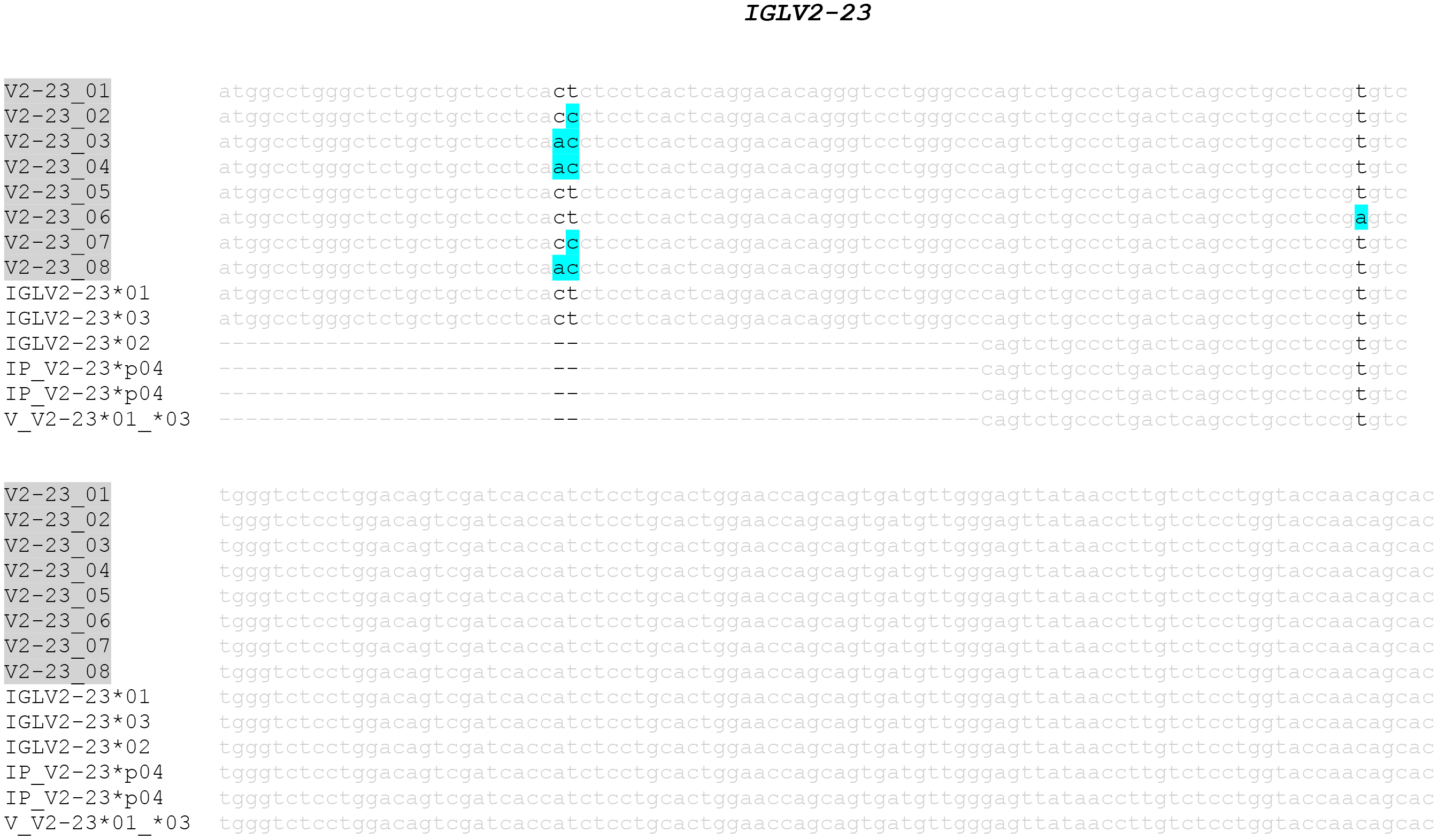
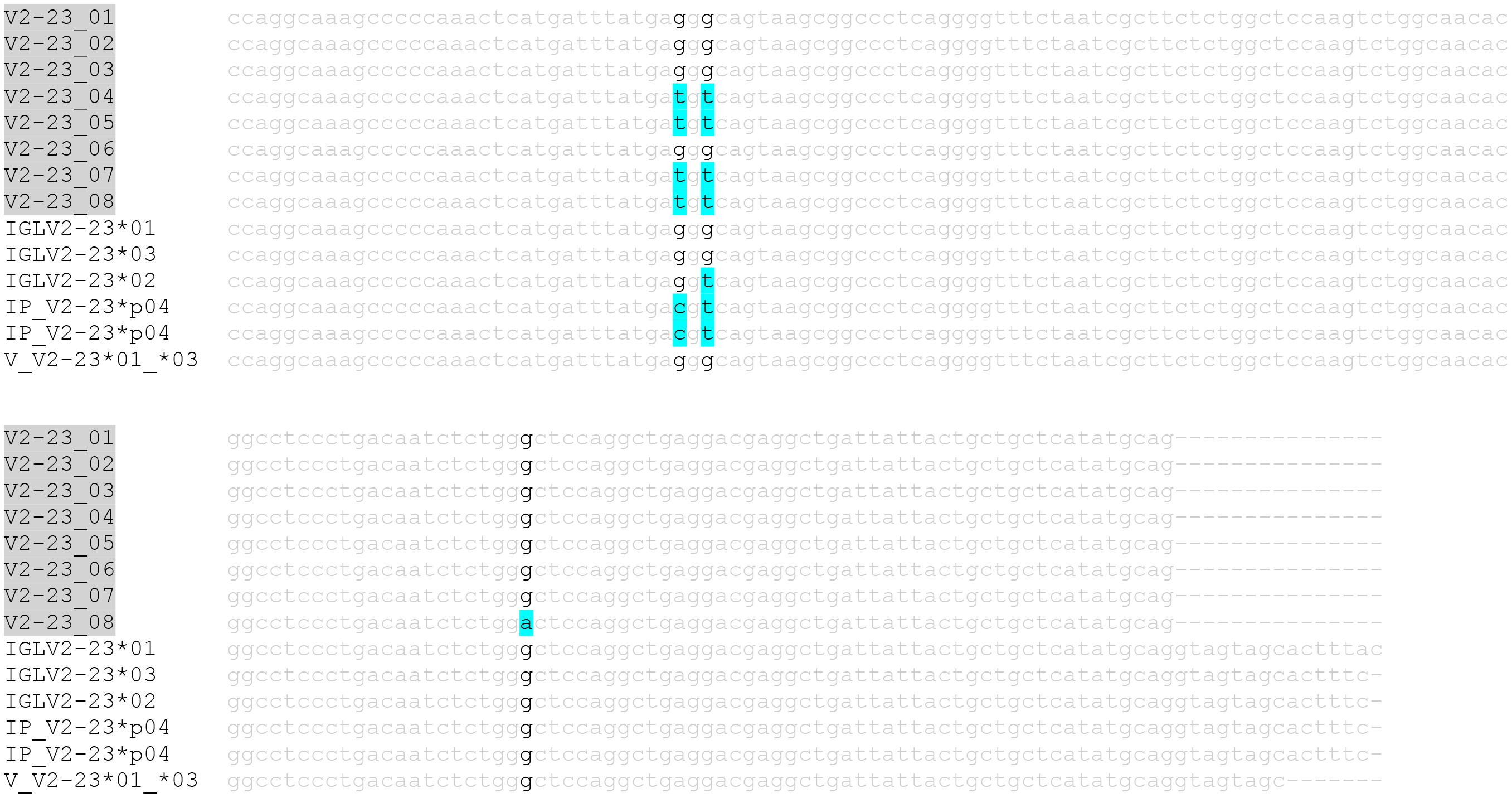
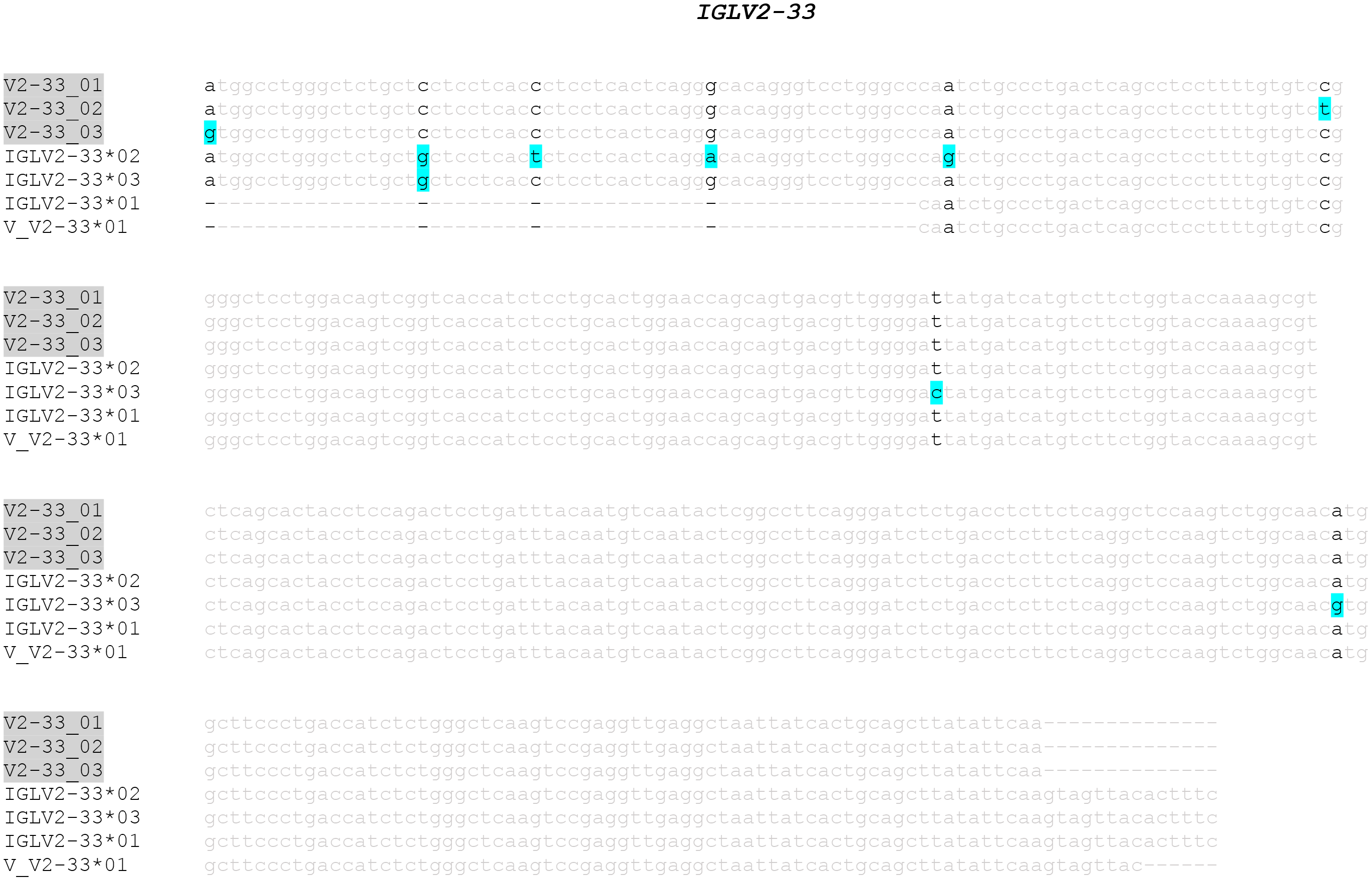
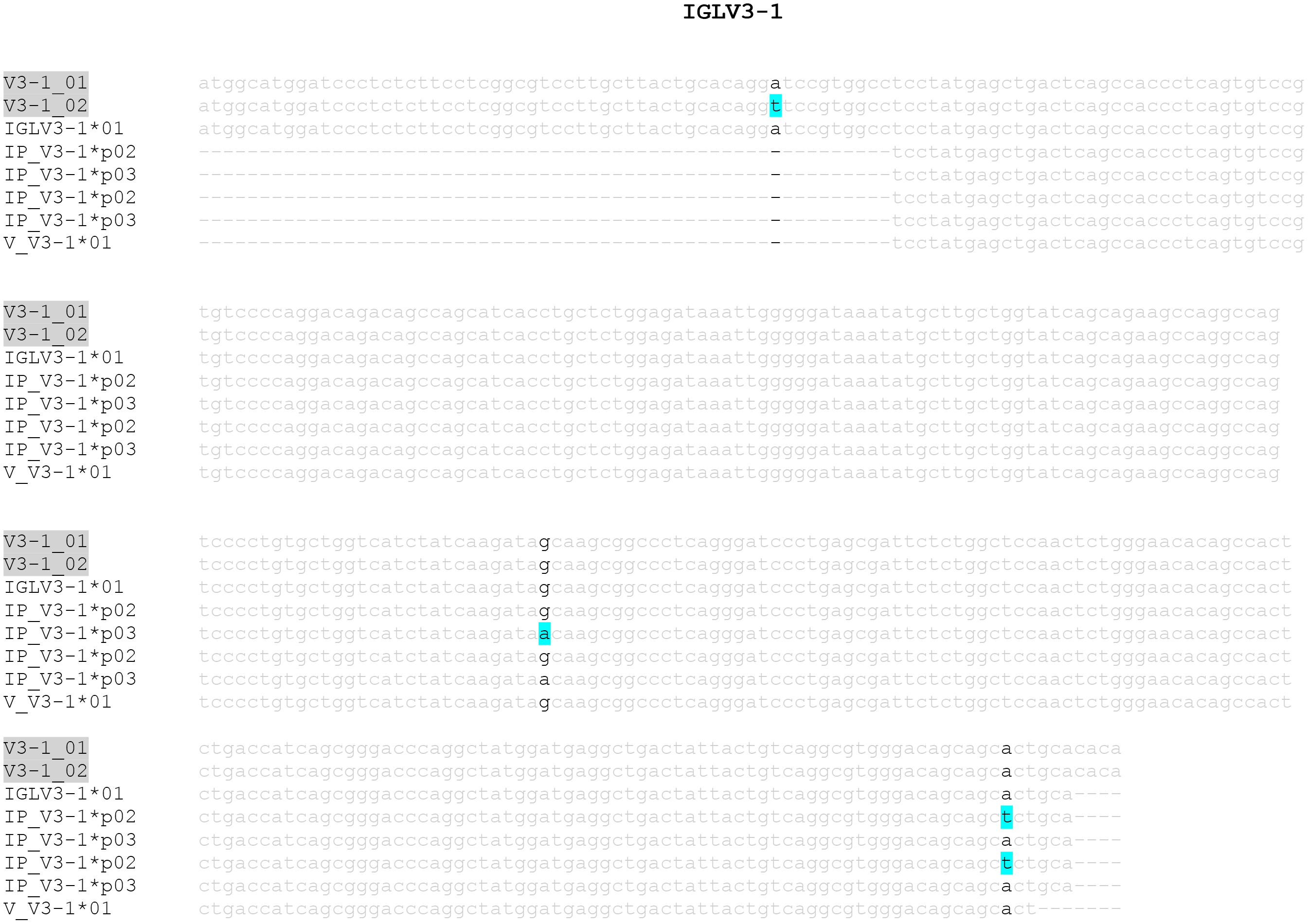
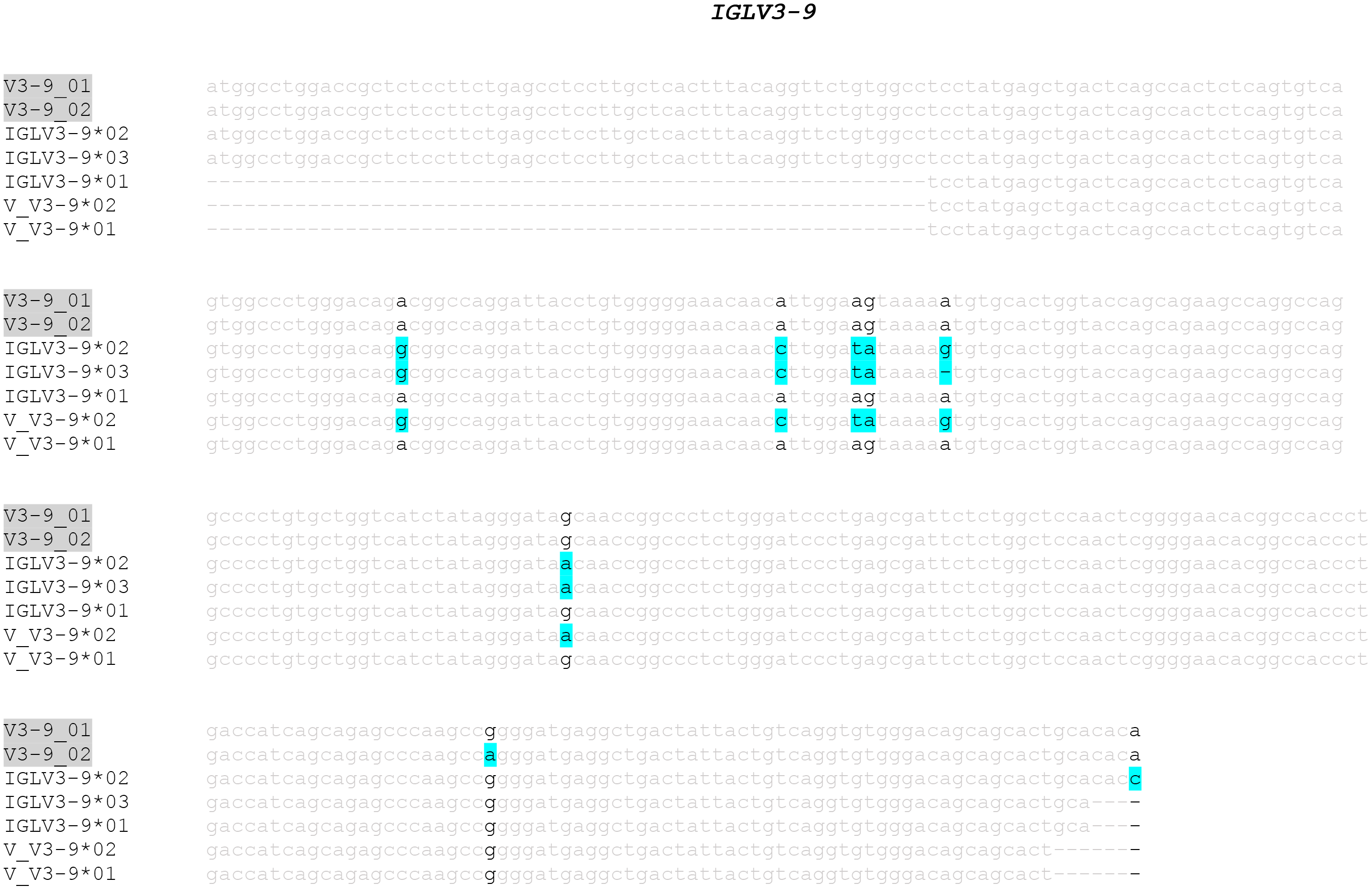
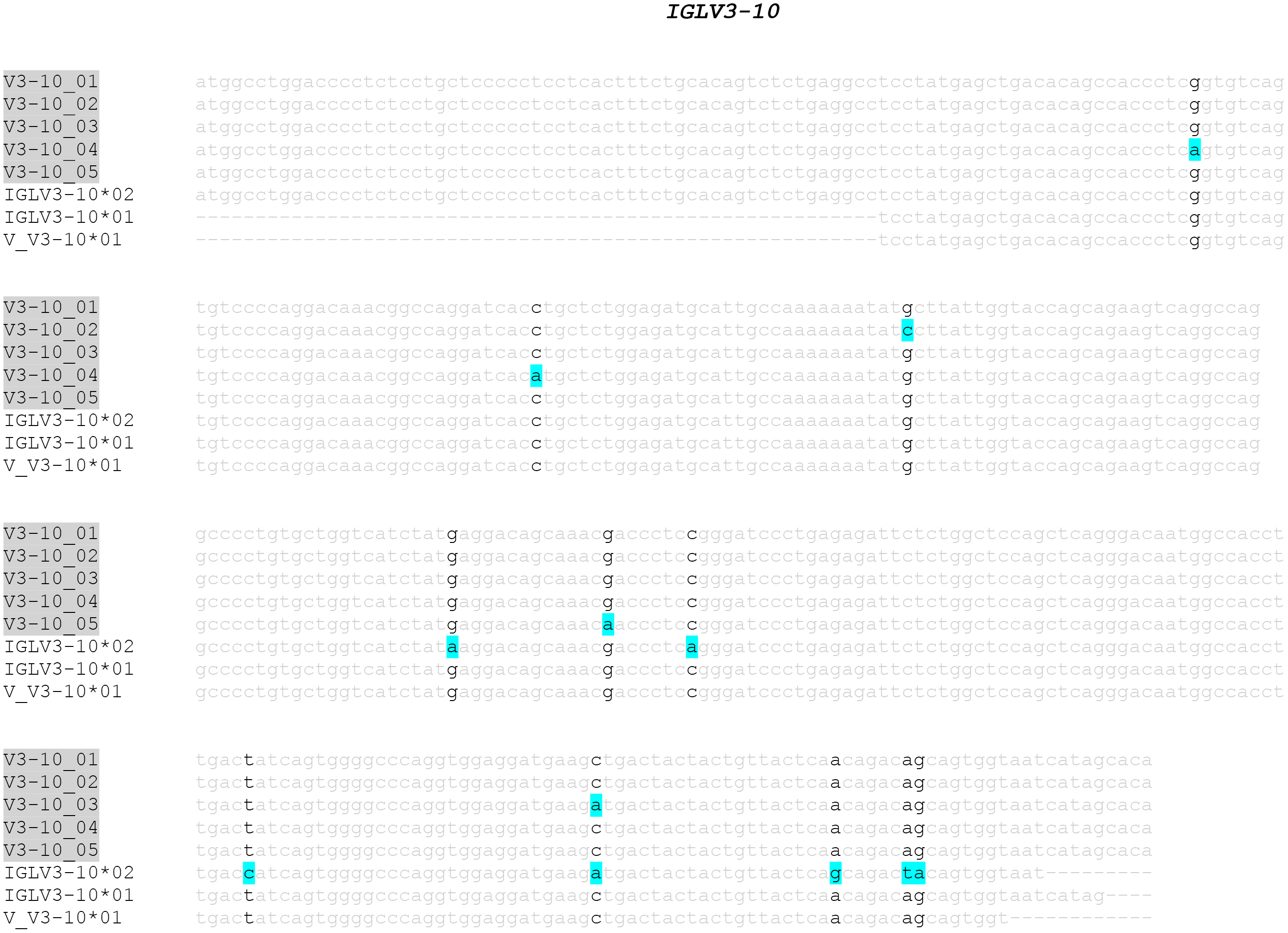
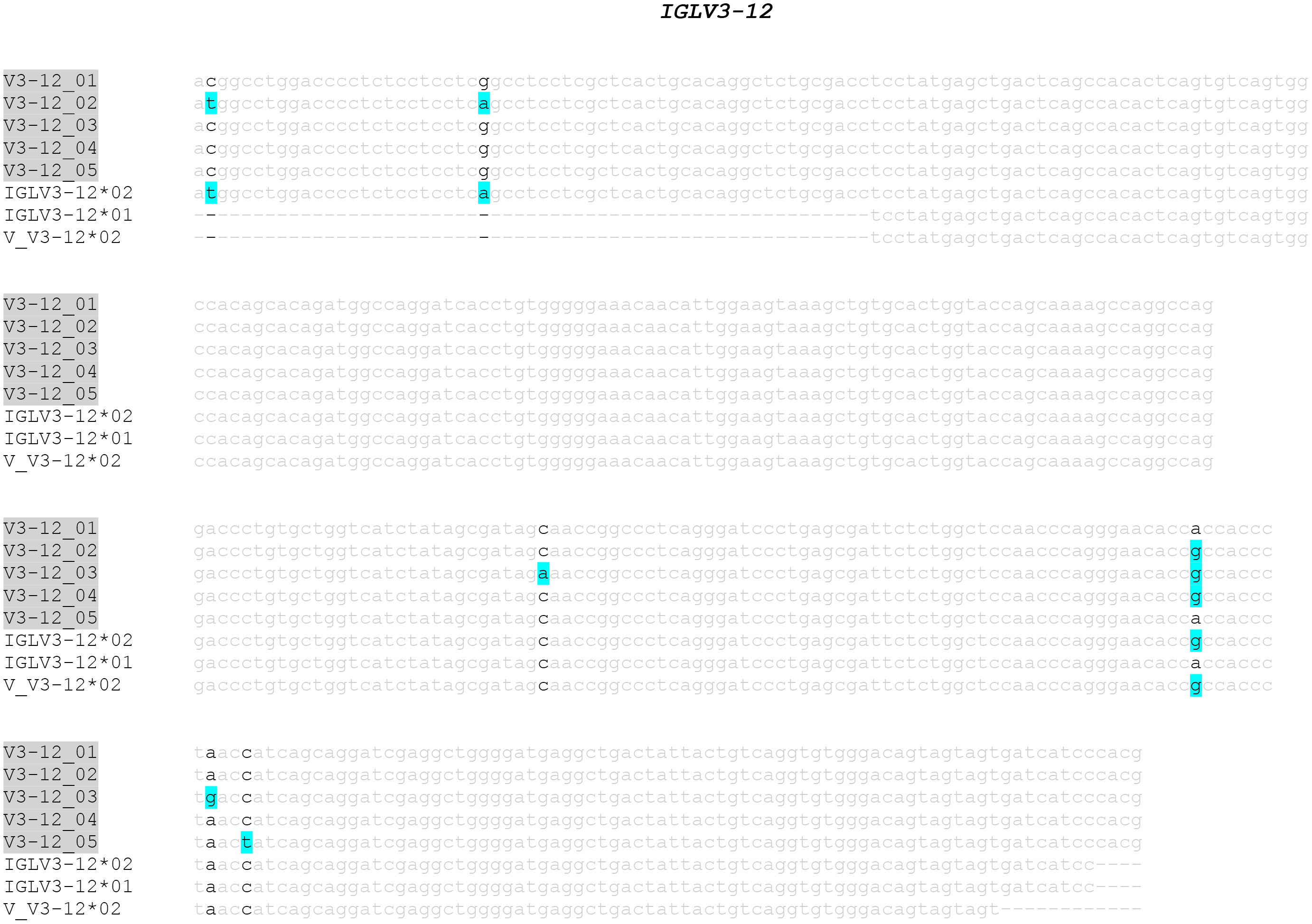
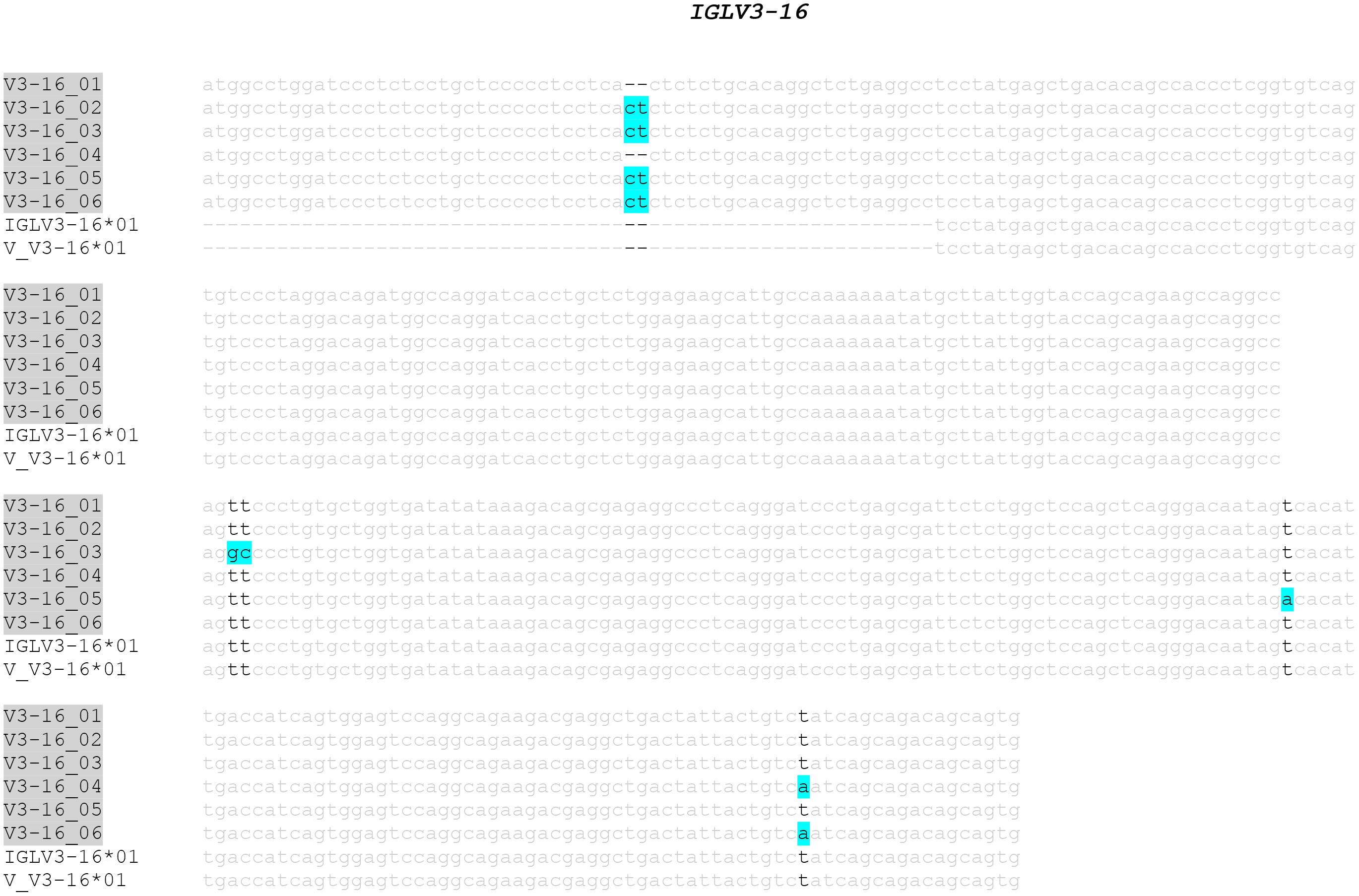
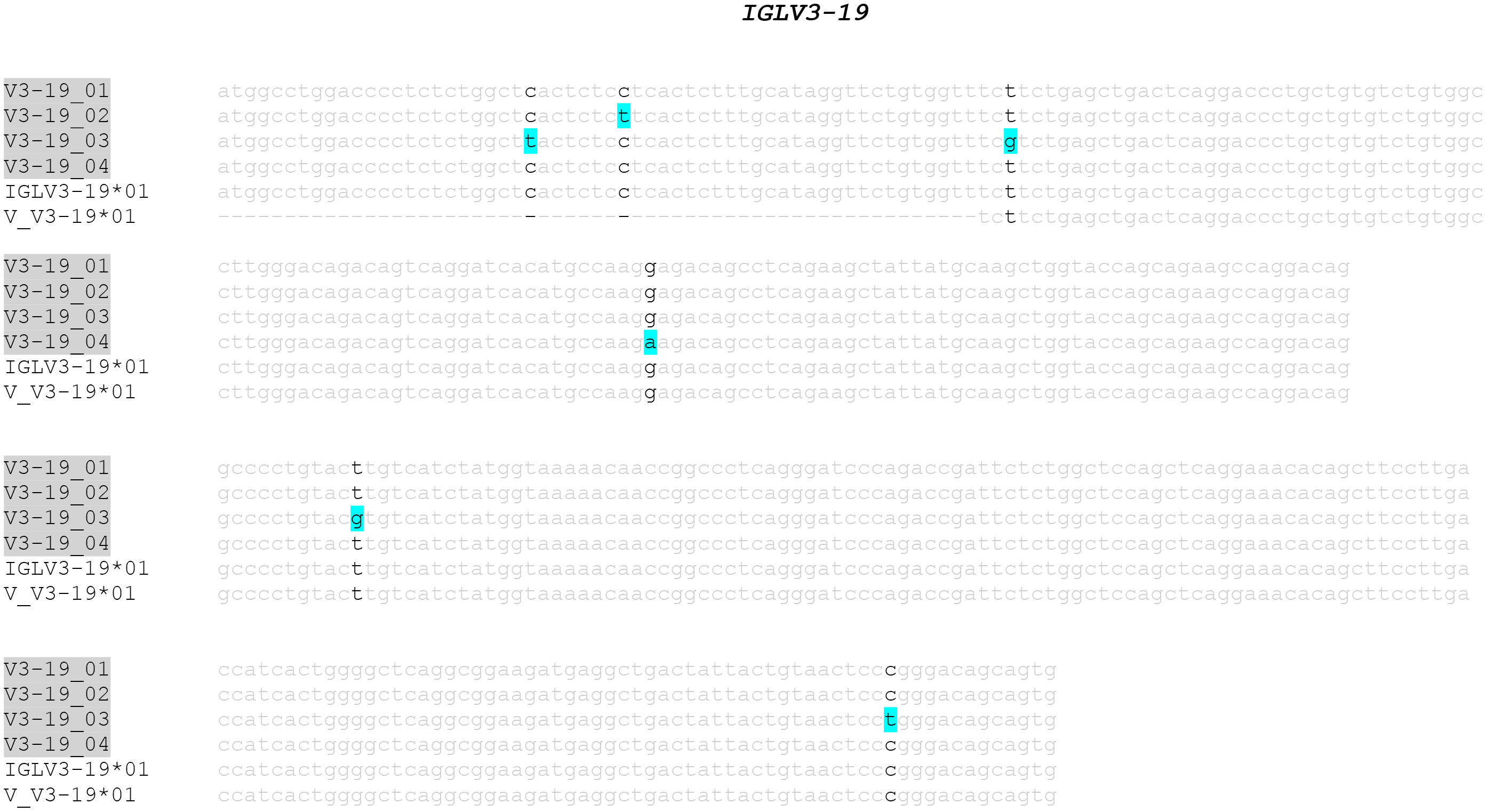
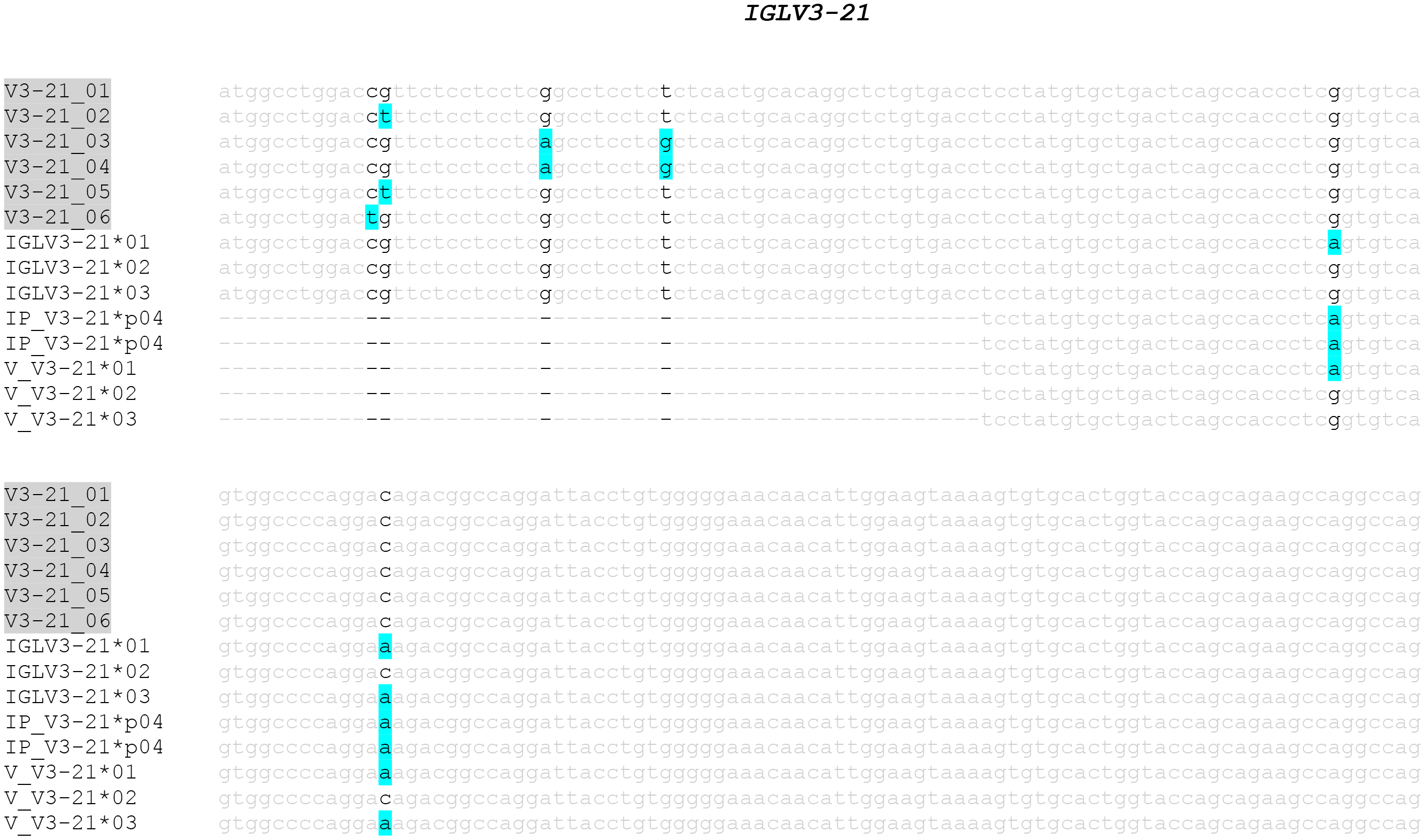
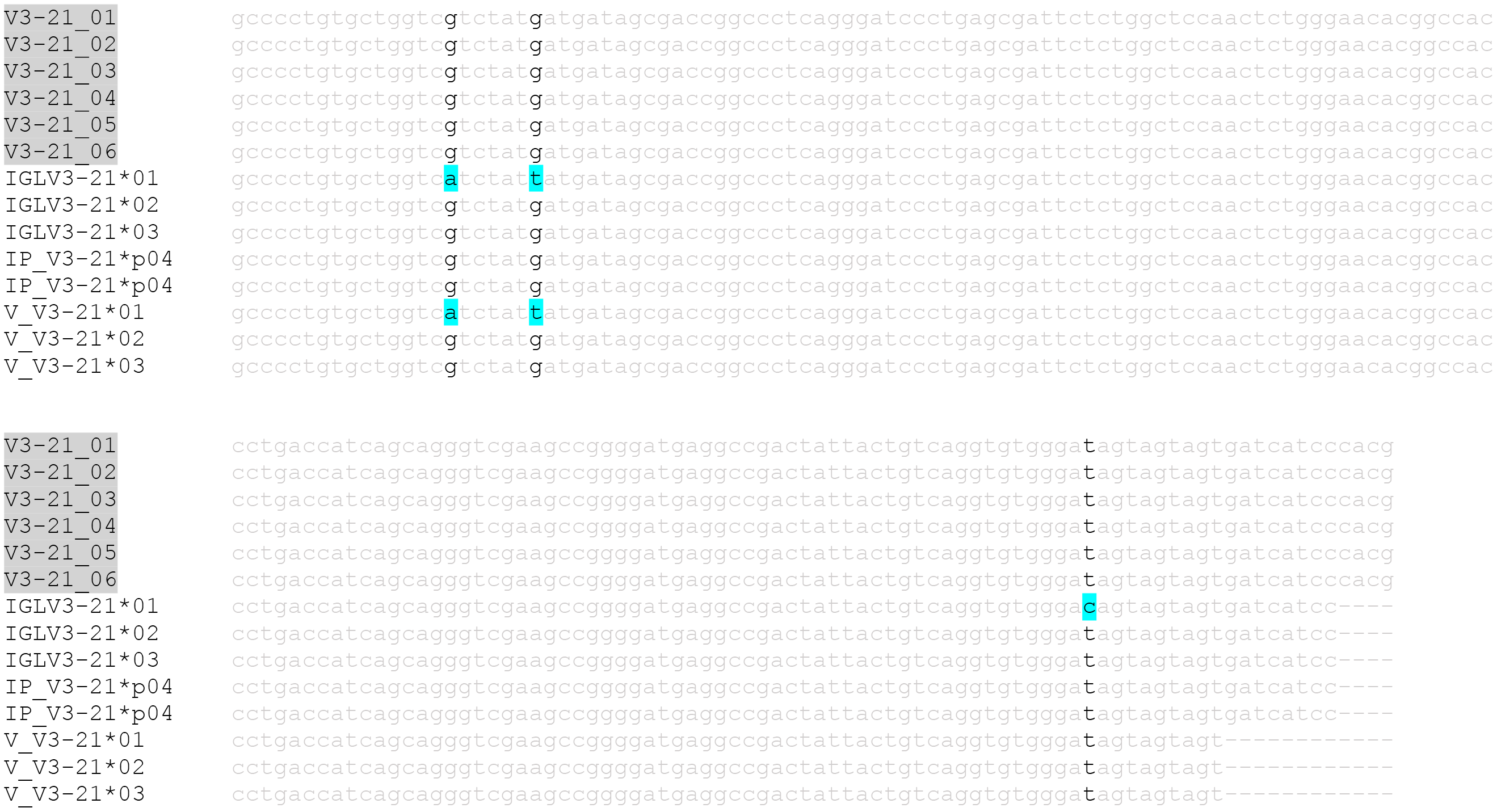
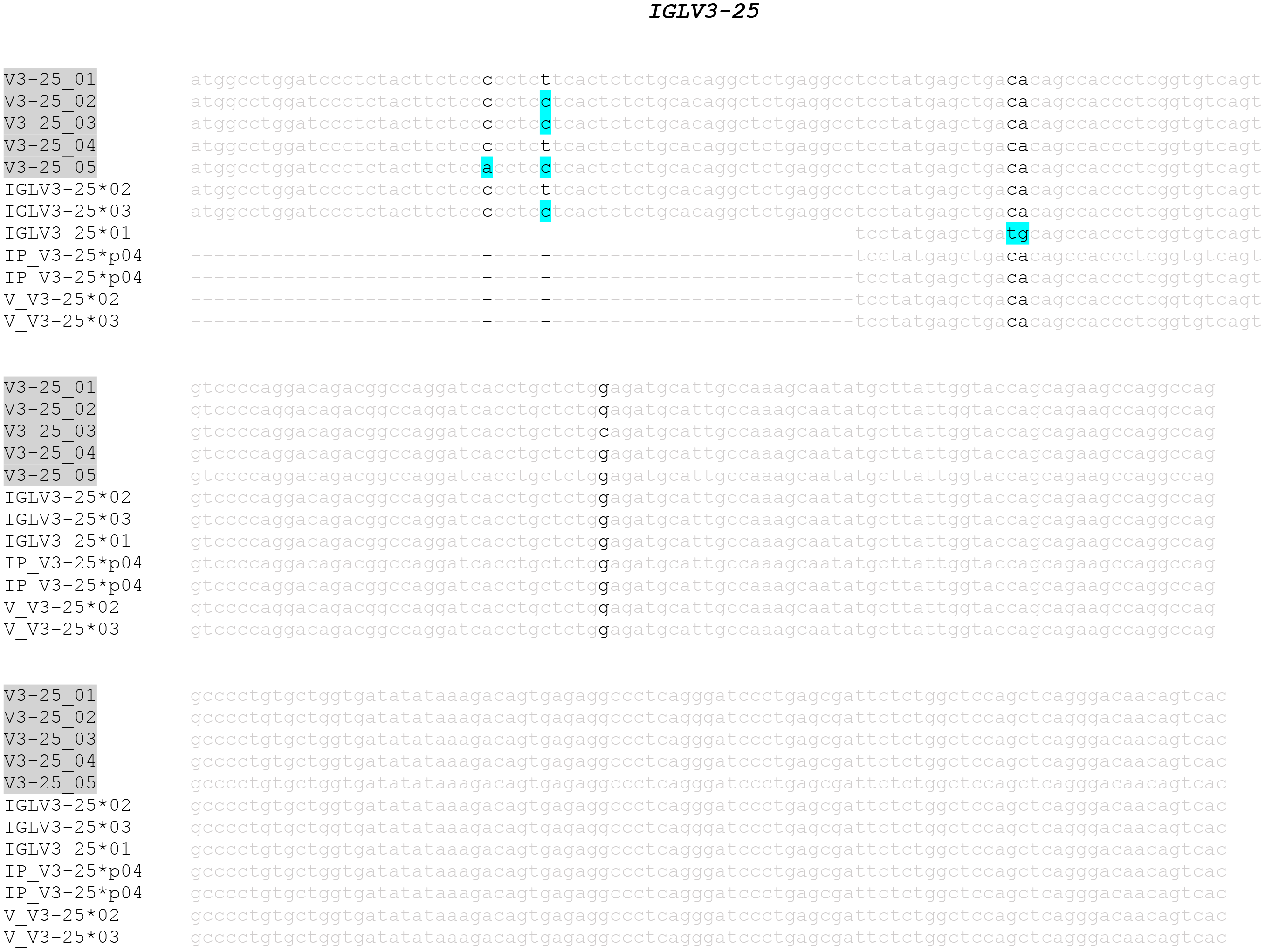
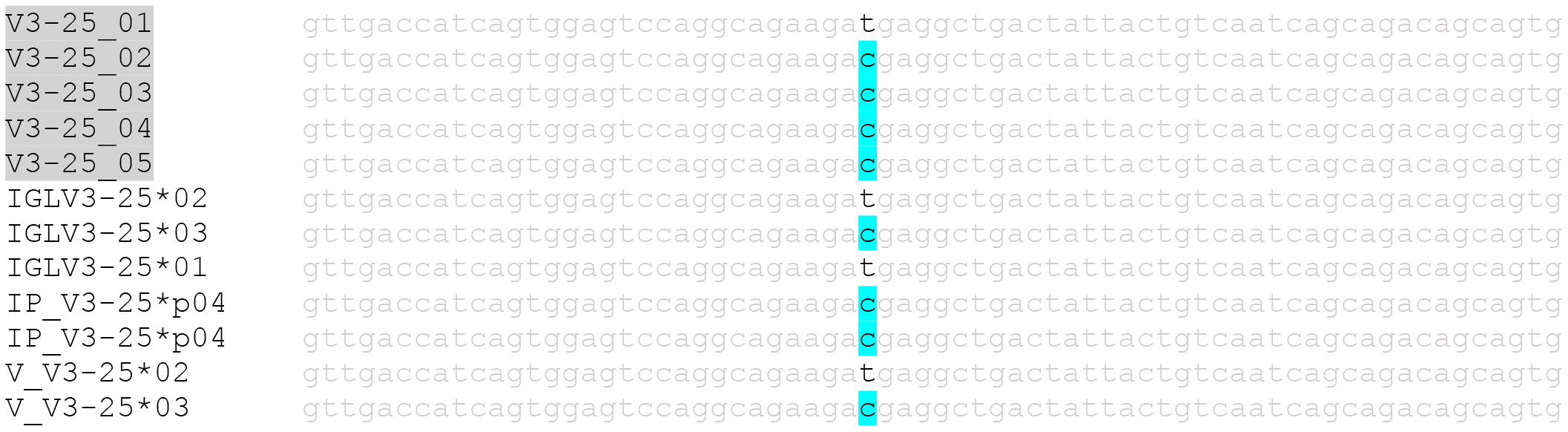
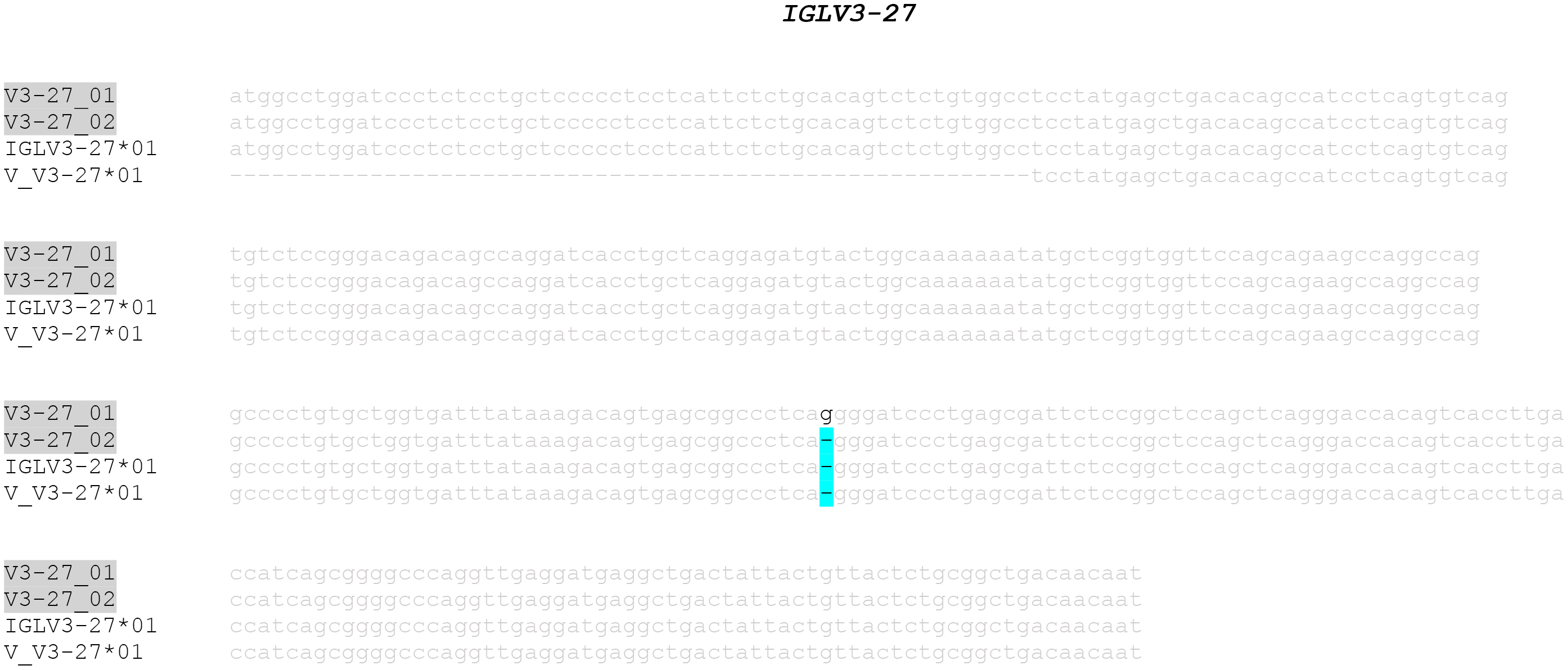
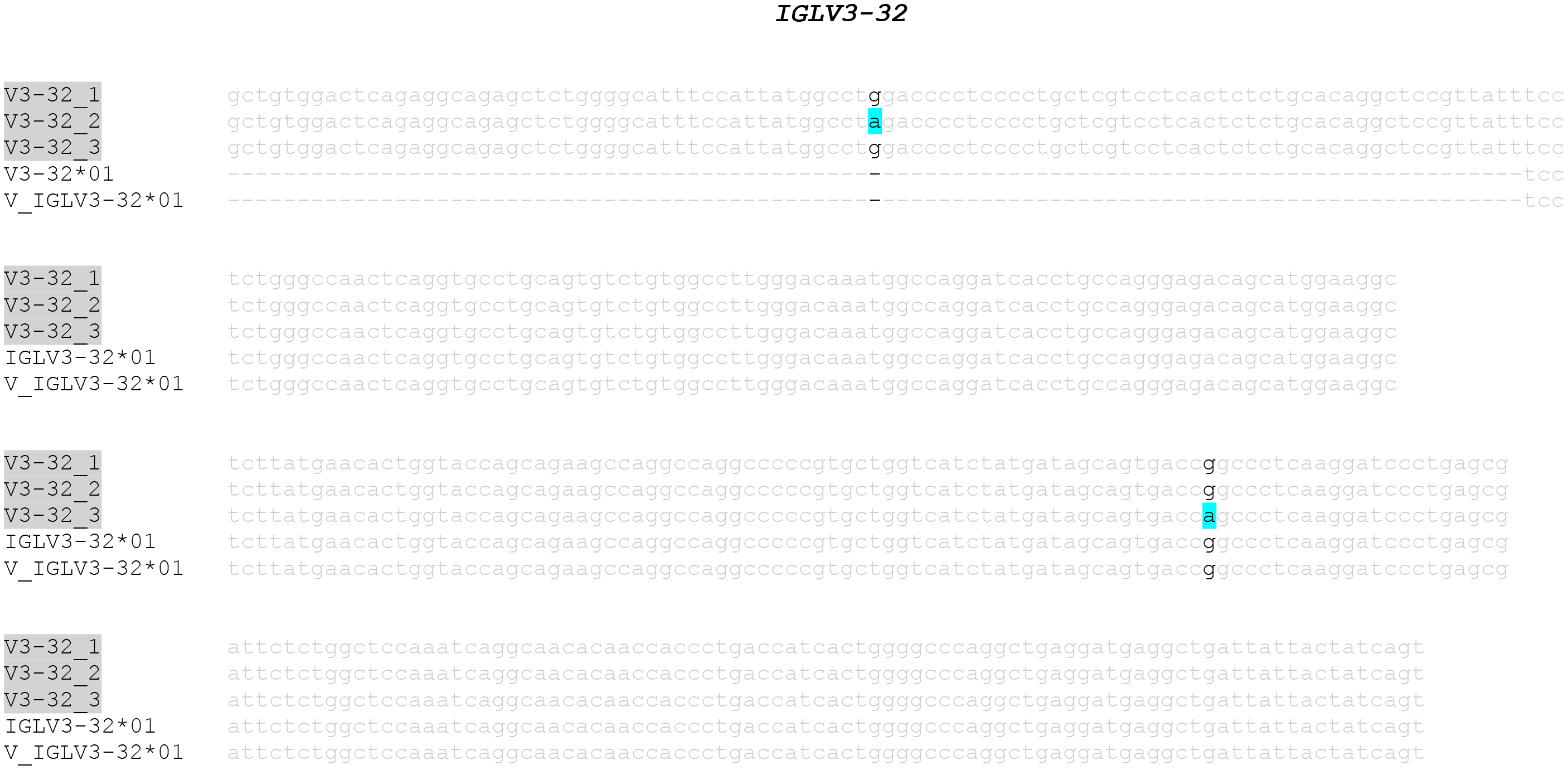
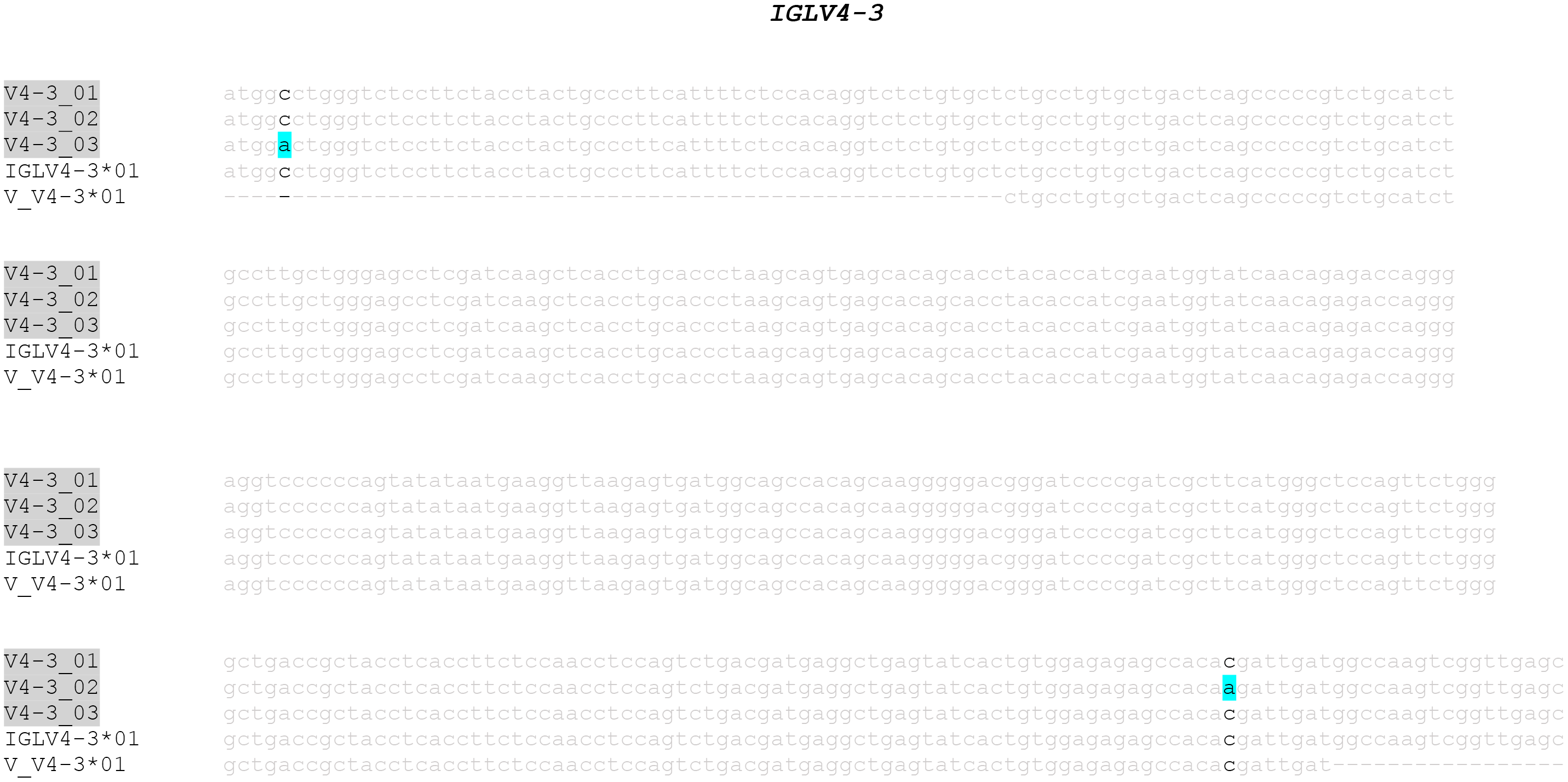
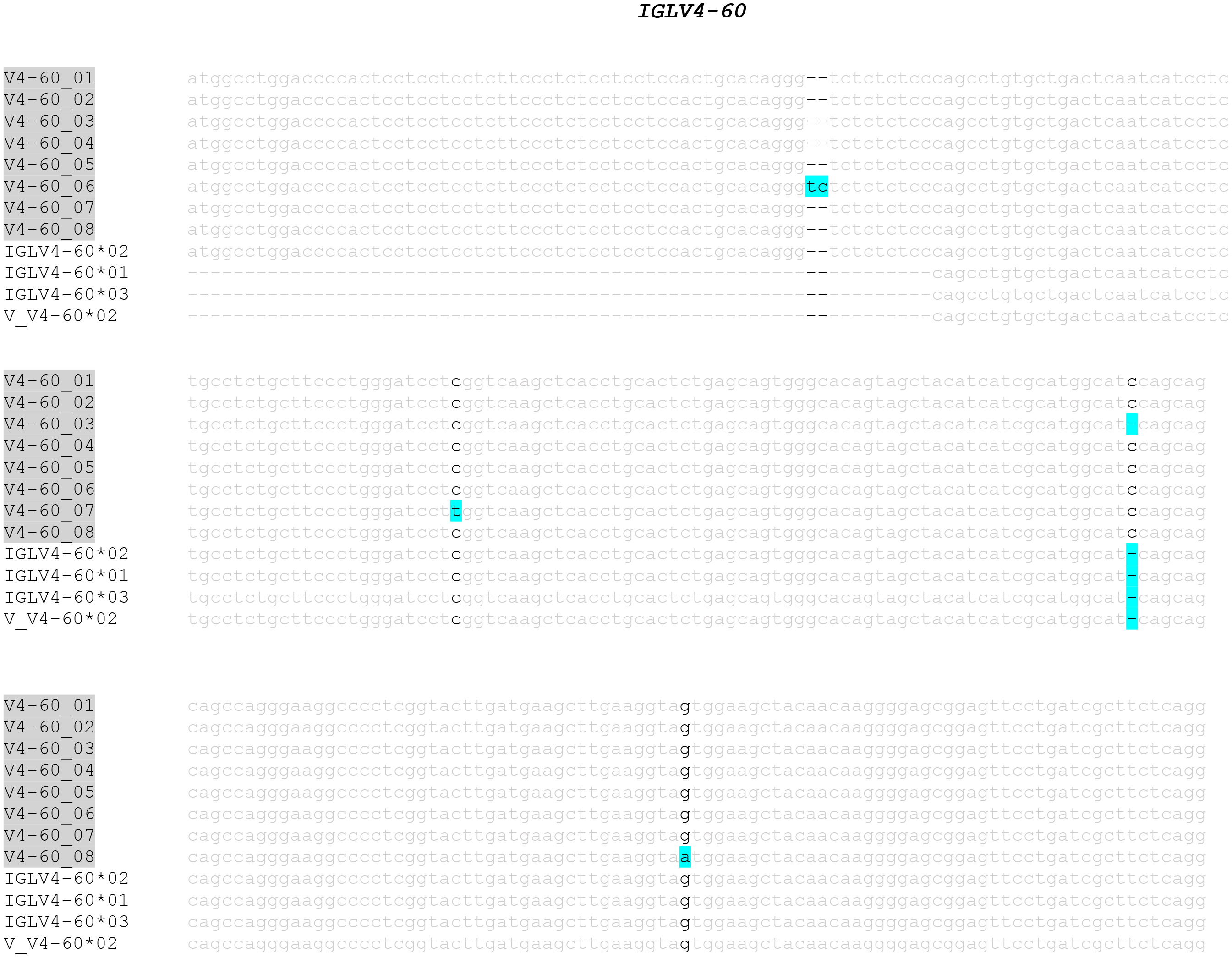
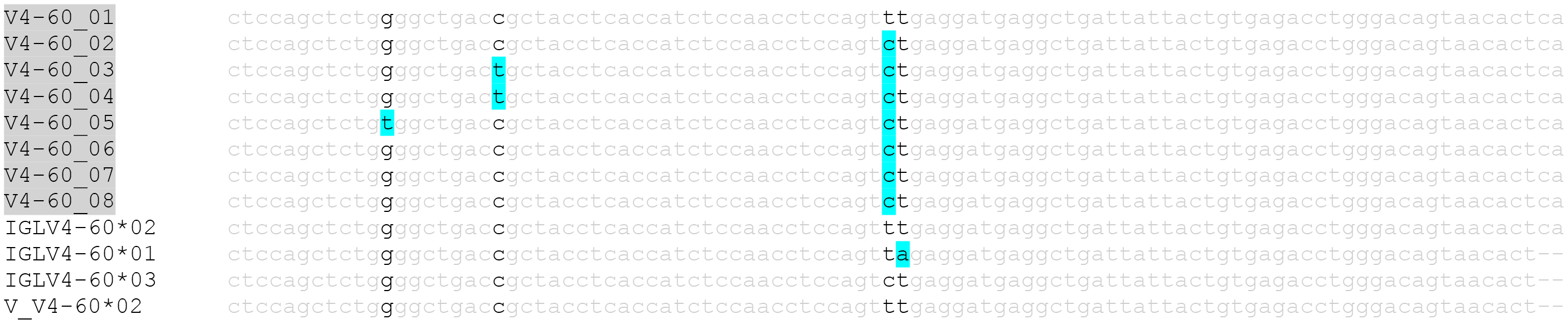
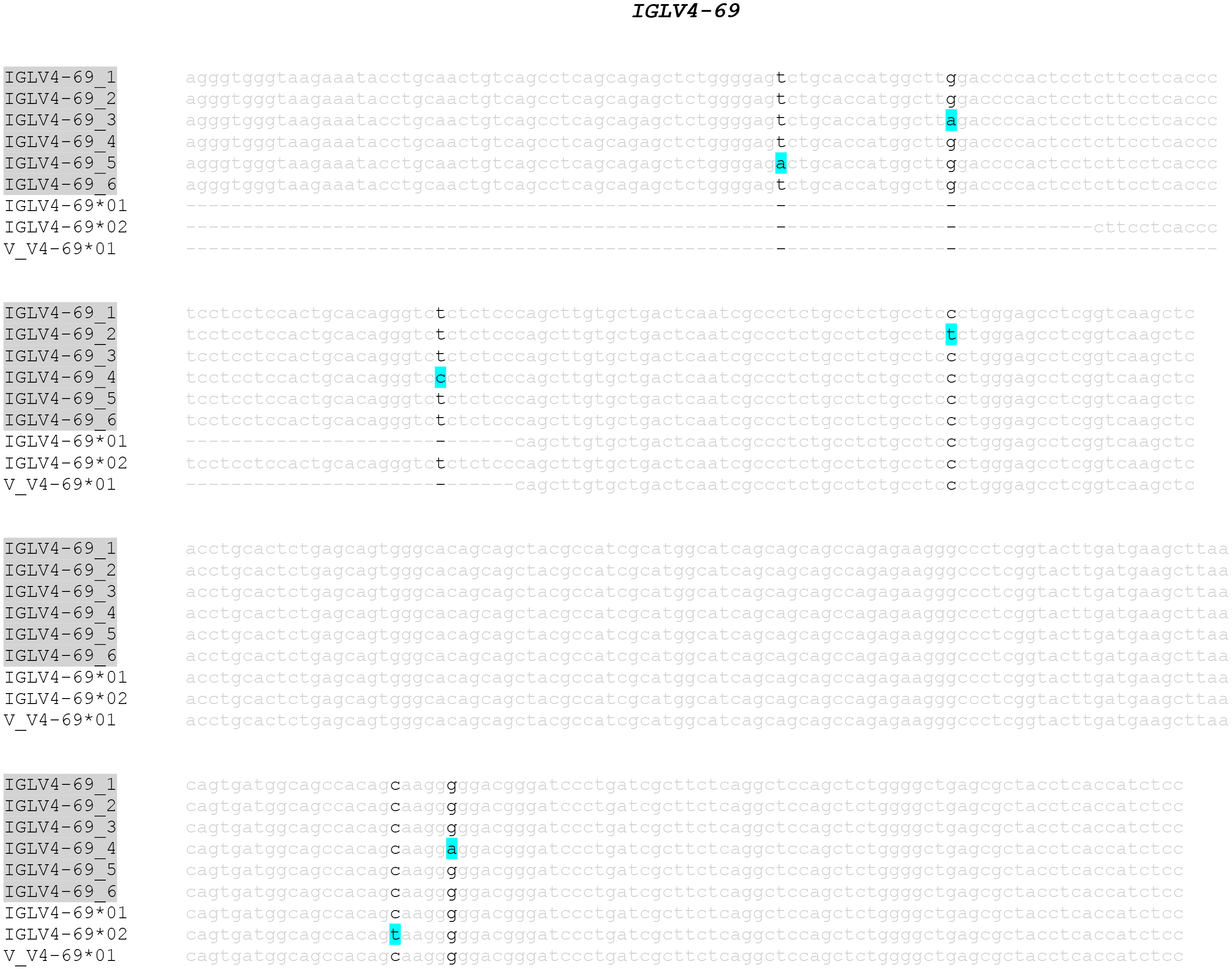
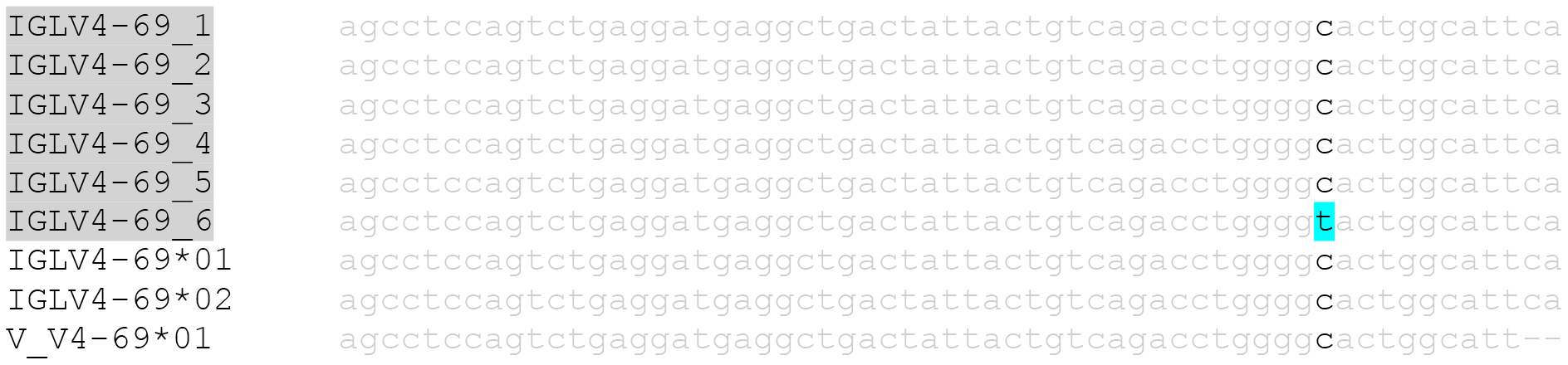
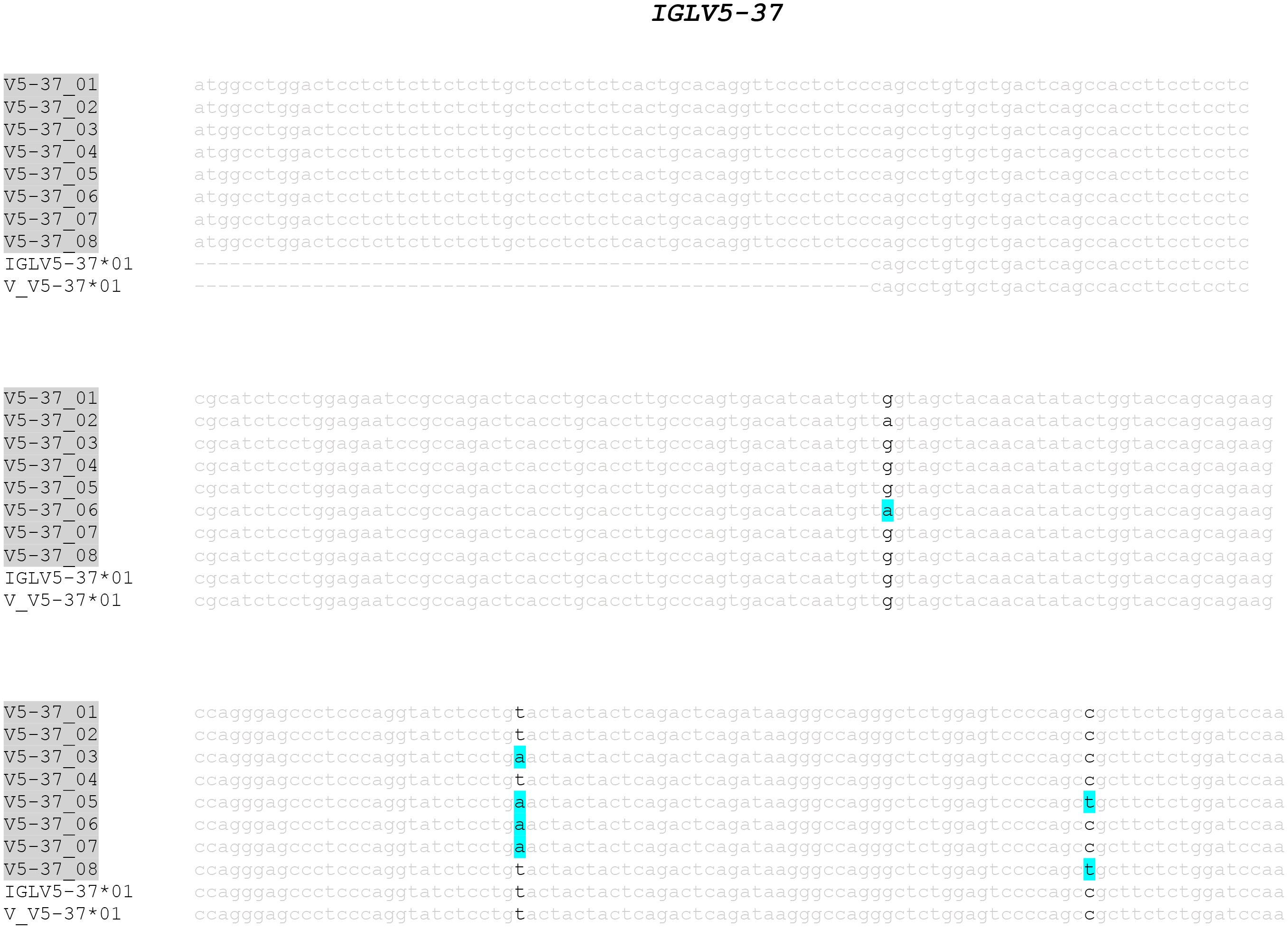
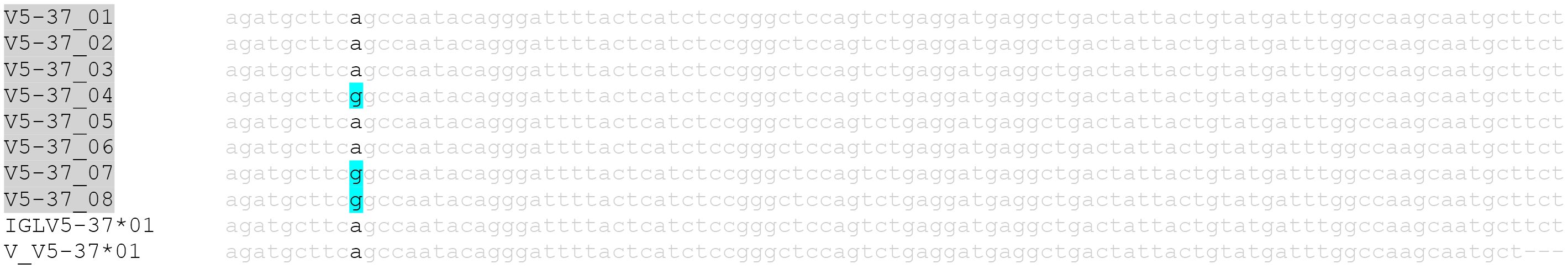
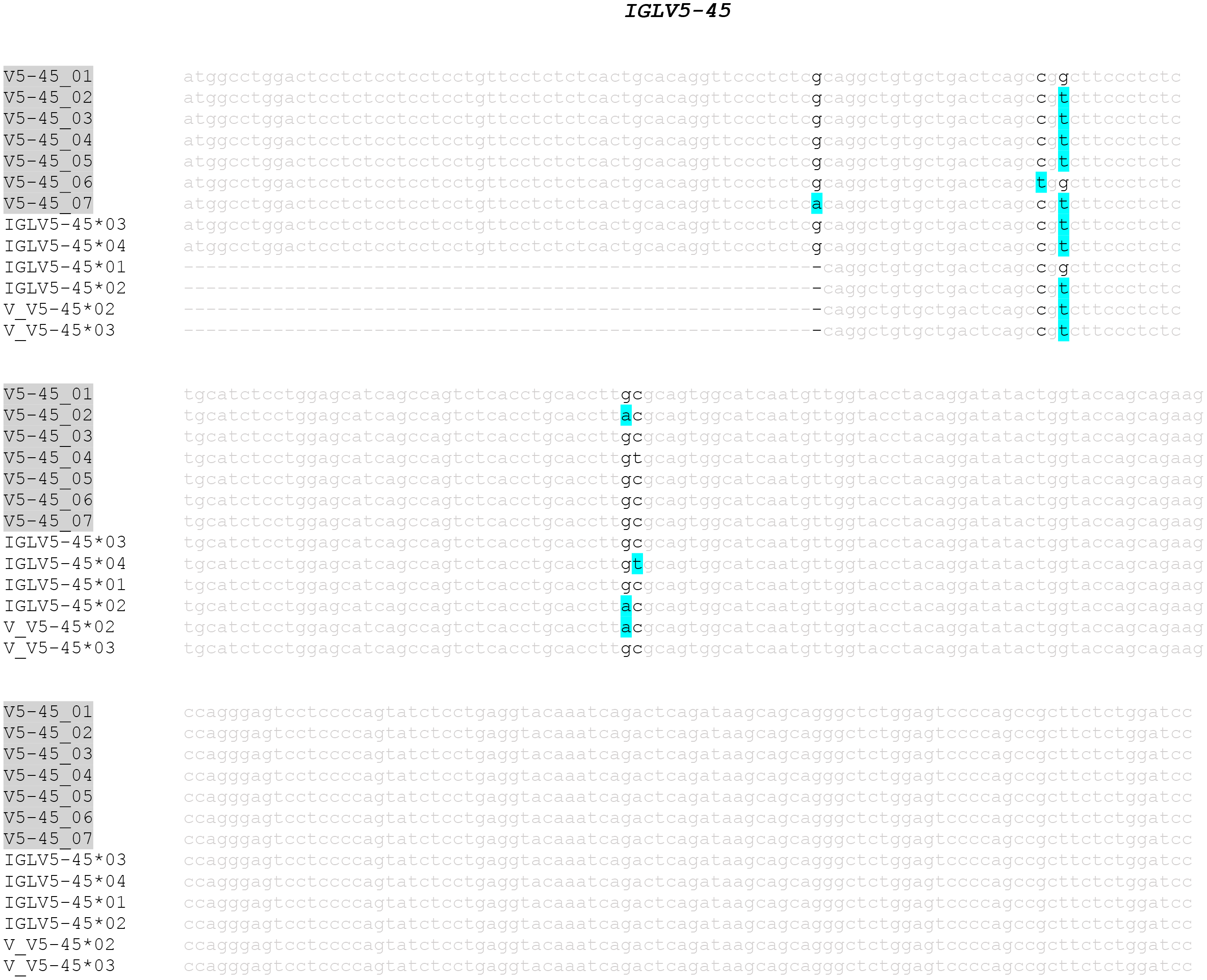
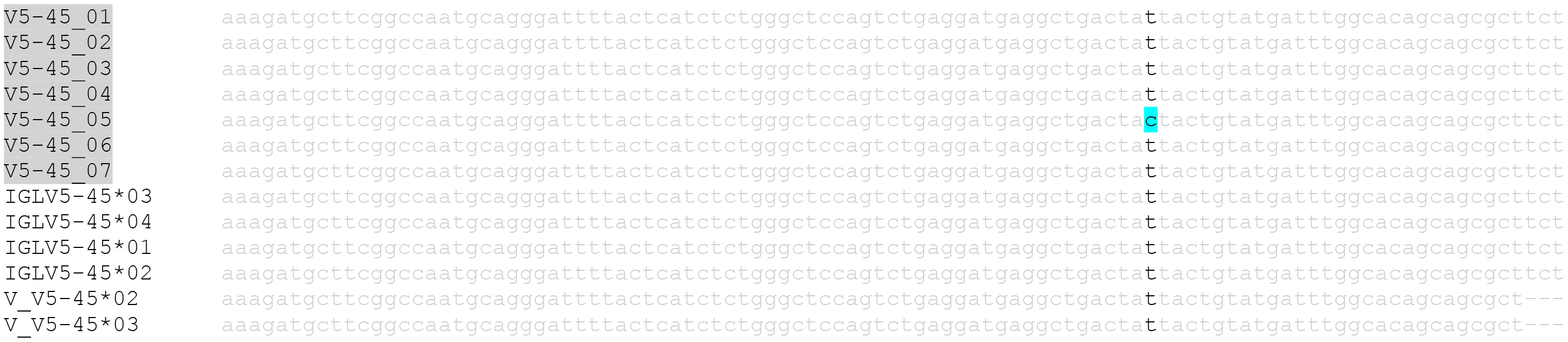
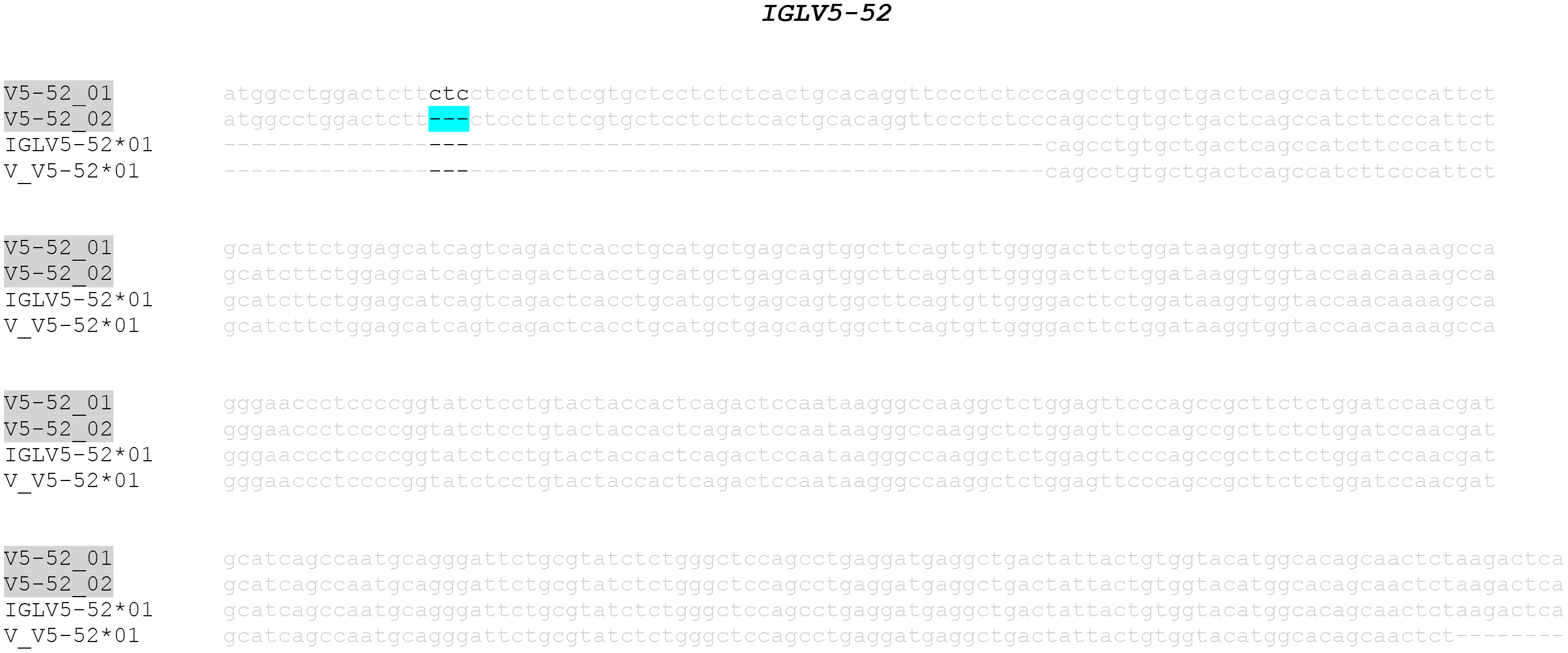
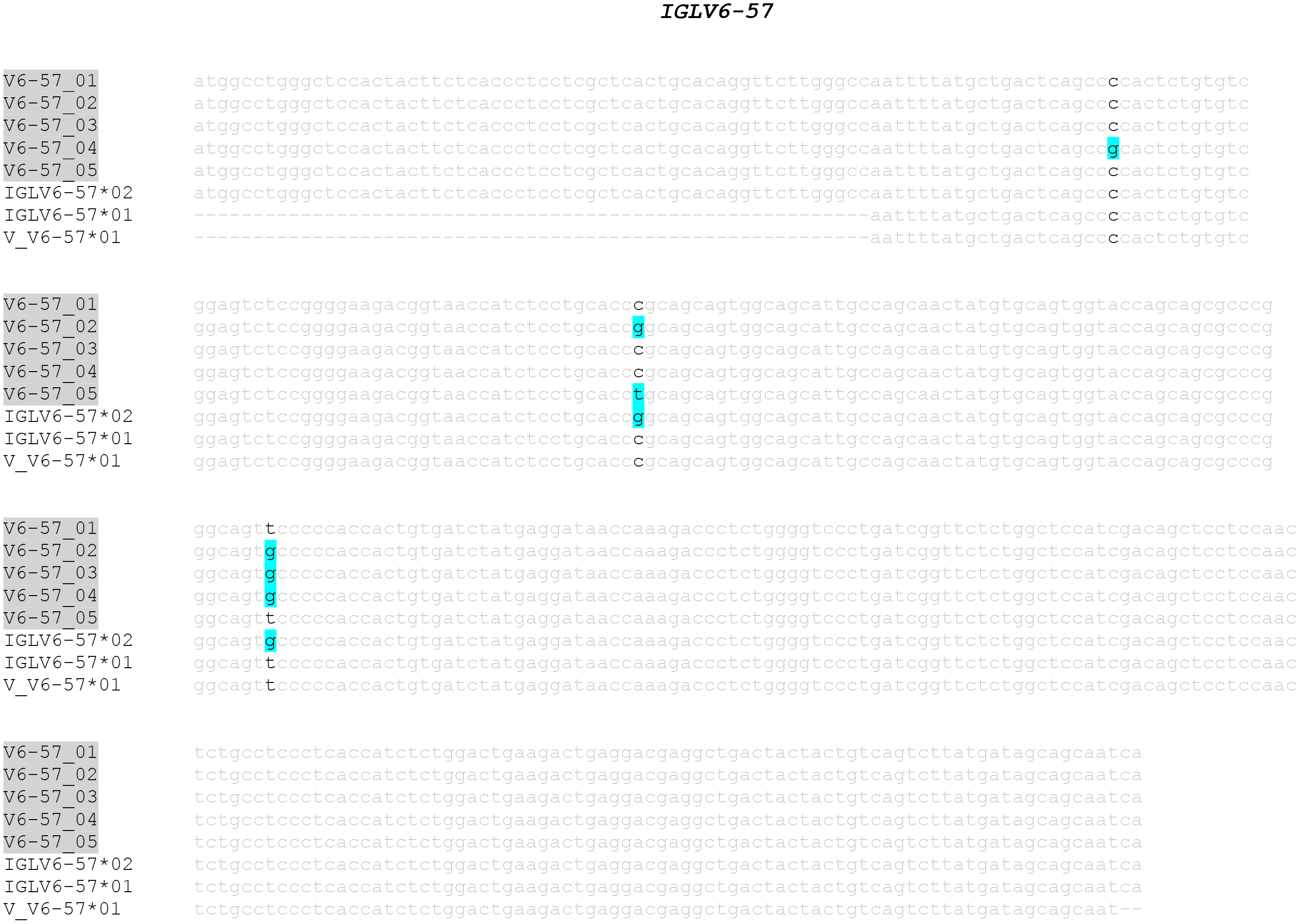
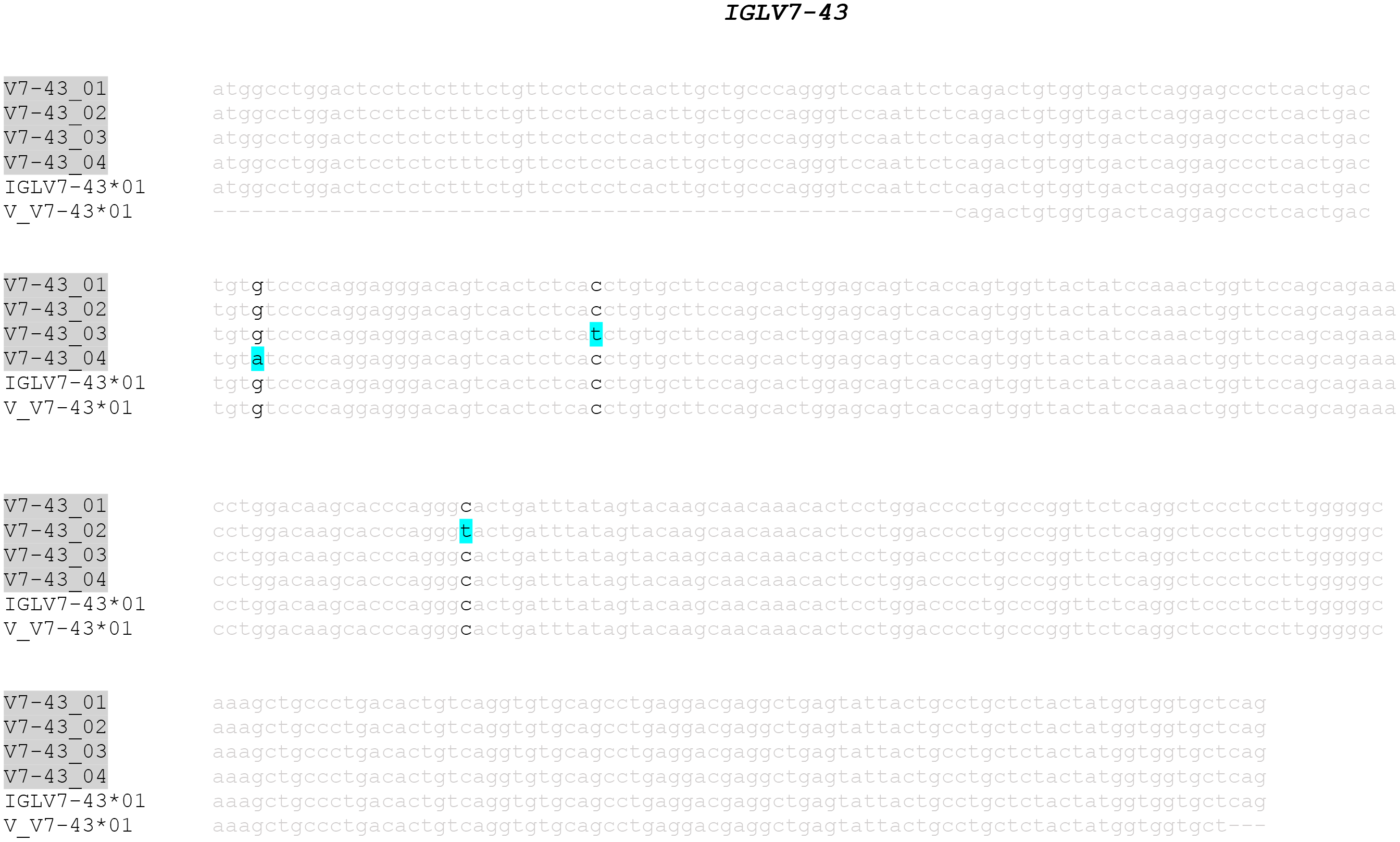
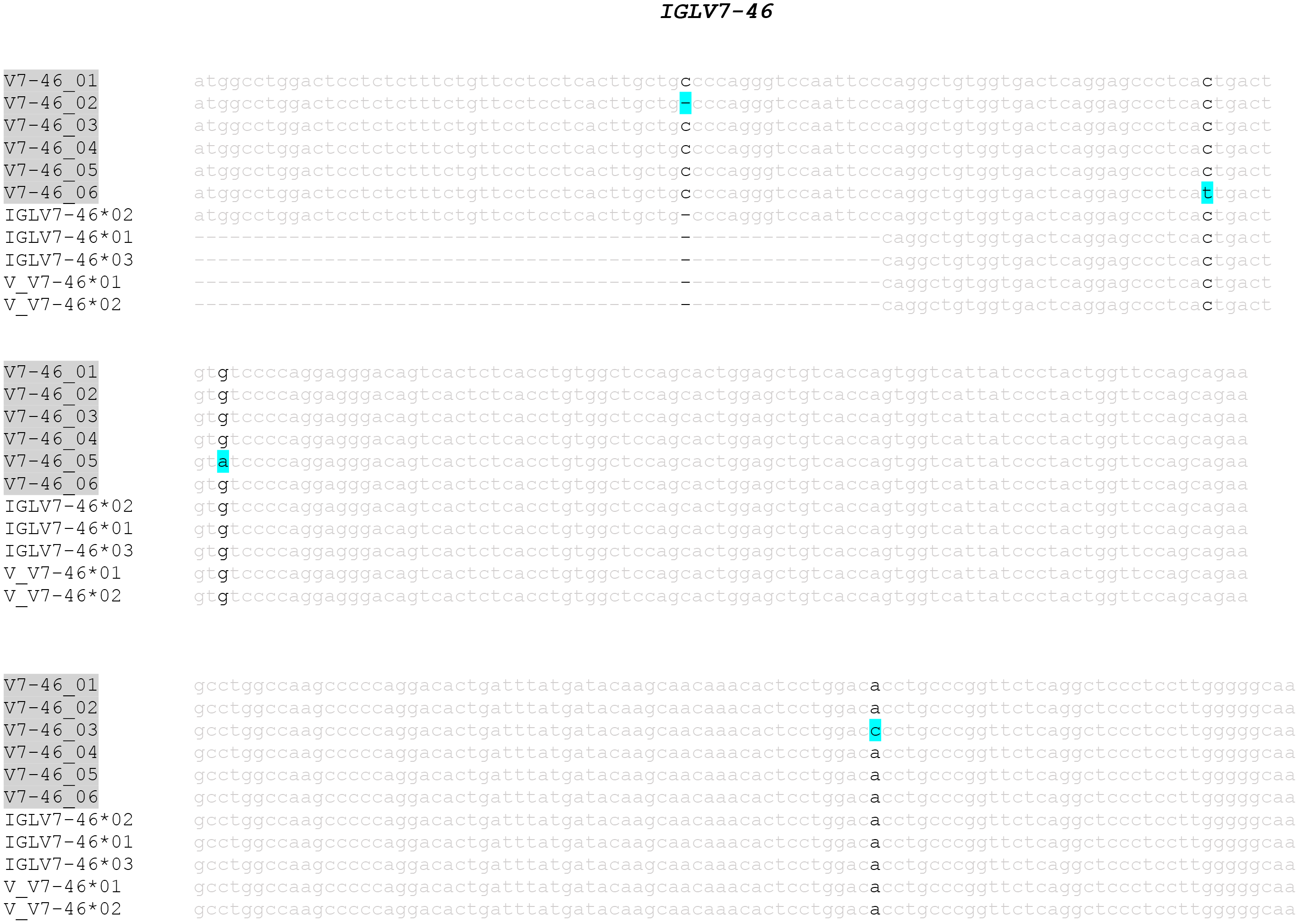
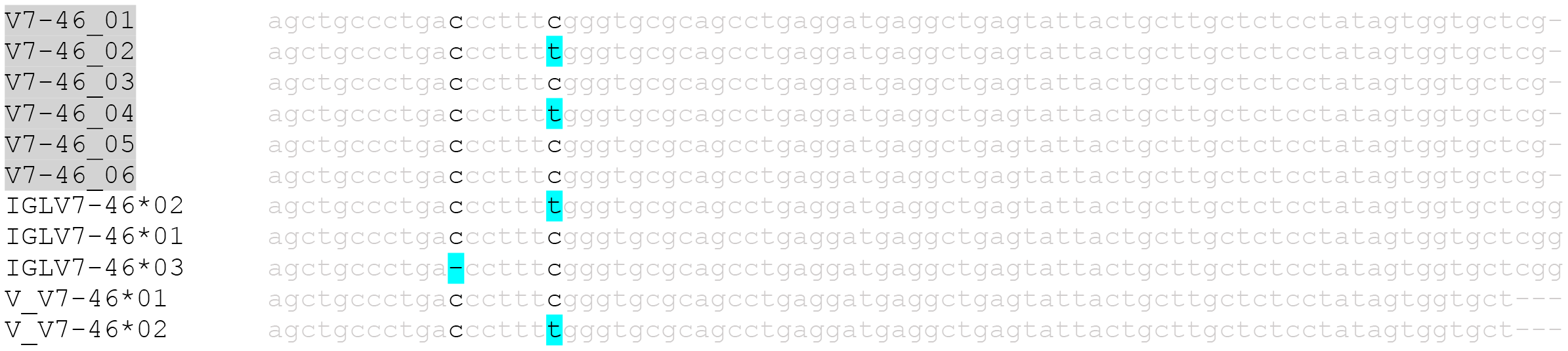
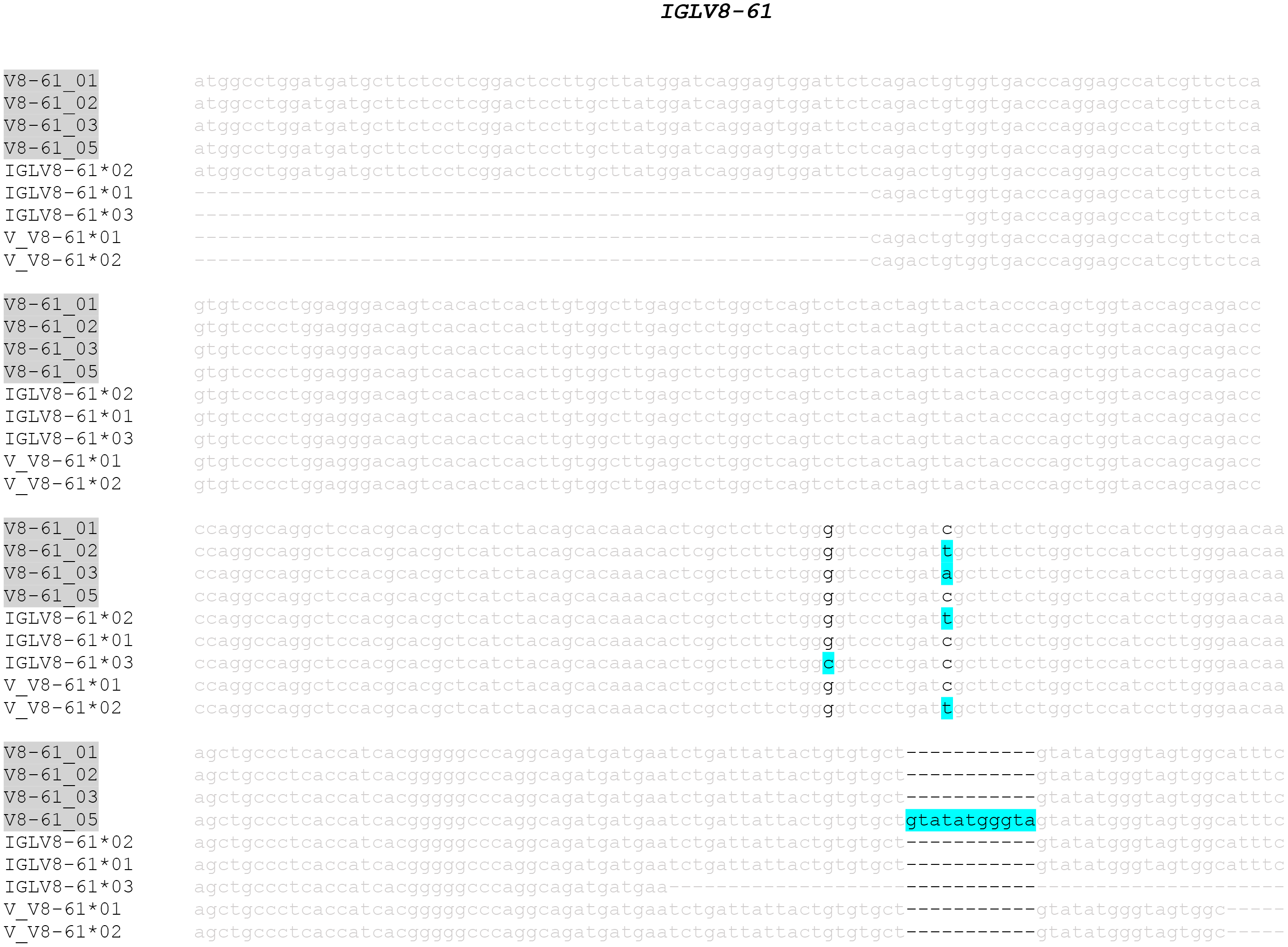
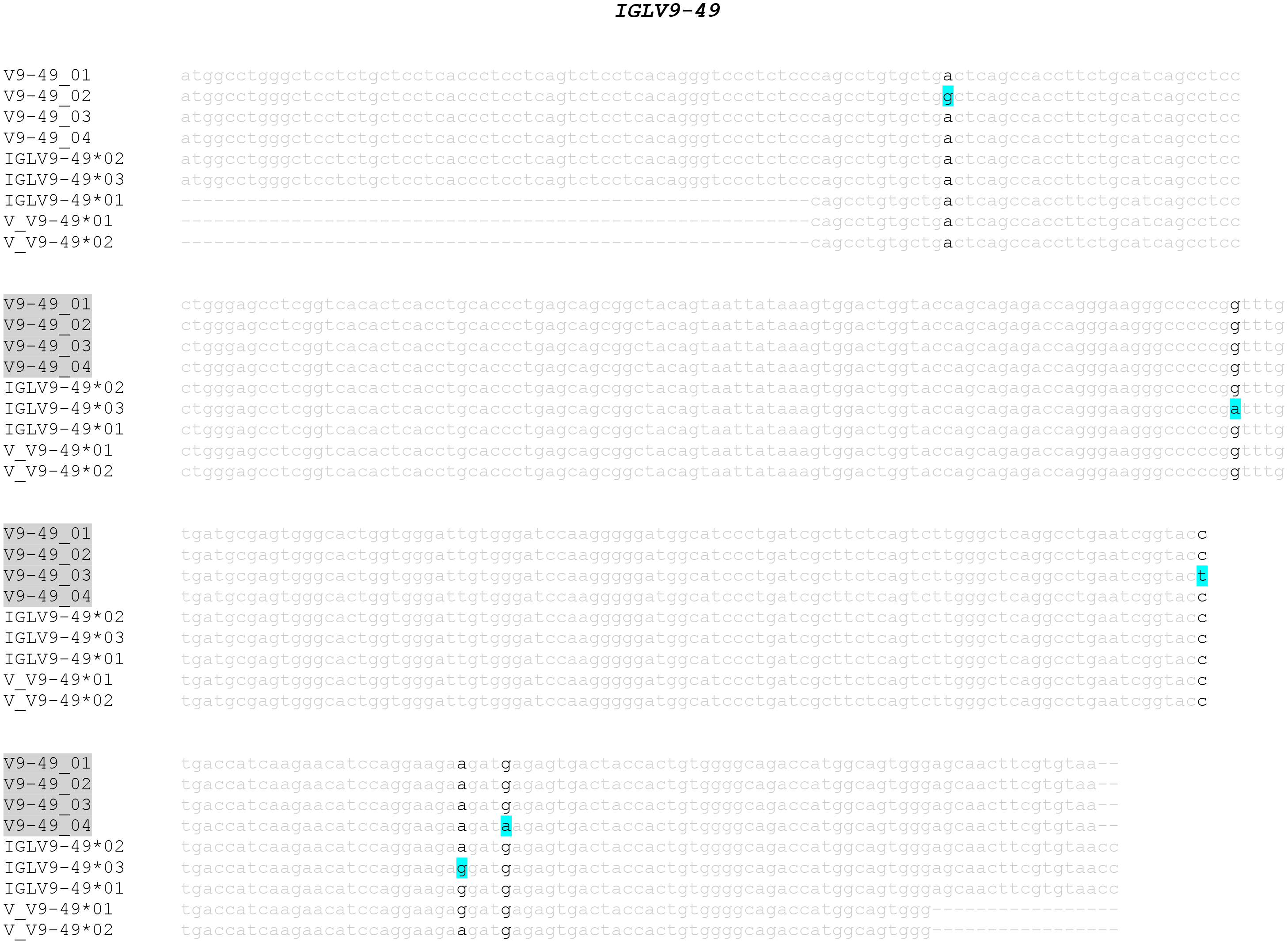
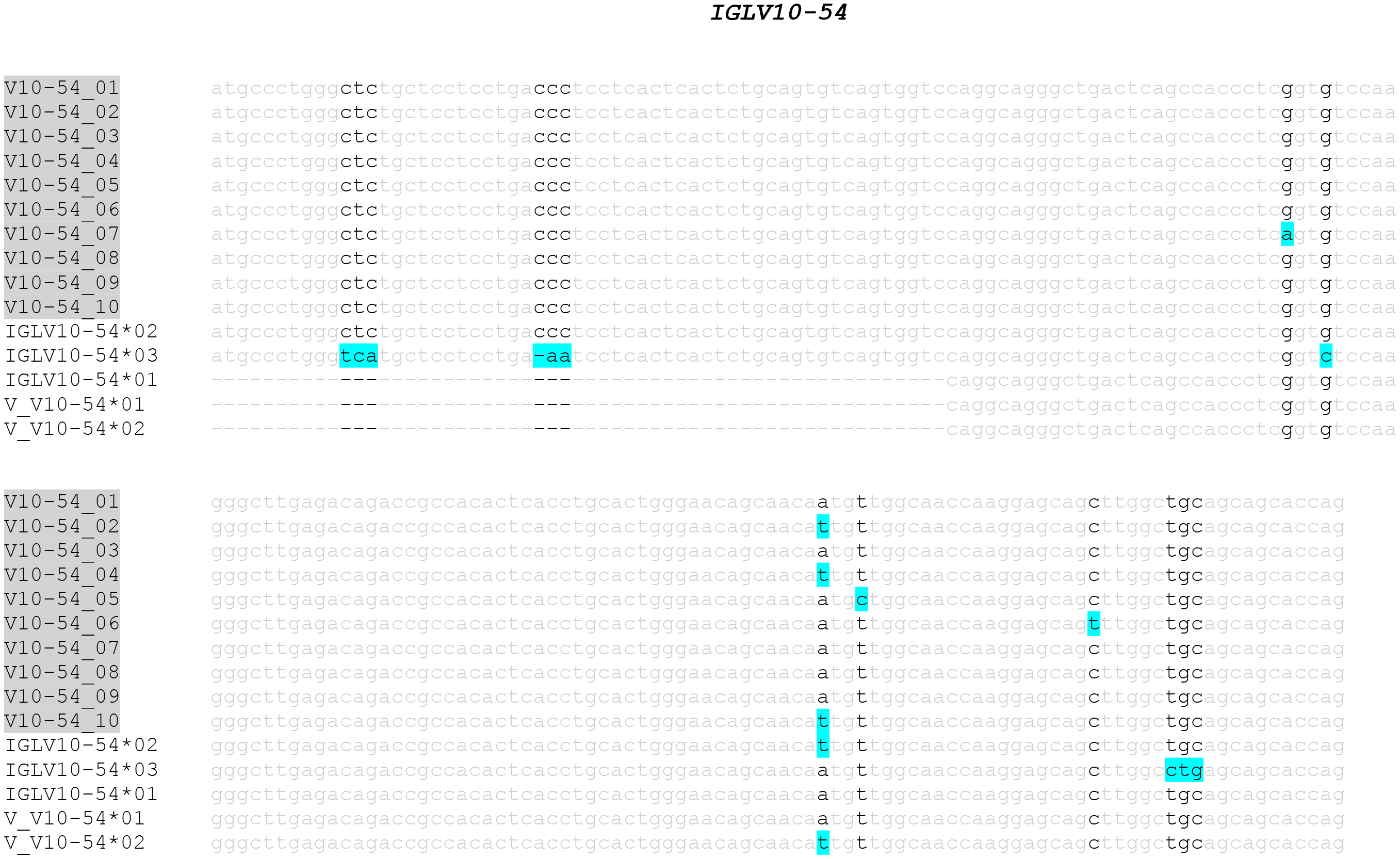
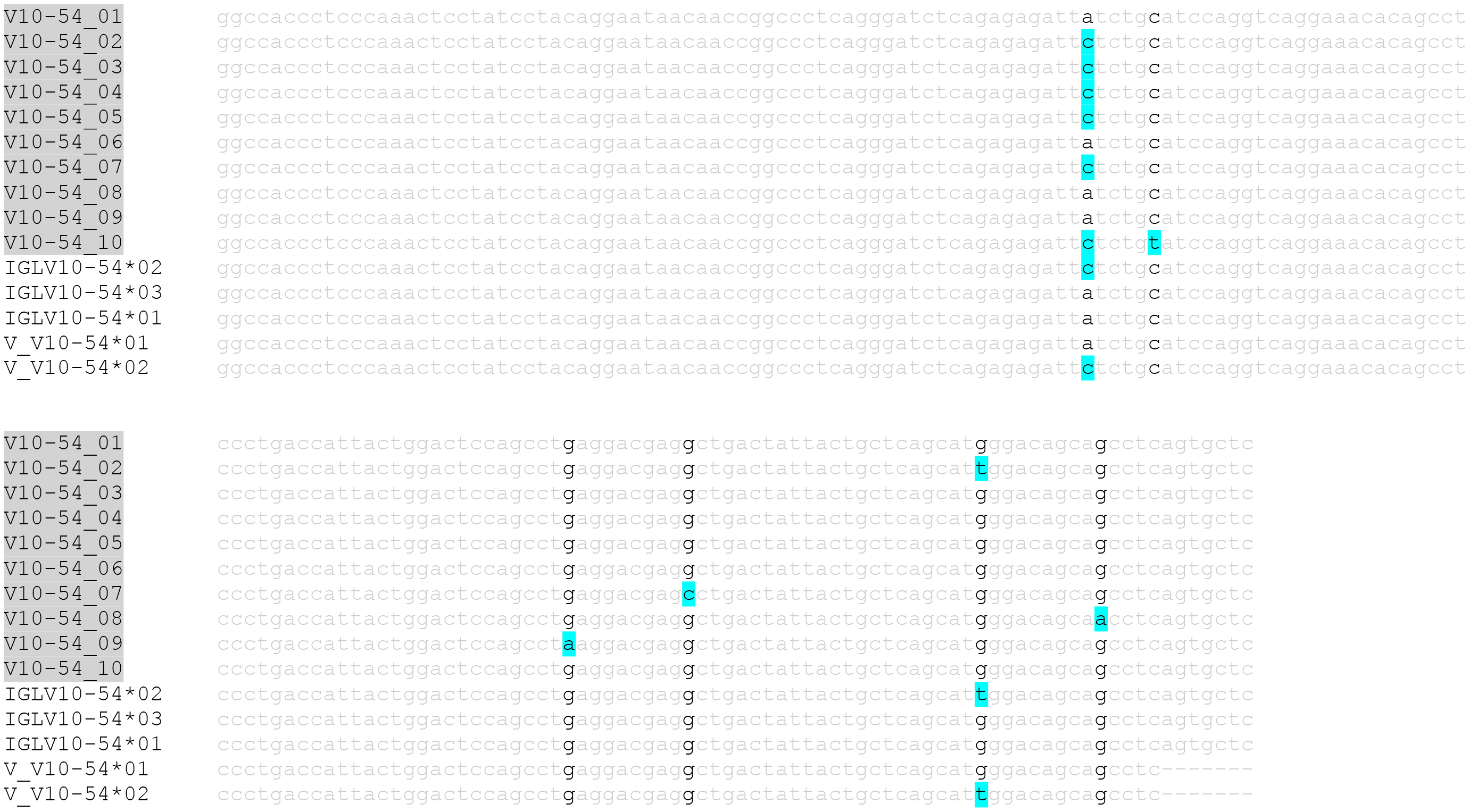
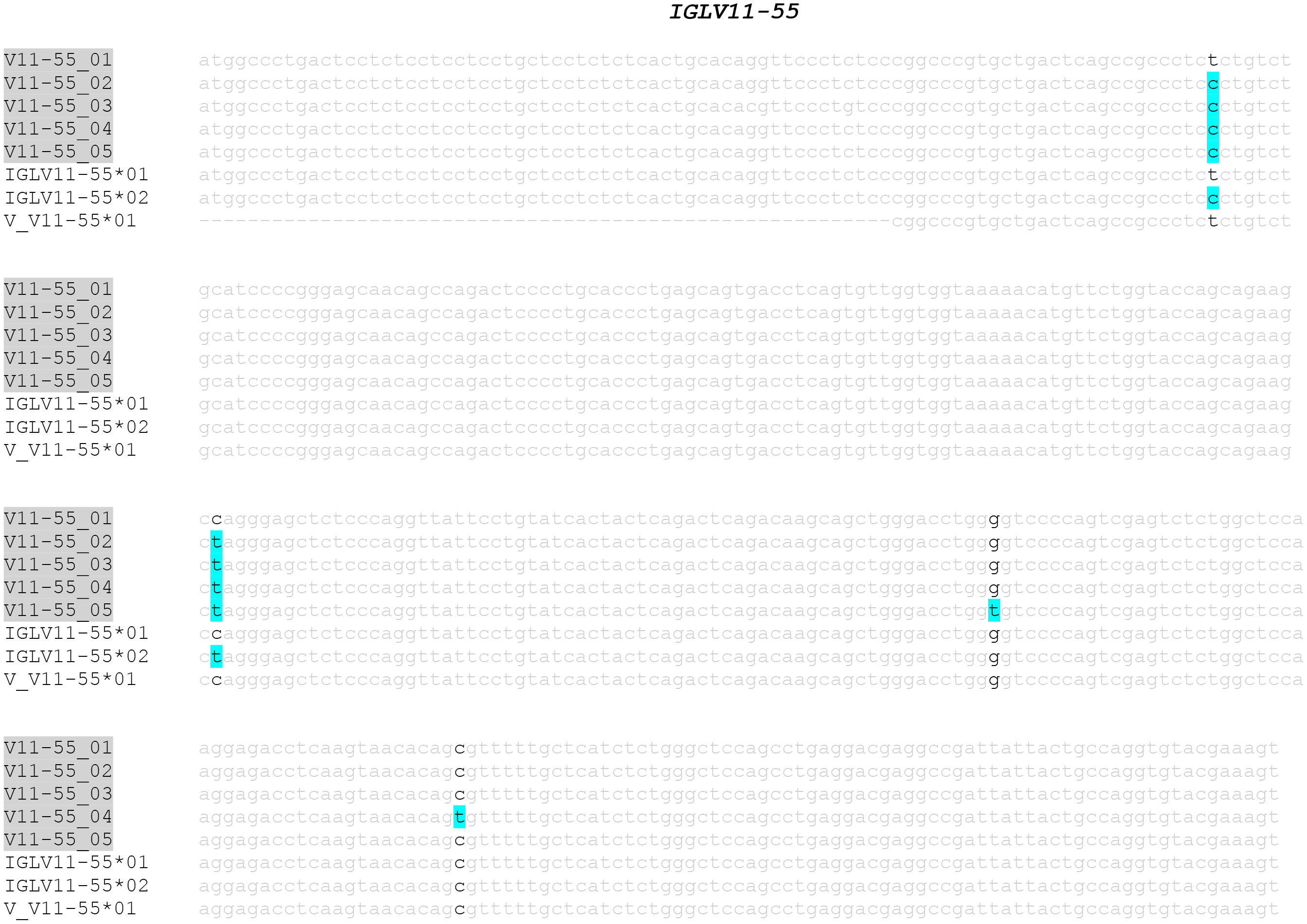
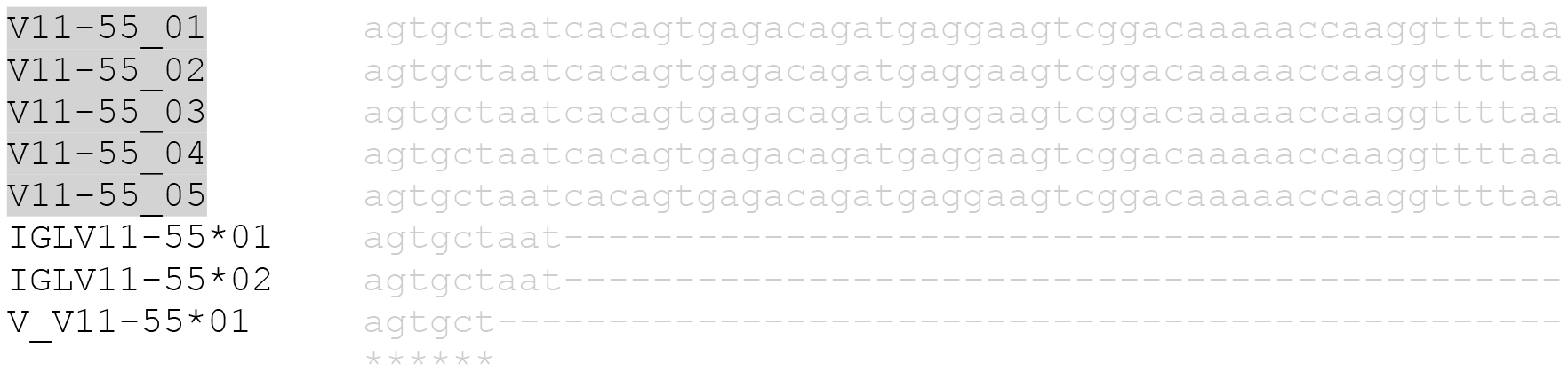
Per gene alignment of alleles from pmIG, IMGT, IgPdb and VBASE2 databases for *IGHV*, *IGKV* and *IGLV* locus. The name of pmIG alleles are marked with grey background. Yellow background is used for the 104 IMGT alleles mentioned to be erroneous/false positives by Wang et al, 2008.

**Figure S3:**
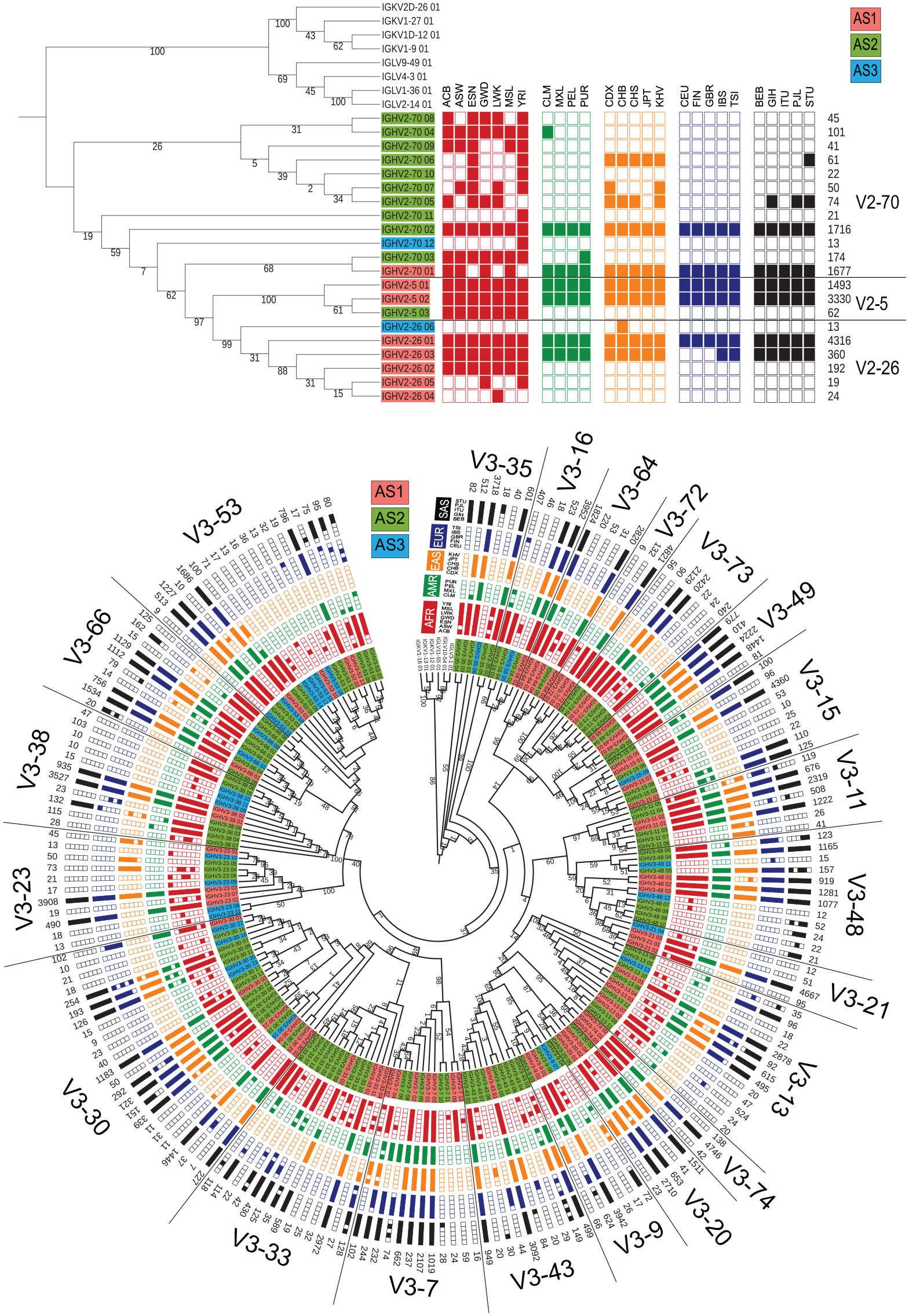
ML tree of the population distribution of *IGHV2 and V3* family alleles.

**Figure S4:**
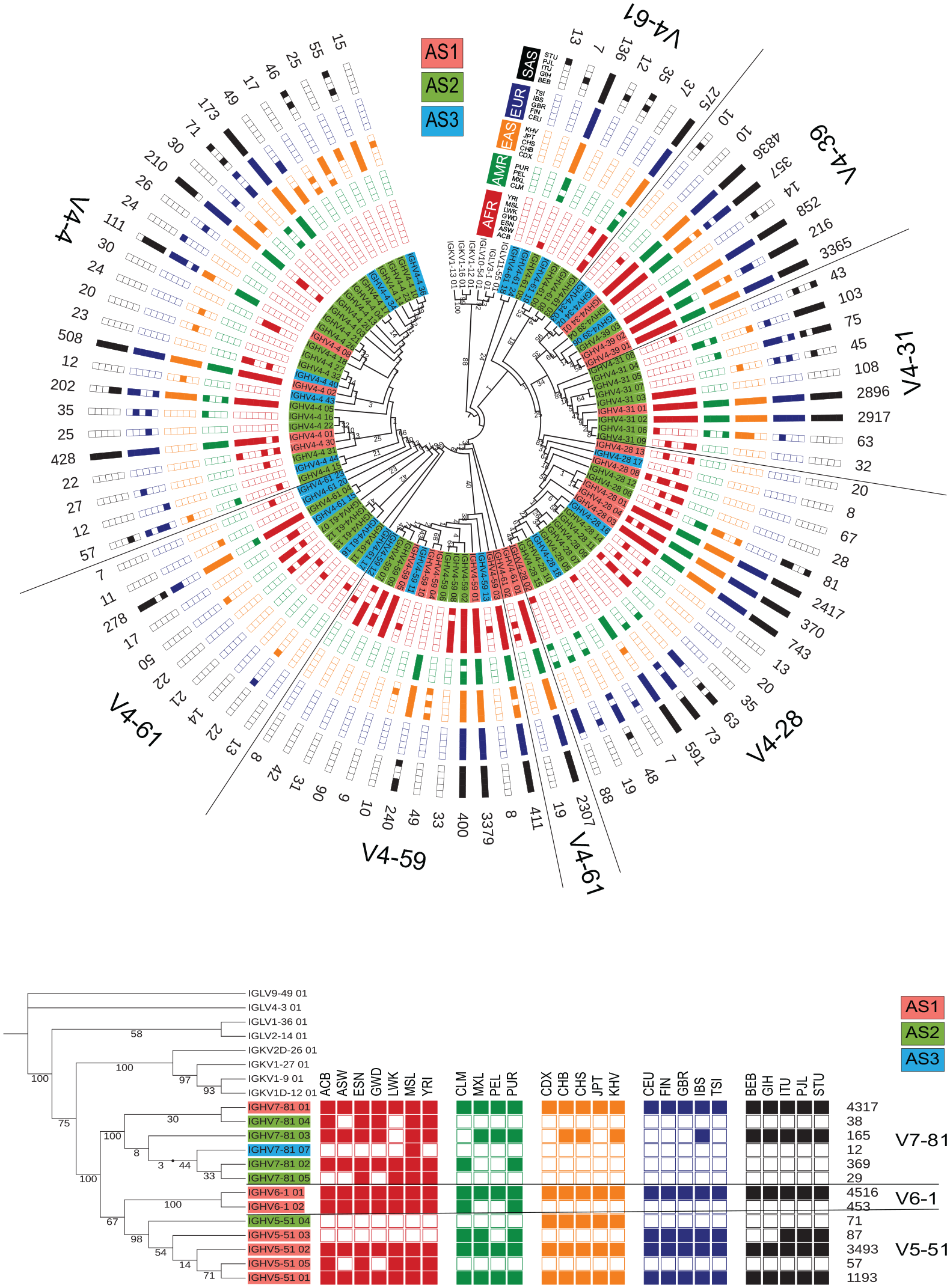
ML tree of the population distribution of *IGHV4, V5, V6* and *V7* family alleles.

**Figure S5:**
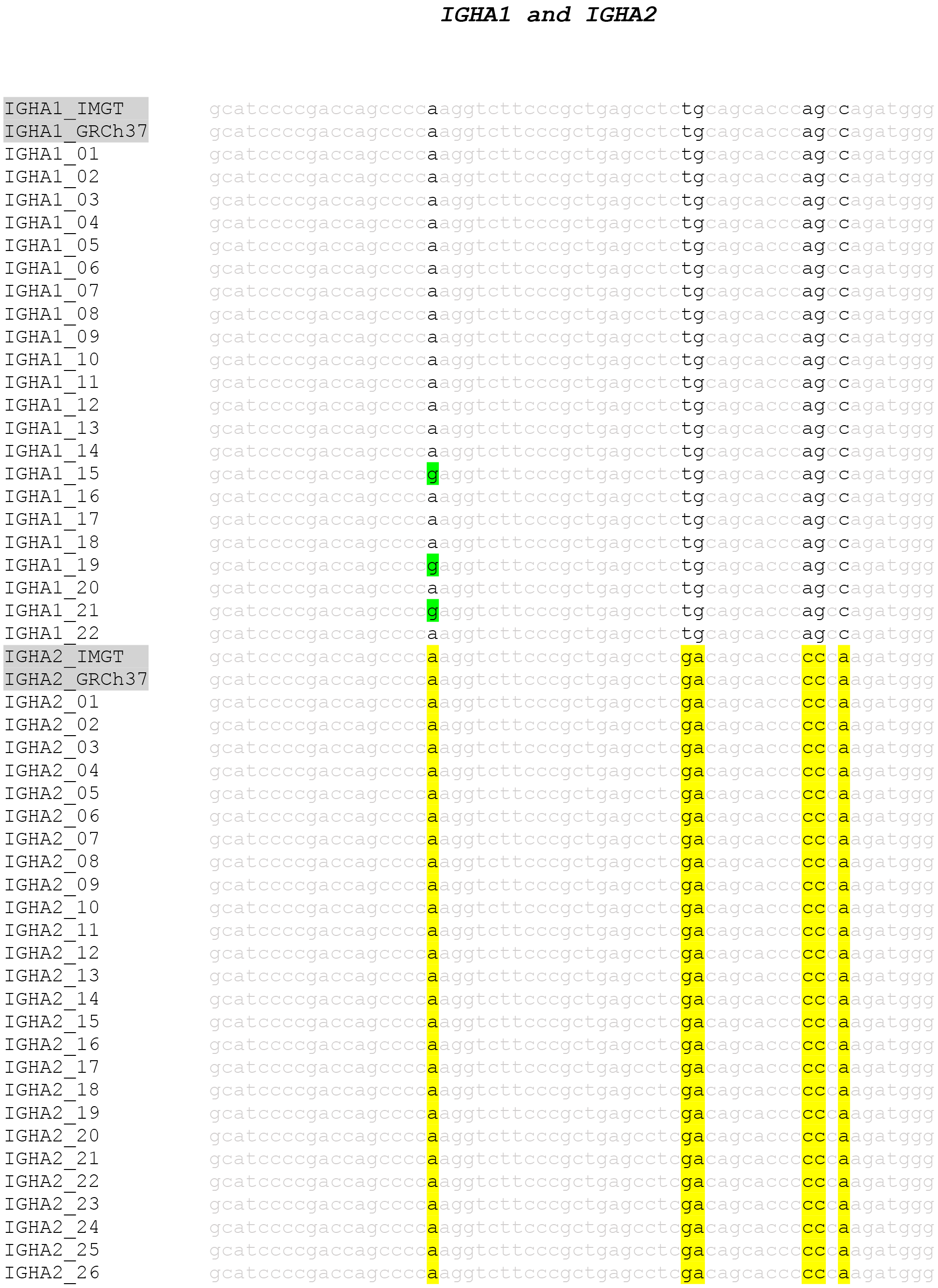
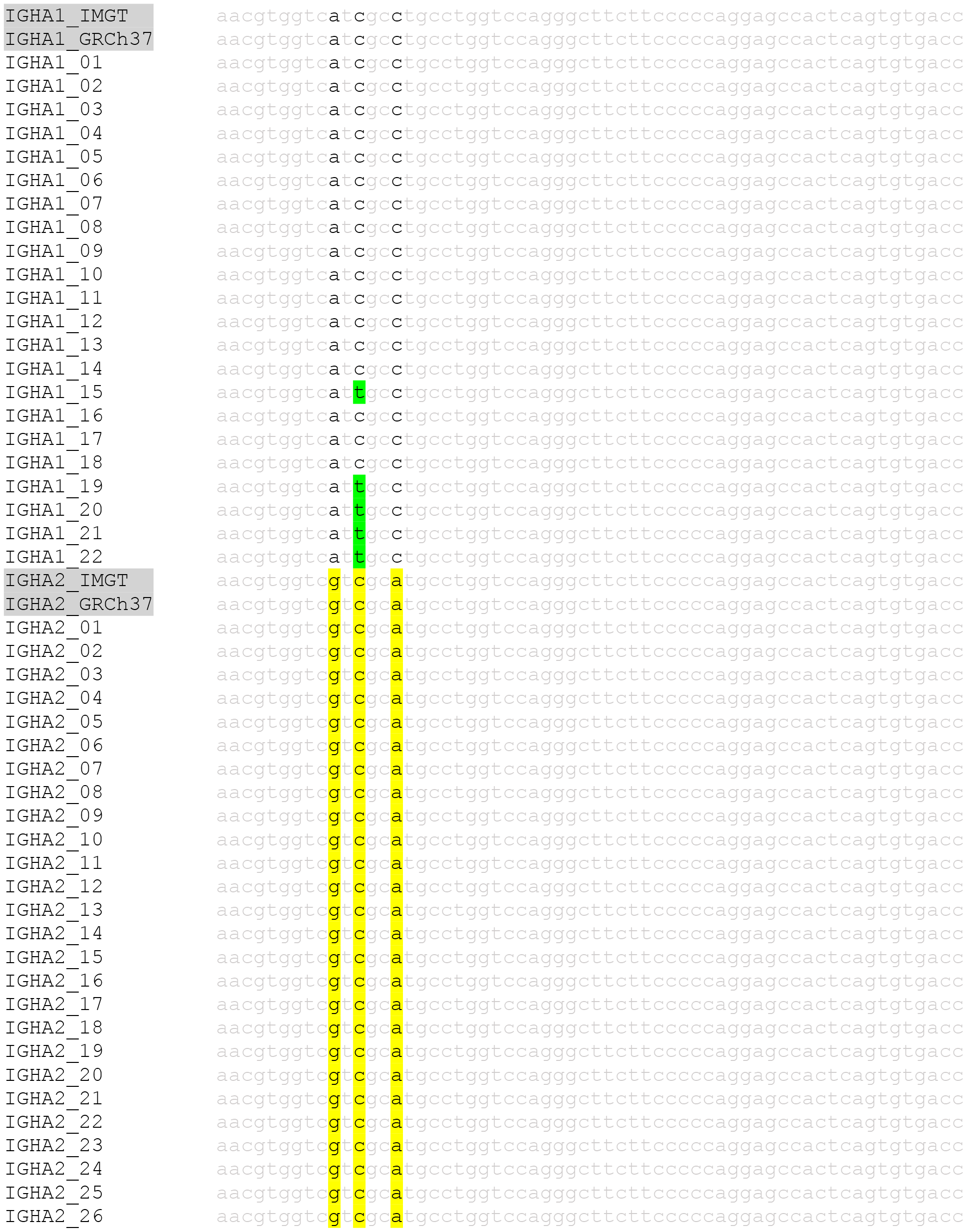
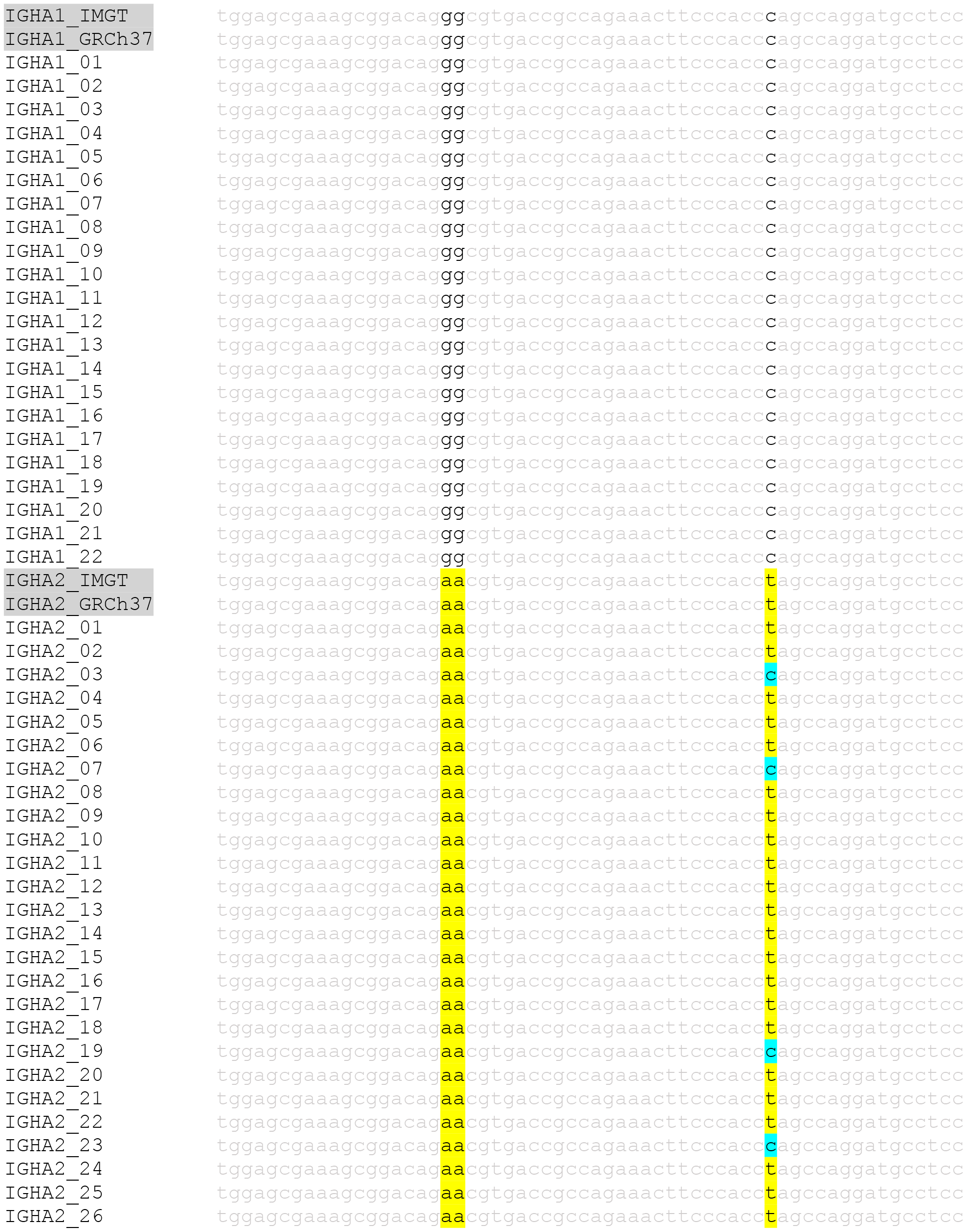
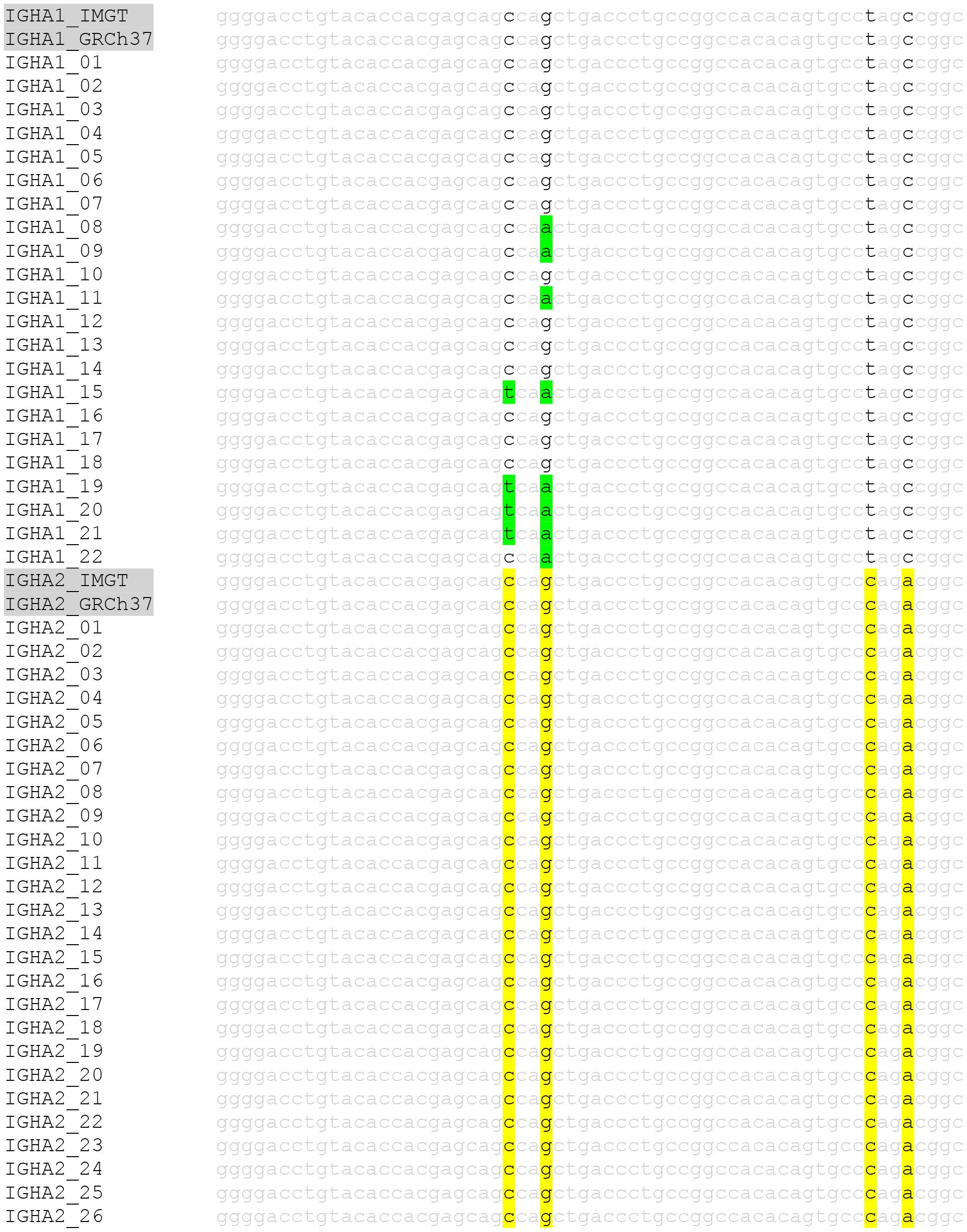
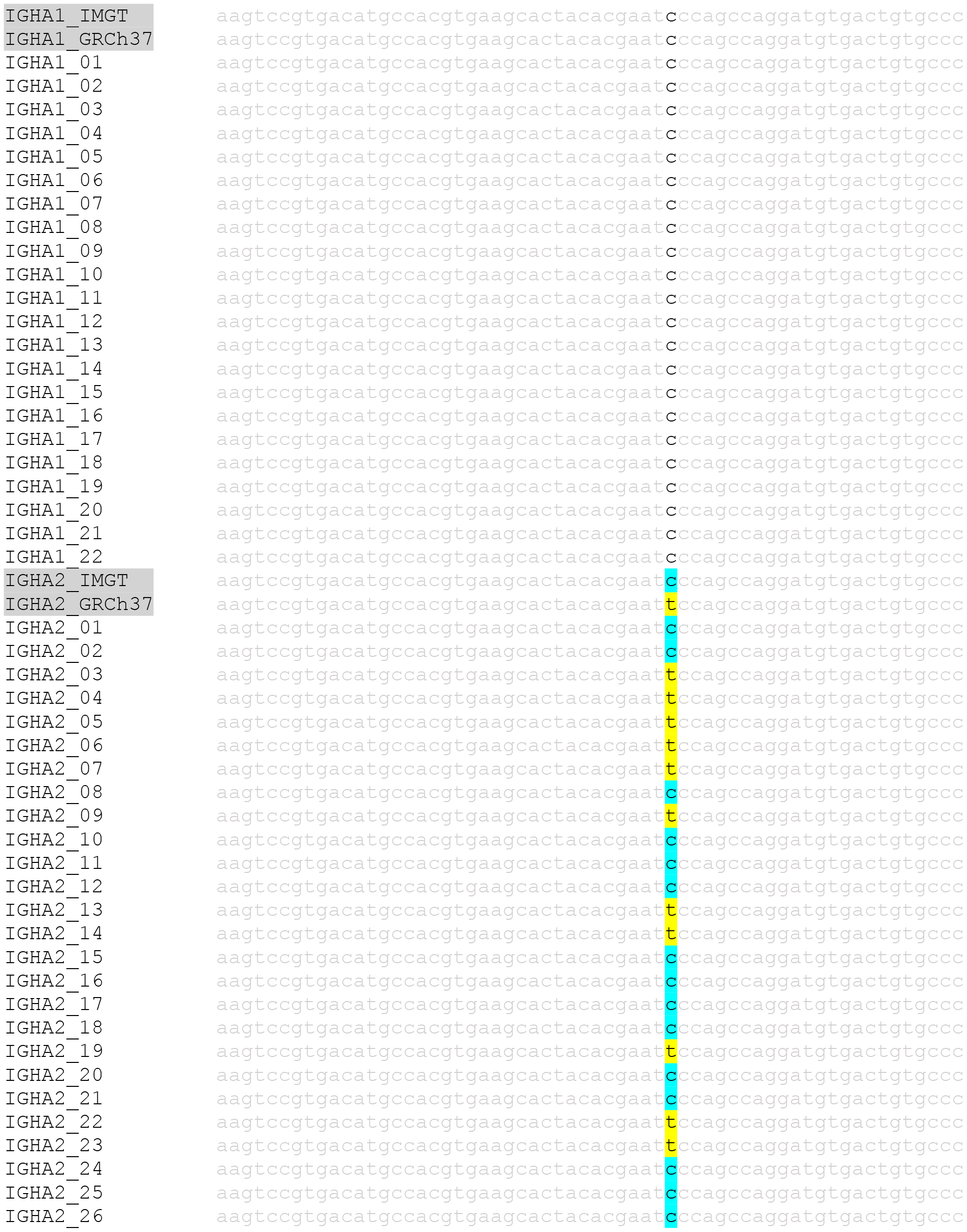
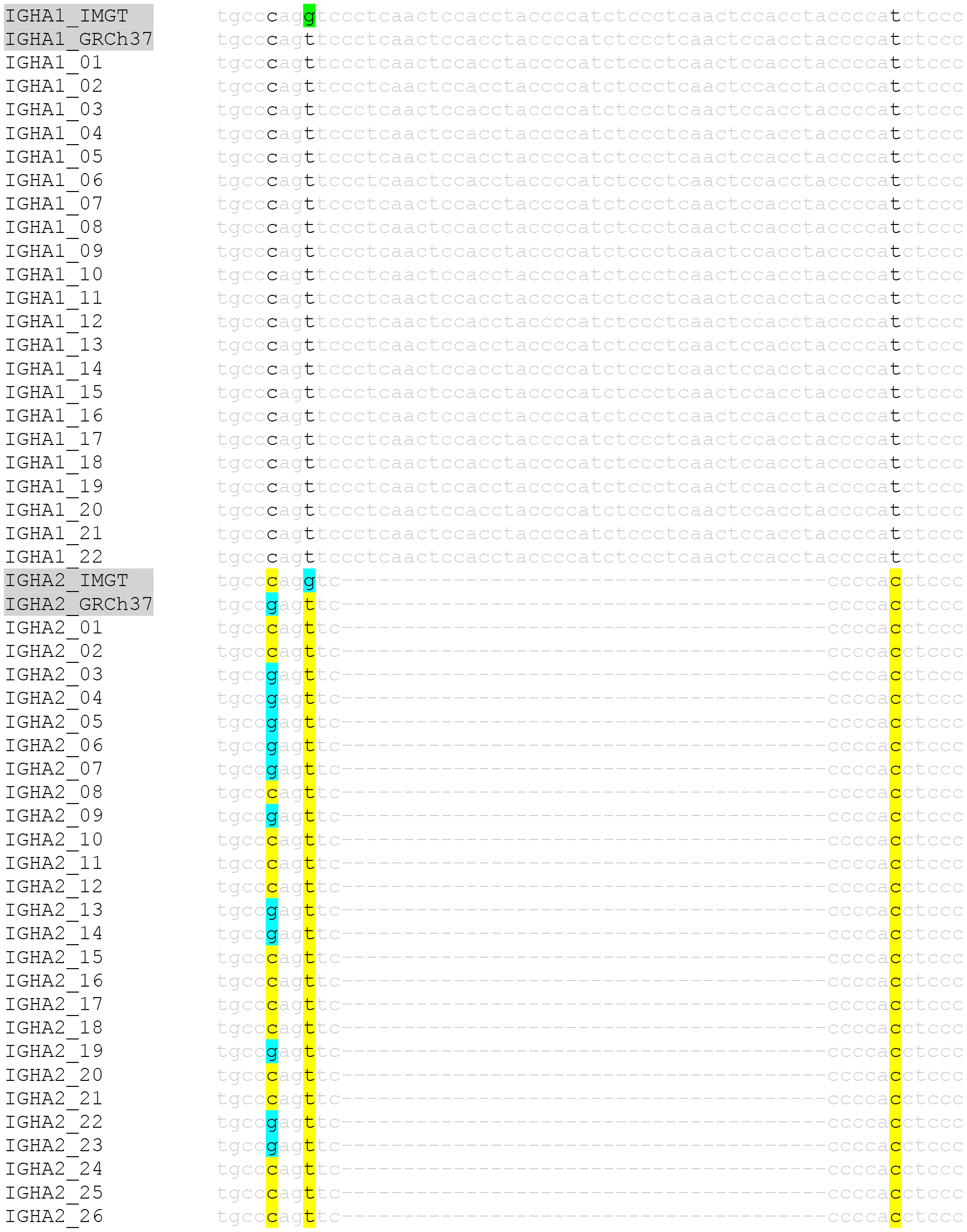
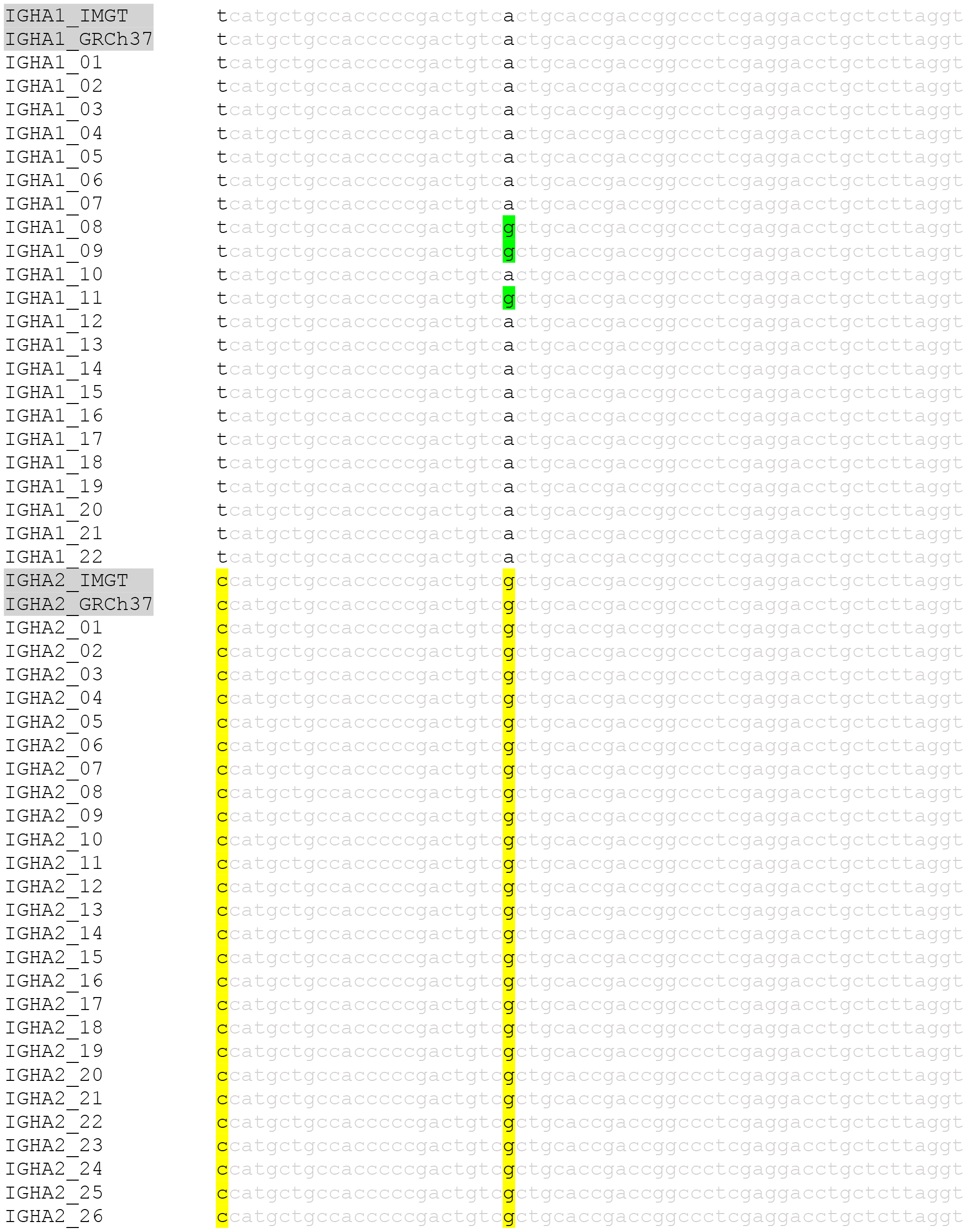
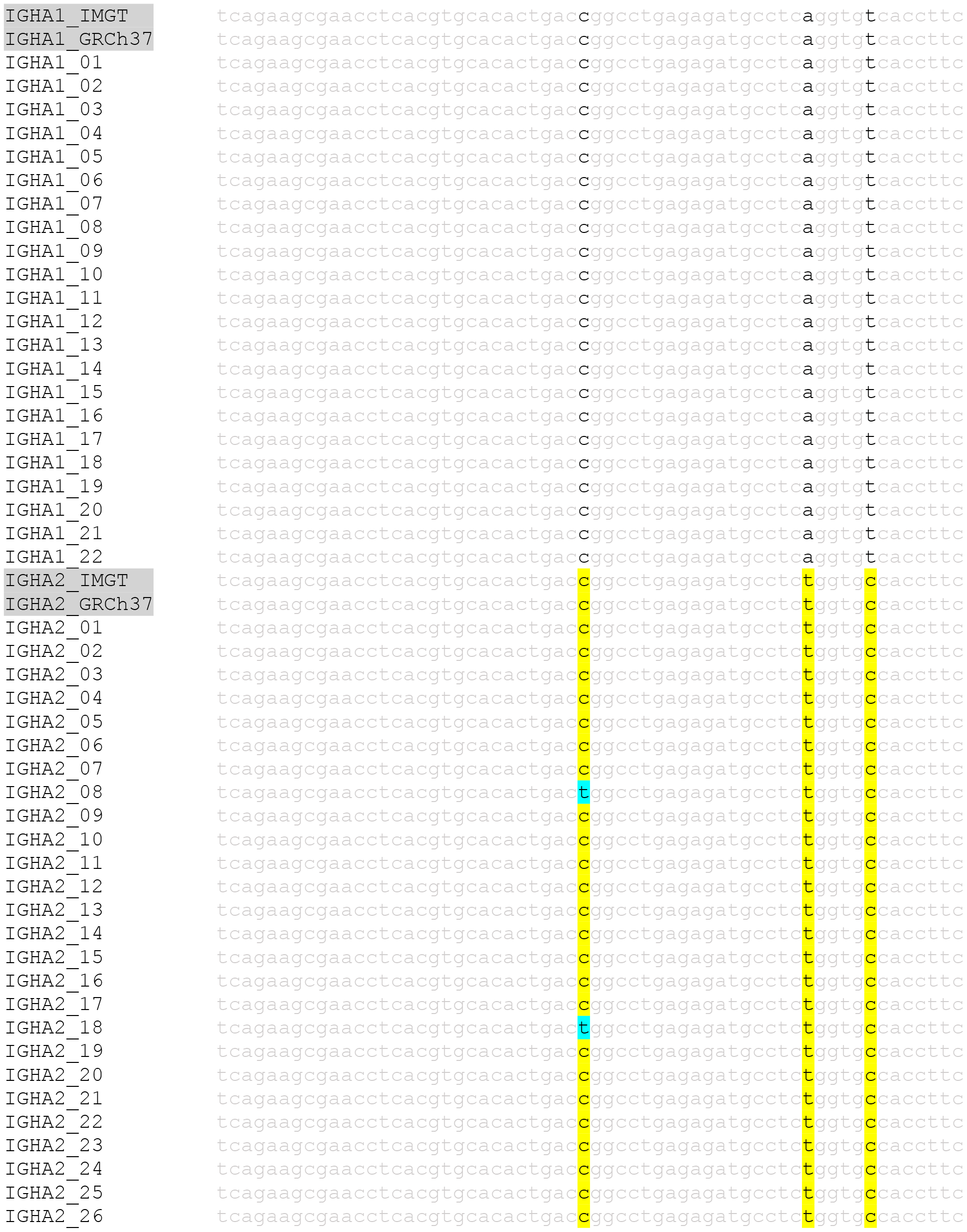
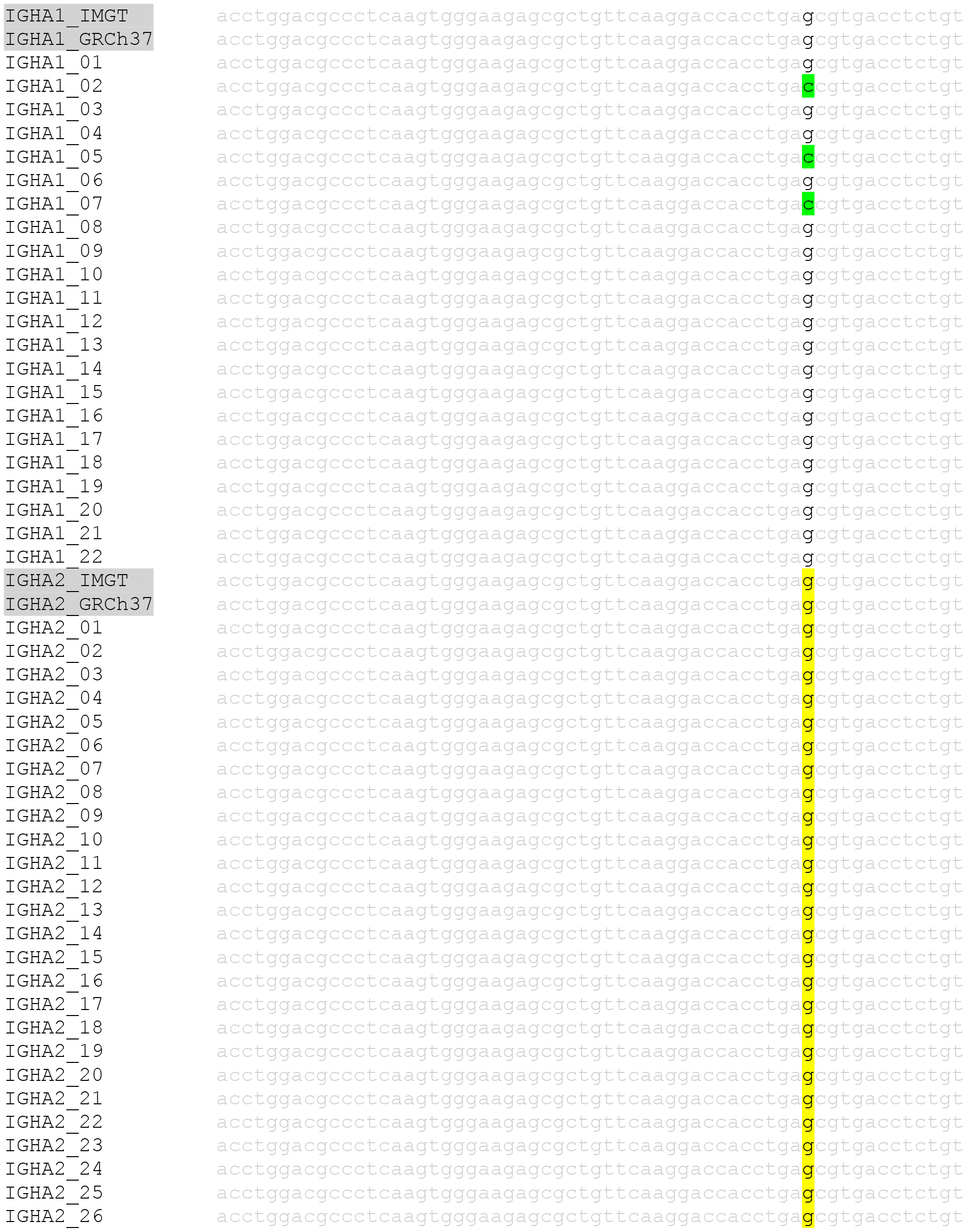
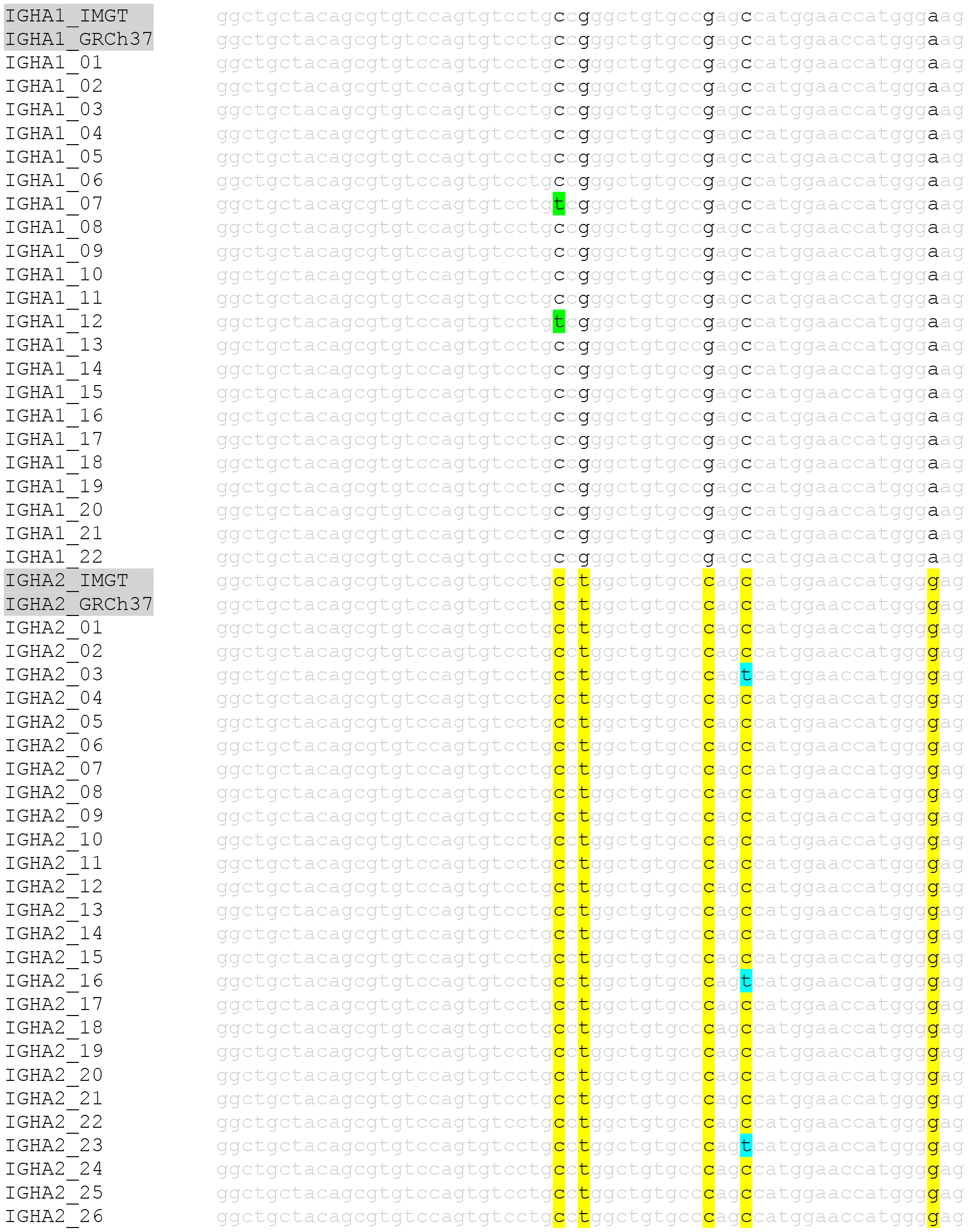
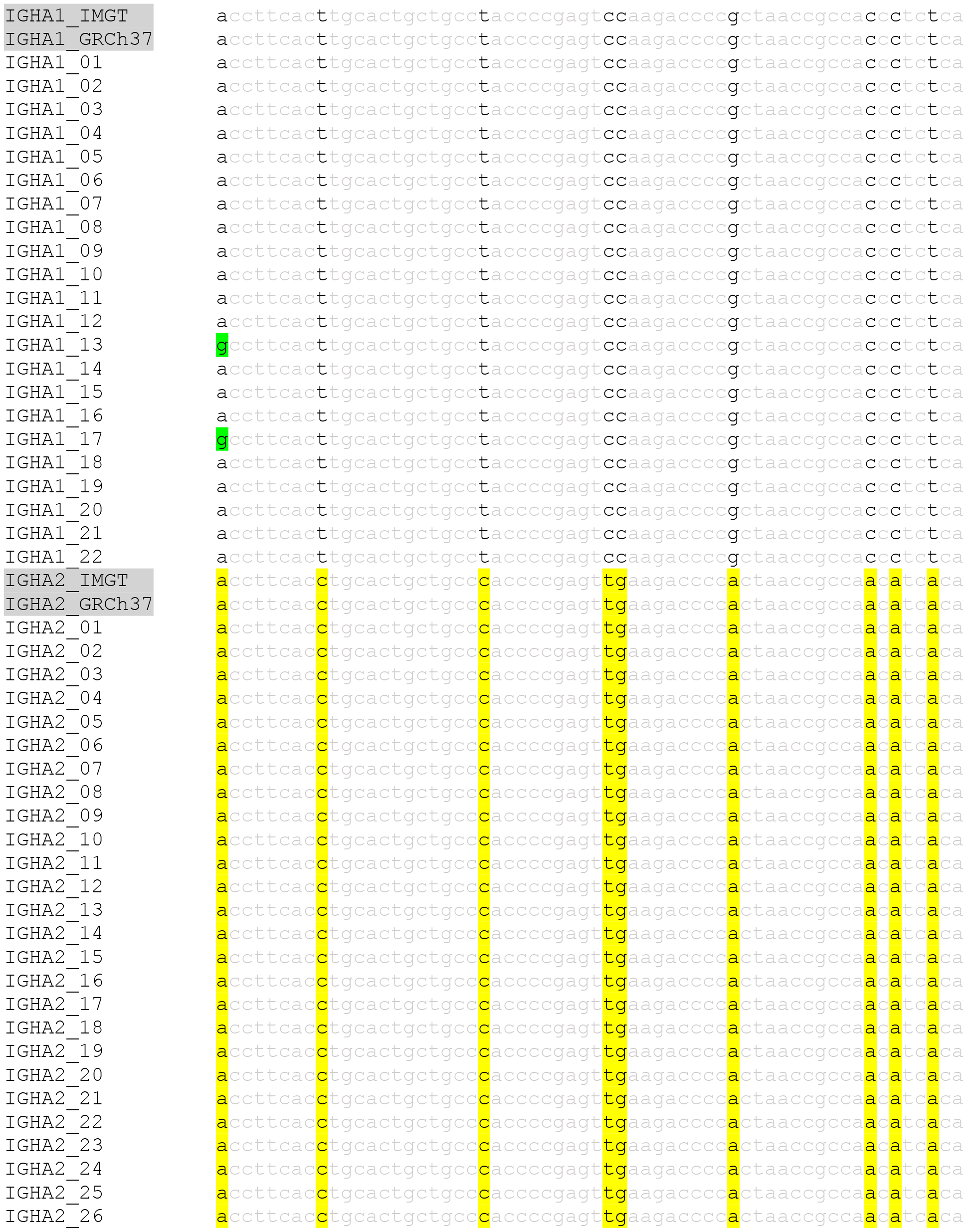
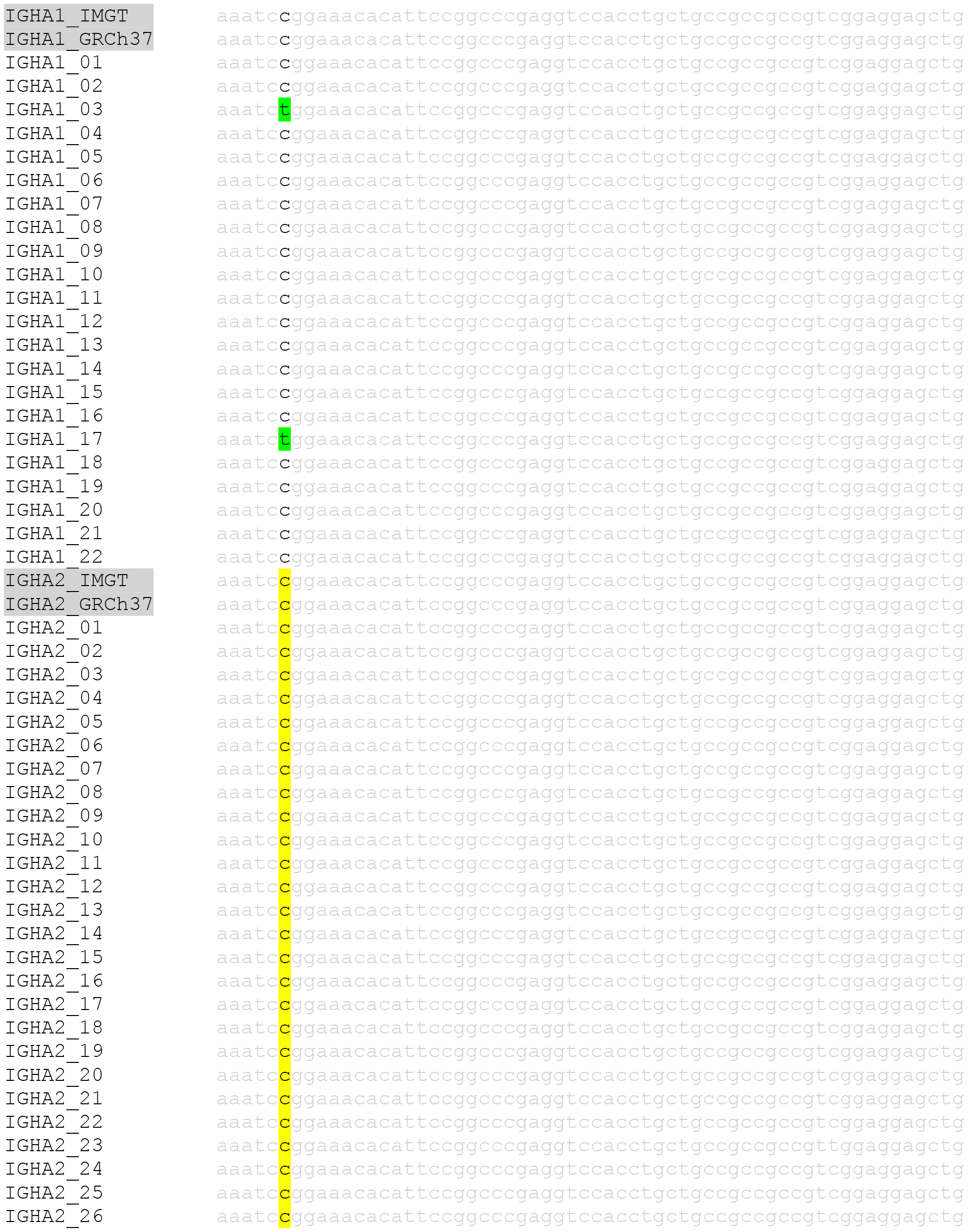
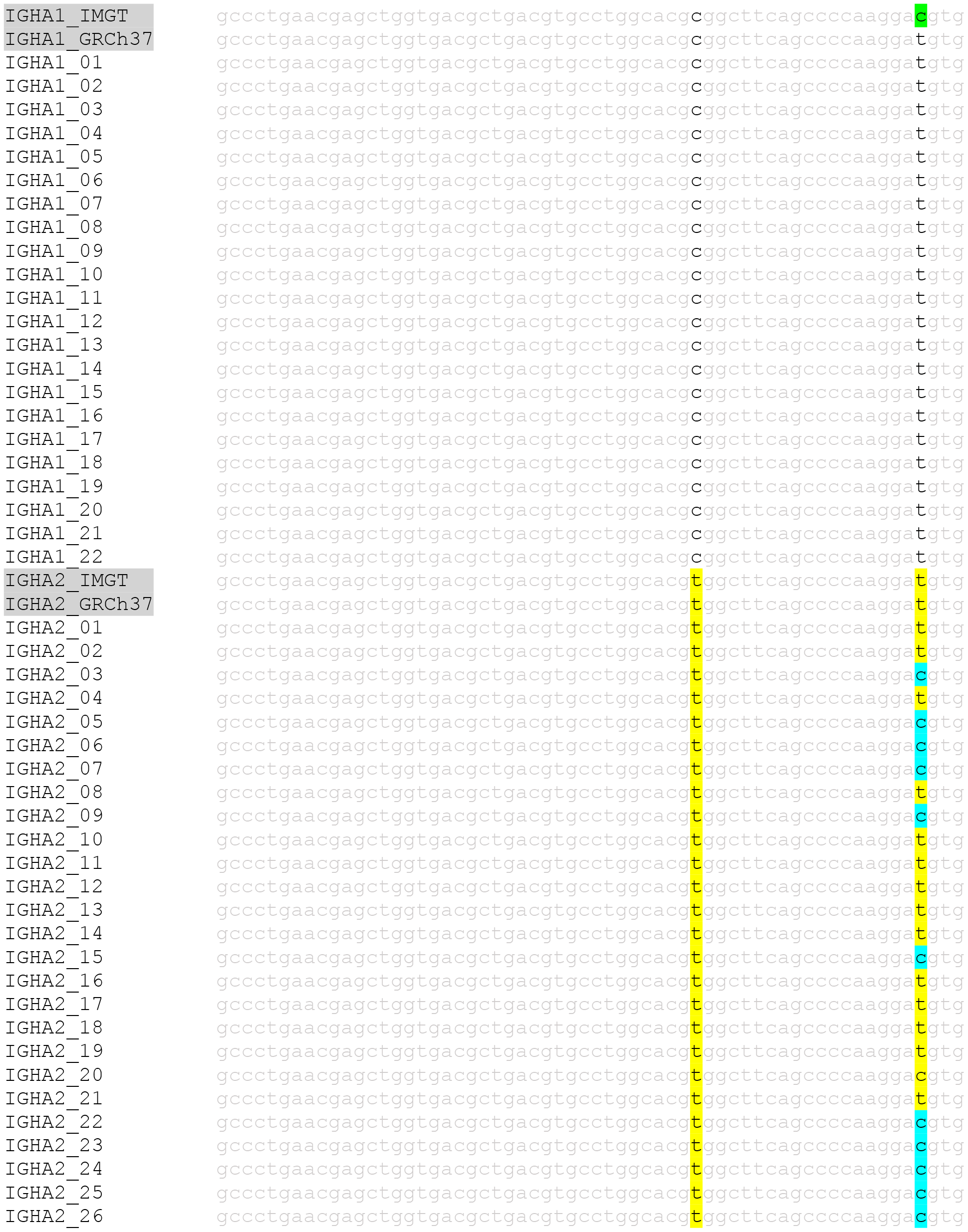
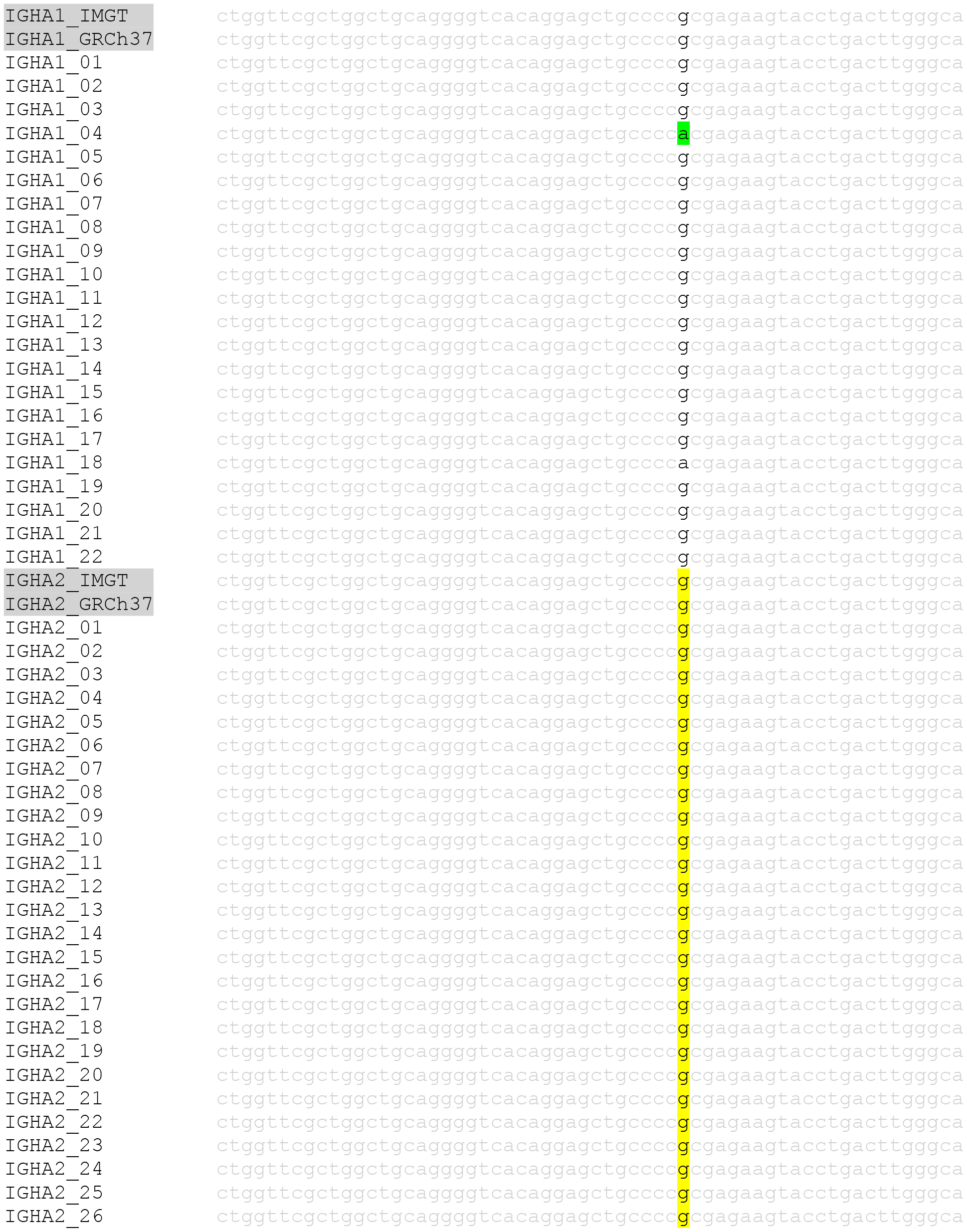
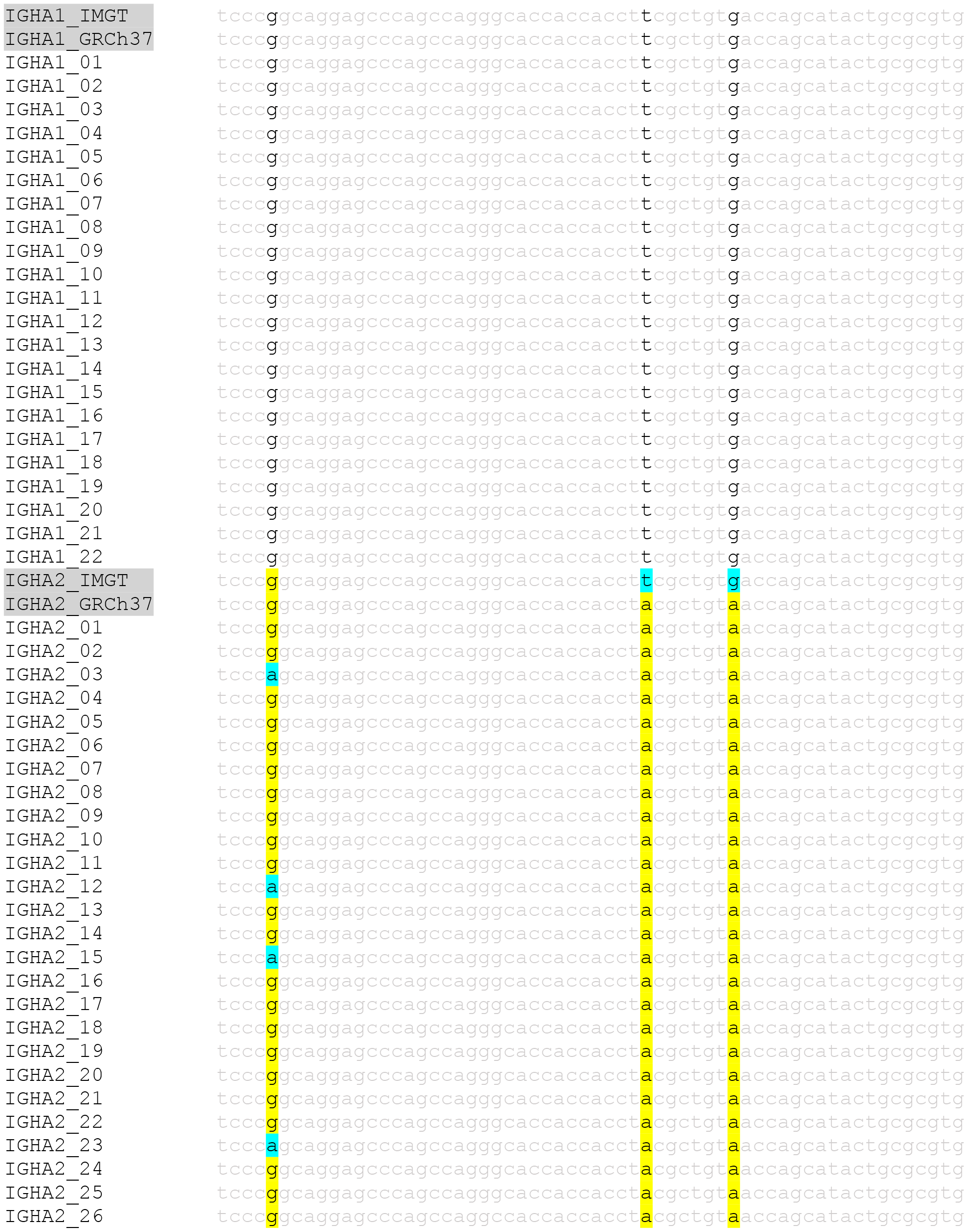
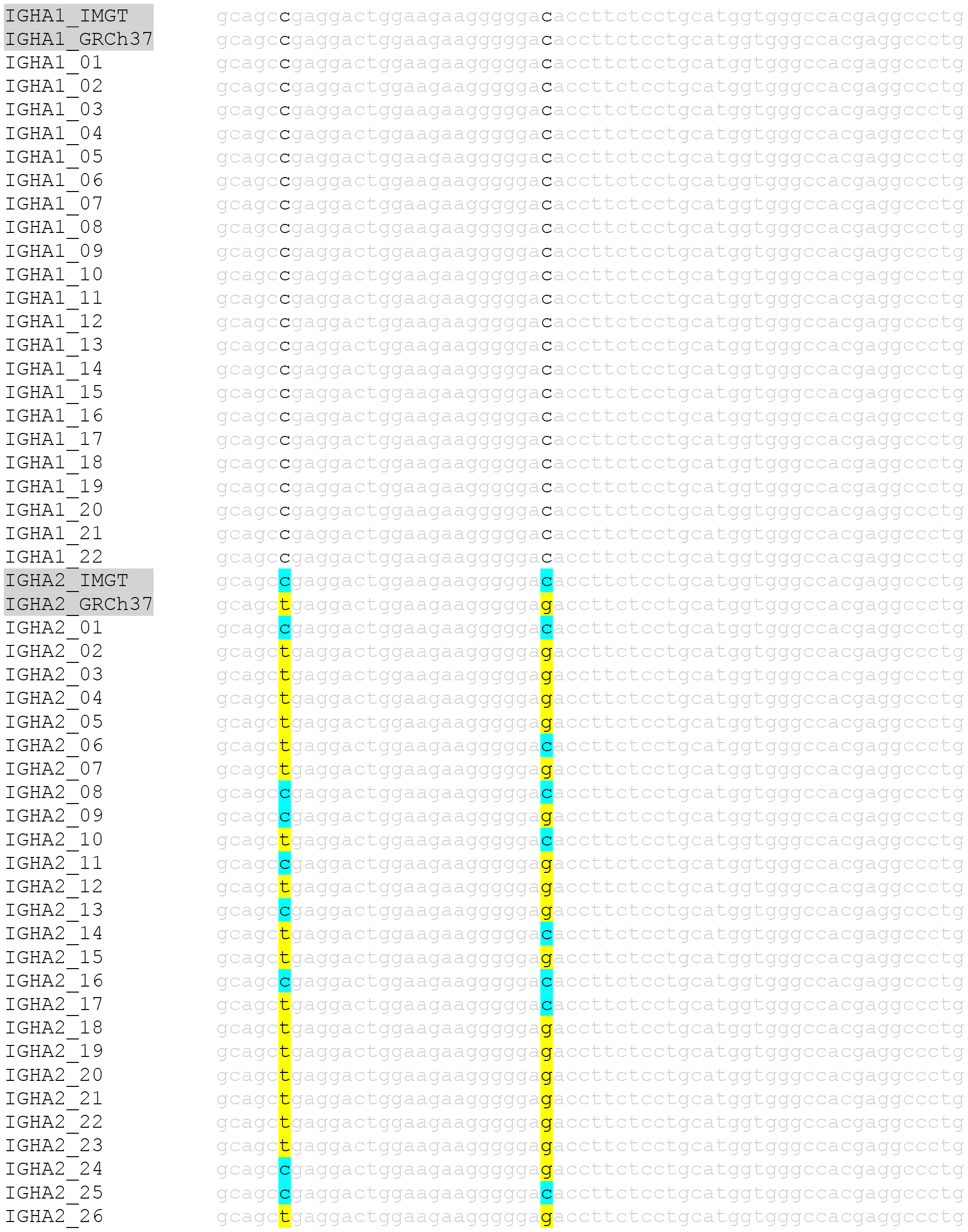
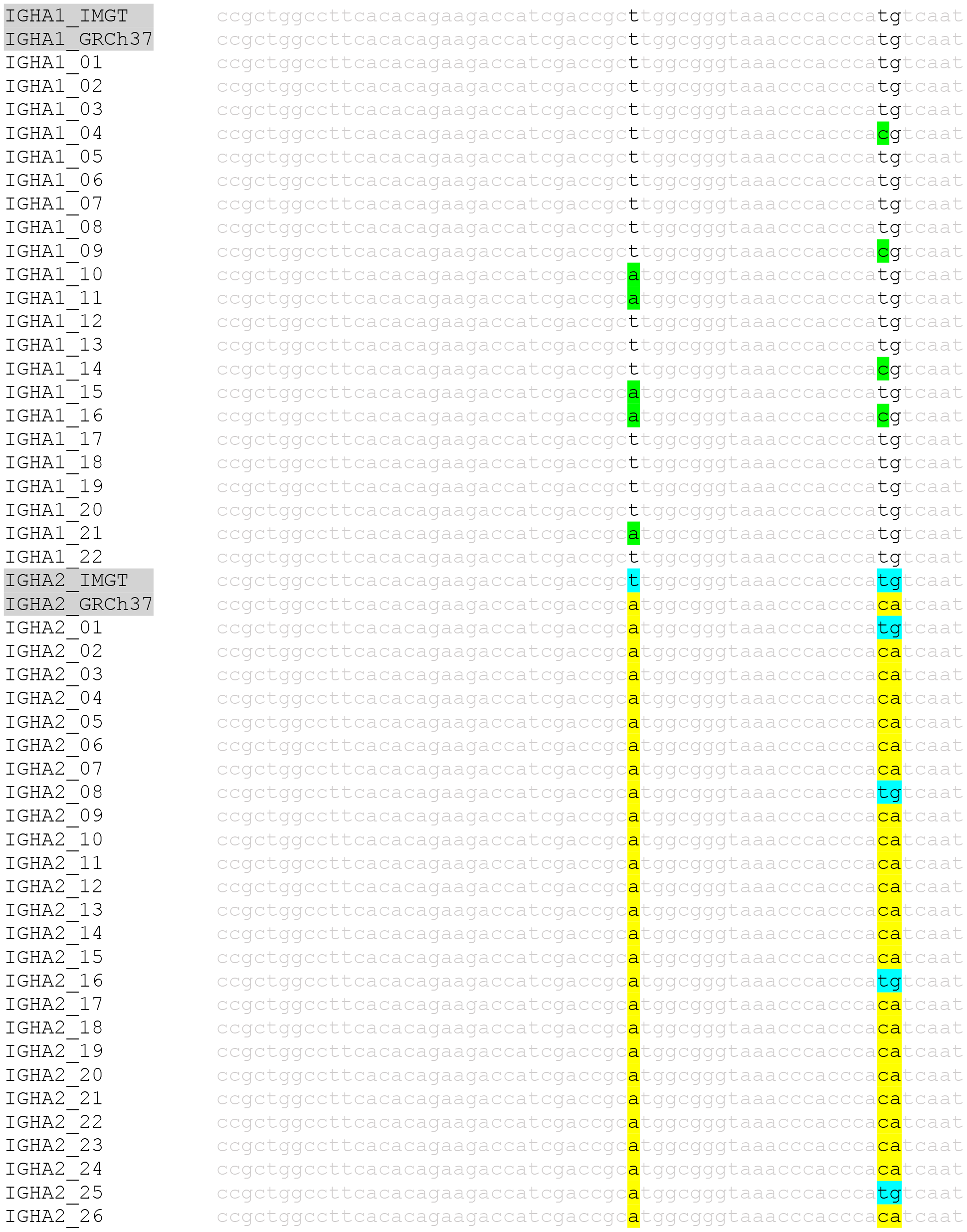
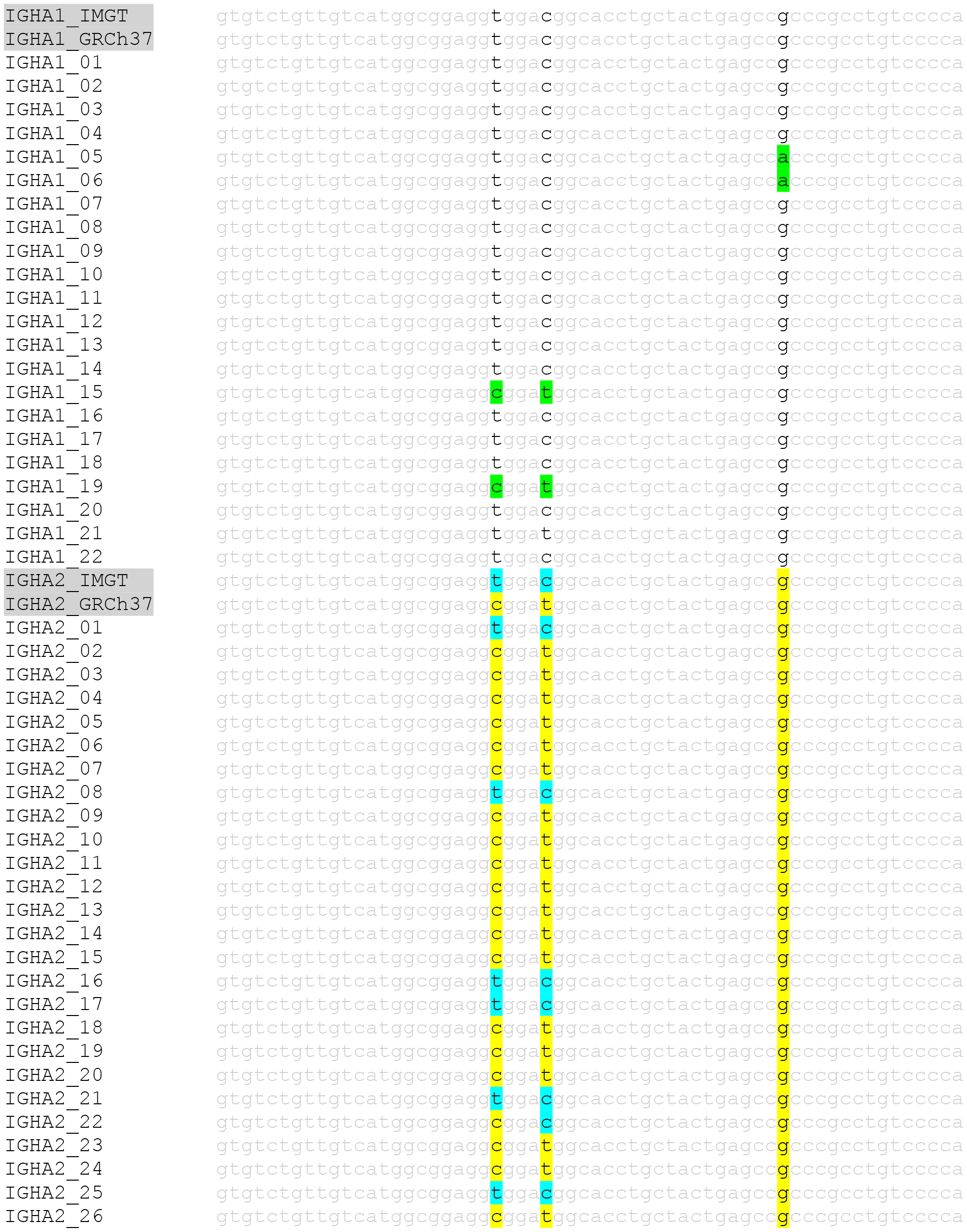
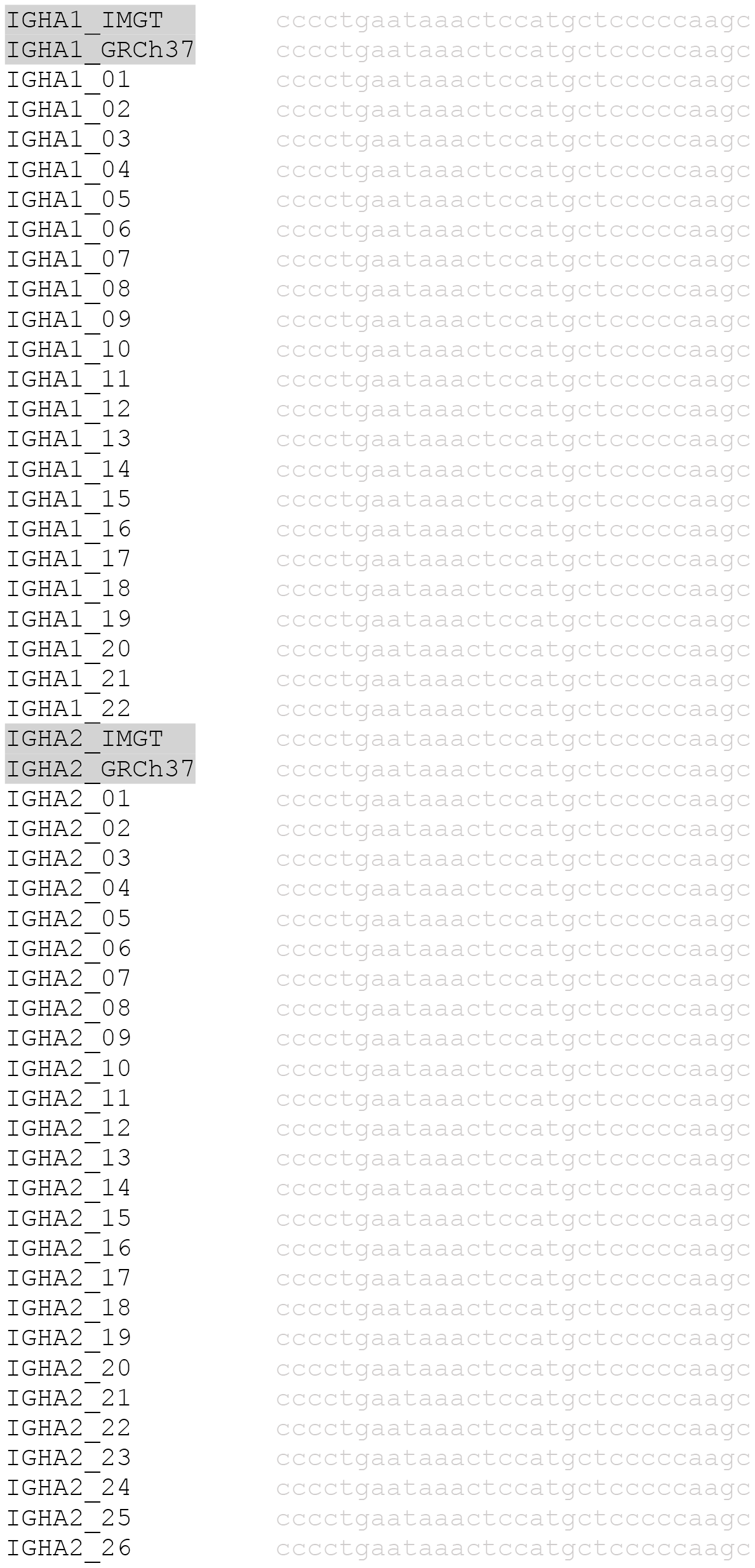
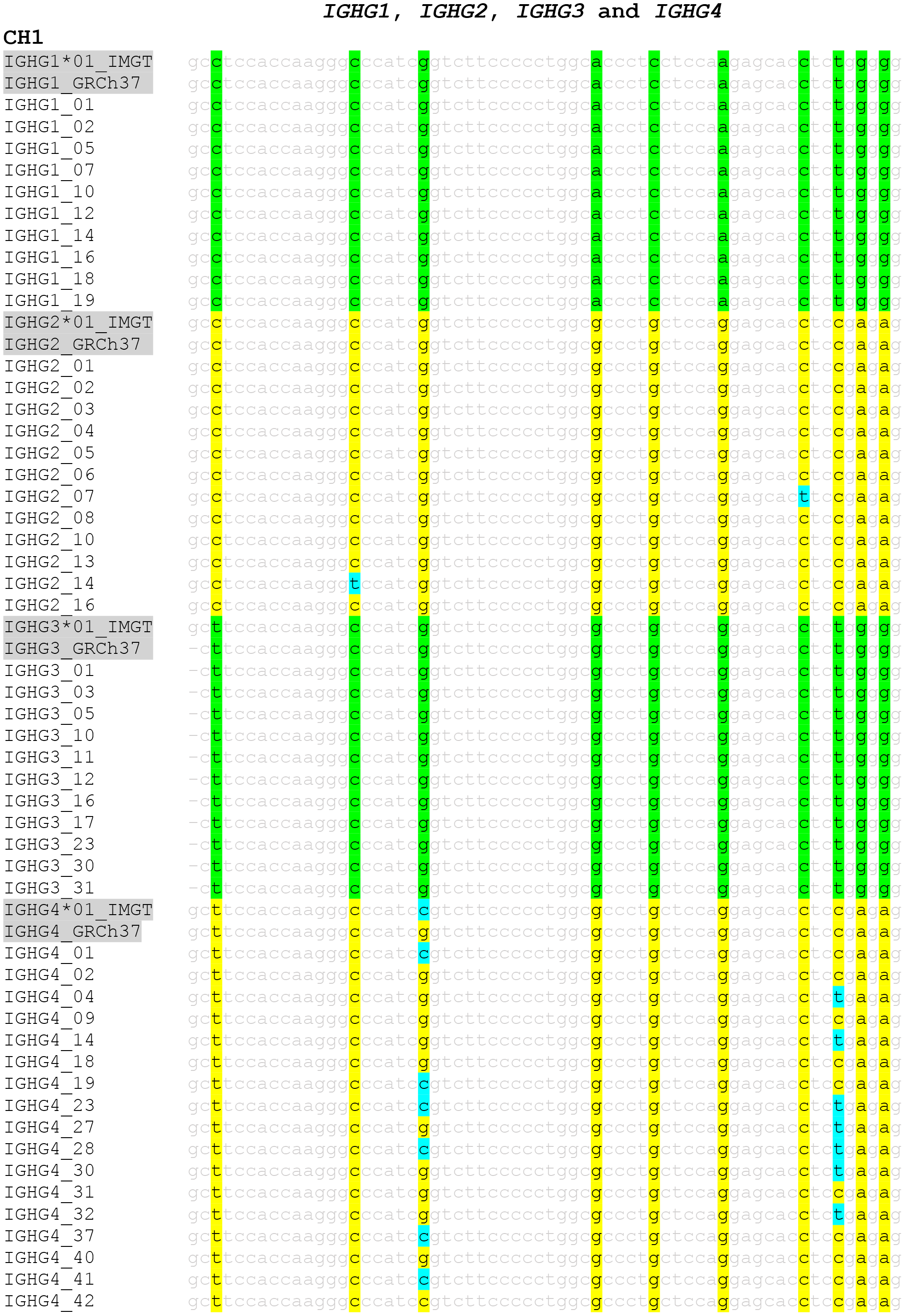
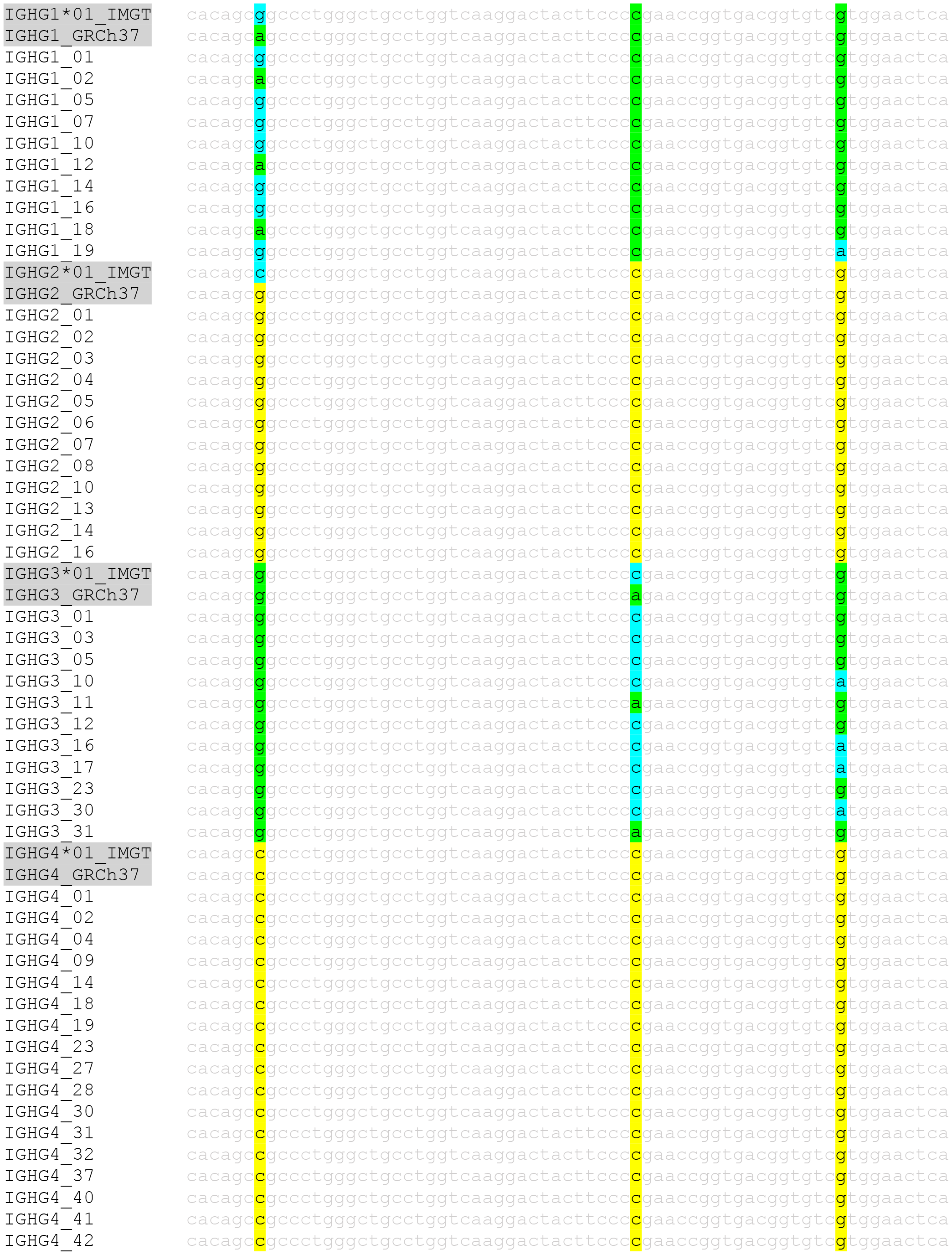
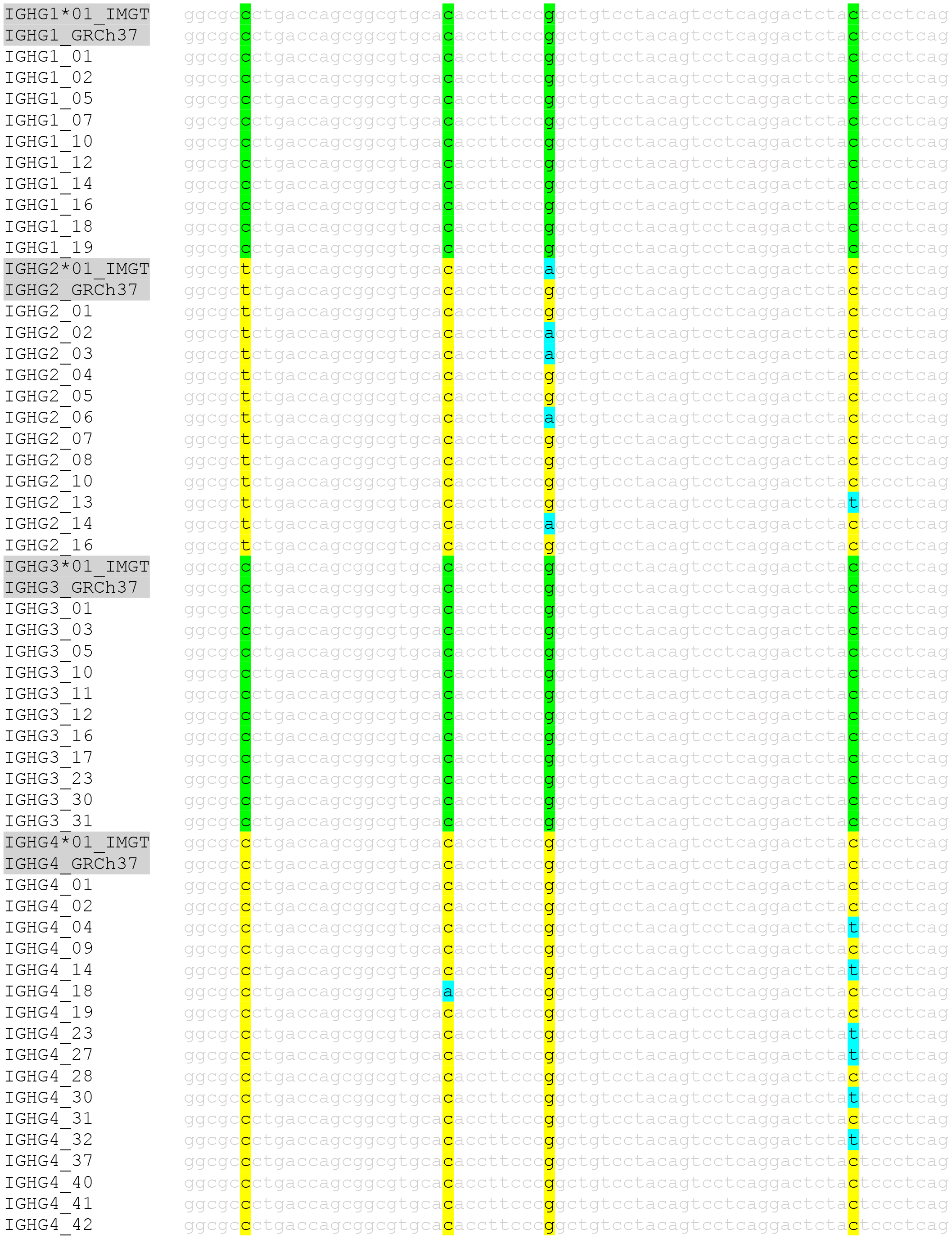
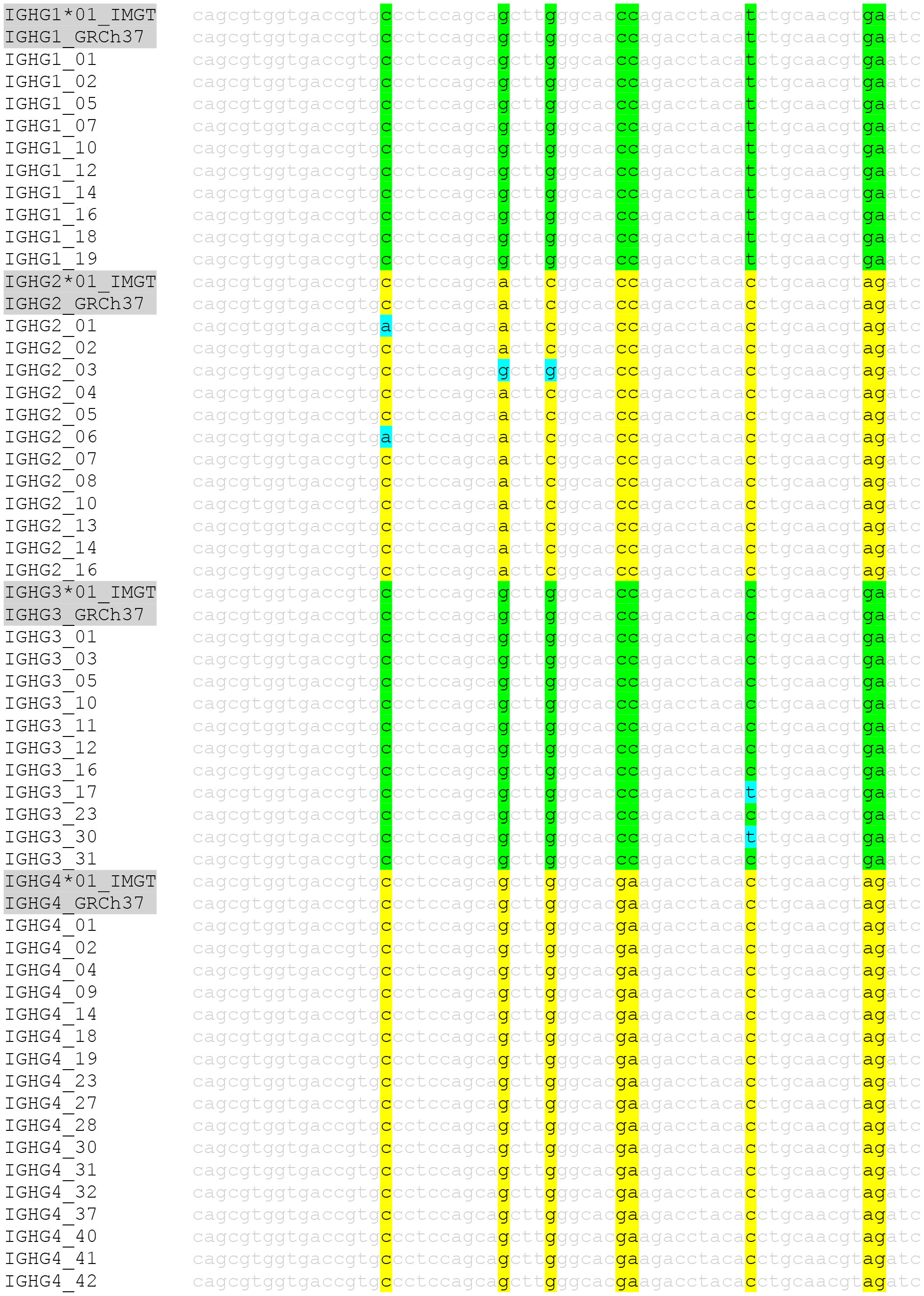
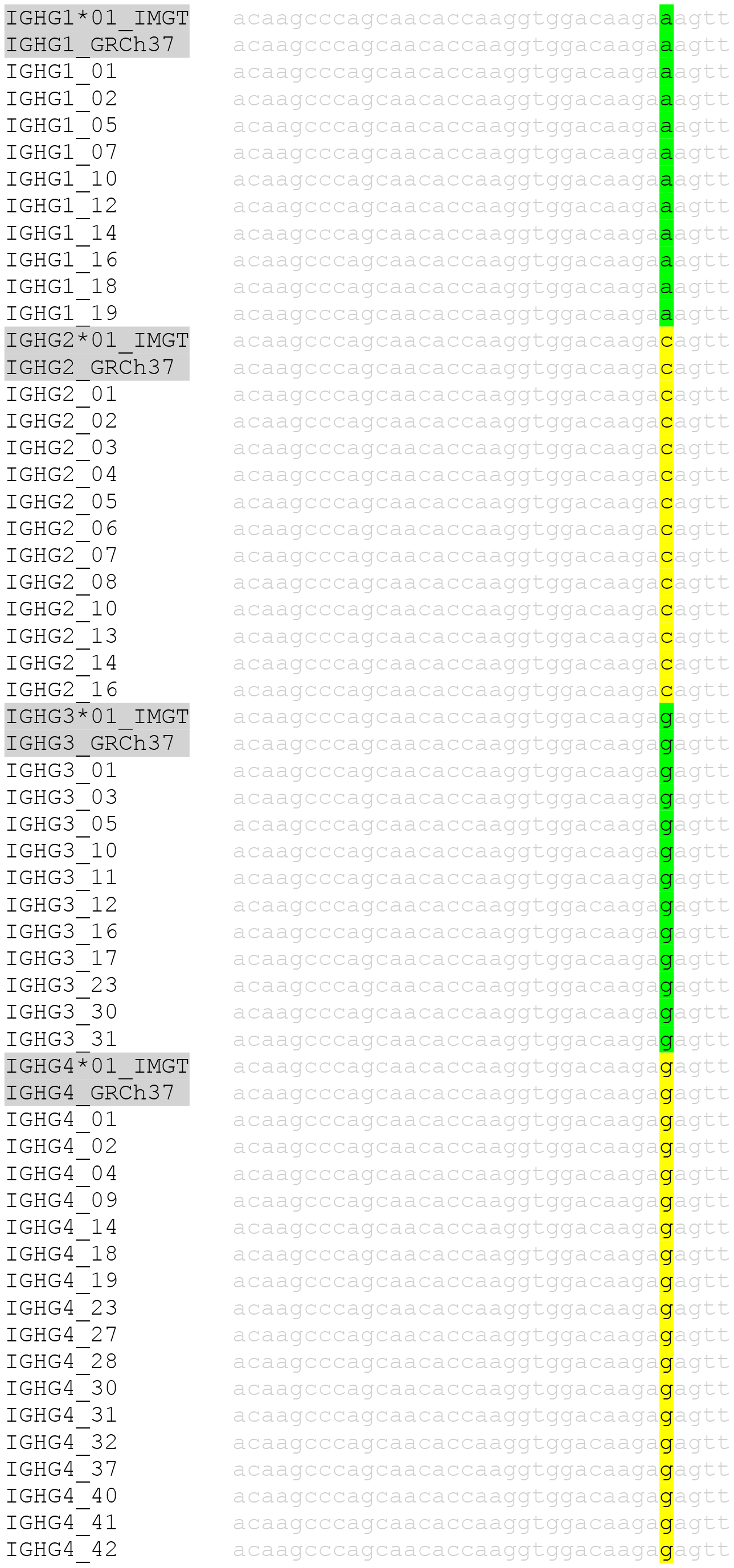
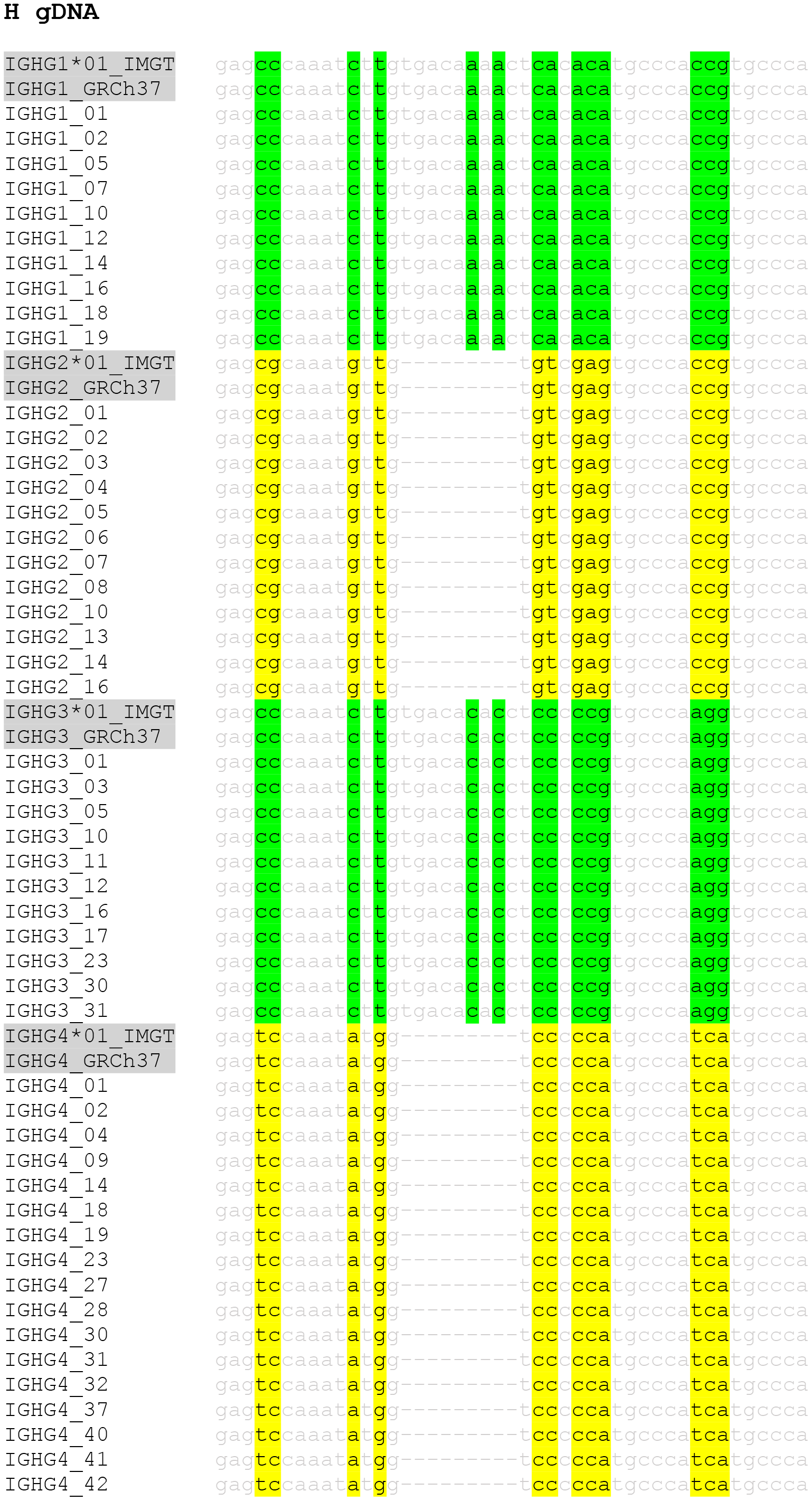
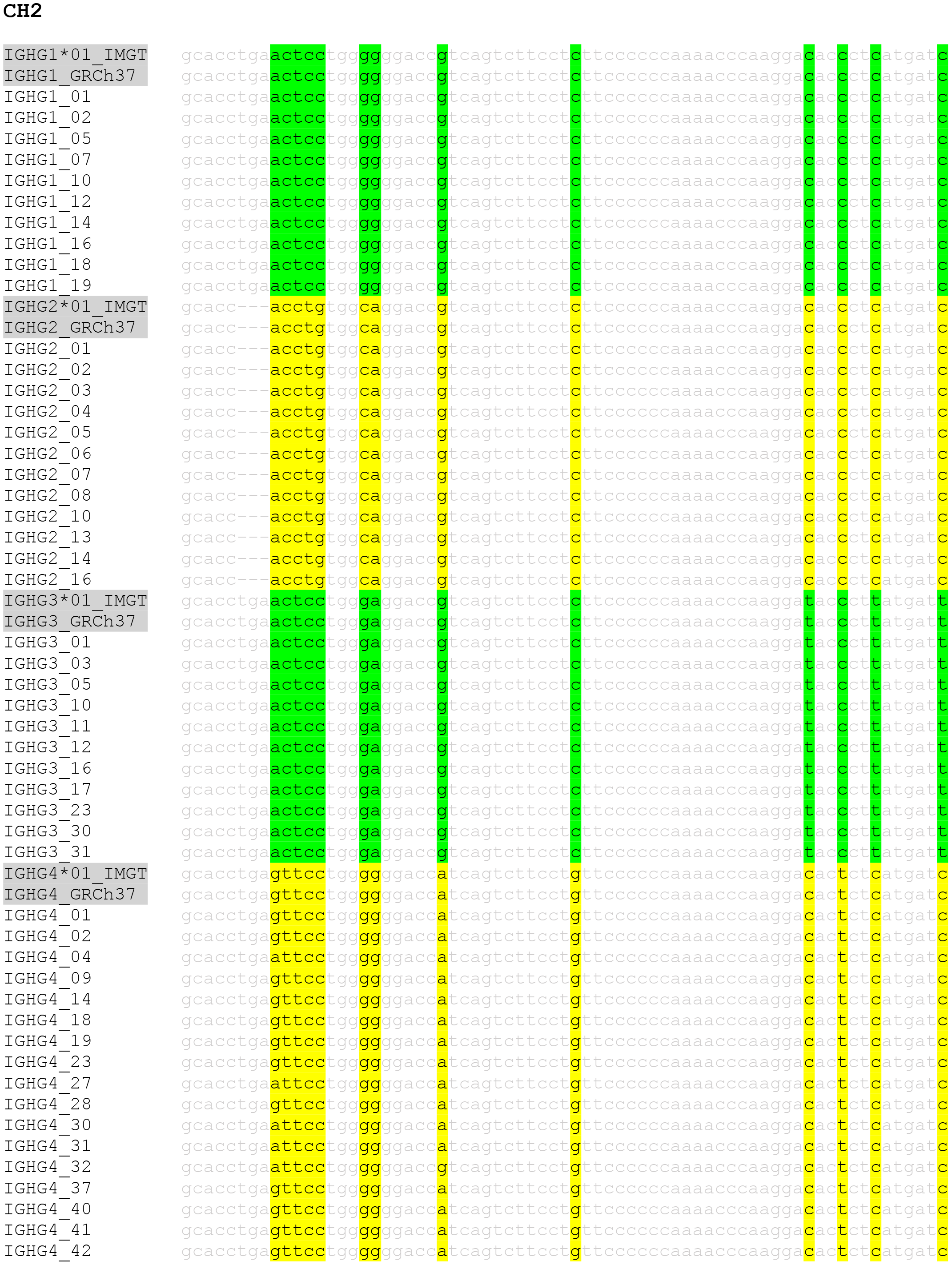
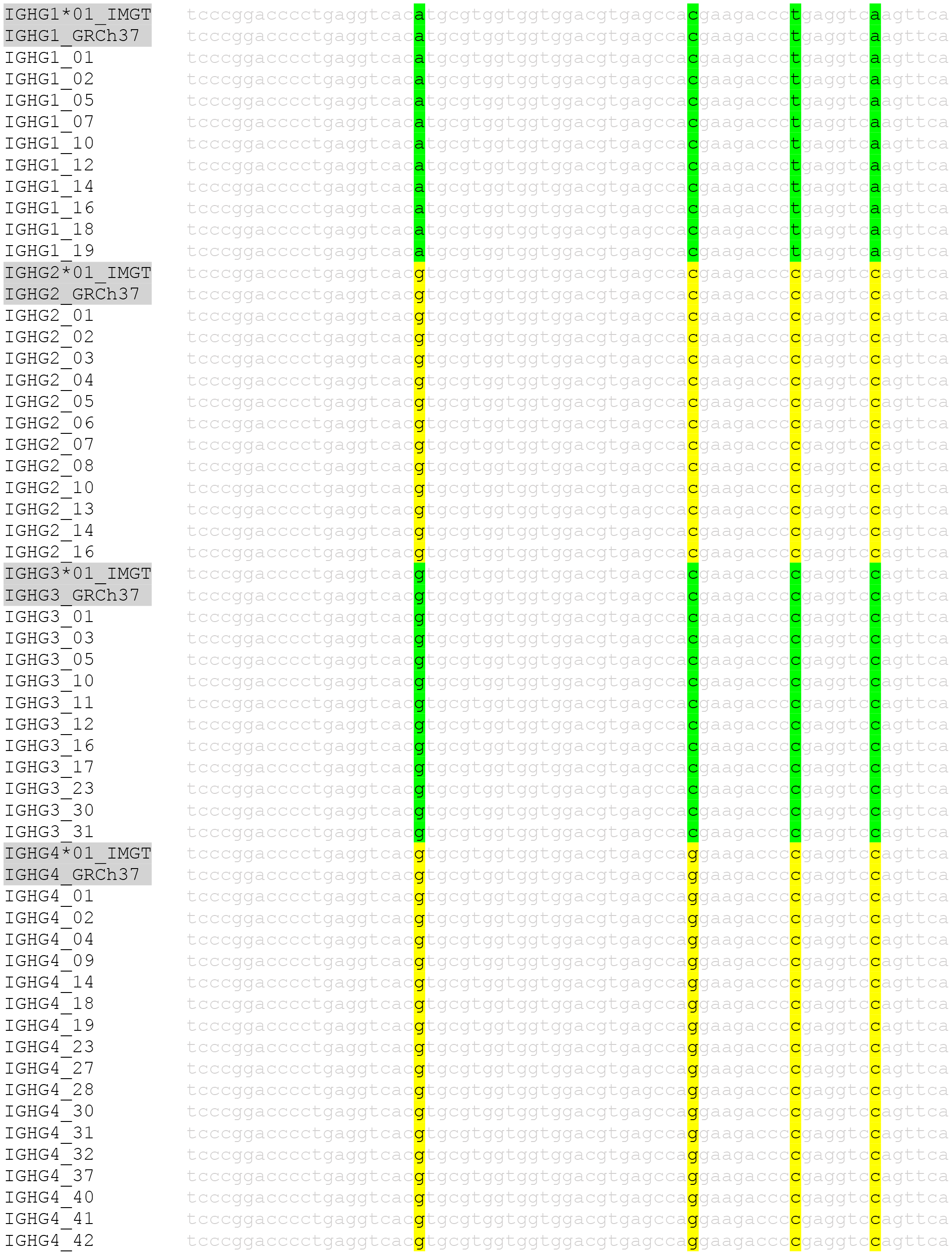
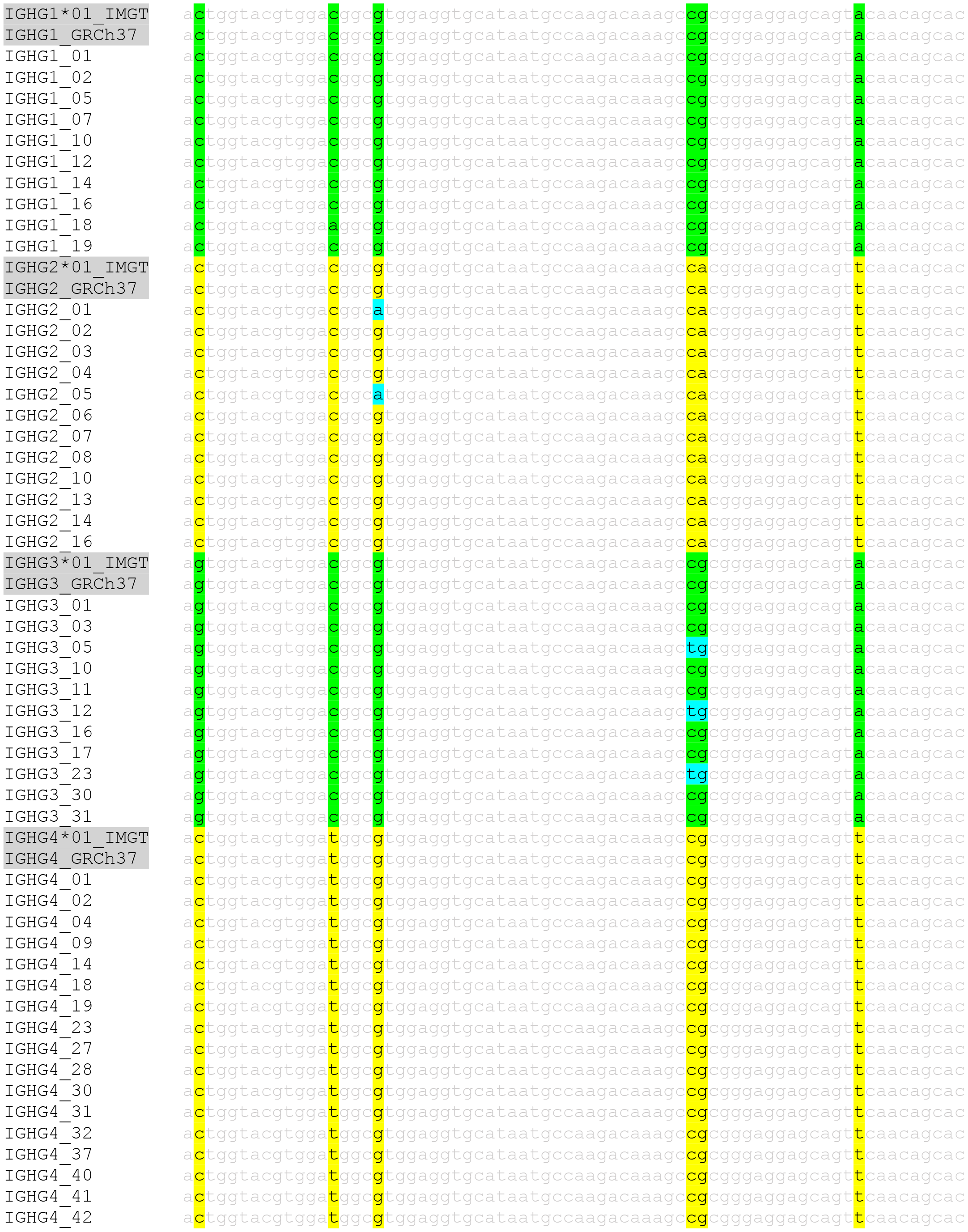
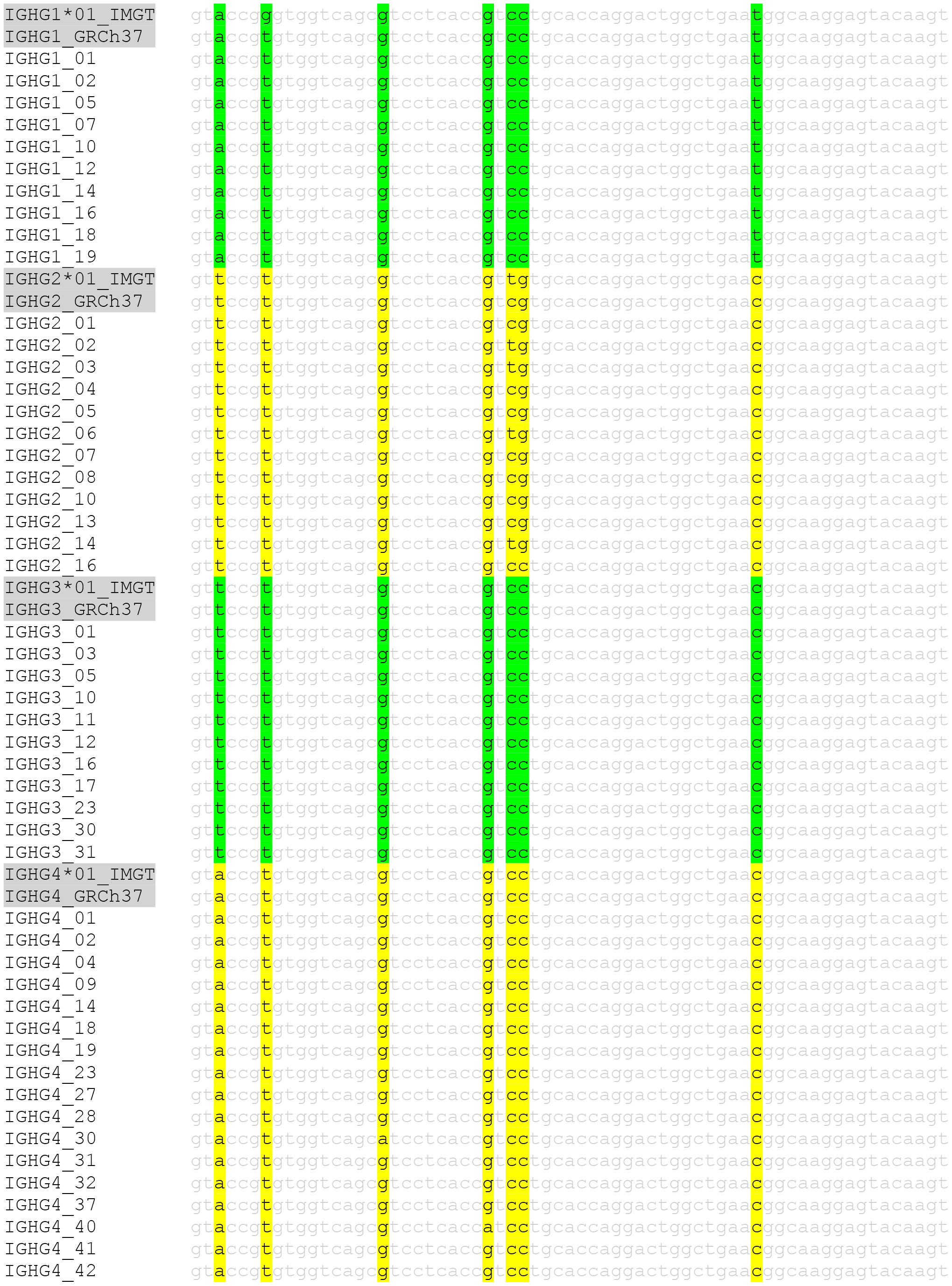
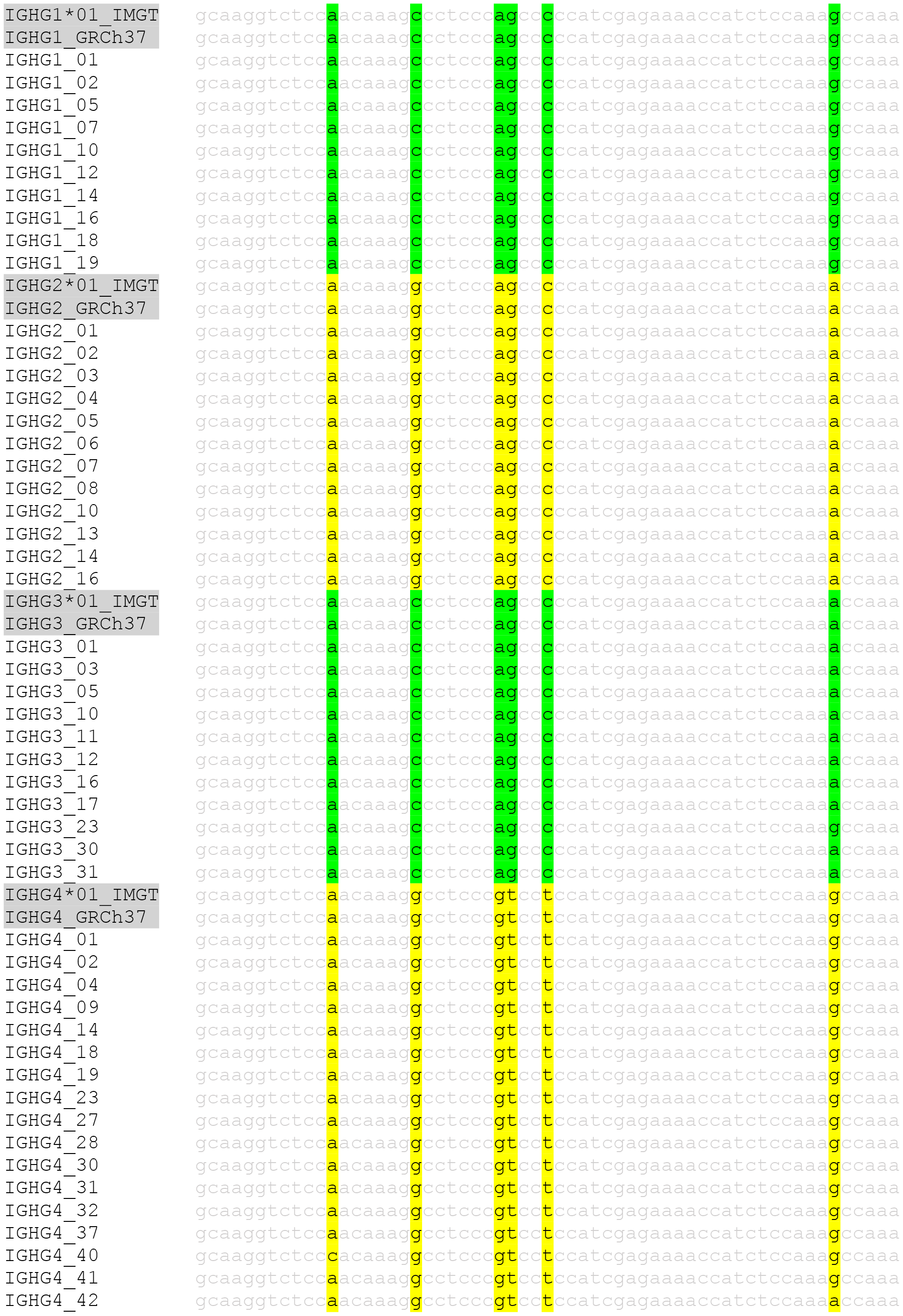
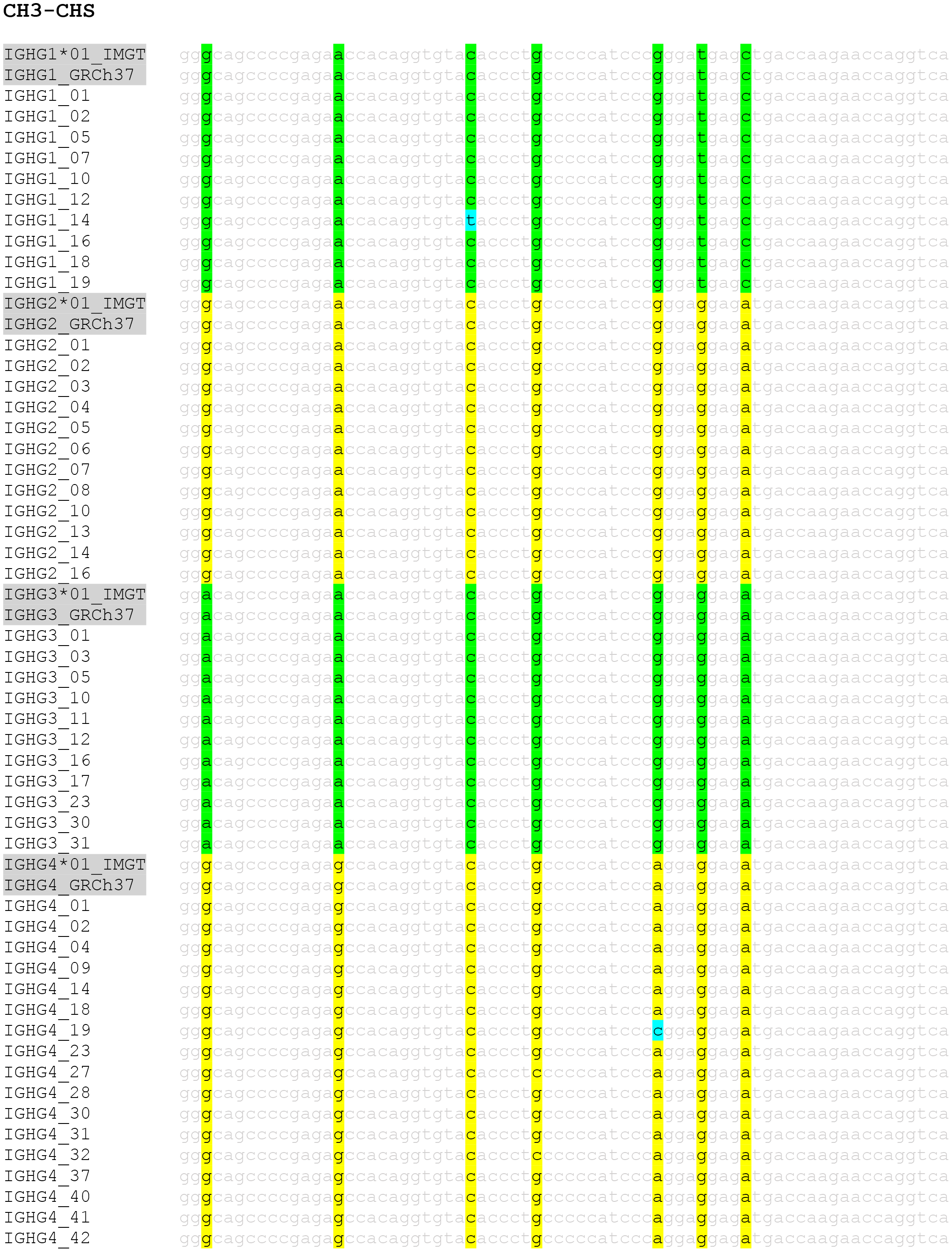
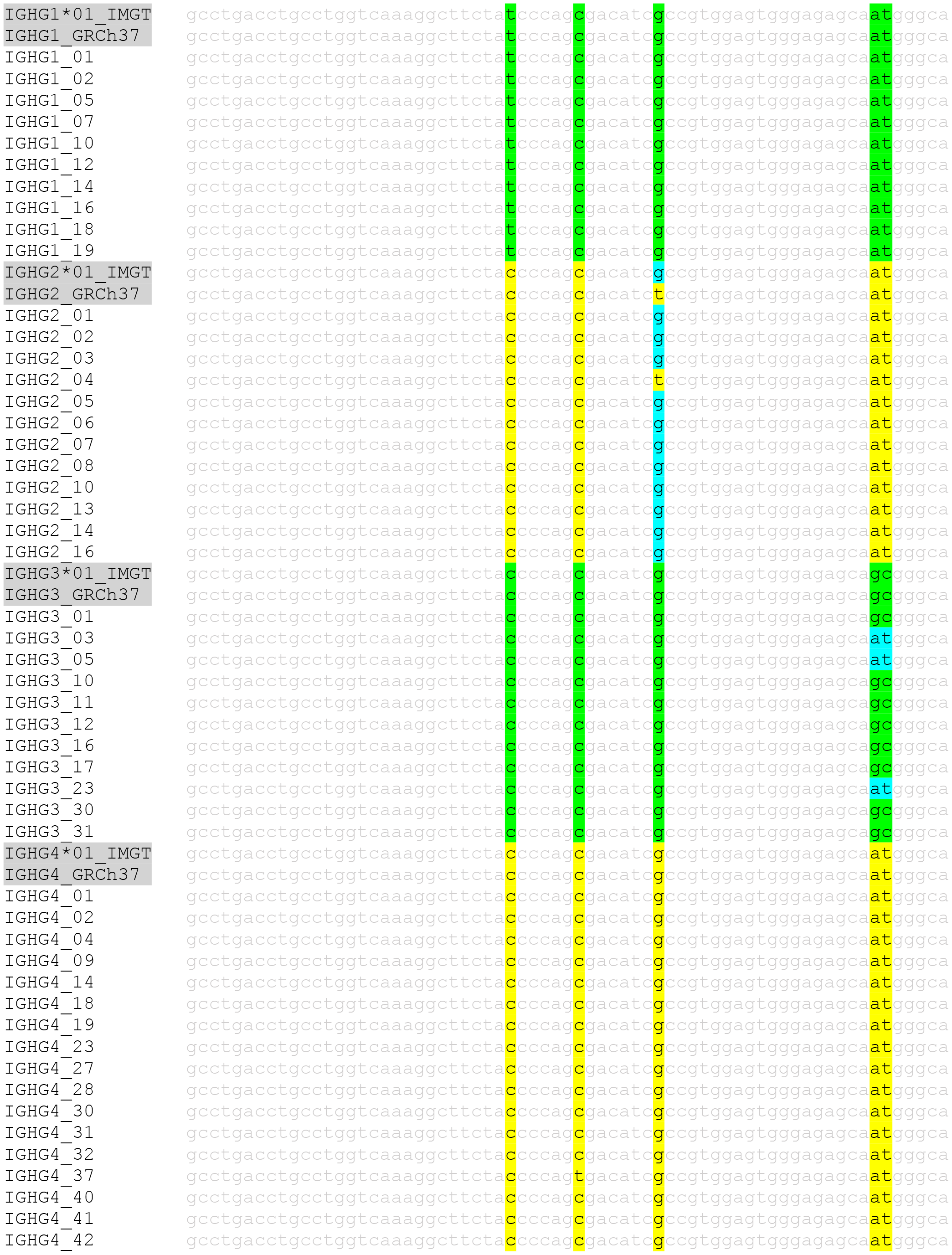
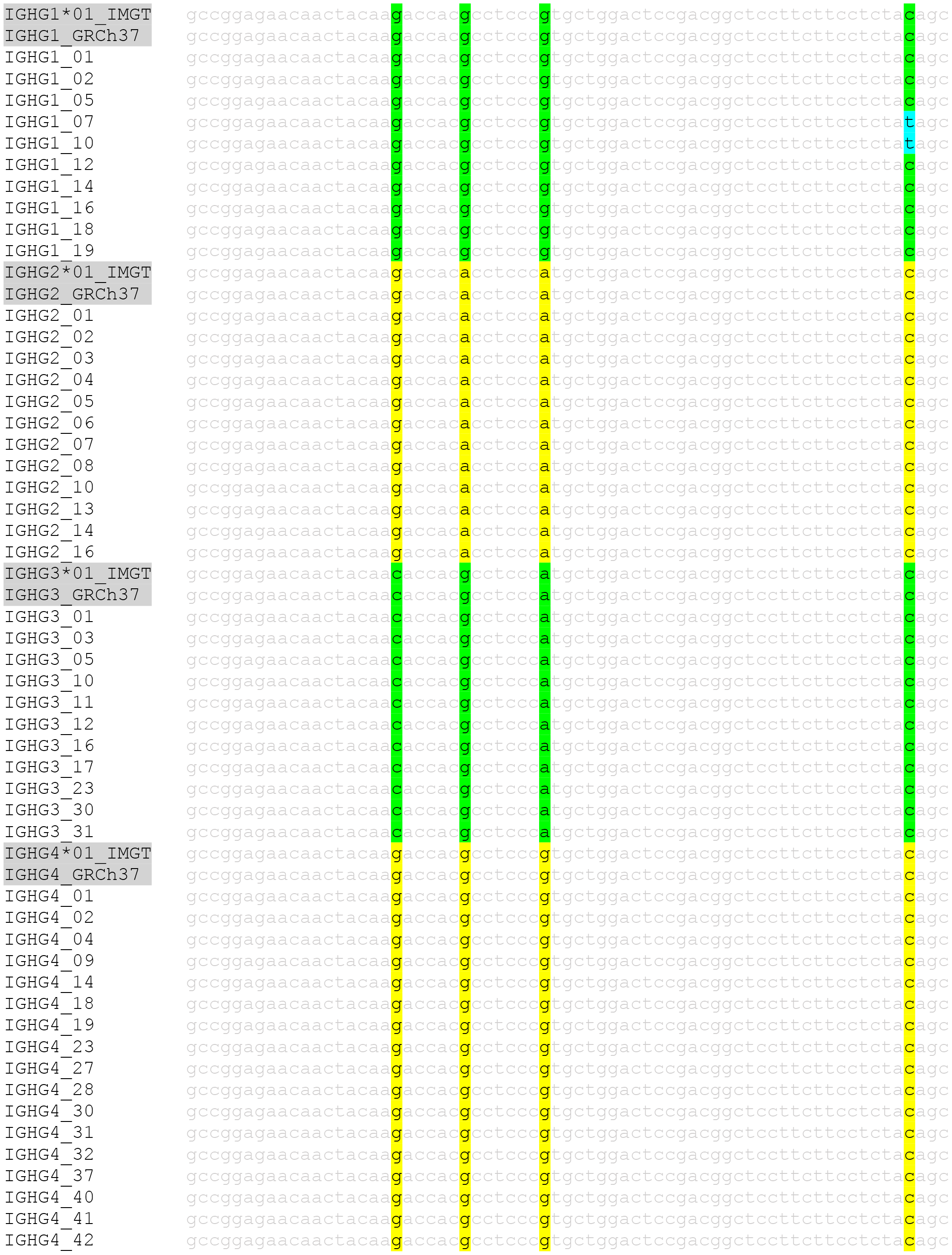
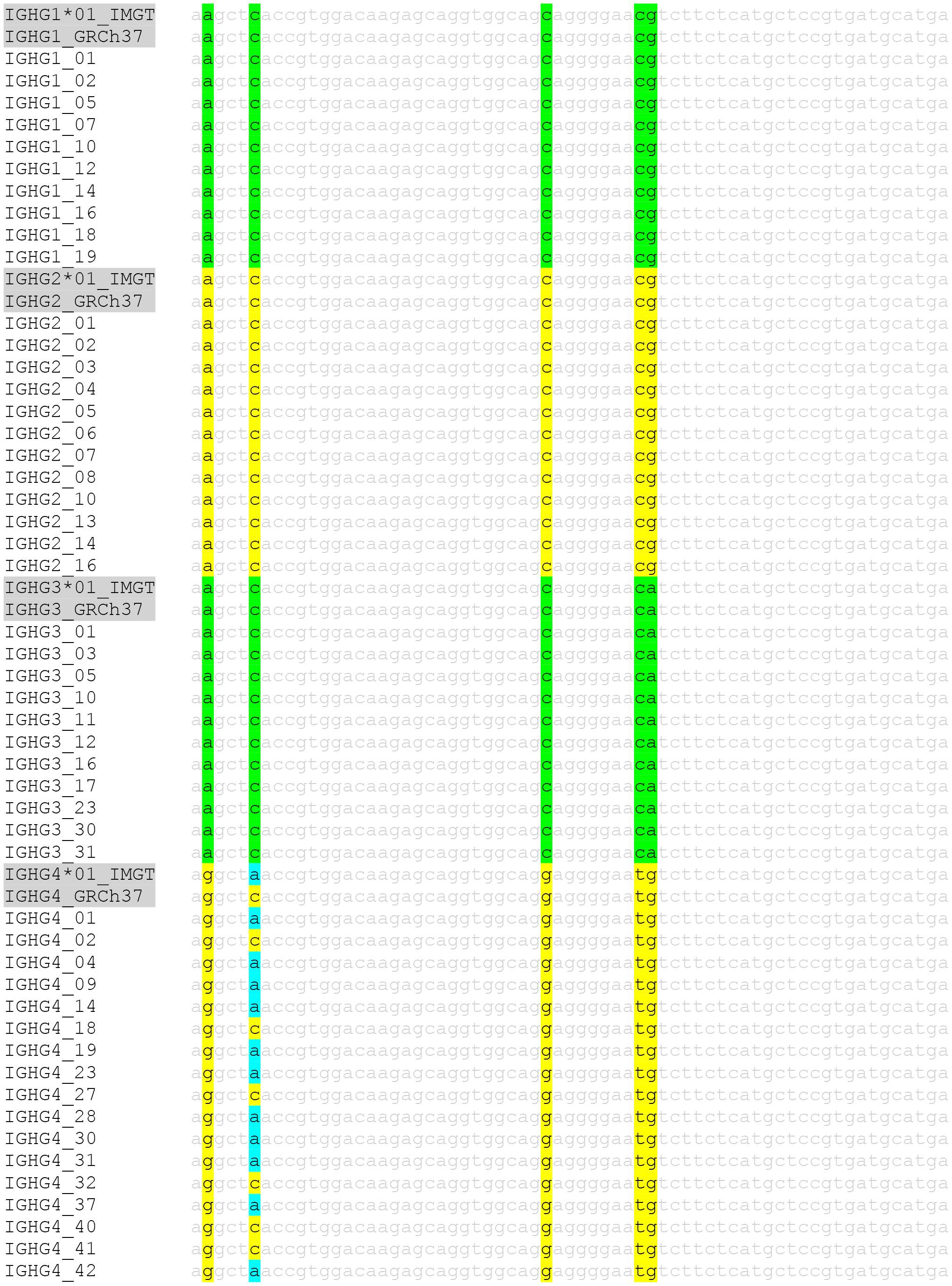
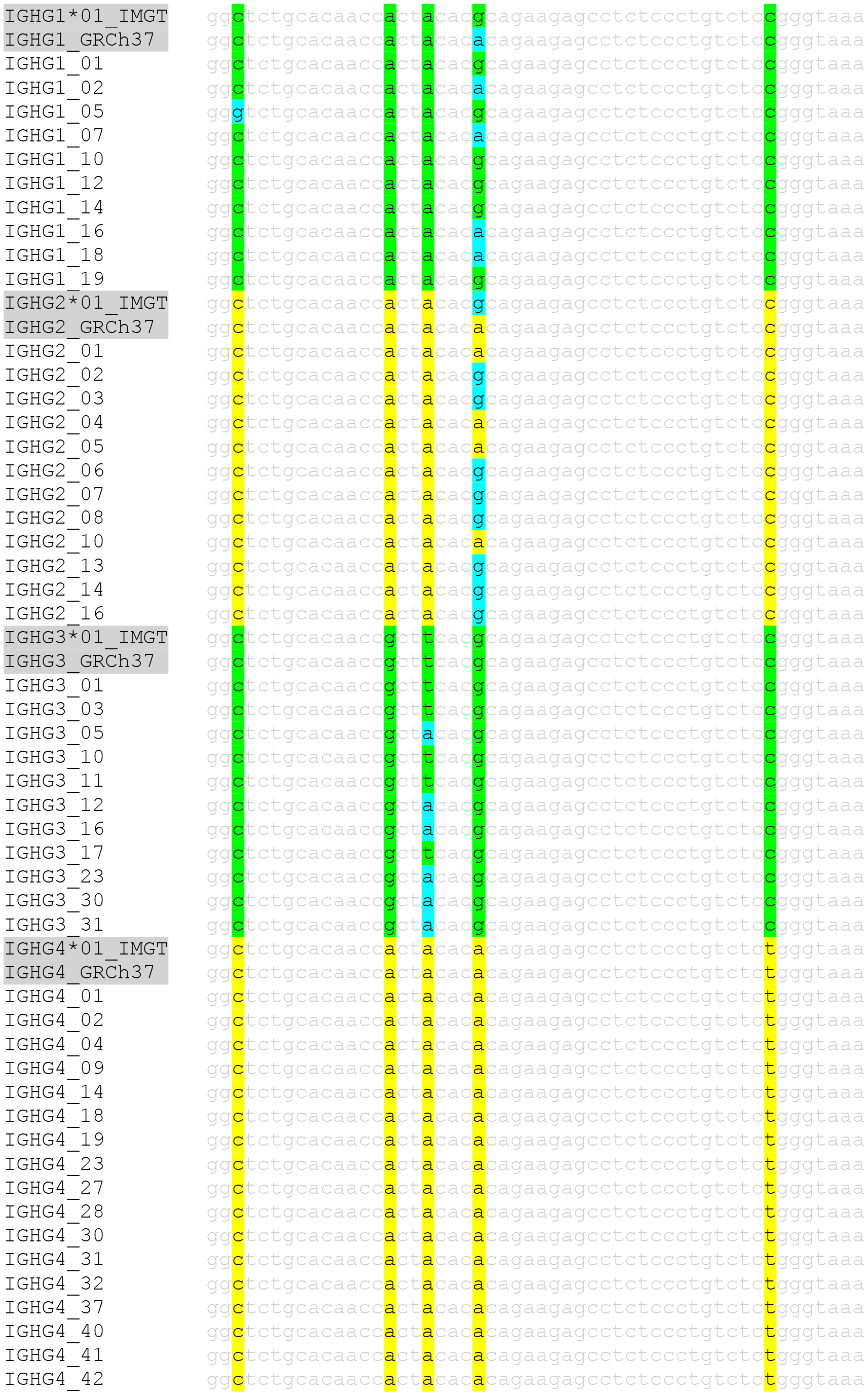
Alignment of CH1, CH2 and CH3 domain for *IGHA1*, *IGHA2* group and *IGHG1*, *IGHG2*, *IGHG3* and *IGHG4* group.

**Figure S6:**
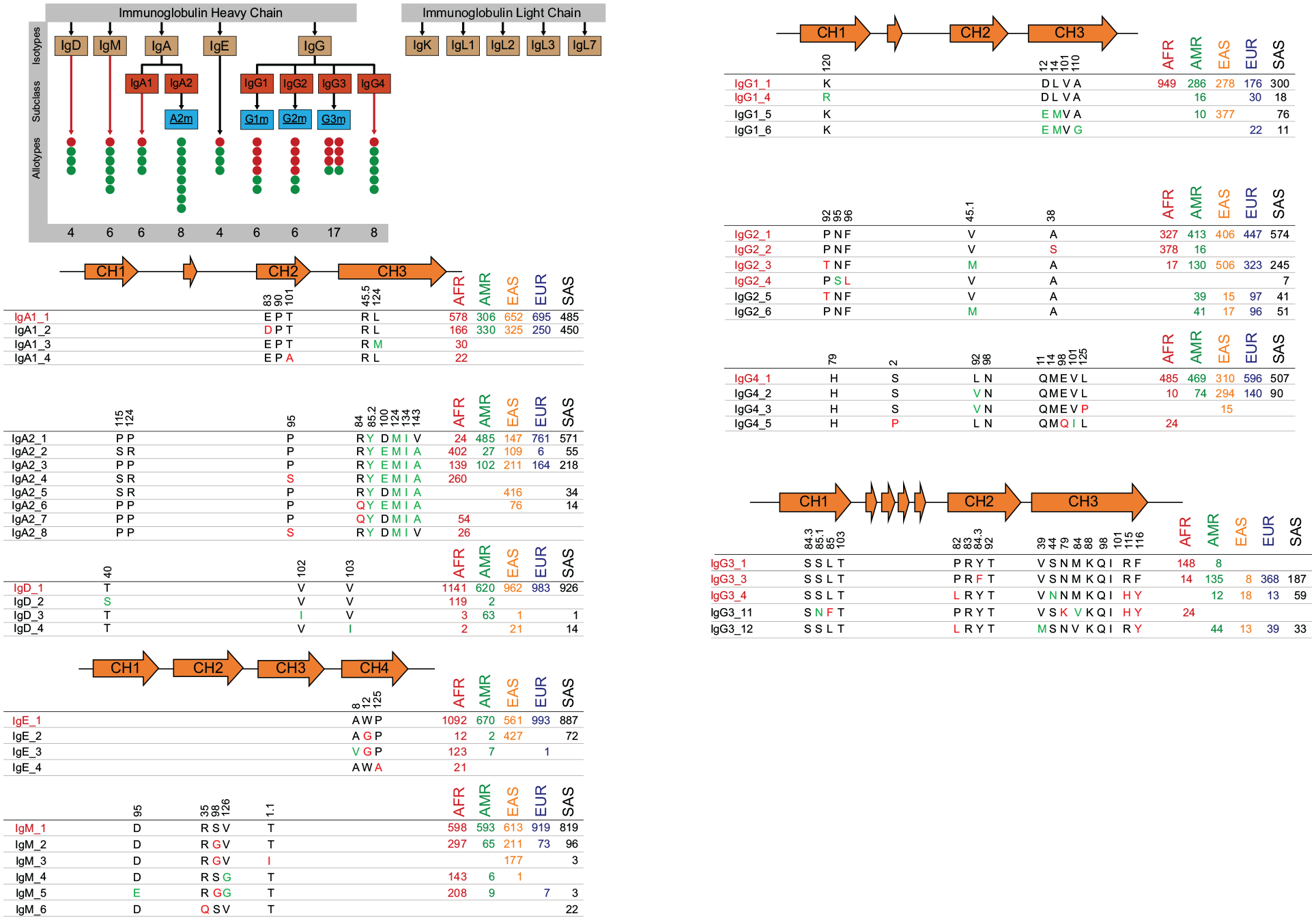
**Isotypes, Subclasses and allotypes of *IGH* locus.** The red circles in the nomenclature plot represents the number of allotypes (protein level translation of nucleotide germline alleles) matched with IMGT (similar to AS1) and green circles represents the new allotypes identified from G1K data. All the *IGH* constant genes are represented here. The three CH domains and one hinge region for IgA1, IgA2 and IgD is shown in 2-D diagram and the mutation existing in each domain is under the respective domain. The IgG3 protein has four hinges and three CH domains. IgM and IgE have four CH domains. The position of the amino-acid mutated is mentioned for the respective domain. The determinants marked in green represents no change in charge or structure whereas red determinant suggests the change in amino-acid properties.

**Figure S7:**
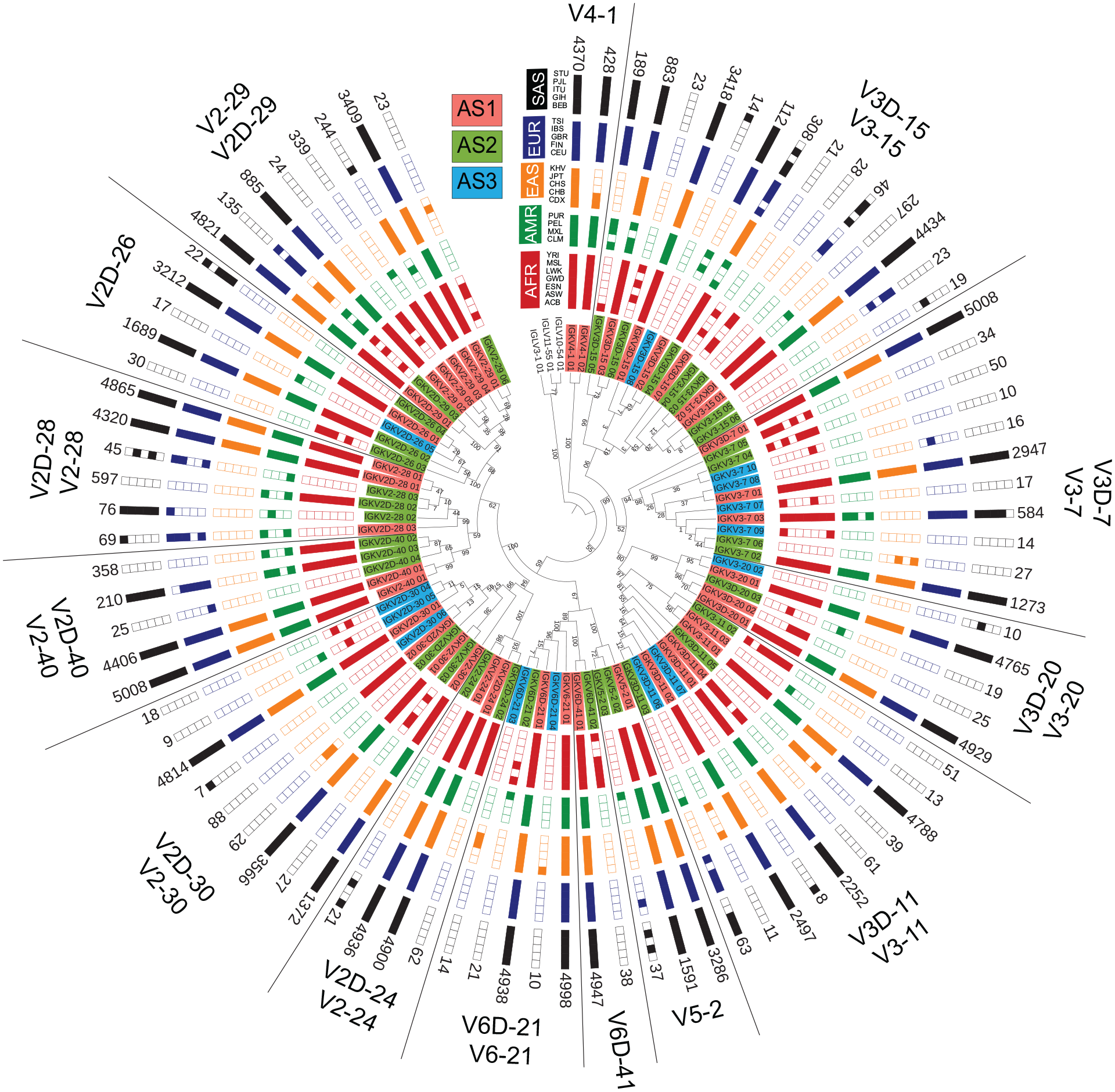
ML tree of the population distribution of *IGKV2, V3, V4, V5,* and *V6* family alleles.

**Figure S8:**
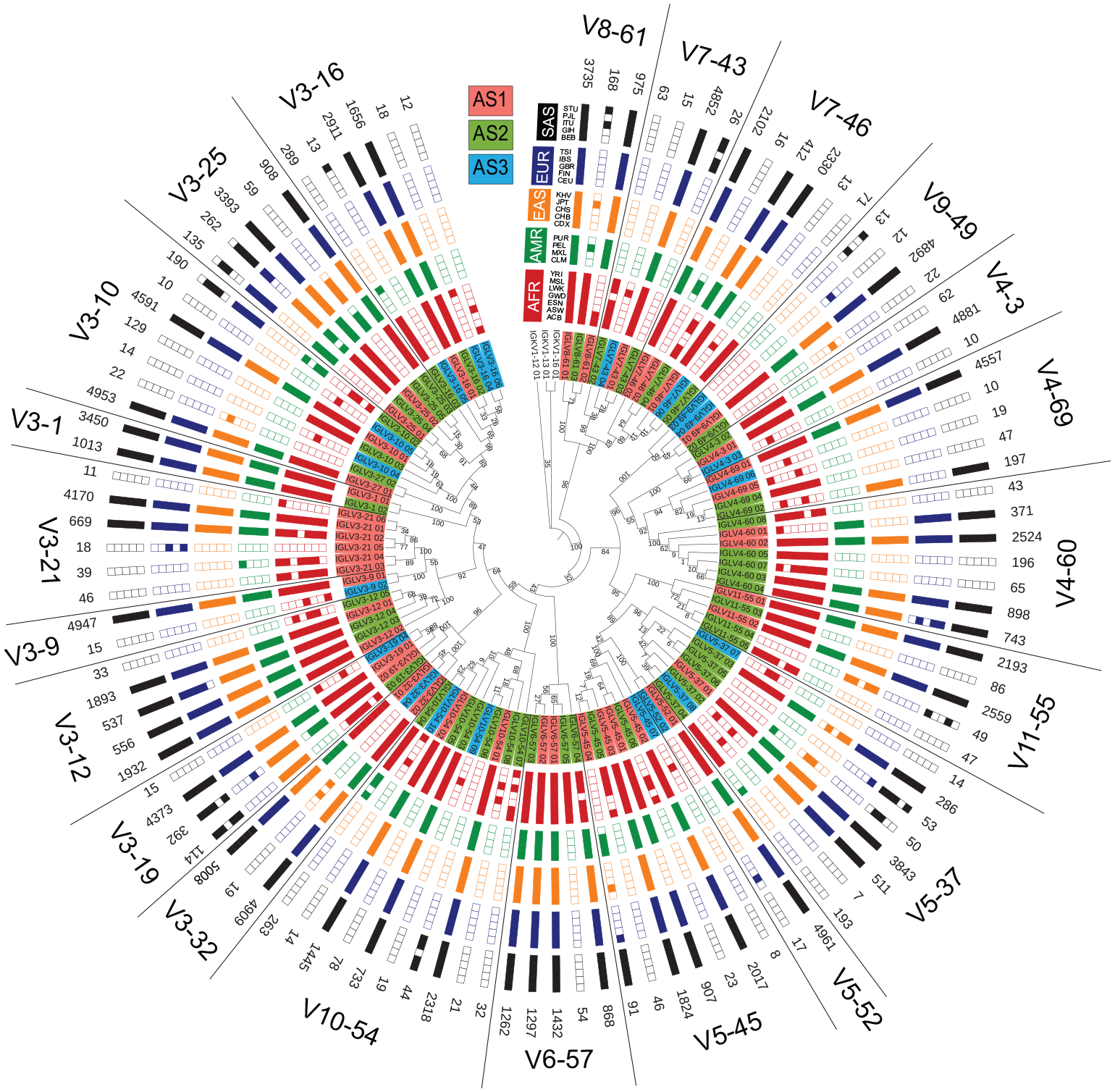
ML tree of the population distribution of *IGLV3, V4, V5, V6, V7, V8, V9, V10 and V11* family alleles.

## Supplementary Tables

Table S1: 1000 Genomes population information for 2504 and 563 individuals. Individual Sample IDs with their source and EBV coverage is mentioned in the table below. The 563 genomes selected are marked with green shade.

Table S2: pmIG functional alleles for *IGH* Locus with confidence level information, database mapping information, full sequence and haplotype support in both population and superpopulation.

Table S3: pmIG functional alleles for *IGK* Locus with confidence level information, database mapping information, full sequence and haplotype support in both population and superpopulation.

Table S4: pmIG functional alleles for *IGL* Locus with confidence level information, database mapping information, full sequence and haplotype support in both population and superpopulation.

Table S4: Deleted pmIG functional alleles for *IGH, IGK and IGL* Locus with confidence level information, database mapping information, full sequence and haplotype support in both population and superpopulation.

Table S5: *IG* pseudogenes with conserved heptamer of RSS sequence.

Table S6: RSS sequences in pmIG functional alleles for *IGH* Locus for each allele.

Table S7: RSS sequences in pmIG functional alleles for *IGK* Locus for each allele.

Table S8: RSS sequences in pmIG functional alleles for *IGL* Locus for each allele.

Note: For Supplementary Tables visit the GitHub page (https://github.com/InduKhatri/pmIG)

